# Biobased, Biodegradable but not bio-neutral: about the effects of polylactic acid nanoparticles on macrophages

**DOI:** 10.1101/2024.07.15.603484

**Authors:** Véronique Collin-Faure, Marianne Vitipon, Hélène Diemer, Sarah Cianférani, Elisabeth Darrouzet, Thierry Rabilloud

**Affiliations:** Chemistry and Biology of Metals, Univ. Grenoble Alpes, CNRS UMR5249, CEA, IRIG-LCBM, F-38054 Grenoble, France; Laboratoire de Spectrométrie de Masse BioOrganique (LSMBO), IPHC UMR 7178, Université de Strasbourg, CNRS, 67087 Strasbourg, France; Infrastructure Nationale de Protéomique ProFI - FR2048, 67087 Strasbourg, France

## Abstract

Plastics are persistent pollutants, because of their slow degradation, which suggests that they may lead to cumulative and/or delayed adverse effects due to their progressive accumulation over time. Macroplastics produced by human activity are released in the environment, where they degrade into micro and nanoplastics that are very easily uptaken by a wide variety of organisms, including humans. Microplastics and nanoplastics being particulates, they are handled in the body by specialized cells such as macrophages (or their evolutionary counterparts), where they can elicit a variety of responses. One solution to alleviate the problems due to biopersistence, such as accumulation over life, would be to use biodegradable plastics. One of the emerging biodegradable plastics being polylactide, we decided to test the responses of macrophages to polylactide nanoparticles, using a combination of untargeted proteomics and targeted validation experiments. Proteomics showed important adaptive changes in the proteome in response to exposure to polylactide nanoparticles. These changes affected for example mitochondrial, cytoskeletal and lysosomal proteins, but also proteins implicated in immune functions or redox homeostasis. Validation experiments showed that many of these changes were homeostatic, with no induced oxidative stress and no gross perturbation of the mitochondrial function. However, polylactide particles altered the immune functions such as phagocytosis (−20%) or cytokine production (2-fold increase for TNF production), which may translate into a decreased ability to macrophages to respond to bacterial infections. Furthermore, polylactide particles also induced moderate cross-toxicity with some quinones such as phenanthrene quinone, a combustion by-product that is a suspected carcinogen.

## 1. Introduction

The wide use of plastics in very diverse areas (e.g. packaging, automotive, textile, electronics, to quote just a few) translates into huge annual production figures, i.e. close to a gigaton/year ^1^. Unfortunately a huge proportion of these plastics, estimated to 500 Mt/year, is released in the environment ^2^, where it has deleterious effects that are more and more documented in detail, on a wide range of marine taxa ^3^, e.g. sea birds ^4^. This pollution was first documented in aquatic marine environments ^5–8^, but is now found in marine sediments^9,10^, freshwater environments ^11–13^, and also in terrestrial ones ^14,15^.

Although plastics are chemically resistant and degrade very slowly, with half lives in the environment amounting in decades ^16^, they progressively fragment, first into microplastics (lesser than 5 mm in size and then into nanoplastics (less than 1 µm in size). These nanoplastics are far more difficult to detect in the environment, and their effects are much less known. It is however anticipated that the smaller they are, the more easily they will cross biological barriers and be able to disturb the homeostasis of living organisms. This hypothesis has prompted extensive research on the effects of micro and nanoplastics on a wide variety of living organisms from various phylla, ranging from worms ^17,18^ to molluscs ^19–21^, crustaceans ^22,23^, insects ^24–26^, and then vertebrates such as fish ^27–29^ and of course mammals ^30^ including humans models (e.g. in ^31–35^).

Multicellular organisms defend themselves against particles, including plastic particles, by a series of mechanisms. The first line of defense is represented by biological barriers (e.g. intestinal or epidermal barriers) and research has been devoted to understand how these barriers interact with plastic particles. When translocation across these barriers occurs ^31^, then a second line of barriers comes into play, and is represented by professional phagocytes. This cell type is encountered in invertebrates (annelids coelomocytes, insect hemocytes) as well as in vertebrates (macrophages, neutrophils). Indeed, it has been shown that this cell type does respond to plastic particles ^18,26,36–42^.

In view of these deleterious effects and of the fact that fabrication of plastics consumes fossil resources and increases the concentration of greenhouse effect gases, the development of biobased and biodegradable plastics has been investigated and is a very active area of research. Plastics that are both biobased and biodegradable belong to two main families. The first family is represented by modified polysaccharides (e.g. modified starch or cellulose derivatives), and the second family is represented by polyhydroxyalkanoates, i.e. polymers of natural hydroxyacids. Indeed polyhydroxyalkanoates are produced by a variety of bacteria both as a protective medium and as a chemical energy storage ^43–47^. One of the simplest poly hydoxyalkanoate, i.e. poly lactic acid (PLA), was first used as a material for surgical sutures ^48,49^, and it was demonstrated that it was biodegradable ^50^. It also gained popularity by the fact that it could be polymerized as nanospheres for drug encapsulation and controlled release, most often as a copolymer of lactic and glycolic acid ^51–53^. However, it was also shown that pure PLA is less toxic than the copolymer ^54,55^, so that PLA is a preferred choice as a model for a safe, biobased and biodegradable plastic available in the nanoplastic format.

Recent work showed that degradation of PLA produced fragmentation and nanoparticles ^56^, even in domestic use ^57^, which further increases the probability that living organisms may encounter PLA nanoparticles, and especially if the use of this plastic increases. We thus decided to investigate the responses of macrophages to PLA nanoparticles, using a combination of proteomic and targeted approaches, as previously done with polystyrene nanoparticles ^41^. In this approach, proteomics is used to gain a wide appraisal of the cell responses, highlighting hypotheses that are then verified by targeted experiments.

## 2. Materials and Methods

### 2.1. Plastic particles

Poly lactic acid (PLA) particles (red Poly-lactic acid 150 nm, fluorescent red labelled, catalog number #RFiP-600-150 batch number #230612-ip) were purchased from Adjuvatis (Lyon, France) and provided as sterile 3% (w/v) suspensions. Deep red fluorescent labelled polystyrene (PS) particles (SkyBlue particles, 100-300 nm, catalog number #FP-0270-2, batch number #AL01) were purchased from Spherotech and provided as non-sterile 0.25% (w/v) suspensions. The particles were used for both the proteomic experiments and the validation experiments. The polystyrene suspensions were pasteurized overnight at 80°C before use in cell culture. The PLA suspension was provided sterile by the supplier. The particles were excited at 560 nm and their emission read at 613/18 nm for the PLA particles and at 695/40 nm for the PS particles.

The particles were diluted at 10 µg/mL in PBS 0.001X and characterized by DLS using a Litesizer 500 (Anton Paar, USA) and Omega Cuvette (225288, Anton Paar, USA). An average value was obtained from repeated measurements for each sample (n = 3) and analyzed with the instrument-associated Kaliope software.

### 2.2. Cell culture

The J774A.1 cell line (mouse macrophages) was purchased from European cell culture collection (Salisbury, UK). Cells were routinely propagated in Dulbecco’s Modified Eagle’s Medium (DMEM) supplemented with 10% fetal bovine serum (FBS) in non-adherent flasks (Cellstar flasks for suspension culture, Greiner Bio One, Les Ulis, France). For routine culture, the cells were seeded at 200,000 cells/ml and passaged two days later, with a cell density ranging from 800,000 to 1,000,000 cells/ml.

For exposure to plastic particles and to limit the effects of cell growth, cells were seeded at 500,000 cells/ml in 6 or 12 wells plates, let settle and recover for 24 hours, and then exposed to the particles at 50µg/ml for 24 hours before harvesting for the experiments. Proteomic experiments were carried out in 6 well plates, and all the other experiments in 12 well plates. The medium volume was adjusted to keep the same height across all cell culture formats. Cells were used at passage numbers from 5 to 15 post-reception from the repository. Cell viability was measured by the propidium iodide method ^58^, or with the SytoxGreen probe (Thermofisher S7020) using the protocol provided by the supplier.

For assessing the persistence of the particles in the cells, we followed the strategy published previously^42^. The cells were seeded in adherent 12-well plates at 400,000 cells/ml in DMEM supplemented with 1% horse serum, in order to limit their proliferation ^59^. After adaptation to the medium for 72 hours (D0), the medium was changed and the cells were exposed to the plastic particles for 24 hours at a concentration of 50 or 100 µg/ml. The particle-containing medium was removed (D1). The remaining cells were cultivated in DMEM supplemented with 1% horse serum for an additional 3 days without splitting, and with a culture medium change every 2 days. Wells were harvested every day starting at D1 during this process and analyzed by flow cytometry to measure the cell-associated fluorescence. In order to normalize for the cell number, a cell numeration was performed on the harvested cells in each well. This process allowed to determine the remaining fraction of particles over time after the initial loading (D1).

### 2.3. Confocal microscopy

J774.A1 cells were seeded onto glass coverslips at a density of 200,000 cells/mL in DMEM supplemented with 1% horse serum and 1% streptomycin-penicillin. The cells were then incubated overnight at 37°C with 5% CO2.

Subsequently, PLA beads were added to the cells at a concentration of 50µg/ml, and further incubation was carried out for 24 hours. The medium was then changed for particle-free medium and the cells were let to recover for 24 hours and 48 hours at 37°C.

After each incubation period (without exposure, immediately after exposure and after the 24 and 48 hours recovery periods) the cells were fixed with 4% paraformaldehyde for 30 minutes at room temperature and permeabilized with 0.1% Triton X-100 for 20 minutes at room temperature. Following permeabilization, the cells were incubated with Phalloidin Atto 488 (1/500 dilution) and DAPI (1/1000 dilution) for 20 minutes and 5 minutes, respectively, at room temperature. Between each step, the cells were washed three times with PBS (1X).

Confocal microscopy experiments were performed using a Zeiss LMS880 confocal microscope (Carl Zeiss, Jena, Germany) equipped with a 20× objective. Laser tracks were set as following:

- Dapi: Excitation at 405nm, Emission at [410 – 467]nm
- Phalloidin Atto 488: Excitation at 488nm, Emission at [494 - 550]nm
- PLA beads: Excitation at 561nm, Emission at [596 – 632]nm

Laser settings were optimized using the 24-hour-exposed cells labeled with Phalloidin and DAPI to maximize fluorescence intensity before imaging each condition. Confocal laser scanning microscopy was employed to visualize the actin structure, nucleus, and PLA beads.

The acquired images were processed using ImageJ software. Specifically, adjustments were made to enhance the quality and uniformity of the images. Green fluorescence intensity in the B1 merged image was increased to ensure consistency across all panels. Similarly, brightness levels in the red channel were uniformly adjusted for all samples, with or without PLA beads.

### 2.4. Proteomics

Proteomics was carried out essentially as described previously ^41^. However, the experimental details are given here for the sake of consistency.

#### 2.4.1. Sample preparation

After exposure to the plastic particles, the cells were harvested by flushing the 6 well plates. They were collected by centrifugation (200g, 5 minutes) and rinsed twice in PBS. The cell pellets were lysed in 100 µl of extraction buffer (4M urea, 2.5% cetyltrimethylammonium chloride, 100mM sodium phosphate buffer pH 3, 150µM methylene blue). The extraction was let to proceed at room temperature for 1 hour, after which the lysate was centrifuged (15,000g, 30 minutes) to pellet the nucleic acids. The supernatants were then stored at −20°C until use.

#### 2.4.2. Shotgun proteomics

For the shotgun proteomic analysis, the samples were included in polyacrylamide plugs according to Muller et al. ^62^ with some modifications to downscale the process ^63^. To this purpose, the photopolymerization system using methylene blue, toluene sulfinate and diphenyliodonium chloride was used ^64^.

As mentioned above, the methylene blue was included in the cell lysis buffer. The other initiator solutions consisted in a 1 M solution of sodium toluene sulfinate in water and in a saturated water solution of diphenyliodonium chloride. The ready-to-use polyacrylamide solution consisted of 1.2 ml of a commercial 40% acrylamide/bis solution (37.5/1) to which 100 µl of diphenyliodonium chloride solution, 100 µl of sodium toluene sulfinate solution and 100 µl of water were added.

To the protein samples (15 µl), 5 µl of acrylamide solution were added and mixed by pipetting in a 500µl conical polypropylene microtube. 100 µl of water-saturated butanol were then layered on top of the samples, and polymerization was carried out under a 1500 lumen 2700K LED lamp for 2 hours, during which the initially blue gel solution discolored. At the end of the polymerization period, the butanol was removed, and the gel plugs were fixed for 2×1 hr with 200 µl of 30% ethanol 2 % phosphoric acid, followed by a 30 minutes wash in 30% ethanol. The fixed gel plugs were then stored at −20°C until use.

Gel plug processing, digestion, peptide extraction and nanoLC-MS/MS was performed as previously described, without the robotic protein handling system and using a Q-Exactive Plus mass spectrometer (Thermo Fisher Scientific, Bremen, Germany). Further details are available in **Methods S1**.

For protein identification, the MS/MS data were interpreted using a local Mascot server with MASCOT 2.6.2 algorithm (Matrix Science, London, UK) against an in-house database containing all Mus musculus and Rattus norvegicus entries from UniProtKB/SwissProt (version 2019_10, 25,156 sequences) and the corresponding 25,156 reverse entries. Spectra were searched with a mass tolerance of 10 ppm for MS and 0.05 Da for MS/MS data, allowing a maximum of one trypsin missed cleavage. Trypsin was specified as enzyme. Acetylation of protein N-termini, carbamidomethylation of cysteine residues and oxidation of methionine residues were specified as variable modifications. Identification results were imported into Proline software version 2.2 (http://profiproteomics.fr/proline) for validation. Peptide Spectrum Matches (PSM) with pretty rank equal to one and a length greater than 7 amino acids were retained. False Discovery Rate was then optimized to be below 1% at PSM level using Mascot Adjusted E-value and below 1% at Protein Level using Mascot Standard score.

Mass spectrometry data are available via ProteomeXchange with the identifier PXD048664.

#### 2.4.3. Label Free Quantification

Peptides abundances were extracted thanks to Proline software version 2.2 (http://profiproteomics.fr/pro-line) using a m/z tolerance of 10 ppm. Alignment of the LC-MS runs was performed using Loess smoothing. Cross Assignment was performed within groups only. Protein Abundances were computed by sum of peptides abundances (normalized using the median).

#### 2.4.4. Data analysis

For the global analysis of the protein abundances data, missing data were imputed with a low, non-null value. The complete abundance dataset was then analyzed by the PAST software ^65^.

Proteins were considered as significantly different if their p value in the Mann-Whitney U-test against control values was inferior to 0.05. No quantitative change threshold value was applied. The selected proteins were then submitted to pathway analysis using the DAVID tool ^66^, with a cutoff value set at a FDR of 0.1.

### 2.5. Mitochondrial transmembrane potential assay

The mitochondrial transmembrane potential assay was performed essentially as described previously ^42^. Rhodamine 123 (Rh123) was added to the cultures at an 80 nM final concentration (to avoid quenching ^60^), and the cultures were further incubated at 37°C for 30 minutes. At the end of this period, the cells were collected, washed in cold PBS containing 0.1% glucose, resuspended in PBS glucose and analyzed for the green fluorescence (excitation 488 nm emission 525nm) on a Melody flow cytometer. As a positive control, butanedione monoxime (BDM) was added at a 30mM final concentration together with the Rh123 ^61^. As a negative control, carbonyl cyanide 4-(trifluoromethoxy)phenylhydrazone (FCCP) was added at 5µM final concentration together with the Rh123 ^61^.

### 2.6. Phagocytosis assay

For this assay, the cells were first exposed to the red fluorescent particles. After 24 hours of exposure, the cells were the exposed to 0.5 µm latex beads (carboxylated surface, yellow green-labelled, from Polysciences excitation 488 nm emission 527/32 nm) for 3 hours. After this second exposure, the cells were collected, rinsed twice with PBS, and analyzed for the two fluorescences (green and red) on a Melody flow cytometer.

### 2.7. Lysosomal assay

For the lysosomal function assay, the Lysosensor method was used, as described previously ^42^. After exposure to plastic beads, the medium was removed, the cell layer was rinsed with complete culture medium and incubated with 1µM Lysosensor Green (Molecular Probes) diluted in warm (37°C) complete culture medium for 1 hour at 37°C. At the end of this period, the cells were collected, washed in cold PBS containing 0.1% glucose, resuspended in PBS glucose and analyzed for the green fluorescence (excitation 488 nm emission 540nm) on a Melody flow cytometer.

### 2.8. Cytokine release assays

Cells were first exposed to nanoplastics (50µg/ml) for 24 hours. At the end of this exposure period, the culture medium was removed, the cell layer was rinsed with culture medium and fresh medium was added to the wells. In half of the wells LPS (1 ng/ml) was added. After another 24 hours, the medium was collected and analyzed for proinflammatory cytokines. Tumor necrosis factor (catalog number 558299, BD Biosciences, Le Pont de Claix, France) and interleukin 6 (IL-6) (catalog number 558301, BD Biosciences, Le Pont de Claix) levels were measured using the Cytometric Bead Array Mouse Inflammation Kit (catalog number 558266, BD Biosciences, Le Pont de Claix), and analyzed with FCAP Array software (3.0, BD Biosciences) according to the manufacturer’s instructions.

### 2.9. Assay for oxidative stress

For the oxidative stress assay, a protocol based on the oxidation of dihydrorhodamine 123 (DHR123) was used, essentially as described previously ^42^. After exposure to plastic beads, the cells were treated in PBS containing 500 ng/ml DHR123 for 20 minutes at 37°C. The cells were then harvested, washed in cold PBS containing 0.1% glucose, resuspended in PBS glucose and analyzed for the green fluorescence (same parameters as rhodamine 123) on a Melody flow cytometer. Menadione (applied on the cells for 2 hours prior to treatment with DHR123) was used as a positive control in a concentration range of 25-50µM.

### 2.10. Cross-toxicity assays

For these assays, the cells were plated at 400,000 cells/ml in a 12-well non adherent plate. After 24 hours, 50µg/ml of beads were added, followed 6 hours later by a variable concentration of the toxicant to be tested, pre-dissolved in ethanol so that the final ethanol concentration in the culture well did not exceed 1% in volume. Toxicants tested included menadione and 9,10-phenanthrene quinone.

## 3. Results

### 3.1. Plastics beads characterization

Plastic beads were characterized by DLS and electrophoretic mobility in order to determine their average hydrodynamic size and zeta potential. The results are presented in Table 1

**Table 1:**
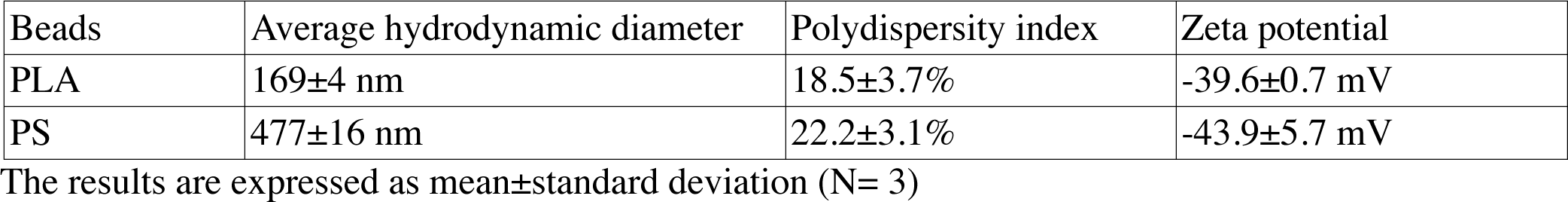
Characterization of the beads parameters.

| Beads | Average hydrodynamic diameter | Polydispersity index | Zeta potential |
| --- | --- | --- | --- |
| PLA | 169±4 nm | 18.5±3.7% | -39.6±0.7 mV |
| PS | 477±16 nm | 22.2±3.1% | -43.9±5.7 mV |

The results are expressed as mean±standard deviation (N= 3)

The PS beads proves larger in size through these measurements than the nominal diameter given by the supplier (0.1-0.3 µm, mean 0.26 µm). This larger size however did not prevent a bead/cell number ratio close to 1000 beads/cell (at 50µg/ml) for the largest beads and higher than 10,000 beads/cell for the smaller ones.

### 3.2. Viability of the plastics-treated cells

First, the toxic effects of the PLA beads on J774A.1 cells was determined after a 24 hours exposure. The results, shown in **Fig. 1a**, demonstrated a very low toxicity of the PLA beads, with a LD50 that reached 400µg/ml. As we planned to use polystyrene beads as positive controls for non-degradable plastics in subsequent experiments, we decided to use a dose that was half of the LD20 (100µg/ml) for the PS beads, i.e. 50µg/ml. This dose showed very low toxicity for the PLA beads, but was however the first concentration for which the viability was slightly but statistically significantly lower than the one of unexposed, control cells. As the particles were labelled, we could follow their internalization and their degradation. As shown on **Fig. 1b**, the amount of internalized PLA particles increased with increasing concentration in the medium. However, the amount of intracellular fluorescence rapidly decreased over time, suggesting a rather fast degradation of the particles.

**Figure 1:**
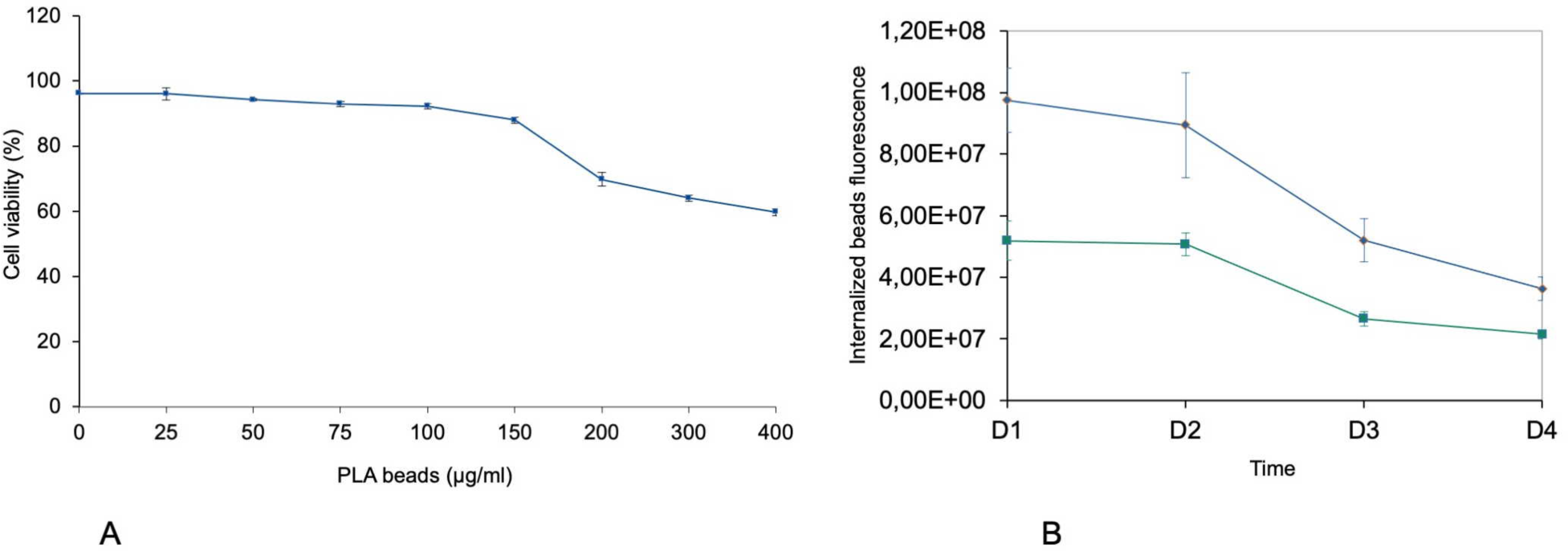
Viability and intracellular fluorescence of cells treated with PLA particles. In panel 1A, cells were treated with PLA particles for 24 hours, and their viability measured by a flow cytometry fluorophore exclusion assay (sytox green). Results are displayed as mean± standard deviation (N=4) In panel 1B, cells were treated with 50µg/ml (green curve) or 100µg/ml (blue curve) PLA particles for 24 hours, and the cell-associated fluorescence was measured immediately after exposure (D1), or up to 3 days post-exposure (D2 to D4). Results are displayed as mean± standard deviation (N=4).

This was further confirmed by confocal microscopy, as shown in Fig. 2. Disappearance of the particles and appearance of a fuzzy intracytoplasmic fluorescence was observed, indicating degradation of the particles.

**Figure 2:**
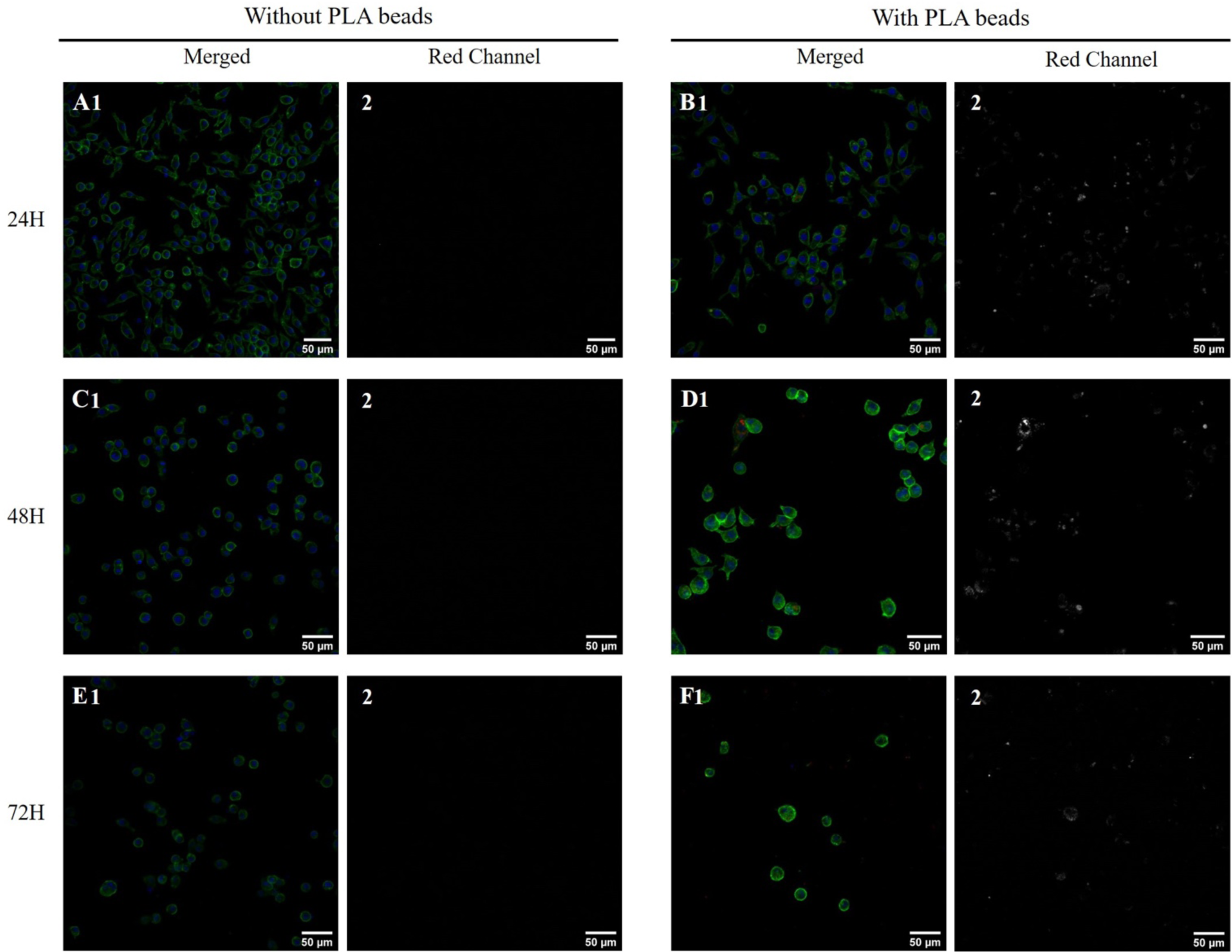
Confocal Microscopy Imaging of J774.A1 Cells. Confocal microscopy images of J774.A1 cells stained with Phalloidin Atto 488 (green) and DAPI (blue) after exposure to PLA beads for 24, 48, and 72 hours. (A1,C1,E1) Show the overlay of actin cytoskeleton (green) and cell nuclei (blue) captured at each time point. (B1,D1,F1) present the overlay of Phalloidin, DAPI staining and PLA beads. The second column of images (A2,B2,C2,D2,E2,F2) represents the red channel, allowing the observation of the PLA beads (white).

Magnification x20

### 3.3. Analysis of the proteomic results

The shotgun proteomic analysis was able to detect and quantify 2869 proteins (**Table S1**). A first global analysis of the complete protein list by principle coordinates analysis showed that the two groups (control and particle-treated) appeared separated on the diagram (**Fig. 3**), indicating significative changes in the proteome, even if the chosen concentration was quite remote of the toxicity threshold. The fact that the two proteomes were significantly different, on a statistical point of view, was further substantiated by an analysis of similarities ^67^, which lead to a Bonferroni-corrected post-hoc p value of 0.009.

**Figure 3:**
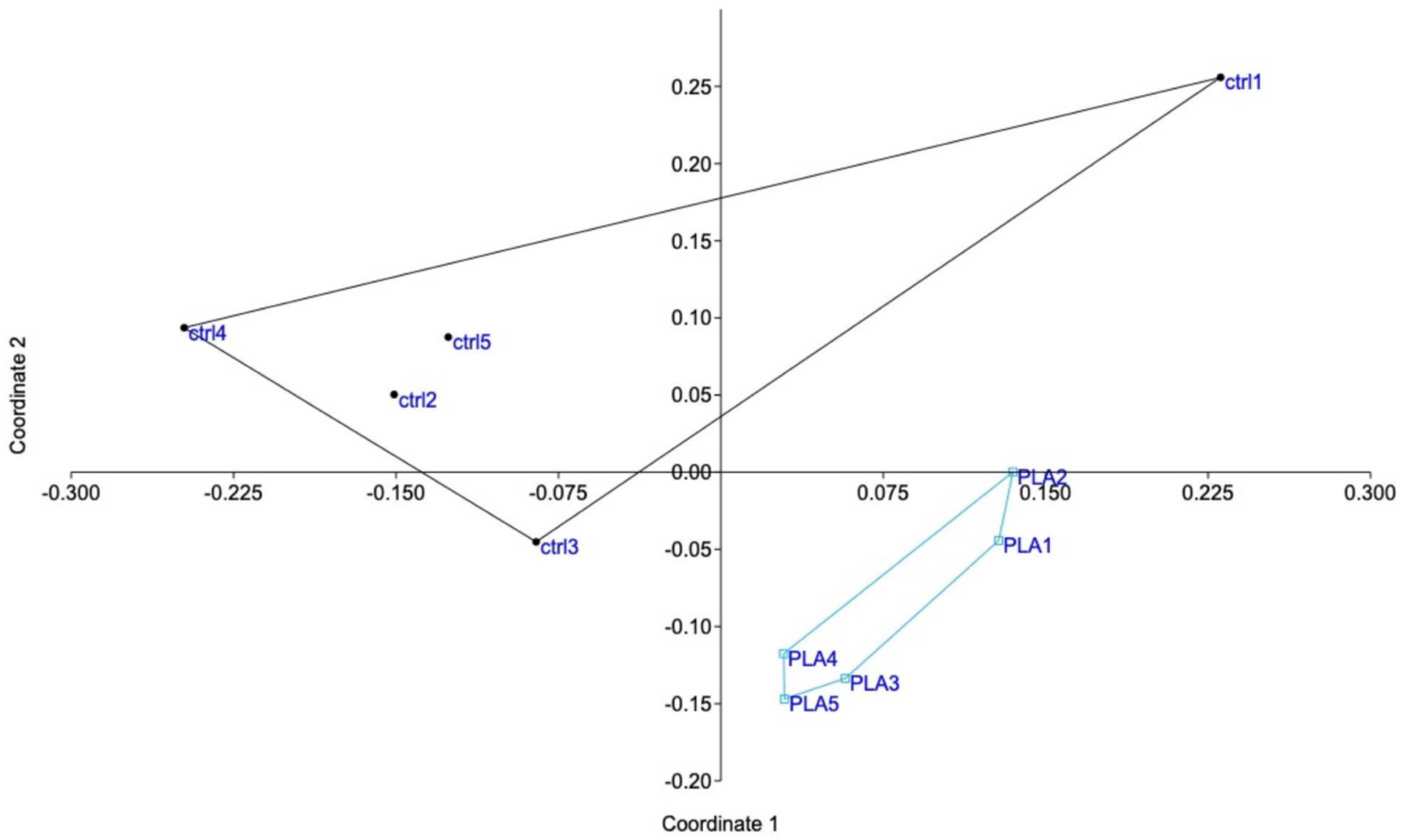
Global analysis of the proteomic data. The complete proteomic data table (2869 proteins) was analysed by Principal Coordinates Analysis, using the PAST software. The mathematical distance used for the calculations was the Gower distance. The results are represented as the X-Y diagram of the first two axes of the Principal Coordinates Analysis, representing 54% of the total variance. Eigenvalue scale.

Proteins modulated by the internalization of PLA particles were selected on the basis of a Mann-Whitney U test ≤ 2 in the comparison of plastic-treated cells compared to unexposed controls. This resulted in the selection of 346 modulated proteins (**Table S2**). In order to gain further insight into the significance of the observed changes, this list of modulated proteins was used to perform pathways analyses by the DAVID software, and the results are shown in **Table S3**. Some of the pathways highlighted by this analysis indicated a global stress response (e.g. translation, nucleotide binding, carbon metabolism, endoplasmic reticulum), which is expected for any cellular stress, while other pathways appeared more specific of cellular internalization of particles (e.g. mitochondria, lysosomes).

As these pathway analysis softwares proceed by aggregation of proteins sharing the same annotations, they require a minimum number of proteins to build a cluster. Thus, the list of modulated proteins was manually combed in addition to this global analysis, in order to retrieve isolated protein changes, which are discarded by pathway analyses but may be of interest in the frame of macrophage physiology. This comprehensive analysis of the proteomic results led us to perform validation experiments on several functions.

### 3.4. Mitochondria and endoplasmic reticulum

The list of the mitochondrial proteins modulated by cells treatment with PLA nanoparticles is presented in **Table S4**, and included 41 proteins, of which 32 were increased in abundance in response to PLA nanoparticles. Among them, 11 subunits of the respiratory chain were highlighted (8 increases, 3 decreases). Although the changes observed in protein abundances were usually of low magnitude for each protein, the changes occurring on several subunits of the same complexes may suggest a functional alteration. To test this hypothesis, we tested the mitochondrial transmembrane potential as a proxy of mitochondrial function. The results, shown in **Fig. 4a**, indicated that PLA particles did not induce any change in the mitochondrial transmembrane potential, while PS particles induced a small but not statistically significant increase in the mitochondrial transmembrane potential, as described previously ^41,42^. Regarding endoplasmic reticulum, 30 proteins were modulated by treatment with PLA nanoparticles, as shown in **Table S5**. These included proteins involved in the endoplasmic reticulum stress response, as the VapB protein (Uniprot Q9QY76) or the tyrosine phosphatase Ptpn1 (Uniprot P35821). This prompted us to test whether endoplasmic reticulum stress response may play a role in PLA nanoparticles toxicity or not. To this purpose we used salubrinal, an inhibitor of the endoplasmic reticulum stress response ^68^, which has been shown to counteract the lethal effects of this stress on cells ^69^. The results, shown in **Fig. 4b**, indicated no effect of salubrinal on cell survival up to treatment with 200µg/ml PLA particles for 24 hours.

**Figure 4:**
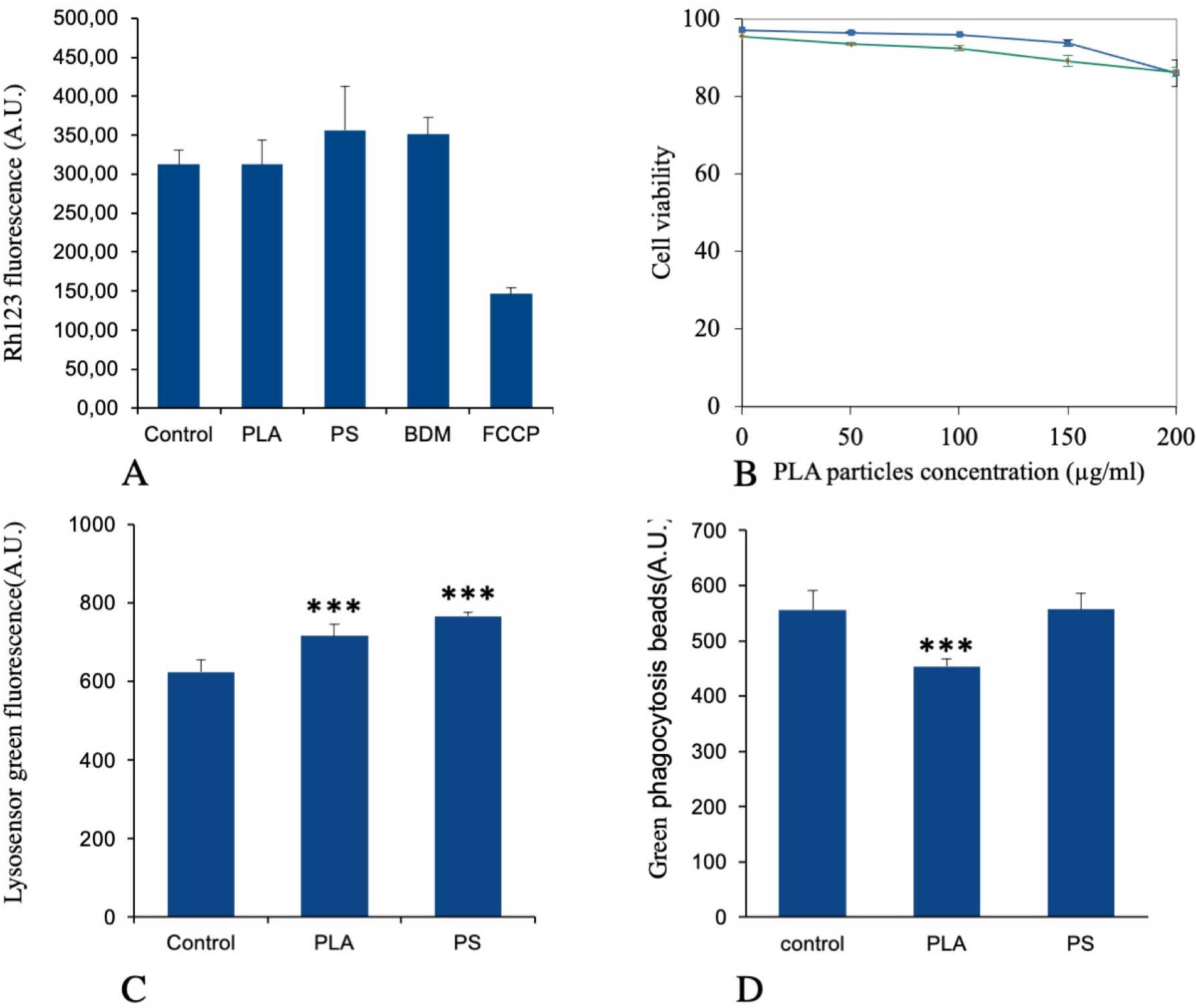
Mitochondria, lysosomes and phagocytosis experiments. For panels A, C and D, the cells were treated for 24 hours with 50µg/ml PLA or PS particles, then various physiological parameters were tested. Panel A: mitochondrial transmembrane potential (rhodamine 123 method). All cells were positive for rhodamine 123 internalization in mitochondria, and the mean fluorescence is the displayed parameter. Results are displayed as mean± standard deviation (N=6) Panel B: test of the endoplasmic stress response in PLA toxicity. Cells were pre-treated with 4µM salubrinal for 4 hours, and various concentrations of PLA beads were then added for a further 18 hours in culture. At the end of the experiment, the cell viability was measured. Results are displayed as survival curves, with the standard deviations at each tested point (N=4). Blue curve: cells untreated with salubrinal. Green curve: salubrinal-treated cells. Panel C: lysosomal proton pumping (Lysosensor method). All cells were positive for lysosensor internalization in lysosomes, and the mean fluorescence is the displayed parameter. Results are displayed as mean± standard deviation (N=6). Significance marks: *** p<0.001 (Student t-test method, comparison between control and each treatment) Panel D: phagocytosis. Cells were first treated for 24 hours with 50µg/ml PLA particles. After removal of the PLA-containing cell culture medium, the cells were treated with green fluorophore labelled carboxylated polystyrene beads for 3 hours. The mean fluorescence, indicating the amount of green beads internalized, is the displayed parameter. Results are displayed as mean± standard deviation (N=6). Significance marks: *** p<0.001 (Student t-test method, comparison between control and each treatment)

### 3.5. Lysosomes, phagocytosis

For this important macrophage function, we gathered the proteins selected under the “lysosome” and “actin cytoskeleton” pathways, to which we added some proteins picked manually on the basis of their annotations in the Uniprot database, such as the two “engulfment and cell motility” proteins (accession numbers Q8BPU7 and Q8BHL5, respectively). This led to a set of 36 proteins, which is presented in **Table S6**. Two of the V-type proton ATPase subunits were present in this list (among the 11 V-type proton ATPase subunits that were detected in the whole proteomic analysis) so that we wanted to investigate by the lysosensor probe the lysosome acidification in response to treatment with PLA nanoplastics, PS nanoparticles being used as a control for non-degradable plastics. The results, shown in **Fig. 4c**, indicated a small but statistically significant (p<0.001) increase in the lysosensor signal, indicating an increase in the number of functional lysosomes and/or an increase proton pumping in the existing lysosomes.

When the phagocytic capacity of plastics-treated cells was probed, as described in **Fig. 4d**, a small (−19%) but significant (p< 0.001) decrease was observed for PLA-treated cells, while PS-treated cells were not altered compared to control cells.

### 3.6. Immunity-related proteins, inflammation

As this function was not detected through the classical pathway analysis, opposite to what was observed for polystyrene particles ^41^, we hand-picked on the basis of Uniprot annotations the proteins that could be associated with this pathway in the shortlist of proteins which expression was significantly modulated in response to exposure to PLA particles. This led to a set of 22 proteins, which is presented in **Table S7**. This relatively low number of proteins may explain why the “innate immunity” annotation was not selected in the classical pathway analysis. Among the selected proteins, several were annotated as regulators of immune response and cytokine secretion (P09581, P10810, P11680, Q60875, Q6P549, Q9D8Y7). This prompted us to investigate the pro-inflammatory response of PLA-exposed macrophages, either in response to PLA alone or to a succession of exposures to PLA and to LPS, by measuring the secretion of TNF-alpha and IL-6. The results, shown in **Fig. 5**, indicated that the LPS-induced IL-6 secretion was decreased close to twofold upon prior exposure to PLA (and PS) particles (**Fig. 5a**) No Il-6 secretion could be measured in response to plastic alone. In the case of TNF, the opposite response was observed. A statistically significant increase could be detected for the basal levels after exposure to plastics alone (**Fig. 5b**) and this increase was more pronounced in response to PLA than to PS. After successive exposure to plastics and LPS, the same trend was observed again, of course at much higher levels (**Fig. 5c**).

**Figure 5:**
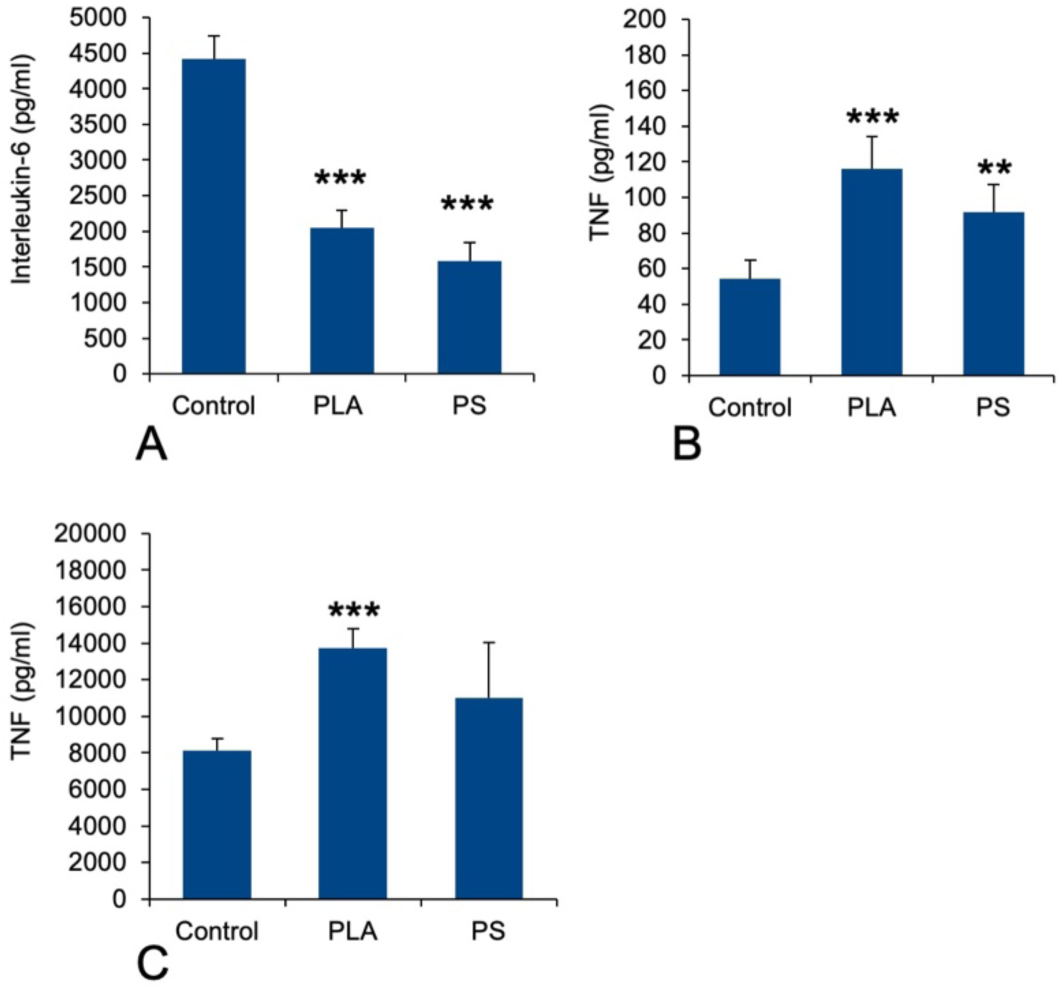
Cytokine release. The cells were first treated for 24 hours with 50µg/ml PLA or PS particles. The medium was then removed and the cells were treated (or not) with 1ng/ml lipopolysaccharide in complete cell culture medium for 24 hours. The cell medium was then collected for secreted TNF and IL-6 measurements. Results are displayed as mean± standard deviation (N=6). Significance marks: *** p<0.001 (Student t-test method, comparison between control and each treatment) Panel A: IL-6 release (after stimulation with LPS) Panel B: TNF-alpha release (without stimulation with LPS) Panel C: TNF-alpha release (after stimulation with LPS)

### 3.7. Redox homeostasis related proteins, oxidative stress

As an increase in oxidative stress was observed in response to polystyrene particles ^42^, we hand-picked on the basis of Uniprot annotations proteins that could be associated with redox metabolism at large (excluding central metabolism) in the shortlist of proteins which expression was significantly modulated in response to exposure to PLA particles. This led to a set of 13 proteins, which is presented in **Table S8**. This relatively low number of proteins may explain why the “redox homeostasis” annotation was not selected in the classical pathway analysis.

However, we decided to investigate the level of cellular oxidative stress in response to exposure to PLA (and PS nanoparticles). The results, shown in **Fig. 6**, indicated no increase in oxidative stress in response to PLA nanoparticles, while a small but significant increase was observed in response to PS nanoparticles, as previously described ^42^.

**Figure 6:**
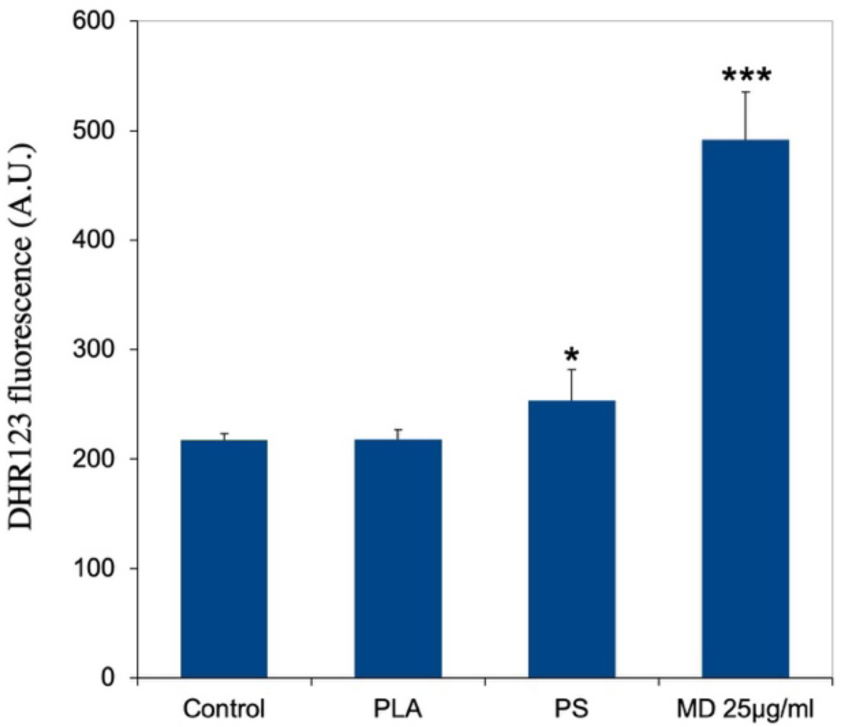
Cellular oxidative stress, measured with the dihydrorhodamine 123 (DHR123) indicator. The cells were exposed to PLA or PS beads for 24 hours, and finally for 20 minutes to the DHR 123 probe. Menadione (25µg/ml for 2 hours) was used as a positive oxidative stress control. Results are displayed as mean± standard deviation (N=6). Significance marks: * p<0.05; *** p<0.001 (Student t-test method, comparison between control and each treatment)

Furthermore, the proteins highlighted in **Table S8** included a few proteins included in quinine metabolism, such as aflatoxin B1 aldehyde reductase member 2, and quinone oxidoreductase. This prompted us to seek for cross toxicity between quinones and plastic beads. To this purpose, we performed comparative cell viability on cells first treated (or not) with nanoplastics, then with variable concentrations of polycyclic quinones. The results, shown on **Fig. 7**, showed an increased toxicity of phenanthrene quinone on plastic-treated cells, while such increased toxicity was not observed with menadione.

**Figure 7:**
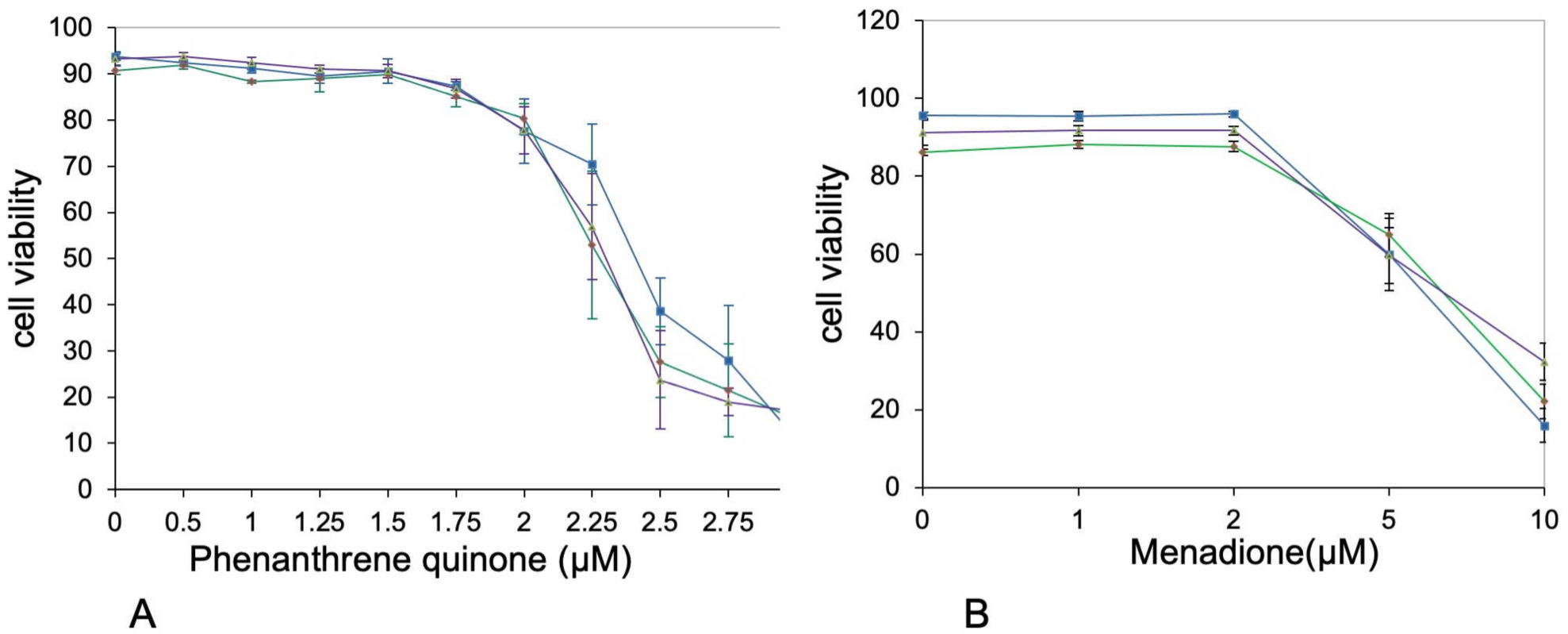
Cross toxicities. The cells were first exposed to PLA or PS beads (50µg/ml). After 6 hours, the secondary toxicant, i.e. 9,10-phenanthrene quinone or menadione, was added at various concentrations and the cells were cultured for an additional 18 hours. After a total of 24 hours, the cell viability was measured. Blue curve: control cells (unexposed to plastics) Green curve: PLA-exposed cells Purple curve: PS-exposed cells Panel A: cross toxicity with 9,10-phenanthrene quinone. The significance of the difference in viability between cells untreated with plastics and plastic-treated cells at 2.25 and 2.5µM 9,10-phenanthrene quinone is p<0.1. Panel B: cross toxicity with menadione. No difference in toxicity between cells untreated with plastics and plastic-treated cells was detected.

## 4. Discussion

The wide release of plastics in the environment is more than suspected to have major deleterious consequences on life. These can be due to mechanical effects of macroplastics ^3^, but microplastics arising from the fragmentation of macroplastics are also known to have deleterious effects, e.g. on oysters ^19^ on seabirds ^4^. Microplastics fragment even further into nanoplastics, which have been associated with liver fibrosis in rodents ^70^, and more recently with increased cardiovascular risk in humans ^71^. These deleterious effects are the indirect consequences of the very low biodegradability of conventional, oil-derived plastics, which makes the particles resulting from their fragmentation long lasting in the environment and within living organisms. Even though this low biodegradability will make this concern a problem for many years to come, one way not to aggravate it would be to switch from poorly-degradable plastics to biodegradable ones, so that the persistent mechanical damage as well as the bioaccumulation issues will be alleviated. Among the few chemical solutions available to produce biodegradable plastics, polymers of organic alkyl hydroxyesters are a promising area of research. Among this class of chemicals, polylactic acid (PLA) represents an interesting choice, as it is both biobased and biodegradable ^50^. However, it is also known to fragment easily into nanoparticles ^56,57^. As the global biodegradability in the environment does not necessarily translate into the absence of accumulation in all cell types, it is necessary to obtain data on the internalization, degradation and effects of PLA nanoparticles on cell types of interest. This is required to figure out if more or less transient effects can be expected on certain cell types. Within the cell types of interest in multicellular organisms, macrophages are important to test, as they are present from invertebrates to vertebrates and play a key role in inflammation, which can be deleterious on a long time frame ^72^.

First of all, we investigated the acute toxicity of PLA nanoparticles toward macrophages. The LD20 proved to be rather high (150µg/ml) i.e. around twice the value observed for polystyrene ^73^. More importantly, PLA particles were actively degraded in macrophages. Half of the fluorescence had disappeared from cells exposed to PLA nanoparticles 48 hours after the end of exposure. As this disappearance of fluorescence, as measured by flow cytometry, implies degradation of the PLA shell of the nanoparticles, then release of the fluorophore within the cell then excretion of the free fluorophore, the real degradation of the PLA shell is probably even faster than this figure. This shows a very active degradation of the PLA particles within macrophages. This shall not come as a surprise, as PLA has been shown to be very sensitive to proteases (which often also have an esterase activity) ^74^, proteases being quite abundant in lysosomes. This easy degradation of PLA by mammalian enzymes may be attributed to the fact that lactic acid is an alpha hydroxy acid, quite close in its structure to the alpha amino acids that make proteins.

We then investigated the effects of the PLA particles on macrophages immediately after exposure using a proteomic screen. Out of 2869 proteins quantified, the proteomic screen highlighted 346 proteins modulated upon treatment with PLA, covering several pathways.

Interestingly, we did not see any modulation of metabolic proteins, although lactate has been shown to induce major metabolic reprogramming ^75^. In the same trend, we did not detect a switch to an inflammatory phenotype, as described when macrophages are confronted with a high lactate concentration ^76^. There can be several explanations to this fact. The first one could just be that immediately after exposure, there is not a sufficient intracellular release of lactate to induce these phenomena. Moreover, as particles are present in the lysosomes and degraded there, a control of the speed of lactate release from the lysosomes to the cytosol may also occur. Finally, regarding the central metabolism, it should be kept in mind that it is heavily regulated by post-translational modifications ^77–79^, which do not appear easily in a global shotgun proteomic screen as in this work.

Nevertheless we detected changes in the abundances of many proteins related to organelles such as mitochondria, lysosomes and endoplasmic reticulum (ER). Regarding ER proteins, on the 30 that showed a significant changes in their abundance, 24 were increased in response to PLA treatment and only 6 decreased. As ER stress may be a factor explaining toxicity, we tested this hypothesis with the drug salubrinal, which is an inhibitor of ER stress ^68^, and has been used to demonstrate the implication of ER stress in toxicity ^69^. As we did not detect any effect of salubrinal in the cell viability of cells treated with up to 200µg/ml PLA, we can conclude that this pathway is not a major determinant of PLA toxicity in macrophages. In the same trend, we could not detect any sign of mitochondrial depolarization after treatment with PLA nanoparticles, although the abundance of more than 40 mitochondrial proteins is changed, with 32 increases and 9 decreases. It must therefore be concluded that the observed changes are homeostatic, and reflect how the cells adapt to the presence of PLA particles.

Regarding lysosomes, 36 proteins were modulated in their abundances, with 23 increases and 13 decreases. Many of the observed decreases are indeed lysosome-associated cytoskeletal proteins, such as tropomyosins, ARP complex subunits, gelsolin and myosin 9. Among the increases were detected two subunits of the lysosomal proton pump. This prompted us to test some lysosomal functions with the lysosensor probe. As neutral red, lysosensor probes are pumped in acidic organelles ^80^. The lysosensor signal thus reflects both the activity of the proton pump and the integrity of the lysosomes, as damaged lysosomes cannot maintain a proton gradient and thus exhibit a reduced lysosensor signal ^81^. We detected indeed a small but significant increase in the intensity of the lysosensor signal for PLA-treated cells, but also for PS-treated cells, suggesting that this effect is not specific for a given nanoparticle but more a generic effect linked to particle internalization per se.

We then tested more specific functions of the macrophage, starting with phagocytosis. To this purpose, we first exposed cells to the nanoplastics of interest, and then to a fluorescently-labelled test bead for a relatively short time, in order to determine whether plastic-treated cells are still active for phagocytosis or not. In this case we observed a specific decrease in the phagocytic activity of PLA-treated beads, which did not occur when the cells were pre-treated with PS beads, as described earlier ^42^. A possible explanation for this phenomenon is the induction of the phosphatase SHIP2 (Q6P549) in response to PLA treatment. This phosphatase is known to decrease Fc-gamma-R-mediated phagocytosis in macrophages ^82^, and the beads that we use to probe phagocytosis are incubated with serum, a process known to lead to opsonization ^83^.

Regarding cytokines production, which is another important specialized function of macrophages, the proteomic analysis provided interesting, and somewhat contradictory, hypotheses. On the one hand some proteins increasing the inflammatory response were increased in their abundance in response to treatment with PLA. Examples are the M-CSF receptor (P09581) which mediates the pro-inflammatory effects of M-CSF, the ARHGEF2 protein (Q60875), which plays a role in the signal transduction in response to peptidoglycans ^84^, and Rab-10, which plays a role in the recycling of TLR4 at the membrane and therefore in the response to LPS ^85^.

On the other hand, some proteins acting directly or indirectly as negative regulators of the inflammatory response were also increased in their abundance in response to treatment with PLA. Examples are the SHIP2 protein (Q6P549), which down regulates the signal transduction pathway downstream of the M-CSF receptor, the Tnfaip8l2 protein (Q9D8Y7), which acts as a negative regulator downstream of some TLRs ^86^, and the TMEM43 protein (Q9DBS1), which acts as a negative regulator of the cGAS/STING interferon response pathway ^87^. Interestingly, the cGAS/STING pathway has been implicated in the PS particles-induced liver inflammation and fibrosis ^70^.

In view of this complex landscape of responses, we decided to perform simple validation experiments in measuring the secretion of the pro-inflammatory cytokines IL-6 and TNF-alpha in response to exposure to PLA or PS nanoparticles, with and without terminal activation with LPS. The results were not consistent between the two cytokines, indicating that their regulations are not strictly similar. For IL-6, we did not detect any secretion without LPS-stimulation, indicating that neither PLA or PS alone induced its secretion. However, when cells were exposed to plastics and then to LPS, an important reduction in IL-6 secretion was observed. This means in turn that plastic-exposed macrophages are less performing in their antibacterial response, compared to plastic-free macrophages. For TNF alpha, the same trend was observed with or without LPS activation, i.e. an increase in TNF secretion upon exposure to plastic, the effect being more pronounced for PLA than for PS. This effect is consistent with the one already described on RAW264.7 ^40^, i.e. another mouse macrophage cell line. These diverging results between IL-6 and TNF-alpha showed how complex the inflammatory response to plastics can be. The important point to be underlined here is that although biodegradable, PLA induces at least a transient pro-inflammatory response at the TNF level.

Finally, we investigated the redox metabolism of plastic-treated macrophages. We started with the ROS production, as overproduction of hydrogen peroxide, which is diffusible, may contribute to damage the adjacent tissues ^88^. In macrophages, hydrogen peroxide is produced by the combined action of NADPH oxidase, which produces superoxide, and superoxide dismutase, which transforms superoxide into hydrogen peroxide. Here again, the proteomic screen revealed inconsistent results. The levels of the NADPH oxidase subunits Ncf1 and Ncf4 were constant, while the level of cytochrome b245 (Q61093), the protein that transfers single electrons to oxygen atoms to produce superoxide increased, and the level of superoxide dismutase decreased. Thus, no clear trend emerged from the proteomic screen. However, targeted validation experiments showed that PLA nanoparticles did not induce any overproduction of ROS in cells, while PS nanoparticles induced a moderate but significant overproduction, as analyzed in more detail previously ^42^. Thus in this regard, PLA particles may induce less oxidative damage to the surrounding tissue than PS particles.

In the proteins grouped under the “redox homeostasis” header, two proteins drew our attention. First the AKR7A2 oxidoreductase (P45376) and then crystallin zeta, also known as the QOR quinone oxidoreductase (P47199). The latter protein is different from the more well-known NQO1 quinone reductase (Q64669). Opposite to NQO1, which catalytic mechanism ensures a two-electron reduction process ^89^, both QOR (which is known to produce semiquinones ^90^) and AKR7A2 are able to induce redox cycling with their preferential substrate phenanthrene 9-10 quinone ^90,91^. This redox cycling is known to induce in turn adverse effects such as cell death and malignant transformation ^92^. We therefore decided to investigate whether treatment of macrophages with PLA could induce a cross-toxicity with phenanthrene 9-10 quinone. As a control, we used menadione, which is not a substrate for at least crystallin zeta ^90^. Although the toxicity shown by phenanthrene 9-10 quinone was characterized by high standard deviations, which decreased the statistical significance of the results, we could observe a differential toxicity of phenanthrene 9-10 quinone in cells treated with plastics (either PLA or PS) and control cells. This differential toxicity was not observed with menadione.

## 5. Conclusions

The first important conclusion of this work on PLA particles is that they are biodegradable within macrophages. However, this does not mean that they do not show immediate consequences on the physiology of macrophages, as shown by the proteome alteration that they induce. Comparison with PS nanoparticles shows that there are both common consequences and specific ones. Among the specific consequences are the inhibition of phagocytosis by the PLA nanoparticles and the absence of detectable oxidative stress. Among the shared ones are the impact on cytokine production, the induction of lysosomal acidification and the cross toxicity with phenanthrene quinone. Phenanthrene quinone is found in combustion products and is also one of the compounds produced by the metabolic activation of phenanthrene ^93^, a combustion product itself. This suggests that some nanoplastics may induce cross-effects with chemicals present in, e.g., diesel exhaust particles or cigarette smoke, which may be another mechanism underlying noxious effects of nanoplastics.

## Author Contributions

VCF: methodology, investigation. MV: methodology, investigation, writing (review and editing). HD: methodology, investigation, writing (review and editing). SC: resources, funding acquisition, writing (review and editing). ED: formal analysis, writing (review and editing). TR: conceptualization, formal analysis, funding acquisition, writing (original draft).

## Funding

This work used the flow cytometry facility and the microscopy facility MuLife of IRIG/DBSCI, funded by CEA Nanobio and by GRAL LabEX (ANR-10-LABX-49-01), a project of the University Grenoble Alpes graduate school (Ecoles Universitaires de Recherche) CBH-EUR-GS (ANR-17-EURE-0003), as well as the platforms of the French Proteomic Infrastructure (ProFI) project (grant ANR-10-INBS-08-03).

This work was carried out in the frame of the PlasticHeal project, which has received funding from the European Union’s Horizon 2020 research and innovation programme under grant agreement No. 965196.

This work was also supported by the ANR Plastox project (grant ANR-21-CE34-0028-04)

## Conflict of Interest

There are no conflicts of interest to declare

## Supporting information

supplemental methods (shotgun proteomic analysis)

## Notes

https://www.doi.org/10.6019/PXD048664

https://www.ebi.ac.uk/biostudies/studies/S-BSST1470

### Competing Interest Statement

The authors have declared no competing interest.

