## supplemental Table 1 for "Biobased, Biodegradable but not bio-neutral: about the effects of polylactic acid nanoparticles on macrophages"

Supplementary Table 1: list of proteins detected and quantified by the proteomic screen

| acc number | gene_name | description | protein_set _score | coverage | MW | #peptides | abundance_Ct | abundance_ctr | abundance_Ct | abundance_Ct | abundance_Ct | abundance_PL | abundance_PL | abundance_PL | abundance_PL | abundance_PL | U PLA | ratio PLA/ctrl |
|---|---|---|---|---|---|---|---|---|---|---|---|---|---|---|---|---|---|---|
| A0JPM9 | Eif3j | Eukaryotic translation initiation factor 3 subunit J (Rattus norvegicus) | 149.93 | 12.74 | 29187 | 3 | 5504758 | 1521614.38 | 829474.25 | 4087191.5 | 4690523 | 4207197.5 | 4031471 | 3498181 | 3448141.5 | 3997800.5 | 11 | **1.153** |
| A0JPP1 | Drap1 | Dr1-associated corepressor (Rattus norvegicus) | 124.11 | 11.22 | 22326 | 3 | 3629375.25 | 2162194 | 1640813 | 1449228.5 | 1138232.25 | 2166073.5 | 2607142.5 | 2245114.5 | 2567565.5 | 2134798.5 | 6 | **1.170** |
| A0MZ67 | Shtn1 | Shootin-1 (Rattus norvegicus) | 143.66 | 4.74 | 71446 | 3 | 2208127 | 2488481.5 | 2873893.5 | 2911975.5 | 1834460.12 | 2603137.5 | 2177412.5 | 2408826.5 | 640198.56 | 3104386.25 | 11 | **0.888** |
| A1A5S1 | Prpf6 | Pre-mRNA-processing factor 6 (Rattus norvegicus) | 83.66 | 2.34 | 106764 | 2 | 0 | 1905176.25 | 2008816.38 | 1715037.88 | 1009300 | 1476328.5 | 1410347.75 | 1740729.75 | 1135194.75 | 815394.5 | 10 | **0.991** |
| A1EC95 | Heatr6 | HEAT repeat-containing protein 6 (Rattus norvegicus) | 44.46 | 0.96 | 136505 | 1 | 0 | 0 | 123824.69 | 0 | 361638.34 | 385905.12 | 422063.59 | 213089.27 | 216220.89 | 264684.41 | 6 | **3.094** |
| A1L134 | Aup1 | Ancient ubiquitous protein 1 (Rattus norvegicus) | 88.37 | 9.51 | 46199 | 2 | 781307.38 | 1087021.12 | 1673866.62 | 381025.06 | 173095.19 | 1451203.25 | 969692.12 | 1315251.5 | 1267189.5 | 1138123 | 6 | **1.499** |
| A1L314 | Mpeg1 | Macrophage-expressed gene 1 protein (Mus musculus) | 922.75 | 40.81 | 78390 | 17 | 41420120 | 38384404 | 40366676 | 35873584 | 34609164 | 47871508 | 52164452 | 53467204 | 46966720 | 50276012 | 0 | **1.315** |
| A2A432 | Cul4b | Cullin-4B (Mus musculus) | 321.79 | 8.04 | 110699 | 6 | 4197291 | 4922409.5 | 6130902.5 | 4662547.5 | 3195784 | 5092460 | 4864116.5 | 6137487 | 5810568 | 6197554 | 4 | **1.216** |
| A2A5R2 | Arfgef2 | Brefeldin A-inhibited guanine nucleotide-exchange protein 2 (Mus musculus) | 47.35 | 1.28 | 202240 | 1 | 0 | 263307.47 | 297531.88 | 208591.09 | 224647.02 | 0 | 0 | 239591.3 | 300679.34 | 257578.52 | 14 | **0.803** |
| A2AAY5 | Sh3pxd2b | SH3 and PX domain-containing protein 2B (Mus musculus) b | 133.49 | 3.08 | 101517 | 3 | 6033651 | 5347541.5 | 5826108 | 5066451 | 4744459 | 6913103 | 6087302 | 4592920.5 | 4224098.5 | 3639960.5 | 10 | **0.942** |
| A2ADY9 | Ddi2 | Protein DDI1 homolog 2 (Mus musculus) | 101.06 | 8.27 | 44591 | 2 | 3162293 | 1362458.62 | 1476211.5 | 2504601.75 | 2573145 | 895893.56 | 970992.19 | 1839460.38 | 1942067.38 | 2428644.25 | 6 | **0.729** |
| A2AGH6 | Med12 | Mediator of RNA polymerase II transcription subunit 12 (Mus musculus) | 53 | 0.37 | 244561 | 1 | 0 | 567538.62 | 515575.91 | 0 | 0 | 534547.88 | 549878.06 | 460524.94 | 557430.31 | 0 | 13 | **1.941** |
| A2AGT5 | Ckap5 | Cytoskeleton-associated protein 5 (Mus musculus) | 492.92 | 5.51 | 225635 | 10 | 8627341 | 7451486 | 6734907 | 7413412 | 5305783.5 | 6673925.5 | 6524361 | 7189875 | 7479148 | 7598155.5 | 12 | **0.998** |
| A2AIV8 | Card9 | Caspase recruitment domain-containing protein 9 (Mus musculus) | 70.85 | 1.87 | 62462 | 1 | 684006.5 | 376365.88 | 320576.25 | 0 | 213718.61 | 437209.56 | 595201.25 | 446443.06 | 519830.84 | 627632.19 | 5 | **1.647** |
| A2AKQ0 | Slc35d1 | UDP-glucuronic acid/UDP-N-acetylgalactosamine transporter (Mus musculus) | 40.94 | 5.92 | 39272 | 1 | 208064.23 | 134456.62 | 37279.64 | 0 | 0 | 121531.62 | 144911.53 | 105306.96 | 0 | 0 | 15 | **0.979** |
| A2AN08 | Ubr4 | E3 ubiquitin-protein ligase UBR4 (Mus musculus) | 696.8 | 3.38 | 572290 | 13 | 6315274.5 | 7720871.5 | 6704901.5 | 5454841 | 5373147 | 8515579 | 7860598.5 | 9443227 | 8384951 | 6094102.5 | 3 | **1.277** |
| A2AWP8 | Arhgef10l | Rho guanine nucleotide exchange factor 10-like protein (Mus musculus) 0l | 49.66 | 1.25 | 139987 | 1 | 109898.29 | 189148.44 | 134745.66 | 169324.16 | 196983.2 | 159068.23 | 232805.98 | 170743.42 | 176425.86 | 118697.96 | 11 | **1.072** |
| A2BDX3 | Mocs3 | Adenylyltransferase and sulfurtransferase MOCS3 (Mus musculus) | 80.54 | 4.57 | 49375 | 2 | 469940.44 | 820130.06 | 892049.38 | 750048.31 | 731808.94 | 546488.12 | 733885.56 | 0 | 420310.22 | 1025124.81 | 8 | **0.744** |
| A2RRU1 | Gys1 | Glycogen [starch] synthase, muscle (Rattus norvegicus) | 110.76 | 6.5 | 84072 | 3 | 1742992 | 1701412.75 | 1929784.75 | 2413375.25 | 2952200 | 1932299.38 | 1887582.25 | 2818227.5 | 2791148.5 | 2732649 | 8 | **1.132** |
| A2RSY6 | Trmt1l | TRMT1-like protein (Mus musculus) | 226.06 | 8.52 | 80861 | 4 | 2723967.75 | 3806853 | 3989451 | 3607882 | 905068.06 | 3622147.25 | 3010241.75 | 3589017.75 | 3591607.5 | 3511317 | 11 | **1.152** |
| A2RUW1 | Tollip | Toll-interacting protein (Rattus norvegicus) | 132.23 | 13.14 | 30315 | 3 | 4837403.5 | 3774933.5 | 4771638 | 3652738 | 4043455 | 5195159 | 5005635 | 5068564 | 4764839 | 4542623.5 | 4 | **1.166** |
| A2VCX1 | Tiprl | TIP41-like protein (Rattus norvegicus) | 227.54 | 22.88 | 31225 | 5 | 1210124 | 2238762 | 2583513.5 | 1818904 | 3185867.5 | 2008129.25 | 1259619.88 | 2287510.25 | 1719261.25 | 1239240.38 | 8 | **0.771** |
| A3KGS3 | Ralgapa2 | Ral GTPase-activating protein subunit alpha-2 (Mus musculus) 2 | 167.92 | 1.44 | 210288 | 3 | 144726.97 | 1022570.12 | 1257880 | 813322 | 762810.06 | 868635.5 | 869574.06 | 1272496.5 | 964232.75 | 870083.69 | 8 | **1.211** |
| A4L9P7 | Pds5a | Sister chromatid cohesion protein PDS5 homolog A (Rattus norvegicus) | 131.05 | 1.95 | 150286 | 2 | 455915.78 | 377345.12 | 0 | 1341187.25 | 501383.56 | 312126.91 | 715968.94 | 212828.5 | 619461.5 | 201749.52 | 11 | **0.771** |
| A6H5Z3 | Exoc6b | Exocyst complex component 6B (Mus musculus) | 90.21 | 2.59 | 94130 | 2 | 1032259.88 | 1252740.12 | 1046875.19 | 860729.19 | 1031610.25 | 886875.56 | 827808.88 | 849894.56 | 1193454.62 | 996921.12 | 6 | **0.910** |
| A6H8H2 | Dennd4c | DENN domain-containing protein 4C (Mus musculus) | 107.5 | 1.26 | 211458 | 2 | 519606.59 | 671318 | 765408 | 242725.09 | 762200.5 | 902250.75 | 0 | 892104.62 | 887758.5 | 817230 | 5 | **1.182** |
| A6X935 | Itih4 | Inter alpha-trypsin inhibitor, heavy chain 4 (Mus musculus) | 58.31 | 1.91 | 104659 | 2 | 19059844 | 21073892 | 19697220 | 33472202 | 9191843 | 14372952 | 14025199 | 11169314 | 5921494 | 13458838 | 4 | **0.575** |
| B0BN18 | Pfdn2 | Prefoldin subunit 2 (Rattus norvegicus) | 232.48 | 30.52 | 16580 | 4 | 7302642.5 | 3788149.5 | 3715274 | 6017942.5 | 6887994 | 5474323 | 4750602 | 4526944.5 | 4852866 | 4827677 | 10 | **0.882** |
| B0BN56 | Mrps31 | 28S ribosomal protein S31, mitochondrial (Rattus norvegicus) | 46.6 | 5.94 | 43962 | 1 | 29475.67 | 0 | 0 | 64959.22 | 65945.13 | 198806.78 | 0 | 0 | 0 | 0 | 19 | **1.240** |
| B0BNG0 | Emc2 | ER membrane protein complex subunit 2 (Rattus norvegicus) | 95.56 | 8.42 | 34870 | 2 | 3250350.25 | 1315664.88 | 3056724 | 1354164.62 | 2617846.5 | 2586661.75 | 2356335.75 | 2965753.5 | 2693221.5 | 2826033 | 12 | **1.158** |
| B0BNM9 | GLTP | Glycolipid transfer protein (Rattus norvegicus) | 138.16 | 16.75 | 23704 | 3 | 2021334.25 | 1165056.38 | 711667.12 | 552274.25 | 845206.19 | 1677691.75 | 1996504.25 | 1875775.25 | 1583353.75 | 1604889.38 | 5 | **1.650** |
| B1ARD6 | Slfn9 | Schlafen family member 9 (Mus musculus) | 132.07 | 2.53 | 103992 | 2 | 937568.25 | 4150037.75 | 5257197 | 4408678 | 4076805.5 | 3232252.5 | 3823063.5 | 3878379.25 | 2758091 | 3609545.5 | 5 | **0.919** |
| B1AZI6 | Thoc2 | THO complex subunit 2 (Mus musculus) | 445.41 | 6.71 | 182773 | 9 | 4391891.5 | 5312444 | 4400884.5 | 2753359.5 | 3069123.5 | 5171351 | 4704745 | 4654073 | 5119213.5 | 4626623 | 5 | **1.218** |
| B2GV87 | Ptpre | Receptor-type tyrosine-protein phosphatase epsilon (Rattus norvegicus) | 45.13 | 1.86 | 80734 | 1 | 1130364.75 | 739373.62 | 891582.12 | 603119.25 | 122826.52 | 941525.38 | 1066123.5 | 386130.75 | 1231133.75 | 995526.88 | 7 | **1.325** |
| B2RQC6 | Cad | CAD protein (Mus musculus) | 1674.28 | 19.55 | 243238 | 29 | 34312688 | 36805200 | 40667672 | 43969188 | 34691472 | 42503040 | 37344564 | 42465540 | 39242860 | 41035380 | 7 | **1.064** |
| B2RRE7 | Otud4 | OTU domain-containing protein 4 (Mus musculus) | 48 | 0.72 | 123055 | 1 | 681377.38 | 549618 | 481957.41 | 688516.25 | 670655.5 | 641911.44 | 835534.44 | 837077.38 | 526395.62 | 590910.94 | 10 | **1.117** |
| B2RWS6 | Ep300 | Histone acetyltransferase p300 (Mus musculus) | 43.74 | 0.75 | 263305 | 1 | 125811.41 | 315285.03 | 216869 | 211540.09 | 261833.62 | 0 | 0 | 208860.36 | 267428.16 | 185023.38 | 7 | **0.585** |
| B2RX12 | Abcc3 | Canalicular multispecific organic anion transporter 2 (Mus musculus) | 316.69 | 6.76 | 169124 | 7 | 2740816.75 | 3401197.5 | 3081148.75 | 3256246.5 | 1968142.88 | 2793716.5 | 3656663.75 | 3504520.5 | 3574254.25 | 3473836.75 | 3 | **1.177** |
| B2RXC1 | Trappc11 | Trafficking protein particle complex subunit 11 (Mus musculus) 1 | 93.67 | 2.56 | 128396 | 2 | 1281884.75 | 804301.69 | 686682.25 | 130987.43 | 800756.62 | 1015746.38 | 610377.88 | 708656.25 | 771227.62 | 626427.69 | 10 | **1.008** |
| B2RXR6 | Ankrd44 | Serine/threonine-protein phosphatase 6 regulatory ankyrin repeat subunit B (Mus musculus) | 81.17 | 1.51 | 107385 | 1 | 957390.56 | 1115858.38 | 1204737.88 | 1416866.25 | 1132155.12 | 967845.12 | 1057927.5 | 1089370.88 | 1110661.25 | 942554 | 4 | **0.887** |
| B2RXS4 | Plxnb2 | Plexin-B2 (Mus musculus) | 828.13 | 12.87 | 206230 | 17 | 10504924 | 10259854 | 12093153 | 12613137 | 11380884 | 12596358 | 12259158 | 14630655 | 11918932 | 14995643 | 4 | **1.168** |
| B2RY56 | Rbm25 | RNA-binding protein 25 (Mus musculus) | 285.02 | 9.9 | 99552 | 6 | 10320339 | 10325358 | 10371586 | 8962380 | 11129582 | 6843760.5 | 9884895 | 9840864 | 11026370 | 10400348 | 10 | **0.939** |
| B2RYG6 | Otub1 | Ubiquitin thioesterase OTUB1 (Rattus norvegicus) | 680.92 | 58.67 | 31270 | 11 | 13419175 | 27688568 | 34040992 | 25481080 | 23725286 | 27184202 | 26647230 | 28075154 | 29667892 | 30684912 | 7 | **1.144** |
| B2RYI0 | Wdr91 | WD repeat-containing protein 91 (Rattus norvegicus) | 134.36 | 3.08 | 83146 | 2 | 1550099.75 | 1262201.5 | 1048265.69 | 1191137.25 | 1577575.12 | 1490716.25 | 1089008.88 | 1464716.88 | 1080846.75 | 1306821.5 | 11 | **0.970** |
| B3GNI6 | Sept11 | Septin-11 (Rattus norvegicus) | 248.29 | 15.55 | 49695 | 5 | 3600248.25 | 4369434.5 | 4253937.5 | 2959582 | 4025761.5 | 2883788.5 | 3959114.5 | 4832458 | 3518860.75 | 3201707.25 | 9 | **0.958** |
| B5DEN9 | Vps28 | Vacuolar protein sorting-associated protein 28 homolog (Rattus norvegicus) | 61.14 | 6.58 | 26110 | 1 | 2709986.5 | 890031 | 1067848.62 | 865771.38 | 946301.19 | 690484.19 | 615722 | 649359.75 | 622875.38 | 736974.12 | 0 | **0.512** |
| B5DF89 | Cul3 | Cullin-3 (Rattus norvegicus) | 321.13 | 10.55 | 88932 | 5 | 226297.42 | 2076801 | 1878366 | 3441296 | 2453515.5 | 3490766.5 | 2927802 | 2108823.5 | 3368850.5 | 2165689.5 | 6 | **1.396** |
| B5DF91 | Elavl1 | ELAV-like protein 1 (Rattus norvegicus) | 384.1 | 21.78 | 36169 | 6 | 32706928 | 19577440 | 21154428 | 19854178 | 20074860 | 26620532 | 27138722 | 19155394 | 21752068 | 20752228 | 10 | **1.018** |
| B5DFC8 | Eif3c | Eukaryotic translation initiation factor 3 subunit C (Rattus norvegicus) | 1088.69 | 18.66 | 105435 | 18 | 66018952 | 54116384 | 57920100 | 53691692 | 49902720 | 64122088 | 58314580 | 57566404 | 61901356 | 59313208 | 6 | **1.069** |
| B5DFN2 | Ahcyl1 | S-adenosylhomocysteine hydrolase-like protein 1 (Rattus norvegicus) | 402.27 | 19.46 | 53753 | 8 | 6521812 | 7753339.5 | 6880196 | 5358355.5 | 5448415 | 6543600 | 6173019 | 6632335 | 6206754 | 8122431.5 | 10 | **1.054** |
| B7ZMP1 | Xpnpep3 | Xaa-Pro aminopeptidase 3 (Mus musculus) | 70.75 | 3.36 | 56677 | 1 | 444498.88 | 519460.75 | 522980.22 | 380251.56 | 409459.31 | 439517.28 | 495489.03 | 416509.28 | 542852.19 | 458142.84 | 10 | **1.033** |
| B9EJ86 | Osbpl8 | Oxysterol-binding protein-related protein 8 (Mus musculus) | 952.48 | 25.08 | 101269 | 16 | 17439232 | 15670853 | 17732248 | 14771491 | 16407440 | 22990444 | 18855972 | 22578428 | 20475144 | 19126650 | 0 | **1.268** |
| B9EKI3 | Tmf1 | TATA element modulatory factor (Mus musculus) | 42.29 | 1.37 | 121803 | 1 | 676838.5 | 842472.69 | 603472.94 | 1000464.06 | 903099.94 | 818315.62 | 1145150.75 | 702854.12 | 720230.38 | 734617.69 | 12 | **1.024** |
| C0HKD8 | Mfap1a | Microfibrillar-associated protein 1A (Mus musculus) | 60.42 | 3.19 | 51954 | 1 | 1428921.12 | 1120158 | 1438291.75 | 790745.12 | 1190402.5 | 1623167.62 | 1121182.75 | 1094464.25 | 1190081.25 | 991804.31 | 11 | **1.009** |
| C0HKE1 | Hist1h2ab | Histone H2A type 1-B (Mus musculus) ab | 380.32 | 63.08 | 14135 | 6 | 1041337600 | 288229824 | 388707040 | 108272368 | 343947168 | 765011840 | 679616896 | 599502912 | 651189248 | 626936640 | 5 | **1.531** |
| D0QMC3 | Mndal | Myeloid cell nuclear differentiation antigen-like protein (Mus musculus) | 229.81 | 8.55 | 60526 | 4 | 6582555.5 | 10480319 | 12134507 | 9432182 | 8788045 | 10039328 | 9372165 | 9574838 | 9136640 | 9529092 | 12 | **1.005** |
| D2XV59 | Gtpbp1 | GTP-binding protein 1 (Rattus norvegicus) | 214.64 | 9.42 | 72489 | 4 | 2288624.5 | 16943914 | 15297326 | 12083438 | 13279657 | 2331594.5 | 2129651.5 | 14954381 | 12170598 | 15085469 | 9 | **0.779** |
| D3YXK2 | Safb | Scaffold attachment factor B1 (Mus musculus) | 433.3 | 11.1 | 105104 | 8 | 19626642 | 17518570 | 11680603 | 16606633 | 14913515 | 14594501 | 8662833 | 15123393 | 15679690 | 15573105 | 7 | **0.867** |
| D3Z6Q9 | Bin2 | Bridging integrator 2 (Mus musculus) | 438.08 | 19.84 | 52554 | 8 | 5279807 | 6133981.5 | 6651461 | 123010016 | 124643768 | 6313610.5 | 7796790.5 | 9300609 | 8235834 | 8656041 | 11 | **0.152** |
| D3Z7P3 | Gls | Glutaminase kidney isoform, mitochondrial (Mus musculus) | 522.54 | 16.47 | 73964 | 9 | 8127890.5 | 7357695.5 | 8832253 | 8113201 | 8586824 | 7856163 | 6782478 | 8352695.5 | 8161045.5 | 8069847 | 8 | **0.956** |
| D3ZBE5 | Nek7 | Serine/threonine-protein kinase Nek7 (Rattus norvegicus) | 79.27 | 7.95 | 34529 | 2 | 4272802 | 2799211.5 | 3591407.25 | 2996215.5 | 3107011 | 3976556 | 3923384.25 | 3918337.25 | 3768186 | 4133117.75 | 5 | **1.176** |
| D3ZC96 | Ttc39b | Tetratricopeptide repeat protein 39B (Rattus norvegicus) | 47.22 | 1.78 | 70168 | 1 | 666408.25 | 709636.88 | 738484.62 | 328159.34 | 0 | 509231.31 | 742869.62 | 748933.25 | 787043 | 739262.75 | 3 | **1.444** |
| D3ZCL3 | Snrpc | U1 small nuclear ribonucleoprotein C (Rattus norvegicus) | 81.24 | 13.21 | 17364 | 2 | 4349829 | 1525571.12 | 837407.56 | 0 | 3743401.5 | 3403446 | 4082432 | 4209325 | 4792808.5 | 4313347 | 5 | **1.989** |
| D3ZDK7 | Pgp | Glycerol-3-phosphate phosphatase (Rattus norvegicus) | 260.2 | 18.69 | 34601 | 4 | 6712070 | 4534441.5 | 800992.25 | 3021625.25 | 5365149 | 4136812.75 | 4876112.5 | 4355517.5 | 4563100 | 3288934 | 12 | **1.038** |
| D3ZKD3 | Alkbh5 | RNA demethylase ALKBH5 (Rattus norvegicus) | 54.01 | 2.78 | 44453 | 1 | 1486394.25 | 1482609.62 | 1513203.5 | 1749510.88 | 754244.81 | 1805046.62 | 1511494.5 | 1713156.75 | 1944836.62 | 2107734.5 | 3 | **1.300** |
| D3ZLZ7 | Impdh1 | Inosine-5'-monophosphate dehydrogenase 1 (Rattus norvegicus) | 126.48 | 7.2 | 55293 | 3 | 1146596.38 | 1093065.12 | 293664.09 | 812097.5 | 1102323 | 927764.75 | 1090324.88 | 699838.38 | 914008.94 | 804930.38 | 8 | **0.998** |
| D3ZTX0 | Tmed7 | Transmembrane emp24 domain-containing protein 7 (Rattus norvegicus) | 226.09 | 27.43 | 25479 | 5 | 2126612 | 7326040 | 5253437 | 8387936 | 4423989 | 5606112 | 3929140.5 | 6390063.5 | 6092528 | 6382606 | 12 | **1.032** |
| D3ZYR1 | Fcho2 | F-BAR domain only protein 2 (Rattus norvegicus) | 48.7 | 2.97 | 88712 | 1 | 0 | 685786.5 | 521718.19 | 723607.69 | 550825.75 | 0 | 0 | 828123.62 | 636714.19 | 611154.38 | 14 | **0.836** |
| D4A208 | Srgap2 | SLIT-ROBO Rho GTPase-activating protein 2 (Rattus norvegicus) | 375.76 | 6.91 | 120878 | 7 | 4558247 | 4084222.25 | 3907515.5 | 6070298 | 4629985.5 | 4945878 | 6315725 | 7336522 | 6473749 | 6845962 | 1 | **1.373** |
| D4A2Z8 | Dhx36 | ATP-dependent DNA/RNA helicase DHX36 (Rattus norvegicus) | 132.56 | 2.4 | 113843 | 2 | 2101936.5 | 1331210.38 | 1407370.88 | 7369266.5 | 1591809.12 | 1471741.75 | 1963684.12 | 1961429.62 | 1421615 | 1718857.88 | 12 | **0.619** |
| D4A4T9 | Chordc1 | Cysteine and histidine-rich domain-containing protein 1 (Rattus norvegicus) | 320.13 | 28.7 | 37362 | 6 | 7835527.5 | 7730264.5 | 8971823 | 11056443 | 11220126 | 9617487 | 12101152 | 11621472 | 10524760 | 11451546 | 4 | **1.182** |
| D4AE41 | Rbmxl1 | RNA binding motif protein, X-linked-like-1 (Rattus norvegicus) | 354.06 | 19.33 | 42250 | 6 | 9483069 | 13205487 | 8633114 | 14336086 | 9781665 | 7158564.5 | 5755399.5 | 5660710.5 | 6195424.5 | 6983642.5 | 0 | **0.573** |
| E9PVA8 | Gcn1 | eIF-2-alpha kinase activator GCN1 (Mus musculus) | 707.07 | 7.49 | 293021 | 15 | 5347299 | 8991495 | 7331956.5 | 9634840 | 6482552 | 7491262 | 6844023.5 | 8684962 | 6973270 | 7708436 | 12 | **0.998** |
| E9PVX6 | Mki67 | Proliferation marker protein Ki-67 (Mus musculus) | 1393.63 | 11.61 | 350864 | 27 | 37926544 | 27105856 | 35640560 | 22549970 | 30168000 | 32879794 | 32535742 | 28762954 | 32373688 | 34255856 | 11 | **1.048** |
| E9Q555 | Rnf213 | E3 ubiquitin-protein ligase RNF213 (Mus musculus) | 719.92 | 3.94 | 584784 | 15 | 11124430 | 9763944 | 8335191.5 | 8254021.5 | 10206034 | 10826404 | 9327232 | 10433300 | 10828683 | 10336492 | 7 | **1.085** |
| E9Q5C9 | Nolc1 | Nucleolar and coiled-body phosphoprotein 1 (Mus musculus) | 40.44 | 1.71 | 73698 | 1 | 2257599.5 | 1078538.88 | 1291335.25 | 379989.94 | 1138884.25 | 1187488.88 | 1763347 | 1255459.62 | 1285523.38 | 1078103.88 | 11 | **1.069** |
| E9Q634 | Myo1e | Unconventional myosin-Ie (Mus musculus) | 854.97 | 18.43 | 126818 | 16 | 17856690 | 15635516 | 19211716 | 13847555 | 16946492 | 23137486 | 21423142 | 24289554 | 23303390 | 24706076 | 0 | **1.400** |
| E9Q7E2 | Arid2 | AT-rich interactive domain-containing protein 2 (Mus musculus) | 48 | 0.49 | 195988 | 1 | 894495.12 | 656693.38 | 513883.09 | 0 | 0 | 854832.12 | 816365 | 832477.31 | 0 | 658733.88 | 10 | **1.531** |
| E9Q7G0 | Numa1 | Nuclear mitotic apparatus protein 1 (Mus musculus) | 806.23 | 10.17 | 235630 | 15 | 14788139 | 10622805 | 9797877 | 10116118 | 12472913 | 10383432 | 12519295 | 10442732 | 12270817 | 11128369 | 11 | **0.982** |
| E9Q8D0 | Dnajc21 | DnaJ homolog subfamily C member 21 (Mus musculus) | 62.01 | 2.45 | 61735 | 1 | 838275.38 | 435405.62 | 586940.81 | 0 | 671869.38 | 0 | 304406.97 | 406047.25 | 394398 | 0 | 6 | **0.436** |
| E9Q9A9 | Oas2 | 2'-5'-oligoadenylate synthase 2 (Mus musculus) | 41.29 | 1.75 | 85051 | 1 | 0 | 293288.94 | 301098.88 | 285091.66 | 379468.44 | 0 | 0 | 190916.14 | 201204.8 | 207952.34 | 6 | **0.477** |
| E9QAM5 | Helz2 | Helicase with zinc finger domain 2 (Mus musculus) | 56.65 | 1.12 | 331563 | 2 | 585997.88 | 449945.25 | 417255.88 | 0 | 0 | 457810.62 | 609384.31 | 195252.39 | 0 | 0 | 15 | **0.869** |
| F1LNJ2 | Snrnp200 | U5 small nuclear ribonucleoprotein 200 kDa helicase (Rattus norvegicus) 0 | 1743.73 | 22.67 | 244875 | 31 | 25023626 | 30036780 | 31213572 | 34415444 | 37526704 | 26935866 | 27889696 | 29564602 | 28950044 | 35189380 | 8 | **0.939** |
| F1LP64 | Trip12 | E3 ubiquitin-protein ligase TRIP12 (Rattus norvegicus) | 429.46 | 5.38 | 223928 | 8 | 3318712.25 | 8124399 | 6910472 | 8063329 | 5345267.5 | 6472792.5 | 6498261.5 | 6866717.5 | 6938921 | 6659937 | 11 | **1.053** |
| F1LQ48 | Hnrnpl | Heterogeneous nuclear ribonucleoprotein L (Rattus norvegicus) | 1228.89 | 61.32 | 67903 | 22 | 58479396 | 57385048 | 53120976 | 58498020 | 50014416 | 55152688 | 52417752 | 43415528 | 44222116 | 44747424 | 3 | **0.865** |

| | | | | | | | | | | | | | | | | | |
|---|---|---|---|---|---|---|---|---|---|---|---|---|---|---|---|---|---|
| F1LSG8 | Vps50 | Syndetin (Rattus norvegicus) | 142.95 | 2.7 | 111188 | 2 | 755557.44 | 555829.44 | 492933.75 | 0 | 0 | 123499.7 | 566431.31 | 802605.69 | 831686.12 | 596753.69 | 6 | **1.619** |
| F1M775 | Diaph1 | Protein diaphanous homolog 1 (Rattus norvegicus) | 243.69 | 4.74 | 140409 | 4 | 1706313.5 | 1300200 | 1571182.75 | 1786169.25 | 922449.69 | 2154315.5 | 2197711.5 | 1628156.88 | 2020454.5 | 1447364.25 | 5 | **1.297** |
| F6ZDS4 | Tpr | Nucleoprotein TPR (Mus musculus) | 978.85 | 9.67 | 273990 | 18 | 26902440 | 14430411 | 20883642 | 15469246 | 17714320 | 22058934 | 18721266 | 18824976 | 19620660 | 18067018 | 9 | **1.020** |
| F7BJB9 | Morc3 | MORC family CW-type zinc finger protein 3 (Mus musculus) | 95.64 | 3.82 | 106522 | 2 | 0 | 627653 | 848178 | 474589.91 | 1042068.19 | 0 | 1173709.12 | 1053550.25 | 472421.28 | 1326352 | 9 | **1.345** |
| G3V6S8 | Srsf6 | Serine/arginine-rich splicing factor 6 (Rattus norvegicus) | 495.3 | 25.96 | 39025 | 9 | 74764440 | 72048344 | 73911256 | 71743128 | 43444192 | 59742624 | 65116440 | 68362896 | 64321008 | 71728880 | 5 | **0.980** |
| G3V7W1 | Pdcd6 | Programmed cell death protein 6 (Rattus norvegicus) | 101.06 | 9.95 | 21905 | 2 | 225873.25 | 1276603.88 | 1190396.38 | 1563754.5 | 1275907.62 | 989409.62 | 878295.56 | 869158.81 | 958462.5 | 838151.88 | 5 | **0.819** |
| G5E829 | Atp2b1 | Plasma membrane calcium-transporting ATPase 1 (Mus musculus) | 671.03 | 13.93 | 134747 | 12 | 7816870 | 7632992 | 8968369 | 8379600.5 | 8474393 | 10707785 | 8999851 | 10356291 | 10555959 | 10510940 | 0 | **1.239** |
| O08528 | Hk2 | Hexokinase-2 (Mus musculus) | 1428.24 | 31.41 | 102535 | 22 | 58215216 | 55515092 | 56726132 | 52468956 | 48778392 | 58966864 | 56162244 | 60414516 | 58757920 | 61771996 | 2 | **1.090** |
| O08529 | Capn2 | Calpain-2 catalytic subunit (Mus musculus) | 733.3 | 23.57 | 79872 | 13 | 14718324 | 19486724 | 15259352 | 19367016 | 16423187 | 12680778 | 13257809 | 14941984 | 15164113 | 17365472 | 5 | **0.861** |
| O08539 | Bin1 | Myc box-dependent-interacting protein 1 (Mus musculus) | 346.58 | 19.9 | 64470 | 7 | 7847207.5 | 6257028 | 6255147.5 | 5809333 | 5669139.5 | 6787688 | 5910718 | 6608997 | 6120033.5 | 6423647 | 9 | **1.000** |
| O08547 | Sec22b | Vesicle-trafficking protein SEC22b (Mus musculus) | 286.61 | 28.84 | 24741 | 5 | 10928426 | 14928759 | 13852077 | 19014588 | 11611427 | 9694480 | 8581584 | 6525659 | 8643225 | 8019364.5 | 0 | **0.590** |
| O08553 | Dpysl2 | Dihydropyrimidinase-related protein 2 (Mus musculus) | 1822.4 | 72.9 | 62278 | 28 | 115468768 | 117006960 | 134334112 | 110017176 | 120595136 | 122370656 | 115490040 | 129444256 | 129051192 | 135113168 | 6 | **1.057** |
| O08579 | Emd | Emerin (Mus musculus) | 257.2 | 33.98 | 29436 | 5 | 8494672 | 7990216 | 7785098 | 5493867 | 7621716 | 6163435 | 5539652 | 8096838 | 7207839.5 | 8276920 | 11 | **0.944** |
| O08583 | Alyref | THO complex subunit 4 (Mus musculus) | 253.9 | 27.06 | 26940 | 4 | 11875945 | 6986969 | 5255737 | 6430219 | 8536843 | 4592606 | 6962494 | 4919321 | 6307370 | 5463366.5 | 4 | **0.723** |
| O08585 | Clta | Clathrin light chain A (Mus musculus) | 178.08 | 13.19 | 25604 | 4 | 30323726 | 15042925 | 16595675 | 15091126 | 20624784 | 19738180 | 21693906 | 18503400 | 22027104 | 23633564 | 7 | **1.081** |
| O08599 | Stxbp1 | Syntaxin-binding protein 1 (Mus musculus) | 105.01 | 3.7 | 67569 | 2 | 0 | 272471.41 | 415138.09 | 237682.11 | 230505.19 | 0 | 0 | 334162.56 | 238085.16 | 231628.78 | 12 | **0.696** |
| O08618 | Prpsap2 | Phosphoribosyl pyrophosphate synthase-associated protein 2 (Rattus norvegicus) | 291.9 | 22.76 | 40867 | 6 | 6267149.5 | 5727799 | 7281254.5 | 5878331 | 5226058.5 | 6625062.5 | 7453654 | 6496671 | 5442271.5 | 6705405.5 | 7 | **1.077** |
| O08648 | Map3k4 | Mitogen-activated protein kinase kinase kinase 4 (Mus musculus) | 37.03 | 0.88 | 179834 | 1 | 0 | 140029.78 | 146302.52 | 149874.52 | 123126.82 | 0 | 0 | 123843.54 | 170180.19 | 126436.07 | 12 | **0.752** |
| O08663 | Metap2 | Methionine aminopeptidase 2 (Mus musculus) | 316.1 | 21.34 | 52922 | 6 | 5178562.5 | 5977512.5 | 6283525 | 8044466 | 5319393.5 | 5436212.5 | 2536130 | 5252437.5 | 5867760.5 | 4901548 | 5 | **0.779** |
| O08697 | Arl2 | ADP-ribosylation factor-like protein 2 (Rattus norvegicus) | 247.82 | 26.09 | 20836 | 3 | 1958014.5 | 2592854.5 | 2360315.75 | 476129.06 | 580398.56 | 2893671.25 | 0 | 2196246.5 | 2682504.25 | 410649.22 | 12 | **1.027** |
| O08700 | Vps45 | Vacuolar protein sorting-associated protein 45 (Rattus norvegicus) | 86.36 | 4.56 | 64894 | 2 | 366911.5 | 1319302.5 | 925188.12 | 860836.81 | 549144.25 | 670474 | 305799.78 | 793920.88 | 400964.88 | 629271.25 | 7 | **0.696** |
| O08709 | Prdx6 | Peroxiredoxin-6 (Mus musculus) | 1188.9 | 83.48 | 24871 | 19 | 138697856 | 134043440 | 144528752 | 167852048 | 156982416 | 154086016 | 123240424 | 133244616 | 151961904 | 141545008 | 8 | **0.949** |
| O08715 | Akap1 | A-kinase anchor protein 1, mitochondrial (Mus musculus) | 99.93 | 3.27 | 92195 | 2 | 427368.44 | 433210.28 | 405868.62 | 0 | 0 | 464461.16 | 476194.62 | 436089.59 | 577892.31 | 710110.75 | 1 | **2.104** |
| O08734 | Bak1 | Bcl-2 homologous antagonist/killer (Mus musculus) | 156.37 | 19.62 | 23295 | 3 | 1231320.88 | 2089871 | 3528660 | 986332.69 | 729239.69 | 3218961.5 | 2421250 | 4028904.5 | 3898303 | 4415748 | 2 | **2.100** |
| O08738 | Casp6 | Caspase-6 (Mus musculus) | 98.15 | 6.88 | 31595 | 2 | 0 | 0 | 0 | 1066014.88 | 942038.75 | 0 | 848123.44 | 836598.88 | 756462 | 962628.25 | 15 | **1.695** |
| O08739 | Ampd3 | AMP deaminase 3 (Mus musculus) | 141.57 | 2.35 | 88652 | 3 | 1587033.38 | 1227860.5 | 1560577.12 | 392102.62 | 506085.25 | 2277428.25 | 2160807.25 | 1773903.12 | 1542427.25 | 1566284.88 | 3 | **1.767** |
| O08749 | Dld | Dihydrolipoyl dehydrogenase, mitochondrial (Mus musculus) | 714.58 | 33.2 | 54272 | 12 | 28236668 | 28013464 | 30186468 | 40122924 | 24712818 | 35878484 | 35723424 | 35020008 | 28861456 | 35996368 | 6 | **1.134** |
| O08756 | Hsd17b10 | 3-hydroxyacyl-CoA dehydrogenase type-2 (Mus musculus) 0 | 632.09 | 53.26 | 27419 | 9 | 21046912 | 18570530 | 20168830 | 19449818 | 18220626 | 16050046 | 20522536 | 20274432 | 18628518 | 18760204 | 12 | **0.967** |
| O08759 | Ube3a | Ubiquitin-protein ligase E3A (Mus musculus) | 168.64 | 5.98 | 99819 | 4 | 3158144 | 2013167.88 | 2211002 | 1637479.38 | 2672513.75 | 3428251 | 3370940 | 2696237.5 | 1764238 | 2883953.75 | 6 | **1.210** |
| O08784 | Tcof1 | Treacle protein (Mus musculus) | 422.11 | 9.39 | 135001 | 9 | 21203750 | 8861584 | 12283888 | 4868213.5 | 12072568 | 13794380 | 12803603 | 14258613 | 15207697 | 16678118 | 5 | **1.227** |
| O08788 | Dctn1 | Dynactin subunit 1 (Mus musculus) | 368.37 | 6.09 | 141676 | 6 | 1881717.75 | 1703108.38 | 3022439 | 1727598.25 | 2424169 | 1902789.62 | 2492758.5 | 2502646 | 2349289.5 | 2821408 | 7 | **1.122** |
| O08795 | Prkcsh | Glucosidase 2 subunit beta (Mus musculus) | 409.18 | 18.23 | 58793 | 7 | 14825971 | 13577900 | 12878836 | 12407479 | 13973062 | 11904046 | 11560019 | 11202170 | 11984042 | 11919059 | 0 | **0.866** |
| O08807 | Prdx4 | Peroxiredoxin-4 (Mus musculus) | 734.19 | 53.65 | 31053 | 13 | 262358320 | 300338912 | 282387040 | 300220864 | 327576224 | 338961600 | 280084768 | 314852416 | 336442208 | 329300416 | 5 | **1.086** |
| O08810 | Eftud2 | 116 kDa U5 small nuclear ribonucleoprotein component (Mus musculus) | 1106.59 | 27.6 | 109361 | 20 | 155994544 | 80197848 | 83870528 | 89003280 | 91648328 | 91623640 | 101447224 | 96202872 | 87642560 | 99135232 | 8 | **0.951** |
| O08836 | Igbp1 | Immunoglobulin-binding protein 1 (Rattus norvegicus) | 59.2 | 3.53 | 39135 | 1 | 174723.19 | 176671.42 | 324098.66 | 131661.95 | 0 | 245986.66 | 537986.44 | 99802.85 | 284827.69 | 0 | 11 | **1.448** |
| O08837 | Cdc5l | Cell division cycle 5-like protein (Rattus norvegicus) | 412.95 | 11.47 | 92218 | 7 | 8692509 | 5726117 | 4550436 | 4250674.5 | 6445177 | 8670712 | 7188518.5 | 8212540 | 6650664 | 7373091.5 | 5 | **1.284** |
| O08915 | Aip | AH receptor-interacting protein (Mus musculus) | 64.99 | 4.24 | 37605 | 1 | 2236891 | 1809814.88 | 1921080.62 | 1411382.38 | 2401121 | 1631451.75 | 2524677.25 | 2237079.5 | 2088012.25 | 2081947.88 | 9 | **1.080** |
| O08917 | Flot1 | Flotillin-1 (Mus musculus) | 61.85 | 2.57 | 47513 | 1 | 0 | 0 | 0 | 722393.75 | 0 | 607924.31 | 1148950.75 | 716847.12 | 900081.69 | 632022 | 9 | **5.545** |
| O08992 | Sdcbp | Syntenin-1 (Mus musculus) | 224.11 | 17.73 | 32379 | 3 | 1817779 | 1783844 | 2342503.25 | 831902.31 | 1628373.88 | 4090364.75 | 4056952 | 3978907 | 5898522 | 4764695.5 | 0 | **2.712** |
| O08997 | Atox1 | Copper transport protein ATOX1 (Mus musculus) | 239.64 | 32.35 | 7338 | 3 | 20809244 | 9854306 | 9907746 | 8330644 | 13404692 | 13360765 | 11547526 | 12678648 | 13579308 | 12054000 | 9 | **1.015** |
| O09043 | Napsa | Napsin-A (Mus musculus) | 98.39 | 6.44 | 45544 | 2 | 7019860 | 4114608.5 | 4652887.5 | 3512031 | 4137120.25 | 3893056.25 | 5851816 | 3927178 | 4387210.5 | 4994261 | 12 | **0.984** |
| O09061 | Psmb1 | Proteasome subunit beta type-1 (Mus musculus) | 774.55 | 61.67 | 26372 | 10 | 44704064 | 38091176 | 39576612 | 30672388 | 36403592 | 44287632 | 35625180 | 26411302 | 34558108 | 38922924 | 9 | **0.949** |
| O09106 | Hdac1 | Histone deacetylase 1 (Mus musculus) | 483.37 | 25.1 | 55075 | 7 | 15320164 | 20314342 | 14951380 | 15147043 | 16187964 | 16081330 | 14462711 | 17210102 | 16986936 | 17150910 | 10 | **1.000** |
| O09110 | Map2k3 | Dual specificity mitogen-activated protein kinase kinase 3 (Mus musculus) | 205.79 | 14.41 | 39296 | 4 | 3581602.5 | 893733.31 | 3420099.25 | 1810836.25 | 2531529.75 | 4373626 | 3832487.5 | 4156146.25 | 4221544 | 4934146 | 0 | **1.758** |
| O09111 | Ndufb11 | NADH dehydrogenase [ubiquinone] 1 beta subcomplex subunit 11, mitochondrial (Mus mu | 48.13 | 14.57 | 17444 | 1 | 544344.19 | 595573.25 | 424509.75 | 533108.19 | 561995.69 | 371308.31 | 542629.69 | 377717.31 | 393806.53 | 391157.78 | 2 | **0.781** |
| O09117 | Sypl1 | Synaptophysin-like protein 1 (Mus musculus) | 88.71 | 4.21 | 28899 | 2 | 4146290.5 | 4403219.5 | 4686254.5 | 5576336 | 4837793.5 | 890258.31 | 4482081.5 | 4673382.5 | 4218568 | 4944931.5 | 9 | **0.812** |
| O09131 | Gsto1 | Glutathione S-transferase omega-1 (Mus musculus) | 109.63 | 16.67 | 27498 | 3 | 4523821.5 | 2278196 | 2981939.5 | 2116887.75 | 4074243 | 4178055.25 | 3892212.75 | 2770056 | 3620950.5 | 2687170.75 | 11 | **1.073** |
| O09159 | Man2b1 | Lysosomal alpha-mannosidase (Mus musculus) | 632.59 | 13.82 | 114648 | 10 | 22631720 | 23016558 | 25024536 | 16373330 | 24080298 | 20609582 | 18686608 | 22062820 | 23805030 | 22975646 | 8 | **0.973** |
| O09167 | Rpl21 | 60S ribosomal protein L21 (Mus musculus) | 193.12 | 20 | 18562 | 3 | 54000160 | 26897780 | 38478156 | 19610310 | 34999692 | 57949616 | 48540964 | 44007800 | 40969988 | 43037484 | 4 | **1.348** |
| O09172 | Gclm | Glutamate--cysteine ligase regulatory subunit (Mus musculus) | 619.61 | 41.24 | 30535 | 10 | 43647504 | 37433592 | 41427840 | 28078038 | 27744814 | 46867536 | 45581000 | 41769748 | 40119928 | 39186504 | 5 | **1.197** |
| O35071 | Kif1c | Kinesin-like protein KIF1C (Mus musculus) | 47 | 1.18 | 122434 | 1 | 0 | 0 | 65255.03 | 0 | 0 | 118530.86 | 0 | 256684.61 | 170324.25 | 240192 | 11 | **12.041** |
| O35075 | Dscr3 | Down syndrome critical region protein 3 homolog (Mus musculus) | 51.46 | 6.4 | 32970 | 1 | 650354.25 | 587723.38 | 476224.53 | 624933.44 | 709926.19 | 465299.62 | 407144.88 | 0 | 0 | 572671.56 | 2 | **0.474** |
| O35092 | Timm17a | Mitochondrial import inner membrane translocase subunit Tim17-A (Rattus norvegicus) | 78.05 | 12.87 | 18038 | 1 | 2529535.25 | 1751816.62 | 1676883.88 | 1272355 | 2163523.25 | 1980162.5 | 1875641.5 | 1568476 | 2170878 | 2071575.88 | 11 | **1.029** |
| O35093 | Timm23 | Mitochondrial import inner membrane translocase subunit Tim23 (Rattus norvegicus) | 77.31 | 8.13 | 21907 | 1 | 735879.25 | 0 | 0 | 873569.94 | 818832.56 | 615274.38 | 580353.19 | 524965 | 521096.22 | 0 | 10 | **0.923** |
| O35095 | Ncdn | Neurochondrin (Rattus norvegicus) | 41.8 | 2.06 | 78923 | 1 | 0 | 0 | 0 | 0 | 0 | 0 | 0 | 515251.34 | 539235.56 | 465957.91 | 20 | **#DIV/0 !** |
| O35114 | Scarb2 | Lysosome membrane protein 2 (Mus musculus) | 108.5 | 5.23 | 54044 | 2 | 154844.95 | 253787.28 | 0 | 0 | 0 | 203487.2 | 0 | 517522.44 | 400546.5 | 253993.45 | 9 | **3.366** |
| O35127 | Grcc10 | Protein C10 (Mus musculus) | 39.12 | 14.29 | 13194 | 1 | 0 | 0 | 320162.84 | 174967.67 | 161401.81 | 111536.76 | 0 | 344555.88 | 499956.41 | 282219.47 | 10 | **1.886** |
| O35129 | Phb2 | Prohibitin-2 (Mus musculus) | 775.45 | 52.51 | 33296 | 13 | 50936140 | 56689912 | 48100476 | 59008900 | 42174260 | 52956696 | 34238476 | 41289924 | 35185668 | 30460422 | 3 | **0.756** |
| O35130 | Emg1 | Ribosomal RNA small subunit methyltransferase NEP1 (Mus musculus) | 256.37 | 22.95 | 26974 | 4 | 2288961.25 | 2867279.5 | 3376357 | 3105882.25 | 2223970.75 | 3487237 | 2283682.75 | 2838038.25 | 2814471 | 2826710.25 | 12 | **1.028** |
| O35134 | Polr1a | DNA-directed RNA polymerase I subunit RPA1 (Mus musculus) | 217 | 3.26 | 194110 | 4 | 1893800.25 | 2214213.75 | 2743660.5 | 1883398 | 2424231.75 | 1130869.12 | 710847.06 | 2410929.5 | 2779964.5 | 2693467.25 | 12 | **0.872** |
| O35141 | Rassf5 | Ras association domain-containing protein 5 (Rattus norvegicus) | 47.35 | 3.63 | 46680 | 1 | 0 | 0 | 135916.5 | 0 | 0 | 222859.94 | 214894.03 | 179906.09 | 0 | 0 | 16 | **4.544** |
| O35142 | Copb2 | Coatomer subunit beta' (Rattus norvegicus) | 1513.15 | 41.77 | 102551 | 26 | 43168188 | 57072396 | 55630944 | 55622336 | 45960128 | 61312976 | 52848044 | 60900132 | 66171720 | 69036728 | 3 | **1.205** |
| O35207 | Cdk2ap1 | Cyclin-dependent kinase 2-associated protein 1 (Mus musculus) | 65.87 | 11.4 | 12354 | 1 | 0 | 0 | 376184.03 | 263781.06 | 196525.28 | 273724.84 | 0 | 374256.56 | 403978.44 | 364977.91 | 9 | **1.694** |
| O35215 | Ddt | D-dopachrome decarboxylase (Mus musculus) | 101.91 | 22.03 | 13077 | 2 | 2534427.75 | 1159739.12 | 969962.25 | 4900360 | 1862892.38 | 2179534.75 | 203348.48 | 1945795 | 2302921.5 | 2259201.75 | 12 | **0.778** |
| O35218 | Cpsf2 | Cleavage and polyadenylation specificity factor subunit 2 (Mus musculus) | 277.63 | 6.52 | 88383 | 5 | 2511579.5 | 1352349.5 | 1342829.5 | 1202764.12 | 759809 | 2509944.25 | 1558803 | 2524895.75 | 1338657 | 2417839.25 | 6 | **1.444** |
| O35226 | Psmd4 | 26S proteasome non-ATPase regulatory subunit 4 (Mus musculus) | 244.27 | 14.1 | 40704 | 4 | 8758211 | 7135502 | 9490966 | 6921804.5 | 2990188.5 | 9950875 | 12271609 | 9334207 | 10221968 | 9533990 | 1 | **1.454** |
| O35239 | Ptpn9 | Tyrosine-protein phosphatase non-receptor type 9 (Mus musculus) | 98.46 | 4.55 | 67970 | 2 | 0 | 685508 | 843327.88 | 812270.88 | 598757.25 | 0 | 395758.03 | 891728.88 | 753173.25 | 853636.38 | 11 | **0.985** |
| O35264 | Pafah1b2 | Platelet-activating factor acetylhydrolase IB subunit beta (Rattus norvegicus) 2 | 106.75 | 12.23 | 25581 | 2 | 360676.38 | 2506323.5 | 1420885.75 | 2648786.5 | 1705123.75 | 2201711.5 | 2291797.5 | 1908879.25 | 1759729.62 | 332801.84 | 12 | **0.983** |
| O35286 | Dhx15 | Pre-mRNA-splicing factor ATP-dependent RNA helicase DHX15 (Mus musculus) | 1498.1 | 31.07 | 91007 | 26 | 51902548 | 62364720 | 57064948 | 65433008 | 60719192 | 62554704 | 53456928 | 59119568 | 57597640 | 66824708 | 11 | **1.007** |
| O35295 | Purb | Transcriptional activator protein Pur-beta (Mus musculus) | 117.75 | 15.12 | 33901 | 2 | 2042624.5 | 642832.69 | 598863.31 | 0 | 1724727.88 | 2389338.75 | 2015820.5 | 2504713.5 | 2190395.5 | 1846649.5 | 2 | **2.185** |
| O35303 | Dnm1l | Dynamin-1-like protein (Rattus norvegicus) | 459.75 | 15.23 | 83908 | 7 | 2699549 | 6644695 | 7589436.5 | 6980706 | 6932898 | 8140386.5 | 8592178 | 9146497 | 8832190 | 7542495 | 1 | **1.370** |
| O35309 | Nmi | N-myc-interactor (Mus musculus) | 141.55 | 16.56 | 35236 | 3 | 1843450.88 | 3153863 | 3307292 | 2471595.75 | 2124730 | 2405433 | 2930180 | 1340701 | 2286181.5 | 2196048 | 9 | **0.865** |
| O35326 | Srsf5 | Serine/arginine-rich splicing factor 5 (Mus musculus) | 397.93 | 27.88 | 30891 | 7 | 56995440 | 42209324 | 46104460 | 42619488 | 19309708 | 43881416 | 40712880 | 46102204 | 47960752 | 47662856 | 10 | **1.092** |
| O35343 | Kpna4 | Importin subunit alpha-3 (Mus musculus) | 501.02 | 25.91 | 57923 | 9 | 11019406 | 12167423 | 11949402 | 11075872 | 9863869 | 17805050 | 16195701 | 15403492 | 15043157 | 13494040 | 0 | **1.390** |
| O35344 | Kpna3 | Importin subunit alpha-4 (Mus musculus) | 431.09 | 18.62 | 57773 | 9 | 15403126 | 15917767 | 16646241 | 15620210 | 12512923 | 17131198 | 17395650 | 17042038 | 14838634 | 17742196 | 4 | **1.106** |
| O35345 | Kpna6 | Importin subunit alpha-7 (Mus musculus) | 382.84 | 18.1 | 59964 | 6 | 4764762 | 5723627 | 5833330 | 5296230 | 4699324 | 4787771 | 4718428 | 4706785 | 4579508.5 | 4557047.5 | 4 | **0.887** |
| O35350 | Capn1 | Calpain-1 catalytic subunit (Mus musculus) | 383 | 11.92 | 82106 | 7 | 5373070 | 6352969 | 7889988 | 3574670 | 6233404 | 6243877 | 6643934 | 7201307 | 6286605.5 | 8176442.5 | 6 | **1.174** |
| O35375 | Nrp2 | Neuropilin-2 (Mus musculus) | 132.42 | 3.87 | 104631 | 3 | 8775626 | 2409488.25 | 4353239 | 2872341.5 | 3848369 | 4262183 | 4198001 | 5035199 | 4454680 | 6232738 | 7 | **1.086** |
| O35379 | Abcc1 | Multidrug resistance-associated protein 1 (Mus musculus) | 272.02 | 5.37 | 171185 | 6 | 1556851.88 | 1396159.25 | 2259833.5 | 1450025.38 | 1222834 | 2994807.25 | 1933289 | 3601483.5 | 2435304 | 3243533.75 | 1 | **1.802** |
| O35381 | Anp32a | Acidic leucine-rich nuclear phosphoprotein 32 family member A (Mus musculus) | 539.99 | 28.74 | 28538 | 9 | 41174436 | 41002816 | 52619424 | 37306416 | 47466244 | 40381564 | 36584956 | 50494372 | 38551580 | 51606728 | 10 | **0.991** |
| O35382 | Exoc4 | Exocyst complex component 4 (Mus musculus) | 160.53 | 5.13 | 110545 | 3 | 1845657.75 | 1093885 | 2310414.25 | 913737.69 | 0 | 960317.62 | 1661980 | 1426893.25 | 416443.44 | 1917374.62 | 12 | **1.036** |
| O35405 | Pld3 | Phospholipase D3 (Mus musculus) | 126.44 | 9.22 | 54389 | 2 | 460195.81 | 891286.81 | 1339470.62 | 563072.38 | 401621.06 | 890598.69 | 0 | 847152.25 | 990469.5 | 1048178.69 | 11 | **1.033** |
| O35435 | Dhodh | Dihydroorotate dehydrogenase (quinone), mitochondrial (Mus musculus) | 65.69 | 2.28 | 42700 | 1 | 604045 | 853272.56 | 570406.88 | 1057999.12 | 671411.31 | 123072.54 | 0 | 338422.19 | 730156.12 | 396379.25 | 3 | **0.423** |
| O35459 | Ech1 | Delta(3,5)-Delta(2,4)-dienoyl-CoA isomerase, mitochondrial (Mus musculus) | 375.39 | 19.88 | 36118 | 6 | 13721496 | 9963191 | 11869011 | 11168562 | 19448604 | 11386583 | 10464962 | 14703772 | 14930808 | 12889030 | 11 | **0.973** |
| O35465 | Fkbp8 | Peptidyl-prolyl cis-trans isomerase FKBP8 (Mus musculus) | 434.44 | 27.11 | 43529 | 7 | 14658320 | 10175525 | 11813610 | 10248845 | 10475697 | 13210549 | 12524477 | 11747018 | 11522918 | 11761032 | 8 | **1.059** |
| O35495 | Cdk14 | Cyclin-dependent kinase 14 (Mus musculus) | 96.49 | 3.41 | 52996 | 2 | 0 | 443961.53 | 559775.81 | 379982.53 | 371236.91 | 730088.62 | 472757.53 | 607657 | 487629.88 | 453865.06 | 3 | **1.568** |
| O35509 | Rab11b | Ras-related protein Rab-11B (Rattus norvegicus) | 428.27 | 37.61 | 24489 | 8 | 40248320 | 76864664 | 81692552 | 74151096 | 73374688 | 55654868 | 71356120 | 87735696 | 90038536 | 91412048 | 8 | **1.144** |
| O35522 | Psmb9 | Proteasome subunit beta type-9 (Mus musculus) bactrianus (Southwestern Asian house mou | 209.5 | 21 | 23427 | 5 | 9876936 | 14141506 | 14785920 | 15765350 | 13633892 | 12348320 | 15203445 | 13115092 | 15438930 | 14978992 | 11 | **1.042** |

| | | | | | | | | | | | | | | | | | | |
|---|---|---|---|---|---|---|---|---|---|---|---|---|---|---|---|---|---|---|
| O35532 | Msmo1 | Methylsterol monooxygenase 1 (Rattus norvegicus) | 53.73 | 5.8 | 34964 | 1 | 240641.92 | 252552.5 | 103838.56 | 130314.24 | 96704.22 | 501879.34 | 483151.97 | 70943.28 | 111073.84 | 0 | 12 | **1.416** |
| O35543 | Hpgds | Hematopoietic prostaglandin D synthase (Rattus norvegicus) | 39.94 | 4.52 | 23297 | 1 | 1680373.5 | 1308605.25 | 0 | 1107750.88 | 1210799.62 | 1183580.25 | 1448158.75 | 1281289.25 | 1418878.38 | 1084921.5 | 11 | **1.209** |
| O35593 | Psmd14 | 26S proteasome non-ATPase regulatory subunit 14 (Mus musculus) | 399.85 | 38.06 | 34577 | 6 | 12696569 | 19872692 | 16479362 | 17572584 | 16381160 | 12242326 | 12113963 | 15330802 | 16155768 | 15128259 | 3 | **0.855** |
| O35601 | Fyb1 | FYN-binding protein 1 (Mus musculus) | 212.6 | 4.4 | 90055 | 4 | 1865621.88 | 3338581.75 | 2992108.5 | 3219814 | 4687228.5 | 3587027 | 3980639.75 | 4112916.75 | 4227931.5 | 4040220.5 | 5 | **1.239** |
| O35604 | Npc1 | NPC intracellular cholesterol transporter 1 (Mus musculus) | 104.87 | 3.45 | 142885 | 3 | 1955991.5 | 2049957.38 | 1260163.62 | 2058745 | 4404391 | 1597900 | 1558137.5 | 1887765.12 | 1296708.12 | 1930220.38 | 5 | **0.705** |
| O35609 | Scamp3 | Secretory carrier-associated membrane protein 3 (Mus musculus) | 49.17 | 4.58 | 38458 | 1 | 891906.19 | 0 | 0 | 0 | 1031617.06 | 681000.25 | 1022219.94 | 1016960.44 | 880165.44 | 0 | 14 | **1.872** |
| O35638 | Stag2 | Cohesin subunit SA-2 (Mus musculus) | 91.12 | 2.11 | 141281 | 2 | 241069.48 | 683503.69 | 489126.94 | 722500.44 | 456731.16 | 189684.84 | 0 | 293570.78 | 286461.31 | 333872.94 | 3 | **0.426** |
| O35639 | Anxa3 | Annexin A3 (Mus musculus) | 1350.51 | 61.61 | 36384 | 20 | 117912520 | 127235496 | 124105480 | 129778568 | 134606976 | 144579360 | 112003344 | 116621176 | 123643304 | 122070112 | 7 | **0.977** |
| O35643 | Ap1b1 | AP-1 complex subunit beta-1 (Mus musculus) | 1862.21 | 40.93 | 103935 | 28 | 30247816 | 39140588 | 45932076 | 43905444 | 39368400 | 42155664 | 38882864 | 56005844 | 51810984 | 54087228 | 6 | **1.223** |
| O35658 | C1qbp | Complement component 1 Q subcomponent-binding protein, mitochondrial (Mus musculus) | 332.99 | 24.46 | 31013 | 4 | 3313173.5 | 13957845 | 11402733 | 15113661 | 12432894 | 8526703 | 6655149.5 | 6431356 | 7527134.5 | 7114819 | 5 | **0.645** |
| O35664 | Ifnar2 | Interferon alpha/beta receptor 2 (Mus musculus) | 44.45 | 1.95 | 56578 | 1 | 0 | 588855.75 | 504597.75 | 0 | 542160.19 | 0 | 0 | 624846.25 | 673031.81 | 534335.75 | 13 | **1.120** |
| O35682 | Myadm | Myeloid-associated differentiation marker (Mus musculus) | 127.96 | 8.44 | 35285 | 2 | 2170235.25 | 577450.06 | 0 | 376960.38 | 0 | 1390240.38 | 1063065.5 | 832515.75 | 812200 | 1034325.69 | 6 | **1.643** |
| O35685 | Nudc | Nuclear migration protein nudC (Mus musculus) | 992.37 | 51.81 | 38358 | 18 | 71430304 | 63473532 | 74473432 | 63480860 | 80645816 | 76935800 | 65660008 | 69038336 | 74231160 | 71909504 | 11 | **1.012** |
| O35691 | Pnn | Pinin (Mus musculus) | 79.98 | 2.34 | 82436 | 2 | 3731096.5 | 3569692.25 | 3505013 | 3188274.5 | 2821476 | 2031043.62 | 3700032.25 | 4820162.5 | 4841245 | 4663028 | 6 | **1.193** |
| O35704 | Sptlc1 | Serine palmitoyltransferase 1 (Mus musculus) | 48.97 | 3.81 | 52535 | 1 | 179902.11 | 334286.84 | 265295.97 | 271683.91 | 0 | 130305.87 | 260013.7 | 208954.31 | 295994.56 | 191972.02 | 11 | **1.034** |
| O35737 | Hnrnph1 | Heterogeneous nuclear ribonucleoprotein H (Mus musculus) | 645.83 | 31.85 | 49199 | 10 | 36858160 | 39402900 | 31801212 | 42106152 | 36520696 | 33048684 | 31789092 | 39403368 | 38381432 | 39445960 | 12 | **0.975** |
| O35828 | Coro7 | Coronin-7 (Rattus norvegicus) | 196.19 | 7.05 | 100782 | 4 | 8313893 | 7230458.5 | 7118883 | 7119447 | 6188417 | 8309919 | 8426868 | 8374701 | 7374888.5 | 6820661.5 | 6 | **1.093** |
| O35841 | Api5 | Apoptosis inhibitor 5 (Mus musculus) | 810.1 | 34.13 | 56785 | 13 | 25974118 | 21383742 | 23191004 | 20958978 | 16246554 | 22354296 | 23018796 | 25027904 | 23507296 | 23861970 | 7 | **1.093** |
| O35855 | Bcat2 | Branched-chain-amino-acid aminotransferase, mitochondrial (Mus musculus) | 217.75 | 11.96 | 44127 | 4 | 14489939 | 12583109 | 14423989 | 13199741 | 11997085 | 11780188 | 11969700 | 12258327 | 12916100 | 12742996 | 5 | **0.925** |
| O35857 | Timm44 | Mitochondrial import inner membrane translocase subunit TIM44 (Mus musculus) | 493.01 | 21.46 | 51091 | 9 | 7320577 | 8127194 | 5801815.5 | 6920609.5 | 6650774 | 8711473 | 6452049 | 7482268 | 7105276.5 | 7027355 | 9 | **1.056** |
| O35864 | Cops5 | COP9 signalosome complex subunit 5 (Mus musculus) | 349.26 | 21.26 | 37549 | 7 | 10847557 | 10901956 | 13385044 | 9139152 | 7315845 | 15038064 | 13684781 | 13169667 | 13612852 | 12810917 | 2 | **1.324** |
| O35887 | Calu | Calumenin (Mus musculus) | 505.72 | 31.75 | 37064 | 7 | 9194570 | 13115308 | 10514737 | 12734605 | 12065465 | 12286664 | 13455838 | 11084420 | 12082074 | 12625177 | 9 | **1.068** |
| O35889 | Afdn | Afadin (Rattus norvegicus) | 42.96 | 0.98 | 207678 | 1 | 0 | 78473.39 | 122232.48 | 118392.07 | 99917.62 | 80654.53 | 0 | 165770.44 | 154207.52 | 119535.7 | 9 | **1.241** |
| O35900 | Lsm2 | U6 snRNA-associated Sm-like protein LSm2 (Mus musculus) | 261.26 | 55.79 | 10835 | 5 | 3914340.5 | 7616122 | 6535731 | 5334927 | 5748480 | 4848332.5 | 6772688 | 5570797 | 5843179 | 6199505 | 12 | **1.003** |
| O35945 | Aldh1a7 | Aldehyde dehydrogenase, cytosolic 1 (Mus musculus) | 103.04 | 3.59 | 54588 | 2 | 27490564 | 18308660 | 13489729 | 14773782 | 14328028 | 22012422 | 16906326 | 20646094 | 20528860 | 21792472 | 6 | **1.153** |
| O35955 | Psmb10 | Proteasome subunit beta type-10 (Mus musculus) | 228.22 | 15.02 | 29063 | 3 | 7603417 | 4649171 | 5670138.5 | 6088379 | 6648596 | 6256136 | 6041871 | 6044875.5 | 6765447 | 6935150 | 10 | **1.045** |
| O35963 | Rab33b | Ras-related protein Rab-33B (Mus musculus) | 159.88 | 10.48 | 25767 | 3 | 925365.19 | 730312.75 | 0 | 209969.62 | 785420.88 | 1067989.75 | 964213.69 | 726743.44 | 1206509.38 | 985300.88 | 3 | **1.867** |
| O35964 | Sh3gl1 | Endophilin-A2 (Rattus norvegicus) | 159.15 | 10.33 | 41492 | 3 | 3451404.75 | 4668639 | 4506118 | 3531856.75 | 2409688.25 | 3748349.25 | 3954760 | 4111086 | 4757845 | 3970172 | 8 | **1.106** |
| O35986 | Zranb2 | Zinc finger Ran-binding domain-containing protein 2 (Rattus norvegicus) | 170.09 | 12.42 | 37350 | 4 | 4425681.5 | 7265259.5 | 9410981 | 5434527 | 9603744 | 6158634 | 8700472 | 8822704 | 9567847 | 9365119 | 10 | **1.179** |
| O54692 | Zw10 | Centromere/kinetochore protein zw10 homolog (Mus musculus) | 251.03 | 9.88 | 88063 | 5 | 1520201 | 2221459.5 | 2020323.5 | 1419322.5 | 1411225.12 | 2528023.75 | 2935945.5 | 2750049 | 2643862 | 2636045 | 0 | **1.570** |
| O54734 | Ddost | Dolichyl-diphosphooligosaccharide--protein glycosyltransferase 48 kDa subunit (Mus musc | 744.76 | 39.46 | 49028 | 12 | 28784130 | 44321128 | 43774636 | 40072132 | 38350576 | 39413052 | 35313192 | 34006208 | 43590472 | 47723348 | 12 | **1.024** |
| O54774 | Ap3d1 | AP-3 complex subunit delta-1 (Mus musculus) | 140.71 | 3.84 | 135081 | 3 | 1036916.5 | 1141704.75 | 1846796.5 | 1868683.5 | 677740.81 | 2072867 | 965819 | 1361660.25 | 2246384 | 2015838.5 | 6 | **1.318** |
| O54781 | Srpk2 | SRSF protein kinase 2 (Mus musculus) | 50.4 | 1.91 | 76757 | 1 | 202895.52 | 340897.38 | 296048.59 | 0 | 0 | 509208.25 | 0 | 0 | 0 | 0 | 19 | **0.606** |
| O54782 | Man2b2 | Epididymis-specific alpha-mannosidase (Mus musculus) | 97.25 | 3.83 | 115610 | 2 | 0 | 0 | 138448.53 | 0 | 582669.06 | 456950.59 | 125457.58 | 361823.47 | 556273 | 399583.41 | 9 | **2.635** |
| O54825 | Bysl | Bystin (Mus musculus) | 200.96 | 9.63 | 49784 | 4 | 3107314.5 | 4574074.5 | 5005694.5 | 3897666.5 | 2754964.5 | 4371604 | 4929425.5 | 4564541.5 | 4216059 | 3391106.25 | 10 | **1.110** |
| O54833 | Csnk2a2 | Casein kinase II subunit alpha' (Mus musculus) | 310.33 | 23.43 | 41215 | 6 | 4668629.5 | 5984685 | 9971373 | 5968385 | 5660282 | 7195965 | 7414905 | 9264829 | 8981888 | 10032689 | 4 | **1.330** |
| O54835 | Smad9 | Mothers against decapentaplegic homolog 9 (Rattus norvegicus) | 80.37 | 5.53 | 48992 | 2 | 2525646.5 | 1478763.88 | 2149803.5 | 1027556.5 | 3093524.25 | 2279994.25 | 2202723.25 | 2148498.75 | 2738029.75 | 1768603.75 | 11 | **1.084** |
| O54888 | Polr1b | DNA-directed RNA polymerase I subunit RPA2 (Rattus norvegicus) | 44.48 | 1.06 | 127719 | 1 | 99803.48 | 297115.47 | 273594.72 | 0 | 0 | 171087.64 | 258809.22 | 297802.94 | 0 | 0 | 15 | **1.085** |
| O54910 | Nfkbie | NF-kappa-B inhibitor epsilon (Mus musculus) | 60.89 | 3.3 | 39200 | 1 | 0 | 0 | 366901.28 | 400466.31 | 373320.22 | 0 | 0 | 400585.94 | 353281.59 | 475617.09 | 14 | **1.078** |
| O54940 | Bnip2 | BCL2/adenovirus E1B 19 kDa protein-interacting protein 2 (Mus musculus) | 66.62 | 7.67 | 37769 | 2 | 1146863.12 | 738038.19 | 937399.44 | 0 | 0 | 1132143.75 | 1067980.12 | 751289.56 | 57013.05 | 0 | 13 | **1.066** |
| O54941 | Smarce1 | SWI/SNF-related matrix-associated actin-dependent regulator of chromatin subfamily E mer | 54.7 | 3.16 | 46638 | 1 | 1675307.38 | 1310047 | 1383652.38 | 1772800.62 | 1943508.88 | 0 | 1941372.62 | 2232333.5 | 0 | 0 | 12 | **0.516** |
| O54950 | Prkag1 | 5'-AMP-activated protein kinase subunit gamma-1 (Mus musculus) | 393.82 | 26.97 | 37520 | 6 | 2945779 | 5664837 | 6841320 | 7044781.5 | 5776935 | 5814787.5 | 5952352 | 5916308.5 | 6288043 | 6437328.5 | 10 | **1.076** |
| O54962 | Banf1 | Barrier-to-autointegration factor (Mus musculus) | 152.34 | 44.94 | 10103 | 3 | 13910853 | 7552971.5 | 7619131 | 7729859.5 | 13527872 | 10813649 | 10103294 | 9751771 | 9385278 | 8964889 | 10 | **0.974** |
| O54984 | Asna1 | ATPase Asna1 (Mus musculus) | 274.95 | 32.18 | 38823 | 6 | 16447682 | 11584972 | 12692844 | 14322501 | 12646926 | 13530298 | 12622327 | 11200262 | 12147967 | 10465710 | 5 | **0.886** |
| O54988 | Slk | STE20-like serine/threonine-protein kinase (Mus musculus) | 462.66 | 8.92 | 141457 | 9 | 11472101 | 4963881 | 10839852 | 11711427 | 7703362.5 | 12039852 | 8992776 | 11583190 | 11301591 | 12852152 | 6 | **1.216** |
| O55012 | Picalm | Phosphatidylinositol-binding clathrin assembly protein (Rattus norvegicus) | 972.19 | 37.19 | 69286 | 16 | 30591922 | 38556988 | 41113356 | 32466186 | 35503640 | 40225940 | 40329552 | 48798456 | 42583884 | 43808300 | 2 | **1.210** |
| O55022 | Pgrmc1 | Membrane-associated progesterone receptor component 1 (Mus musculus) | 78.18 | 7.18 | 21694 | 1 | 2227765.75 | 1428915.75 | 1166617.88 | 1845458.88 | 0 | 1177382.88 | 1949989.25 | 1369625.38 | 2100754.75 | 1758265 | 10 | **1.253** |
| O55023 | Impa1 | Inositol monophosphatase 1 (Mus musculus) | 561.25 | 42.6 | 30436 | 11 | 31203516 | 35220940 | 36634624 | 36676984 | 38596872 | 34593924 | 23545428 | 37730648 | 35722360 | 34621088 | 8 | **0.932** |
| O55043 | Arhgef7 | Rho guanine nucleotide exchange factor 7 (Rattus norvegicus) | 98.45 | 4.49 | 73140 | 2 | 2462730.75 | 2139590 | 1718597.88 | 1751880.75 | 2053723.88 | 2363064 | 1867992.25 | 2440419.25 | 2640386.75 | 2393632.5 | 6 | **1.156** |
| O55047 | Tlk2 | Serine/threonine-protein kinase tousled-like 2 (Mus musculus) | 150.05 | 3.9 | 82261 | 3 | 0 | 580915.12 | 485092 | 465500.56 | 722568.19 | 401911.12 | 204988.05 | 771473.12 | 622963.25 | 946911.56 | 9 | **1.308** |
| O55098 | Stk10 | Serine/threonine-protein kinase 10 (Mus musculus) | 386.24 | 9.73 | 111906 | 7 | 7211931 | 3958672.25 | 7706414 | 9457672 | 4668200.5 | 6302622 | 7184439.5 | 8913564 | 9303241 | 9914079 | 8 | **1.261** |
| O55101 | Syngr2 | Synaptogyrin-2 (Mus musculus) | 54.27 | 4.46 | 24778 | 1 | 3110087.5 | 4368730.5 | 4182091.75 | 4224840 | 2969239 | 6056100.5 | 5033463.5 | 5334115 | 5359933 | 4792124.5 | 0 | **1.409** |
| O55106 | Strn | Striatin (Mus musculus) | 40.16 | 1.03 | 85966 | 1 | 0 | 1371431.25 | 1101560.38 | 0 | 0 | 1203994.25 | 1416149.25 | 1160418.38 | 1613696.25 | 1637015.5 | 5 | **2.843** |
| O55125 | Nipsnap1 | Protein NipSnap homolog 1 (Mus musculus) 1 | 54.52 | 3.17 | 33363 | 1 | 5279957 | 1878379.75 | 2384794 | 2272690 | 1442339.25 | 3623801.5 | 3590695 | 2877993.25 | 3848260.25 | 3420678.5 | 5 | **1.309** |
| O55131 | Sept7 | Septin-7 (Mus musculus) | 388.18 | 22.48 | 50550 | 7 | 6259523.5 | 6926517.5 | 6044650 | 9351994 | 5904303 | 6494834 | 4201939.5 | 4970819 | 5071446.5 | 5629432 | 3 | **0.765** |
| O55135 | Eif6 | Eukaryotic translation initiation factor 6 (Mus musculus) | 425.9 | 40.82 | 26511 | 7 | 26860422 | 26679722 | 23226784 | 26780026 | 27615304 | 25326512 | 23676988 | 23515464 | 22361688 | 25231180 | 4 | **0.916** |
| O55142 | Rpl35a | 60S ribosomal protein L35a (Mus musculus) | 149.86 | 31.82 | 12554 | 4 | 19679548 | 27364646 | 32233748 | 35912268 | 22422812 | 23301492 | 11864810 | 14075509 | 14870770 | 13835862 | 2 | **0.566** |
| O55143 | Atp2a2 | Sarcoplasmic/endoplasmic reticulum calcium ATPase 2 (Mus musculus) | 1805.17 | 33.43 | 114858 | 30 | 81937488 | 80366992 | 85646248 | 76896376 | 68561264 | 78245488 | 82004200 | 83176784 | 87506744 | 83674944 | 6 | **1.054** |
| O55171 | Acot2 | Acyl-coenzyme A thioesterase 2, mitochondrial (Rattus norvegicus) | 110.74 | 6.4 | 49701 | 2 | 1278078.25 | 2106839 | 2643897.5 | 1826450.38 | 1951164.88 | 2703715.25 | 1632816.38 | 2302430.25 | 2174864.25 | 2412946.5 | 7 | **1.145** |
| O55173 | Pdpk1 | 3-phosphoinositide-dependent protein kinase 1 (Rattus norvegicus) | 75.64 | 2.86 | 63593 | 1 | 487869.69 | 615063.12 | 392944.09 | 539556.06 | 113798.79 | 421138.44 | 372899.88 | 351659.03 | 577048.62 | 860477.44 | 12 | **1.202** |
| O55201 | Supt5h | Transcription elongation factor SPT5 (Mus musculus) | 469.99 | 12.48 | 120664 | 9 | 5366546.5 | 3999549 | 7459454 | 4277688 | 6545547 | 7818873.5 | 6469751 | 10284548 | 6329978 | 8296746 | 4 | **1.418** |
| O55234 | Psmb5 | Proteasome subunit beta type-5 (Mus musculus) | 96.7 | 8.33 | 28532 | 2 | 2729750.75 | 2891165 | 3966650 | 2691113.5 | 1472838.5 | 2164445.5 | 3304250.25 | 1827414.88 | 1901979.12 | 2114083.5 | 8 | **0.823** |
| O70126 | Aurkb | Aurora kinase B (Mus musculus) | 57.09 | 3.77 | 39384 | 1 | 659920.69 | 717389.19 | 1141522.75 | 973799.88 | 1015285.75 | 624712.94 | 974862.19 | 775611.88 | 784297.25 | 663347.88 | 8 | **0.848** |
| O70133 | Dhx9 | ATP-dependent RNA helicase A (Mus musculus) | 1821.86 | 26.16 | 149475 | 31 | 64600044 | 90062208 | 85556192 | 85777144 | 81483184 | 93586888 | 89941752 | 102238248 | 105379328 | 109645208 | 1 | **1.229** |
| O70145 | Ncf2 | Neutrophil cytosol factor 2 (Mus musculus) | 832.97 | 38.48 | 59485 | 17 | 70212152 | 58194048 | 61757128 | 61209092 | 52748392 | 59396476 | 71168768 | 65631764 | 63726524 | 62204116 | 6 | **1.059** |
| O70152 | Dpm1 | Dolichol-phosphate mannosyltransferase subunit 1 (Mus musculus) | 228.22 | 22.69 | 29175 | 4 | 510428.72 | 4094986.5 | 3604479.25 | 3874272 | 3707708.5 | 3942115.5 | 4334019.5 | 4054786.5 | 4242061 | 3580890.5 | 6 | **1.276** |
| O70194 | Eif3d | Eukaryotic translation initiation factor 3 subunit D (Mus musculus) | 902.62 | 43.98 | 63989 | 16 | 30257812 | 28733616 | 31908856 | 24466882 | 35363644 | 35905060 | 35441632 | 36722988 | 35843788 | 34077192 | 1 | **1.181** |
| O70199 | Ugdh | UDP-glucose 6-dehydrogenase (Rattus norvegicus) | 64.61 | 1.83 | 54892 | 1 | 488118.03 | 353019.62 | 553627.38 | 419302.88 | 434611.06 | 566901.31 | 313413.69 | 384570.16 | 333665.75 | 356058.84 | 7 | **0.869** |
| O70201 | Birc5 | Baculoviral IAP repeat-containing protein 5 (Mus musculus) | 59.11 | 10 | 16298 | 1 | 381492.56 | 214509.08 | 157344.08 | 180149.78 | 216275.58 | 0 | 0 | 68763.7 | 197880.81 | 210483.72 | 5 | **0.415** |
| O70251 | Eef1b | Elongation factor 1-beta (Mus musculus) | 767.41 | 73.33 | 24694 | 12 | 166143120 | 178143616 | 178698080 | 164655200 | 188859872 | 161078448 | 94472600 | 188834704 | 205642768 | 200481568 | 11 | **0.970** |
| O70252 | Hmox2 | Heme oxygenase 2 (Mus musculus) | 426.79 | 35.87 | 35739 | 8 | 56279632 | 11220643 | 10686552 | 11028855 | 8027231.5 | 8243059 | 11343573 | 11207832 | 12601423 | 9001672 | 12 | **0.539** |
| O70305 | Atxn2 | Ataxin-2 (Mus musculus) | 38.88 | 0.93 | 136485 | 1 | 589800.94 | 470031.66 | 0 | 340663.97 | 502889.81 | 442686.38 | 399880.56 | 0 | 611885.88 | 526150.44 | 12 | **1.041** |
| O70310 | Nmt1 | Glycylpeptide N-tetradecanoyltransferase 1 (Mus musculus) | 513.56 | 25.2 | 56888 | 10 | 26127238 | 18780386 | 22356246 | 16725510 | 19596388 | 24572082 | 18656260 | 20357418 | 19947126 | 21439794 | 11 | **1.013** |
| O70318 | Epb41l2 | Band 4.1-like protein 2 (Mus musculus) | 362.89 | 10.63 | 109940 | 8 | 7466906 | 7358596.5 | 7394816 | 6191099 | 6093989 | 8208987 | 9529214 | 10044652 | 7998844 | 8041848.5 | 0 | **1.270** |
| O70351 | Hsd17b10 | 3-hydroxyacyl-CoA dehydrogenase type-2 (Rattus norvegicus) 0 | 208.35 | 24.14 | 27246 | 4 | 15243306 | 14093532 | 12913633 | 13111396 | 12951566 | 7295599 | 13477136 | 14663388 | 13815700 | 14381532 | 11 | **0.931** |
| O70370 | Ctss | Cathepsin S (Mus musculus) | 399.87 | 28.53 | 38475 | 6 | 8529381 | 7603918 | 5952036 | 6770676 | 8034471 | 8745192 | 7303789 | 9446527 | 11044375 | 8743523 | 3 | **1.228** |
| O70378 | Emc8 | ER membrane protein complex subunit 8 (Mus musculus) | 215.66 | 26.09 | 23348 | 4 | 989504.88 | 4133602.5 | 5054308 | 4200347.5 | 4294902.5 | 6690644.5 | 5764991 | 5035526.5 | 4921810 | 4806910.5 | 3 | **1.458** |
| O70435 | Psma3 | Proteasome subunit alpha type-3 (Mus musculus) | 876.56 | 51.76 | 28405 | 14 | 69644760 | 80142392 | 85187688 | 75511256 | 82845760 | 73878904 | 80741864 | 77312600 | 85521816 | 90503144 | 9 | **1.037** |
| O70439 | Stx7 | Syntaxin-7 (Mus musculus) | 92.15 | 10.73 | 29821 | 2 | 5177416 | 2096778.25 | 2412157.5 | 2676686.25 | 3178525 | 2108874.5 | 3274750.25 | 2913574.75 | 3386917 | 4074386.25 | 9 | **1.014** |
| O70456 | Sfn | 14-3-3 protein sigma (Mus musculus) | 292.03 | 16.13 | 27706 | 6 | 173566096 | 340880768 | 444959232 | 337409280 | 247154160 | 229723680 | 191927888 | 208030624 | 271227584 | 278527232 | 7 | **0.764** |
| O70467 | Prmt3 | Protein arginine N-methyltransferase 3 (Rattus norvegicus) | 108.47 | 4.55 | 59420 | 2 | 5372180 | 909756.94 | 2632525.75 | 3345317.25 | 2650715 | 4004987 | 4129365.25 | 3521766.25 | 3460400.25 | 3916510.75 | 5 | **1.276** |
| O70469 | Dok2 | Docking protein 2 (Mus musculus) | 144.34 | 8.25 | 45522 | 3 | 5437873.5 | 4064867.75 | 4855744 | 3329450.5 | 3889205 | 5249082.5 | 2580248.25 | 3890904.25 | 2889763.5 | 4245560 | 9 | **0.874** |
| O70492 | Snx3 | Sorting nexin-3 (Mus musculus) | 166.67 | 21.6 | 18757 | 4 | 19226926 | 11573120 | 9048753 | 12554072 | 15444668 | 7037376 | 14944985 | 10806205 | 14121712 | 13076862 | 10 | **0.884** |
| O70493 | Snx12 | Sorting nexin-12 (Mus musculus) | 93.09 | 9.7 | 19116 | 2 | 10838123 | 6252130 | 5007024 | 8294482 | 11648383 | 2912533 | 8781005 | 7441264 | 9021599 | 9018226 | 11 | **0.884** |
| O70503 | Hsd17b12 | Very-long-chain 3-oxoacyl-CoA reductase (Mus musculus) | 754.77 | 41.35 | 34742 | 13 | 35440676 | 35069836 | 34023676 | 32048882 | 31321556 | 38877088 | 35591736 | 38682968 | 38036008 | 36553204 | 0 | **1.118** |
| O70551 | Srpk1 | SRSF protein kinase 1 (Mus musculus) | 544.63 | 20.52 | 73088 | 9 | 7640265 | 7949920.5 | 8513782 | 8951432 | 8041706.5 | 9075918 | 8371455 | 11364323 | 9599410 | 11106507 | 2 | **1.205** |
| O70593 | Sgta | Small glutamine-rich tetratricopeptide repeat-containing protein alpha (Rattus norvegicus) | 53.53 | 4.46 | 34157 | 1 | 822121.19 | 564972.31 | 396215.81 | 539669.31 | 1091608.25 | 517731.56 | 0 | 445945.09 | 440157.44 | 401820.44 | 4 | **0.529** |
| O88271 | Cfdp1 | Craniofacial development protein 1 (Mus musculus) | 204.29 | 21.02 | 32921 | 4 | 6687758.5 | 4933311 | 8295787 | 6039196 | 8740825 | 8309184 | 9451353 | 7337301 | 7305533 | 7504204 | 7 | **1.150** |

| | | | | | | | | | | | | | | | | | | |
|---|---|---|---|---|---|---|---|---|---|---|---|---|---|---|---|---|---|---|
| O88291 | Znf326 | DBIRD complex subunit ZNF326 (Mus musculus) | 47.16 | 2.76 | 65225 | 1 | 590532.94 | 636344.56 | 562589.81 | 724355 | 944810.31 | 587523.38 | 877700.62 | 807453.31 | 793506.56 | 810473.5 | 8 | **1.121** |
| O88342 | Wdr1 | WD repeat-containing protein 1 (Mus musculus) | 1407.82 | 54.29 | 66407 | 22 | 126465192 | 96079592 | 106034592 | 119690984 | 113084696 | 127561112 | 95283544 | 115506400 | 130561544 | 113841120 | 9 | **1.038** |
| O88351 | Ikbkb | Inhibitor of nuclear factor kappa-B kinase subunit beta (Mus musculus) | 61.65 | 2.38 | 86690 | 1 | 0 | 699402.5 | 769958.69 | 738722.5 | 0 | 515778.91 | 556295.12 | 641379.12 | 730497.44 | 821793.25 | 12 | **1.479** |
| O88379 | Baz1a | Bromodomain adjacent to zinc finger domain protein 1A (Mus musculus) | 42.36 | 1.22 | 178459 | 1 | 47168.43 | 0 | 74532.93 | 0 | 0 | 166952.19 | 197876.19 | 140636.23 | 0 | 141726.81 | 8 | **5.318** |
| O88441 | Mtx2 | Metaxin-2 (Mus musculus) | 182.82 | 27.38 | 29758 | 4 | 3920049.5 | 3773200.75 | 2092453.38 | 3554144 | 2332850 | 3816757.5 | 3889242.75 | 3958743.25 | 3611550.75 | 3414607 | 7 | **1.193** |
| O88447 | Klc1 | Kinesin light chain 1 (Mus musculus) | 207.11 | 9.43 | 61450 | 4 | 3794406.25 | 3048875 | 4007074.75 | 834712.38 | 4358873.5 | 3248047.5 | 3018319 | 4152090 | 3687606 | 4658997 | 11 | **1.170** |
| O88455 | Dhcr7 | 7-dehydrocholesterol reductase (Mus musculus) | 117.23 | 7.64 | 53919 | 3 | 653118.44 | 2097323.5 | 1698143.5 | 978678.88 | 437895.47 | 3139201.5 | 2721117.5 | 2518221.5 | 2825251.5 | 2458481.5 | 0 | **2.329** |
| O88456 | Capns1 | Calpain small subunit 1 (Mus musculus) | 561.39 | 63.94 | 28463 | 10 | 13662510 | 13555646 | 13892038 | 13751943 | 13166243 | 14412270 | 16968294 | 17418950 | 16691464 | 18259780 | 0 | **1.231** |
| O88487 | Dync1i2 | Cytoplasmic dynein 1 intermediate chain 2 (Mus musculus) | 399.06 | 17.81 | 68394 | 6 | 4334657.5 | 3991604.5 | 4888876.5 | 5297460.5 | 3403527.5 | 5292114.5 | 5177019.5 | 4884901.5 | 4926055.5 | 5333905.5 | 5 | **1.169** |
| O88508 | Dnmt3a | DNA (cytosine-5)-methyltransferase 3A (Mus musculus) | 263.01 | 7.05 | 101672 | 5 | 3606697.25 | 1889906.88 | 2529447.5 | 332613.97 | 1331621.38 | 3161323.5 | 2320375 | 2184875 | 2884505 | 1804610 | 9 | **1.275** |
| O88531 | Ppt1 | Palmitoyl-protein thioesterase 1 (Mus musculus) | 334.86 | 27.45 | 34490 | 6 | 3937382.75 | 7247936.5 | 3061268 | 7424355 | 9870224 | 7943302 | 2867397.5 | 6445056 | 7885167 | 6424262.5 | 12 | **1.001** |
| O88532 | Zfr | Zinc finger RNA-binding protein (Mus musculus) | 161.41 | 3.07 | 116859 | 2 | 1563539.88 | 733113.12 | 1687918.25 | 746086 | 1830954.88 | 755242.88 | 1673812.25 | 856078.69 | 1054052.38 | 790016.44 | 11 | **0.782** |
| O88543 | Cops3 | COP9 signalosome complex subunit 3 (Mus musculus) | 586.46 | 36.41 | 47832 | 10 | 15758343 | 10846729 | 10237606 | 10011481 | 9635744 | 14927951 | 16293985 | 15739762 | 16279848 | 12821112 | 3 | **1.346** |
| O88544 | Cops4 | COP9 signalosome complex subunit 4 (Mus musculus) | 352.17 | 18.47 | 46285 | 5 | 4608399.5 | 6507748.5 | 5607081 | 6943923.5 | 5749110 | 5856315.5 | 6304034.5 | 4755053.5 | 7047479 | 6895633.5 | 9 | **1.049** |
| O88545 | Cops6 | COP9 signalosome complex subunit 6 (Mus musculus) | 91.65 | 6.17 | 35880 | 2 | 4112387.5 | 3121738.25 | 3395250 | 5924685.5 | 4941973 | 4777682 | 2415760.75 | 4790491 | 4972674.5 | 2938369.25 | 10 | **0.926** |
| O88554 | Parp2 | Poly [ADP-ribose] polymerase 2 (Mus musculus) | 82.87 | 3.94 | 63397 | 2 | 716841.94 | 448879.94 | 594861.75 | 78885.63 | 621123.12 | 305739.91 | 210509.11 | 592713.75 | 445641.94 | 481506.56 | 7 | **0.827** |
| O88569 | Hnrnpa2b1 | Heterogeneous nuclear ribonucleoproteins A2/B1 (Mus musculus) b1 | 1644.85 | 66.86 | 37403 | 24 | 188563184 | 259040192 | 289462112 | 261573216 | 300718848 | 240482080 | 210766192 | 249277232 | 234918544 | 237534208 | 5 | **0.903** |
| O88587 | Comt | Catechol O-methyltransferase (Mus musculus) | 328.45 | 30.19 | 29486 | 6 | 3745454 | 5030349 | 3255307.25 | 3987454.5 | 3785124.5 | 5341302.5 | 4225386.5 | 4711092.5 | 4421194 | 3469932.5 | 7 | **1.119** |
| O88637 | Pcyt2 | Ethanolamine-phosphate cytidylyltransferase (Rattus norvegicus) | 117.49 | 8.42 | 45220 | 2 | 2170653 | 2246806 | 2778077.75 | 2222191.5 | 2740206 | 2582717.5 | 2244561.5 | 1683141.25 | 2177507 | 2652376.5 | 9 | **0.933** |
| O88653 | Lamtor3 | Ragulator complex protein LAMTOR3 (Mus musculus) | 48.84 | 8.87 | 13553 | 1 | 0 | 545494.31 | 810019.69 | 736665.69 | 376879.72 | 549039.62 | 534926.94 | 942350.31 | 413159.81 | 721722.31 | 10 | **1.280** |
| O88668 | Creg1 | Protein CREG1 (Mus musculus) | 178.78 | 16.36 | 24452 | 3 | 7192028 | 6644835.5 | 6028073 | 5022762.5 | 6652136 | 6204781.5 | 7407932 | 5871739.5 | 6504688.5 | 4739909 | 10 | **0.974** |
| O88685 | Psmc3 | 26S proteasome regulatory subunit 6A (Mus musculus) | 851.58 | 44.34 | 49549 | 17 | 34027012 | 30938756 | 33388602 | 26267942 | 35814324 | 38204540 | 36082420 | 32450510 | 35489972 | 34060396 | 5 | **1.099** |
| O88712 | Ctbp1 | C-terminal-binding protein 1 (Mus musculus) | 340.96 | 22 | 47745 | 7 | 19469652 | 13807223 | 14827522 | 14358538 | 14405323 | 15890924 | 15537774 | 17637522 | 15327391 | 18888314 | 5 | **1.083** |
| O88736 | Hsd17b7 | 3-keto-steroid reductase (Mus musculus) | 56.11 | 3.59 | 37317 | 1 | 1553666.12 | 0 | 0 | 367913.91 | 443217.59 | 1614688.38 | 1631343.25 | 1516283.25 | 1261935.25 | 1212704.38 | 4 | **3.060** |
| O88738 | Birc6 | Baculoviral IAP repeat-containing protein 6 (Mus musculus) | 635.37 | 3.75 | 532170 | 13 | 27505318 | 8821942 | 9746726 | 10394895 | 6516752 | 15084635 | 14818257 | 15889752 | 18276486 | 10509403 | 5 | **1.184** |
| O88746 | Tom1 | Target of Myb protein 1 (Mus musculus) | 175.28 | 8.74 | 54325 | 3 | 3029526 | 2356535.5 | 2278502.75 | 2394317.5 | 208835.39 | 1572635.62 | 2488780 | 1828562.62 | 1790857.75 | 1691212.75 | 8 | **0.913** |
| O88763 | Pik3c3 | Phosphatidylinositol 3-kinase catalytic subunit type 3 (Rattus norvegicus) | 40.03 | 1.35 | 101534 | 1 | 0 | 0 | 297882 | 206581.41 | 180435.73 | 0 | 0 | 166442.48 | 170449.45 | 133217.53 | 11 | **0.686** |
| O88777 | Psen2 | Presenilin-2 (Rattus norvegicus) | 109.64 | 3.35 | 50051 | 1 | 405938 | 590002.94 | 472975.41 | 0 | 403324.19 | 463451.47 | 620519.12 | 569880 | 507180.06 | 464939.69 | 6 | **1.403** |
| O88783 | F5 | Coagulation factor V (Mus musculus) | 138.2 | 1.15 | 247230 | 2 | 558729.75 | 489988.22 | 559757.62 | 666163.19 | 703236.06 | 619584.5 | 560187.75 | 776107 | 716395.44 | 719521.75 | 4 | **1.139** |
| O88796 | Rpp30 | Ribonuclease P protein subunit p30 (Mus musculus) | 113.31 | 8.58 | 29473 | 2 | 2559679.5 | 1960837.38 | 2020995.5 | 1683857.75 | 1934987.25 | 2046221.5 | 1963297 | 2145834.75 | 1758130.88 | 1834947.25 | 12 | **0.959** |
| O88811 | Stam2 | Signal transducing adapter molecule 2 (Mus musculus) | 42.52 | 2.1 | 57455 | 1 | 602349.62 | 1212494.75 | 580496.69 | 950268.06 | 694076.5 | 847975.25 | 492651.69 | 696916.69 | 655512.56 | 774717.19 | 11 | **0.858** |
| O88828 | Polr2f | DNA-directed RNA polymerases I, II, and III subunit RPABC2 (Rattus norvegicus) | 40.31 | 5.51 | 14464 | 1 | 3364148 | 1972634 | 2219021.25 | 1980337.12 | 2525487.25 | 2169742 | 2573580.25 | 2053281.62 | 2295801.75 | 2720458 | 10 | **0.979** |
| O88829 | St3gal5 | Lactosylceramide alpha-2,3-sialyltransferase (Mus musculus) | 244.57 | 10.63 | 47360 | 5 | 5344305 | 5884965 | 5173378 | 4955515 | 5132620 | 4900940 | 5020943 | 5355918 | 4573325.5 | 5187524.5 | 8 | **0.945** |
| O88842 | Fgd3 | FYVE, RhoGEF and PH domain-containing protein 3 (Mus musculus) | 74.59 | 3.14 | 80624 | 1 | 326916.91 | 259938.62 | 299453.69 | 0 | 0 | 453976 | 661363.69 | 118839.82 | 0 | 0 | 14 | **1.392** |
| O88844 | Idh1 | Isocitrate dehydrogenase [NADP] cytoplasmic (Mus musculus) | 764.36 | 36.71 | 46674 | 13 | 55087552 | 46048608 | 48879536 | 36803372 | 34427228 | 51306912 | 48622940 | 46137936 | 34367724 | 43523444 | 12 | **1.012** |
| O88879 | Apaf1 | Apoptotic protease-activating factor 1 (Mus musculus) | 152.82 | 2.32 | 141003 | 2 | 414678.47 | 194467.95 | 1393303.5 | 161917.42 | 183229.94 | 584363.44 | 412575.69 | 172102.77 | 178348.98 | 158444.09 | 9 | **0.641** |
| O88958 | Gnpda1 | Glucosamine-6-phosphate isomerase 1 (Mus musculus) | 120.8 | 9.69 | 32549 | 2 | 3292145.25 | 2662754.75 | 2926280.5 | 3188292.75 | 3107003.75 | 3660438.5 | 2977663.5 | 2664318.75 | 2796244.5 | 3236924.75 | 12 | **1.010** |
| O88986 | Gcat | 2-amino-3-ketobutyrate coenzyme A ligase, mitochondrial (Mus musculus) | 53.82 | 5.05 | 44931 | 1 | 0 | 281518.72 | 456549.59 | 304416.34 | 560815.44 | 481309.09 | 0 | 543066 | 0 | 775900.56 | 12 | **1.123** |
| O89001 | Cpd | Carboxypeptidase D (Mus musculus) | 114.53 | 1.74 | 152406 | 2 | 1509683.88 | 3148116.5 | 1429146.25 | 3015761.5 | 1997888.5 | 1885962.62 | 2293999.25 | 2161526 | 603844.75 | 881445.81 | 8 | **0.705** |
| O89017 | Lgmn | Legumain (Mus musculus) | 145.45 | 6.67 | 49373 | 2 | 1979969.88 | 4133870.25 | 3491845 | 2944143.5 | 3969392 | 2330942.75 | 3205558.25 | 2634162.75 | 3219468.5 | 3134523 | 8 | **0.879** |
| O89023 | Tpp1 | Tripeptidyl-peptidase 1 (Mus musculus) | 198.22 | 15.3 | 61342 | 4 | 4880736 | 8419109 | 1959472.25 | 6492341.5 | 7109269 | 7557760.5 | 9237867 | 6474669 | 6337931 | 6076718 | 10 | **1.236** |
| O89046 | Coro1b | Coronin-1B (Rattus norvegicus) | 101.1 | 4.96 | 53845 | 2 | 7149385 | 5918471.5 | 3368823 | 3259281 | 3805236.5 | 5220325 | 5846304.5 | 5717919.5 | 6265996.5 | 6126766.5 | 8 | **1.242** |
| O89050 | Mkln1 | Muskelin (Mus musculus) | 51.01 | 1.63 | 84878 | 1 | 748362.81 | 0 | 688520.44 | 0 | 946657.69 | 501526.94 | 910720.25 | 707921.94 | 737728.06 | 610889.94 | 12 | **1.455** |
| O89051 | Itm2b | Integral membrane protein 2B (Mus musculus) | 110.78 | 7.52 | 30260 | 2 | 634869.38 | 934779.06 | 1085783 | 952022.69 | 1092636.62 | 1117109.62 | 1062526.5 | 1057870.12 | 1119909.12 | 926297.5 | 8 | **1.124** |
| O89053 | Coro1a | Coronin-1A (Mus musculus) | 1319.81 | 54.01 | 50989 | 21 | 534041728 | 375772864 | 417225024 | 377370080 | 449265728 | 530581696 | 368444448 | 484512064 | 458700352 | 478893792 | 9 | **1.078** |
| O89079 | Cope | Coatomer subunit epsilon (Mus musculus) | 436.32 | 36.04 | 34567 | 6 | 6866686 | 7706228.5 | 10779593 | 11073111 | 7429408 | 9483940 | 8117951.5 | 10818914 | 10136240 | 11116534 | 7 | **1.133** |
| O89086 | Rbm3 | RNA-binding protein 3 (Mus musculus) | 254.23 | 53.59 | 16605 | 5 | 8438786 | 15201176 | 16116072 | 14905163 | 13672338 | 9790935 | 8798340 | 10124271 | 8578986 | 7421003.5 | 4 | **0.654** |
| O89110 | Casp8 | Caspase-8 (Mus musculus) | 432.24 | 19.17 | 55357 | 8 | 4353906 | 5803089 | 5325289.5 | 7948822 | 5548503.5 | 6048130.5 | 6066675 | 5656695 | 7229566 | 6319739 | 6 | **1.081** |
| P00158 | Mt-Cyb | Cytochrome b (Mus musculus) | 39.39 | 2.36 | 43210 | 1 | 0 | 205075.23 | 705092.31 | 0 | 212503.64 | 969426.94 | 753008.12 | 0 | 871509.5 | 0 | 11 | **2.311** |
| P00375 | Dhfr | Dihydrofolate reductase (Mus musculus) | 316.6 | 37.43 | 21606 | 6 | 8354471 | 9564058 | 10981812 | 10147198 | 11953822 | 10480797 | 9116044 | 8250990 | 11778810 | 9912461 | 10 | **0.971** |
| P00397 | Mtco1 | Cytochrome c oxidase subunit 1 (Mus musculus) | 57.91 | 2.72 | 56910 | 1 | 211920.12 | 1038802.31 | 1178125 | 465283.16 | 636615.38 | 1453482.5 | 833832.12 | 1535903.5 | 1230163.25 | 1440565 | 2 | **1.839** |
| P00405 | Mtco2 | Cytochrome c oxidase subunit 2 (Mus musculus) | 362.18 | 23.35 | 25976 | 6 | 38443164 | 46967184 | 37643528 | 52048016 | 31732512 | 37851488 | 46134820 | 48592256 | 43022716 | 45586340 | 10 | **1.069** |
| P00493 | Hprt1 | Hypoxanthine-guanine phosphoribosyltransferase (Mus musculus) | 719.46 | 51.38 | 24570 | 11 | 26606110 | 26900532 | 31584256 | 27910168 | 28636656 | 30811956 | 27216996 | 27796368 | 31327772 | 30777092 | 9 | **1.044** |
| P01027 | C3 | Complement C3 (Mus musculus) | 495.78 | 5.59 | 186484 | 10 | 39720388 | 119013968 | 85524936 | 176874816 | 90823152 | 53379092 | 53566904 | 45840560 | 47475208 | 45515164 | 5 | **0.480** |
| P01887 | B2m | Beta-2-microglobulin (Mus musculus) | 56.08 | 8.4 | 13779 | 1 | 867237.44 | 1244057.38 | 735033.69 | 1333927.12 | 1549120.5 | 817071.06 | 1213595.75 | 1009310.19 | 1159046.25 | 963536.19 | 9 | **0.901** |
| P01899 | H2-D1 | H-2 class I histocompatibility antigen, D-B alpha chain (Mus musculus) | 369.49 | 23.76 | 40836 | 7 | 38885724 | 44173092 | 43661080 | 42083164 | 42607668 | 41218424 | 45896816 | 43383764 | 43781788 | 44733968 | 7 | **1.036** |
| P01900 | H2-D1 | H-2 class I histocompatibility antigen, D-D alpha chain (Mus musculus) | 543.81 | 26.85 | 41110 | 9 | 44228204 | 47712884 | 48819868 | 45192988 | 45425848 | 45569848 | 52578004 | 44997644 | 48712756 | 46765052 | 9 | **1.031** |
| P01902 | H2-K1 | H-2 class I histocompatibility antigen, K-D alpha chain (Mus musculus) | 462.83 | 25.54 | 41490 | 9 | 32347170 | 37218520 | 32862386 | 34468668 | 34564600 | 35374788 | 32456466 | 34870520 | 34441744 | 34873712 | 10 | **1.003** |
| P02262 | | Histone H2A type 1 (Rattus norvegicus) | 352.07 | 57.69 | 14077 | 5 | 1040942208 | 287317888 | 389023872 | 106579608 | 343381632 | 765498688 | 679671936 | 600474688 | 652105344 | 627719296 | 5 | **1.534** |
| P02301 | H3f3c | Histone H3.3C (Mus musculus) | 322.8 | 50.74 | 15315 | 6 | 1452368384 | 909234688 | 1228443904 | 994521280 | 1196124032 | 1003872512 | 959959680 | 858984640 | 895109376 | 740945152 | 3 | **0.771** |
| P02798 | Mt2 | Metallothionein-2 (Mus musculus) | 54.81 | 19.67 | 6115 | 1 | 2393418.5 | 1581477.38 | 1545835.25 | 1181107.5 | 2541113.75 | 1339220.5 | 1571179.12 | 1226286.75 | 1904896.5 | 1485658.12 | 8 | **0.814** |
| P02802 | Mt1 | Metallothionein-1 (Mus musculus) | 188.01 | 34.43 | 6018 | 3 | 4930482.5 | 3016108.5 | 2222204 | 1653540.38 | 4064597.5 | 2982866 | 3504440.75 | 3754651.5 | 5599991.5 | 3027923.25 | 9 | **1.188** |
| P03911 | Mtnd4 | NADH-ubiquinone oxidoreductase chain 4 (Mus musculus) | 119.87 | 6.32 | 51882 | 3 | 159151.61 | 1168524.62 | 1090450.25 | 959225.25 | 752494.88 | 1764551.25 | 123807.04 | 1467414.88 | 1555210 | 1039471.94 | 7 | **1.441** |
| P03921 | Mtnd5 | NADH-ubiquinone oxidoreductase chain 5 (Mus musculus) | 145.72 | 6.59 | 68475 | 3 | 2060673.75 | 2186605.25 | 1603184.38 | 1341622.75 | 392239.09 | 1926327.25 | 2446561.25 | 1081639.88 | 1050116 | 1074281.62 | 11 | **0.999** |
| P03958 | Ada | Adenosine deaminase (Mus musculus) | 87.85 | 7.67 | 39992 | 2 | 0 | 0 | 673208.5 | 0 | 0 | 635509.94 | 1102388 | 644757.31 | 1256489 | 1334687.5 | 8 | **7.388** |
| P03975 | Iap | IgE-binding protein (Mus musculus) | 593.11 | 23.52 | 62747 | 10 | 20999216 | 17572656 | 18099556 | 15463192 | 13372818 | 18575776 | 20176416 | 18278530 | 18729764 | 19406726 | 5 | **1.113** |
| P03995 | Gfap | Glial fibrillary acidic protein (Mus musculus) | 166.35 | 6.05 | 49900 | 3 | 58089500 | 84765360 | 95236016 | 96953024 | 82705712 | 46233808 | 67078580 | 79949584 | 85561856 | 75538536 | 6 | **0.848** |
| P04184 | Tk1 | Thymidine kinase, cytosolic (Mus musculus) | 250.66 | 24.89 | 25776 | 5 | 7086316.5 | 8309174 | 7564118.5 | 7012133 | 7669179 | 6511499.5 | 7510161 | 5741195 | 6269874.5 | 6676438 | 2 | **0.869** |
| P04186 | Cfb | Complement factor B (Mus musculus) | 97.17 | 2.23 | 85004 | 1 | 1330466.38 | 2200167 | 1937148.5 | 2567504.25 | 2588772.25 | 1469616.5 | 1351676 | 1264413 | 1457486.62 | 1188178.88 | 3 | **0.634** |
| P04202 | Tgfb1 | Transforming growth factor beta-1 proprotein (Mus musculus) | 47.23 | 5.13 | 44310 | 1 | 314609.41 | 0 | 348469.81 | 286761.5 | 187589.41 | 0 | 495355.97 | 396885.28 | 0 | 283868.28 | 13 | **1.034** |
| P04256 | Hnrnpa1 | Heterogeneous nuclear ribonucleoprotein A1 (Rattus norvegicus) | 1480.95 | 58.75 | 34212 | 22 | 99473448 | 156288640 | 177085792 | 134635216 | 152753968 | 154278784 | 135263184 | 137484768 | 125535648 | 137118480 | 10 | **0.958** |
| P04550 | Ptms | Parathymosin (Rattus norvegicus) | 188.05 | 22.55 | 11559 | 3 | 25735378 | 16352121 | 21517764 | 14610602 | 17826192 | 18840224 | 22750632 | 22450344 | 24588584 | 21447746 | 7 | **1.146** |
| P04692 | Tpm1 | Tropomyosin alpha-1 chain (Rattus norvegicus) | 351.92 | 15.85 | 32681 | 8 | 102405760 | 79210584 | 79201632 | 63748472 | 60275044 | 71617680 | 82993024 | 70489888 | 84005712 | 71963960 | 11 | **0.990** |
| P04916 | Rbp4 | Retinol-binding protein 4 (Rattus norvegicus) | 44.52 | 3.48 | 23220 | 1 | 0 | 4982805.5 | 4381361.5 | 11071416 | 6038478.5 | 0 | 0 | 0 | 0 | 0 | 15 | **0.000** |
| P05063 | Aldoc | Fructose-bisphosphate aldolase C (Mus musculus) | 990.22 | 46.28 | 39395 | 17 | 289117024 | 224758496 | 256275712 | 229579776 | 269885632 | 333192608 | 325243232 | 299800960 | 326499136 | 318251200 | 0 | **1.263** |
| P05064 | Aldoa | Fructose-bisphosphate aldolase A (Mus musculus) | 2177.77 | 85.44 | 39356 | 33 | 1042111808 | 962607552 | 1011378176 | 926057088 | 1028945536 | 1160460928 | 947779840 | 993951296 | 1023979712 | 1019461312 | 11 | **1.035** |
| P05132 | Prkaca | cAMP-dependent protein kinase catalytic subunit alpha (Mus musculus) | 471.41 | 30.48 | 40571 | 8 | 21879984 | 21198976 | 21461836 | 19651344 | 15691775 | 26440928 | 17814222 | 25867236 | 23612200 | 24882852 | 4 | **1.188** |
| P05201 | Got1 | Aspartate aminotransferase, cytoplasmic (Mus musculus) | 1336.71 | 60.29 | 46248 | 22 | 117103864 | 134346400 | 118649360 | 137585264 | 109520224 | 110442216 | 103308520 | 99980704 | 105250800 | 105921064 | 1 | **0.850** |
| P05202 | Got2 | Aspartate aminotransferase, mitochondrial (Mus musculus) | 1460.56 | 58.84 | 47411 | 26 | 305252704 | 235851216 | 244066400 | 279071552 | 218971776 | 274528608 | 243717984 | 258757104 | 258085008 | 263150416 | 11 | **1.012** |
| P05213 | Tuba1b | Tubulin alpha-1B chain (Mus musculus) | 2448.36 | 74.72 | 50152 | 36 | 653985280 | 960073472 | 839683008 | 920406656 | 858907968 | 785021312 | 710738496 | 782241920 | 825162624 | 845222912 | 6 | **0.933** |
| P05555 | Itgam | Integrin alpha-M (Mus musculus) | 1165.56 | 20.99 | 127481 | 17 | 50307568 | 55625972 | 54072076 | 50085580 | 41733024 | 58325884 | 52198704 | 51223872 | 55039172 | 62503844 | 5 | **1.109** |
| P05712 | Rab2a | Ras-related protein Rab-2A (Rattus norvegicus) | 467.14 | 44.81 | 23536 | 9 | 10369232 | 18095260 | 17365738 | 20898884 | 16654089 | 16491658 | 14302152 | 16344049 | 17673192 | 16656934 | 8 | **0.977** |
| P05765 | Rps21 | 40S ribosomal protein S21 (Rattus norvegicus) | 268.72 | 57.83 | 9127 | 4 | 34097428 | 17930064 | 22278390 | 17869752 | 29200140 | 32078282 | 25036784 | 28846536 | 24938880 | 27862112 | 9 | **1.143** |
| P06151 | Ldha | L-lactate dehydrogenase A chain (Mus musculus) | 2414.12 | 86.14 | 36499 | 35 | 1362660608 | 1484729984 | 1499876480 | 1387953280 | 1302912384 | 1708862848 | 1523867904 | 1505928832 | 1469969408 | 1479607168 | 4 | **1.092** |
| P06214 | Alad | Delta-aminolevulinic acid dehydratase (Rattus norvegicus) | 284.51 | 19.7 | 36032 | 5 | 8010210 | 5065472 | 5632653.5 | 4010891.75 | 5691865.5 | 7191285 | 4715142 | 3757224.5 | 3120905.5 | 4031601.25 | 6 | **0.803** |
| P06339 | H2-T23 | H-2 class I histocompatibility antigen, D-37 alpha chain (Mus musculus) | 183.77 | 8.68 | 40875 | 3 | 13901631 | 14529112 | 13552805 | 15194174 | 13624856 | 13383388 | 14129388 | 13415324 | 14259084 | 14246950 | 9 | **0.981** |
| P06685 | Atp1a1 | Sodium/potassium-transporting ATPase subunit alpha-1 (Rattus norvegicus) | 1629.15 | 27.66 | 113054 | 25 | 125921832 | 92176576 | 98264088 | 93576776 | 89514896 | 106571528 | 112685440 | 114645136 | 109246544 | 117412872 | 5 | **1.122** |
| P06745 | Gpi | Glucose-6-phosphate isomerase (Mus musculus) | 2093.14 | 61.83 | 62767 | 34 | 240326224 | 205848752 | 242658032 | 221203296 | 244266256 | 265550224 | 252102112 | 270212128 | 279561632 | 261053536 | 0 | **1.151** |

| | | | | | | | | | | | | | | | | | |
|---|---|---|---|---|---|---|---|---|---|---|---|---|---|---|---|---|---|
| P06761 | Hspa5 | Endoplasmic reticulum chaperone BiP (Rattus norvegicus) | 2988.42 | 62.08 | 72347 | 43 | 834373056 | 968720640 | 875070656 | 1010361152 | 866570560 | 800013632 | 775890496 | 826494528 | 904672704 | 888094336 | 6 | **0.921** |
| P06762 | Hmox1 | Heme oxygenase 1 (Rattus norvegicus) | 83.16 | 7.96 | 33006 | 2 | 3171267 | 2152082.5 | 626794.06 | 0 | 2605123.5 | 3260851.25 | 2936128 | 820745.94 | 3312419.5 | 2031193.88 | 7 | **1.445** |
| P06795 | Abcb1b | Multidrug resistance protein 1B (Mus musculus) | 76.99 | 1.72 | 140994 | 1 | 129495.48 | 119119.99 | 187363.22 | 0 | 109211.33 | 220433.33 | 217494.84 | 293832.41 | 250583.89 | 190829.22 | 0 | **2.152** |
| P06797 | Ctsl | Cathepsin L1 (Mus musculus) | 198.14 | 14.37 | 37547 | 3 | 3719251.5 | 3494819.5 | 4076310.5 | 3814310.75 | 5319546.5 | 3272075.75 | 4181364.75 | 4169006 | 4299606 | 4281599 | 9 | **0.989** |
| P06800 | Ptprc | Receptor-type tyrosine-protein phosphatase C (Mus musculus) | 1360.7 | 23.51 | 144837 | 24 | 46689524 | 67207472 | 64880500 | 69301320 | 67512768 | 50838480 | 59399516 | 66191848 | 66842772 | 69964552 | 11 | **0.993** |
| P06801 | Me1 | NADP-dependent malic enzyme (Mus musculus) | 199.32 | 11.54 | 63954 | 3 | 2092376 | 1468467.5 | 1811592.12 | 1660795.5 | 1773850.75 | 2884075 | 2215113.25 | 2397950 | 2035250.25 | 1883195.12 | 2 | **1.296** |
| P07091 | S100a4 | Protein S100-A4 (Mus musculus) | 374.39 | 42.57 | 11721 | 6 | 105417704 | 91662048 | 108165168 | 94597776 | 74628984 | 98327896 | 100396024 | 75729360 | 126587320 | 80186896 | 12 | **1.014** |
| P07356 | Anxa2 | Annexin A2 (Mus musculus) | 1481.47 | 67.26 | 38676 | 24 | 197898688 | 242911072 | 243364816 | 243485408 | 235647136 | 203033456 | 218620608 | 214174384 | 227333312 | 259950672 | 9 | **0.965** |
| P07742 | Rrm1 | Ribonucleoside-diphosphate reductase large subunit (Mus musculus) | 654.91 | 22.6 | 90210 | 12 | 11111167 | 12701613 | 13693502 | 11832731 | 9114881 | 14005490 | 12669551 | 11581960 | 11620768 | 11033736 | 12 | **1.042** |
| P07901 | Hsp90aa1 | Heat shock protein HSP 90-alpha (Mus musculus) 1 | 2850.7 | 59.62 | 84788 | 44 | 541586240 | 636753728 | 575622912 | 706375488 | 575365248 | 592581184 | 568659712 | 577605632 | 591946304 | 617726208 | 12 | **0.971** |
| P08003 | Pdia4 | Protein disulfide-isomerase A4 (Mus musculus) | 2023.13 | 50.31 | 71982 | 33 | 125582728 | 131723232 | 131073264 | 144485472 | 140080192 | 135609264 | 142703968 | 145368864 | 141645008 | 148557408 | 4 | **1.061** |
| P08030 | Aprt | Adenine phosphoribosyltransferase (Mus musculus) | 853.12 | 75 | 19724 | 12 | 47590992 | 58077788 | 59437944 | 56958340 | 59567484 | 58088528 | 43275256 | 49543388 | 44138748 | 49220956 | 5 | **0.867** |
| P08101 | Fcgr2 | Low affinity immunoglobulin gamma Fc region receptor II (Mus musculus) | 185.68 | 14.24 | 36695 | 4 | 9299542 | 8675824 | 8485995 | 8261789 | 7948324.5 | 9310710 | 11419042 | 6497470 | 7386535 | 6786685.5 | 10 | **0.970** |
| P08103 | Hck | Tyrosine-protein kinase HCK (Mus musculus) | 224.23 | 7.63 | 59129 | 4 | 3846644.75 | 4677206 | 4425508 | 4877067 | 4127357.5 | 4060741.75 | 4425778.5 | 4785899.5 | 4810374 | 4954428.5 | 8 | **1.049** |
| P08113 | Hsp90b1 | Endoplasmin (Mus musculus) | 2910.68 | 53.74 | 92476 | 46 | 500644352 | 513478912 | 491748192 | 486387872 | 499512544 | 595737152 | 475380960 | 587404608 | 580325440 | 584509888 | 5 | **1.133** |
| P08121 | Col3a1 | Collagen alpha-1(III) chain (Mus musculus) | 121.31 | 1.91 | 138943 | 2 | 0 | 3084623.5 | 2536533 | 3360617 | 2549808.75 | 1103007.38 | 1457105.5 | 1182192.38 | 611931.12 | 1153192.25 | 5 | **0.478** |
| P08207 | S100a10 | Protein S100-A10 (Mus musculus) | 128.75 | 27.84 | 11186 | 2 | 7231029 | 6328998 | 6563731 | 5955377 | 6221763.5 | 3580083.75 | 4476409 | 4470479 | 4727259.5 | 12751378 | 5 | **0.929** |
| P08228 | Sod1 | Superoxide dismutase [Cu-Zn] (Mus musculus) | 207.36 | 25.32 | 15943 | 3 | 35989644 | 21394508 | 23559204 | 25583612 | 37062376 | 19427500 | 20034056 | 20427674 | 20957460 | 21156188 | 0 | **0.710** |
| P08249 | Mdh2 | Malate dehydrogenase, mitochondrial (Mus musculus) | 1294.35 | 65.68 | 35611 | 20 | 184394672 | 251144736 | 234576336 | 245180528 | 223067872 | 195104528 | 154962544 | 179700592 | 189820672 | 183278000 | 2 | **0.793** |
| P08556 | Nras | GTPase NRas (Mus musculus) | 98.3 | 19.05 | 21199 | 2 | 3345554.25 | 4755133.5 | 5716865 | 5627112 | 4914942 | 4502278 | 3473212.25 | 4654683 | 4272891.5 | 3536487.5 | 5 | **0.839** |
| P08649 | C4 | Complement C4 (Rattus norvegicus) | 370.05 | 3.68 | 192163 | 7 | 17382548 | 26587448 | 21658132 | 29866274 | 20663538 | 21240858 | 16710738 | 18140266 | 15872952 | 16533257 | 3 | **0.762** |
| P08752 | Gnai2 | Guanine nucleotide-binding protein G(i) subunit alpha-2 (Mus musculus) | 1520 | 63.38 | 40489 | 21 | 68502640 | 85048656 | 97562240 | 83799400 | 80849344 | 89878488 | 83578472 | 95472992 | 87235248 | 97589496 | 6 | **1.091** |
| P08753 | Gnai3 | Guanine nucleotide-binding protein G(k) subunit alpha (Rattus norvegicus) | 880.24 | 46.05 | 40522 | 13 | 18968000 | 38572008 | 40307136 | 36489280 | 33936584 | 36988904 | 33700448 | 37685784 | 37853168 | 40025116 | 11 | **1.107** |
| P08775 | Polr2a | DNA-directed RNA polymerase II subunit RPB1 (Mus musculus) | 253.96 | 5.74 | 217176 | 6 | 1958891.5 | 5300913 | 5154773 | 3155136 | 3541033.5 | 3962638.5 | 3921455.75 | 3621217.75 | 4000473.25 | 4594881.5 | 10 | **1.052** |
| P08905 | Lyz2 | Lysozyme C-2 (Mus musculus) | 156.84 | 27.03 | 16689 | 2 | 5689185 | 4450389.5 | 2502938.75 | 5203031.5 | 6528614 | 2085914.62 | 2497829.5 | 2760278.5 | 3930143.5 | 3146039.25 | 3 | **0.592** |
| P09041 | Pgk2 | Phosphoglycerate kinase 2 (Mus musculus) | 487.17 | 19.42 | 44853 | 8 | 132895728 | 63334732 | 60005244 | 52467512 | 68792688 | 113015400 | 102956544 | 95024552 | 104819992 | 98577016 | 5 | **1.363** |
| P09055 | Itgb1 | Integrin beta-1 (Mus musculus) | 707.21 | 25.56 | 88231 | 15 | 26048682 | 20074998 | 20959790 | 15950080 | 15900980 | 23680812 | 25766408 | 26751436 | 23022218 | 24257182 | 4 | **1.248** |
| P09103 | P4hb | Protein disulfide-isomerase (Mus musculus) | 2083.86 | 58.74 | 57058 | 31 | 232263232 | 240645360 | 207470880 | 279870944 | 194404240 | 243277184 | 214730928 | 207028688 | 189209600 | 183643088 | 7 | **0.899** |
| P09330 | Prps2 | Ribose-phosphate pyrophosphokinase 2 (Rattus norvegicus) | 740.23 | 46.86 | 34813 | 13 | 31539022 | 43842928 | 40090012 | 39757140 | 36951836 | 33920144 | 36562308 | 32642884 | 35231368 | 34025084 | 5 | **0.897** |
| P09405 | Ncl | Nucleolin (Mus musculus) | 2877.89 | 46.39 | 76723 | 48 | 646970880 | 737028160 | 748263296 | 632975744 | 739612416 | 797281856 | 748400128 | 855526400 | 842248256 | 862190976 | 0 | **1.171** |
| P09411 | Pgk1 | Phosphoglycerate kinase 1 (Mus musculus) | 3209.94 | 78.42 | 44550 | 49 | 1085991424 | 964527232 | 916579456 | 893784064 | 973289856 | 1147771008 | 1004080256 | 1053363648 | 1111173888 | 1132560896 | 2 | **1.127** |
| P09495 | Tpm4 | Tropomyosin alpha-4 chain (Rattus norvegicus) | 820.98 | 37.9 | 28510 | 14 | 163723840 | 144634464 | 128235168 | 126418848 | 143287216 | 114016120 | 131063656 | 109925624 | 128251768 | 118314888 | 4 | **0.852** |
| P09527 | Rab7a | Ras-related protein Rab-7a (Rattus norvegicus) | 992.92 | 74.4 | 23504 | 15 | 123756480 | 117489120 | 112909216 | 107603096 | 110388048 | 139209680 | 111998872 | 106945584 | 102500768 | 107330640 | 7 | **0.993** |
| P09528 | Fth1 | Ferritin heavy chain (Mus musculus) | 436.21 | 63.74 | 21067 | 8 | 55953120 | 55961524 | 59794052 | 44794384 | 64347288 | 53081964 | 45170396 | 53389172 | 55352084 | 50757192 | 5 | **0.918** |
| P09581 | Csf1r | Macrophage colony-stimulating factor 1 receptor (Mus musculus) | 601.33 | 17.4 | 109179 | 12 | 21018306 | 18947928 | 22020824 | 16419172 | 16340135 | 24114492 | 25730374 | 25241064 | 32219154 | 28738920 | 0 | **1.436** |
| P09671 | Sod2 | Superoxide dismutase [Mn], mitochondrial (Mus musculus) | 385.13 | 39.64 | 24603 | 7 | 51529304 | 48392372 | 46012536 | 49110516 | 50424676 | 48414508 | 42593080 | 53800560 | 54897712 | 53769248 | 8 | **1.033** |
| P0C0A1 | Vps25 | Vacuolar protein-sorting-associated protein 25 (Rattus norvegicus) | 46.1 | 8.52 | 20762 | 1 | 744418.62 | 0 | 1019478.94 | 1197363.38 | 803205.88 | 950707.19 | 854086.38 | 583677.25 | 819944.12 | 0 | 11 | **0.852** |
| P0C0A2 | Vps36 | Vacuolar protein-sorting-associated protein 36 (Rattus norvegicus) | 160.83 | 9.84 | 43718 | 3 | 3132760.5 | 3496615.25 | 3435456.5 | 5440697.5 | 2153306.5 | 3636666 | 4101284.25 | 4813686 | 4425550 | 4049208 | 5 | **1.191** |
| P0C0R5 | Pik3r4 | Phosphoinositide 3-kinase regulatory subunit 4 (Rattus norvegicus) | 61.67 | 0.96 | 152447 | 1 | 0 | 591130.62 | 541424.12 | 699585.5 | 371763.41 | 548925 | 513669.78 | 497640.25 | 509976.47 | 360402.16 | 10 | **1.103** |
| P0C0S6 | H2afz | Histone H2A.Z (Mus musculus) | 357.44 | 53.91 | 13553 | 5 | 992355840 | 300446016 | 405370176 | 109288096 | 338662976 | 768992704 | 693296000 | 604213568 | 671353664 | 634424000 | 5 | **1.571** |
| P0C5H9 | Manf | Mesencephalic astrocyte-derived neurotrophic factor (Rattus norvegicus) | 43.84 | 6.15 | 20388 | 1 | 2969458.5 | 2207440.75 | 1150477.38 | 2723193.5 | 3006472 | 1226643.38 | 2859669.75 | 1467867.25 | 2353976 | 1671067 | 8 | **0.794** |
| P0C6B1 | Slc35e1 | Solute carrier family 35 member E1 (Rattus norvegicus) | 63.38 | 2.2 | 44384 | 1 | 0 | 912554.19 | 656731.5 | 638579.31 | 617077.31 | 813210.06 | 593785.94 | 821130.31 | 734137.06 | 856335.81 | 8 | **1.352** |
| P0CC09 | Hist2h2aa3 | Histone H2A type 2-A (Rattus norvegicus) aa3 | 496.91 | 57.69 | 14095 | 6 | 997109248 | 290309312 | 397757504 | 108372584 | 344307392 | 791304768 | 688014592 | 621136384 | 650972416 | 647093632 | 5 | **1.590** |
| P0DOV2 | Ifi204 | Interferon-activable protein 204 (Mus musculus) | 124.33 | 4.36 | 69438 | 2 | 2589524.5 | 4854451 | 5602693 | 4335980.5 | 3787169.75 | 5363833 | 4090626 | 4457601.5 | 4482090.5 | 4214975 | 11 | **1.068** |
| P0DP26 | Calm1 | Calmodulin-1 (Mus musculus) | 297.11 | 38.26 | 16838 | 4 | 5527006.5 | 15336732 | 8480538 | 15422295 | 16837778 | 11104074 | 13375993 | 9993856 | 12968568 | 9652162 | 10 | **0.927** |
| P10107 | Anxa1 | Annexin A1 (Mus musculus) | 1659.47 | 72.83 | 38734 | 28 | 119728728 | 105414328 | 89715216 | 96736544 | 82960744 | 107874512 | 107202912 | 104869584 | 90166616 | 92755264 | 10 | **1.017** |
| P10126 | Eef1a1 | Elongation factor 1-alpha 1 (Mus musculus) | 2483.35 | 63.64 | 50114 | 42 | 2466219520 | 2516683776 | 2550276096 | 2682799104 | 2620179968 | 2851312128 | 2375665152 | 2733583104 | 2658929408 | 2718250752 | 6 | **1.039** |
| P10417 | Bcl2 | Apoptosis regulator Bcl-2 (Mus musculus) | 51.37 | 4.24 | 26407 | 1 | 0 | 1043929.5 | 1094112.38 | 794864.44 | 769036.44 | 951172.38 | 0 | 673059.19 | 718223.5 | 722547.75 | 7 | **0.828** |
| P10536 | Rab1b | Ras-related protein Rab-1B (Rattus norvegicus) | 861.21 | 71.14 | 22163 | 12 | 23743284 | 45924180 | 60672008 | 45128976 | 18981588 | 36235184 | 40164060 | 47238744 | 47885904 | 39779632 | 11 | **1.087** |
| P10605 | Ctsb | Cathepsin B (Mus musculus) | 912.86 | 56.05 | 37280 | 14 | 82568344 | 84070256 | 81974736 | 73851120 | 111395160 | 68566552 | 76420832 | 75171376 | 85456320 | 81649512 | 7 | **0.893** |
| P10630 | Eif4a2 | Eukaryotic initiation factor 4A-II (Mus musculus) | 1004.69 | 28.01 | 46402 | 15 | 314967296 | 269545536 | 290669728 | 288932448 | 284920032 | 326535168 | 307431360 | 298790848 | 305671936 | 295416512 | 4 | **1.059** |
| P10639 | Txn | Thioredoxin (Mus musculus) | 437.86 | 55.24 | 11675 | 7 | 46038504 | 79123352 | 86549416 | 72360944 | 91166664 | 94373376 | 69620384 | 86403584 | 80471760 | 87492736 | 9 | **1.115** |
| P10649 | Gstm1 | Glutathione S-transferase Mu 1 (Mus musculus) | 466.46 | 42.66 | 25970 | 8 | 39585776 | 24870022 | 22483554 | 16058366 | 16494454 | 28419150 | 35125892 | 20565414 | 19077802 | 25358200 | 9 | **1.076** |
| P10711 | Tcea1 | Transcription elongation factor A protein 1 (Mus musculus) | 613.05 | 35.22 | 33880 | 9 | 17035754 | 7453654 | 6478488 | 7056481.5 | 14745885 | 13362729 | 14119662 | 13137980 | 14059078 | 13214768 | 10 | **1.287** |
| P10810 | Cd14 | Monocyte differentiation antigen CD14 (Mus musculus) | 515.2 | 20.49 | 39204 | 7 | 16801956 | 20428604 | 18556168 | 21670858 | 19473342 | 15003135 | 13969148 | 13923828 | 16672033 | 15345096 | 0 | **0.773** |
| P10818 | Cox6a1 | Cytochrome c oxidase subunit 6A1, mitochondrial (Rattus norvegicus) | 46.62 | 18.92 | 12301 | 1 | 0 | 0 | 63753.31 | 437139.19 | 242924.45 | 89327.95 | 0 | 0 | 0 | 0 | 17 | **0.120** |
| P10833 | Rras | Ras-related protein R-Ras (Mus musculus) | 204.38 | 21.1 | 23764 | 5 | 1228481.38 | 3480239.5 | 3084049.25 | 2652499.5 | 1478654.38 | 3824553.5 | 987800.75 | 4304741 | 3584362.5 | 3546857.5 | 5 | **1.363** |
| P10852 | Slc3a2 | 4F2 cell-surface antigen heavy chain (Mus musculus) | 684.55 | 29.85 | 58337 | 12 | 35131460 | 29372646 | 33723908 | 32223972 | 31917884 | 43692020 | 38452636 | 42802300 | 42271232 | 42973972 | 0 | **1.295** |
| P10853 | Hist1h2bf | Histone H2B type 1-F/J/L (Mus musculus) bf | 1085.69 | 62.7 | 13936 | 17 | 1530760704 | 645957376 | 432904192 | 604414080 | 656620800 | 784460160 | 866814144 | 647187904 | 657506944 | 700600384 | 6 | **0.945** |
| P10854 | Hist1h2bm | Histone H2B type 1-M (Mus musculus) bm | 1136.43 | 62.7 | 13936 | 18 | 1533204224 | 651593920 | 436002048 | 606636544 | 660084224 | 785857792 | 870168768 | 647045376 | 658995840 | 700295424 | 8 | **0.942** |
| P10922 | H1f0 | Histone H1.0 (Mus musculus) | 196.31 | 20.62 | 20861 | 4 | 28009318 | 12082644 | 11312744 | 12587980 | 8580469 | 13466413 | 20132910 | 11730436 | 18370042 | 15897070 | 7 | **1.097** |
| P11030 | Dbi | Acyl-CoA-binding protein (Rattus norvegicus) | 67.82 | 20.69 | 10027 | 1 | 7136550.5 | 4026707.75 | 3304629.25 | 3465705 | 6339626 | 4523395.5 | 2714021.5 | 3971303.75 | 4895309 | 4196571 | 11 | **0.836** |
| P11031 | Sub1 | Activated RNA polymerase II transcriptional coactivator p15 (Mus musculus) | 211.66 | 44.09 | 14427 | 5 | 13193966 | 14891411 | 21269348 | 15254901 | 20806000 | 14037646 | 15739196 | 13783407 | 17317032 | 16807756 | 11 | **0.909** |
| P11103 | Parp1 | Poly [ADP-ribose] polymerase 1 (Mus musculus) | 1010.12 | 22.31 | 113100 | 16 | 33654196 | 29450012 | 30741168 | 28525054 | 30586070 | 36586468 | 33840212 | 30465906 | 34471140 | 36159124 | 3 | **1.121** |
| P11152 | Lpl | Lipoprotein lipase (Mus musculus) | 42.24 | 3.59 | 53109 | 1 | 367128.19 | 295677.62 | 310697.94 | 518800.19 | 289952.03 | 278275.28 | 223802.84 | 176484.03 | 120022.11 | 156361.39 | 0 | **0.536** |
| P11157 | Rrm2 | Ribonucleoside-diphosphate reductase subunit M2 (Mus musculus) | 452.82 | 27.44 | 45096 | 7 | 6736523 | 7712483.5 | 9194521 | 8953686 | 8625940 | 7969329.5 | 6525913.5 | 6529465 | 7447987.5 | 6582203 | 3 | **0.850** |
| P11167 | Slc2a1 | Solute carrier family 2, facilitated glucose transporter member 1 (Rattus norvegicus) | 290.61 | 13.21 | 53963 | 6 | 4831800 | 6579184 | 6660739.5 | 7801286 | 6867642 | 8764583 | 7670329 | 6290730 | 7210874.5 | 5751007 | 10 | **1.090** |
| P11240 | Cox5a | Cytochrome c oxidase subunit 5A, mitochondrial (Rattus norvegicus) | 204.85 | 32.19 | 16130 | 4 | 35023172 | 20856900 | 14939632 | 21112752 | 19411632 | 19459548 | 24372432 | 16675182 | 23333242 | 20785416 | 12 | **0.940** |
| P11345 | Raf1 | RAF proto-oncogene serine/threonine-protein kinase (Rattus norvegicus) | 89.78 | 3.55 | 72928 | 2 | 0 | 1252074.62 | 1138108.88 | 1119595.88 | 955603.88 | 1380226.12 | 1214304.75 | 1132678.25 | 998543.94 | 1111118.62 | 9 | **1.307** |
| P11438 | Lamp1 | Lysosome-associated membrane glycoprotein 1 (Mus musculus) | 623.69 | 28.33 | 43865 | 9 | 51990440 | 72632288 | 85978304 | 69628408 | 88067248 | 93866744 | 83007336 | 89963920 | 98172808 | 100342136 | 2 | **1.264** |
| P11440 | Cdk1 | Cyclin-dependent kinase 1 (Mus musculus) | 764.52 | 49.16 | 34107 | 14 | 22547130 | 17055754 | 21492424 | 15336100 | 21691624 | 25143346 | 29980882 | 30479112 | 30259872 | 29939620 | 0 | **1.486** |
| P11466 | Crot | Peroxisomal carnitine O-octanoyltransferase (Rattus norvegicus) | 44.28 | 2.78 | 70302 | 1 | 0 | 264519 | 0 | 0 | 0 | 0 | 0 | 0 | 0 | 0 | 30 | **0.000** |
| P11497 | Acaca | Acetyl-CoA carboxylase 1 (Rattus norvegicus) | 213.76 | 3.41 | 265194 | 5 | 2755209.75 | 3191019.75 | 1777254.62 | 1932338.88 | 2761432 | 2255438.75 | 1553178.62 | 1729046.12 | 2572295.25 | 2629102 | 6 | **0.865** |
| P11499 | Hsp90ab1 | Heat shock protein HSP 90-beta (Mus musculus) 1 | 3869.41 | 68.78 | 83281 | 58 | 1148492032 | 1288080896 | 1238298240 | 1412441216 | 1139300992 | 1280189440 | 1210246400 | 1218915712 | 1285920128 | 1341196544 | 11 | **1.018** |
| P11680 | Cfp | Properdin (Mus musculus) | 141.6 | 5.6 | 50327 | 2 | 2010312 | 1652909 | 1698509.88 | 2088846.5 | 2311353.5 | 1347231.25 | 1518248.75 | 1339010.5 | 1528172.25 | 1627252.75 | 0 | **0.754** |
| P11688 | Itga5 | Integrin alpha-5 (Mus musculus) | 136.93 | 3.32 | 115043 | 3 | 6179218 | 3901717 | 4844803.5 | 2583303.5 | 4677529 | 6231262 | 5837991 | 5460236 | 4943875 | 6174393 | 4 | **1.291** |
| P11835 | Itgb2 | Integrin beta-2 (Mus musculus) | 1155.14 | 28.02 | 85026 | 17 | 28184676 | 21639284 | 21394980 | 22863270 | 23146522 | 20602424 | 27063126 | 22846852 | 25994430 | 23127928 | 12 | **1.021** |
| P11881 | Itpr1 | Inositol 1,4,5-trisphosphate receptor type 1 (Mus musculus) | 100.44 | 0.73 | 313167 | 2 | 344649.69 | 1096207 | 1153530.62 | 1194980.75 | 735206.75 | 1344451.62 | 1522035.38 | 1136168.88 | 1010053 | 1075355.12 | 8 | **1.346** |
| P11915 | Scp2 | Non-specific lipid-transfer protein (Rattus norvegicus) | 91.73 | 3.47 | 58813 | 2 | 3845159 | 1262718.75 | 2371180 | 2193997 | 2614102.25 | 2664917 | 2482230.5 | 2592191.25 | 2759292.5 | 2564341.25 | 8 | **1.063** |
| P11928 | Oas1a | 2'-5'-oligoadenylate synthase 1A (Mus musculus) | 85.12 | 10.08 | 42429 | 2 | 1742244.25 | 624245.75 | 1723920.75 | 0 | 856371.38 | 2041777.5 | 2201820 | 1293864.75 | 2506547.25 | 2098335.75 | 2 | **2.050** |
| P11980 | Pkm | Pyruvate kinase PKM (Rattus norvegicus) | 2860.88 | 55.37 | 57818 | 40 | 1603333376 | 1454209664 | 1555886720 | 1608545920 | 1458948480 | 1815413248 | 1659335552 | 1510676736 | 1539624832 | 1552870272 | 9 | **1.052** |
| P11983 | Tcp1 | T-complex protein 1 subunit alpha (Mus musculus) | 2407.39 | 71.04 | 60449 | 34 | 153128080 | 173340240 | 160716672 | 176988400 | 151747920 | 162758288 | 168309440 | 151867040 | 159282704 | 159433616 | 11 | **0.983** |
| P12075 | Cox5b | Cytochrome c oxidase subunit 5B, mitochondrial (Rattus norvegicus) | 48.43 | 8.53 | 13915 | 1 | 2200902 | 1234282 | 970030.62 | 1578980.88 | 0 | 1752240.25 | 1717052.75 | 1394253 | 0 | 1565396.25 | 11 | **1.074** |
| P12265 | Gusb | Beta-glucuronidase (Mus musculus) | 1044.45 | 37.65 | 74195 | 20 | 53648020 | 44741432 | 55198400 | 43546464 | 63777448 | 55175232 | 57912896 | 53642380 | 60167756 | 60381912 | 8 | **1.101** |
| P12367 | Prkar2a | cAMP-dependent protein kinase type II-alpha regulatory subunit (Mus musculus) | 150.97 | 9.48 | 45389 | 3 | 1966679.25 | 1943527.75 | 962502.12 | 1310065.75 | 1414886.25 | 2215557.25 | 2281403 | 1849227.75 | 1649087.25 | 1755849.62 | 6 | **1.283** |
| P12382 | Pfkl | ATP-dependent 6-phosphofructokinase, liver type (Mus musculus) | 683.31 | 22.05 | 85360 | 12 | 27340018 | 36731856 | 38302792 | 35369868 | 33576940 | 37913024 | 31963400 | 38629448 | 34408440 | 38348536 | 8 | **1.058** |
| P12785 | Fasn | Fatty acid synthase (Rattus norvegicus) | 1863.96 | 16.25 | 272650 | 33 | 41925288 | 56612492 | 62239652 | 58579320 | 55185476 | 50736796 | 54882916 | 65450656 | 70184576 | 71658464 | 8 | **1.140** |
| P12970 | Rpl7a | 60S ribosomal protein L7a (Mus musculus) | 1087.08 | 52.26 | 29977 | 18 | 117315728 | 95617720 | 98626464 | 102276992 | 97726656 | 116446312 | 124404056 | 114129888 | 121277032 | 113869552 | 3 | **1.154** |

| | | | | | | | | | | | | | | | | | |
|---|---|---|---|---|---|---|---|---|---|---|---|---|---|---|---|---|---|
| P13020 | Gsn | Gelsolin (Mus musculus) | 1472.83 | 42.56 | 85942 | 24 | 137309024 | 121233416 | 117093624 | 129330256 | 144686176 | 111846048 | 111196136 | 102997000 | 104405832 | 100769120 | 0 | **0.818** |
| P13084 | Npm1 | Nucleophosmin (Rattus norvegicus) | 1072.21 | 54.45 | 32560 | 15 | 172933056 | 118701856 | 169417792 | 126606176 | 194715520 | 110338656 | 126292024 | 148157984 | 166923120 | 161036928 | 7 | **0.911** |
| P13439 | Umps | Uridine 5'-monophosphate synthase (Mus musculus) | 764.21 | 33.89 | 52292 | 13 | 16793204 | 17240022 | 17915862 | 14988556 | 17422246 | 19153388 | 19969980 | 18963840 | 18851082 | 19160568 | 0 | **1.139** |
| P13471 | Rps14 | 40S ribosomal protein S14 (Rattus norvegicus) | 554.14 | 66.23 | 16259 | 10 | 65601056 | 44755120 | 53597988 | 55293564 | 63058316 | 72247256 | 53513064 | 44999112 | 62392456 | 45622232 | 11 | **0.987** |
| P13597 | Icam1 | Intercellular adhesion molecule 1 (Mus musculus) | 219.88 | 12.29 | 58844 | 5 | 4163169.5 | 3805277.25 | 4075741.25 | 3871283.75 | 4054873 | 6224373 | 1406409.75 | 4009729 | 2793353 | 2268108 | 7 | **0.836** |
| P13832 | Rlc-a | Myosin regulatory light chain RLC-A (Rattus norvegicus) | 381.85 | 57.56 | 19895 | 7 | 20414408 | 25507946 | 19998292 | 31913778 | 21289080 | 27878108 | 29407432 | 24967904 | 27547418 | 27289204 | 6 | **1.151** |
| P13864 | Dnmt1 | DNA (cytosine-5)-methyltransferase 1 (Mus musculus) | 1109.34 | 17.04 | 183189 | 21 | 35741824 | 32496392 | 32704620 | 33504218 | 33286866 | 32856892 | 30001202 | 34048568 | 32317016 | 33064700 | 8 | **0.968** |
| P14046 | A1i3 | Alpha-1-inhibitor 3 (Rattus norvegicus) | 217.4 | 2.03 | 163773 | 4 | 57492500 | 111411656 | 98712768 | 144272320 | 122000944 | 86958200 | 87075136 | 80530048 | 71395256 | 77400256 | 5 | **0.756** |
| P14069 | S100a6 | Protein S100-A6 (Mus musculus) | 127.6 | 26.97 | 10051 | 2 | 20374508 | 12076886 | 5154510.5 | 14591929 | 17146406 | 9874882 | 3194262.5 | 0 | 11217901 | 3154057.5 | 2 | **0.396** |
| P14106 | C1qb | Complement C1q subcomponent subunit B (Mus musculus) | 233.55 | 21.74 | 26717 | 5 | 9973442 | 6522290 | 10937561 | 11520158 | 12293603 | 13126660 | 10648389 | 12134538 | 11803712 | 12129249 | 6 | **1.168** |
| P14115 | Rpl27a | 60S ribosomal protein L27a (Mus musculus) | 280.9 | 22.97 | 16605 | 5 | 104491960 | 61935052 | 92013640 | 47929644 | 107449208 | 145907136 | 123162736 | 125160640 | 119520352 | 125282816 | 0 | **1.544** |
| P14131 | Rps16 | 40S ribosomal protein S16 (Mus musculus) | 643 | 64.38 | 16445 | 13 | 203663424 | 213866912 | 216630304 | 276147136 | 205374752 | 213739296 | 180077968 | 205423600 | 211865616 | 189981584 | 6 | **0.897** |
| P14148 | Rpl7 | 60S ribosomal protein L7 (Mus musculus) | 1270.7 | 50.37 | 31420 | 23 | 342045792 | 196462352 | 268268144 | 167569968 | 254372592 | 344234208 | 333991296 | 321995136 | 342357760 | 322602176 | 3 | **1.355** |
| P14152 | Mdh1 | Malate dehydrogenase, cytoplasmic (Mus musculus) | 914.85 | 62.28 | 36511 | 14 | 162605792 | 114923096 | 143823760 | 113458232 | 123506120 | 155943344 | 136316656 | 142092768 | 142921584 | 150246784 | 8 | **1.105** |
| P14206 | Rpsa | 40S ribosomal protein SA (Mus musculus) | 1240.03 | 61.02 | 32838 | 17 | 255166880 | 232166384 | 259659456 | 234321552 | 231987424 | 242648544 | 229031504 | 239576768 | 252173616 | 239735280 | 12 | **0.992** |
| P14211 | Calr | Calreticulin (Mus musculus) | 1359.04 | 56.49 | 47995 | 22 | 428453440 | 493344800 | 525046656 | 501126144 | 484212608 | 502171520 | 446975168 | 571331840 | 597422080 | 603246976 | 5 | **1.119** |
| P14576 | Srp54 | Signal recognition particle 54 kDa protein (Mus musculus) | 200.39 | 16.27 | 55721 | 5 | 5059462 | 2862921.25 | 3455309 | 2977644.5 | 2246373.75 | 3159256 | 3765474.5 | 3893492 | 3329550.25 | 2920225 | 9 | **1.028** |
| P14604 | Echs1 | Enoyl-CoA hydratase, mitochondrial (Rattus norvegicus) | 41.38 | 4.48 | 31516 | 1 | 2084452.25 | 1941222.38 | 1752297.88 | 1134605.25 | 1312450.38 | 1870815.38 | 1873633.62 | 1552989.62 | 1173208.38 | 1332033 | 11 | **0.949** |
| P14685 | Psmd3 | 26S proteasome non-ATPase regulatory subunit 3 (Mus musculus) | 382.69 | 15.47 | 60718 | 7 | 8208220 | 5889570.5 | 9070015 | 9180806 | 10315656 | 13768109 | 11647393 | 12172161 | 11776871 | 11873422 | 0 | **1.435** |
| P14733 | Lmnb1 | Lamin-B1 (Mus musculus) | 1312.09 | 42.01 | 66786 | 23 | 55711460 | 64499460 | 66322732 | 63252284 | 55981752 | 51290252 | 56346776 | 54807004 | 54479140 | 52467972 | 2 | **0.881** |
| P14824 | Anxa6 | Annexin A6 (Mus musculus) | 961.14 | 28.68 | 75885 | 17 | 19312798 | 34315124 | 31875422 | 46931636 | 39790384 | 21574576 | 19461836 | 23489936 | 24900244 | 32167930 | 6 | **0.706** |
| P14869 | Rplp0 | 60S acidic ribosomal protein P0 (Mus musculus) | 983.74 | 67.82 | 34216 | 14 | 268193328 | 239912016 | 249531456 | 209677696 | 228598000 | 297958240 | 252526256 | 270779200 | 256935488 | 264138192 | 3 | **1.122** |
| P15379 | Cd44 | CD44 antigen (Mus musculus) | 330.33 | 7.97 | 85617 | 5 | 14555408 | 15601497 | 19006968 | 11093886 | 9357158 | 17067014 | 16233621 | 18377266 | 16872566 | 18322442 | 5 | **1.248** |
| P15429 | Eno3 | Beta-enolase (Rattus norvegicus) | 595.22 | 25.35 | 47014 | 7 | 86043616 | 79776240 | 84670912 | 72520544 | 86937328 | 96939448 | 40968268 | 77351520 | 74887128 | 72844864 | 8 | **0.885** |
| P15532 | Nme1 | Nucleoside diphosphate kinase A (Mus musculus) | 594.19 | 73.68 | 17208 | 12 | 105969528 | 153749408 | 147206048 | 145329296 | 143620064 | 153842832 | 138792016 | 139196832 | 148269888 | 142376624 | 12 | **1.038** |
| P15535 | B4galt1 | Beta-1,4-galactosyltransferase 1 (Mus musculus) | 89.78 | 4.76 | 44411 | 2 | 0 | 0 | 0 | 448113.28 | 0 | 717992.38 | 447492.03 | 628484.62 | 500348.94 | 548856.94 | 7 | **6.345** |
| P15651 | Acads | Short-chain specific acyl-CoA dehydrogenase, mitochondrial (Rattus norvegicus) | 222.93 | 16.99 | 44765 | 5 | 8825169 | 20134118 | 16159194 | 26264350 | 20489548 | 11413730 | 9752915 | 8635826 | 9564444 | 9718832 | 4 | **0.534** |
| P15791 | Camk2d | Calcium/calmodulin-dependent protein kinase type II subunit delta (Rattus norvegicus) | 304.22 | 14.45 | 60081 | 6 | 7907762.5 | 5995462.5 | 8686771 | 5127845.5 | 6703307 | 10411372 | 9885130 | 9753898 | 10177372 | 10862253 | 0 | **1.484** |
| P15864 | Hist1h1c | Histone H1.2 (Mus musculus) c | 649.22 | 37.26 | 21267 | 11 | 492401088 | 174547904 | 134995264 | 175541200 | 232151312 | 151878464 | 248853216 | 102409880 | 248613664 | 158671760 | 10 | **0.753** |
| P15865 | Hist1h1e | Histone H1.4 (Rattus norvegicus) e | 589.96 | 31.96 | 21987 | 10 | 479070080 | 168897072 | 130669272 | 169195488 | 225084720 | 147468032 | 242094272 | 99156328 | 242262656 | 153885472 | 10 | **0.754** |
| P15920 | Atp6v0a2 | V-type proton ATPase 116 kDa subunit a isoform 2 (Mus musculus) 2 | 48.56 | 1.4 | 98145 | 1 | 85253.11 | 212059.91 | 191151.25 | 353121.97 | 234840.94 | 112317.59 | 308471.12 | 249084.58 | 136038.5 | 267728.81 | 11 | **0.997** |
| P15943 | Aplp2 | Amyloid-like protein 2 (Rattus norvegicus) | 51.26 | 2.61 | 86883 | 1 | 113305.31 | 285332.19 | 215538.23 | 97582.06 | 126245.92 | 347543.69 | 382887.44 | 78500.88 | 203960.73 | 118189.7 | 10 | **1.350** |
| P16036 | Slc25a3 | Phosphate carrier protein, mitochondrial (Rattus norvegicus) | 748.41 | 39.33 | 39445 | 13 | 73843920 | 83044224 | 86774984 | 83256488 | 67842736 | 91299040 | 94364648 | 106143904 | 103554408 | 106365792 | 0 | **1.271** |
| P16045 | Lgals1 | Galectin-1 (Mus musculus) | 931.97 | 86.67 | 14866 | 12 | 254062784 | 210025776 | 217129280 | 231286224 | 295040192 | 258054544 | 227240608 | 240903712 | 236371168 | 236729488 | 10 | **0.993** |
| P16086 | Sptan1 | Spectrin alpha chain, non-erythrocytic 1 (Rattus norvegicus) | 291.67 | 3.11 | 284637 | 5 | 1771989.38 | 1287998.5 | 919346.12 | 2747557.5 | 2152267.5 | 1553058.5 | 1621881.25 | 2483228 | 2282806.75 | 1121158.5 | 12 | **1.021** |
| P16110 | Lgals3 | Galectin-3 (Mus musculus) | 624.38 | 39.02 | 27515 | 10 | 80114712 | 123871952 | 126223392 | 130263408 | 136863232 | 93495840 | 88521744 | 99762560 | 112543528 | 96229632 | 5 | **0.821** |
| P16125 | Ldhb | L-lactate dehydrogenase B chain (Mus musculus) | 120.06 | 5.99 | 36572 | 2 | 165973296 | 153237504 | 164026304 | 140478976 | 128787896 | 148049472 | 129154256 | 133465368 | 146579696 | 149916480 | 8 | **0.940** |
| P16254 | Srp14 | Signal recognition particle 14 kDa protein (Mus musculus) | 56.44 | 11.82 | 12510 | 1 | 300212.75 | 593145.44 | 868807.44 | 0 | 1223260.25 | 736445.06 | 1124452 | 1024172.19 | 1345170.38 | 308902.38 | 7 | **1.520** |
| P16332 | Mut | Methylmalonyl-CoA mutase, mitochondrial (Mus musculus) | 288.76 | 10.29 | 82844 | 6 | 4087549 | 3767616.75 | 3313395.5 | 3889818.5 | 4823309 | 1880161.12 | 3121980 | 5249158 | 5340837 | 4904885.5 | 10 | **1.031** |
| P16409 | Myl3 | Myosin light chain 3 (Rattus norvegicus) | 189.17 | 12 | 22156 | 3 | 0 | 27272994 | 28269292 | 30496032 | 30480174 | 0 | 0 | 29047146 | 32614280 | 32467632 | 12 | **0.808** |
| P16446 | Pitpna | Phosphatidylinositol transfer protein alpha isoform (Rattus norvegicus) | 476.7 | 29.89 | 31907 | 10 | 15310409 | 16092923 | 18422536 | 16798758 | 8824565 | 17682646 | 17664332 | 18885556 | 17654638 | 19072704 | 3 | **1.206** |
| P16460 | Ass1 | Argininosuccinate synthase (Mus musculus) | 898.19 | 38.11 | 46584 | 16 | 37640172 | 52905360 | 55448624 | 43394492 | 48023424 | 54043792 | 54886668 | 52898208 | 52534928 | 54829420 | 7 | **1.134** |
| P16675 | Ctsa | Lysosomal protective protein (Mus musculus) | 1004.02 | 39.66 | 53844 | 15 | 62066828 | 87174920 | 97157352 | 71821712 | 86371936 | 78898696 | 67236480 | 79744064 | 78812576 | 81647736 | 9 | **0.955** |
| P16858 | Gapdh | Glyceraldehyde-3-phosphate dehydrogenase (Mus musculus) | 2507.1 | 81.68 | 35810 | 37 | 2346152704 | 2698561792 | 2890210304 | 2611060992 | 2677200896 | 2589618688 | 2252451072 | 2155498496 | 2656034560 | 2532390912 | 4 | **0.922** |
| P17047 | Lamp2 | Lysosome-associated membrane glycoprotein 2 (Mus musculus) | 339.57 | 15.18 | 45681 | 6 | 64129616 | 115369040 | 107971856 | 97282288 | 109275696 | 115975536 | 105945264 | 112729472 | 117514992 | 113902752 | 5 | **1.146** |
| P17074 | Rps19 | 40S ribosomal protein S19 (Rattus norvegicus) | 464.71 | 49.66 | 16085 | 8 | 114188632 | 59904852 | 46755224 | 73214632 | 42204548 | 69943208 | 106588880 | 65775104 | 69837096 | 57768896 | 10 | **1.100** |
| P17078 | Rpl35 | 60S ribosomal protein L35 (Rattus norvegicus) | 141.22 | 21.95 | 14553 | 3 | 37742420 | 8061734 | 23956934 | 5304917 | 12978617 | 54858616 | 39872980 | 39386296 | 32307782 | 33601092 | 2 | **2.272** |
| P17095 | Hmga1 | High mobility group protein HMG-I/HMG-Y (Mus musculus) | 51.23 | 14.95 | 11614 | 1 | 1336059 | 753171.94 | 950016.62 | 875054.62 | 1232856.12 | 696213.94 | 675834.38 | 611730.88 | 625623.19 | 680603.69 | 0 | **0.639** |
| P17136 | Snrpb | Small nuclear ribonucleoprotein-associated protein B (Rattus norvegicus) | 214.41 | 13.42 | 23656 | 4 | 23788106 | 26357236 | 18829104 | 31934980 | 18690836 | 20294394 | 13428986 | 14447216 | 14935335 | 13789072 | 2 | **0.643** |
| P17182 | Eno1 | Alpha-enolase (Mus musculus) | 2725.87 | 75.58 | 47141 | 36 | 840917632 | 1023320512 | 1075350784 | 1049897280 | 1050764544 | 923075200 | 976624896 | 962868352 | 1078850176 | 1049699264 | 10 | **0.990** |
| P17220 | Psma2 | Proteasome subunit alpha type-2 (Rattus norvegicus) | 499.67 | 49.57 | 25927 | 9 | 17930552 | 16766846 | 12001180 | 13270282 | 12326676 | 14152602 | 10904048 | 11540240 | 12753614 | 13242483 | 7 | **0.866** |
| P17225 | Ptbp1 | Polypyrimidine tract-binding protein 1 (Mus musculus) | 1175.92 | 51.8 | 56478 | 15 | 55977224 | 48075660 | 42085456 | 52589484 | 46571140 | 48751908 | 50200240 | 37958228 | 43433360 | 33117340 | 7 | **0.870** |
| P17426 | Ap2a1 | AP-2 complex subunit alpha-1 (Mus musculus) | 723.09 | 20.47 | 107664 | 15 | 22033118 | 24017876 | 25352158 | 17020830 | 17845682 | 26340406 | 23894972 | 27950960 | 26210074 | 27218944 | 2 | **1.239** |
| P17427 | Ap2a2 | AP-2 complex subunit alpha-2 (Mus musculus) | 776.72 | 24.41 | 104017 | 16 | 16892286 | 18941624 | 22613622 | 17830256 | 14445135 | 21525546 | 20319612 | 22602284 | 21449810 | 22364106 | 5 | **1.193** |
| P17433 | Spi1 | Transcription factor PU.1 (Mus musculus) | 68.11 | 3.68 | 31349 | 1 | 1284659.12 | 1124484.25 | 1066339 | 0 | 1394889.38 | 1293886 | 1685786.25 | 1238496.5 | 1443631.25 | 1290697.38 | 4 | **1.428** |
| P17439 | Gba | Glucosylceramidase (Mus musculus) | 57.18 | 2.52 | 57622 | 1 | 3171942.75 | 4108049.25 | 3780039.75 | 3426538.75 | 3655244.25 | 2102779.75 | 2411952.5 | 2273608 | 3052751.75 | 2861801.75 | 0 | **0.700** |
| P17710 | Hk1 | Hexokinase-1 (Mus musculus) | 1585.46 | 28.64 | 108303 | 25 | 88925240 | 89174744 | 86651464 | 79497096 | 91116552 | 84118448 | 77870192 | 92132864 | 93878304 | 89820032 | 10 | **1.006** |
| P17742 | Ppia | Peptidyl-prolyl cis-trans isomerase A (Mus musculus) | 1182.45 | 75.61 | 17971 | 19 | 297214272 | 202226464 | 188589056 | 300255680 | 286361856 | 265889552 | 210665616 | 224985744 | 239593408 | 228873840 | 10 | **0.918** |
| P17751 | Tpi1 | Triosephosphate isomerase (Mus musculus) | 1352.34 | 71.57 | 32192 | 19 | 650032640 | 565414784 | 546710080 | 596203840 | 550234432 | 677155904 | 595798336 | 611715712 | 592575936 | 537346368 | 10 | **1.036** |
| P17918 | Pcna | Proliferating cell nuclear antigen (Mus musculus) | 1303.69 | 72.8 | 28785 | 18 | 130809768 | 157053312 | 144063424 | 144970608 | 116766040 | 130227784 | 136607760 | 143528368 | 148448528 | 144759888 | 12 | **1.014** |
| P18052 | Ptpra | Receptor-type tyrosine-protein phosphatase alpha (Mus musculus) | 48.59 | 1.45 | 93698 | 1 | 638966.56 | 635995.88 | 0 | 839445.31 | 609899.31 | 681024.69 | 181448.06 | 712925.25 | 770756.75 | 662418.88 | 8 | **1.104** |
| P18155 | Mthfd2 | Bifunctional methylenetetrahydrofolate dehydrogenase/cyclohydrolase, mitochondrial (Mus | 433.62 | 41.43 | 37863 | 9 | 15698362 | 9859296 | 8794305 | 8845407 | 9580124 | 9935885 | 10080469 | 11385098 | 10467972 | 10815968 | 5 | **0.998** |
| P18242 | Ctsd | Cathepsin D (Mus musculus) | 1480.33 | 48.78 | 44954 | 24 | 341039424 | 291833280 | 239003328 | 418200896 | 431196416 | 267192224 | 271063936 | 305947904 | 346540448 | 322748128 | 9 | **0.879** |
| P18266 | Gsk3b | Glycogen synthase kinase-3 beta (Rattus norvegicus) | 63.4 | 10.24 | 46742 | 2 | 1953752.88 | 1907788.5 | 1832926.38 | 1396373.38 | 1505124.5 | 1831499 | 1552626.62 | 2226361.5 | 1713622 | 2144829 | 9 | **1.102** |
| P18292 | F2 | Prothrombin (Rattus norvegicus) | 48.8 | 1.46 | 70412 | 1 | 1155446.75 | 2343524.25 | 1955254.25 | 2409455.25 | 2280818 | 2162900 | 2234059.25 | 2089834.5 | 2285670.5 | 2142753 | 11 | **1.076** |
| P18298 | Mat2a | S-adenosylmethionine synthase isoform type-2 (Rattus norvegicus) | 543.94 | 33.67 | 43716 | 11 | 26329818 | 22280820 | 28346842 | 21245164 | 24792388 | 29500216 | 28148622 | 24406266 | 31021068 | 28589072 | 4 | **1.152** |
| P18572 | Bsg | Basigin (Mus musculus) | 196.92 | 12.08 | 42445 | 4 | 9690908 | 6232871 | 7635540.5 | 5772358 | 7237908 | 6676445 | 9904650 | 5899842 | 7388969.5 | 7067200 | 12 | **1.010** |
| P18596 | Atp2a3 | Sarcoplasmic/endoplasmic reticulum calcium ATPase 3 (Rattus norvegicus) | 673.77 | 10.18 | 116284 | 9 | 30412192 | 27498088 | 28449190 | 21252536 | 20660818 | 27919684 | 31170804 | 30928662 | 32519616 | 31818752 | 2 | **1.203** |
| P18653 | Rps6ka1 | Ribosomal protein S6 kinase alpha-1 (Mus musculus) | 244.56 | 8.56 | 81595 | 5 | 2919490.75 | 4273064.5 | 4706000 | 4671630.5 | 5136124 | 4535426 | 1894869.75 | 4914768.5 | 4911632 | 4423944 | 12 | **0.953** |
| P18654 | Rps6ka3 | Ribosomal protein S6 kinase alpha-3 (Mus musculus) | 283.95 | 8.24 | 83694 | 6 | 7272886.5 | 24607856 | 16356618 | 19987472 | 12119599 | 8327713 | 12035907 | 13886969 | 16021162 | 12901275 | 8 | **0.786** |
| P18760 | Cfl1 | Cofilin-1 (Mus musculus) | 903.92 | 73.49 | 18560 | 13 | 337229376 | 307062656 | 263853440 | 389920672 | 351883424 | 400762976 | 313827424 | 371769120 | 345359296 | 370646848 | 7 | **1.092** |
| P18826 | Phka1 | Phosphorylase b kinase regulatory subunit alpha, skeletal muscle isoform (Mus musculus) | 43.63 | 1.21 | 138825 | 1 | 0 | 541806.31 | 0 | 256069.3 | 320684.66 | 0 | 0 | 0 | 0 | 0 | 20 | **0.000** |
| P19096 | Fasn | Fatty acid synthase (Mus musculus) | 3353.21 | 30.99 | 272428 | 56 | 84290864 | 96697336 | 99600000 | 105418624 | 98992464 | 95255840 | 98753880 | 111872976 | 112135296 | 118616736 | 7 | **1.106** |
| P19139 | Csnk2a1 | Casein kinase II subunit alpha (Rattus norvegicus) | 527.81 | 27.37 | 45073 | 8 | 15374872 | 15528830 | 18372356 | 16062830 | 17269048 | 19344792 | 16762326 | 19236556 | 19296768 | 21095606 | 2 | **1.159** |
| P19253 | Rpl13a | 60S ribosomal protein L13a (Mus musculus) | 382.13 | 33.5 | 23464 | 8 | 129596624 | 41287772 | 103043784 | 21551392 | 82081200 | 176754832 | 145114224 | 130725104 | 132589856 | 122500672 | 1 | **1.874** |
| P19783 | Cox4i1 | Cytochrome c oxidase subunit 4 isoform 1, mitochondrial (Mus musculus) | 193.41 | 30.77 | 19530 | 4 | 29238976 | 19268346 | 16543776 | 22187826 | 15621185 | 13743194 | 18480258 | 17692108 | 19654988 | 17880448 | 9 | **0.850** |
| P19973 | Lsp1 | Lymphocyte-specific protein 1 (Mus musculus) | 650.95 | 43.94 | 36714 | 11 | 52032028 | 57537824 | 42369108 | 60852192 | 70633760 | 42922476 | 46288516 | 40786744 | 48717840 | 44545620 | 4 | **0.788** |
| P20060 | Hexb | Beta-hexosaminidase subunit beta (Mus musculus) | 480.25 | 22.01 | 61116 | 10 | 16232692 | 21637548 | 26506658 | 23988450 | 27170076 | 24582340 | 22086642 | 23506682 | 24415104 | 24534674 | 12 | **1.031** |
| P20069 | Pmpca | Mitochondrial-processing peptidase subunit alpha (Rattus norvegicus) | 45.74 | 2.1 | 58608 | 1 | 431796.97 | 567034.69 | 577726.62 | 367372.84 | 305874.03 | 802898.81 | 639472.31 | 1030475.25 | 653124.69 | 557675.81 | 2 | **1.637** |
| P20108 | Prdx3 | Thioredoxin-dependent peroxide reductase, mitochondrial (Mus musculus) | 386.69 | 36.19 | 28127 | 7 | 32942818 | 30321688 | 28302956 | 33121882 | 30182782 | 36242400 | 38828284 | 36547288 | 36405884 | 39131900 | 0 | **1.208** |
| P20152 | Vim | Vimentin (Mus musculus) | 2960.12 | 75.97 | 53688 | 45 | 902193280 | 809042752 | 763185600 | 843862016 | 821002624 | 824063104 | 878815104 | 890914304 | 914463040 | 901752576 | 5 | **1.065** |
| P20611 | Acp2 | Lysosomal acid phosphatase (Rattus norvegicus) | 74 | 2.84 | 48320 | 1 | 934547.19 | 981434.06 | 1389781.25 | 1225550.25 | 1012822.12 | 1181649.12 | 1074172.75 | 1181313.38 | 1295966.12 | 1070798 | 9 | **1.047** |
| P20664 | Prim1 | DNA primase small subunit (Mus musculus) | 42.82 | 2.64 | 49295 | 1 | 797168.56 | 598570.88 | 696887.25 | 831623 | 897172.56 | 893441.5 | 689277.81 | 1207895.5 | 963490.5 | 1387412.75 | 5 | **1.345** |
| P21107 | Tpm3 | Tropomyosin alpha-3 chain (Mus musculus) | 515.89 | 23.16 | 32994 | 10 | 147905312 | 133635584 | 121570352 | 122139024 | 155233120 | 107267856 | 120087392 | 99904976 | 127599144 | 103653104 | 2 | **0.821** |
| P21279 | Gnaq | Guanine nucleotide-binding protein G(q) subunit alpha (Mus musculus) | 232.25 | 15.04 | 42158 | 5 | 1378977.5 | 3857699.5 | 3877771 | 3142698.25 | 2488307 | 3583171 | 3390838 | 2923882 | 3044637.25 | 2795341 | 12 | **1.067** |
| P21300 | Akr1b7 | Aldose reductase-related protein 1 (Mus musculus) | 294.17 | 13.61 | 35988 | 6 | 62078320 | 49405016 | 62170744 | 61436804 | 62954216 | 64963608 | 59956496 | 74925752 | 78019344 | 85224336 | 4 | **1.218** |
| P21575 | Dnm1 | Dynamin-1 (Rattus norvegicus) | 122.55 | 3.12 | 97295 | 3 | 2931718 | 2387117 | 2403440.5 | 2371863.5 | 3008155.5 | 2155366 | 2296744.75 | 2826490.5 | 2867120.75 | 2984723 | 10 | **1.002** |
| P21619 | Lmnb2 | Lamin-B2 (Mus musculus) | 166.12 | 4.36 | 67318 | 3 | 14147860 | 15446658 | 16440657 | 14339795 | 16068318 | 15652751 | 16677992 | 15750056 | 15637502 | 15203122 | 9 | **1.032** |

| | | | | | | | | | | | | | | | | | | |
|---|---|---|---|---|---|---|---|---|---|---|---|---|---|---|---|---|---|---|
| P21670 | Psma4 | Proteasome subunit alpha type-4 (Rattus norvegicus) | 608.78 | 53.64 | 29498 | 10 | 49329720 | 47299352 | 50596352 | 43745880 | 49234428 | 58955472 | 43827464 | 54176052 | 58124584 | 58470420 | 4 | **1.139** |
| P21708 | Mapk3 | Mitogen-activated protein kinase 3 (Rattus norvegicus) | 412.45 | 30.26 | 43081 | 8 | 16165972 | 10813989 | 16322060 | 17832298 | 13345007 | 12672908 | 14018353 | 15416684 | 9429307 | 16439216 | 9 | **0.913** |
| P21956 | Mfge8 | Lactadherin (Mus musculus) | 488.98 | 29.16 | 51241 | 11 | 25654940 | 22850990 | 23211134 | 20431676 | 14985210 | 25469334 | 21897672 | 22390734 | 20118124 | 23998552 | 12 | **1.063** |
| P21958 | Tap1 | Antigen peptide transporter 1 (Mus musculus) | 285.01 | 12.15 | 78864 | 5 | 1944817.25 | 1883085.75 | 2514395 | 1699571.12 | 1856765 | 1792396.62 | 2977387.5 | 2975106.5 | 3214189.5 | 3254743 | 4 | **1.436** |
| P22366 | Myd88 | Myeloid differentiation primary response protein MyD88 (Mus musculus) | 119.17 | 8.11 | 33753 | 2 | 548165.69 | 784114.69 | 1833955 | 0 | 936021 | 1754406.5 | 974717.81 | 869198.88 | 863524.38 | 836023.31 | 8 | **1.291** |
| P22437 | Ptgs1 | Prostaglandin G/H synthase 1 (Mus musculus) | 765.27 | 32.56 | 69042 | 13 | 15647191 | 27365518 | 30694222 | 26330834 | 19216296 | 24590980 | 21090336 | 22302506 | 21281664 | 22528614 | 10 | **0.937** |
| P22509 | Fbl | rRNA 2'-O-methyltransferase fibrillarin (Rattus norvegicus) | 693.53 | 51.38 | 34222 | 13 | 55220932 | 55583680 | 50693032 | 45378620 | 53576392 | 49118836 | 57049784 | 50170940 | 50512084 | 46912552 | 9 | **0.974** |
| P22682 | Cbl | E3 ubiquitin-protein ligase CBL (Mus musculus) | 115.49 | 3.18 | 100564 | 2 | 1401825.62 | 1562541.12 | 1153903.12 | 923538.62 | 899416.44 | 1765901.75 | 1775363.25 | 1509359.75 | 1425991.12 | 1255828.5 | 4 | **1.301** |
| P22892 | Ap1g1 | AP-1 complex subunit gamma-1 (Mus musculus) | 576.51 | 19.1 | 91350 | 12 | 5827508 | 11036488 | 9778925 | 15096436 | 11614203 | 8327085 | 7494572 | 11364956 | 9449453 | 10112892 | 8 | **0.876** |
| P22907 | Hmbs | Porphobilinogen deaminase (Mus musculus) | 287.28 | 21.61 | 39344 | 4 | 4353905 | 3842782 | 4070556.75 | 3436519.75 | 3822892.5 | 3699720.5 | 4003379 | 3323157 | 3174188 | 3038320 | 4 | **0.883** |
| P22985 | Xdh | Xanthine dehydrogenase/oxidase (Rattus norvegicus) | 293.63 | 5.41 | 146243 | 6 | 3079745 | 5422345 | 4226458 | 6119297.5 | 3254817 | 4359371 | 3532856.5 | 3511576.5 | 3018726.5 | 3648916 | 9 | **0.818** |
| P23116 | Eif3a | Eukaryotic translation initiation factor 3 subunit A (Mus musculus) | 1132.69 | 19.94 | 161936 | 20 | 26527452 | 29295266 | 32238562 | 36046496 | 25298564 | 33699824 | 32349908 | 34578172 | 32460094 | 33680060 | 5 | **1.116** |
| P23198 | Cbx3 | Chromobox protein homolog 3 (Mus musculus) | 639.69 | 42.62 | 20855 | 9 | 17595912 | 33307666 | 24206872 | 38888064 | 28126976 | 21887598 | 23043632 | 21322976 | 25314868 | 22166228 | 6 | **0.800** |
| P23249 | Mov10 | Putative helicase MOV-10 (Mus musculus) | 236.94 | 6.18 | 113583 | 4 | 1860291.88 | 1767005.12 | 2764009.5 | 3020526 | 1257186.38 | 2724427 | 2391321.75 | 2573994 | 2382639.5 | 2634074.5 | 10 | **1.191** |
| P23385 | Grm1 | Metabotropic glutamate receptor 1 (Rattus norvegicus) | 38.59 | 0.58 | 133236 | 1 | 147772.77 | 455547.31 | 464401.31 | 0 | 0 | 833028.38 | 624443.75 | 785761.69 | 747558.69 | 549459.38 | 1 | **3.316** |
| P23475 | Xrcc6 | X-ray repair cross-complementing protein 6 (Mus musculus) | 54.5 | 1.81 | 69484 | 1 | 1044012.12 | 1268079.88 | 762859.5 | 822909.81 | 0 | 924533.12 | 839034.19 | 758772.38 | 815163.44 | 730391.25 | 10 | **1.044** |
| P23492 | Pnp | Purine nucleoside phosphorylase (Mus musculus) | 704.43 | 50.52 | 32277 | 11 | 34748096 | 29190266 | 30037812 | 28673224 | 25830444 | 25291460 | 32912816 | 32428882 | 31449174 | 32268226 | 9 | **1.040** |
| P23591 | Tsta3 | GDP-L-fucose synthase (Mus musculus) | 319.3 | 20.25 | 35878 | 6 | 13434782 | 9988364 | 11924326 | 9994230 | 9307805 | 11714957 | 10958142 | 10775770 | 12495234 | 11018272 | 9 | **1.042** |
| P23780 | Glb1 | Beta-galactosidase (Mus musculus) | 463.79 | 15.77 | 73121 | 8 | 22764638 | 23385564 | 23645104 | 22185520 | 23333822 | 25374072 | 27860266 | 25617050 | 24268578 | 33071734 | 0 | **1.181** |
| P24049 | Rpl17 | 60S ribosomal protein L17 (Rattus norvegicus) | 519.85 | 40.22 | 21397 | 9 | 68248096 | 51718272 | 67720912 | 39396828 | 58072240 | 68284160 | 82296536 | 85728816 | 77283832 | 86099560 | 0 | **1.402** |
| P24051 | Rps27l | 40S ribosomal protein S27-like (Rattus norvegicus) | 134.95 | 25 | 9477 | 2 | 15217438 | 5085674.5 | 9410728 | 5041158 | 11086985 | 12870316 | 10730236 | 10611675 | 10265952 | 9708004 | 9 | **1.182** |
| P24270 | Cat | Catalase (Mus musculus) | 724.89 | 35.86 | 59795 | 13 | 20324756 | 17961902 | 17897028 | 17947264 | 19463672 | 21888046 | 19992578 | 18350236 | 18497716 | 19964640 | 6 | **1.054** |
| P24288 | Bcat1 | Branched-chain-amino-acid aminotransferase, cytosolic (Mus musculus) | 103.29 | 6.22 | 42791 | 2 | 2769221.25 | 2654319.5 | 2067999.25 | 1364070 | 1652454.12 | 2292048.5 | 2313274.25 | 2379450.75 | 2192757 | 2112119.5 | 10 | **1.074** |
| P24369 | Ppib | Peptidyl-prolyl cis-trans isomerase B (Mus musculus) | 553.91 | 41.2 | 23713 | 8 | 37477724 | 54631160 | 52873608 | 51718620 | 50598856 | 48192196 | 46781288 | 43209732 | 50254840 | 49484824 | 5 | **0.962** |
| P24452 | Capg | Macrophage-capping protein (Mus musculus) | 1114.25 | 44.32 | 39240 | 15 | 496035456 | 313135360 | 344037536 | 321234464 | 337447968 | 343757248 | 398474976 | 329669600 | 328798528 | 342622496 | 11 | **0.962** |
| P24527 | Lta4h | Leukotriene A-4 hydrolase (Mus musculus) | 670.27 | 30.11 | 69051 | 14 | 17731188 | 16733538 | 16982998 | 20981266 | 10796016 | 18048840 | 15866509 | 18912206 | 17888020 | 15252364 | 11 | **1.033** |
| P24547 | Impdh2 | Inosine-5'-monophosphate dehydrogenase 2 (Mus musculus) | 1099.27 | 30.93 | 55815 | 15 | 74971072 | 89268904 | 74491744 | 95979424 | 80457944 | 64838928 | 60690248 | 60187756 | 63744396 | 63975000 | 0 | **0.755** |
| P24668 | M6pr | Cation-dependent mannose-6-phosphate receptor (Mus musculus) | 323.25 | 24.82 | 31172 | 4 | 9403911 | 11008829 | 10391964 | 12275437 | 11470308 | 4984128.5 | 7456842 | 7402903.5 | 7188308 | 7946741 | 0 | **0.641** |
| P24860 | Ccnb1 | G2/mitotic-specific cyclin-B1 (Mus musculus) | 38.37 | 1.86 | 48052 | 1 | 986240.19 | 831260.31 | 690876.25 | 0 | 0 | 939458.62 | 976079.31 | 1147791.88 | 0 | 0 | 13 | **1.221** |
| P25093 | Fah | Fumarylacetoacetase (Rattus norvegicus) | 41.64 | 2.15 | 45976 | 1 | 1656855.88 | 3312706.75 | 2680932.75 | 3571315 | 3255428.5 | 534575.81 | 1723114 | 2337149.75 | 1893783.62 | 2168163.25 | 4 | **0.598** |
| P25113 | Pgam1 | Phosphoglycerate mutase 1 (Rattus norvegicus) | 1406.39 | 76.38 | 28832 | 20 | 244777024 | 171882112 | 196667056 | 139394208 | 164923040 | 258071152 | 202336960 | 205612832 | 200576240 | 192038192 | 5 | **1.154** |
| P25206 | Mcm3 | DNA replication licensing factor MCM3 (Mus musculus) | 1601.56 | 39.04 | 91546 | 25 | 89971112 | 80576016 | 71570856 | 76796864 | 78305688 | 82668128 | 75580944 | 72862560 | 75360304 | 75547864 | 8 | **0.962** |
| P25286 | Atp6v0a1 | V-type proton ATPase 116 kDa subunit a isoform 1 (Rattus norvegicus) 1 | 48.53 | 1.31 | 96328 | 1 | 395829.69 | 463001.47 | 549744.31 | 0 | 399165.66 | 525461.12 | 849855.25 | 502241.28 | 463960.84 | 528004.81 | 4 | **1.587** |
| P25444 | Rps2 | 40S ribosomal protein S2 (Mus musculus) | 925.21 | 57.34 | 31231 | 17 | 202255088 | 150130080 | 168148656 | 154744528 | 160197488 | 212384992 | 171200416 | 190910064 | 172331168 | 183934560 | 4 | **1.114** |
| P25799 | Nfkb1 | Nuclear factor NF-kappa-B p105 subunit (Mus musculus) | 339.5 | 6.49 | 105615 | 6 | 9944098 | 5702794 | 7872977 | 8038838 | 7727170 | 9669992 | 8817122 | 8643724 | 9487941 | 9428709 | 5 | **1.172** |
| P25911 | Lyn | Tyrosine-protein kinase Lyn (Mus musculus) | 462.4 | 16.41 | 58812 | 7 | 9007806 | 11413495 | 11704219 | 9717936 | 6670838 | 10491726 | 11460481 | 10890296 | 11469683 | 11633778 | 7 | **1.153** |
| P26039 | Tln1 | Talin-1 (Mus musculus) | 7562.97 | 60.92 | 269821 | 107 | 489147488 | 388984896 | 416562272 | 374639616 | 423618848 | 482942272 | 485743264 | 508235488 | 478057664 | 485630368 | 4 | **1.166** |
| P26040 | Ezr | Ezrin (Mus musculus) | 1018.6 | 31.57 | 69407 | 19 | 248971824 | 224644192 | 255522352 | 264647360 | 258546736 | 304079520 | 214008112 | 264562624 | 279680640 | 255264976 | 9 | **1.052** |
| P26041 | Msn | Moesin (Mus musculus) | 2746.84 | 67.76 | 67767 | 44 | 572791424 | 486213376 | 488097472 | 528847424 | 545030784 | 637699968 | 516044256 | 532847072 | 559972160 | 546328704 | 7 | **1.066** |
| P26043 | Rdx | Radixin (Mus musculus) | 1076.47 | 31.39 | 68543 | 19 | 282476416 | 227228016 | 261159360 | 271589408 | 277978656 | 327172480 | 230755184 | 274973056 | 293936896 | 266046864 | 9 | **1.055** |
| P26151 | Fcgr1 | High affinity immunoglobulin gamma Fc receptor I (Mus musculus) | 44.82 | 3.71 | 44888 | 1 | 0 | 0 | 577697.69 | 713483.56 | 826204.94 | 0 | 0 | 411341.19 | 500562.38 | 322258.03 | 11 | **0.583** |
| P26231 | Ctnna1 | Catenin alpha-1 (Mus musculus) | 49.43 | 3.2 | 100106 | 1 | 0 | 189819.52 | 139536.5 | 138261.23 | 358515.72 | 111471.58 | 177387.28 | 147211.48 | 177893.84 | 195269.47 | 11 | **0.980** |
| P26284 | Pdha1 | Pyruvate dehydrogenase E1 component subunit alpha, somatic form, mitochondrial (Rattus | 153.9 | 11.79 | 43227 | 3 | 7851119 | 5043665 | 5043657 | 4352342 | 4615256 | 6724592 | 6118933 | 5760381 | 6175793.5 | 5747285.5 | 5 | **1.135** |
| P26350 | Ptma | Prothymosin alpha (Mus musculus) | 599.43 | 38.74 | 12254 | 7 | 122256512 | 93595720 | 125496904 | 81353488 | 160468336 | 151063120 | 138114176 | 166319808 | 141638848 | 167620032 | 3 | **1.311** |
| P26369 | U2af2 | Splicing factor U2AF 65 kDa subunit (Mus musculus) | 753.17 | 44.42 | 53517 | 13 | 21100032 | 37640004 | 38304592 | 30342754 | 37715912 | 40927160 | 35720096 | 39839728 | 38219400 | 39472392 | 4 | **1.176** |
| P26443 | Glud1 | Glutamate dehydrogenase 1, mitochondrial (Mus musculus) | 1550.01 | 56.09 | 61337 | 24 | 76582504 | 88675664 | 94727144 | 79147880 | 84112664 | 91316904 | 88496032 | 92329704 | 94097352 | 99289064 | 5 | **1.100** |
| P26516 | Psmd7 | 26S proteasome non-ATPase regulatory subunit 7 (Mus musculus) | 540.26 | 32.4 | 36540 | 8 | 20825272 | 17392656 | 16044315 | 16652721 | 15911290 | 18517568 | 18930984 | 15245002 | 14785754 | 13238426 | 8 | **0.930** |
| P26638 | Sars | Serine--tRNA ligase, cytoplasmic (Mus musculus) | 1630.54 | 58.01 | 58389 | 26 | 82977272 | 128654920 | 127862776 | 114832736 | 104074928 | 119505856 | 103042984 | 91536160 | 89485872 | 93307928 | 7 | **0.890** |
| P26645 | Marcks | Myristoylated alanine-rich C-kinase substrate (Mus musculus) | 92.51 | 11 | 29661 | 2 | 1585933.5 | 355274 | 1301347 | 364756.16 | 569662.38 | 1338218.25 | 1697442 | 1462974.38 | 1332596.88 | 1208902.75 | 5 | **1.685** |
| P26817 | Grk2 | Beta-adrenergic receptor kinase 1 (Rattus norvegicus) | 537.38 | 20.46 | 79785 | 11 | 8636781 | 11305770 | 12326856 | 10335655 | 12068338 | 11856403 | 11771955 | 11412409 | 12401517 | 12133582 | 7 | **1.090** |
| P26883 | Fkbp1a | Peptidyl-prolyl cis-trans isomerase FKBP1A (Mus musculus) | 122.8 | 12.96 | 11923 | 2 | 8464670 | 2598613.75 | 2480752.25 | 2791960 | 3579226.75 | 3157712 | 3769915.5 | 3348736 | 2997620.25 | 3033824 | 9 | **0.819** |
| P27008 | Parp1 | Poly [ADP-ribose] polymerase 1 (Rattus norvegicus) | 1103.1 | 26.82 | 112660 | 19 | 33632480 | 29519652 | 32013806 | 31745962 | 33501988 | 37560104 | 35456028 | 32426152 | 37514352 | 38853228 | 2 | **1.133** |
| P27046 | Man2a1 | Alpha-mannosidase 2 (Mus musculus) | 99.05 | 2 | 131631 | 2 | 881464.75 | 457981.31 | 978865.25 | 209304.59 | 620789.69 | 1431853.5 | 1120476.75 | 1568426.5 | 471953.81 | 1035170.5 | 3 | **1.788** |
| P27546 | Map4 | Microtubule-associated protein 4 (Mus musculus) | 649.18 | 17.69 | 117429 | 14 | 17500540 | 13946866 | 16345959 | 18098484 | 19475562 | 13078224 | 15925507 | 33160344 | 11960294 | 33582132 | 11 | **1.262** |
| P27601 | Gna13 | Guanine nucleotide-binding protein subunit alpha-13 (Mus musculus) | 149.72 | 10.08 | 44055 | 3 | 1563268.5 | 1844319.75 | 1828145 | 619531.19 | 1912326.5 | 1558978.5 | 2408800.75 | 2029595 | 981508.38 | 1662785.75 | 11 | **1.113** |
| P27612 | Plaa | Phospholipase A-2-activating protein (Mus musculus) | 682.58 | 22.42 | 87221 | 12 | 6440452.5 | 12589025 | 6642127.5 | 8203678.5 | 5373889 | 7676569.5 | 9357395 | 11646830 | 9390414 | 7704931.5 | 7 | **1.166** |
| P27641 | Xrcc5 | X-ray repair cross-complementing protein 5 (Mus musculus) | 137.58 | 4.92 | 83057 | 3 | 1495267.12 | 1460830.62 | 3162719.5 | 1322137.75 | 2367345.5 | 2405388 | 2453206.75 | 2620096.25 | 1810418.5 | 1738283.25 | 7 | **1.124** |
| P27659 | Rpl3 | 60S ribosomal protein L3 (Mus musculus) | 1290.3 | 45.66 | 46110 | 21 | 177553904 | 67905984 | 112421144 | 43960944 | 81630760 | 228427376 | 189000432 | 175959728 | 124459520 | 130318200 | 3 | **1.754** |
| P27661 | H2afx | Histone H2AX (Mus musculus) | 259.04 | 47.55 | 15143 | 4 | 1037882688 | 285787840 | 388033664 | 102809592 | 341053952 | 760674880 | 676448576 | 598636544 | 650314240 | 626507264 | 5 | **1.537** |
| P27773 | Pdia3 | Protein disulfide-isomerase A3 (Mus musculus) | 1985.09 | 60.79 | 56678 | 32 | 334439680 | 382407968 | 405523840 | 353795296 | 374145216 | 393464736 | 358379360 | 356650624 | 365179232 | 387993056 | 11 | **1.006** |
| P27808 | Mgat1 | Alpha-1,3-mannosyl-glycoprotein 2-beta-N-acetylglucosaminyltransferase (Mus musculus) | 64.75 | 3.58 | 51690 | 1 | 0 | 0 | 0 | 640127.44 | 0 | 522149.81 | 531925.19 | 610584.56 | 822159.88 | 714838.56 | 9 | **5.002** |
| P27867 | Sord | Sorbitol dehydrogenase (Rattus norvegicus) | 40.55 | 4.76 | 38235 | 1 | 0 | 0 | 0 | 0 | 0 | 0 | 602493.62 | 0 | 0 | 0 | 30 | **#DIV/0 !** |
| P27870 | Vav1 | Proto-oncogene vav (Mus musculus) | 1245.08 | 33.14 | 98137 | 21 | 31803336 | 42735344 | 46231476 | 45830920 | 37897936 | 39597516 | 37355928 | 46994304 | 45201436 | 47500692 | 9 | **1.059** |
| P28037 | Aldh1l1 | Cytosolic 10-formyltetrahydrofolate dehydrogenase (Rattus norvegicus) | 84.31 | 1.66 | 98874 | 2 | 2305100.75 | 2606109.5 | 2539724.25 | 1609125.12 | 2345551 | 1399214.88 | 784280.75 | 1674104.88 | 1949231 | 1668761.25 | 3 | **0.655** |
| P28063 | Psmb8 | Proteasome subunit beta type-8 (Mus musculus) | 634.03 | 39.49 | 30260 | 9 | 35551112 | 37489312 | 44590012 | 43317048 | 40026168 | 43414584 | 37027408 | 38279528 | 37513192 | 40359992 | 12 | **0.978** |
| P28271 | Aco1 | Cytoplasmic aconitate hydratase (Mus musculus) | 393.92 | 12.71 | 98126 | 7 | 2584584.5 | 2871273 | 3264935.5 | 4953886 | 4221996 | 2809597.75 | 2396498.75 | 3047620.5 | 3367921.25 | 1838683.25 | 6 | **0.752** |
| P28352 | Apex1 | DNA-(apurinic or apyrimidinic site) lyase (Mus musculus) | 335.54 | 23.03 | 35490 | 5 | 9285191 | 8006389 | 5551001.5 | 7360100.5 | 9660736 | 8651703 | 8165460 | 7415341.5 | 8831518 | 6191313 | 12 | **0.985** |
| P28474 | Adh5 | Alcohol dehydrogenase class-3 (Mus musculus) | 1035.8 | 56.42 | 39548 | 15 | 32228196 | 33263348 | 42677924 | 28192330 | 31312074 | 35563956 | 31482968 | 36613876 | 36213336 | 35405788 | 7 | **1.045** |
| P28650 | Adssl1 | Adenylosuccinate synthetase isozyme 1 (Mus musculus) | 332.35 | 15.1 | 50254 | 5 | 3789175.25 | 2498592.5 | 4206216.5 | 5149086 | 4224750 | 5893679.5 | 5148959 | 6038140.5 | 5684797 | 6299952.5 | 1 | **1.463** |
| P28656 | Nap1l1 | Nucleosome assembly protein 1-like 1 (Mus musculus) | 432.62 | 24.55 | 45345 | 7 | 19317178 | 22616568 | 25057832 | 34577040 | 28599304 | 28012756 | 25193150 | 27109300 | 26958836 | 24959110 | 11 | **1.016** |
| P28658 | Atxn10 | Ataxin-10 (Mus musculus) | 864.61 | 38.95 | 53707 | 14 | 24178402 | 27679250 | 32109072 | 27831612 | 20337186 | 25473606 | 27529914 | 25123952 | 26631178 | 28213928 | 12 | **1.006** |
| P28667 | Marcksl1 | MARCKS-related protein (Mus musculus) 1 | 262.4 | 51.5 | 20165 | 5 | 17676088 | 8013404 | 9455987 | 6750526.5 | 9057112 | 11644740 | 14134917 | 12196109 | 15154915 | 12704523 | 5 | **1.292** |
| P28740 | Kif2a | Kinesin-like protein KIF2A (Mus musculus) | 154.82 | 4.26 | 79756 | 3 | 1085798.25 | 2097604.75 | 2407000.5 | 2487980.75 | 2197979.25 | 2337709.5 | 1964698.5 | 2108502 | 2444249.25 | 2199151 | 12 | **1.076** |
| P28798 | Grn | Granulins (Mus musculus) | 492.09 | 17.49 | 63458 | 8 | 22164552 | 9248667 | 8024311 | 10079544 | 15196245 | 10105605 | 11482560 | 10921055 | 11376388 | 10409796 | 10 | **0.839** |
| P28867 | Prkcd | Protein kinase C delta type (Mus musculus) | 438.53 | 16.77 | 77547 | 9 | 15129857 | 14128751 | 14228078 | 14799178 | 10085208 | 14467931 | 11980612 | 14408092 | 13711943 | 15448046 | 12 | **1.024** |
| P29066 | Arrb1 | Beta-arrestin-1 (Rattus norvegicus) | 255.16 | 17.7 | 47020 | 6 | 13261310 | 13514175 | 14337694 | 11637640 | 12642640 | 14904214 | 8770240 | 10327352 | 11184610 | 12813849 | 7 | **0.887** |
| P29067 | Arrb2 | Beta-arrestin-2 (Rattus norvegicus) | 552.34 | 28.05 | 46340 | 10 | 24666414 | 24121054 | 23920696 | 22350918 | 21964262 | 25019284 | 16933284 | 18566328 | 19814110 | 19247566 | 5 | **0.851** |
| P29314 | Rps9 | 40S ribosomal protein S9 (Rattus norvegicus) | 579.81 | 38.14 | 22591 | 12 | 228979376 | 182722496 | 266521472 | 143469008 | 159696432 | 286916448 | 231264560 | 208268352 | 211674256 | 219986704 | 7 | **1.180** |
| P29341 | Pabpc1 | Polyadenylate-binding protein 1 (Mus musculus) | 1242.22 | 36.79 | 70671 | 21 | 140809664 | 115520936 | 111518272 | 123817000 | 121176048 | 140659968 | 115584504 | 116558632 | 112896208 | 110030792 | 9 | **0.972** |
| P29351 | Ptpn6 | Tyrosine-protein phosphatase non-receptor type 6 (Mus musculus) | 2114.3 | 67.06 | 67559 | 34 | 122960872 | 112083472 | 119935088 | 84941992 | 106584472 | 134939328 | 138383008 | 131659712 | 129615992 | 137961712 | 0 | **1.231** |
| P29391 | Ftl1 | Ferritin light chain 1 (Mus musculus) | 727.12 | 72.13 | 20802 | 11 | 94888904 | 76983872 | 90089464 | 62962596 | 90216984 | 77231472 | 104166424 | 92785720 | 97640512 | 90256688 | 5 | **1.113** |
| P29411 | Ak3 | GTP:AMP phosphotransferase AK3, mitochondrial (Rattus norvegicus) | 41.23 | 7.93 | 25438 | 1 | 345013.84 | 374987.06 | 101672.95 | 160111.23 | 0 | 420629.12 | 626991 | 366657.31 | 0 | 246936.09 | 8 | **1.692** |
| P29416 | Hexa | Beta-hexosaminidase subunit alpha (Mus musculus) | 834.52 | 36.93 | 60613 | 16 | 31121928 | 39981544 | 34205656 | 32129998 | 40060248 | 42781176 | 32754510 | 41431380 | 43588716 | 45389792 | 3 | **1.160** |
| P29418 | Atp5f1e | ATP synthase subunit epsilon, mitochondrial (Rattus norvegicus) | 43.37 | 15.69 | 5767 | 1 | 3374864.75 | 2991149 | 2933104.75 | 2777129.5 | 2643576.25 | 3653230.75 | 3398464.25 | 3583707 | 3156163 | 3447913 | 1 | **1.171** |
| P29452 | Casp1 | Caspase-1 (Mus musculus) | 467.99 | 21.64 | 45640 | 8 | 15225848 | 12600576 | 12978158 | 10549168 | 10128478 | 14829327 | 13808055 | 15347686 | 15135982 | 15534574 | 3 | **1.214** |
| P29595 | Nedd8 | NEDD8 (Mus musculus) | 136.88 | 34.57 | 8972 | 2 | 6383147 | 3903158 | 3244339.5 | 3171939.75 | 4589564 | 5810793 | 4417385 | 4527183.5 | 3708676.75 | 4122576.5 | 10 | **1.061** |
| P29758 | Oat | Ornithine aminotransferase, mitochondrial (Mus musculus) | 1085.39 | 59.23 | 48355 | 18 | 70752504 | 80133880 | 72871616 | 91934712 | 76143536 | 78954184 | 54339840 | 63933268 | 64924240 | 66741896 | 3 | **0.839** |
| P30204 | Msr1 | Macrophage scavenger receptor types I and II (Mus musculus) | 378.79 | 13.97 | 50170 | 6 | 18134764 | 19641024 | 22479556 | 18625132 | 22057796 | 17786734 | 19460908 | 21431948 | 15411966 | 20927166 | 8 | **0.941** |

| | | | | | | | | | | | | | | | | | |
|---|---|---|---|---|---|---|---|---|---|---|---|---|---|---|---|---|---|
| P30285 | Cdk4 | Cyclin-dependent kinase 4 (Mus musculus) | 169.19 | 12.54 | 33751 | 3 | 2919480 | 2224512.75 | 2485558.5 | 2137587.75 | 1229078.38 | 3064783 | 2978120 | 2760906.5 | 2258306 | 2324182 | 5 | **1.217** |
| P30355 | Alox5ap | Arachidonate 5-lipoxygenase-activating protein (Mus musculus) | 80.99 | 14.91 | 18136 | 2 | 5789801.5 | 8128315 | 8560608 | 7789619.5 | 5895297 | 7752725 | 8207238 | 10221972 | 8344263 | 8908511 | 5 | **1.201** |
| P30416 | Fkbp4 | Peptidyl-prolyl cis-trans isomerase FKBP4 (Mus musculus) | 1400.39 | 67.69 | 51572 | 24 | 119489184 | 118585592 | 133151496 | 118606192 | 116362952 | 126306712 | 110204224 | 117983952 | 116941384 | 118904168 | 9 | **0.974** |
| P30681 | Hmgb2 | High mobility group protein B2 (Mus musculus) | 510.18 | 29.05 | 24162 | 7 | 48220824 | 31680020 | 35195056 | 29798464 | 44610704 | 36256260 | 40439212 | 36363272 | 32620440 | 36090184 | 11 | **0.959** |
| P30993 | C5ar1 | C5a anaphylatoxin chemotactic receptor 1 (Mus musculus) | 108.98 | 6.27 | 39023 | 2 | 3566930.5 | 4103693.75 | 3866001.5 | 4359313 | 3918211 | 5081924.5 | 4751952 | 4281760.5 | 4566730.5 | 4557687 | 1 | **1.173** |
| P30999 | Ctnnd1 | Catenin delta-1 (Mus musculus) | 322.25 | 9.38 | 104925 | 7 | 7071932 | 5388926 | 4470927.5 | 7363121 | 6272860 | 18111664 | 7522262.5 | 8635880 | 6850968.5 | 5891443.5 | 5 | **1.538** |
| P31230 | Aimp1 | Aminoacyl tRNA synthase complex-interacting multifunctional protein 1 (Mus musculus) | 234.62 | 19.68 | 33997 | 4 | 5592174 | 5365329 | 3280816.25 | 4236441 | 7138327 | 6647929 | 5706743.5 | 4230244 | 5670466 | 4307540 | 10 | **1.037** |
| P31266 | Rbpj | Recombining binding protein suppressor of hairless (Mus musculus) | 416.87 | 19.01 | 58537 | 8 | 33263554 | 36306948 | 30352102 | 32918280 | 29600056 | 31827160 | 28944510 | 28900452 | 28174276 | 29329920 | 2 | **0.906** |
| P31750 | Akt1 | RAC-alpha serine/threonine-protein kinase (Mus musculus) | 179.75 | 7.92 | 55707 | 3 | 632120.38 | 767150.69 | 1103209.62 | 302557.44 | 294533.53 | 684659.38 | 803575.75 | 972107.94 | 658462.31 | 756456.69 | 8 | **1.250** |
| P31938 | Map2k1 | Dual specificity mitogen-activated protein kinase kinase 1 (Mus musculus) | 679.48 | 39.44 | 43474 | 12 | 51118904 | 35324532 | 41756720 | 35168952 | 29163284 | 44647312 | 38836780 | 46287316 | 45275716 | 46270444 | 6 | **1.150** |
| P31996 | Cd68 | Macrosialin-CD68 (Mus musculus) | 185.55 | 16.87 | 34818 | 4 | 136995072 | 107297440 | 112937000 | 103680072 | 94193328 | 116831480 | 121022064 | 101644960 | 104626184 | 108858112 | 11 | **0.996** |
| P32067 | Ssb | Lupus La protein homolog (Mus musculus) | 601.23 | 27.71 | 47756 | 10 | 27303742 | 19735020 | 20793444 | 21576688 | 24254452 | 33553444 | 23265068 | 26620446 | 25682784 | 25057440 | 5 | **1.180** |
| P32089 | Slc25a1 | Tricarboxylate transport protein, mitochondrial (Rattus norvegicus) | 211.26 | 13.18 | 33835 | 4 | 8254761 | 7525206 | 7193562 | 8262498.5 | 7787176 | 6776535 | 5652780 | 9005459 | 9774588 | 9155810 | 10 | **1.034** |
| P32198 | Cpt1a | Carnitine O-palmitoyltransferase 1, liver isoform (Rattus norvegicus) | 55.08 | 2.2 | 88125 | 1 | 0 | 0 | 37947.41 | 176119.61 | 157638.56 | 0 | 216077.69 | 122339.21 | 216999.75 | 127636.93 | 10 | **1.838** |
| P32233 | Drg1 | Developmentally-regulated GTP-binding protein 1 (Mus musculus) | 286.71 | 22.62 | 40512 | 6 | 7734277 | 8978432 | 8478290 | 13537839 | 7420836.5 | 8260912.5 | 9454215 | 8657632 | 8102438 | 5571350 | 11 | **0.868** |
| P32921 | Wars | Tryptophan--tRNA ligase, cytoplasmic (Mus musculus) | 556.78 | 34.93 | 54358 | 11 | 11451138 | 10498130 | 10772734 | 13004744 | 11253158 | 12208728 | 12621674 | 14522067 | 13087104 | 12036169 | 3 | **1.132** |
| P33174 | Kif4 | Chromosome-associated kinesin KIF4 (Mus musculus) | 159.79 | 2.44 | 139519 | 3 | 1180726.75 | 1316970.38 | 1475925 | 1750907.88 | 950823.69 | 1372259.62 | 1352325.88 | 1578004.25 | 1378928.25 | 990290.19 | 11 | **0.999** |
| P33610 | Prim2 | DNA primase large subunit (Mus musculus) | 241.28 | 10.1 | 58408 | 4 | 2711906.75 | 2907621.5 | 3000782 | 985862.62 | 2136931 | 4173638.5 | 4220267 | 4138193.75 | 2558000.75 | 3847729 | 3 | **1.613** |
| P33611 | Pola2 | DNA polymerase alpha subunit B (Mus musculus) | 130.76 | 8.67 | 66214 | 3 | 220414.69 | 376659.94 | 650155.62 | 373162.62 | 302220.75 | 289777.16 | 186272.36 | 245026.53 | 0 | 244224.22 | 3 | **0.502** |
| P34022 | Ranbp1 | Ran-specific GTPase-activating protein (Mus musculus) | 153.73 | 13.3 | 23596 | 3 | 10248635 | 8114277 | 6230199 | 9743123 | 8823060 | 11733255 | 9170987 | 7168289 | 7286774.5 | 6454204 | 11 | **0.969** |
| P34884 | Mif | Macrophage migration inhibitory factor (Mus musculus) | 230.07 | 55.65 | 12504 | 3 | 35564568 | 22031636 | 22115030 | 23820588 | 12153679 | 30571026 | 32185462 | 21471014 | 34032368 | 30774780 | 8 | **1.288** |
| P34902 | Il2rg | Cytokine receptor common subunit gamma (Mus musculus) | 41.08 | 2.71 | 42241 | 1 | 0 | 0 | 0 | 0 | 2919505.75 | 0 | 0 | 0 | 537625.5 | 406271.81 | 23 | **0.323** |
| P34960 | Mmp12 | Macrophage metalloelastase (Mus musculus) | 337.67 | 18.18 | 54971 | 8 | 4796958 | 6754042.5 | 7870068.5 | 6300580 | 5226438 | 9984937 | 6713616 | 8079629.5 | 7571843 | 7874636.5 | 3 | **1.300** |
| P35123 | Usp4 | Ubiquitin carboxyl-terminal hydrolase 4 (Mus musculus) | 446.72 | 13.31 | 108343 | 9 | 4866512.5 | 4794789 | 5224010 | 4818294 | 3877631.5 | 3213756 | 4297001.5 | 4755313.5 | 4868226 | 4225744 | 7 | **0.906** |
| P35276 | Rab3d | Ras-related protein Rab-3D (Mus musculus) | 360.31 | 31.51 | 24416 | 7 | 4529818 | 5530433 | 6501572 | 6352498 | 5999658.5 | 5507897 | 5331396 | 5269414 | 5602449.5 | 6958965 | 10 | **0.992** |
| P35278 | Rab5c | Ras-related protein Rab-5C (Mus musculus) | 613.7 | 71.76 | 23413 | 10 | 45990892 | 59634992 | 44562328 | 73551768 | 59126536 | 48445924 | 45115424 | 47584276 | 45082324 | 48183356 | 8 | **0.829** |
| P35279 | Rab6a | Ras-related protein Rab-6A (Mus musculus) | 351.43 | 26.92 | 23590 | 5 | 2213874.75 | 7241597 | 9406182 | 9942022 | 9642482 | 9637693 | 8236862 | 9293338 | 9338224 | 11187316 | 11 | **1.241** |
| P35280 | Rab8a | Ras-related protein Rab-8A (Rattus norvegicus) | 481.25 | 39.13 | 23668 | 9 | 0 | 1317332.38 | 855015 | 2743280.25 | 1506396.88 | 2316891 | 1277253.38 | 1257444.12 | 2468832.25 | 2008226.75 | 9 | **1.453** |
| P35282 | Rab21 | Ras-related protein Rab-21 (Mus musculus) | 268.5 | 30.18 | 24106 | 5 | 1832508.88 | 5792059 | 5231207 | 6491861 | 5733705.5 | 4333081 | 4783314.5 | 7093857 | 5903180.5 | 4746411 | 12 | **1.071** |
| P35286 | Rab13 | Ras-related protein Rab-13 (Rattus norvegicus) | 134.73 | 12.81 | 22901 | 3 | 4315624.5 | 2780952.25 | 3005489.75 | 2138173.75 | 2801024.75 | 3572940.25 | 4615157 | 3635103.75 | 2831173.5 | 4210859.5 | 5 | **1.254** |
| P35293 | Rab18 | Ras-related protein Rab-18 (Mus musculus) | 317.98 | 38.83 | 23035 | 6 | 7188657 | 11684764 | 10406542 | 10455340 | 8399639 | 9198858 | 7851246 | 9827557 | 10665351 | 10258200 | 11 | **0.993** |
| P35433 | Ppat | Amidophosphoribosyltransferase (Rattus norvegicus) | 41.05 | 1.35 | 57437 | 1 | 560332.25 | 1797292.62 | 2128774.5 | 1403959.38 | 1597780.5 | 0 | 0 | 1933294.5 | 2538519.25 | 2177446 | 11 | **0.888** |
| P35441 | Thbs1 | Thrombospondin-1 (Mus musculus) | 41.89 | 0.6 | 129647 | 1 | 9623173 | 12527290 | 7505677.5 | 14171931 | 13968960 | 5076641.5 | 6605606 | 0 | 5831837.5 | 0 | 1 | **0.303** |
| P35444 | Comp | Cartilage oligomeric matrix protein (Rattus norvegicus) | 137.22 | 4.5 | 82664 | 3 | 3974323.5 | 7422113 | 6610481 | 9422822 | 5879394.5 | 5212048 | 6037767.5 | 4911849.5 | 4964199 | 5083495.5 | 6 | **0.787** |
| P35559 | Ide | Insulin-degrading enzyme (Rattus norvegicus) | 396.64 | 9.91 | 117710 | 9 | 7369183 | 7490916 | 9375926 | 9680624 | 7353714.5 | 7617533.5 | 7660247.5 | 9182765 | 8890259 | 9830780 | 8 | **1.046** |
| P35564 | Canx | Calnexin (Mus musculus) | 1288.68 | 45.52 | 67278 | 22 | 77678680 | 86358784 | 79166520 | 88791504 | 96543488 | 72865120 | 74561656 | 62470120 | 70893888 | 63852496 | 0 | **0.804** |
| P35585 | Ap1m1 | AP-1 complex subunit mu-1 (Mus musculus) | 299.23 | 21.51 | 48543 | 7 | 7362663 | 5916001 | 5217146 | 4293039.5 | 4131353.5 | 10264432 | 6553768 | 8167726 | 7058384 | 6045153.5 | 3 | **1.415** |
| P35601 | Rfc1 | Replication factor C subunit 1 (Mus musculus) | 56.85 | 1.06 | 125985 | 1 | 319207.03 | 436636.66 | 409769.75 | 383541.66 | 319218.44 | 313999.5 | 111147.67 | 229361.92 | 211382.39 | 300486.72 | 0 | **0.624** |
| P35700 | Prdx1 | Peroxiredoxin-1 (Mus musculus) | 1476.28 | 88.44 | 22176 | 24 | 1199074176 | 1304044800 | 1360288128 | 1190462080 | 1323993216 | 1259546496 | 1362418560 | 1457817984 | 1435685888 | 1435288448 | 3 | **1.090** |
| P35738 | Bckdhb | 2-oxoisovalerate dehydrogenase subunit beta, mitochondrial (Rattus norvegicus) | 62.4 | 4.36 | 42823 | 1 | 0 | 349661.47 | 176620.23 | 342030.34 | 210720.73 | 0 | 0 | 89625.74 | 98813.91 | 37258.1 | 6 | **0.209** |
| P35821 | Ptpn1 | Tyrosine-protein phosphatase non-receptor type 1 (Mus musculus) | 319.23 | 16.67 | 49593 | 6 | 6173386.5 | 5347889 | 4585074 | 4436308 | 4585390 | 9322156 | 9237753 | 8835321 | 8582800 | 7997213 | 0 | **1.750** |
| P35922 | Fmr1 | Synaptic functional regulator FMR1 (Mus musculus) | 167.62 | 6.03 | 68989 | 3 | 12831766 | 9898607 | 9810141 | 10232626 | 9231560 | 9342007 | 10593791 | 10231866 | 10226051 | 11249252 | 10 | **0.993** |
| P35979 | Rpl12 | 60S ribosomal protein L12 (Mus musculus) | 609.89 | 63.64 | 17805 | 9 | 173810128 | 132304072 | 151462032 | 149069888 | 150558464 | 144582944 | 146355760 | 149692480 | 134242048 | 136800208 | 6 | **0.940** |
| P35980 | Rpl18 | 60S ribosomal protein L18 (Mus musculus) | 568.15 | 45.74 | 21645 | 10 | 227873856 | 58860572 | 117696416 | 56769324 | 99147224 | 275154048 | 171961248 | 213435232 | 186478432 | 198641488 | 4 | **1.866** |
| P35991 | Btk | Tyrosine-protein kinase BTK (Mus musculus) | 77.36 | 4.25 | 76437 | 2 | 193059.52 | 0 | 507260.97 | 549914.69 | 488665.22 | 801944.19 | 0 | 375301.59 | 735002.56 | 536727.94 | 9 | **1.408** |
| P36371 | Tap2 | Antigen peptide transporter 2 (Mus musculus) | 284.82 | 9.4 | 77445 | 5 | 2132111 | 2500809.75 | 2584775.5 | 1387628.88 | 1609249.62 | 2436308.5 | 2998408 | 3508412 | 3163531 | 3181686.5 | 2 | **1.497** |
| P36506 | Map2k2 | Dual specificity mitogen-activated protein kinase kinase 2 (Rattus norvegicus) | 352.74 | 22.75 | 44282 | 7 | 22238710 | 15961588 | 17654502 | 13863578 | 8171548.5 | 17761548 | 17463572 | 18443024 | 19046604 | 20721586 | 6 | **1.200** |
| P36536 | Sar1a | GTP-binding protein SAR1a (Mus musculus) | 302.29 | 31.31 | 22371 | 5 | 9183993 | 5352560.5 | 8663322 | 4623321.5 | 9015756 | 10864958 | 7853355 | 8573837 | 7570602.5 | 7743746.5 | 12 | **1.157** |
| P36552 | Cpox | Oxygen-dependent coproporphyrinogen-III oxidase, mitochondrial (Mus musculus) | 221.43 | 9.71 | 49715 | 4 | 2027712 | 3822680 | 3475124.5 | 3655774.75 | 2878798 | 1668691.88 | 1732985 | 2679493.5 | 2936214 | 3191744.5 | 5 | **0.770** |
| P36860 | Ralb | Ras-related protein Ral-B (Rattus norvegicus) | 145.14 | 15.53 | 23317 | 3 | 1622788.88 | 2668082.5 | 1653143.12 | 3632086.5 | 1954599.88 | 1497320.38 | 1512595.5 | 1445566.5 | 1585637.12 | 1327120 | 0 | **0.639** |
| P36876 | Ppp2r2a | Serine/threonine-protein phosphatase 2A 55 kDa regulatory subunit B alpha isoform (Rattus | 354.15 | 14.99 | 51678 | 6 | 7293400 | 15041357 | 15840211 | 12496525 | 10936088 | 13608892 | 12212341 | 13667293 | 14686993 | 14820902 | 11 | **1.120** |
| P36916 | Gnl1 | Guanine nucleotide-binding protein-like 1 (Mus musculus) | 312.03 | 12.52 | 68771 | 6 | 5378000 | 2281665 | 3699758 | 2451282.25 | 2894679 | 4247382 | 4898252.5 | 4586660.5 | 3762232.25 | 3741809.25 | 5 | **1.271** |
| P36993 | Ppm1b | Protein phosphatase 1B (Mus musculus) | 134.62 | 10.26 | 42795 | 3 | 2703121.75 | 2926719.5 | 3095647 | 2445665 | 2228373.25 | 3386224 | 3428101.5 | 2949653 | 3109145.25 | 3571932.25 | 1 | **1.227** |
| P37040 | Por | NADPH--cytochrome P450 reductase (Mus musculus) | 290.92 | 11.5 | 77044 | 5 | 10497676 | 10856993 | 10978945 | 11615057 | 7230780.5 | 9869389 | 10853528 | 8764830 | 9829127 | 10008266 | 6 | **0.964** |
| P37913 | Lig1 | DNA ligase 1 (Mus musculus) | 567.63 | 16.81 | 102290 | 12 | 14394593 | 11726776 | 13635073 | 9504897 | 10018383 | 15964004 | 15390490 | 16686196 | 14452246 | 15630995 | 0 | **1.318** |
| P38060 | Hmgcl | Hydroxymethylglutaryl-CoA lyase, mitochondrial (Mus musculus) | 544.72 | 42.46 | 34239 | 9 | 18698162 | 15575742 | 16567485 | 18471820 | 16958546 | 18783540 | 15248786 | 19341302 | 20030526 | 20610612 | 5 | **1.090** |
| P38647 | Hspa9 | Stress-70 protein, mitochondrial (Mus musculus) | 1768.49 | 44.33 | 73461 | 27 | 327212768 | 270179360 | 268121792 | 264085088 | 257844896 | 279472320 | 324960480 | 275672192 | 299124768 | 299090432 | 5 | **1.065** |
| P38718 | Mpc2 | Mitochondrial pyruvate carrier 2 (Rattus norvegicus) | 78.97 | 14.17 | 14258 | 2 | 1176104.12 | 3149599.5 | 4011664 | 5338119 | 3901484 | 3397278.25 | 3443327.75 | 3391135.5 | 5551226 | 4146779 | 10 | **1.134** |
| P39052 | Dnm2 | Dynamin-2 (Rattus norvegicus) | 778.7 | 21.49 | 98230 | 14 | 17643294 | 15568552 | 19079504 | 18986912 | 18212404 | 20751972 | 18105928 | 21177238 | 20447200 | 22200792 | 3 | **1.147** |
| P39749 | Fen1 | Flap endonuclease 1 (Mus musculus) | 575.37 | 26.46 | 42315 | 10 | 34750676 | 28747420 | 26833568 | 28797728 | 29406678 | 27534182 | 35828352 | 28613184 | 31233504 | 30586898 | 10 | **1.035** |
| P40124 | Cap1 | Adenylyl cyclase-associated protein 1 (Mus musculus) | 2390.11 | 72.15 | 51565 | 37 | 335397280 | 304102272 | 373593184 | 262246464 | 296346784 | 381014304 | 337150784 | 333769632 | 400158624 | 358562176 | 4 | **1.152** |
| P40142 | Tkt | Transketolase (Mus musculus) | 2152.29 | 62.12 | 67630 | 33 | 429328896 | 397428512 | 438889440 | 417566464 | 438042144 | 493665056 | 435067040 | 450342752 | 456969824 | 450808160 | 2 | **1.078** |
| P40240 | Cd9 | CD9 antigen (Mus musculus) | 74.12 | 11.06 | 25258 | 1 | 415955 | 0 | 552779.12 | 785008.5 | 1216860.75 | 1045157.19 | 1239543.25 | 1446330.75 | 1197740.25 | 675762.38 | 4 | **1.887** |
| P40336 | Vps26a | Vacuolar protein sorting-associated protein 26A (Mus musculus) | 451.16 | 39.45 | 38114 | 10 | 7953466.5 | 14515345 | 17517474 | 11306741 | 10209316 | 13713023 | 11828149 | 10938958 | 9845946 | 11235733 | 11 | **0.936** |
| P41105 | Rpl28 | 60S ribosomal protein L28 (Mus musculus) | 363.84 | 45.99 | 15733 | 8 | 75073712 | 56685020 | 80793944 | 51264476 | 73146128 | 103587728 | 103633664 | 110740744 | 121299488 | 105383216 | 0 | **1.616** |
| P41123 | Rpl13 | 60S ribosomal protein L13 (Rattus norvegicus) | 411.12 | 38.86 | 24309 | 8 | 171428640 | 85497224 | 123545904 | 67057160 | 132039312 | 189232496 | 133854296 | 156121024 | 157604528 | 164285440 | 4 | **1.382** |
| P41216 | Acsl1 | Long-chain-fatty-acid--CoA ligase 1 (Mus musculus) | 969.55 | 29.47 | 77951 | 17 | 27884500 | 27578900 | 32582066 | 31202822 | 28475402 | 33017736 | 29519922 | 31115076 | 28012310 | 34243840 | 7 | **1.055** |
| P41241 | Csk | Tyrosine-protein kinase CSK (Mus musculus) | 444.74 | 22.67 | 50716 | 9 | 13331900 | 17351114 | 17807220 | 12090852 | 11431334 | 12165940 | 12861956 | 13392681 | 13786856 | 14992502 | 12 | **0.933** |
| P41245 | Mmp9 | Matrix metalloproteinase-9 (Mus musculus) | 275.23 | 11.37 | 80535 | 6 | 3055487.25 | 4689053 | 5517637 | 4381816 | 4577701.5 | 3240077.5 | 2231929.5 | 4313779 | 4857580.5 | 4303701.5 | 7 | **0.853** |
| P41498 | Acp1 | Low molecular weight phosphotyrosine protein phosphatase (Rattus norvegicus) | 306.17 | 43.04 | 18152 | 5 | 23992742 | 14833806 | 13367166 | 14941412 | 13986900 | 18920846 | 17140490 | 15583193 | 18316800 | 18719016 | 5 | **1.093** |
| P41565 | Idh3g | Isocitrate dehydrogenase [NAD] subunit gamma 1, mitochondrial (Rattus norvegicus) | 226.17 | 12.98 | 42851 | 5 | 13299294 | 14295729 | 12443544 | 11886308 | 10939259 | 12447963 | 11662999 | 12689938 | 12063663 | 11719586 | 10 | **0.964** |
| P42125 | Eci1 | Enoyl-CoA delta isomerase 1, mitochondrial (Mus musculus) | 239.95 | 18.69 | 32250 | 3 | 4917455 | 4546290 | 3644504.5 | 5110930 | 4108446.75 | 3278440 | 3939527 | 2366267 | 2937554.5 | 2029824.75 | 1 | **0.652** |
| P42208 | Sept2 | Septin-2 (Mus musculus) | 528.32 | 28.81 | 41526 | 7 | 34149696 | 32773972 | 27215738 | 32077298 | 29446714 | 29705172 | 27365902 | 28994554 | 26852662 | 25268220 | 4 | **0.888** |
| P42225 | Stat1 | Signal transducer and activator of transcription 1 (Mus musculus) | 343.93 | 10.28 | 87197 | 6 | 14375734 | 12849240 | 16707891 | 15184298 | 16103348 | 9970681 | 11439567 | 14237404 | 13329545 | 13517070 | 3 | **0.831** |
| P42227 | Stat3 | Signal transducer and activator of transcription 3 (Mus musculus) | 523.78 | 21.04 | 88054 | 10 | 4857057 | 5909592.5 | 6851488.5 | 5195042 | 5455524 | 8408922 | 7185163 | 5860043 | 8697097 | 6947801.5 | 2 | **1.312** |
| P42567 | Eps15 | Epidermal growth factor receptor substrate 15 (Mus musculus) | 168.05 | 4.91 | 98471 | 4 | 2558024.25 | 2705232 | 3797290.75 | 3767888.75 | 3307030.75 | 3244642 | 3793753.5 | 3528011 | 3391361 | 3561937 | 10 | **1.086** |
| P42667 | Sec11a | Signal peptidase complex catalytic subunit SEC11A (Rattus norvegicus) | 50.45 | 6.7 | 20599 | 1 | 1404856.12 | 1326305.5 | 1059775.62 | 0 | 238688.48 | 1017182.06 | 1746505.25 | 1692877.38 | 1362544.62 | 1430862 | 4 | **1.799** |
| P42859 | Htt | Huntingtin (Mus musculus) | 157.85 | 1.28 | 344690 | 3 | 870606.75 | 310700.5 | 981652.44 | 172000.56 | 0 | 728281.31 | 833505.75 | 641994.94 | 773413.5 | 723956.44 | 10 | **1.585** |
| P42932 | Cct8 | T-complex protein 1 subunit theta (Mus musculus) | 2317.05 | 68.98 | 59555 | 36 | 269770656 | 243524864 | 254066624 | 259580048 | 241206496 | 271552704 | 234651840 | 245309568 | 244187008 | 250323936 | 11 | **0.983** |
| P43247 | Msh2 | DNA mismatch repair protein Msh2 (Mus musculus) | 647.62 | 22.25 | 104151 | 14 | 9538493 | 8595724 | 12152574 | 12602264 | 9845775 | 9777203 | 12447326 | 13896200 | 14311094 | 13526158 | 4 | **1.213** |
| P43274 | Hist1h1e | Histone H1.4 (Mus musculus) e | 612.34 | 36.07 | 21977 | 11 | 362550016 | 138104032 | 106472184 | 137115344 | 173959664 | 115149072 | 195029216 | 90009040 | 185102672 | 121629184 | 10 | **0.770** |
| P43275 | Hist1h1a | Histone H1.1 (Mus musculus) a | 343.19 | 31.92 | 21785 | 6 | 242913120 | 75311056 | 60694680 | 79287600 | 109859944 | 79312808 | 120564008 | 55104136 | 116035016 | 75083360 | 12 | **0.785** |
| P43276 | Hist1h1b | Histone H1.5 (Mus musculus) b | 352.2 | 29.6 | 22576 | 7 | 53705996 | 15091147 | 13530040 | 26334396 | 36164560 | 17413246 | 23873796 | 11983674 | 22895080 | 15703262 | 8 | **0.634** |
| P43277 | Hist1h1d | Histone H1.3 (Mus musculus) d | 645.85 | 35.75 | 22100 | 11 | 506499456 | 178984144 | 138029808 | 178520624 | 236371728 | 154905968 | 255524032 | 104680360 | 253839984 | 161631520 | 10 | **0.751** |
| P43346 | Dck | Deoxycytidine kinase (Mus musculus) | 106.53 | 8.85 | 30367 | 2 | 2879030 | 2729410 | 3015542.75 | 2523721.5 | 2826454 | 1963163.88 | 2179975.25 | 2625566 | 2473278.75 | 2844336 | 4 | **0.865** |
| P43883 | Plin2 | Perilipin-2 (Mus musculus) | 63.34 | 3.29 | 46646 | 1 | 1384305.5 | 881956.19 | 821113.12 | 0 | 0 | 1027648.81 | 1207031.25 | 0 | 0 | 0 | 17 | **0.724** |
| P45376 | Akr1b1 | Aldose reductase (Mus musculus) | 914.68 | 57.28 | 35732 | 17 | 200260416 | 211552176 | 208355568 | 230077408 | 188115200 | 185150976 | 162397440 | 175486128 | 181519920 | 169752496 | 0 | **0.842** |
| P45377 | Akr1b8 | Aldose reductase-related protein 2 (Mus musculus) | 1109.12 | 64.87 | 36121 | 18 | 108310480 | 104183856 | 120352712 | 106592312 | 108640512 | 124519248 | 116796752 | 131509056 | 130770264 | 141474400 | 1 | **1.177** |

| | | | | | | | | | | | | | | | | | |
|---|---|---|---|---|---|---|---|---|---|---|---|---|---|---|---|---|---|
| P45878 | Fkbp2 | Peptidyl-prolyl cis-trans isomerase FKBP2 (Mus musculus) | 110.07 | 19.29 | 15344 | 2 | 0 | 1324432.75 | 1161821 | 4001551 | 2108591 | 1179702.38 | 890577.62 | 749691.69 | 1086739.38 | 751832.88 | 6 | **0.542** |
| P45952 | Acadm | Medium-chain specific acyl-CoA dehydrogenase, mitochondrial (Mus musculus) | 182.98 | 10.93 | 46481 | 4 | 11569391 | 10128812 | 9682616 | 10659141 | 9518494 | 10073664 | 9216589 | 8794430 | 8898458 | 9192944 | 2 | **0.896** |
| P46061 | Rangap1 | Ran GTPase-activating protein 1 (Mus musculus) | 1322.06 | 43.8 | 63531 | 22 | 33144054 | 37737064 | 39888584 | 34846988 | 29888310 | 43291300 | 36276740 | 41780948 | 38457888 | 42836316 | 3 | **1.155** |
| P46062 | Sipa1 | Signal-induced proliferation-associated protein 1 (Mus musculus) | 69.94 | 1.45 | 112066 | 1 | 680652.75 | 972805.12 | 1238600.75 | 944643.5 | 972601.44 | 1076709 | 1252362.62 | 981859.06 | 864324.5 | 1245469.5 | 6 | **1.127** |
| P46413 | Gss | Glutathione synthetase (Rattus norvegicus) | 148.12 | 8.02 | 52345 | 3 | 2199341.5 | 1854750.75 | 2024639.5 | 1644986.5 | 2283196.5 | 2344125.5 | 2545010 | 2169482 | 2239048.5 | 2307636 | 3 | **1.160** |
| P46460 | Nsf | Vesicle-fusing ATPase (Mus musculus) | 1003.67 | 38.17 | 82613 | 20 | 37287516 | 38522364 | 42628572 | 37673332 | 39227268 | 41088100 | 35847784 | 42434716 | 44636792 | 43530212 | 7 | **1.062** |
| P46462 | Vcp | Transitional endoplasmic reticulum ATPase (Rattus norvegicus) | 1942.47 | 54.22 | 89349 | 32 | 122066880 | 97636904 | 85985064 | 89196376 | 96560840 | 107006712 | 104258016 | 87330424 | 101116480 | 81807112 | 12 | **0.980** |
| P46467 | Vps4b | Vacuolar protein sorting-associated protein 4B (Mus musculus) | 386.74 | 24.77 | 49419 | 8 | 7002616.5 | 8725945 | 6233409 | 10988353 | 10958634 | 10524971 | 5268784 | 8937301 | 8763798 | 8124841 | 11 | **0.948** |
| P46471 | Psmc2 | 26S proteasome regulatory subunit 7 (Mus musculus) | 956.74 | 42.03 | 48648 | 16 | 25161098 | 27779066 | 26059404 | 24639772 | 25546876 | 27489546 | 26269146 | 31963046 | 31624096 | 33968688 | 2 | **1.171** |
| P46664 | Adss | Adenylosuccinate synthetase isozyme 2 (Mus musculus) | 915.3 | 36.84 | 50021 | 14 | 41135340 | 40909848 | 43130752 | 47870920 | 44007004 | 43867708 | 41637664 | 42167736 | 41076720 | 44951584 | 12 | **0.985** |
| P46978 | Stt3a | Dolichyl-diphosphooligosaccharide--protein glycosyltransferase subunit STT3A (Mus musc | 531.06 | 18.44 | 80598 | 11 | 7206030.5 | 23439968 | 27081800 | 21611200 | 15670516 | 12382392 | 16461691 | 19737664 | 21469718 | 20196690 | 9 | **0.950** |
| P47197 | Akt2 | RAC-beta serine/threonine-protein kinase (Rattus norvegicus) | 99.59 | 3.33 | 55543 | 2 | 623734.31 | 705300.31 | 663919.44 | 806159.25 | 624174.94 | 575464.81 | 518814.47 | 567084.38 | 799016.88 | 606103.88 | 4 | **0.896** |
| P47199 | Cryz | Quinone oxidoreductase (Mus musculus) | 99.8 | 10.27 | 35269 | 2 | 1223611.12 | 1148962.12 | 1197404.12 | 1110846.38 | 1129542.38 | 1383507.88 | 1938565.12 | 2411163.25 | 2821552.5 | 2705563 | 0 | **1.938** |
| P47226 | Tes | Testin (Mus musculus) | 311.21 | 19.62 | 47983 | 7 | 15920796 | 3224125 | 7851813 | 5900517 | 6803646 | 12874794 | 16854698 | 13013414 | 13049506 | 14384290 | 4 | **1.768** |
| P47245 | Nrdc | Nardilysin (Rattus norvegicus) | 484.72 | 12.06 | 132976 | 11 | 16209932 | 7184836 | 9914651 | 9984664 | 9546030 | 6287344 | 6508002.5 | 5740381 | 8236811.5 | 6232689 | 1 | **0.625** |
| P47713 | Pla2g4a | Cytosolic phospholipase A2 (Mus musculus) | 897.13 | 30.08 | 85222 | 16 | 57781224 | 44431580 | 52024032 | 38148708 | 32747750 | 58228152 | 62296808 | 56840340 | 39667816 | 40064360 | 7 | **1.142** |
| P47738 | Aldh2 | Aldehyde dehydrogenase, mitochondrial (Mus musculus) | 1903.83 | 68.4 | 56538 | 30 | 216342160 | 227137056 | 196844288 | 204379648 | 194457360 | 219241472 | 204465840 | 209058784 | 210081776 | 226385120 | 8 | **1.029** |
| P47740 | Aldh3a2 | Fatty aldehyde dehydrogenase (Mus musculus) | 115.94 | 5.99 | 53971 | 2 | 520445.75 | 499299.19 | 321359.53 | 531501.69 | 470492.16 | 560222.31 | 797985.5 | 489890.44 | 547453 | 491266.53 | 6 | **1.232** |
| P47753 | Capza1 | F-actin-capping protein subunit alpha-1 (Mus musculus) | 577.85 | 52.8 | 32940 | 11 | 39613384 | 59324736 | 46225212 | 110606144 | 42583712 | 39783152 | 44120692 | 43784992 | 42656652 | 41216608 | 8 | **0.709** |
| P47754 | Capza2 | F-actin-capping protein subunit alpha-2 (Mus musculus) | 616.66 | 60.84 | 32967 | 11 | 39875656 | 61669316 | 44922536 | 111517888 | 47934744 | 44617644 | 41321936 | 43345668 | 44399192 | 46380508 | 6 | **0.719** |
| P47758 | Srprb | Signal recognition particle receptor subunit beta (Mus musculus) | 76.24 | 7.06 | 29579 | 1 | 1082529.5 | 944435.56 | 646461.19 | 1456378.5 | 592110.88 | 0 | 540801.56 | 682109.19 | 672899.31 | 877597 | 6 | **0.587** |
| P47802 | Mtx1 | Metaxin-1 (Mus musculus) | 141.28 | 14.2 | 35624 | 3 | 794645.69 | 1350218.25 | 1524077.25 | 1435659.25 | 1542706.25 | 561027.44 | 637877.62 | 1497233.25 | 2029424.88 | 1729663 | 12 | **0.971** |
| P47811 | Mapk14 | Mitogen-activated protein kinase 14 (Mus musculus) | 489.99 | 33.06 | 41287 | 8 | 17543418 | 19377108 | 19189076 | 18517402 | 17304688 | 16248858 | 18492740 | 19749518 | 19090958 | 18854784 | 12 | **1.005** |
| P47856 | Gfpt1 | Glutamine--fructose-6-phosphate aminotransferase [isomerizing] 1 (Mus musculus) | 237.68 | 7.46 | 78539 | 5 | 4759992 | 3705621.5 | 4094261.75 | 2318894 | 4654402 | 4064612.75 | 4853619.5 | 5452217 | 5199457.5 | 5097430.5 | 3 | **1.263** |
| P47857 | Pfkm | ATP-dependent 6-phosphofructokinase, muscle type (Mus musculus) | 320.78 | 9.36 | 85269 | 6 | 9349342 | 18114278 | 16873060 | 14200266 | 13941593 | 15206898 | 12785986 | 18253386 | 16379966 | 19562652 | 8 | **1.134** |
| P47911 | Rpl6 | 60S ribosomal protein L6 (Mus musculus) | 599.55 | 36.15 | 33510 | 11 | 101578112 | 57555052 | 78836336 | 50309180 | 79611112 | 104963904 | 85089312 | 78742408 | 85159616 | 81165704 | 6 | **1.183** |
| P47915 | Rpl29 | 60S ribosomal protein L29 (Mus musculus) | 150.12 | 16.88 | 17587 | 4 | 59171780 | 30871738 | 40866676 | 24416830 | 39839744 | 43092368 | 58443964 | 62100180 | 57710888 | 60503104 | 3 | **1.444** |
| P47955 | Rplp1 | 60S acidic ribosomal protein P1 (Mus musculus) | 311.18 | 79.82 | 11475 | 4 | 58736504 | 45607936 | 44820776 | 42355464 | 15672762 | 45686444 | 58306600 | 50183152 | 54970216 | 52398296 | 5 | **1.262** |
| P47962 | Rpl5 | 60S ribosomal protein L5 (Mus musculus) | 605.19 | 48.48 | 34401 | 10 | 111420768 | 97677856 | 97977288 | 99362184 | 82884192 | 99290832 | 97776488 | 99363200 | 93285688 | 95064744 | 11 | **0.991** |
| P47964 | Rpl36 | 60S ribosomal protein L36 (Mus musculus) | 87.03 | 18.1 | 12216 | 2 | 22056376 | 20064696 | 23002440 | 17040904 | 21473388 | 25654976 | 25622186 | 24047528 | 27286908 | 25125134 | 0 | **1.233** |
| P47968 | Rpia | Ribose-5-phosphate isomerase (Mus musculus) | 164.85 | 10.89 | 32451 | 4 | 2552192.25 | 3596075.5 | 3591613.5 | 3679499.5 | 2910767.5 | 1036262.75 | 1257973.62 | 1216294 | 1012087.88 | 1490179.5 | 0 | **0.368** |
| P48024 | Eif1 | Eukaryotic translation initiation factor 1 (Mus musculus) | 70.68 | 11.5 | 12747 | 1 | 0 | 197912.66 | 290735.72 | 0 | 47189.72 | 267515.66 | 379743.97 | 272618.03 | 0 | 184466.8 | 10 | **2.061** |
| P48025 | Syk | Tyrosine-protein kinase SYK (Mus musculus) | 38.77 | 1.43 | 71376 | 1 | 0 | 0 | 0 | 0 | 243431.53 | 74575.58 | 0 | 94927.79 | 67466.88 | 180936.92 | 15 | **1.717** |
| P48036 | Anxa5 | Annexin A5 (Mus musculus) | 2179.27 | 78.06 | 35752 | 33 | 186966448 | 302060832 | 262135312 | 338333344 | 257377168 | 307269216 | 273526528 | 271795232 | 272361952 | 271376352 | 9 | **1.037** |
| P48193 | Epb41 | Protein 4.1 (Mus musculus) | 91.88 | 3.73 | 95911 | 2 | 1322466.25 | 1506873.75 | 1761248 | 1171680.12 | 2082459.5 | 1416954.38 | 1487513.25 | 1550411.75 | 1015137.12 | 950827.31 | 7 | **0.818** |
| P48441 | Idua | Alpha-L-iduronidase (Mus musculus) | 39.23 | 1.58 | 71254 | 1 | 0 | 522693.56 | 572783.38 | 99768.87 | 440900.16 | 446804.75 | 0 | 197244.17 | 405369.06 | 476887.22 | 11 | **0.933** |
| P48678 | Lmna | Prelamin-A/C (Mus musculus) | 2502.06 | 56.69 | 74238 | 41 | 356644256 | 272778400 | 280669120 | 270486464 | 300195680 | 293701632 | 311305984 | 278115392 | 277137792 | 276130464 | 12 | **0.970** |
| P48722 | Hspa4l | Heat shock 70 kDa protein 4L (Mus musculus) | 229.89 | 6.44 | 94382 | 4 | 11495837 | 12815690 | 13208098 | 13010715 | 11893420 | 10418190 | 4304682.5 | 11232996 | 10210708 | 9732598 | 0 | **0.735** |
| P48729 | CSNK1A1 | Casein kinase I isoform alpha Homo sapiens OX=9606 | 94.36 | 6.53 | 38915 | 2 | 1540187.5 | 2253166.75 | 2843244.5 | 2289992.25 | 2789662.75 | 2480121 | 2093999.5 | 2176685 | 2263752.75 | 1992149.75 | 8 | **0.939** |
| P48758 | Cbr1 | Carbonyl reductase [NADPH] 1 (Mus musculus) | 65.8 | 6.86 | 30641 | 2 | 2803155.5 | 1508261.25 | 0 | 1297599.5 | 1523888.62 | 2091828.12 | 2160794.25 | 1718389.88 | 1936881.12 | 1733865.88 | 5 | **1.352** |
| P48771 | Cox7a2 | Cytochrome c oxidase subunit 7A2, mitochondrial (Mus musculus) | 62.93 | 12.05 | 9291 | 1 | 1819240.62 | 1409216.12 | 1434799.5 | 875781.69 | 1043038.88 | 0 | 1454881.62 | 1245722.75 | 1609827.62 | 1544142.12 | 11 | **0.889** |
| P48774 | Gstm5 | Glutathione S-transferase Mu 5 (Mus musculus) | 412.73 | 33.93 | 26635 | 7 | 36356824 | 24990564 | 24976540 | 20495530 | 15511298 | 28676848 | 33599708 | 23299032 | 38791996 | 38702132 | 5 | **1.333** |
| P48962 | Slc25a4 | ADP/ATP translocase 1 (Mus musculus) | 1110.48 | 61.07 | 32904 | 19 | 294963008 | 252604928 | 251443296 | 300774848 | 186277760 | 268004096 | 258523296 | 303924384 | 285785600 | 307415968 | 6 | **1.107** |
| P48999 | Alox5 | Arachidonate 5-lipoxygenase (Mus musculus) | 1004.08 | 31.75 | 77967 | 18 | 63671472 | 27720002 | 32377776 | 25431732 | 27402278 | 67485936 | 56977036 | 34634700 | 31334096 | 39624668 | 5 | **1.303** |
| P49138 | Mapkapk2 | MAP kinase-activated protein kinase 2 (Mus musculus) 2 | 140.29 | 6.99 | 44050 | 3 | 6210879.5 | 6280554 | 6447896 | 5180264 | 6400917 | 4911477 | 4731123 | 8517831 | 8695881 | 8439814 | 10 | **1.156** |
| P49186 | Mapk9 | Mitogen-activated protein kinase 9 (Rattus norvegicus) | 95.56 | 4.96 | 48017 | 2 | 9671131 | 9151841 | 8970676 | 7726482.5 | 8054215 | 7574157 | 8069501 | 9697737 | 8399406 | 8836435 | 11 | **0.977** |
| P49242 | Rps3a | 40S ribosomal protein S3a (Rattus norvegicus) | 1066.14 | 56.06 | 29945 | 18 | 300981920 | 192126576 | 244173120 | 174974240 | 193749744 | 264039184 | 239519024 | 266250336 | 276805056 | 275329120 | 6 | **1.195** |
| P49300 | Clec10a | C-type lectin domain family 10 member A (Mus musculus) | 612.63 | 32.57 | 34596 | 9 | 25421644 | 18339096 | 17772658 | 16279950 | 19817112 | 18658224 | 20626856 | 18616828 | 19293330 | 18870872 | 9 | **0.984** |
| P49442 | Inpp1 | Inositol polyphosphate 1-phosphatase (Mus musculus) | 149.14 | 8.08 | 43346 | 3 | 2615158 | 3345181 | 2645321.5 | 3919097 | 3646322.75 | 2964261.5 | 2687938.5 | 2666195.5 | 3166182 | 3624401 | 11 | **0.934** |
| P49615 | Cdk5 | Cyclin-dependent-like kinase 5 (Mus musculus) | 132.7 | 8.22 | 33288 | 3 | 680543.5 | 680670.56 | 835709.75 | 1203185.12 | 1481614 | 1856611.12 | 1480266.5 | 1535710.5 | 1760438 | 1704054.5 | 1 | **1.708** |
| P49710 | Hcls1 | Hematopoietic lineage cell-specific protein (Mus musculus) | 489.91 | 19.75 | 54240 | 8 | 40338440 | 24547326 | 23608064 | 26020656 | 28416484 | 26452492 | 27245880 | 24265832 | 25505842 | 25283944 | 11 | **0.901** |
| P49717 | Mcm4 | DNA replication licensing factor MCM4 (Mus musculus) | 706.52 | 18.79 | 96736 | 13 | 20974454 | 17893554 | 17033292 | 17233130 | 17101672 | 20475682 | 19432716 | 19524970 | 16811130 | 15674426 | 12 | **1.019** |
| P49718 | Mcm5 | DNA replication licensing factor MCM5 (Mus musculus) | 1277.33 | 37.93 | 82343 | 25 | 90731640 | 80506824 | 90768592 | 72746728 | 80483648 | 83241552 | 93960864 | 88003784 | 83126952 | 93477896 | 6 | **1.064** |
| P49793 | Nup98 | Nuclear pore complex protein Nup98-Nup96 (Rattus norvegicus) | 338.02 | 4.3 | 197283 | 6 | 3153646 | 3200333.5 | 3704253 | 3453852.5 | 3186220.5 | 3580218 | 3379403.75 | 3622938.25 | 3417459 | 4204915.5 | 6 | **1.090** |
| P49962 | Srp9 | Signal recognition particle 9 kDa protein (Mus musculus) | 65.99 | 12.79 | 10194 | 1 | 1496860.38 | 1141091.12 | 1237549.5 | 1066658.75 | 1209625.25 | 1267459.88 | 1334366.25 | 1129305.62 | 1247529 | 1618520 | 7 | **1.072** |
| P50171 | Hsd17b8 | Estradiol 17-beta-dehydrogenase 8 (Mus musculus) | 173.47 | 17.76 | 26588 | 4 | 5457797 | 3295419.5 | 4191708 | 844281.94 | 4491145 | 4693965.5 | 2915605 | 4597430 | 2906779 | 3708792.5 | 12 | **1.030** |
| P50247 | Ahcy | Adenosylhomocysteinase (Mus musculus) | 1776.5 | 69.44 | 47688 | 28 | 187698352 | 156558544 | 190122336 | 157942528 | 164455440 | 192367536 | 187203232 | 181704976 | 208020432 | 220958080 | 4 | **1.156** |
| P50396 | Gdi1 | Rab GDP dissociation inhibitor alpha (Mus musculus) | 590.09 | 31.32 | 50522 | 10 | 43331144 | 44430984 | 41801848 | 47513908 | 31852952 | 39405388 | 32219080 | 42791352 | 40600464 | 44230696 | 8 | **0.954** |
| P50399 | Gdi2 | Rab GDP dissociation inhibitor beta (Rattus norvegicus) | 2353.98 | 82.7 | 50537 | 35 | 136826144 | 179674208 | 158304080 | 195803920 | 156212768 | 156487696 | 136418608 | 153875808 | 154112944 | 156698336 | 6 | **0.916** |
| P50408 | Atp6v1f | V-type proton ATPase subunit F (Rattus norvegicus) | 132.17 | 29.41 | 13370 | 2 | 2240411 | 1976212.25 | 2033730.5 | 2275273.25 | 2665422.75 | 2075345 | 1379539.5 | 2148159.75 | 2605853.5 | 2453343.5 | 12 | **0.953** |
| P50428 | Arsa | Arylsulfatase A (Mus musculus) | 96.85 | 7.31 | 53748 | 2 | 3908339.5 | 3285120 | 996265.81 | 2500024.5 | 2599576 | 2587097.75 | 2915159 | 2584797.25 | 2725770 | 2737057.25 | 12 | **1.020** |
| P50429 | Arsb | Arylsulfatase B (Mus musculus) | 523.99 | 18.73 | 59647 | 10 | 62762132 | 73687840 | 54415776 | 51338816 | 68095728 | 44189600 | 68086744 | 48212920 | 56922892 | 55786940 | 7 | **0.880** |
| P50431 | Shmt1 | Serine hydroxymethyltransferase, cytosolic (Mus musculus) | 285.11 | 12.55 | 52601 | 5 | 6311569 | 8750483 | 6068127 | 6880388.5 | 8443801 | 8339127.5 | 8062292 | 7953023 | 8647549 | 9932381 | 7 | **1.178** |
| P50516 | Atp6v1a | V-type proton ATPase catalytic subunit A (Mus musculus) | 2251.45 | 66.45 | 68326 | 34 | 123593824 | 120118504 | 131405120 | 106950920 | 107660016 | 142176208 | 144122560 | 145873040 | 138922928 | 158581376 | 0 | **1.237** |
| P50518 | Atp6v1e1 | V-type proton ATPase subunit E 1 (Mus musculus) 1 | 222.25 | 19.47 | 26157 | 4 | 8998090 | 3051607.5 | 6215975 | 2546323.25 | 5736166.5 | 6903900 | 7648374.5 | 9083834 | 9555134 | 9302507 | 2 | **1.601** |
| P50543 | S100a11 | Protein S100-A11 (Mus musculus) | 243.1 | 23.47 | 11083 | 3 | 4234637.5 | 3541057.25 | 4128157.75 | 2777850.5 | 4485630 | 3878243 | 4923254 | 4730332.5 | 5376501.5 | 5471857 | 3 | **1.272** |
| P50544 | Acadvl | Very long-chain specific acyl-CoA dehydrogenase, mitochondrial (Mus musculus) | 285.49 | 11.74 | 70875 | 5 | 3866223.5 | 4938236 | 4640147 | 4087640 | 4118536 | 5531239.5 | 4014652.5 | 5164625.5 | 5886706 | 5325654.5 | 4 | **1.197** |
| P50580 | Pa2g4 | Proliferation-associated protein 2G4 (Mus musculus) | 977.81 | 53.3 | 43699 | 16 | 72237904 | 86088040 | 89955136 | 92289120 | 85512544 | 76259424 | 74063280 | 76613584 | 76948632 | 84855768 | 5 | **0.912** |
| P50637 | Tspo | Translocator protein (Mus musculus) | 185.75 | 41.42 | 18841 | 4 | 817732.81 | 2381274.5 | 2802880.75 | 3546712.75 | 3315689 | 2633248 | 3416458.5 | 1598578.38 | 2874686.5 | 2140097.75 | 11 | **0.984** |
| P50904 | Rasa1 | Ras GTPase-activating protein 1 (Rattus norvegicus) | 137.23 | 3.08 | 115440 | 3 | 1188856.88 | 1431815.5 | 1704927.25 | 1852181 | 318603.16 | 1732778.25 | 1357828.62 | 1690605.75 | 1601826.88 | 2003959.38 | 8 | **1.291** |
| P51125 | Cast | Calpastatin (Mus musculus) | 44.78 | 2.28 | 84922 | 1 | 894935.31 | 651097.94 | 765881.44 | 459139.91 | 833573.38 | 650336.88 | 673213.06 | 617775.31 | 1232116.62 | 933933.12 | 11 | **1.139** |
| P51146 | Rab4b | Ras-related protein Rab-4B (Rattus norvegicus) | 116.69 | 10.33 | 23629 | 2 | 1438135.12 | 1239416.25 | 1348244.25 | 0 | 0 | 1209507.5 | 1284266.75 | 1474086.12 | 1032037.5 | 1021362.5 | 12 | **1.496** |
| P51174 | Acadl | Long-chain specific acyl-CoA dehydrogenase, mitochondrial (Mus musculus) | 1014.47 | 35.58 | 47908 | 17 | 57214516 | 62122968 | 73089544 | 80890824 | 71087256 | 60573632 | 66252320 | 62074324 | 64231748 | 65959924 | 8 | **0.927** |
| P51175 | Ppox | Protoporphyrinogen oxidase (Mus musculus) | 45.95 | 1.68 | 50871 | 1 | 589906.31 | 0 | 0 | 0 | 0 | 555692.5 | 0 | 586693.12 | 582542.12 | 575978.19 | 15 | **3.900** |
| P51410 | Rpl9 | 60S ribosomal protein L9 (Mus musculus) | 895.61 | 68.75 | 21881 | 13 | 92734640 | 104911224 | 131240792 | 119724792 | 123934976 | 123362992 | 87052440 | 186112208 | 191293408 | 171733872 | 7 | **1.327** |
| P51432 | Plcb3 | 1-phosphatidylinositol 4,5-bisphosphate phosphodiesterase beta-3 (Mus musculus) | 157.64 | 3 | 139487 | 3 | 596573.56 | 1856523 | 1704946.12 | 1656080.5 | 1752592.5 | 2058011.75 | 1787691.25 | 1618788.88 | 1750454.25 | 1827594.88 | 8 | **1.195** |
| P51569 | Gla | Alpha-galactosidase A (Mus musculus) | 244.05 | 12.65 | 47643 | 4 | 35074740 | 6213487.5 | 5488720 | 22310334 | 21537518 | 7247148 | 6237335 | 16547576 | 17815600 | 18045816 | 10 | **0.727** |
| P51660 | Hsd17b4 | Peroxisomal multifunctional enzyme type 2 (Mus musculus) | 771.92 | 25.31 | 79482 | 13 | 27567706 | 21698626 | 15720178 | 14297225 | 15303890 | 21602100 | 17054290 | 15881905 | 16732988 | 16609819 | 10 | **0.929** |
| P51807 | Dynlt1 | Dynein light chain Tctex-type 1 (Mus musculus) | 70.29 | 15.93 | 12483 | 1 | 888594 | 671046 | 684163.06 | 821232.06 | 762337.5 | 902846.06 | 0 | 772534.5 | 643375.31 | 568396.69 | 8 | **0.754** |
| P51829 | Adcy7 | Adenylate cyclase type 7 (Mus musculus) | 68.86 | 1.55 | 122708 | 1 | 0 | 528602.44 | 543524.62 | 743732.75 | 1186811.38 | 447122.31 | 484338.38 | 411112.34 | 754794.69 | 510454.44 | 8 | **0.869** |
| P51859 | Hdgf | Hepatoma-derived growth factor (Mus musculus) | 498.79 | 49.79 | 26269 | 9 | 22487344 | 17452954 | 14623261 | 15211983 | 21552122 | 18638408 | 18083090 | 17332646 | 20719316 | 18052198 | 11 | **1.016** |
| P51863 | Atp6v0d1 | V-type proton ATPase subunit d 1 (Mus musculus) 1 | 554.93 | 37.89 | 40301 | 9 | 29988552 | 31465618 | 31707054 | 31204254 | 25180682 | 33653404 | 31910808 | 28891080 | 30298256 | 31959864 | 7 | **1.048** |
| P51870 | Cyp4f5 | Cytochrome P450 4F5 (Rattus norvegicus) | 46.72 | 3.8 | 60681 | 1 | 0 | 0 | 0 | 265183.34 | 247921.72 | 0 | 0 | 376716.97 | 371162.19 | 319071.44 | 13 | **2.079** |
| P51881 | Slc25a5 | ADP/ATP translocase 2 (Mus musculus) | 1429.35 | 72.82 | 32931 | 22 | 327527584 | 327796768 | 299790208 | 343976224 | 235484256 | 311383680 | 304464128 | 384223008 | 370283328 | 386995744 | 6 | **1.145** |
| P51912 | Slc1a5 | Neutral amino acid transporter B(0) (Mus musculus) | 322.9 | 12.12 | 58483 | 6 | 7707645 | 7925704 | 8275023.5 | 6870187.5 | 9555320 | 11217212 | 9603862 | 10337615 | 9190203 | 9548128 | 2 | **1.237** |
| P52287 | Gnb3 | Guanine nucleotide-binding protein G(I)/G(S)/G(T) subunit beta-3 (Rattus norvegicus) | 133.65 | 7.35 | 37180 | 3 | 20842434 | 21514070 | 31825480 | 14076119 | 21880664 | 29951864 | 31428488 | 26204872 | 21971208 | 25483416 | 5 | **1.226** |
| P52293 | Kpna2 | Importin subunit alpha-1 (Mus musculus) | 1243.72 | 41.02 | 57928 | 18 | 48701308 | 42390980 | 43375148 | 43097008 | 41734324 | 42467528 | 44820976 | 49991224 | 45361392 | 50948376 | 5 | **1.065** |
| P52332 | Jak1 | Tyrosine-protein kinase JAK1 (Mus musculus) | 43.03 | 2.25 | 133367 | 1 | 0 | 105495.39 | 125910.72 | 207947.38 | 234160.48 | 0 | 0 | 321686.41 | 350134.59 | 122873.03 | 13 | **1.180** |

| | | | | | | | | | | | | | | | | | |
|---|---|---|---|---|---|---|---|---|---|---|---|---|---|---|---|---|---|
| P52431 | Pold1 | DNA polymerase delta catalytic subunit (Mus musculus) | 343.61 | 7.78 | 123790 | 7 | 5277158 | 4455993.5 | 3514637.75 | 2381879 | 3734563.5 | 3422791 | 2822492.25 | 4401062 | 4540128 | 3998268.25 | 12 | **0.991** |
| P52432 | Polr1c | DNA-directed RNA polymerases I and III subunit RPAC1 (Mus musculus) | 56.63 | 3.18 | 39107 | 1 | 1737923 | 1720670.5 | 1508010.88 | 1691376.25 | 1652004.75 | 1877218.38 | 1749964.25 | 1700167.25 | 1747242.75 | 1517209.62 | 6 | **1.034** |
| P52479 | Usp10 | Ubiquitin carboxyl-terminal hydrolase 10 (Mus musculus) | 223.08 | 10.23 | 87022 | 6 | 4543738 | 7162058.5 | 6455174 | 7072285.5 | 5037876.5 | 7429257.5 | 4956853 | 7337452 | 6224699.5 | 6145386 | 10 | **1.060** |
| P52480 | Pkm | Pyruvate kinase PKM (Mus musculus) | 3903.86 | 74.39 | 57845 | 51 | 1934376576 | 1819444480 | 1819975808 | 2015923840 | 1857499136 | 2254509312 | 1982101888 | 1931985152 | 1949321600 | 1972399104 | 5 | **1.068** |
| P52483 | Ube2e3 | Ubiquitin-conjugating enzyme E2 E3 (Mus musculus) | 94.4 | 19.81 | 22913 | 2 | 269521.31 | 1991058.38 | 2166192.75 | 1921072.38 | 1836373.25 | 1583210.12 | 1640017.38 | 1456520.5 | 1269788.88 | 1528938.5 | 5 | **0.914** |
| P52633 | Stat6 | Signal transducer and transcription activator 6 (Mus musculus) | 224.55 | 10.87 | 93497 | 7 | 2313669.5 | 3437223.75 | 3780984 | 3158594.25 | 1665580.5 | 3933363 | 4036456.25 | 4407526 | 1856231.75 | 3595335 | 5 | **1.242** |
| P52825 | Cpt2 | Carnitine O-palmitoyltransferase 2, mitochondrial (Mus musculus) | 101.64 | 4.41 | 73981 | 2 | 130291.18 | 848366.81 | 1173433 | 634751.62 | 1336864 | 1018931.44 | 994909.19 | 1194302.12 | 1220890.5 | 1500083.5 | 6 | **1.438** |
| P52873 | Pc | Pyruvate carboxylase, mitochondrial (Rattus norvegicus) | 40.91 | 0.85 | 129777 | 1 | 0 | 3015962.75 | 2072846.62 | 0 | 1784065.12 | 0 | 0 | 2184504 | 911111.81 | 561386.62 | 13 | **0.532** |
| P53395 | Dbt | Lipoamide acyltransferase component of branched-chain alpha-keto acid dehydrogenase cor | 135.42 | 6.02 | 53247 | 2 | 1651408.75 | 1420769.62 | 2035327.38 | 1022555.81 | 1376893.75 | 2248312 | 2405552 | 1999527.75 | 2318225 | 2093684.5 | 1 | **1.474** |
| P53534 | Pygb | Glycogen phosphorylase, brain form (Rattus norvegicus) | 846.89 | 23.39 | 96174 | 15 | 19277264 | 22276080 | 21108562 | 20785528 | 21747234 | 21975428 | 20510864 | 20394376 | 22505410 | 22798764 | 9 | **1.028** |
| P53563 | Bcl2l1 | Bcl-2-like protein 1 (Rattus norvegicus) | 102.75 | 8.15 | 26158 | 2 | 3761144.5 | 3458364.25 | 3755021.5 | 3309615 | 3519322.5 | 3618658 | 4708578 | 4480489 | 4783961 | 4752746.5 | 2 | **1.255** |
| P53676 | Ap3m1 | AP-3 complex subunit mu-1 (Rattus norvegicus) | 219.21 | 17.46 | 46981 | 5 | 7388759 | 5563036 | 6030761.5 | 6207951.5 | 4713140.5 | 5965865 | 6331709.5 | 6861002 | 5969706 | 6899427 | 9 | **1.071** |
| P53702 | Hccs | Cytochrome c-type heme lyase (Mus musculus) | 40.72 | 4.78 | 30978 | 1 | 0 | 0 | 0 | 0 | 0 | 843657.06 | 1028062.94 | 863914.75 | 1106974.38 | 1009008.44 | 10 | **#DIV/0 !** |
| P53811 | Pitpnb | Phosphatidylinositol transfer protein beta isoform (Mus musculus) | 150.31 | 14.02 | 31487 | 4 | 10250852 | 8633376 | 9225876 | 8085220 | 3343226 | 11036751 | 9035820 | 9278283 | 9417622 | 9854462 | 5 | **1.230** |
| P53986 | Slc16a1 | Monocarboxylate transporter 1 (Mus musculus) | 239.18 | 13.79 | 53267 | 4 | 0 | 1358879 | 2377096.5 | 1348842 | 1143457.12 | 1260284.25 | 1255269 | 1478425.5 | 1383177.88 | 1478412 | 9 | **1.101** |
| P53995 | Anapc1 | Anaphase-promoting complex subunit 1 (Mus musculus) | 50.54 | 0.72 | 215994 | 1 | 0 | 397830.44 | 500168.41 | 363648.03 | 307073.78 | 390368.03 | 419596.41 | 469713.94 | 386921.91 | 399407.72 | 7 | **1.317** |
| P53996 | Cnbp | Cellular nucleic acid-binding protein (Mus musculus) | 404.66 | 45.51 | 19592 | 7 | 29215590 | 22461844 | 24143020 | 13612230 | 30117704 | 26864104 | 24628954 | 26418294 | 23680954 | 28128750 | 11 | **1.085** |
| P54001 | P4ha1 | Prolyl 4-hydroxylase subunit alpha-1 (Rattus norvegicus) | 463.4 | 18.73 | 60898 | 8 | 8633017 | 9917150 | 10224831 | 8430589 | 10225026 | 10528331 | 10617057 | 11567671 | 11891528 | 12961898 | 0 | **1.214** |
| P54071 | Idh2 | Isocitrate dehydrogenase [NADP], mitochondrial (Mus musculus) | 1128.32 | 46.9 | 50906 | 20 | 95877496 | 89171984 | 96971504 | 80154424 | 87490952 | 102565792 | 98649512 | 92310536 | 82440424 | 94304024 | 8 | **1.046** |
| P54116 | Stom | Erythrocyte band 7 integral membrane protein (Mus musculus) | 433.69 | 32.39 | 31375 | 7 | 31463906 | 32544132 | 32789124 | 33437360 | 30951756 | 30224964 | 28261794 | 36556124 | 38228464 | 39461296 | 10 | **1.072** |
| P54276 | Msh6 | DNA mismatch repair protein Msh6 (Mus musculus) | 79.36 | 0.88 | 151084 | 1 | 241108.66 | 248141.3 | 399385.19 | 0 | 370573.31 | 355728.41 | 324256.84 | 216958.81 | 0 | 0 | 10 | **0.712** |
| P54310 | Lipe | Hormone-sensitive lipase (Mus musculus) | 80.42 | 5.4 | 83348 | 2 | 339239.22 | 608861.06 | 433370.66 | 113507.59 | 113495.98 | 350322.94 | 619112.5 | 455582.31 | 163469.8 | 133483.28 | 9 | **1.071** |
| P54311 | Gnb1 | Guanine nucleotide-binding protein G(I)/G(S)/G(T) subunit beta-1 (Rattus norvegicus) | 783.17 | 47.35 | 37377 | 12 | 41000780 | 40645532 | 56381436 | 32301790 | 34523988 | 50590956 | 60532348 | 56560448 | 54373636 | 59714928 | 2 | **1.375** |
| P54313 | Gnb2 | Guanine nucleotide-binding protein G(I)/G(S)/G(T) subunit beta-2 (Rattus norvegicus) | 1051.01 | 70.29 | 37331 | 18 | 62765012 | 62692520 | 81031344 | 49397060 | 55579712 | 67562064 | 79779776 | 81924192 | 79735552 | 84389344 | 3 | **1.263** |
| P54645 | Prkaa1 | 5'-AMP-activated protein kinase catalytic subunit alpha-1 (Rattus norvegicus) | 245.67 | 9.48 | 63973 | 4 | 4234222 | 3387856.5 | 4110951.75 | 2294786.75 | 3017470.5 | 4237008.5 | 3714205.25 | 4043688.25 | 3656831 | 3744516.25 | 8 | **1.138** |
| P54726 | Rad23a | UV excision repair protein RAD23 homolog A (Mus musculus) | 166.66 | 10.74 | 39706 | 2 | 2449056.5 | 2558080.5 | 2429857.25 | 3201502.25 | 976675.88 | 1415902.5 | 2052679.5 | 665589 | 2012414 | 703443.69 | 3 | **0.590** |
| P54728 | Rad23b | UV excision repair protein RAD23 homolog B (Mus musculus) | 144.87 | 9.86 | 43513 | 3 | 2841036.75 | 3262418 | 2752214 | 3969022.5 | 2341452.5 | 1837409.75 | 2438585.5 | 1033679.69 | 2519630.25 | 1089020.75 | 2 | **0.588** |
| P54775 | Psmc4 | 26S proteasome regulatory subunit 6B (Mus musculus) | 847.34 | 52.39 | 47408 | 16 | 42078664 | 29401656 | 35966024 | 26872104 | 37744840 | 45029512 | 44639424 | 38543776 | 40404496 | 40318024 | 3 | **1.214** |
| P54822 | Adsl | Adenylosuccinate lyase (Mus musculus) | 691.83 | 40.29 | 54866 | 12 | 35490248 | 23895996 | 26040196 | 19742748 | 27015964 | 36429508 | 32995406 | 33012102 | 33325158 | 33321142 | 4 | **1.279** |
| P54823 | Ddx6 | Probable ATP-dependent RNA helicase DDX6 (Mus musculus) | 680.24 | 32.71 | 54192 | 11 | 20986300 | 24170308 | 22914412 | 26006326 | 21909856 | 23217472 | 22865882 | 25366534 | 23165318 | 25806480 | 9 | **1.038** |
| P54921 | Napa | Alpha-soluble NSF attachment protein (Rattus norvegicus) | 424.85 | 31.19 | 33193 | 7 | 11948720 | 10514654 | 9949963 | 10626314 | 11791412 | 11846855 | 11645363 | 13773662 | 13860144 | 13576710 | 3 | **1.180** |
| P54923 | Adprh | [Protein ADP-ribosylarginine] hydrolase (Mus musculus) | 98.05 | 6.91 | 40068 | 2 | 3426518.5 | 2290017.75 | 2532025.75 | 2775032 | 1280085.25 | 2696558.5 | 3025249.5 | 2592464.5 | 1014540.88 | 1140124.62 | 10 | **0.851** |
| P54987 | Acod1 | Cis-aconitate decarboxylase (Mus musculus) | 116.37 | 6.56 | 53758 | 2 | 576714.81 | 1331774.88 | 1554799 | 1005220.88 | 1291764.75 | 1663621.38 | 1515799.5 | 1538973.38 | 1385239.25 | 1301283.88 | 5 | **1.286** |
| P55194 | Sh3bp1 | SH3 domain-binding protein 1 (Mus musculus) | 123.07 | 5 | 74172 | 2 | 193854.72 | 332379.91 | 0 | 216621.17 | 1353592.62 | 0 | 69668.85 | 656521.12 | 1443078.75 | 1038254.62 | 11 | **1.530** |
| P55264 | Adk | Adenosine kinase (Mus musculus) | 258.42 | 16.07 | 40149 | 4 | 2110437.5 | 3175321.5 | 6878129 | 3043065 | 314539.38 | 3074840.5 | 3002948.5 | 5454088 | 3536416.75 | 2183131.5 | 10 | **1.111** |
| P55266 | Adar | Double-stranded RNA-specific adenosine deaminase (Rattus norvegicus) | 57.98 | 1.79 | 129911 | 1 | 0 | 0 | 0 | 182727.64 | 356368.53 | 327145.78 | 0 | 311652.91 | 375154.78 | 381590.38 | 10 | **2.589** |
| P55302 | Lrpap1 | Alpha-2-macroglobulin receptor-associated protein (Mus musculus) | 125.14 | 6.67 | 42215 | 2 | 4653143.5 | 3889179 | 2787174 | 2979247.5 | 4754609 | 3869944 | 3912393 | 3756568.5 | 3313280 | 2975638.5 | 10 | **0.935** |
| P55770 | Snu13 | NHP2-like protein 1 (Rattus norvegicus) | 297.49 | 47.66 | 14174 | 6 | 7882679 | 1799852.88 | 5333933.5 | 5459888.5 | 6285971 | 10954834 | 10088873 | 9575441 | 7288532 | 6998806.5 | 2 | **1.678** |
| P56135 | Atp5mf | ATP synthase subunit f, mitochondrial (Mus musculus) | 122.35 | 23.86 | 10344 | 3 | 9312084 | 7083682 | 7895358.5 | 6919994 | 6071396.5 | 7665558 | 7155366 | 6737506 | 5893402.5 | 6243729 | 8 | **0.904** |
| P56183 | Rrp1 | Ribosomal RNA processing protein 1 homolog A (Mus musculus) | 220.17 | 13.36 | 54777 | 4 | 1719924.88 | 1716124.88 | 2034712.5 | 295834.59 | 421134.38 | 2516586 | 2695452.25 | 2720812.75 | 3089063 | 2650115 | 0 | **2.210** |
| P56380 | Nudt2 | Bis(5'-nucleosyl)-tetraphosphatase [asymmetrical] (Mus musculus) | 183.5 | 27.89 | 16989 | 4 | 1460745.75 | 2140873 | 2745478.25 | 1652172.5 | 1057880.62 | 1426504.5 | 2810223.5 | 2981260.25 | 2547103.5 | 1100163.25 | 9 | **1.200** |
| P56383 | Atp5mc2 | ATP synthase F(0) complex subunit C2, mitochondrial (Mus musculus) | 57.58 | 21.23 | 15476 | 1 | 308962.47 | 244382.19 | 271012.88 | 129464.12 | 134610.95 | 0 | 236036.5 | 51000.69 | 83735.15 | 85013.41 | 2 | **0.419** |
| P56391 | Cox6b1 | Cytochrome c oxidase subunit 6B1 (Mus musculus) | 80.54 | 16.28 | 10071 | 1 | 2352645 | 1397014 | 1050468.5 | 1139132 | 1484622.88 | 978643.62 | 1394389.25 | 999858.88 | 1410784.5 | 1012858.75 | 5 | **0.781** |
| P56394 | Cox17 | Cytochrome c oxidase copper chaperone (Mus musculus) | 70.53 | 25.4 | 6784 | 1 | 1341764.38 | 850076.88 | 992468.38 | 859699.25 | 1210447.5 | 1425722 | 1194260 | 1194943.62 | 1259520.38 | 1123161.38 | 7 | **1.179** |
| P56399 | Usp5 | Ubiquitin carboxyl-terminal hydrolase 5 (Mus musculus) | 1409.92 | 39.51 | 95833 | 25 | 61450492 | 61930816 | 63732252 | 55557228 | 57560720 | 57564384 | 65725036 | 64314192 | 64561428 | 64964092 | 3 | **1.056** |
| P56477 | Irf5 | Interferon regulatory factor 5 (Mus musculus) | 242.17 | 11.27 | 56005 | 4 | 3400471 | 1935870.38 | 3226028 | 1466319.25 | 2091884.75 | 3706225 | 3188995.5 | 2968063.5 | 2849179 | 665881.19 | 11 | **1.104** |
| P56480 | Atp5f1b | ATP synthase subunit beta, mitochondrial (Mus musculus) | 2673.85 | 79.58 | 56300 | 38 | 404212128 | 406746432 | 499733536 | 423109856 | 465373824 | 504886208 | 502116032 | 528180928 | 517253664 | 542219200 | 0 | **1.180** |
| P56500 | Ucp2 | Mitochondrial uncoupling protein 2 (Rattus norvegicus) | 53.24 | 7.12 | 33377 | 1 | 170098.66 | 214761.5 | 335659.97 | 0 | 0 | 251660.98 | 282329.47 | 246926.34 | 0 | 255585.08 | 10 | **1.439** |
| P56542 | Dnase2 | Deoxyribonuclease-2-alpha (Mus musculus) | 68.61 | 5.38 | 38811 | 1 | 739968.5 | 0 | 596335.25 | 657888.5 | 1395876.5 | 931662.94 | 0 | 583196.81 | 824765.94 | 592827.38 | 11 | **0.865** |
| P56546 | Ctbp2 | C-terminal-binding protein 2 (Mus musculus) | 584.59 | 35.28 | 48957 | 12 | 27503994 | 23014648 | 24990928 | 22710224 | 19718972 | 27515054 | 24649298 | 26062472 | 23894394 | 28171490 | 5 | **1.105** |
| P56558 | Ogt | UDP-N-acetylglucosamine--peptide N-acetylglucosaminyltransferase 110 kDa subunit (Rat | 510.32 | 11.39 | 115606 | 9 | 7321050 | 3980479 | 6197141 | 2964884.5 | 3994011.25 | 5360632 | 6214121 | 6637884 | 5811452.5 | 5674827 | 8 | **1.214** |
| P56959 | Fus | RNA-binding protein FUS (Mus musculus) | 558.93 | 23.36 | 52673 | 9 | 45141112 | 31401000 | 35700340 | 31478206 | 36573320 | 39959884 | 32837478 | 34178944 | 32011168 | 35225720 | 12 | **0.966** |
| P57113 | Gstz1 | Maleylacetoacetate isomerase (Rattus norvegicus) | 39.23 | 6.02 | 23961 | 1 | 313471 | 0 | 0 | 560289.44 | 495650.19 | 0 | 0 | 507495.16 | 445576.22 | 417811.34 | 15 | **1.001** |
| P57716 | Ncstn | Nicastrin (Mus musculus) | 351.91 | 10.17 | 78492 | 7 | 28305288 | 13573066 | 14016807 | 13562004 | 16860382 | 28499644 | 30216056 | 27631942 | 29551486 | 27819056 | 2 | **1.665** |
| P57746 | Atp6v1d | V-type proton ATPase subunit D (Mus musculus) | 140.28 | 10.93 | 28369 | 2 | 2101392.75 | 1222122.62 | 1182464.88 | 1638904.5 | 1610059 | 1978383 | 1418378.88 | 1046051.31 | 1848949.5 | 841682.94 | 10 | **0.920** |
| P57759 | Erp29 | Endoplasmic reticulum resident protein 29 (Mus musculus) | 459.74 | 31.68 | 28823 | 8 | 9356362 | 20669274 | 18584198 | 24822476 | 18326178 | 15838570 | 16677240 | 16076460 | 17407272 | 16236881 | 5 | **0.896** |
| P57776 | Eef1d | Elongation factor 1-delta (Mus musculus) | 975.25 | 59.43 | 31293 | 15 | 173208736 | 168607760 | 195805168 | 136514496 | 177562128 | 172478192 | 132256128 | 164184352 | 165908576 | 163287504 | 5 | **0.937** |
| P57780 | Actn4 | Alpha-actinin-4 (Mus musculus) | 3138.57 | 65.13 | 104977 | 47 | 143928192 | 161818000 | 168974224 | 143433920 | 169940704 | 169755840 | 164793360 | 159886416 | 167735232 | 163659408 | 10 | **1.048** |
| P57784 | Snrpa1 | U2 small nuclear ribonucleoprotein A' (Mus musculus) | 157.89 | 20 | 28357 | 3 | 3001737.5 | 1063896.25 | 1074335.25 | 1139921.5 | 1398276.38 | 2004768.5 | 2488723 | 1884778.75 | 1250211 | 1860269.5 | 6 | **1.236** |
| P58021 | Tm9sf2 | Transmembrane 9 superfamily member 2 (Mus musculus) | 127.65 | 5.89 | 75330 | 2 | 1840302.75 | 2786191.25 | 2056500.75 | 1542429.25 | 1134436 | 2739870 | 2721631.5 | 3212441.5 | 2189240.75 | 3060590.75 | 3 | **1.488** |
| P58044 | Idi1 | Isopentenyl-diphosphate Delta-isomerase 1 (Mus musculus) | 291.06 | 32.6 | 26289 | 6 | 3740328 | 9713030 | 9281428 | 11047648 | 9363257 | 9820572 | 4083823.75 | 9136112 | 9101428 | 8766414 | 8 | **0.948** |
| P58058 | Nadk | NAD kinase (Mus musculus) | 58.63 | 3.87 | 48597 | 1 | 0 | 0 | 0 | 541379.81 | 995469.12 | 0 | 0 | 512554.22 | 0 | 651815.94 | 19 | **0.758** |
| P58137 | Acot8 | Acyl-coenzyme A thioesterase 8 (Mus musculus) | 38.54 | 5 | 35827 | 1 | 491139.47 | 0 | 0 | 532379.88 | 0 | 0 | 0 | 0 | 501789.47 | 394794.06 | 19 | **0.876** |
| P58195 | Plscr1 | Phospholipid scramblase 1 (Rattus norvegicus) | 43.4 | 5.97 | 36711 | 1 | 0 | 240396.27 | 114880.58 | 233089 | 433405.31 | 0 | 0 | 367155.56 | 293880.03 | 285313.88 | 13 | **0.926** |
| P58242 | Smpdl3b | Acid sphingomyelinase-like phosphodiesterase 3b (Mus musculus) | 272.97 | 16.67 | 51600 | 5 | 4667244 | 5335564 | 6174566.5 | 3402151.5 | 5134807.5 | 4073272 | 4736844 | 3883791.5 | 4416478.5 | 3748554.25 | 6 | **0.844** |
| P58252 | Eef2 | Elongation factor 2 (Mus musculus) | 3513.26 | 68.18 | 95314 | 55 | 600991424 | 658130496 | 691897088 | 621285312 | 630809280 | 799924032 | 726322496 | 726290816 | 726312320 | 705636800 | 0 | **1.150** |
| P58321 | Uchl4 | Ubiquitin carboxyl-terminal hydrolase isozyme L4 (Mus musculus) | 71.1 | 6.01 | 26450 | 1 | 0 | 491093.06 | 121831.16 | 749471.12 | 125553.37 | 0 | 0 | 103331.31 | 219273.39 | 178434.89 | 10 | **0.337** |
| P58389 | Ptpa | Serine/threonine-protein phosphatase 2A activator (Mus musculus) | 426.56 | 41.8 | 36710 | 8 | 3827226.75 | 11538749 | 12067936 | 9980822 | 9039668 | 10439694 | 11635052 | 12622349 | 11577291 | 12326347 | 4 | **1.261** |
| P58681 | Tlr7 | Toll-like receptor 7 (Mus musculus) | 525.49 | 10.86 | 121837 | 10 | 8897982 | 13519894 | 12911018 | 13875489 | 12175586 | 10504195 | 11339133 | 15185228 | 11799680 | 14710678 | 12 | **1.035** |
| P58742 | Aaas | Aladin (Mus musculus) | 111.71 | 4.03 | 59431 | 2 | 5016758 | 3992421 | 4011061 | 3318630.75 | 2724794.25 | 3060639 | 5059693 | 4518846 | 3808417.75 | 4361277 | 9 | **1.092** |
| P58774 | Tpm2 | Tropomyosin beta chain (Mus musculus) | 223.42 | 11.97 | 32837 | 5 | 103467384 | 80768384 | 80122184 | 64474120 | 62327388 | 73647616 | 84942016 | 71464680 | 85585200 | 73206872 | 11 | **0.994** |
| P58854 | Tubgcp3 | Gamma-tubulin complex component 3 (Mus musculus) | 42.72 | 0.77 | 103469 | 1 | 663269.69 | 0 | 0 | 200143.61 | 0 | 472450.06 | 505394.78 | 630954.75 | 446422.44 | 490270.69 | 8 | **2.948** |
| P59016 | Vps33b | Vacuolar protein sorting-associated protein 33B (Mus musculus) | 56.44 | 1.78 | 70526 | 1 | 104216.51 | 0 | 0 | 0 | 0 | 0 | 304335.5 | 78097.31 | 133017.45 | 34995.79 | 13 | **5.282** |
| P59017 | Bcl2l13 | Bcl-2-like protein 13 (Mus musculus) | 168.53 | 13.36 | 46719 | 3 | 1261159.75 | 2771217 | 2160222 | 3488853.25 | 1251142.62 | 5083090 | 1731737.62 | 2276488.5 | 2715399.75 | 2938299.5 | 8 | **1.349** |
| P59235 | Nup43 | Nucleoporin Nup43 (Mus musculus) | 43.83 | 3.68 | 41990 | 1 | 1468892 | 1204420.88 | 1211616 | 1428863.88 | 1312147 | 1349684 | 940964.38 | 1099242.12 | 1172237.38 | 1218241.75 | 5 | **0.872** |
| P59325 | Eif5 | Eukaryotic translation initiation factor 5 (Mus musculus) | 216.32 | 10.49 | 48968 | 4 | 9225646 | 8908591 | 5684259.5 | 12436730 | 6105998 | 6596240 | 7349370 | 6523022.5 | 8385446 | 6732428.5 | 10 | **0.840** |
| P59326 | Ythdf1 | YTH domain-containing family protein 1 (Mus musculus) | 77.97 | 2.33 | 60879 | 1 | 737986.44 | 2334407 | 4974634.5 | 2708708.25 | 1252844.88 | 1385603.75 | 2853733.75 | 278730.94 | 959835.19 | 1169759.12 | 8 | **0.554** |
| P59438 | Hps5 | Hermansky-Pudlak syndrome 5 protein homolog (Mus musculus) | 86.91 | 0.98 | 126337 | 1 | 0 | 495335.78 | 325642.16 | 437737.56 | 329273.5 | 0 | 0 | 364452.25 | 338106.47 | 308851.75 | 10 | **0.637** |
| P59999 | Arpc4 | Actin-related protein 2/3 complex subunit 4 (Mus musculus) | 620.23 | 75.6 | 19667 | 10 | 122111408 | 156472144 | 140645488 | 190976544 | 138710496 | 123081648 | 99155448 | 101584352 | 104115128 | 93652256 | 1 | **0.696** |
| P60060 | Sec61g | Protein transport protein Sec61 subunit gamma (Mus musculus) | 49.62 | 17.65 | 7741 | 1 | 699390.06 | 453386.5 | 0 | 345432.84 | 0 | 443763.78 | 451539.31 | 0 | 735884.62 | 0 | 15 | **1.089** |
| P60122 | Ruvbl1 | RuvB-like 1 (Mus musculus) | 815.84 | 47.59 | 50214 | 14 | 46779008 | 38495928 | 46818200 | 36746616 | 40955236 | 46438196 | 35706144 | 45659444 | 45487368 | 45424832 | 12 | **1.043** |
| P60229 | Eif3e | Eukaryotic translation initiation factor 3 subunit E (Mus musculus) | 1158.44 | 48.09 | 52221 | 20 | 58898800 | 62009772 | 67856560 | 55119684 | 54408684 | 70420104 | 71584424 | 69852352 | 68584512 | 72131216 | 0 | **1.182** |
| P60330 | Espl1 | Separin (Mus musculus) | 53.41 | 0.61 | 233035 | 1 | 0 | 212355.73 | 94681.43 | 1069204.5 | 860005.75 | 205814.34 | 173435.7 | 0 | 1636977.75 | 1778939.5 | 11 | **1.697** |
| P60335 | Pcbp1 | Poly(rC)-binding protein 1 (Mus musculus) | 1052.76 | 60.96 | 37498 | 14 | 215596608 | 186229456 | 188040240 | 183873552 | 184982016 | 200809808 | 179478048 | 193263536 | 189771232 | 190643888 | 9 | **0.995** |
| P60517 | Gabarap | Gamma-aminobutyric acid receptor-associated protein (Rattus norvegicus) | 117.55 | 20.51 | 13918 | 2 | 10824572 | 7183982.5 | 7314528 | 7099702 | 8649134 | 9546427 | 6962355 | 7745781.5 | 9590865 | 8569122 | 11 | **1.033** |
| P60521 | Gabarapl2 | Gamma-aminobutyric acid receptor-associated protein-like 2 (Mus musculus) l2 | 233.58 | 46.15 | 13667 | 5 | 15781775 | 10967620 | 11447432 | 11664954 | 13103038 | 14084139 | 10723446 | 11860068 | 13036043 | 12301042 | 12 | **0.985** |
| P60603 | Romo1 | Reactive oxygen species modulator 1 (Mus musculus) | 68.71 | 21.52 | 8183 | 1 | 1085766.12 | 1096754.25 | 1224121.12 | 1058383.62 | 1356971 | 1100306.38 | 1453951.25 | 1401712.88 | 1542871.62 | 2015828.25 | 2 | **1.291** |
| P60670 | Nploc4 | Nuclear protein localization protein 4 homolog (Mus musculus) | 324.29 | 13.82 | 68017 | 6 | 7438159 | 3412542.5 | 5509140.5 | 7690554 | 3603182 | 3663706.5 | 3844421.5 | 3939395.5 | 3433198.75 | 5030327 | 9 | **0.720** |

| | | | | | | | | | | | | | | | | | |
|---|---|---|---|---|---|---|---|---|---|---|---|---|---|---|---|---|---|
| P60710 | Actb | Actin, cytoplasmic 1 (Mus musculus) | 4134.671 | 91.47 | 41737 | 58 | 6286627840 | 6780435968 | 6642248192 | 6230931456 | 6671910400 | 6746650624 | 5772912640 | 6527685632 | 5861831168 | 6529829888 | 8 | **0.964** |
| P60762 | Morf4l1 | Mortality factor 4-like protein 1 (Mus musculus) | 41.89 | 4.97 | 41493 | 1 | 79565.21 | 525338.06 | 374339.03 | 204563.27 | 264985.88 | 694319.81 | 555217.25 | 0 | 429927.81 | 212174.28 | 9 | **1.306** |
| P60766 | Cdc42 | Cell division control protein 42 homolog (Mus musculus) | 278.02 | 26.7 | 21259 | 4 | 25764776 | 15298575 | 15956102 | 17584966 | 19533118 | 20266624 | 21072808 | 17067514 | 20196800 | 18347632 | 8 | **1.030** |
| P60843 | Eif4a1 | Eukaryotic initiation factor 4A-I (Mus musculus) | 2306.16 | 71.92 | 46154 | 33 | 441859424 | 420403840 | 459000864 | 436052512 | 440689184 | 498719680 | 485018848 | 472670848 | 475270624 | 475272576 | 0 | **1.095** |
| P60867 | Rps20 | 40S ribosomal protein S20 (Mus musculus) | 160.97 | 25.21 | 13373 | 3 | 53051292 | 38187008 | 50332244 | 36231288 | 49989560 | 51097360 | 30940106 | 48095616 | 49450704 | 49787660 | 10 | **1.007** |
| P60892 | Prps1 | Ribose-phosphate pyrophosphokinase 1 (Rattus norvegicus) | 753.99 | 46.54 | 34834 | 13 | 34090940 | 46209164 | 47179744 | 41831920 | 37988152 | 35728636 | 39836524 | 37276980 | 39758960 | 39488788 | 8 | **0.927** |
| P61021 | Rab5b | Ras-related protein Rab-5B (Mus musculus) | 231.37 | 21.4 | 23707 | 4 | 11358316 | 27471378 | 18472750 | 27254268 | 17166088 | 20418808 | 17644862 | 22723306 | 20247346 | 23186732 | 11 | **1.025** |
| P61027 | Rab10 | Ras-related protein Rab-10 (Mus musculus) | 452.67 | 43.5 | 22541 | 9 | 14393470 | 9581477 | 15355400 | 7725858 | 15018065 | 18835202 | 19114772 | 17025148 | 15457588 | 17774304 | 0 | **1.421** |
| P61028 | Rab8b | Ras-related protein Rab-8B (Mus musculus) | 646.44 | 49.28 | 23603 | 11 | 13176939 | 11049601 | 13994494 | 12992504 | 12936688 | 14158461 | 14862581 | 15294314 | 14070254 | 16447691 | 0 | **1.167** |
| P61078 | Ube2d3 | Ubiquitin-conjugating enzyme E2 D3 (Rattus norvegicus) | 126.41 | 28.57 | 16687 | 2 | 7478825 | 6064331 | 3133867.5 | 5098482 | 2484142.25 | 4699109 | 4526112 | 5112729 | 5455086 | 4471082 | 12 | **1.000** |
| P61082 | Ube2m | NEDD8-conjugating enzyme Ubc12 (Mus musculus) | 216 | 24.04 | 20900 | 4 | 13635080 | 11072371 | 10174262 | 11147468 | 14301934 | 12588866 | 12127214 | 11634567 | 11513978 | 9023251 | 12 | **0.943** |
| P61087 | Ube2k | Ubiquitin-conjugating enzyme E2 K (Mus musculus) | 421.2 | 53.5 | 22407 | 7 | 12666450 | 14290062 | 13655464 | 13882630 | 12918791 | 9376058 | 11453376 | 9987658 | 12171281 | 11731070 | 0 | **0.812** |
| P61089 | Ube2n | Ubiquitin-conjugating enzyme E2 N (Mus musculus) | 297.65 | 56.58 | 17138 | 6 | 19469872 | 11139816 | 8866094 | 11079006 | 12191527 | 11264992 | 11641206 | 10767022 | 13183063 | 10348604 | 12 | **0.912** |
| P61161 | Actr2 | Actin-related protein 2 (Mus musculus) | 849.73 | 41.62 | 44761 | 14 | 40547148 | 54874412 | 55663796 | 36690712 | 46681604 | 48545356 | 48896344 | 56595148 | 55789932 | 52396920 | 6 | **1.118** |
| P61164 | Actr1a | Alpha-centractin (Mus musculus) | 432.74 | 35.9 | 42614 | 9 | 16819324 | 17118420 | 17362944 | 13137759 | 14623905 | 17544130 | 18149930 | 16491862 | 16863070 | 14855169 | 8 | **1.061** |
| P61202 | Cops2 | COP9 signalosome complex subunit 2 (Mus musculus) | 71.72 | 3.84 | 51597 | 1 | 1407371.25 | 459414.19 | 878779.81 | 0 | 0 | 802057.88 | 1080176 | 559921.5 | 0 | 439171.34 | 14 | **1.049** |
| P61211 | Arl1 | ADP-ribosylation factor-like protein 1 (Mus musculus) | 340.77 | 40.33 | 20412 | 5 | 7643054.5 | 14219881 | 12248646 | 14362690 | 5680015 | 12644188 | 11356402 | 11994664 | 11203654 | 11587542 | 11 | **1.086** |
| P61222 | Abce1 | ATP-binding cassette sub-family E member 1 (Mus musculus) | 1140.1 | 42.9 | 67314 | 20 | 31509978 | 35673000 | 35584104 | 38097528 | 34945692 | 44934236 | 40108204 | 36035056 | 36346680 | 39193476 | 2 | **1.118** |
| P61226 | Rap2b | Ras-related protein Rap-2b (Mus musculus) | 213.27 | 38.8 | 20504 | 4 | 6977261.5 | 5748387.5 | 6537171.5 | 6356885.5 | 6051019 | 6678365 | 8084610.5 | 6238486 | 7002476 | 7094616.5 | 4 | **1.108** |
| P61255 | Rpl26 | 60S ribosomal protein L26 (Mus musculus) | 539.21 | 46.21 | 17258 | 10 | 121931128 | 55334548 | 95767352 | 35705252 | 78441152 | 148068928 | 129112016 | 122781304 | 114250808 | 124847072 | 1 | **1.651** |
| P61290 | Psme3 | Proteasome activator complex subunit 3 (Mus musculus) | 651.74 | 31.5 | 29506 | 10 | 27266926 | 24039592 | 25530164 | 17153520 | 15764322 | 24526118 | 26300478 | 31859512 | 30683296 | 31145508 | 3 | **1.317** |
| P61314 | Rpl15 | 60S ribosomal protein L15 (Rattus norvegicus) | 828.49 | 53.92 | 24146 | 16 | 126769208 | 92650016 | 162411872 | 61326404 | 104152648 | 181754352 | 166057856 | 178336944 | 149728624 | 176729904 | 1 | **1.558** |
| P61327 | Magoh | Protein mago nashi homolog (Mus musculus) | 233.06 | 51.37 | 17164 | 5 | 4147400.5 | 9379835 | 10199843 | 6970849 | 2078774.75 | 7809235 | 6654943.5 | 7995107 | 7668638.5 | 8174086 | 11 | **1.169** |
| P61354 | Rpl27 | 60S ribosomal protein L27 (Rattus norvegicus) | 207.71 | 36.03 | 15798 | 4 | 79432648 | 43244140 | 57365832 | 33791828 | 70703392 | 104157888 | 82697232 | 78775760 | 78306688 | 69544576 | 4 | **1.453** |
| P61406 | Smg6 | Telomerase-binding protein EST1A (Mus musculus) | 45.16 | 1.34 | 160496 | 1 | 118593.41 | 315689.03 | 257785.77 | 0 | 0 | 116565.34 | 183254.81 | 0 | 0 | 0 | 14 | **0.433** |
| P61514 | Rpl37a | 60S ribosomal protein L37a (Mus musculus) | 326.65 | 58.7 | 10275 | 6 | 12902440 | 23397550 | 39570580 | 9397125 | 13606862 | 11406545 | 10850126 | 12198218 | 13762724 | 12668385 | 7 | **0.616** |
| P61589 | Rhoa | Transforming protein RhoA (Rattus norvegicus) | 632.57 | 61.66 | 21782 | 10 | 44591496 | 74145304 | 61628120 | 84657704 | 59311156 | 53918984 | 53007380 | 45364856 | 45677816 | 41844592 | 4 | **0.739** |
| P61620 | Sec61a1 | Protein transport protein Sec61 subunit alpha isoform 1 (Mus musculus) | 183.61 | 12.39 | 52265 | 3 | 3871470.75 | 4916545.5 | 8284310 | 6510234.5 | 3722310.5 | 1806698.5 | 4304469 | 4204379 | 7165849 | 804145.06 | 8 | **0.670** |
| P61750 | Arf4 | ADP-ribosylation factor 4 (Mus musculus) | 519.33 | 47.22 | 20397 | 9 | 35919336 | 70099080 | 70755608 | 61720992 | 54972380 | 74015312 | 61708252 | 71270728 | 67084864 | 72602816 | 5 | **1.181** |
| P61759 | Vbp1 | Prefoldin subunit 3 (Mus musculus) | 260.42 | 25.51 | 22436 | 5 | 3709844.75 | 3519042.25 | 5539470 | 7843426.5 | 5773669 | 3571314 | 5002103 | 5089575 | 5773155 | 4944445 | 10 | **0.924** |
| P61924 | Copz1 | Coatomer subunit zeta-1 (Mus musculus) | 314.78 | 35.03 | 20198 | 6 | 8374864 | 7623195 | 10032201 | 9490974 | 8926332 | 9797798 | 11161667 | 13634442 | 11006759 | 12690767 | 1 | **1.311** |
| P61928 | Rpl37 | 60S ribosomal protein L37 (Rattus norvegicus) | 85.67 | 16.49 | 11078 | 2 | 11113326 | 3262924.5 | 4623327.5 | 2939877.5 | 4950344 | 5850713 | 6882582 | 5267544.5 | 6747688 | 7538957.5 | 5 | **1.201** |
| P61957 | Sumo2 | Small ubiquitin-related modifier 2 (Mus musculus) | 44.83 | 12.63 | 10871 | 1 | 128558136 | 4462890 | 7506877 | 4679599.5 | 8342970.5 | 38796896 | 9656708 | 8725380 | 8232302.5 | 8117533 | 7 | **0.479** |
| P61965 | Wdr5 | WD repeat-containing protein 5 (Mus musculus) | 102.77 | 8.98 | 36588 | 2 | 1686300.38 | 2626552.5 | 375116.62 | 2112039.5 | 2182381.5 | 2256868.5 | 2250756.5 | 1968201.75 | 2125242 | 2095910.38 | 10 | **1.191** |
| P61967 | Ap1s1 | AP-1 complex subunit sigma-1A (Mus musculus) | 87.89 | 10.13 | 18733 | 1 | 0 | 620103.25 | 395629.84 | 0 | 0 | 660211.19 | 684260.62 | 833795.44 | 721518.44 | 941984.56 | 3 | **3.782** |
| P61971 | Nutf2 | Nuclear transport factor 2 (Mus musculus) | 127.28 | 30.71 | 14478 | 2 | 572795.81 | 0 | 2757435 | 1214945.5 | 2587369 | 2045221.75 | 2259552.75 | 1605168 | 1993721.5 | 2334042.5 | 10 | **1.435** |
| P61979 | Hnrnpk | Heterogeneous nuclear ribonucleoprotein K (Mus musculus) | 1235.49 | 54.86 | 50976 | 19 | 197075728 | 170576896 | 151539088 | 166170576 | 179736000 | 173947344 | 127738024 | 184338080 | 162619280 | 168967888 | 10 | **0.945** |
| P61982 | Ywhag | 14-3-3 protein gamma (Mus musculus) | 1271.69 | 74.09 | 28303 | 21 | 223306480 | 413007680 | 561194624 | 428218912 | 323818016 | 294276032 | 271007104 | 274025696 | 347511552 | 353427072 | 7 | **0.790** |
| P62046 | Lrch1 | Leucine-rich repeat and calponin homology domain-containing protein 1 (Mus musculus) | 39.98 | 1.83 | 79076 | 1 | 0 | 542886.31 | 527006.19 | 536687.38 | 539800.25 | 0 | 0 | 499848 | 420677.47 | 449718.22 | 6 | **0.638** |
| P62082 | Rps7 | 40S ribosomal protein S7 (Mus musculus) | 384.1 | 40.21 | 22127 | 8 | 107740056 | 69892968 | 72858000 | 65829072 | 65043760 | 102221672 | 93646024 | 64465064 | 74364224 | 64244344 | 12 | **1.046** |
| P62137 | Ppp1ca | Serine/threonine-protein phosphatase PP1-alpha catalytic subunit (Mus musculus) | 829.23 | 46.97 | 37540 | 13 | 28337002 | 50250320 | 49092280 | 51637880 | 40256652 | 34421888 | 30794572 | 40844104 | 37817936 | 37001376 | 6 | **0.824** |
| P62138 | Ppp1ca | Serine/threonine-protein phosphatase PP1-alpha catalytic subunit (Rattus norvegicus) | 852.45 | 46.97 | 37512 | 13 | 31280584 | 52831416 | 52958520 | 53807544 | 43955832 | 38424432 | 35143976 | 43304040 | 40244636 | 39332804 | 5 | **0.837** |
| P62141 | Ppp1cb | Serine/threonine-protein phosphatase PP1-beta catalytic subunit (Mus musculus) | 835.59 | 41.59 | 37187 | 12 | 22492144 | 44917872 | 46909448 | 44108112 | 37828940 | 31888492 | 29694026 | 36951416 | 34245980 | 32306450 | 5 | **0.841** |
| P62192 | Psmc1 | 26S proteasome regulatory subunit 4 (Mus musculus) | 317.12 | 14.32 | 49185 | 5 | 3407894 | 5277445.5 | 2857703.25 | 8429317 | 5476441 | 3515387 | 3526133 | 4442504 | 6130947 | 5222117.5 | 12 | **0.897** |
| P62196 | Psmc5 | 26S proteasome regulatory subunit 8 (Mus musculus) | 625.23 | 33 | 45626 | 10 | 18421670 | 18902336 | 18192314 | 18424240 | 17694188 | 17972642 | 18351068 | 19878462 | 20628660 | 17678136 | 12 | **1.031** |
| P62242 | Rps8 | 40S ribosomal protein S8 (Mus musculus) | 779.29 | 55.29 | 24205 | 12 | 206453376 | 246492432 | 247513952 | 222724064 | 221581248 | 244575856 | 219989776 | 226417136 | 230026384 | 241895648 | 12 | **1.016** |
| P62245 | Rps15a | 40S ribosomal protein S15a (Mus musculus) | 468.46 | 48.46 | 14840 | 9 | 210792368 | 143387840 | 161626160 | 124912768 | 131953576 | 194240112 | 178508048 | 167662192 | 177402928 | 174826960 | 5 | **1.155** |
| P62254 | Ube2g1 | Ubiquitin-conjugating enzyme E2 G1 (Mus musculus) | 45.56 | 5.88 | 19509 | 1 | 691445.88 | 440232.81 | 505804.69 | 384727.09 | 483294.62 | 526307.25 | 479354.5 | 303782.34 | 528899 | 432920.97 | 11 | **0.907** |
| P62257 | Ube2h | Ubiquitin-conjugating enzyme E2 H (Mus musculus) | 47.94 | 8.2 | 20655 | 1 | 779751.94 | 1221833 | 1190315.38 | 0 | 995404.94 | 0 | 855913.56 | 1158545.5 | 1250839.75 | 0 | 13 | **0.780** |
| P62259 | Ywhae | 14-3-3 protein epsilon (Mus musculus) | 1828.28 | 85.88 | 29174 | 29 | 366488960 | 593255808 | 745424128 | 600643648 | 503812288 | 476216896 | 434584608 | 470609280 | 534675104 | 569086208 | 7 | **0.885** |
| P62267 | Rps23 | 40S ribosomal protein S23 (Mus musculus) | 185.49 | 32.87 | 15808 | 4 | 13642948 | 16765073 | 16003087 | 6728655 | 20155702 | 20753148 | 19387780 | 21904372 | 15653696 | 19837108 | 5 | **1.331** |
| P62270 | Rps18 | 40S ribosomal protein S18 (Mus musculus) | 577.44 | 63.16 | 17719 | 11 | 187163264 | 96340576 | 149741472 | 89378752 | 150802352 | 258723488 | 177854336 | 213596160 | 210569680 | 219841328 | 1 | **1.605** |
| P62274 | Rps29 | 40S ribosomal protein S29 (Mus musculus) | 90.72 | 32.14 | 6677 | 2 | 2440522.75 | 894117 | 1911949.5 | 1211972.5 | 1812306.12 | 3590463.25 | 3193292.5 | 3789460.5 | 3318915.5 | 3244327.25 | 0 | **2.072** |
| P62278 | Rps13 | 40S ribosomal protein S13 (Rattus norvegicus) | 547.34 | 49.01 | 17222 | 10 | 151355488 | 101129048 | 125911512 | 93379536 | 135813216 | 139114704 | 117341680 | 128473216 | 144512672 | 119903904 | 10 | **1.069** |
| P62281 | Rps11 | 40S ribosomal protein S11 (Mus musculus) | 438.46 | 50.63 | 18431 | 9 | 126808624 | 62411640 | 76966960 | 39773296 | 96685944 | 146796192 | 107006320 | 142986144 | 130696304 | 136590720 | 1 | **1.649** |
| P62305 | Snrpe | Small nuclear ribonucleoprotein E (Mus musculus) | 223.3 | 52.17 | 10804 | 4 | 20304018 | 1980248.5 | 28442004 | 18999440 | 23424250 | 26152002 | 1499531.62 | 25468378 | 25043976 | 23253196 | 10 | **1.089** |
| P62307 | Snrpf | Small nuclear ribonucleoprotein F (Mus musculus) | 139.72 | 39.53 | 9725 | 3 | 1202907 | 3841594.5 | 2451128 | 4978175 | 3383659 | 3742711 | 3353298.25 | 3747718.25 | 3817009.5 | 3802527.25 | 11 | **1.164** |
| P62315 | Snrpd1 | Small nuclear ribonucleoprotein Sm D1 (Mus musculus) | 72.67 | 20.17 | 13282 | 2 | 5592538.5 | 2807406.5 | 3763332.25 | 3209013.5 | 4028243 | 4095044.75 | 4696901 | 5438594 | 4018120 | 4657870 | 6 | **1.181** |
| P62317 | Snrpd2 | Small nuclear ribonucleoprotein Sm D2 (Mus musculus) | 219.61 | 52.54 | 13527 | 6 | 19334964 | 14274662 | 15175456 | 13665881 | 13195634 | 20293524 | 14065274 | 13866350 | 11130812 | 12795606 | 9 | **0.954** |
| P62320 | Snrpd3 | Small nuclear ribonucleoprotein Sm D3 (Mus musculus) | 222.82 | 37.3 | 13916 | 4 | 23640020 | 36689264 | 37760056 | 38422364 | 38081780 | 36078444 | 30241146 | 29253598 | 31098790 | 29774496 | 5 | **0.896** |
| P62322 | Lsm5 | U6 snRNA-associated Sm-like protein LSm5 (Mus musculus) | 84.35 | 20.88 | 9937 | 1 | 1593432.12 | 1248490 | 1331043.25 | 1348588.25 | 1583084.25 | 1098364.62 | 1394425.12 | 1067686.62 | 1130603.88 | 1198899.5 | 3 | **0.829** |
| P62329 | Tmsb4x | Thymosin beta-4 (Rattus norvegicus) | 96.79 | 54.55 | 5053 | 2 | 16998264 | 4315539.5 | 2335190.25 | 5046894.5 | 6569912.5 | 3141399.75 | 2843846 | 2606739 | 2780597.25 | 1986998.5 | 4 | **0.379** |
| P62331 | Arf6 | ADP-ribosylation factor 6 (Mus musculus) | 324.98 | 30.29 | 20082 | 5 | 3630183.5 | 8433076 | 8751130 | 8380612.5 | 6147184 | 8859244 | 6737897 | 7820023.5 | 7109375.5 | 7640629 | 12 | **1.080** |
| P62334 | Psmc6 | 26S proteasome regulatory subunit 10B (Mus musculus) | 541.38 | 26.48 | 44173 | 9 | 15664647 | 12366426 | 12818725 | 10502664 | 15533898 | 16603395 | 17135284 | 16917364 | 17501328 | 15265783 | 2 | **1.247** |
| P62482 | Kcnab2 | Voltage-gated potassium channel subunit beta-2 (Mus musculus) | 340.27 | 35.69 | 41021 | 7 | 8496692 | 11336599 | 12575081 | 12404320 | 12596424 | 9323069 | 9971737 | 10326833 | 10076978 | 10224986 | 5 | **0.870** |
| P62484 | Abi2 | Abl interactor 2 (Mus musculus) | 39.29 | 2.02 | 49387 | 1 | 2211017.25 | 2973919.75 | 3022161 | 2881898 | 2973286.5 | 2805596.25 | 3008468 | 2706766.5 | 3079485.75 | 2800854 | 12 | **1.024** |
| P62488 | Polr2g | DNA-directed RNA polymerase II subunit RPB7 (Mus musculus) | 117.61 | 13.95 | 19294 | 2 | 2281928.5 | 2523908.5 | 2362691.5 | 2318151.5 | 1938829.38 | 1984743 | 2029847.5 | 2178260.5 | 2129112.5 | 2182230.5 | 5 | **0.919** |
| P62634 | Cnbp | Cellular nucleic acid-binding protein (Rattus norvegicus) | 413.58 | 45.2 | 19463 | 7 | 30756444 | 23536432 | 25241560 | 14247597 | 31289282 | 27846886 | 25546296 | 27605362 | 24942338 | 29587270 | 11 | **1.084** |
| P62702 | Rps4x | 40S ribosomal protein S4, X isoform (Mus musculus) | 1123.59 | 57.41 | 29598 | 19 | 321324224 | 236730048 | 263240448 | 245674960 | 282069376 | 333190848 | 308168320 | 306538656 | 310972288 | 298665056 | 4 | **1.155** |
| P62715 | Ppp2cb | Serine/threonine-protein phosphatase 2A catalytic subunit beta isoform (Mus musculus) | 578.05 | 39.48 | 35575 | 10 | 27747714 | 34258512 | 34702800 | 41949752 | 28565568 | 30004386 | 31167152 | 34635608 | 36838472 | 37130872 | 10 | **1.015** |
| P62717 | Rpl18a | 60S ribosomal protein L18a (Mus musculus) | 427.4 | 44.89 | 20732 | 9 | 95099880 | 62016392 | 81118952 | 66292820 | 87760032 | 134338560 | 95750448 | 104621480 | 93815296 | 93176264 | 2 | **1.330** |
| P62748 | Hpcal1 | Hippocalcin-like protein 1 (Mus musculus) | 47.72 | 4.15 | 22338 | 1 | 2322021.5 | 631261.56 | 1354973.25 | 0 | 327669.5 | 1898365.5 | 1723907.25 | 1629955.5 | 1633805 | 1539107.75 | 5 | **1.817** |
| P62751 | Rpl23a | 60S ribosomal protein L23a (Mus musculus) | 395.31 | 41.03 | 17695 | 9 | 49045152 | 46638444 | 35049904 | 83678200 | 53910048 | 24310240 | 32218388 | 21121768 | 28703338 | 23479546 | 0 | **0.484** |
| P62754 | Rps6 | 40S ribosomal protein S6 (Mus musculus) | 507.33 | 27.71 | 28681 | 7 | 39950884 | 34261640 | 37748876 | 41700332 | 33450190 | 43419340 | 38635484 | 37746840 | 34795016 | 33717204 | 12 | **1.006** |
| P62774 | Mtpn | Myotrophin (Mus musculus) | 304.58 | 51.69 | 12861 | 6 | 15650788 | 10371032 | 11989412 | 14364844 | 13511479 | 17411326 | 15666806 | 15289413 | 16119808 | 15160860 | 2 | **1.209** |
| P62804 | Hist1h4b | Histone H4 (Rattus norvegicus) b | 1184.97 | 64.08 | 11367 | 16 | 3137517056 | 2408995072 | 2629014016 | 2107695744 | 2576225024 | 2668030464 | 2563976704 | 2626671104 | 2585066240 | 2592203264 | 10 | **1.014** |
| P62814 | Atp6v1b2 | V-type proton ATPase subunit B, brain isoform (Mus musculus) 2 | 1401.58 | 59.3 | 56551 | 21 | 71675416 | 73685808 | 70061752 | 72908104 | 69048608 | 69804928 | 71608712 | 67020204 | 66751208 | 67912088 | 3 | **0.960** |
| P62821 | Rab1A | Ras-related protein Rab-1A (Mus musculus) | 865.29 | 73.66 | 22678 | 13 | 27036852 | 57584120 | 72631440 | 60541544 | 26709756 | 43667596 | 32047196 | 53154744 | 52542692 | 43853208 | 10 | **0.921** |
| P62827 | Ran | GTP-binding nuclear protein Ran (Mus musculus) | 525.99 | 44.91 | 24423 | 10 | 118824968 | 104191680 | 98806816 | 112938640 | 104752368 | 127636584 | 104910088 | 95516080 | 101172400 | 99923568 | 10 | **0.981** |
| P62830 | Rpl23 | 60S ribosomal protein L23 (Mus musculus) | 385.19 | 52.14 | 14865 | 7 | 150858496 | 122548872 | 129114256 | 120829464 | 136965856 | 124572176 | 122199464 | 112847984 | 115761872 | 111353904 | 3 | **0.889** |
| P62835 | Rap1a | Ras-related protein Rap-1A (Mus musculus) | 814.56 | 68.48 | 20987 | 12 | 70004464 | 77174736 | 82061968 | 80480728 | 74380072 | 78377144 | 45542996 | 66315660 | 65562476 | 67513672 | 3 | **0.842** |
| P62843 | Rps15 | 40S ribosomal protein S15 (Mus musculus) | 449.67 | 37.24 | 17040 | 7 | 3598181.5 | 5838287 | 12363676 | 4164032.5 | 12516668 | 3348345.5 | 5959548 | 6539737.5 | 6250918 | 5312485 | 11 | **0.712** |
| P62849 | Rps24 | 40S ribosomal protein S24 (Mus musculus) | 382.47 | 40.6 | 15423 | 7 | 53417852 | 40017040 | 48641152 | 33226728 | 47324096 | 54043912 | 54686296 | 48778844 | 50962476 | 44036668 | 5 | **1.134** |
| P62852 | Rps25 | 40S ribosomal protein S25 (Mus musculus) | 306.65 | 29.6 | 13742 | 5 | 195341728 | 148162848 | 127009488 | 130989696 | 137836848 | 133498464 | 82119808 | 124347296 | 145295408 | 138748640 | 8 | **0.844** |
| P62855 | Rps26 | 40S ribosomal protein S26 (Mus musculus) | 152.66 | 20.87 | 13015 | 2 | 74593336 | 46956036 | 62245512 | 26808010 | 36491272 | 61364924 | 70178016 | 67943696 | 73313144 | 71341656 | 6 | **1.393** |
| P62858 | Rps28 | 40S ribosomal protein S28 (Mus musculus) | 215.52 | 46.38 | 7841 | 3 | 6857129.5 | 14112508 | 13916952 | 15537972 | 18433580 | 11042894 | 8723351 | 10561442 | 11474466 | 11185416 | 5 | **0.770** |
| P62862 | Fau | 40S ribosomal protein S30 (Mus musculus) | 85.91 | 18.64 | 6648 | 2 | 25502096 | 32231592 | 35570708 | 19117670 | 30294192 | 19908426 | 26885706 | 20276202 | 26766362 | 22200596 | 7 | **0.813** |
| P62869 | Elob | Elongin-B (Mus musculus) | 389.04 | 60.17 | 13170 | 7 | 28298118 | 18968932 | 18909668 | 17161032 | 20695780 | 15218190 | 21863058 | 19838426 | 20898664 | 17808646 | 12 | **0.919** |

| | | | | | | | | | | | | | | | | | | |
|---|---|---|---|---|---|---|---|---|---|---|---|---|---|---|---|---|---|---|
| P62878 | Rbx1 | E3 ubiquitin-protein ligase RBX1 (Mus musculus) | 189.7 | 25 | 12274 | 3 | 6242756 | 5953402.5 | 6579879 | 8095491.5 | 6093074 | 5755668.5 | 6160762 | 6007144 | 5863008 | 7093014 | 7 | **0.937** |
| P62889 | Rpl30 | 60S ribosomal protein L30 (Mus musculus) | 404.48 | 51.3 | 12784 | 5 | 58407132 | 40996760 | 40180156 | 37573120 | 36301408 | 56121040 | 51682232 | 46700224 | 49814576 | 47603172 | 5 | **1.180** |
| P62892 | Rpl39 | 60S ribosomal protein L39 (Mus musculus) | 48.98 | 19.61 | 6407 | 1 | 20021838 | 11994938 | 395574.88 | 10078160 | 23389008 | 23838960 | 393502.97 | 21805618 | 36700152 | 29023640 | 6 | **1.696** |
| P62897 | Cycs | Cytochrome c, somatic (Mus musculus) | 159.35 | 40.95 | 11605 | 4 | 35206976 | 35992556 | 35044396 | 37254512 | 47193508 | 28160890 | 27197172 | 29130132 | 33288050 | 33042912 | 0 | **0.791** |
| P62900 | Rpl31 | 60S ribosomal protein L31 (Mus musculus) | 94.84 | 16.8 | 14463 | 2 | 25741710 | 7417131 | 13461023 | 8084467 | 17349888 | 29347004 | 19683800 | 30651364 | 26755888 | 23623280 | 2 | **1.805** |
| P62907 | Rpl10a | 60S ribosomal protein L10a (Rattus norvegicus) | 465.9 | 35.94 | 24831 | 9 | 137034912 | 57971880 | 77435176 | 52720136 | 55103416 | 96855960 | 105750464 | 90133856 | 83882320 | 75984272 | 6 | **1.190** |
| P62908 | Rps3 | 40S ribosomal protein S3 (Mus musculus) | 1231.97 | 73.66 | 26674 | 23 | 370527104 | 360646592 | 379153824 | 374249664 | 370418912 | 338525056 | 356167200 | 377481760 | 365265984 | 369252512 | 6 | **0.974** |
| P62911 | Rpl32 | 60S ribosomal protein L32 (Mus musculus) | 414.35 | 40 | 15860 | 8 | 23640616 | 27058126 | 28924496 | 28352192 | 23418288 | 29841242 | 21803986 | 14650178 | 23725894 | 14015393 | 7 | **0.792** |
| P62914 | Rpl11 | 60S ribosomal protein L11 (Rattus norvegicus) | 543.92 | 52.81 | 20252 | 9 | 115509512 | 64642288 | 93078992 | 51795104 | 110164992 | 115695384 | 110738960 | 110835344 | 105866760 | 80098040 | 7 | **1.202** |
| P62915 | Gtf2b | Transcription initiation factor IIB (Mus musculus) | 37.16 | 3.48 | 34819 | 1 | 0 | 906680.62 | 1094165.25 | 803609.81 | 739065.31 | 0 | 0 | 0 | 0 | 0 | 15 | **0.000** |
| P62918 | Rpl8 | 60S ribosomal protein L8 (Mus musculus) | 463.6 | 33.07 | 28025 | 10 | 159883504 | 61436336 | 106301568 | 49602768 | 117352520 | 188220448 | 133291328 | 171669632 | 187292112 | 184569968 | 1 | **1.749** |
| P62944 | Ap2b1 | AP-2 complex subunit beta (Rattus norvegicus) | 1675.99 | 38.21 | 104553 | 28 | 36320704 | 45696172 | 51452096 | 46959536 | 44654064 | 49902248 | 47225652 | 58886612 | 54680268 | 55738036 | 2 | **1.184** |
| P62959 | Hint1 | Histidine triad nucleotide-binding protein 1 (Rattus norvegicus) | 51.44 | 11.11 | 13777 | 1 | 20134562 | 8821902 | 11212770 | 6667217.5 | 13397861 | 17717178 | 15403560 | 16450185 | 17132268 | 16640399 | 5 | **1.384** |
| P62960 | Ybx1 | Nuclease-sensitive element-binding protein 1 (Mus musculus) | 645.32 | 41.3 | 35730 | 8 | 17748946 | 15735166 | 26437748 | 13636236 | 27756752 | 22345462 | 21529538 | 26351492 | 23560532 | 26822344 | 9 | **1.190** |
| P62962 | Pfn1 | Profilin-1 (Mus musculus) | 1044.67 | 91.43 | 14957 | 16 | 461648320 | 603694976 | 500311840 | 744934592 | 514216448 | 500650048 | 406209216 | 478674816 | 440258048 | 309961312 | 3 | **0.756** |
| P62982 | Rps27a | Ubiquitin-40S ribosomal protein S27a (Rattus norvegicus) | 615.99 | 55.13 | 17951 | 10 | 255552064 | 202875280 | 195630528 | 211911984 | 230277104 | 185225632 | 208061136 | 191922448 | 211514032 | 213873776 | 7 | **0.922** |
| P62984 | Uba52 | Ubiquitin-60S ribosomal protein L40 (Mus musculus) | 499.94 | 50 | 14728 | 9 | 248713456 | 198430000 | 191271360 | 211520448 | 226200784 | 176355136 | 199284432 | 183993376 | 203649712 | 205948624 | 6 | **0.901** |
| P62996 | Tra2b | Transformer-2 protein homolog beta (Mus musculus) | 336.01 | 23.61 | 33666 | 5 | 10031407 | 10442159 | 9484616 | 12879595 | 9426679 | 7578763 | 7777982 | 7569084.5 | 7418938 | 7933221 | 0 | **0.732** |
| P63001 | Rac1 | Ras-related C3 botulinum toxin substrate 1 (Mus musculus) | 356.25 | 41.15 | 21450 | 6 | 43167488 | 20672080 | 25455688 | 26020262 | 26032926 | 29645602 | 31406554 | 27076400 | 26475480 | 27878200 | 5 | **1.008** |
| P63004 | Pafah1b1 | Platelet-activating factor acetylhydrolase IB subunit alpha (Rattus norvegicus) 1 | 607.97 | 36.1 | 46670 | 12 | 24098346 | 22178674 | 26668980 | 20817088 | 23883970 | 35999868 | 28456912 | 32016546 | 29843050 | 32858630 | 0 | **1.353** |
| P63017 | Hspa8 | Heat shock cognate 71 kDa protein (Mus musculus) | 3711.36 | 71.67 | 70871 | 52 | 1969635328 | 1860678528 | 1930356992 | 1737074432 | 1795090176 | 1868648064 | 1889526272 | 1929000960 | 2006865792 | 2062857088 | 6 | **1.050** |
| P63028 | Tpt1 | Translationally-controlled tumor protein (Mus musculus) | 517.69 | 73.26 | 19462 | 8 | 31055520 | 72188992 | 70188024 | 60365928 | 65882668 | 61772956 | 59654024 | 62717340 | 84370120 | 87417080 | 10 | **1.188** |
| P63030 | Mpc1 | Mitochondrial pyruvate carrier 1 (Mus musculus) | 63.07 | 19.27 | 12455 | 1 | 368092.56 | 530636.69 | 207650 | 603806.38 | 651704.44 | 0 | 86142.9 | 344872.53 | 535809.5 | 460889.25 | 6 | **0.604** |
| P63036 | Dnaja1 | DnaJ homolog subfamily A member 1 (Rattus norvegicus) | 334.41 | 29.22 | 44868 | 7 | 11185840 | 8052902 | 7992770 | 6414867.5 | 8268863 | 7298115.5 | 7092628.5 | 9739523 | 8731433 | 7945487 | 11 | **0.974** |
| P63038 | Hspd1 | 60 kDa heat shock protein, mitochondrial (Mus musculus) | 3435.89 | 76.79 | 60955 | 49 | 506479328 | 615583744 | 554901888 | 681385920 | 551899008 | 525978176 | 486942272 | 501036768 | 482019520 | 509978464 | 2 | **0.861** |
| P63073 | Eif4e | Eukaryotic translation initiation factor 4E (Mus musculus) | 170.81 | 15.21 | 25053 | 4 | 20002894 | 12878180 | 15600070 | 5864253 | 11976024 | 16474283 | 14899636 | 13879176 | 13645930 | 13729272 | 9 | **1.095** |
| P63081 | Atp6v0c | V-type proton ATPase 16 kDa proteolipid subunit (Rattus norvegicus) | 575.3 | 73.55 | 15808 | 8 | 6240467.5 | 9443288 | 9548256 | 7259751.5 | 7463246.5 | 6985806 | 13626207 | 9406089 | 10233377 | 9672998 | 6 | **1.250** |
| P63085 | Mapk1 | Mitogen-activated protein kinase 1 (Mus musculus) | 589.07 | 39.94 | 41276 | 11 | 21773208 | 13617778 | 17585540 | 22990250 | 17008000 | 24712926 | 22050808 | 17811790 | 16182200 | 22852254 | 8 | **1.114** |
| P63087 | Ppp1cc | Serine/threonine-protein phosphatase PP1-gamma catalytic subunit (Mus musculus) | 862.03 | 49.23 | 36984 | 13 | 25760326 | 48405104 | 50939840 | 47712928 | 41050212 | 34147496 | 32493964 | 39451596 | 36620792 | 35266272 | 5 | **0.832** |
| P63094 | Gnas | Guanine nucleotide-binding protein G(s) subunit alpha isoforms short (Mus musculus) | 288.62 | 24.87 | 45664 | 6 | 3175463 | 3552615.5 | 2777751.25 | 3261942.5 | 2474360 | 2974474.75 | 3518805.75 | 3573331 | 3056236.5 | 3481654 | 8 | **1.089** |
| P63101 | Ywhaz | 14-3-3 protein zeta/delta (Mus musculus) | 1394.18 | 82.86 | 27771 | 23 | 345415296 | 574782400 | 725068160 | 632237888 | 483233696 | 513360672 | 466576608 | 446122944 | 525710080 | 563107200 | 8 | **0.911** |
| P63158 | Hmgb1 | High mobility group protein B1 (Mus musculus) | 490.76 | 34.88 | 24894 | 8 | 50741992 | 36228768 | 36731052 | 38624784 | 49037476 | 41592452 | 44422376 | 39787300 | 48716148 | 46445104 | 10 | **1.045** |
| P63166 | Sumo1 | Small ubiquitin-related modifier 1 (Mus musculus) | 131.77 | 26.73 | 11557 | 3 | 11940676 | 8192686 | 7062874 | 6331483 | 7675814.5 | 9189547 | 9742381 | 8756117 | 8746128 | 9390164 | 5 | **1.112** |
| P63168 | Dynll1 | Dynein light chain 1, cytoplasmic (Mus musculus) | 152.4 | 37.08 | 10366 | 2 | 7553385.5 | 5618105 | 5166280.5 | 1056669.88 | 5055491.5 | 5702599 | 7937462 | 6171253 | 5354978 | 1628344.88 | 8 | **1.096** |
| P63174 | Rpl38 | 60S ribosomal protein L38 (Rattus norvegicus) | 63.08 | 17.14 | 8218 | 1 | 592478.31 | 0 | 679462.75 | 144588.17 | 893722.44 | 1378028.12 | 780213.06 | 955536.19 | 932789.62 | 706693.62 | 2 | **2.057** |
| P63213 | Gng2 | Guanine nucleotide-binding protein G(I)/G(S)/G(O) subunit gamma-2 (Mus musculus) | 55.69 | 16.9 | 7850 | 1 | 1947774.5 | 1237191.25 | 1354436.25 | 0 | 0 | 1105758.5 | 1176695 | 1146189.75 | 1075869 | 1246098 | 11 | **1.267** |
| P63242 | Eif5a | Eukaryotic translation initiation factor 5A-1 (Mus musculus) | 1080.15 | 88.31 | 16832 | 16 | 147629616 | 113328768 | 134630816 | 106566552 | 147868480 | 87418008 | 168589616 | 176032832 | 168043936 | 162538544 | 5 | **1.173** |
| P63245 | Rack1 | Receptor of activated protein C kinase 1 (Rattus norvegicus) | 1549.04 | 88.96 | 35077 | 23 | 311757312 | 376458400 | 399713632 | 377517984 | 358237088 | 388687936 | 331124352 | 379885376 | 356216352 | 370224352 | 12 | **1.001** |
| P63254 | Crip1 | Cysteine-rich protein 1 (Mus musculus) | 52.95 | 12.99 | 8550 | 1 | 1387865.12 | 921872.69 | 1108275.12 | 1231125.12 | 1530465.38 | 1225379.25 | 990709.06 | 1168178.88 | 1443314.88 | 1390165.88 | 12 | **1.006** |
| P63259 | Actg1 | Actin, cytoplasmic 2 (Rattus norvegicus) | 4115.571 | 91.47 | 41793 | 58 | 6278696960 | 6772372992 | 6636289536 | 6220765184 | 6659268608 | 6739843072 | 5767645696 | 6518506496 | 5854031360 | 6524017664 | 8 | **0.964** |
| P63276 | Rps17 | 40S ribosomal protein S17 (Mus musculus) | 617.33 | 60 | 15524 | 10 | 53709456 | 39998872 | 51002076 | 33950156 | 46372416 | 68977208 | 62889144 | 64148724 | 55409628 | 52745572 | 1 | **1.352** |
| P63280 | Ube2i | SUMO-conjugating enzyme UBC9 (Mus musculus) | 232.19 | 36.08 | 18007 | 4 | 8883179 | 5264130.5 | 7094593.5 | 3834108.75 | 6517859 | 7608760.5 | 7771314 | 7575358.5 | 6577395 | 7250221.5 | 6 | **1.164** |
| P63323 | Rps12 | 40S ribosomal protein S12 (Mus musculus) | 480.59 | 59.09 | 14525 | 7 | 46834248 | 39382612 | 37348132 | 39374796 | 47493756 | 42958580 | 35601496 | 31055230 | 41959056 | 41384192 | 9 | **0.917** |
| P63325 | Rps10 | 40S ribosomal protein S10 (Mus musculus) | 377.57 | 38.18 | 18916 | 6 | 9387493 | 10705245 | 15832968 | 12689162 | 11063507 | 14768840 | 6156654 | 12748768 | 9568273 | 11950700 | 12 | **0.925** |
| P63328 | Ppp3ca | Serine/threonine-protein phosphatase 2B catalytic subunit alpha isoform (Mus musculus) | 414.29 | 22.65 | 58644 | 8 | 13149105 | 12563917 | 14157003 | 13028932 | 12945610 | 13723948 | 14118115 | 13463516 | 17176052 | 16234172 | 3 | **1.135** |
| P67778 | Phb | Prohibitin (Mus musculus) | 1315.28 | 79.04 | 29820 | 21 | 175098816 | 198187120 | 186439312 | 186280512 | 177001744 | 172639168 | 183632384 | 186423408 | 192501264 | 198204080 | 11 | **1.011** |
| P67871 | Csnk2b | Casein kinase II subunit beta (Mus musculus) | 145.47 | 33.49 | 24942 | 4 | 7790321.5 | 5883408 | 5782546.5 | 5900441 | 4732770.5 | 4644221.5 | 5334377.5 | 5008448 | 5199544 | 5340488 | 4 | **0.848** |
| P67984 | Rpl22 | 60S ribosomal protein L22 (Mus musculus) | 361.87 | 40.62 | 14759 | 5 | 36282100 | 37211976 | 45418824 | 35698700 | 37412744 | 44792368 | 51265056 | 45413856 | 46996028 | 48804476 | 2 | **1.236** |
| P68033 | Actc1 | Actin, alpha cardiac muscle 1 (Mus musculus) | 1468.71 | 42.18 | 42019 | 25 | 4810661376 | 4936100352 | 4867966464 | 4606652928 | 4999840256 | 5029294080 | 4370959872 | 4802103808 | 4429135872 | 4890064896 | 9 | **0.971** |
| P68037 | Ube2l3 | Ubiquitin-conjugating enzyme E2 L3 (Mus musculus) | 340.16 | 57.79 | 17862 | 5 | 14630102 | 16349985 | 10217968 | 15659965 | 17237336 | 14070658 | 12649017 | 14171157 | 14026264 | 13533487 | 5 | **0.924** |
| P68101 | Eif2s1 | Eukaryotic translation initiation factor 2 subunit 1 (Rattus norvegicus) | 1096.02 | 59.05 | 36108 | 20 | 92657888 | 104288880 | 102048088 | 106827608 | 100397776 | 88260616 | 80902240 | 101476464 | 105289160 | 105293536 | 10 | **0.951** |
| P68181 | Prkacb | cAMP-dependent protein kinase catalytic subunit beta (Mus musculus) | 376.41 | 25.64 | 40708 | 7 | 20101616 | 21031964 | 18898692 | 19041636 | 14261365 | 23344456 | 13919063 | 21815530 | 20718948 | 21695516 | 6 | **1.087** |
| P68254 | Ywhaq | 14-3-3 protein theta (Mus musculus) | 1305.41 | 62.86 | 27778 | 21 | 221849248 | 421300224 | 544407424 | 426079392 | 325624544 | 307945248 | 266629712 | 287293952 | 352697184 | 368102112 | 7 | **0.816** |
| P68368 | Tuba4a | Tubulin alpha-4A chain (Mus musculus) | 1854.76 | 56.92 | 49924 | 28 | 442284960 | 636692800 | 559723712 | 674426368 | 579264576 | 530693376 | 497191392 | 509201408 | 530089440 | 538566336 | 5 | **0.901** |
| P68369 | Tuba1a | Tubulin alpha-1A chain (Mus musculus) | 2475.301 | 74.94 | 50136 | 37 | 557232128 | 849307328 | 750751040 | 815794624 | 749000448 | 657374336 | 604448000 | 673241472 | 714862336 | 743142656 | 5 | **0.912** |
| P68372 | Tubb4b | Tubulin beta-4B chain (Mus musculus) | 2345.67 | 76.85 | 49831 | 35 | 759401280 | 739857984 | 640619072 | 763690112 | 690282496 | 739434304 | 683520256 | 741315968 | 697387328 | 681297536 | 9 | **0.986** |
| P68373 | Tuba1c | Tubulin alpha-1C chain (Mus musculus) | 2451.07 | 70.6 | 49909 | 36 | 593824320 | 900713664 | 793039552 | 858660288 | 795973376 | 724901760 | 665308288 | 727965312 | 772390912 | 788349440 | 5 | **0.933** |
| P68433 | Hist1h3a | Histone H3.1 (Mus musculus) a | 388.9 | 50.74 | 15404 | 7 | 1451920768 | 908605888 | 1229817216 | 993050624 | 1196505856 | 1003660800 | 959959680 | 858912640 | 894889664 | 740863296 | 3 | **0.771** |
| P68510 | Ywhah | 14-3-3 protein eta (Mus musculus) | 1329.05 | 72.76 | 28212 | 22 | 227610576 | 410318400 | 527363968 | 410807584 | 314529248 | 294832160 | 262805440 | 288625728 | 354298528 | 366529888 | 7 | **0.829** |
| P69897 | Tubb5 | Tubulin beta-5 chain (Rattus norvegicus) | 2671.031 | 83.11 | 49671 | 38 | 998627904 | 955196160 | 844891328 | 987510272 | 909499648 | 961708672 | 899565632 | 969964864 | 932297600 | 860140864 | 10 | **0.985** |
| P70168 | Kpnb1 | Importin subunit beta-1 (Mus musculus) | 2134.08 | 51.94 | 97184 | 36 | 66267620 | 98286336 | 95953784 | 118263976 | 77951360 | 80241896 | 91301184 | 87484048 | 82658192 | 90397792 | 10 | **0.946** |
| P70188 | Kifap3 | Kinesin-associated protein 3 (Mus musculus) | 107.75 | 4.29 | 91291 | 2 | 66574.5 | 205496.14 | 262255.97 | 309584.94 | 259146.14 | 198078.48 | 185321.89 | 370700 | 276978.12 | 301020.56 | 10 | **1.208** |
| P70195 | Psmb7 | Proteasome subunit beta type-7 (Mus musculus) | 129.49 | 18.41 | 29891 | 3 | 10925470 | 2438127 | 5712621 | 6915441.5 | 6498209 | 9697720 | 8972314 | 7527803 | 8061980 | 8241195 | 5 | **1.308** |
| P70202 | Lxn | Latexin (Mus musculus) | 57.13 | 4.05 | 25492 | 1 | 3912160.75 | 3240432 | 3762417 | 3092352.75 | 2789479 | 4022636.5 | 3300440.75 | 3297236.75 | 3401710 | 3577396.75 | 8 | **1.048** |
| P70206 | Plxna1 | Plexin-A1 (Mus musculus) | 202.86 | 2.32 | 211099 | 4 | 3213746 | 3626339.75 | 3564156.75 | 2557019 | 1550397.25 | 3953580 | 3327248 | 1263613.75 | 1081119.88 | 1672130.12 | 9 | **0.779** |
| P70218 | Map4k1 | Mitogen-activated protein kinase kinase kinase kinase 1 (Mus musculus) | 103.93 | 3.39 | 91535 | 2 | 0 | 0 | 313624.81 | 0 | 339069.41 | 0 | 0 | 117746.78 | 372215.09 | 392639.03 | 15 | **1.352** |
| P70227 | Itpr3 | Inositol 1,4,5-trisphosphate receptor type 3 (Mus musculus) | 46.45 | 0.49 | 304275 | 1 | 80593.76 | 0 | 111997.92 | 0 | 0 | 228943.98 | 172049.75 | 184465.38 | 0 | 0 | 13 | **3.040** |
| P70248 | Myo1f | Unconventional myosin-If (Mus musculus) | 491.15 | 8.55 | 125947 | 8 | 8799587 | 9474237 | 9687511 | 9019736 | 9407401 | 11333380 | 11035171 | 12668711 | 11679069 | 11706360 | 0 | **1.259** |
| P70268 | Pkn1 | Serine/threonine-protein kinase N1 (Mus musculus) | 233.66 | 5.6 | 104411 | 5 | 8715615 | 4745030 | 5377662.5 | 4130582.25 | 4038770.75 | 6406633.5 | 5980470.5 | 4104147.75 | 3314050.5 | 4398014.5 | 11 | **0.896** |
| P70279 | Surf6 | Surfeit locus protein 6 (Mus musculus) | 63.19 | 2.82 | 41235 | 1 | 128530.63 | 795119.75 | 670185.81 | 809146.25 | 838082.88 | 534230.25 | 458613.44 | 425076.16 | 516127.91 | 395539.28 | 5 | **0.719** |
| P70288 | Hdac2 | Histone deacetylase 2 (Mus musculus) | 334.97 | 16.39 | 55302 | 5 | 8486180 | 12020512 | 10513314 | 10979874 | 9461836 | 9911301 | 9315955 | 11028257 | 10812637 | 10433662 | 12 | **1.001** |
| P70290 | Mpp1 | 55 kDa erythrocyte membrane protein (Mus musculus) | 172.77 | 7.94 | 52227 | 3 | 3283482.5 | 2140025.25 | 3145904.5 | 2486244.75 | 2565489.75 | 1764711.75 | 3212058 | 3222710.5 | 3202277.5 | 3243091.5 | 9 | **1.075** |
| P70296 | Pebp1 | Phosphatidylethanolamine-binding protein 1 (Mus musculus) | 255.86 | 36.36 | 20830 | 5 | 20848998 | 16355486 | 13407594 | 20014630 | 16209303 | 17427512 | 14432002 | 14547431 | 11443541 | 12619514 | 5 | **0.812** |
| P70303 | Ctps2 | CTP synthase 2 (Mus musculus) | 63.39 | 2.73 | 65514 | 1 | 386537.41 | 294631.88 | 377680.03 | 372957.41 | 349363.72 | 375385.09 | 350346.72 | 430006.19 | 536170.75 | 530059.12 | 5 | **1.247** |
| P70315 | Was | Wiskott-Aldrich syndrome protein homolog (Mus musculus) | 133.04 | 6.35 | 54192 | 3 | 2978818.5 | 3499156.5 | 1953142 | 2935446.75 | 2557603 | 3607737.5 | 3288342.5 | 3098200 | 3352713.25 | 3173808.5 | 4 | **1.186** |
| P70318 | Tial1 | Nucleolysin TIAR (Mus musculus) | 108.93 | 4.85 | 43389 | 2 | 3680462.5 | 4843167 | 3579978 | 5741653 | 4532267 | 2825753 | 3167625 | 3111571 | 3383272 | 3431324.5 | 0 | **0.711** |
| P70333 | Hnrnph2 | Heterogeneous nuclear ribonucleoprotein H2 (Mus musculus) | 473.17 | 26.95 | 49280 | 8 | 23544270 | 28579680 | 22716074 | 28250596 | 24312314 | 22411350 | 28581134 | 28378514 | 29944068 | 29731366 | 6 | **1.091** |
| P70388 | Rad50 | DNA repair protein RAD50 (Mus musculus) | 145.74 | 4.04 | 153488 | 3 | 333198.03 | 541809.75 | 367633.53 | 234292.31 | 470133.56 | 825844 | 521437.72 | 628440.19 | 555867 | 647313.19 | 1 | **1.633** |
| P70398 | Usp9x | Probable ubiquitin carboxyl-terminal hydrolase FAF-X (Mus musculus) | 199.17 | 2.23 | 290711 | 4 | 0 | 596683.69 | 943294.38 | 1459870.5 | 793781.69 | 1057634.75 | 286034.81 | 758558.06 | 933388.75 | 1442673.25 | 11 | **1.180** |
| P70444 | Bid | BH3-interacting domain death agonist (Mus musculus) | 46.65 | 9.74 | 21952 | 1 | 46303.12 | 550704.81 | 89329.87 | 750215.12 | 567861.81 | 96696.02 | 118941.9 | 364002.66 | 268506.34 | 310618.59 | 10 | **0.578** |
| P70460 | Vasp | Vasodilator-stimulated phosphoprotein (Mus musculus) | 282.66 | 25.07 | 39667 | 7 | 18055258 | 12027784 | 10640491 | 12105600 | 13460676 | 14958766 | 11843356 | 12660050 | 12544206 | 11715646 | 12 | **0.961** |
| P70470 | Lypla1 | Acyl-protein thioesterase 1 (Rattus norvegicus) | 144.2 | 16.09 | 24709 | 2 | 1619632.38 | 0 | 0 | 0 | 0 | 1259061.75 | 1445746.25 | 1004123.94 | 1135635.25 | 971357.25 | 11 | **3.591** |
| P70541 | Eif2b3 | Translation initiation factor eIF-2B subunit gamma (Rattus norvegicus) | 197.19 | 11.28 | 50435 | 4 | 2704882.5 | 4459946 | 4433400.5 | 4540666.5 | 4924377 | 2825532 | 3512707 | 5211114 | 5645075 | 5887603 | 8 | **1.096** |
| P70566 | Tmod2 | Tropomodulin-2 (Rattus norvegicus) | 51.56 | 1.99 | 39492 | 1 | 7528254 | 22142466 | 14463138 | 23826268 | 20168076 | 1391951.75 | 15429553 | 15283340 | 15916996 | 13661744 | 7 | **0.700** |
| P70583 | Dut | Deoxyuridine 5'-triphosphate nucleotidohydrolase (Rattus norvegicus) | 137.71 | 16.59 | 22003 | 3 | 7735037.5 | 11455496 | 10676431 | 12307155 | 10274558 | 10355704 | 7965910 | 8377317.5 | 7244869 | 5191804 | 4 | **0.746** |
| P70619 | Gsr | Glutathione reductase (Rattus norvegicus) | 441.06 | 22.41 | 46301 | 6 | 35378672 | 49030936 | 40286736 | 55128144 | 47405076 | 47371672 | 42312604 | 43399984 | 41079472 | 42652448 | 10 | **0.954** |
| P70665 | Siae | Sialate O-acetylesterase (Mus musculus) | 104.8 | 4.99 | 60775 | 2 | 2012178.5 | 0 | 0 | 1029314.56 | 1484797.62 | 1718135.75 | 2730220 | 1177020.25 | 1169223.25 | 1293380.38 | 8 | **1.787** |
| P70670 | Naca | Nascent polypeptide-associated complex subunit alpha, muscle-specific form (Mus musculu | 558.23 | 3.52 | 220499 | 8 | 41641012 | 65270412 | 61072680 | 68917192 | 75515136 | 63089704 | 50145168 | 57941792 | 51582916 | 52981764 | 6 | **0.883** |

| | | | | | | | | | | | | | | | | | |
|---|---|---|---|---|---|---|---|---|---|---|---|---|---|---|---|---|---|
| P70671 | Irf3 | Interferon regulatory factor 3 (Mus musculus) | 40.63 | 5.73 | 46852 | 1 | 177267.28 | 0 | 120336.45 | 0 | 0 | 0 | 158544.25 | 0 | 0 | 0 | 22 | **0.533** |
| P70677 | Casp3 | Caspase-3 (Mus musculus) | 270.11 | 25.27 | 31475 | 5 | 7595767 | 7719009.5 | 7443517 | 7289133.5 | 4831503.5 | 7495745 | 6683626.5 | 6294714 | 6476673.5 | 6615066 | 7 | **0.962** |
| P70697 | Urod | Uroporphyrinogen decarboxylase (Mus musculus) | 125.75 | 7.9 | 40692 | 2 | 1446897 | 2721500.25 | 2358886.5 | 2454930 | 2403989.75 | 1377301.75 | 1490272.25 | 2601473.25 | 2664881 | 2393269 | 11 | **0.925** |
| P70698 | Ctps1 | CTP synthase 1 (Mus musculus) | 395.86 | 14.21 | 66682 | 6 | 7857447.5 | 8452078 | 6328662 | 7763257 | 7634030 | 9410839 | 7863897 | 9971275 | 9083334 | 8889502 | 1 | **1.189** |
| P70699 | Gaa | Lysosomal alpha-glucosidase (Mus musculus) | 874.27 | 21.09 | 106248 | 15 | 13009021 | 9213706 | 15477415 | 13559162 | 17455130 | 10259434 | 11675503 | 14847395 | 13739850 | 11994965 | 9 | **0.910** |
| P70704 | Atp8a1 | Phospholipid-transporting ATPase IA (Mus musculus) | 40.92 | 0.69 | 131413 | 1 | 8997087 | 3798856.5 | 2904947 | 5176719.5 | 4579527 | 2432238 | 3021963.25 | 1812270 | 2240359 | 2921013.25 | 2 | **0.488** |
| P80313 | Cct7 | T-complex protein 1 subunit eta (Mus musculus) | 2202.12 | 65.62 | 59652 | 35 | 213906304 | 212684320 | 212987504 | 207152496 | 168338112 | 207638032 | 211301600 | 180296656 | 164369136 | 197049280 | 6 | **0.946** |
| P80314 | Cct2 | T-complex protein 1 subunit beta (Mus musculus) | 2533.16 | 71.96 | 57477 | 35 | 127843808 | 136972192 | 143661136 | 128805512 | 153406448 | 164674176 | 144381872 | 158076128 | 155357584 | 149504816 | 2 | **1.118** |
| P80315 | Cct4 | T-complex protein 1 subunit delta (Mus musculus) | 2360.89 | 72.17 | 58066 | 35 | 266959648 | 262477632 | 267382768 | 241521680 | 263680816 | 264549744 | 230609440 | 242381376 | 285692640 | 246814960 | 10 | **0.975** |
| P80316 | Cct5 | T-complex protein 1 subunit epsilon (Mus musculus) | 1483.83 | 61.18 | 59624 | 24 | 135085024 | 140743936 | 140526496 | 146924000 | 126894312 | 128886024 | 128860280 | 130320056 | 131508096 | 134073736 | 5 | **0.947** |
| P80317 | Cct6a | T-complex protein 1 subunit zeta (Mus musculus) | 1075.41 | 41.43 | 58004 | 17 | 121064440 | 134810896 | 134401456 | 139929024 | 126149232 | 129817816 | 123301024 | 122545680 | 129270816 | 125366304 | 7 | **0.960** |
| P80318 | Cct3 | T-complex protein 1 subunit gamma (Mus musculus) | 1805.03 | 55.6 | 60630 | 33 | 217010480 | 188533056 | 215274224 | 162401952 | 189723056 | 220839072 | 205859264 | 223914608 | 218148064 | 230878800 | 2 | **1.130** |
| P80349 | | Fuctinin-3 (Rattus norvegicus) | 52.86 | 38.46 | 4480 | 1 | 0 | 0 | 30382.67 | 0 | 0 | 0 | 0 | 226144.45 | 113498.44 | 135952.78 | 16 | **15.654** |
| P80386 | Prkab1 | 5'-AMP-activated protein kinase subunit beta-1 (Rattus norvegicus) | 40.25 | 5.93 | 30394 | 1 | 214478.94 | 456043.28 | 306549.22 | 350908.12 | 318858.19 | 331072.69 | 124412.39 | 522551.38 | 362877.84 | 238968.86 | 12 | **0.959** |
| P81117 | Nucb2 | Nucleobindin-2 (Mus musculus) | 56.84 | 2.14 | 50304 | 1 | 1692641.25 | 1392926.75 | 896539.12 | 1025427.62 | 0 | 866753.94 | 1116945.5 | 352484.56 | 1300252.12 | 0 | 9 | **0.726** |
| P82343 | Renbp | N-acylglucosamine 2-epimerase (Mus musculus) | 130.04 | 10 | 49771 | 3 | 2626777.25 | 4721174 | 5376092 | 4965427.5 | 5486900 | 3535803.5 | 2992867.75 | 4139923.5 | 3979427 | 4775441.5 | 6 | **0.838** |
| P83732 | Rpl24 | 60S ribosomal protein L24 (Rattus norvegicus) | 234.25 | 19.11 | 17779 | 4 | 36727396 | 26582906 | 19510832 | 19136190 | 30893572 | 39673188 | 39459616 | 29711366 | 25437844 | 27068860 | 7 | **1.215** |
| P83741 | Wnk1 | Serine/threonine-protein kinase WNK1 (Mus musculus) | 437.74 | 5.13 | 250934 | 10 | 11938602 | 10183865 | 11896301 | 9683717 | 13421974 | 12432727 | 13336086 | 11094149 | 12733392 | 10592071 | 9 | **1.054** |
| P83868 | Ptges3 | Prostaglandin E synthase 3 (Rattus norvegicus) | 192.13 | 26.25 | 18721 | 3 | 13611028 | 14572796 | 17552614 | 12527433 | 13294126 | 23001664 | 17376210 | 22000266 | 18184640 | 22433478 | 1 | **1.439** |
| P83877 | Txnl4a | Thioredoxin-like protein 4A (Mus musculus) | 73.83 | 12.68 | 16786 | 1 | 0 | 231970.75 | 334301.34 | 126813.27 | 260005.94 | 351683.12 | 312150.09 | 225581.41 | 235950.28 | 202363.27 | 9 | **1.393** |
| P83882 | Rpl36a | 60S ribosomal protein L36a (Mus musculus) | 181.47 | 25.47 | 12441 | 4 | 11972288 | 13115583 | 22306826 | 13838656 | 25345194 | 21115436 | 22119658 | 31547562 | 32774316 | 31580018 | 4 | **1.607** |
| P83917 | Cbx1 | Chromobox protein homolog 1 (Mus musculus) | 304.94 | 30.27 | 21418 | 4 | 6492450.5 | 8669239 | 6606536 | 8680786 | 7091089 | 6507255.5 | 7128973 | 6551016 | 7691298 | 5886437 | 8 | **0.899** |
| P83940 | Eloc | Elongin-C (Mus musculus) | 192.04 | 30.36 | 12473 | 2 | 1675784 | 1799360.12 | 1660002.12 | 1606532.38 | 1299713.75 | 1201393.62 | 1795337.75 | 1656620.75 | 1849445.25 | 2069844.75 | 9 | **1.066** |
| P83953 | Kpna1 | Importin subunit alpha-5 (Rattus norvegicus) | 453.17 | 20.45 | 60137 | 8 | 9134448 | 10305955 | 9216941 | 9320071 | 7877458.5 | 7913181 | 10566650 | 9120754 | 7756189 | 9033744 | 8 | **0.968** |
| P84078 | Arf1 | ADP-ribosylation factor 1 (Mus musculus) | 715.88 | 60.77 | 20697 | 11 | 39348128 | 95839120 | 95602224 | 94391264 | 80460104 | 93490720 | 78652912 | 93395880 | 91822944 | 97639040 | 12 | **1.122** |
| P84083 | Arf5 | ADP-ribosylation factor 5 (Rattus norvegicus) | 557.3 | 51.67 | 20530 | 9 | 30653430 | 76357976 | 74192672 | 70161128 | 60412720 | 76045872 | 63904228 | 73582800 | 70089552 | 74849280 | 10 | **1.150** |
| P84089 | Erh | Enhancer of rudimentary homolog (Mus musculus) | 181.85 | 46.15 | 12259 | 2 | 507275.41 | 628986.94 | 559891.31 | 478389.94 | 927071.94 | 383088.66 | 491823.03 | 387171.62 | 486247.94 | 651825.12 | 6 | **0.774** |
| P84091 | Ap2m1 | AP-2 complex subunit mu (Mus musculus) | 429.87 | 28.05 | 49655 | 8 | 10002883 | 13945726 | 14656140 | 13416661 | 13023676 | 13320534 | 8579368 | 13569248 | 12364033 | 14428068 | 10 | **0.957** |
| P84096 | Rhog | Rho-related GTP-binding protein RhoG (Mus musculus) | 374.01 | 40.84 | 21309 | 5 | 11658137 | 5483047 | 7182876 | 4415646.5 | 7890135 | 7325730.5 | 8081055 | 9354073 | 9831420 | 8776263 | 6 | **1.184** |
| P84099 | Rpl19 | 60S ribosomal protein L19 (Mus musculus) | 273.97 | 18.37 | 23466 | 4 | 21326104 | 12248624 | 15669608 | 10902060 | 15615862 | 27162728 | 18679608 | 21250476 | 20142082 | 21003152 | 4 | **1.429** |
| P84104 | Srsf3 | Serine/arginine-rich splicing factor 3 (Mus musculus) | 285.07 | 31.71 | 19330 | 5 | 35770416 | 39939236 | 28566510 | 43552932 | 36416392 | 28696608 | 24137610 | 23705708 | 14943968 | 23141692 | 1 | **0.622** |
| P84228 | Hist1h3b | Histone H3.2 (Mus musculus) b | 448.87 | 50.74 | 15388 | 8 | 1453005568 | 924865088 | 1236678400 | 1008597696 | 1197097344 | 1004275392 | 959959680 | 859621888 | 895114368 | 741153664 | 2 | **0.766** |
| P84817 | Fis1 | Mitochondrial fission 1 protein (Rattus norvegicus) | 251.57 | 25.66 | 16995 | 3 | 1330152.25 | 1862384.5 | 1978001.25 | 1624005.62 | 2071632.75 | 2204155.5 | 2245717.25 | 2766192.5 | 2566265.5 | 2547056 | 0 | **1.391** |
| P85094 | Isoc2a | Isochorismatase domain-containing protein 2A (Mus musculus) | 248.26 | 40.78 | 22417 | 4 | 3020936.75 | 2177100.25 | 2622966.5 | 579806.38 | 2271531.5 | 2596221.5 | 2724543.25 | 1959850.62 | 2719112 | 2241403.5 | 11 | **1.147** |
| P85972 | Vcl | Vinculin (Rattus norvegicus) | 459.83 | 10.69 | 116615 | 9 | 7768797 | 11644487 | 6856749 | 6119338.5 | 8440495 | 4673091 | 5082130 | 7575026.5 | 6232913 | 7682457.5 | 5 | **0.765** |
| P86182 | Ccdc22 | Coiled-coil domain-containing protein 22 (Rattus norvegicus) | 44.87 | 2.07 | 70855 | 1 | 345777.66 | 186280.12 | 262239.94 | 258424.48 | 241253.44 | 379464.25 | 414181.81 | 357731.38 | 309000.75 | 277456.28 | 2 | **1.343** |
| P97287 | Mcl1 | Induced myeloid leukemia cell differentiation protein Mcl-1 homolog (Mus musculus) | 80.48 | 6.34 | 35217 | 1 | 0 | 378622.94 | 496026.94 | 347836.5 | 441642.78 | 417866.62 | 0 | 555089.12 | 343613.38 | 552613.56 | 11 | **1.123** |
| P97298 | Serpinf1 | Pigment epithelium-derived factor (Mus musculus) 1 | 301.99 | 15.35 | 46234 | 5 | 6232210 | 12452724 | 10537165 | 14646486 | 12543689 | 6632242.5 | 7889910 | 5501916.5 | 6152079.5 | 5326374 | 2 | **0.558** |
| P97300 | Nptn | Neuroplastin (Mus musculus) | 89.24 | 6.3 | 44373 | 2 | 6036735 | 3657154.25 | 2308595.5 | 3173747 | 2854874.5 | 3737473 | 2880634 | 2149994.5 | 2092832 | 2217876.5 | 6 | **0.725** |
| P97310 | Mcm2 | DNA replication licensing factor MCM2 (Mus musculus) | 1580.09 | 40.27 | 102078 | 28 | 59470956 | 53643152 | 58772140 | 62156128 | 53391416 | 61912424 | 60149564 | 59458016 | 62438516 | 60264740 | 5 | **1.058** |
| P97311 | Mcm6 | DNA replication licensing factor MCM6 (Mus musculus) | 1365.64 | 32.4 | 92867 | 24 | 47725780 | 46455640 | 40578956 | 40953048 | 42920692 | 46133808 | 45824752 | 51106520 | 51917080 | 48238488 | 4 | **1.112** |
| P97315 | Csrp1 | Cysteine and glycine-rich protein 1 (Mus musculus) | 200.97 | 25.39 | 20583 | 3 | 6916446.5 | 5104188.5 | 6152360 | 7176738 | 7465257 | 6836558 | 5212408 | 7093198 | 7561485 | 8180020 | 9 | **1.063** |
| P97355 | Sms | Spermine synthase (Mus musculus) | 231.2 | 14.21 | 41313 | 5 | 1624861.25 | 3197776.25 | 3237563 | 3410036 | 2400440.25 | 1870883.5 | 1289870.25 | 1824463.38 | 1724325.5 | 1084998.5 | 3 | **0.562** |
| P97360 | Etv6 | Transcription factor ETV6 (Mus musculus) | 79.77 | 4.95 | 56406 | 2 | 995637.94 | 694702 | 739423.25 | 564853.75 | 660714.44 | 781408.38 | 1092770.62 | 973889.75 | 704606.62 | 604169.19 | 8 | **1.137** |
| P97369 | Ncf4 | Neutrophil cytosol factor 4 (Mus musculus) | 964.35 | 44.84 | 38707 | 15 | 52184884 | 61154656 | 56502432 | 62058840 | 62449848 | 59083220 | 48136060 | 55752204 | 61093556 | 55028112 | 6 | **0.948** |
| P97370 | Atp1b3 | Sodium/potassium-transporting ATPase subunit beta-3 (Mus musculus) | 151.09 | 14.39 | 31776 | 3 | 7787533.5 | 2225183 | 2190058.5 | 2640599.75 | 6100079 | 4648562.5 | 6522394 | 6499596 | 6590516 | 5782795 | 7 | **1.435** |
| P97371 | Psme1 | Proteasome activator complex subunit 1 (Mus musculus) | 634.1 | 42.97 | 28673 | 10 | 14385135 | 12542088 | 11980092 | 12696956 | 18399084 | 13617209 | 17408944 | 14126992 | 17436428 | 16495913 | 7 | **1.130** |
| P97372 | Psme2 | Proteasome activator complex subunit 2 (Mus musculus) | 1002.06 | 62.76 | 27057 | 14 | 47224916 | 62101708 | 65891908 | 54433940 | 52385360 | 56506776 | 66724404 | 64446096 | 62851192 | 63184680 | 5 | **1.112** |
| P97376 | Frg1 | Protein FRG1 (Mus musculus) | 57.99 | 6.2 | 29127 | 1 | 1180579.25 | 1280718 | 993617.75 | 0 | 1218764.25 | 724184.69 | 719588.62 | 723669.94 | 770610.31 | 883639 | 5 | **0.818** |
| P97377 | Cdk2 | Cyclin-dependent kinase 2 (Mus musculus) | 281.01 | 22.54 | 38978 | 6 | 4014900.5 | 4371166.5 | 3879721 | 3559576.5 | 4821577 | 4585952.5 | 4167169.25 | 5436352 | 5117690 | 6117347 | 3 | **1.231** |
| P97379 | G3bp2 | Ras GTPase-activating protein-binding protein 2 (Mus musculus) | 166.43 | 8.71 | 54088 | 3 | 10965440 | 9614618 | 9021899 | 4897115 | 114653.55 | 8249412 | 10976647 | 8769758 | 9393835 | 9514809 | 10 | **1.355** |
| P97384 | Anxa11 | Annexin A11 (Mus musculus) | 320.74 | 15.11 | 54079 | 7 | 4214381 | 4831773.5 | 4746256 | 7322786 | 5578962 | 5911113 | 5047644 | 5755047 | 4861722 | 6473705 | 7 | **1.051** |
| P97386 | Lig3 | DNA ligase 3 (Mus musculus) | 186.68 | 3.45 | 113072 | 3 | 1478221.5 | 2768243.5 | 872928.25 | 2517825 | 2643023 | 2545610.25 | 0 | 2631835.75 | 2554374.5 | 2495929.25 | 11 | **0.995** |
| P97429 | Anxa4 | Annexin A4 (Mus musculus) | 1024.3 | 51.72 | 35916 | 15 | 59255600 | 73223160 | 71870152 | 68003088 | 63781152 | 72924496 | 58483128 | 66302904 | 69845512 | 71973104 | 12 | **1.010** |
| P97452 | Bop1 | Ribosome biogenesis protein BOP1 (Mus musculus) | 199.6 | 10.38 | 82546 | 4 | 3675355.5 | 4791292.5 | 5689692 | 2083653.25 | 2480891.25 | 2766712 | 5235954.5 | 3663574.25 | 4085984.25 | 4152278.5 | 11 | **1.063** |
| P97461 | Rps5 | 40S ribosomal protein S5 (Mus musculus) | 775.31 | 54.41 | 22889 | 11 | 156658880 | 84235064 | 94919408 | 88416336 | 107443760 | 125020904 | 118510424 | 102425312 | 112038768 | 102958704 | 7 | **1.055** |
| P97470 | Ppp4c | Serine/threonine-protein phosphatase 4 catalytic subunit (Mus musculus) | 156.01 | 16.94 | 35080 | 3 | 3726598.75 | 5582711 | 6257145 | 3952520.75 | 5993129 | 4846534 | 4446849 | 4688410 | 4464959 | 5678270 | 11 | **0.946** |
| P97494 | Gclc | Glutamate--cysteine ligase catalytic subunit (Mus musculus) | 401.19 | 14.91 | 72571 | 7 | 9051773 | 7777894 | 8433269 | 7119801.5 | 7818564 | 8583082 | 9198885 | 8040261.5 | 7921559 | 7946490 | 7 | **1.037** |
| P97496 | Smarcc1 | SWI/SNF complex subunit SMARCC1 (Mus musculus) | 264.38 | 6.43 | 122890 | 5 | 4059841 | 4835200 | 3870547 | 4874515.5 | 4008589.25 | 4394964.5 | 4496380 | 4208362 | 4288043 | 4625060 | 10 | **1.017** |
| P97531 | Trip10 | Cdc42-interacting protein 4 (Rattus norvegicus) | 66.65 | 3.84 | 62799 | 1 | 335118.53 | 344899.66 | 0 | 312392.69 | 229304.14 | 166655.2 | 397201.28 | 104956.9 | 430396.41 | 480693.88 | 8 | **1.293** |
| P97532 | Mpst | 3-mercaptopyruvate sulfurtransferase (Rattus norvegicus) | 123.51 | 7.74 | 32940 | 2 | 3703089 | 1645879.75 | 1569312.62 | 3634448 | 3530131.5 | 3039054.5 | 3715401.75 | 3210453.5 | 1891891.62 | 2745461 | 12 | **1.037** |
| P97536 | Cand1 | Cullin-associated NEDD8-dissociated protein 1 (Rattus norvegicus) | 950.56 | 19.43 | 136362 | 18 | 31942042 | 38366916 | 30864720 | 33855376 | 28394342 | 33790356 | 31967758 | 36452588 | 36931408 | 37244848 | 7 | **1.079** |
| P97576 | Grpel1 | GrpE protein homolog 1, mitochondrial (Rattus norvegicus) | 209.36 | 14.29 | 24297 | 4 | 3708725.75 | 4078263.5 | 4067755.5 | 3597665.75 | 5262176 | 4089841 | 5419328.5 | 5621821 | 6099237 | 4737838 | 2 | **1.254** |
| P97690 | Smc3 | Structural maintenance of chromosomes protein 3 (Rattus norvegicus) | 519.6 | 12.59 | 138448 | 11 | 12718098 | 11935422 | 13191649 | 9142992 | 9158750 | 10845412 | 13911393 | 9337899 | 8079748 | 10449573 | 11 | **0.937** |
| P97760 | Polr2c | DNA-directed RNA polymerase II subunit RPB3 (Mus musculus) | 112.4 | 14.91 | 31443 | 2 | 222271.03 | 0 | 0 | 177841.73 | 207241.55 | 134212.39 | 0 | 111478.08 | 225723.11 | 90549.01 | 13 | **0.925** |
| P97770 | Thumpd3 | THUMP domain-containing protein 3 (Mus musculus) | 252.96 | 10.89 | 56431 | 5 | 1094792 | 1125407 | 2285634.75 | 290966.78 | 420878.75 | 2449214 | 742398.44 | 1968073.12 | 90738.84 | 2277759.75 | 10 | **1.443** |
| P97797 | Sirpa | Tyrosine-protein phosphatase non-receptor type substrate 1 (Mus musculus) | 146.2 | 8.97 | 56425 | 3 | 5187338 | 3378402.25 | 2455716 | 1508970.75 | 4187135.25 | 3634341.75 | 3583905.5 | 3185635.5 | 1677736.88 | 3068213 | 11 | **0.906** |
| P97807 | Fh | Fumarate hydratase, mitochondrial (Mus musculus) | 1486.48 | 55.03 | 54357 | 23 | 95058816 | 91331192 | 87320608 | 111664840 | 103442856 | 94978432 | 108559304 | 96611752 | 109902680 | 100724008 | 9 | **1.045** |
| P97814 | Pstpip1 | Proline-serine-threonine phosphatase-interacting protein 1 (Mus musculus) | 175.61 | 10.6 | 47590 | 3 | 3298289.5 | 2720597 | 3148040.5 | 2566720.5 | 2667500 | 2910745 | 2881597 | 2790923.5 | 2657393.75 | 3135563 | 12 | **0.998** |
| P97820 | Map4k4 | Mitogen-activated protein kinase kinase kinase kinase 4 (Mus musculus) | 103.38 | 1.87 | 140602 | 2 | 240295.31 | 547130.69 | 952075.06 | 845931.12 | 0 | 816068.81 | 593732.5 | 1006094.31 | 799359 | 1027046.88 | 6 | **1.641** |
| P97821 | Ctsc | Dipeptidyl peptidase 1 (Mus musculus) | 401.24 | 16.45 | 52376 | 6 | 28958576 | 23759194 | 27080090 | 24100440 | 28117452 | 20214556 | 25578926 | 23192922 | 24543050 | 26397802 | 6 | **0.908** |
| P97822 | Anp32e | Acidic leucine-rich nuclear phosphoprotein 32 family member E (Mus musculus) | 589.8 | 30.38 | 29622 | 7 | 21962734 | 24678106 | 28632266 | 18881974 | 26902646 | 38945192 | 32865268 | 37635564 | 36352756 | 36551836 | 0 | **1.506** |
| P97834 | Gps1 | COP9 signalosome complex subunit 1 (Rattus norvegicus) | 339.55 | 19.96 | 53428 | 7 | 4089036.5 | 11001200 | 12076607 | 7148200 | 7332231 | 12356017 | 10944530 | 9278188 | 10174922 | 7314660.5 | 9 | **1.202** |
| P97855 | G3bp1 | Ras GTPase-activating protein-binding protein 1 (Mus musculus) | 783.63 | 36.77 | 51829 | 15 | 45660328 | 44494084 | 40878564 | 35215328 | 33590968 | 40958272 | 45076504 | 42495684 | 39637592 | 42761864 | 10 | **1.055** |
| P97858 | Slc35b1 | Solute carrier family 35 member B1 (Mus musculus) | 67.6 | 3.11 | 35883 | 1 | 960815.94 | 1012539.81 | 1057267.25 | 1053471.38 | 909420.5 | 1597779.88 | 1334737.62 | 1345261.5 | 1467780.12 | 1394650 | 0 | **1.430** |
| P97874 | Gak | Cyclin-G-associated kinase (Rattus norvegicus) | 66.2 | 1.69 | 143703 | 1 | 363287.91 | 115411.01 | 0 | 165924.42 | 89402.75 | 0 | 0 | 0 | 0 | 0 | 15 | **0.000** |
| P97930 | Dtymk | Thymidylate kinase (Mus musculus) | 200.85 | 25.47 | 23915 | 4 | 1947245.5 | 4361854 | 4593203.5 | 782484.56 | 180500.75 | 3930290 | 4286431 | 2738772.75 | 2931329 | 2852268.5 | 10 | **1.411** |
| P98078 | Dab2 | Disabled homolog 2 (Mus musculus) | 1138.03 | 29.24 | 82312 | 17 | 29504352 | 47623988 | 43156480 | 42968548 | 41766804 | 44334832 | 43889088 | 42222152 | 42237024 | 41542628 | 12 | **1.045** |
| P98086 | C1qa | Complement C1q subcomponent subunit A (Mus musculus) | 145.06 | 11.84 | 25974 | 2 | 3400937 | 3782547.5 | 3328614.5 | 3221944.75 | 3323420.75 | 3206552.5 | 2664780.5 | 2762225.25 | 2604307.5 | 2690862.5 | 0 | **0.817** |
| P99026 | Psmb4 | Proteasome subunit beta type-4 (Mus musculus) | 587.34 | 48.48 | 29116 | 8 | 32410222 | 39153716 | 44993164 | 46437684 | 44226360 | 38692792 | 39664920 | 48849260 | 46176844 | 49099636 | 8 | **1.074** |
| P99027 | Rplp2 | 60S acidic ribosomal protein P2 (Mus musculus) | 234.71 | 59.13 | 11651 | 3 | 13043449 | 7708621 | 3722292.25 | 7624564 | 9760484 | 4395460.5 | 7734829 | 2102695 | 5884447 | 4022299 | 6 | **0.577** |
| P99028 | Uqcrh | Cytochrome b-c1 complex subunit 6, mitochondrial (Mus musculus) | 74.89 | 20.22 | 10435 | 1 | 773749.56 | 703484 | 775824.81 | 0 | 1440367.5 | 703389.5 | 911100.12 | 540401.5 | 995524.44 | 821745.12 | 11 | **1.075** |
| P99029 | Prdx5 | Peroxiredoxin-5, mitochondrial (Mus musculus) | 742.94 | 57.14 | 21897 | 12 | 72874224 | 66296248 | 54633096 | 63041872 | 96160616 | 61489948 | 62889976 | 70444688 | 62472052 | 64588868 | 8 | **0.912** |
| Q00422 | Gabpa | GA-binding protein alpha chain (Mus musculus) | 148.45 | 6.61 | 51344 | 3 | 3061131 | 4149442 | 3542823.5 | 2273805 | 3203495.75 | 3052539.5 | 2418004 | 3810011.75 | 2589136 | 3282706 | 10 | **0.934** |
| Q00519 | Xdh | Xanthine dehydrogenase/oxidase (Mus musculus) | 401.56 | 7.72 | 146562 | 8 | 4854168 | 5422345 | 5361773.5 | 7378425.5 | 4393010.5 | 6460845.5 | 4598764 | 5554357.5 | 4715651.5 | 4313307 | 10 | **0.936** |
| Q00612 | G6pdx | Glucose-6-phosphate 1-dehydrogenase X (Mus musculus) | 1491.82 | 52.43 | 59263 | 27 | 151315824 | 119815992 | 129002496 | 115762976 | 100667536 | 163341104 | 134504944 | 120142024 | 124011944 | 115308984 | 9 | **1.066** |
| Q00899 | Yy1 | Transcriptional repressor protein YY1 (Mus musculus) | 79.39 | 3.62 | 44717 | 2 | 2855569.75 | 2538407 | 2355415.25 | 2601688.5 | 2141931.75 | 2994893 | 3049007.75 | 2378584.5 | 1670443.25 | 1426564.5 | 12 | **0.922** |
| Q00PI9 | Hnrnpul2 | Heterogeneous nuclear ribonucleoprotein U-like protein 2 (Mus musculus) 2 | 531.42 | 17.32 | 84940 | 11 | 28518978 | 30651608 | 33268164 | 25995074 | 24395684 | 29691908 | 19654116 | 32144286 | 32290720 | 34559476 | 9 | **1.039** |

| | | | | | | | | | | | | | | | | | | |
|---|---|---|---|---|---|---|---|---|---|---|---|---|---|---|---|---|---|---|
| Q01205 | Dlst | Dihydrolipoyllysine-residue succinyltransferase component of 2-oxoglutarate dehydrogenas | 680.91 | 27.75 | 48925 | 12 | 41024912 | 49895684 | 42960624 | 40975988 | 41198312 | 41896380 | 44730880 | 41329472 | 45601404 | 42461820 | 8 | **1.000** |
| Q01320 | Top2a | DNA topoisomerase 2-alpha (Mus musculus) | 1157.26 | 17.47 | 172790 | 21 | 18423608 | 22388746 | 24410504 | 19578610 | 25063412 | 23346658 | 20825420 | 27584370 | 27676792 | 24491738 | 6 | **1.128** |
| Q01514 | Gbp1 | Guanylate-binding protein 1 (Mus musculus) | 69.15 | 3.23 | 67712 | 1 | 338373.44 | 72871.95 | 344969.5 | 0 | 395079.81 | 182578.77 | 332963.66 | 319618.44 | 0 | 241464.59 | 9 | **0.935** |
| Q01730 | Rsu1 | Ras suppressor protein 1 (Mus musculus) | 339.48 | 42.6 | 31550 | 6 | 5467974 | 6608809 | 6360709 | 7240437 | 6667082.5 | 7397604 | 4017695.75 | 7542903.5 | 8042763.5 | 6993344 | 6 | **1.051** |
| Q01768 | Nme2 | Nucleoside diphosphate kinase B (Mus musculus) | 788.65 | 73.68 | 17363 | 13 | 107890864 | 181191296 | 177660064 | 167176368 | 171818064 | 155008928 | 142665824 | 161742800 | 173136160 | 168222656 | 8 | **0.994** |
| Q02053 | Uba1 | Ubiquitin-like modifier-activating enzyme 1 (Mus musculus) | 1820.6 | 32.23 | 117809 | 26 | 81540096 | 72532472 | 73934936 | 69024808 | 60303936 | 77074416 | 76357320 | 72871888 | 70500112 | 72098808 | 10 | **1.032** |
| Q02257 | Jup | Junction plakoglobin (Mus musculus) | 138.93 | 4.43 | 81801 | 3 | 203488.58 | 0 | 0 | 3552943 | 0 | 0 | 0 | 0 | 42425.79 | 0 | 21 | **0.011** |
| Q02819 | Nucb1 | Nucleobindin-1 (Mus musculus) | 83.99 | 2.83 | 53409 | 1 | 986124.19 | 330640.28 | 242630.17 | 0 | 0 | 379017.66 | 467758.59 | 231119.55 | 457661.5 | 214328.62 | 10 | **1.122** |
| Q03145 | Epha2 | Ephrin type-A receptor 2 (Mus musculus) | 159.19 | 2.97 | 108852 | 3 | 613926.81 | 0 | 467412.81 | 0 | 0 | 0 | 537523.12 | 474239.53 | 562791.38 | 0 | 16 | **1.456** |
| Q03265 | Atp5f1a | ATP synthase subunit alpha, mitochondrial (Mus musculus) | 2365.82 | 62.75 | 59753 | 35 | 483799360 | 436804800 | 376579136 | 357170080 | 391420608 | 448232320 | 435916480 | 457448736 | 447214464 | 488434112 | 5 | **1.113** |
| Q03346 | Pmpcb | Mitochondrial-processing peptidase subunit beta (Rattus norvegicus) | 254.81 | 12.47 | 54265 | 5 | 9315134 | 6273233 | 6507673 | 5410440.5 | 6673948.5 | 8942186 | 8125055 | 8854573 | 8157672.5 | 8721905 | 5 | **1.252** |
| Q03963 | Eif2ak2 | Interferon-induced, double-stranded RNA-activated protein kinase (Mus musculus) | 317.3 | 16.31 | 58280 | 6 | 2072227.62 | 2478686.25 | 3402842.5 | 2375124.5 | 2272995.5 | 2623882.25 | 3642644.5 | 2767580 | 2537024.5 | 2523428.25 | 4 | **1.118** |
| Q04207 | Rela | Transcription factor p65 (Mus musculus) | 51.5 | 3.1 | 60212 | 1 | 843627.5 | 0 | 0 | 362214.06 | 454550.22 | 0 | 318386.38 | 0 | 405107.44 | 516514.41 | 14 | **0.747** |
| Q04447 | Ckb | Creatine kinase B-type (Mus musculus) | 1387.4 | 56.17 | 42713 | 17 | 117514336 | 102973536 | 100914368 | 96522608 | 96985616 | 111785392 | 109576512 | 99828168 | 99683840 | 94846616 | 12 | **1.002** |
| Q04692 | Smarcad1 | SWI/SNF-related matrix-associated actin-dependent regulator of chromatin subfamily A co | 132.98 | 2.94 | 116451 | 3 | 1303545 | 1224065 | 783457.31 | 350832.59 | 458192.78 | 1401886.75 | 1330087.5 | 1627761 | 1383895.25 | 1602792.12 | 0 | **1.783** |
| Q04750 | Top1 | DNA topoisomerase 1 (Mus musculus) | 592.05 | 18.25 | 90876 | 12 | 23401656 | 17897632 | 21902802 | 15446772 | 18486504 | 19473468 | 20792810 | 19601402 | 19953848 | 21778144 | 10 | **1.046** |
| Q04899 | Cdk18 | Cyclin-dependent kinase 18 (Mus musculus) | 185.5 | 7.54 | 51848 | 3 | 516443.78 | 353274.47 | 390966.78 | 400542 | 485665.94 | 545340.75 | 777635.44 | 543893.38 | 551789.75 | 468779.47 | 2 | **1.345** |
| Q04931 | Ssrp1 | FACT complex subunit SSRP1 (Rattus norvegicus) | 331.63 | 13.26 | 80914 | 6 | 5260386.5 | 6886181 | 6355084.5 | 10980424 | 7845075.5 | 4982594.5 | 5200523 | 6444360 | 5346584.5 | 6450465 | 5 | **0.761** |
| Q05144 | Rac2 | Ras-related C3 botulinum toxin substrate 2 (Mus musculus) | 363.7 | 41.15 | 21441 | 6 | 91847536 | 54897808 | 57596680 | 57033660 | 56775532 | 68515272 | 64411860 | 73416352 | 70195136 | 72100984 | 5 | **1.096** |
| Q05186 | Rcn1 | Reticulocalbin-1 (Mus musculus) | 52.77 | 4 | 38113 | 1 | 326796.44 | 0 | 0 | 0 | 460444.66 | 478787.19 | 0 | 124166.04 | 424500.81 | 352099.78 | 12 | **1.752** |
| Q05816 | Fabp5 | Fatty acid-binding protein 5 (Mus musculus) | 382.58 | 48.15 | 15137 | 6 | 21222334 | 26682604 | 28624108 | 14402062 | 32338888 | 25127510 | 22503334 | 21774266 | 22746420 | 23228764 | 10 | **0.936** |
| Q05910 | Adam8 | Disintegrin and metalloproteinase domain-containing protein 8 (Mus musculus) | 159.48 | 5.21 | 90046 | 3 | 3555995.5 | 2796557.5 | 2584460.25 | 1777756.12 | 2365081.25 | 2239560.5 | 1063794.25 | 3305451.25 | 4020753.75 | 3231096.5 | 11 | **1.060** |
| Q05CL8 | Larp7 | La-related protein 7 (Mus musculus) | 172.22 | 7.19 | 64802 | 3 | 5268646 | 1224404 | 3280721 | 3290794.5 | 4241239 | 3662100.25 | 3916641 | 3795523.25 | 3498752.5 | 3926824 | 10 | **1.086** |
| Q05D44 | Eif5b | Eukaryotic translation initiation factor 5B (Mus musculus) | 904.76 | 23.27 | 137616 | 21 | 30366344 | 33940344 | 34230324 | 29799496 | 26741484 | 28793116 | 36004160 | 32691340 | 32881654 | 35081072 | 8 | **1.067** |
| Q06138 | Cab39 | Calcium-binding protein 39 (Mus musculus) | 523.49 | 28.15 | 39843 | 10 | 4906968 | 10976890 | 11460170 | 14626813 | 11439513 | 9141026 | 10102899 | 11813697 | 11707668 | 12725285 | 11 | **1.039** |
| Q06486 | Csnk1d | Casein kinase I isoform delta (Rattus norvegicus) | 90.57 | 3.37 | 47316 | 2 | 1305916.38 | 1168030.5 | 1253397.25 | 877308.38 | 1139650.5 | 1253107.5 | 832150.06 | 913872.69 | 428464.22 | 765704.38 | 4 | **0.730** |
| Q07076 | Anxa7 | Annexin A7 (Mus musculus) | 439.11 | 17.71 | 49925 | 7 | 15892487 | 15452174 | 13098053 | 12442800 | 7338808 | 13866190 | 13008514 | 13063036 | 9842344 | 10270152 | 9 | **0.935** |
| Q07797 | Lgals3bp | Galectin-3-binding protein (Mus musculus) p | 732.12 | 27.04 | 64491 | 11 | 25707862 | 29071848 | 31179560 | 27672580 | 33248812 | 28926072 | 27490128 | 26540882 | 27655454 | 26409348 | 6 | **0.933** |
| Q07813 | Bax | Apoptosis regulator BAX (Mus musculus) | 579.63 | 51.56 | 21395 | 8 | 19731658 | 20532910 | 20252940 | 21050764 | 19609354 | 21733714 | 21285334 | 24968738 | 24826328 | 24664158 | 0 | **1.161** |
| Q07832 | Plk1 | Serine/threonine-protein kinase PLK1 (Mus musculus) | 43.02 | 1.33 | 68301 | 1 | 836415.38 | 1042527.5 | 1112161.38 | 1111126.5 | 1195985.25 | 2076348.38 | 1164353.25 | 1057197.25 | 1172287.75 | 1257152.5 | 5 | **1.270** |
| Q08024 | Cbfb | Core-binding factor subunit beta (Mus musculus) | 198.71 | 21.39 | 22030 | 3 | 2325480 | 4566283.5 | 4285152.5 | 3956059.75 | 3386145 | 3005585.5 | 3751455.25 | 3539231 | 4489688.5 | 4022414.75 | 12 | **1.016** |
| Q08288 | Lyar | Cell growth-regulating nucleolar protein (Mus musculus) | 141.56 | 7.22 | 43736 | 2 | 1761743.12 | 0 | 1204007.12 | 0 | 0 | 32668.53 | 1035885.44 | 265733.47 | 249355.62 | 333086.09 | 13 | **0.646** |
| Q08877 | Dnm3 | Dynamin-3 (Rattus norvegicus) | 156.63 | 4.26 | 97914 | 4 | 2931718 | 2387117 | 2403440.5 | 4696523 | 4937768 | 4319117 | 4610920 | 5141519 | 4947232 | 5147769 | 4 | **1.392** |
| Q08879 | Fbln1 | Fibulin-1 (Mus musculus) | 81.78 | 2.7 | 78033 | 2 | 2710085.5 | 6079276 | 3334981 | 10005648 | 6245238.5 | 1898972.25 | 1038047.06 | 1824472.5 | 1832106.5 | 1732128.12 | 0 | **0.293** |
| Q09014 | Ncf1 | Neutrophil cytosol factor 1 (Mus musculus) | 603.51 | 32.05 | 44667 | 11 | 19253396 | 45952368 | 40455672 | 48479824 | 46436488 | 22415024 | 29180578 | 28673484 | 37006788 | 32180666 | 5 | **0.745** |
| Q09200 | B4galnt1 | Beta-1,4 N-acetylgalactosaminyltransferase 1 (Mus musculus) 1 | 53.27 | 2.81 | 59212 | 1 | 333595.44 | 284548.84 | 444228.03 | 291453.47 | 325575.06 | 0 | 0 | 315291.28 | 412583.81 | 250763.19 | 7 | **0.583** |
| Q0GNC1 | Inf2 | Inverted formin-2 (Mus musculus) | 379.14 | 8.01 | 138560 | 7 | 1966926 | 3426698 | 3553334.5 | 2583294.5 | 1820954.5 | 3452694 | 3328651 | 2921120 | 4413638.5 | 2701950.5 | 7 | **1.260** |
| Q0P5W1 | Vps8 | Vacuolar protein sorting-associated protein 8 homolog (Mus musculus) | 41.45 | 0.49 | 161146 | 1 | 0 | 382561.31 | 580586.75 | 631844.06 | 422697.94 | 452170.41 | 582507.38 | 612221.94 | 424212.66 | 554180.25 | 8 | **1.301** |
| Q0VBL3 | Rbm15 | RNA-binding protein 15 (Mus musculus) | 68.08 | 1.77 | 105722 | 1 | 0 | 521729.34 | 549368.06 | 386631.06 | 697278.19 | 0 | 0 | 603562.25 | 637975.69 | 549773.56 | 13 | **0.831** |
| Q0VGB7 | Ppp4r2 | Serine/threonine-protein phosphatase 4 regulatory subunit 2 (Mus musculus) | 203.13 | 11.27 | 46479 | 3 | 2230880.75 | 2739081.5 | 2987432.5 | 1421489.25 | 1943370.62 | 1688417 | 2769780.25 | 2634592.25 | 2714994 | 2499195.25 | 11 | **1.087** |
| Q0VGY8 | Tanc1 | Protein TANC1 (Mus musculus) | 432.68 | 6.41 | 200805 | 8 | 5633271.5 | 3597693.75 | 4178627.75 | 2841803 | 2903775.75 | 4433927.5 | 4985822 | 5056471.5 | 3248630.75 | 3524036 | 9 | **1.109** |
| Q10728 | Ppp1r12a | Protein phosphatase 1 regulatory subunit 12A (Rattus norvegicus) a | 162.67 | 3.39 | 115283 | 3 | 1246718.38 | 1497826.88 | 1650108.62 | 1197917.62 | 1462849.5 | 813337.44 | 1447148.62 | 1375177.88 | 1093725.12 | 1243413.38 | 5 | **0.847** |
| Q11011 | Npepps | Puromycin-sensitive aminopeptidase (Mus musculus) | 752.7 | 20.54 | 103325 | 14 | 15837377 | 25355336 | 22086560 | 20717184 | 19258602 | 23120278 | 21210424 | 24345060 | 22235916 | 22529532 | 6 | **1.099** |
| Q11136 | Pepd | Xaa-Pro dipeptidase (Mus musculus) | 756.55 | 30.22 | 55029 | 13 | 25105406 | 31376732 | 32868558 | 34825112 | 33569736 | 33756312 | 30954656 | 31227370 | 31670044 | 32020256 | 10 | **1.012** |
| Q148V7 | Relch | RAB11-binding protein RELCH (Mus musculus) | 76.97 | 1.56 | 134587 | 2 | 334390.91 | 347933.56 | 397419.31 | 0 | 0 | 335639.94 | 409422 | 349412.94 | 0 | 0 | 14 | **1.014** |
| Q1AAU6 | Asap1 | Arf-GAP with SH3 domain, ANK repeat and PH domain-containing protein 1 (Rattus norve | 183.09 | 4.11 | 127088 | 4 | 6321704 | 5960687 | 4990708 | 5244963 | 4531440 | 6181544 | 4494980 | 4806405 | 4761036 | 5027787 | 8 | **0.934** |
| Q1HAQ0 | Lpcat1 | Lysophosphatidylcholine acyltransferase 1 (Rattus norvegicus) | 48.21 | 2.25 | 59762 | 1 | 1108198.38 | 1193512.25 | 1090095.38 | 1230459.38 | 956125.31 | 1070517 | 592579.38 | 1200059.75 | 0 | 0 | 6 | **0.513** |
| Q1HFZ0 | Nsun2 | tRNA (cytosine(34)-C(5))-methyltransferase (Mus musculus) | 383.39 | 13.34 | 85452 | 7 | 6960727 | 7740449 | 7433346 | 6655328.5 | 6510920 | 6931246.5 | 8847379 | 7686420 | 6363663.5 | 7874913 | 9 | **1.068** |
| Q27W01 | Rbm8a | RNA-binding protein 8A (Rattus norvegicus) | 236.96 | 36.78 | 19889 | 4 | 5502102.5 | 10290278 | 8926742 | 8965959 | 10434308 | 8258639 | 7685609.5 | 9213694 | 10621228 | 10417178 | 11 | **1.047** |
| Q2EMV9 | Parp14 | Poly [ADP-ribose] polymerase 14 (Mus musculus) | 214.61 | 3.41 | 203802 | 5 | 0 | 2419903.5 | 3134609 | 2004680 | 2424581 | 2690253 | 649163.25 | 2488909.25 | 2648398 | 2936226.25 | 8 | **1.143** |
| Q2L4X1 | Bzw2 | Basic leucine zipper and W2 domain-containing protein 2 (Mus musculus) molossinus (Jap | 363.79 | 17.66 | 48063 | 7 | 3972799.25 | 11462839 | 10954050 | 5805932 | 10264358 | 13643143 | 10116852 | 14533508 | 12870158 | 13688341 | 3 | **1.527** |
| Q2NL51 | Gsk3a | Glycogen synthase kinase-3 alpha (Mus musculus) | 154.03 | 8.16 | 51661 | 3 | 1609454 | 1978615.25 | 2147777 | 2117494 | 2381284.25 | 2297706 | 2426602 | 2808592.5 | 2750711.75 | 2726932 | 1 | **1.271** |
| Q2PQA9 | Kif5b | Kinesin-1 heavy chain (Rattus norvegicus) | 452.42 | 10.38 | 109531 | 7 | 5130430 | 4791567.5 | 3388108.75 | 4803820 | 4307289 | 4159721.25 | 3903730.5 | 3674091.5 | 10282608 | 4803657 | 11 | **1.196** |
| Q2TBE6 | Pi4k2a | Phosphatidylinositol 4-kinase type 2-alpha (Mus musculus) | 171.33 | 6.26 | 54258 | 3 | 1544865.5 | 1750450 | 1648498.5 | 2026435.5 | 1904699.12 | 2441995.75 | 2000139.62 | 2305551.25 | 1906197.75 | 1555401.75 | 6 | **1.150** |
| Q2YDW2 | Msto1 | Protein misato homolog 1 (Mus musculus) | 146.2 | 8.09 | 61230 | 3 | 2000856.5 | 3104984.25 | 2896252.5 | 1720733.75 | 2406291 | 1472877 | 2014588.62 | 3058486.75 | 2924174.75 | 3087755.5 | 11 | **1.035** |
| Q2YFS3 | Pilra | Paired immunoglobulin-like type 2 receptor alpha (Mus musculus) | 54.43 | 9.93 | 33064 | 1 | 0 | 215283.75 | 470589.38 | 0 | 0 | 159758.66 | 350325.28 | 165077.33 | 0 | 271590.88 | 14 | **1.380** |
| Q31125 | Slc39a7 | Zinc transporter SLC39A7 (Mus musculus) | 117.52 | 8.19 | 50657 | 2 | 955021.69 | 1881875.75 | 2150207 | 2382023 | 1755289 | 1711655.75 | 2232177.5 | 2234136.5 | 1070168.5 | 816688.69 | 10 | **0.884** |
| Q32PX2 | Aimp2 | Aminoacyl tRNA synthase complex-interacting multifunctional protein 2 (Rattus norvegicu | 209.88 | 13.12 | 35443 | 3 | 6143567.5 | 5189878.5 | 5357200 | 6241636.5 | 5222515.5 | 5700466 | 5215906 | 5376985.5 | 4864619 | 4491031.5 | 7 | **0.911** |
| Q32PX7 | Fubp1 | Far upstream element-binding protein 1 (Rattus norvegicus) | 620.72 | 20.81 | 67197 | 11 | 36406380 | 24423182 | 22914486 | 20911196 | 27348144 | 25440590 | 24650804 | 23442120 | 19637338 | 25064366 | 11 | **0.896** |
| Q3B7D1 | Ube2z | Ubiquitin-conjugating enzyme E2 Z (Rattus norvegicus) | 92.98 | 5.06 | 38352 | 2 | 759198.88 | 2838260.25 | 2823850 | 2860965 | 2754047 | 2150729.5 | 2288530 | 2035285 | 2606045 | 2403595.5 | 5 | **0.954** |
| Q3B7Z2 | Osbp | Oxysterol-binding protein 1 (Mus musculus) | 118.12 | 5.09 | 88797 | 3 | 1900412.75 | 1906468.75 | 1935081.38 | 2289589 | 1728887 | 2264267.75 | 2234615.5 | 1482855.88 | 1886164.88 | 1907911.62 | 12 | **1.002** |
| Q3B8Q0 | Mapre2 | Microtubule-associated protein RP/EB family member 2 (Rattus norvegicus) | 81.65 | 3.68 | 36988 | 1 | 576824.75 | 615435.38 | 843545.69 | 829524.88 | 1023305.56 | 733858.19 | 941689.94 | 748901.38 | 887182.12 | 761270.94 | 11 | **1.047** |
| Q3KRC5 | Dus3l | tRNA-dihydrouridine(47) synthase [NAD(P)(+)]-like (Rattus norvegicus) | 124.27 | 3.75 | 71540 | 2 | 993125.88 | 1081720.75 | 887292.62 | 790809.94 | 1037361.62 | 785925.25 | 329268.41 | 646542.44 | 716692.25 | 276667.44 | 0 | **0.575** |
| Q3KRD0 | Dars2 | Aspartate--tRNA ligase, mitochondrial (Rattus norvegicus) | 41.36 | 1.23 | 73953 | 1 | 1522067.62 | 662378.12 | 515875.16 | 1167562.88 | 0 | 834130.44 | 549462.81 | 817777.88 | 833308.69 | 0 | 12 | **0.785** |
| Q3KRE0 | Atad3 | ATPase family AAA domain-containing protein 3 (Rattus norvegicus) | 54 | 2.37 | 66759 | 1 | 106373.8 | 233197.94 | 206277.09 | 239615.2 | 224609 | 315536 | 104639.71 | 317634.22 | 244399.61 | 254955.17 | 5 | **1.225** |
| Q3KRE8 | Tubb2b | Tubulin beta-2B chain (Rattus norvegicus) | 2021.49 | 60.9 | 49953 | 30 | 752213248 | 732550848 | 632396032 | 754876800 | 677501504 | 691840576 | 671098944 | 717031680 | 691234560 | 671003840 | 8 | **0.970** |
| Q3MHC2 | Ints11 | Integrator complex subunit 11 (Rattus norvegicus) | 59 | 1.67 | 67841 | 1 | 1767699.25 | 1112083.62 | 1284802.12 | 1760012.12 | 1495443.38 | 1764356 | 1250121.25 | 1527629.25 | 1378503.75 | 1260104.12 | 11 | **0.968** |
| Q3MIE4 | Vat1 | Synaptic vesicle membrane protein VAT-1 homolog (Rattus norvegicus) | 219.13 | 11.88 | 43119 | 3 | 4660668 | 3929773.75 | 3671687 | 3996211.75 | 4220508 | 3708408 | 4217804 | 3726812.75 | 4470259.5 | 4552490 | 12 | **1.010** |
| Q3SWS8 | Rae1 | mRNA export factor (Rattus norvegicus) | 49.39 | 3.8 | 40919 | 1 | 709500.62 | 755425 | 541184.06 | 616201 | 732137.69 | 716181.81 | 776957.19 | 0 | 703610 | 865276.5 | 10 | **0.913** |
| Q3SWU3 | Hnrnpdl | Heterogeneous nuclear ribonucleoprotein D-like (Rattus norvegicus) | 92.18 | 5.59 | 35294 | 2 | 0 | 447407.69 | 0 | 513079.69 | 130135.61 | 0 | 0 | 0 | 0 | 0 | 20 | **0.000** |
| Q3T1I4 | Prrc1 | Protein PRRC1 (Rattus norvegicus) | 200.42 | 12.64 | 46295 | 3 | 282993.25 | 2631222.75 | 591819.06 | 1201155.62 | 1988891.12 | 1716974.25 | 956204.88 | 2562614.25 | 2472676 | 2582252.75 | 8 | **1.537** |
| Q3T1J9 | Mob1a | MOB kinase activator 1A (Rattus norvegicus) | 174.52 | 14.81 | 25080 | 3 | 12515315 | 9078002 | 10224601 | 9136170 | 9061754 | 11166084 | 4743449 | 10918446 | 13218388 | 10906078 | 8 | **1.019** |
| Q3TAA7 | Stk11ip | Serine/threonine-protein kinase 11-interacting protein (Mus musculus) | 75.23 | 2.99 | 118005 | 2 | 195589.77 | 345122.19 | 337201.91 | 426800.75 | 87487.64 | 176877.52 | 216545.86 | 342483.5 | 250522.78 | 261341.16 | 10 | **0.896** |
| Q3TBD2 | Arhgap45 | Rho GTPase-activating protein 45 (Mus musculus) 5 | 568.72 | 13.08 | 122902 | 10 | 13062787 | 9708082 | 12949353 | 11622820 | 9760884 | 10754667 | 10706235 | 11511787 | 8780046 | 9311292 | 6 | **0.894** |
| Q3TBT3 | Tmem173 | Stimulator of interferon genes protein (Mus musculus) | 63.13 | 6.61 | 42830 | 2 | 2052020.62 | 443376.72 | 2524991.25 | 2355221.5 | 1910911 | 2771048.5 | 2087194.5 | 2599140.75 | 2480985.5 | 2349919 | 5 | **1.323** |
| Q3TC93 | Hs1bp3 | HCLS1-binding protein 3 (Mus musculus) | 54.53 | 2.78 | 43691 | 1 | 535409.06 | 884058.88 | 930627 | 954612.88 | 775469.94 | 662977.38 | 854000.5 | 587412.81 | 649163.12 | 805311.12 | 7 | **0.872** |
| Q3TCH7 | Cul4a | Cullin-4A (Mus musculus) | 284.3 | 9.88 | 87753 | 6 | 4781478.5 | 7080744 | 6909449.5 | 5307375.5 | 5338577.5 | 5199933.5 | 6866233 | 7279280 | 6360220.5 | 7916297.5 | 8 | **1.143** |
| Q3TCN2 | Plbd2 | Putative phospholipase B-like 2 (Mus musculus) | 198.44 | 5.72 | 66289 | 4 | 817454.81 | 1542982.12 | 0 | 840471.12 | 1213164.75 | 1040069.69 | 1654590.5 | 1762398.5 | 1851449.75 | 1792600.38 | 2 | **1.835** |
| Q3TDD9 | Ppp1r21 | Protein phosphatase 1 regulatory subunit 21 (Mus musculus) | 86.41 | 2.44 | 88338 | 1 | 887088.62 | 609371.38 | 647499.56 | 0 | 0 | 535642.69 | 629216.62 | 759106.12 | 668512.5 | 542634.25 | 11 | **1.462** |
| Q3TDN2 | Faf2 | FAS-associated factor 2 (Mus musculus) | 68.57 | 3.37 | 52471 | 1 | 0 | 0 | 0 | 0 | 0 | 0 | 0 | 495071.66 | 0 | 541890.12 | 25 | **#DIV/0 !** |
| Q3TDQ1 | Stt3b | Dolichyl-diphosphooligosaccharide--protein glycosyltransferase subunit STT3B (Mus musc | 158.74 | 4.74 | 93246 | 4 | 2824201.25 | 6198129 | 5240104 | 4940309 | 5696151 | 4130775.25 | 3724268 | 5856052.5 | 6958969.5 | 6782852 | 9 | **1.103** |
| Q3TEA8 | Hp1bp3 | Heterochromatin protein 1-binding protein 3 (Mus musculus) | 242.21 | 11.91 | 60867 | 5 | 10282799 | 7382309.5 | 7659368 | 8317345 | 9435642 | 8690544 | 8836079 | 8017773 | 9428142 | 8540821 | 11 | **1.010** |
| Q3THG9 | Aarsd1 | Alanyl-tRNA editing protein Aarsd1 (Mus musculus) | 294.25 | 20.15 | 44971 | 6 | 8270639.5 | 8887594 | 10853466 | 10037101 | 9562295 | 8775022 | 6582771.5 | 10042494 | 7314448 | 8470148 | 6 | **0.865** |
| Q3THK3 | Gtf2f1 | General transcription factor IIF subunit 1 (Mus musculus) | 71.88 | 3.15 | 57241 | 1 | 1020438.38 | 823996.88 | 0 | 0 | 640976.31 | 709754.81 | 0 | 0 | 0 | 0 | 17 | **0.286** |
| Q3THK7 | Gmps | GMP synthase [glutamine-hydrolyzing] (Mus musculus) | 1298.29 | 52.24 | 76723 | 24 | 44068864 | 34043600 | 36157812 | 33181354 | 35701976 | 41765844 | 36438720 | 41035940 | 40011484 | 38612408 | 5 | **1.080** |
| Q3THS6 | Mat2a | S-adenosylmethionine synthase isoform type-2 (Mus musculus) | 687.44 | 42.03 | 43689 | 14 | 44860708 | 38286280 | 44006872 | 37489016 | 27142270 | 45188768 | 45072640 | 41711096 | 33920884 | 44645060 | 7 | **1.098** |
| Q3TIR1 | Trappc13 | Trafficking protein particle complex subunit 13 (Mus musculus) 3 | 78.03 | 6.47 | 46477 | 1 | 0 | 192327.94 | 348968.97 | 138087.89 | 289239.22 | 0 | 0 | 263515.94 | 125883.96 | 197682.94 | 10 | **0.606** |
| Q3TIR3 | Ric8a | Synembryn-A (Mus musculus) | 44.98 | 1.7 | 59847 | 1 | 0 | 354382.81 | 363939.22 | 482419.41 | 284847.28 | 0 | 0 | 282554.97 | 215199.47 | 84900.38 | 6 | **0.392** |

| | | | | | | | | | | | | | | | | | | |
|---|---|---|---|---|---|---|---|---|---|---|---|---|---|---|---|---|---|---|
| Q3TIU4 | Pde12 | 2',5'-phosphodiesterase 12 (Mus musculus) | 235.83 | 9.7 | 67514 | 4 | 4766882 | 3549997.5 | 2755406 | 3583953 | 3003300.5 | 3288755.25 | 3006403.5 | 3529620.5 | 3245498.25 | 3743991.75 | 12 | **0.952** |
| Q3TIV5 | Zc3h15 | Zinc finger CCCH domain-containing protein 15 (Mus musculus) | 227.12 | 11.27 | 48327 | 4 | 4057767.5 | 2633444 | 3382199.5 | 3401009.5 | 1904849.62 | 3604204.5 | 3127724.5 | 3285259.75 | 3647112 | 3269951 | 11 | **1.101** |
| Q3TIX9 | Usp39 | U4/U6.U5 tri-snRNP-associated protein 2 (Mus musculus) | 590.19 | 28.72 | 65146 | 12 | 12362832 | 7618129 | 8880674 | 8092273 | 7212941 | 9935423 | 12095863 | 11285878 | 10161073 | 9523307 | 5 | **1.200** |
| Q3TKT4 | Smarca4 | Transcription activator BRG1 (Mus musculus) | 145.65 | 2.23 | 181427 | 4 | 34339128 | 46555244 | 36569404 | 48105404 | 6323394 | 32219892 | 30677140 | 4436711 | 3694159.5 | 10341157 | 3 | **0.473** |
| Q3TKY6 | Cwc27 | Spliceosome-associated protein CWC27 homolog (Mus musculus) | 59.6 | 3.2 | 53543 | 1 | 510271.38 | 263432.84 | 498662.06 | 0 | 575640.25 | 455411.34 | 353496.53 | 0 | 0 | 335193.12 | 9 | **0.619** |
| Q3TL44 | Nlrx1 | NLR family member X1 (Mus musculus) | 38.88 | 1.03 | 107831 | 1 | 0 | 0 | 506359.19 | 598727.88 | 338314.25 | 0 | 0 | 369728.72 | 345381.94 | 396396.78 | 14 | **0.770** |
| Q3TLH4 | Prrc2c | Protein PRRC2C (Mus musculus) | 366.48 | 3.51 | 310892 | 7 | 9532738 | 5888545 | 7677910 | 5366596 | 8786728 | 8492806 | 6114395.5 | 6808090 | 8422195 | 8017864 | 12 | **1.016** |
| Q3TMH2 | Scrn3 | Secernin-3 (Mus musculus) | 49.5 | 2.63 | 47661 | 1 | 1809655.88 | 1682473.75 | 1964575.5 | 480410.84 | 279669.12 | 2109675.5 | 2005462.88 | 338299.19 | 351425.09 | 297527.5 | 12 | **0.821** |
| Q3TPX4 | Exoc5 | Exocyst complex component 5 (Mus musculus) | 88.87 | 2.97 | 81738 | 2 | 940419.75 | 2294833.5 | 2217488.25 | 2235272 | 1923180.5 | 1520775 | 2223268.25 | 2403786 | 1890530.75 | 2139554.25 | 12 | **1.059** |
| Q3TRM8 | Hk3 | Hexokinase-3 (Mus musculus) | 1956.47 | 45.88 | 100101 | 31 | 57781496 | 62493720 | 57037204 | 57935980 | 48004472 | 58758080 | 62588992 | 60951296 | 57417248 | 56339324 | 9 | **1.045** |
| Q3TVI8 | Pbxip1 | Pre-B-cell leukemia transcription factor-interacting protein 1 (Mus musculus) | 204.04 | 6.74 | 81164 | 3 | 2374768.5 | 2776942 | 2139301 | 2499906.75 | 806240.5 | 1637743.75 | 1182627.38 | 1901761 | 2441537.25 | 2141970 | 8 | **0.878** |
| Q3TW96 | Uap1l1 | UDP-N-acetylhexosamine pyrophosphorylase-like protein 1 (Mus musculus) | 629.99 | 25.84 | 56614 | 11 | 25733976 | 19517770 | 20362490 | 20695276 | 17958416 | 25308898 | 23068966 | 23944460 | 25222712 | 22772852 | 5 | **1.154** |
| Q3TX08 | Trmt1 | tRNA (guanine(26)-N(2))-dimethyltransferase (Mus musculus) | 90.14 | 4.52 | 72350 | 2 | 1728786 | 752070.38 | 772663.88 | 587046.19 | 734456.38 | 2044849.88 | 1003542.69 | 834581.44 | 1273376.38 | 1431912.12 | 4 | **1.440** |
| Q3TXS7 | Psmd1 | 26S proteasome non-ATPase regulatory subunit 1 (Mus musculus) | 1552.84 | 34.63 | 105730 | 25 | 60259384 | 63004744 | 63124420 | 65827872 | 58826472 | 70017416 | 64679388 | 65449084 | 60538428 | 62590732 | 8 | **1.039** |
| Q3TYS2 | Cybc1 | Cytochrome b-245 chaperone 1 (Mus musculus) | 112.9 | 11.23 | 20921 | 2 | 1860910.88 | 3930811.5 | 4446984 | 4572753 | 4368694 | 6000743 | 4633038 | 4405565 | 4420078.5 | 4833709 | 4 | **1.267** |
| Q3TYX3 | Smyd5 | SET and MYND domain-containing protein 5 (Mus musculus) | 92.83 | 6.97 | 47095 | 2 | 0 | 331656.53 | 649999.19 | 262714.38 | 266925.44 | 289033.06 | 0 | 323281.44 | 346521.56 | 380164.75 | 11 | **0.886** |
| Q3TZZ7 | Esyt2 | Extended synaptotagmin-2 (Mus musculus) | 44.69 | 1.78 | 94139 | 1 | 648793.5 | 349914.53 | 414144.12 | 362730.66 | 409504.75 | 411765.22 | 519958.69 | 386136.75 | 395057.66 | 374391.03 | 12 | **0.955** |
| Q3U0K8 | Ogfod1 | Prolyl 3-hydroxylase OGFOD1 (Mus musculus) | 69.72 | 2.75 | 62734 | 1 | 144818.31 | 292540.97 | 280073.91 | 0 | 0 | 301834.78 | 242531.11 | 283232.38 | 0 | 167384.59 | 11 | **1.387** |
| Q3U0V1 | Khsrp | Far upstream element-binding protein 2 (Mus musculus) | 999.77 | 33.29 | 76775 | 17 | 70874584 | 40020748 | 35044372 | 39076224 | 45549996 | 50109724 | 45297668 | 47146928 | 42463544 | 44270052 | 8 | **0.994** |
| Q3U186 | Rars2 | Probable arginine--tRNA ligase, mitochondrial (Mus musculus) | 132.86 | 4.84 | 65336 | 2 | 1316311 | 1722723.38 | 2177191.25 | 1607789.75 | 1581320.75 | 1682577 | 1583139 | 1971338.25 | 1426464.25 | 1620357.62 | 12 | **0.986** |
| Q3U1J4 | Ddb1 | DNA damage-binding protein 1 (Mus musculus) | 1751.3 | 34.39 | 126853 | 33 | 70929504 | 85254432 | 86696392 | 72983608 | 86193680 | 96732704 | 86977600 | 90916792 | 90156560 | 93162400 | 0 | **1.139** |
| Q3U1V6 | Uevld | Ubiquitin-conjugating enzyme E2 variant 3 (Mus musculus) | 67.8 | 3.82 | 51681 | 2 | 2413583 | 1363812.25 | 1512347.5 | 1805903.75 | 785874.81 | 1783159 | 1656082.75 | 1858644 | 1782498.25 | 1737345.5 | 9 | **1.119** |
| Q3U487 | Hectd3 | E3 ubiquitin-protein ligase HECTD3 (Mus musculus) | 51.29 | 1.63 | 97347 | 1 | 0 | 242367.19 | 0 | 0 | 348451.5 | 0 | 0 | 318638.19 | 262604.06 | 430653.78 | 15 | **1.713** |
| Q3U7R1 | Esyt1 | Extended synaptotagmin-1 (Mus musculus) | 487.36 | 11.9 | 121554 | 8 | 5373417 | 3916422.5 | 5686836.5 | 7901715.5 | 6281188.5 | 6127386 | 6976279.5 | 6908750 | 6673687.5 | 6739070 | 6 | **1.146** |
| Q3U821 | Wdr75 | WD repeat-containing protein 75 (Mus musculus) | 176.43 | 5.06 | 94037 | 3 | 1007243.69 | 1691061.75 | 2001581.25 | 2139355.5 | 1260996.38 | 1255863.38 | 1476796.5 | 1257257 | 2452025 | 1722151.12 | 12 | **1.008** |
| Q3U9G9 | Lbr | Lamin-B receptor (Mus musculus) | 313.66 | 11.82 | 71440 | 6 | 4768862.5 | 3866566 | 4008049.25 | 3997509.75 | 6316783.5 | 4543442.5 | 6371026 | 4544171.5 | 5171761.5 | 3763338.5 | 10 | **1.063** |
| Q3UA06 | Trip13 | Pachytene checkpoint protein 2 homolog (Mus musculus) | 38.34 | 2.31 | 48377 | 1 | 1082075.5 | 1130620.75 | 1540915.38 | 969646.5 | 0 | 1537356.25 | 1383212.75 | 1436573.88 | 1222505.25 | 1220367.38 | 5 | **1.440** |
| Q3UBX0 | Tmem109 | Transmembrane protein 109 (Mus musculus) | 50.36 | 4.94 | 26306 | 1 | 3331605.25 | 2772231.75 | 2210719.5 | 1026353.62 | 484454.09 | 495016.5 | 2887550.25 | 946302.81 | 3053803.25 | 1790292.38 | 12 | **0.934** |
| Q3UDE2 | Ttll12 | Tubulin--tyrosine ligase-like protein 12 (Mus musculus) | 511.89 | 23.63 | 74043 | 10 | 15444505 | 17918522 | 18086784 | 15323082 | 15561519 | 18806434 | 14964853 | 17301778 | 16255100 | 16515503 | 11 | **1.018** |
| Q3UEB3 | Puf60 | Poly(U)-binding-splicing factor PUF60 (Mus musculus) | 790.96 | 33.33 | 60249 | 13 | 24548658 | 26645376 | 26608820 | 20076234 | 26007418 | 34078100 | 30628748 | 33765948 | 32338582 | 33656128 | 0 | **1.328** |
| Q3UFK8 | Frmd8 | FERM domain-containing protein 8 (Mus musculus) | 38.36 | 1.72 | 51828 | 1 | 964880.88 | 274488.31 | 518888.28 | 527483.5 | 304717.72 | 113354.4 | 609893.12 | 635930.94 | 1083200.62 | 563261.88 | 8 | **1.160** |
| Q3UFY7 | Nt5c3b | 7-methylguanosine phosphate-specific 5'-nucleotidase (Mus musculus) | 144.21 | 16.67 | 34425 | 2 | 780870.62 | 1055232.75 | 1169098.62 | 1430541.62 | 1088567.62 | 1931237 | 1264819 | 1222412.62 | 1678097 | 1805563.12 | 2 | **1.430** |
| Q3UGR5 | Hdhd2 | Haloacid dehalogenase-like hydrolase domain-containing protein 2 (Mus musculus) | 159.48 | 16.22 | 28730 | 3 | 1965299.5 | 2100103.75 | 1205310.75 | 2339126.75 | 906614.81 | 2303734.5 | 2510678.5 | 2254660.5 | 2457517.75 | 2209564.5 | 3 | **1.378** |
| Q3UH60 | Dip2b | Disco-interacting protein 2 homolog B (Mus musculus) | 225.73 | 4.07 | 171126 | 4 | 3967617.75 | 4617975.5 | 3686570.75 | 2240776 | 2574834 | 2953555.25 | 3774382.75 | 2976113 | 3010725.75 | 2859599 | 11 | **0.911** |
| Q3UH66 | Wnk2 | Serine/threonine-protein kinase WNK2 (Mus musculus) | 131.06 | 1.44 | 227527 | 3 | 4068116.5 | 2891375 | 3486689.5 | 2737578.5 | 4057095 | 4228751 | 4325061 | 2921931.25 | 5136733 | 3270700 | 6 | **1.153** |
| Q3UHD6 | Snx27 | Sorting nexin-27 (Mus musculus) | 190.68 | 10.95 | 60989 | 4 | 2517909.25 | 0 | 1378059.62 | 2377469.5 | 2163051.25 | 2013848 | 1159552.38 | 2062660.5 | 2117644.5 | 2162615 | 9 | **1.128** |
| Q3UHH8 | Gxylt1 | Glucoside xylosyltransferase 1 (Mus musculus) | 44.03 | 1.98 | 46491 | 1 | 374074.47 | 696062.94 | 789065 | 640963.25 | 534315.44 | 778802.44 | 705211 | 903371.19 | 701771.19 | 583424.06 | 6 | **1.210** |
| Q3UHX2 | Pdap1 | 28 kDa heat- and acid-stable phosphoprotein (Mus musculus) | 167.74 | 14.92 | 20605 | 3 | 7366567 | 1781623.75 | 518644.41 | 800407.94 | 1337123 | 892885.75 | 2799406.75 | 998148.56 | 1516140.25 | 863881.25 | 12 | **0.599** |
| Q3UIA2 | Arhgap17 | Rho GTPase-activating protein 17 (Mus musculus) 7 | 243.67 | 7.68 | 92202 | 5 | 4826562 | 3398990 | 2712266 | 3997811 | 3157647.5 | 4566444.5 | 4544586 | 3504999.25 | 3095209.5 | 4290601 | 9 | **1.105** |
| Q3UIR3 | Dtx3l | E3 ubiquitin-protein ligase DTX3L (Mus musculus) | 61.15 | 2.14 | 83044 | 1 | 284249.5 | 224927.39 | 478999.84 | 352704.31 | 345557.66 | 382615.06 | 425688.03 | 486737.16 | 514749.06 | 428296.53 | 3 | **1.327** |
| Q3UJB9 | Edc4 | Enhancer of mRNA-decapping protein 4 (Mus musculus) | 299.05 | 6.47 | 152484 | 5 | 4494430 | 2113295.25 | 4124441.5 | 2731072 | 5314508.5 | 3239492.75 | 3211124 | 3795970 | 2914909.5 | 3637334.25 | 10 | **0.895** |
| Q3UJD6 | Usp19 | Ubiquitin carboxyl-terminal hydrolase 19 (Mus musculus) | 153.07 | 3.38 | 150549 | 3 | 1108965.5 | 1015479.44 | 1627942.75 | 1270500 | 547870.31 | 1747083.38 | 1186797 | 1686620.38 | 1486844 | 1310752.5 | 4 | **1.332** |
| Q3UKJ7 | Smu1 | WD40 repeat-containing protein SMU1 (Mus musculus) | 359.66 | 19.49 | 57544 | 7 | 6291397 | 6714535.5 | 4906546.5 | 5451260.5 | 4646887 | 5559478 | 3953750.5 | 6380655 | 5967218.5 | 5491659 | 12 | **0.977** |
| Q3ULD5 | Mccc2 | Methylcrotonoyl-CoA carboxylase beta chain, mitochondrial (Mus musculus) | 54.64 | 3.02 | 61379 | 1 | 199526.81 | 157401.73 | 296416.69 | 141198.97 | 174367.41 | 87580.55 | 296831.84 | 208382.66 | 207188.67 | 172609.62 | 10 | **1.004** |
| Q3UM45 | Ppp1r7 | Protein phosphatase 1 regulatory subunit 7 (Mus musculus) | 566.28 | 32.96 | 41292 | 9 | 10800060 | 12116022 | 11205692 | 12723160 | 11495228 | 12392752 | 13426841 | 12162006 | 12258195 | 11654357 | 5 | **1.061** |
| Q3UMB9 | Washc4 | WASH complex subunit 4 (Mus musculus) | 510 | 9.38 | 136370 | 8 | 1862345.12 | 3173424.5 | 3234365 | 1927891.62 | 1816527.5 | 1495368.62 | 2356370.75 | 1646451.25 | 2199913.25 | 1900992.62 | 8 | **0.799** |
| Q3UMR5 | Mcu | Calcium uniporter protein, mitochondrial (Mus musculus) | 72.15 | 4.57 | 39682 | 1 | 0 | 347409.62 | 333271.91 | 330203.47 | 0 | 525499 | 258681.39 | 287080.41 | 195840.27 | 289600.31 | 13 | **1.540** |
| Q3UMY5 | Eml4 | Echinoderm microtubule-associated protein-like 4 (Mus musculus) | 85.26 | 1.62 | 110027 | 1 | 952931.25 | 841997.69 | 1025294.44 | 1262035.25 | 1688902.5 | 665389.69 | 0 | 852573.19 | 888107.5 | 900877.38 | 3 | **0.573** |
| Q3UND0 | Skap2 | Src kinase-associated phosphoprotein 2 (Mus musculus) | 257.88 | 18.99 | 40712 | 6 | 2107906.5 | 8363920.5 | 8715658 | 5015301 | 4993175 | 8144516 | 6318866.5 | 8790310 | 7409656.5 | 8577280 | 7 | **1.344** |
| Q3UPF5 | Zc3hav1 | Zinc finger CCCH-type antiviral protein 1 (Mus musculus) | 40 | 1.16 | 106688 | 1 | 1296532.5 | 1038206.12 | 973408 | 1095291.75 | 991427.94 | 785411 | 1108989.25 | 1019654.31 | 1073826.75 | 1140969.5 | 12 | **0.951** |
| Q3UPL0 | Sec31a | Protein transport protein Sec31A (Mus musculus) | 587.13 | 12.11 | 133569 | 10 | 4411726 | 6439149.5 | 6936811.5 | 6448564 | 6508781 | 6090538.5 | 6236932 | 6996571.5 | 6569364.5 | 6849002 | 10 | **1.065** |
| Q3UQ44 | Iqgap2 | Ras GTPase-activating-like protein IQGAP2 (Mus musculus) | 229.75 | 2.16 | 180528 | 5 | 8297574.5 | 12624501 | 16662658 | 12446482 | 15336705 | 17262984 | 17484360 | 15078880 | 15259038 | 14243006 | 6 | **1.214** |
| Q3UQ84 | Tars2 | Threonine--tRNA ligase, mitochondrial (Mus musculus) | 45.38 | 1.38 | 81700 | 1 | 816132.12 | 544442.75 | 514187 | 595448 | 760193.56 | 641215.31 | 567506.06 | 647395.75 | 936064.19 | 525139.25 | 11 | **1.027** |
| Q3URS9 | Ccdc51 | Coiled-coil domain-containing protein 51 (Mus musculus) | 56.62 | 2.71 | 45132 | 1 | 346921.69 | 386497.44 | 350093.88 | 517183.94 | 433242.12 | 297525 | 412594.19 | 422225.56 | 512705.31 | 702473.94 | 10 | **1.154** |
| Q3UUQ7 | Pgap1 | GPI inositol-deacylase (Mus musculus) | 121.47 | 5.53 | 104578 | 3 | 109433.04 | 43013.34 | 321867.41 | 0 | 0 | 230487.22 | 306551.16 | 354697.47 | 0 | 337974.31 | 8 | **2.593** |
| Q3UVK0 | Ermp1 | Endoplasmic reticulum metallopeptidase 1 (Mus musculus) | 134.47 | 2.56 | 100148 | 2 | 358405.59 | 233326.27 | 300968.72 | 0 | 0 | 1307504.5 | 0 | 382712.56 | 685259.94 | 85895.11 | 9 | **2.757** |
| Q3UVL4 | Vps51 | Vacuolar protein sorting-associated protein 51 homolog (Mus musculus) | 127.05 | 4.35 | 86187 | 3 | 1191989.12 | 3073681.5 | 2651867 | 2106598 | 2547568.25 | 2144782.5 | 1276936.75 | 2360490 | 1288386.62 | 2499997 | 8 | **0.827** |
| Q3UW53 | Fam129a | Protein Niban (Mus musculus) | 1701.28 | 44.38 | 102649 | 31 | 81109184 | 89051328 | 101453448 | 92577976 | 88006408 | 99086768 | 101830768 | 112417904 | 107088328 | 115568224 | 1 | **1.185** |
| Q3UX61 | Naa11 | N-alpha-acetyltransferase 11 (Mus musculus) | 53.11 | 3.67 | 24671 | 1 | 827302 | 1498078.88 | 2443606.5 | 1478649.38 | 1498743.75 | 2781949 | 1737509 | 2030036.88 | 2029892.5 | 2153879.75 | 4 | **1.386** |
| Q3UYV9 | Ncbp1 | Nuclear cap-binding protein subunit 1 (Mus musculus) | 310.45 | 9.49 | 91927 | 6 | 573722.44 | 2347083.5 | 1713562.75 | 2896377.5 | 2256572.5 | 1287062 | 1108976.5 | 1758196.25 | 2085535.88 | 1977929 | 8 | **0.840** |
| Q3UZ39 | Lrrfip1 | Leucine-rich repeat flightless-interacting protein 1 (Mus musculus) | 1135.23 | 34.16 | 79249 | 17 | 23058266 | 24978792 | 23409748 | 23343684 | 27060216 | 21710678 | 20354938 | 22091172 | 21423062 | 24443868 | 3 | **0.903** |
| Q3UZA1 | Rcsd1 | CapZ-interacting protein (Mus musculus) | 412.92 | 22.57 | 44114 | 6 | 7564033 | 12039472 | 12550518 | 7690956 | 10356821 | 7985307 | 7113993 | 8188080 | 8489729 | 8841860 | 8 | **0.809** |
| Q3V0A6 | Gm5592 | Uncharacterized protein C2orf78 homolog (Mus musculus) | 38.24 | 0.82 | 93729 | 1 | 3204431.75 | 2292052 | 3643453.5 | 1887739.38 | 3988163.75 | 4440378.5 | 4255606 | 4982961 | 4012531.5 | 4620919.5 | 0 | **1.486** |
| Q3V0K9 | Pls1 | Plastin-1 (Mus musculus) | 486.04 | 7.46 | 70408 | 8 | 22223824 | 131744808 | 134057464 | 127301248 | 127433112 | 151793712 | 125342056 | 125974328 | 119238312 | 122171744 | 9 | **1.187** |
| Q3V1L4 | Nt5c2 | Cytosolic purine 5'-nucleotidase (Mus musculus) | 425.02 | 17.86 | 64809 | 8 | 9808834 | 8284897 | 9012114 | 6042348 | 7701998 | 10311300 | 8725167 | 10177543 | 9420280 | 10046025 | 3 | **1.192** |
| Q3V1V3 | Esf1 | ESF1 homolog (Mus musculus) | 48.09 | 1.42 | 98048 | 1 | 448779.38 | 514928.72 | 409977.44 | 0 | 546878.38 | 446353.56 | 500110.22 | 443447.25 | 470221.75 | 315221.38 | 11 | **1.133** |
| Q3V3R1 | Mthfd1l | Monofunctional C1-tetrahydrofolate synthase, mitochondrial (Mus musculus) | 1398.94 | 31.12 | 105729 | 26 | 29308136 | 28254766 | 25969334 | 30890300 | 31916630 | 33436486 | 27306894 | 32953984 | 32173038 | 33753204 | 4 | **1.091** |
| Q497V5 | Srbd1 | S1 RNA-binding domain-containing protein 1 (Mus musculus) | 68.27 | 1.32 | 110281 | 1 | 0 | 649830.94 | 550142.69 | 759333.12 | 545100.81 | 0 | 111896.78 | 526908.38 | 620389.75 | 321928.03 | 7 | **0.631** |
| Q498C8 | Rer1 | Protein RER1 (Rattus norvegicus) | 50.41 | 5.61 | 22988 | 1 | 1423167.12 | 950259.31 | 954051.06 | 1064903.88 | 872241.94 | 1262113.25 | 1817037.38 | 1277327.5 | 1044639.56 | 1015910.5 | 6 | **1.219** |
| Q498E0 | Txndc12 | Thioredoxin domain-containing protein 12 (Rattus norvegicus) | 45 | 8.82 | 19019 | 1 | 1287012.88 | 1435979.75 | 1509713.75 | 1060755 | 1661560.75 | 1045075.12 | 1006914.06 | 1501357.38 | 1499987.25 | 1845088.5 | 11 | **0.992** |
| Q498T9 | Lrrc8c | Volume-regulated anion channel subunit LRRC8C (Rattus norvegicus) | 171.64 | 3.86 | 92463 | 2 | 349968.94 | 380758 | 474467.59 | 0 | 0 | 531143.44 | 671805.25 | 647488.56 | 0 | 0 | 11 | **1.535** |
| Q498U4 | Sarnp | SAP domain-containing ribonucleoprotein (Rattus norvegicus) | 313.86 | 34.29 | 23605 | 6 | 12804634 | 3873200 | 3948610 | 6422611 | 9417400 | 5653122 | 6622600 | 6431827 | 6798265.5 | 6207856 | 12 | **0.870** |
| Q499N6 | Ubxn1 | UBX domain-containing protein 1 (Rattus norvegicus) | 159.32 | 9.43 | 33582 | 2 | 1536358.38 | 195549.48 | 120804.35 | 199610.12 | 155741.58 | 1544238 | 383008.75 | 1824922.88 | 261438.66 | 710040.69 | 3 | **2.139** |
| Q499T7 | Cfap20 | Cilia- and flagella-associated protein 20 (Rattus norvegicus) | 59.34 | 5.49 | 19132 | 2 | 292704.06 | 313483.97 | 1174394.38 | 800829.25 | 694402.56 | 681093.38 | 0 | 1062310.62 | 683303.5 | 793415.88 | 11 | **0.983** |
| Q4AE70 | Carm1 | Histone-arginine methyltransferase CARM1 (Rattus norvegicus) | 41.69 | 1.08 | 70341 | 1 | 1826851.5 | 2401627.75 | 2394618 | 3321650.5 | 2493460.25 | 2468044.75 | 2036275.62 | 2156466.25 | 3137756.25 | 2964278.75 | 12 | **1.026** |
| Q4FK66 | Prpf38a | Pre-mRNA-splicing factor 38A (Mus musculus) | 86.09 | 5.77 | 37437 | 2 | 1167750.88 | 3477439.5 | 2894351 | 2649883.75 | 2830066.75 | 3148631 | 2935932.5 | 2527726.75 | 2785051.5 | 3187236.5 | 10 | **1.120** |
| Q4FZF3 | Ddx49 | Probable ATP-dependent RNA helicase DDX49 (Mus musculus) | 50.82 | 2.29 | 54094 | 1 | 540495.12 | 699656.69 | 591003.44 | 0 | 619275.19 | 0 | 0 | 621613.25 | 675896.12 | 596219.56 | 14 | **0.773** |
| Q4FZY0 | Efhd2 | EF-hand domain-containing protein D2 (Rattus norvegicus) | 1025.68 | 58.58 | 26759 | 17 | 119112176 | 132995320 | 127130888 | 140701120 | 127126440 | 133339216 | 100150904 | 125974240 | 134285872 | 126340848 | 10 | **0.958** |
| Q4G009 | Mcts1 | Malignant T-cell-amplified sequence 1 (Rattus norvegicus) | 138.01 | 21.43 | 20550 | 2 | 444015.31 | 1817556.5 | 1101633.25 | 1024551.88 | 415228.97 | 607853.19 | 565317.5 | 578004.69 | 1119879.5 | 1178324.12 | 11 | **0.843** |
| Q4KLG9 | Zfand2b | AN1-type zinc finger protein 2B (Rattus norvegicus) | 39.8 | 2.72 | 27925 | 1 | 593851.19 | 585558.25 | 150457.03 | 323708.31 | 508814.41 | 498066.88 | 520232.41 | 533627.69 | 449487.09 | 556351 | 12 | **1.183** |
| Q4KLH4 | Pspc1 | Paraspeckle component 1 (Rattus norvegicus) | 203.28 | 9 | 58760 | 3 | 14239966 | 9143534 | 11932610 | 9736154 | 12998823 | 16818524 | 14027440 | 17913566 | 17902590 | 18752540 | 1 | **1.471** |
| Q4KLL0 | Tcea1 | Transcription elongation factor A protein 1 (Rattus norvegicus) | 563.16 | 30.56 | 33893 | 8 | 15670610 | 7453654 | 6478488 | 5375097.5 | 12946462 | 12377179 | 13890050 | 11805755 | 12668088 | 11968170 | 9 | **1.309** |
| Q4KLL4 | Tm9sf4 | Transmembrane 9 superfamily member 4 (Rattus norvegicus) | 71.26 | 2.95 | 74675 | 1 | 454580.28 | 650301.38 | 871451.25 | 659275.81 | 0 | 744500.56 | 913805.25 | 1293832 | 853878.06 | 1084226.5 | 2 | **1.855** |
| Q4KLN7 | Arfgap3 | ADP-ribosylation factor GTPase-activating protein 3 (Rattus norvegicus) | 77.66 | 3.43 | 57671 | 2 | 1751095.12 | 2846952.5 | 2350354.5 | 2058777.88 | 1728236.12 | 2177898.75 | 1944896.75 | 2409928.5 | 1808774.62 | 1809856 | 12 | **0.946** |
| Q4KM64 | Jagn1 | Protein jagunal homolog 1 (Rattus norvegicus) | 54.44 | 6.56 | 21102 | 1 | 0 | 719884.56 | 751021.5 | 705206.94 | 673667.19 | 888890.56 | 0 | 1266216.12 | 1018965.31 | 904128.69 | 5 | **1.431** |
| Q4KM65 | Nudt21 | Cleavage and polyadenylation specificity factor subunit 5 (Rattus norvegicus) | 78.04 | 7.05 | 26240 | 2 | 4087442.5 | 597240.56 | 745374.94 | 0 | 536816.69 | 3084044.5 | 2911019.5 | 2268207 | 1217568.62 | 1054641 | 5 | **1.766** |
| Q4KM73 | Cmpk1 | UMP-CMP kinase (Rattus norvegicus) | 199.49 | 27.04 | 22169 | 4 | 9434235 | 6064060 | 5203832 | 7343600.5 | 0 | 6866793 | 5990151 | 5928312 | 5328613 | 4478681 | 10 | **1.019** |
| Q4KML4 | Abracl | Costars family protein ABRACL (Mus musculus) | 310.8 | 46.91 | 9030 | 5 | 12599569 | 14457032 | 16783398 | 14650006 | 15705167 | 15653754 | 14975596 | 16746050 | 18420192 | 16272064 | 6 | **1.106** |

| | | | | | | | | | | | | | | | | | |
|---|---|---|---|---|---|---|---|---|---|---|---|---|---|---|---|---|---|
| Q4LDD4 | Arap1 | Arf-GAP with Rho-GAP domain, ANK repeat and PH domain-containing protein 1 (Mus m | 166.2 | 3.72 | 162276 | 4 | 2632927.75 | 2355005 | 3327428.5 | 2054825.12 | 2393750.75 | 3005829 | 2867770.75 | 3446299 | 4504738 | 3763128 | 2 | **1.378** |
| Q4PJX1 | Odr4 | Protein odr-4 homolog (Mus musculus) | 42.1 | 3.13 | 49998 | 1 | 666121.19 | 1037623.81 | 0 | 1094711.25 | 565714.62 | 0 | 587309.25 | 742867.12 | 632879.5 | 502253.34 | 9 | **0.733** |
| Q4QQR9 | Memo1 | Protein MEMO1 (Rattus norvegicus) | 334.61 | 34.01 | 33679 | 7 | 8367387 | 7366381.5 | 7237779.5 | 8804404 | 8309387.5 | 9213606 | 2409312.25 | 7885345.5 | 8723745 | 8977701 | 9 | **0.928** |
| Q4QQT4 | Ppp2r1b | Serine/threonine-protein phosphatase 2A 65 kDa regulatory subunit A beta isoform (Rattus | 952.18 | 27.79 | 66006 | 14 | 30414374 | 38035604 | 32616214 | 43195724 | 32113824 | 30292814 | 22858140 | 30455058 | 31762784 | 30730108 | 3 | **0.828** |
| Q4QQT6 | Brix1 | Ribosome biogenesis protein BRX1 homolog (Rattus norvegicus) | 239.63 | 17.9 | 41115 | 5 | 8674598 | 9188876 | 9821438 | 8323401 | 9177312 | 12878808 | 8957159 | 10958750 | 11239105 | 10929412 | 3 | **1.216** |
| Q4QRB4 | Tubb3 | Tubulin beta-3 chain (Rattus norvegicus) | 1493.02 | 38.67 | 50419 | 22 | 547593344 | 553280896 | 442939040 | 569307712 | 475627552 | 502208480 | 486642848 | 539377600 | 567297984 | 530170496 | 12 | **1.014** |
| Q4QRL3 | Ccdc88b | Coiled-coil domain-containing protein 88B (Mus musculus) | 42.32 | 0.61 | 166608 | 1 | 1231904 | 523701.03 | 154106.56 | 711841.75 | 800591.5 | 271035.28 | 270808.47 | 1169041.5 | 952341.12 | 887576.06 | 11 | **1.038** |
| Q4V7C7 | Actr3 | Actin-related protein 3 (Rattus norvegicus) | 1321.91 | 73.92 | 47357 | 23 | 198307456 | 205025344 | 215385904 | 228014192 | 206483632 | 194967616 | 191728432 | 192524128 | 192542800 | 206241056 | 2 | **0.929** |
| Q4V898 | Rbmx | RNA-binding motif protein, X chromosome (Rattus norvegicus) | 240.57 | 14.1 | 42258 | 5 | 10834805 | 11528639 | 9023996 | 12974274 | 9924338 | 8433307 | 7292068 | 7069242 | 7543161.5 | 8794702 | 0 | **0.721** |
| Q4V8B7 | Hsdl1 | Inactive hydroxysteroid dehydrogenase-like protein 1 (Rattus norvegicus) | 71.78 | 3.33 | 36908 | 1 | 592306 | 449721.88 | 555048.25 | 0 | 407405.38 | 324099.88 | 398230.25 | 382254.34 | 273437.88 | 419476.12 | 6 | **0.897** |
| Q4V8C8 | Cdc73 | Parafibromin (Rattus norvegicus) | 287.26 | 15.44 | 60603 | 6 | 2605586.5 | 3385201 | 3421354.5 | 2524780.25 | 2359396.25 | 3463529.25 | 2772673.75 | 2239412.75 | 2133223.5 | 2711396.25 | 11 | **0.932** |
| Q4V8K2 | Ctnnbl1 | Beta-catenin-like protein 1 (Rattus norvegicus) | 49.23 | 1.42 | 64948 | 1 | 1051324.12 | 0 | 0 | 0 | 0 | 814983.12 | 745723.75 | 1008621 | 866979.62 | 940236.88 | 11 | **4.163** |
| Q4VAA2 | Cdv3 | Protein CDV3 (Mus musculus) | 145.62 | 9.25 | 29729 | 2 | 5325290 | 3501784 | 1597291.12 | 4976250 | 6432218 | 2549518.5 | 2418795.5 | 3006186 | 2895997 | 2253745.75 | 5 | **0.601** |
| Q4VBE8 | Wdr18 | WD repeat-containing protein 18 (Mus musculus) | 95.96 | 6.5 | 47211 | 2 | 0 | 863808.06 | 1119176.88 | 0 | 858899.56 | 2017834.5 | 980719.19 | 1060148.12 | 1098457.75 | 1272209.62 | 4 | **2.262** |
| Q4VSI4 | Usp7 | Ubiquitin carboxyl-terminal hydrolase 7 (Rattus norvegicus) | 69.63 | 1.18 | 128431 | 1 | 0 | 690507.31 | 829778.5 | 891608.56 | 611532.19 | 442881.25 | 0 | 533057.81 | 520992.59 | 525972.81 | 5 | **0.669** |
| Q501J6 | Ddx17 | Probable ATP-dependent RNA helicase DDX17 (Mus musculus) | 934.04 | 27.23 | 72399 | 16 | 165612656 | 129435256 | 138777584 | 136331184 | 138531360 | 136791520 | 148742112 | 141286448 | 142319968 | 145372016 | 7 | **1.008** |
| Q505F5 | Lrrc47 | Leucine-rich repeat-containing protein 47 (Mus musculus) | 77.57 | 3.1 | 63590 | 1 | 671021.56 | 260074.59 | 416031.81 | 0 | 613441.5 | 649216.56 | 625915.31 | 475536.72 | 665864.81 | 582367.88 | 7 | **1.530** |
| Q505J9 | Atad1 | ATPase family AAA domain-containing protein 1 (Rattus norvegicus) | 93.04 | 6.37 | 40717 | 2 | 0 | 1791564.88 | 1952841.5 | 1367804.5 | 648572.69 | 4247046 | 938425 | 3174438.25 | 3186920 | 2417350.5 | 3 | **2.424** |
| Q52KE7 | Ccnl1 | Cyclin-L1 (Mus musculus) | 78.3 | 2.82 | 60133 | 1 | 56271.7 | 258925.83 | 211764.94 | 161156.03 | 73994.52 | 285698.84 | 315535.06 | 232919.44 | 247204.39 | 240101.33 | 3 | **1.734** |
| Q52KI8 | Srrm1 | Serine/arginine repetitive matrix protein 1 (Mus musculus) | 344.44 | 8.88 | 106862 | 5 | 9669911 | 6467827.5 | 4838035 | 7669569.5 | 9523510 | 5222832.5 | 4581908.5 | 3327135.5 | 4995507 | 5758731 | 3 | **0.626** |
| Q52KR3 | Prune2 | Protein prune homolog 2 (Mus musculus) | 57.36 | 0.81 | 339516 | 2 | 540047 | 307380.84 | 506459.97 | 376961.31 | 410662.66 | 556495.81 | 425455.12 | 665512.31 | 0 | 458437.44 | 9 | **0.983** |
| Q53UA7 | Taok3 | Serine/threonine-protein kinase TAO3 (Rattus norvegicus) | 84.99 | 2.34 | 105474 | 2 | 2485178 | 2384714.25 | 2497752.25 | 3922931.75 | 2219338 | 1743223.88 | 2609313.75 | 1655937.25 | 2011461 | 1718826.88 | 4 | **0.721** |
| Q562B5 | Pgam5 | Serine/threonine-protein phosphatase PGAM5, mitochondrial (Rattus norvegicus) | 102.19 | 5.9 | 32060 | 2 | 3889792.25 | 2780980.5 | 3390709.5 | 1629783.75 | 1713298.5 | 2857467.5 | 3282633 | 2578649 | 1964509 | 2001310 | 12 | **0.946** |
| Q568Z6 | Ist1 | IST1 homolog (Rattus norvegicus) | 45.46 | 5.19 | 39942 | 1 | 320795.03 | 439765.94 | 438936.94 | 335942.41 | 122493.15 | 410257.53 | 567358.31 | 311306.69 | 515440.5 | 435866.12 | 8 | **1.351** |
| Q569A6 | Kdelr1 | ER lumen protein-retaining receptor 1 (Rattus norvegicus) | 82.93 | 10.38 | 24546 | 2 | 185771.39 | 211551.7 | 322982.69 | 0 | 121455.92 | 686753.5 | 315116.19 | 405477.62 | 287732.62 | 76109.71 | 6 | **2.104** |
| Q569Z5 | Ddx46 | Probable ATP-dependent RNA helicase DDX46 (Mus musculus) | 559.03 | 12.98 | 117448 | 11 | 20002456 | 15323773 | 14655870 | 16308816 | 11181548 | 15221487 | 13453340 | 14773798 | 12993709 | 15190117 | 8 | **0.925** |
| Q569Z6 | Thrap3 | Thyroid hormone receptor-associated protein 3 (Mus musculus) | 221.76 | 5.47 | 108178 | 4 | 2280747.25 | 1183738.88 | 1783255.25 | 2036282.25 | 3538712.25 | 1888569.75 | 2223219.75 | 1774727.75 | 2056241.75 | 2080051 | 12 | **0.926** |
| Q56B11 | Pelp1 | Proline-, glutamic acid- and leucine-rich protein 1 (Rattus norvegicus) | 81.27 | 1.06 | 119139 | 1 | 184244.62 | 243490.41 | 104059.9 | 168346.8 | 292790.88 | 352479.94 | 238385.11 | 233243.05 | 217688.45 | 255177.61 | 7 | **1.306** |
| Q571E4 | Galns | N-acetylgalactosamine-6-sulfatase (Mus musculus) | 166.55 | 7.69 | 57673 | 3 | 6050547 | 5404389 | 3617329.5 | 3077360 | 6632030 | 5280490.5 | 4632355 | 4804377.5 | 4812842.5 | 4976136.5 | 10 | **0.989** |
| Q571I9 | Aldh16a1 | Aldehyde dehydrogenase family 16 member A1 (Mus musculus) 1 | 368.43 | 12.72 | 84756 | 6 | 4812438 | 4861150.5 | 5456270.5 | 4508058.5 | 4792862 | 4937386.5 | 4128358 | 5746199.5 | 5488023.5 | 5063835.5 | 7 | **1.038** |
| Q58A65 | Spag9 | C-Jun-amino-terminal kinase-interacting protein 4 (Mus musculus) | 358.56 | 6.43 | 146219 | 7 | 2889142.75 | 5914338.5 | 7910536 | 4178703.75 | 4691779 | 6007663.5 | 6110933.5 | 5562925.5 | 6262985.5 | 7326360 | 6 | **1.222** |
| Q5BJP2 | Cwc15 | Spliceosome-associated protein CWC15 homolog (Rattus norvegicus) | 105.33 | 16.16 | 26638 | 2 | 562845.56 | 1718306.62 | 1379522.88 | 1013148.38 | 1511152.25 | 1375220.12 | 1830635.75 | 1574494.12 | 1455221.5 | 1490100.38 | 8 | **1.249** |
| Q5BJS0 | Dhx30 | ATP-dependent RNA helicase DHX30 (Rattus norvegicus) | 183.3 | 3.1 | 133997 | 3 | 1803916.75 | 1777560 | 1455601.75 | 1291888.75 | 1250856.75 | 1624114.75 | 1916842 | 1638278.5 | 1515994.75 | 1647848.75 | 8 | **1.101** |
| Q5BJX0 | Ntmt1 | N-terminal Xaa-Pro-Lys N-methyltransferase 1 (Rattus norvegicus) | 57.32 | 3.14 | 25464 | 1 | 1701745.12 | 1253153.12 | 1118476 | 1147964 | 1083242.25 | 1625541.38 | 1383735.88 | 1671098 | 1122903.12 | 1373408 | 7 | **1.138** |
| Q5BJY6 | Amdhd2 | N-acetylglucosamine-6-phosphate deacetylase (Rattus norvegicus) | 209.69 | 18.83 | 43539 | 5 | 11012388 | 5191218 | 5166683 | 3588774.5 | 5280698 | 6750389.5 | 5851539.5 | 5333514 | 5572911 | 5342657.5 | 5 | **0.954** |
| Q5BK20 | Jpt2 | Jupiter microtubule associated homolog 2 (Rattus norvegicus) | 95.08 | 14.21 | 20070 | 2 | 4681651.5 | 3267982.75 | 2935719 | 3850664.75 | 4669305.5 | 3873741.5 | 3639329.75 | 3204153 | 3139972.5 | 3238027.25 | 8 | **0.881** |
| Q5DTM8 | Rnf20 | E3 ubiquitin-protein ligase BRE1A (Mus musculus) | 145.76 | 4.21 | 113520 | 2 | 208121.12 | 549304.06 | 570314.75 | 227798.34 | 202043.84 | 346961.72 | 1072108.5 | 774453.69 | 673635.19 | 487266.62 | 4 | **1.909** |
| Q5EZ72 | Enpp7 | Ectonucleotide pyrophosphatase/phosphodiesterase family member 7 (Rattus norvegicus) | 69.06 | 2.28 | 49837 | 1 | 852677.25 | 465745.16 | 0 | 678380.38 | 0 | 0 | 715221.06 | 0 | 783643.44 | 630934.38 | 15 | **1.067** |
| Q5F2E7 | Nufip2 | Nuclear fragile X mental retardation-interacting protein 2 (Mus musculus) | 107.87 | 3.9 | 75657 | 2 | 0 | 624360.88 | 614833.56 | 0 | 724683.44 | 0 | 723741.06 | 694265 | 791095.88 | 896294.44 | 8 | **1.581** |
| Q5FVI6 | Atp6v1c1 | V-type proton ATPase subunit C 1 (Rattus norvegicus) 1 | 813 | 43.98 | 43901 | 16 | 30835556 | 22162240 | 26695450 | 39215800 | 18763346 | 33008434 | 29862826 | 31442902 | 31807896 | 31930632 | 6 | **1.148** |
| Q5FVM3 | Retreg1 | Reticulophagy regulator 1 (Rattus norvegicus) | 100.57 | 4.79 | 52782 | 2 | 1096852.62 | 1293803.88 | 3189037.5 | 821258.06 | 788327.69 | 3228096 | 1628529.88 | 3847404.5 | 3555591 | 3330852.25 | 1 | **2.169** |
| Q5FVM4 | Nono | Non-POU domain-containing octamer-binding protein (Rattus norvegicus) | 868.9 | 35.08 | 54925 | 15 | 98739960 | 99397592 | 119708160 | 81678424 | 96182568 | 96497160 | 98192224 | 102187464 | 126247000 | 103330176 | 8 | **1.062** |
| Q5FVQ0 | Slc39a8 | Zinc transporter ZIP8 (Rattus norvegicus) | 45 | 1.52 | 50171 | 1 | 0 | 623555.31 | 647709.81 | 853610.25 | 3680984.25 | 551254.31 | 308730.88 | 614246.25 | 2759840.75 | 2770414 | 11 | **1.206** |
| Q5FVQ4 | Mlec | Malectin (Rattus norvegicus) | 154.8 | 11 | 32418 | 3 | 4683591.5 | 7371547 | 3046281.25 | 6821777 | 5480973 | 6866030 | 6058107 | 6942579 | 6785399 | 7276558 | 7 | **1.238** |
| Q5FVR4 | Eri1 | 3'-5' exoribonuclease 1 (Rattus norvegicus) | 92.76 | 5.8 | 39580 | 2 | 926489.25 | 2041614.5 | 2703901.5 | 2305315.25 | 2732283.75 | 3014537.5 | 2523567 | 2681228 | 2736984.5 | 2750816 | 4 | **1.280** |
| Q5FWK3 | Arhgap1 | Rho GTPase-activating protein 1 (Mus musculus) | 443.01 | 27.56 | 50411 | 8 | 6215351 | 7722780.5 | 8901978 | 4484900 | 3780669.5 | 10750000 | 9104083 | 10603899 | 9572534 | 10604433 | 0 | **1.628** |
| Q5FWT7 | Ublcp1 | Ubiquitin-like domain-containing CTD phosphatase 1 (Rattus norvegicus) | 97.05 | 8.81 | 36867 | 2 | 834794.12 | 1442068.25 | 948154.56 | 1012358.56 | 979918.38 | 1136268 | 392010.56 | 901432.75 | 971089.38 | 814424.5 | 7 | **0.808** |
| Q5HZF2 | Wbp4 | WW domain-binding protein 4 (Rattus norvegicus) | 62.64 | 4.55 | 42157 | 1 | 0 | 0 | 658863.5 | 0 | 672402.81 | 677306.81 | 705466.19 | 626029.81 | 593754.31 | 505997.94 | 9 | **2.335** |
| Q5HZY2 | Sar1b | GTP-binding protein SAR1b (Rattus norvegicus) | 325.06 | 39.39 | 22410 | 6 | 11983010 | 10411978 | 12387958 | 10027146 | 13523839 | 15760600 | 12258312 | 13874462 | 12501842 | 13350593 | 4 | **1.161** |
| Q5I0E7 | Tmed9 | Transmembrane emp24 domain-containing protein 9 (Rattus norvegicus) | 266.13 | 20 | 27028 | 5 | 17064900 | 18991704 | 19769380 | 20442268 | 18943096 | 17600648 | 13106192 | 18817036 | 19970874 | 18614012 | 7 | **0.925** |
| Q5I0H4 | Tmco1 | Calcium load-activated calcium channel (Rattus norvegicus) | 150.25 | 19.68 | 21175 | 3 | 1978749.88 | 1718210.88 | 2679861 | 3702414.75 | 2654657 | 3156787.75 | 1569595.5 | 2580358.25 | 2740472 | 2056244.5 | 12 | **0.950** |
| Q5KU39 | Vps41 | Vacuolar protein sorting-associated protein 41 homolog (Mus musculus) | 89.24 | 3.17 | 98602 | 2 | 385714.06 | 459648.31 | 458571.69 | 1130844.38 | 823295 | 553515.94 | 854347.62 | 494519.5 | 772878.81 | 922820.62 | 8 | **1.104** |
| Q5M7W1 | Txnip | Thioredoxin-interacting protein (Rattus norvegicus) | 45.24 | 3.05 | 44018 | 1 | 602051.12 | 738547.31 | 629491.5 | 878464.25 | 582950.25 | 781071 | 771473.38 | 249754.22 | 514702.12 | 497267.53 | 8 | **0.820** |
| Q5M823 | Nudcd2 | NudC domain-containing protein 2 (Rattus norvegicus) | 184.33 | 24.2 | 17674 | 3 | 10416389 | 7916261 | 8885842 | 6384751.5 | 6391956 | 8229008.5 | 9468760 | 6764389 | 6675771.5 | 6608915.5 | 12 | **0.944** |
| Q5M9F8 | Scyl1 | N-terminal kinase-like protein (Rattus norvegicus) | 49.2 | 1.73 | 89126 | 1 | 515725.47 | 562856.38 | 549713.25 | 229646.92 | 492512.72 | 368854.94 | 824102 | 564491.5 | 354322.97 | 409347.75 | 12 | **1.073** |
| Q5M9G3 | Caprin1 | Caprin-1 (Rattus norvegicus) | 841.77 | 20.08 | 78121 | 14 | 37501968 | 44529520 | 43860972 | 48948968 | 38877052 | 42752632 | 41139952 | 42505920 | 45316220 | 43212480 | 12 | **1.006** |
| Q5M9G7 | Diexf | Digestive organ expansion factor homolog (Rattus norvegicus) | 51.68 | 1.05 | 87825 | 1 | 435187.97 | 820041.56 | 895940.06 | 909761.62 | 966250.25 | 160360.75 | 1172783.88 | 809274.75 | 824627.94 | 829522.62 | 10 | **0.943** |
| Q5NC05 | Ttf2 | Transcription termination factor 2 (Mus musculus) | 53.21 | 1.49 | 125530 | 1 | 207047.05 | 163481.75 | 142185.42 | 0 | 0 | 175432.66 | 198528.17 | 203087.44 | 0 | 105777.59 | 12 | **1.332** |
| Q5PPH0 | Enoph1 | Enolase-phosphatase E1 (Rattus norvegicus) | 42.1 | 8.43 | 28875 | 1 | 0 | 407906.22 | 292925.69 | 286708.91 | 260047.73 | 666225.06 | 0 | 537463.62 | 607865.06 | 489135.53 | 5 | **1.844** |
| Q5PPL2 | Dcun1d5 | DCN1-like protein 5 (Rattus norvegicus) | 46 | 3.8 | 27568 | 1 | 0 | 388764.84 | 397272.22 | 0 | 0 | 331922.34 | 357308.44 | 302023.53 | 272753.5 | 242652.69 | 13 | **1.917** |
| Q5PQJ7 | Tbcel | Tubulin-specific chaperone cofactor E-like protein (Rattus norvegicus) | 72.81 | 5.9 | 48046 | 1 | 221009.69 | 194249.28 | 178872.34 | 0 | 0 | 339315.53 | 503671.03 | 485409.5 | 0 | 454614.12 | 6 | **3.001** |
| Q5PQL2 | Cnot9 | CCR4-NOT transcription complex subunit 9 (Rattus norvegicus) | 60.94 | 4.01 | 33601 | 1 | 926989.25 | 1052027.62 | 1119831.62 | 1064940.12 | 0 | 1130991.88 | 1227642.5 | 1224762.38 | 1081397 | 979571 | 4 | **1.356** |
| Q5PQL7 | Itm2c | Integral membrane protein 2C (Rattus norvegicus) | 61.28 | 8.18 | 30480 | 1 | 0 | 542820 | 0 | 0 | 0 | 0 | 671388.06 | 702843 | 551376.69 | 525553.62 | 12 | **4.516** |
| Q5PQM0 | Tmem168 | Transmembrane protein 168 (Rattus norvegicus) | 44.55 | 1.29 | 79772 | 1 | 288760.81 | 0 | 0 | 0 | 0 | 451905.56 | 0 | 373972.66 | 258292.64 | 242014.52 | 13 | **4.593** |
| Q5RJQ4 | Sirt2 | NAD-dependent protein deacetylase sirtuin-2 (Rattus norvegicus) | 130.6 | 10.57 | 39319 | 3 | 2806410.5 | 1659799.62 | 1377674.75 | 2135290 | 3303854.75 | 2324145.25 | 2091262.25 | 2796528.75 | 2924063.5 | 2577415 | 10 | **1.127** |
| Q5RJQ8 | Calhm2 | Calcium homeostasis modulator protein 2 (Rattus norvegicus) | 111.67 | 7.43 | 35931 | 2 | 269840.69 | 171684.84 | 525704.25 | 494565.19 | 431061.19 | 442461.88 | 59001.73 | 225456.92 | 544137.88 | 574606.75 | 11 | **0.975** |
| Q5RKH0 | Glyr1 | Putative oxidoreductase GLYR1 (Rattus norvegicus) | 132.37 | 6.52 | 60421 | 2 | 2758397.5 | 1871773 | 2100138 | 1979306.38 | 2554729.5 | 2714188.5 | 2216408.25 | 2137535 | 1922885.5 | 2157811.75 | 11 | **0.990** |
| Q5SGE0 | Lrpprc | Leucine-rich PPR motif-containing protein, mitochondrial (Rattus norvegicus) | 438.61 | 7.33 | 156653 | 9 | 7320142 | 12884452 | 11851657 | 11986494 | 7744228 | 9416882 | 10008411 | 9850205 | 10941008 | 10284152 | 10 | **0.975** |
| Q5SQX6 | Cyfip2 | Cytoplasmic FMR1-interacting protein 2 (Mus musculus) | 680.43 | 10.14 | 145659 | 13 | 13365725 | 18136080 | 16552657 | 19828516 | 11898332 | 14244789 | 14181742 | 16328094 | 13595185 | 13824414 | 10 | **0.905** |
| Q5SSI6 | Utp18 | U3 small nucleolar RNA-associated protein 18 homolog (Mus musculus) | 201.29 | 9.06 | 61218 | 3 | 2572680 | 1408449.25 | 2467554.75 | 1264652.5 | 2410919 | 2291576.25 | 2519546 | 1166769 | 2853438.75 | 2471343.75 | 10 | **1.116** |
| Q5SSL4 | Abr | Active breakpoint cluster region-related protein (Mus musculus) | 263.97 | 7.92 | 97667 | 5 | 3349216 | 3320120.5 | 3619804 | 2797682.5 | 2774822 | 4030968.25 | 3475745.25 | 3502494 | 3830673.25 | 4105693.5 | 2 | **1.194** |
| Q5SSW2 | Psme4 | Proteasome activator complex subunit 4 (Mus musculus) | 37.42 | 0.38 | 211197 | 1 | 241857.95 | 0 | 0 | 0 | 0 | 464560.31 | 0 | 513735.62 | 0 | 332852.53 | 16 | **5.421** |
| Q5SSZ5 | Tns3 | Tensin-3 (Mus musculus) | 43.48 | 0.69 | 155589 | 1 | 1399006.12 | 1026028.5 | 1245143.38 | 1137279 | 925110.75 | 1453623.88 | 1375967.5 | 1528753.5 | 1662789.88 | 1151612.88 | 3 | **1.251** |
| Q5SU73 | Coil | Coilin (Mus musculus) | 51.87 | 1.75 | 61917 | 1 | 627842.5 | 804655.12 | 642572.75 | 869772.94 | 686215.5 | 694207.31 | 521700.09 | 485915.09 | 0 | 404448.78 | 3 | **0.580** |
| Q5SUA5 | Myo1g | Unconventional myosin-Ig (Mus musculus) | 55.94 | 1.07 | 117227 | 1 | 554301.06 | 309710.66 | 786382.19 | 0 | 0 | 651989.06 | 1080138.12 | 680551.19 | 0 | 0 | 13 | **1.462** |
| Q5SUF2 | Luc7l3 | Luc7-like protein 3 (Mus musculus) | 396.35 | 18.06 | 51450 | 5 | 7519460.5 | 4396352 | 5195333 | 4234375.5 | 5621201.5 | 4677195.5 | 5462369 | 5020008.5 | 5318037 | 5490427 | 12 | **0.963** |
| Q5SUQ9 | Ctc1 | CST complex subunit CTC1 (Mus musculus) | 67.66 | 0.91 | 134028 | 1 | 309939.88 | 425486.19 | 400538.88 | 290810.16 | 379814.31 | 451855.69 | 256761.19 | 349551.56 | 407462.62 | 397429.91 | 11 | **1.031** |
| Q5SUR0 | Pfas | Phosphoribosylformylglycinamidine synthase (Mus musculus) | 929.33 | 19.3 | 144629 | 17 | 24770432 | 22050012 | 23613832 | 22757300 | 26476894 | 28162140 | 26525380 | 23179380 | 23547662 | 23178618 | 9 | **1.041** |
| Q5SVQ0 | Kat7 | Histone acetyltransferase KAT7 (Mus musculus) | 221.84 | 8.48 | 70641 | 3 | 0 | 1783864.38 | 1651619.12 | 479712.25 | 250216.45 | 1362998.25 | 1794647.5 | 850649.81 | 791168.94 | 651420.5 | 8 | **1.309** |
| Q5SW19 | Cluh | Clustered mitochondria protein homolog (Mus musculus) | 685.21 | 11.33 | 148068 | 12 | 11663706 | 10318907 | 10645048 | 8711774 | 7788800 | 12065797 | 10644583 | 12503023 | 10580229 | 11591200 | 5 | **1.168** |
| Q5SWD9 | Tsr1 | Pre-rRNA-processing protein TSR1 homolog (Mus musculus) | 227.51 | 7.85 | 92105 | 5 | 3097609.5 | 2478704.25 | 2971127.5 | 2484239.5 | 3032783.5 | 5937110 | 4531672.5 | 3922258.75 | 5652917 | 1760468.38 | 5 | **1.550** |
| Q5U1X1 | Rexo2 | Oligoribonuclease, mitochondrial (Rattus norvegicus) | 194.54 | 15.19 | 26752 | 3 | 4393494 | 6141397 | 6697391 | 3730915.5 | 3869754.25 | 6784821.5 | 5545643 | 7485478.5 | 6965166 | 7763162 | 2 | **1.391** |
| Q5U2M6 | Dcaf8 | DDB1- and CUL4-associated factor 8 (Rattus norvegicus) | 49.39 | 2.71 | 66156 | 1 | 700886.44 | 0 | 330618.66 | 684035.75 | 803460.5 | 380734.34 | 710485.06 | 255899.2 | 436728 | 721919.75 | 12 | **0.995** |
| Q5U2Q3 | | Ester hydrolase C11orf54 homolog (Rattus norvegicus) | 109.61 | 8.89 | 34993 | 2 | 950291.75 | 2581437 | 1574954.25 | 2347289.5 | 2487450.75 | 901992.81 | 0 | 932614.31 | 1231680.88 | 1226933.5 | 2 | **0.432** |
| Q5U2Q7 | Etf1 | Eukaryotic peptide chain release factor subunit 1 (Rattus norvegicus) | 1162.86 | 56.75 | 49031 | 18 | 25851356 | 34863320 | 36024000 | 37073804 | 32158706 | 42606140 | 34616796 | 42064528 | 37698436 | 41445392 | 3 | **1.196** |
| Q5U2R0 | Mat2b | Methionine adenosyltransferase 2 subunit beta (Rattus norvegicus) | 192.02 | 18.26 | 37375 | 4 | 8081794.5 | 6926177.5 | 6436786.5 | 6978748.5 | 5726394 | 8487354 | 6876783 | 7364993 | 6661739.5 | 6580509.5 | 10 | **1.053** |
| Q5U2U7 | Rnmt | mRNA cap guanine-N7 methyltransferase (Rattus norvegicus) | 41.47 | 4.12 | 52817 | 1 | 0 | 305428.22 | 0 | 345899 | 226608.58 | 377252.44 | 419656.09 | 328204.5 | 525377 | 0 | 7 | **1.880** |
| Q5U2V8 | Emc3 | ER membrane protein complex subunit 3 (Rattus norvegicus) | 91.96 | 9.58 | 29980 | 2 | 721420.62 | 860548.75 | 1331321.38 | 0 | 662093.31 | 0 | 810156.25 | 1018192.31 | 909311.38 | 946382.5 | 10 | **1.030** |

| | | | | | | | | | | | | | | | | | | |
|---|---|---|---|---|---|---|---|---|---|---|---|---|---|---|---|---|---|---|
| Q5U2Y1 | Gtf2i | General transcription factor II-I (Rattus norvegicus) | 49.72 | 1.23 | 110215 | 1 | 0 | 0 | 0 | 0 | 0 | 1331510.25 | 0 | 783496.31 | 992866.06 | 707394.69 | 15 | **#DIV/0 !** |
| Q5U2Y6 | Tfip11 | Tuftelin-interacting protein 11 (Rattus norvegicus) | 52.09 | 1.55 | 96152 | 1 | 0 | 543024.81 | 748868.5 | 670592.69 | 678925.56 | 657017.56 | 703297.06 | 870319.88 | 919619.5 | 836063.06 | 4 | **1.509** |
| Q5U316 | Rab35 | Ras-related protein Rab-35 (Rattus norvegicus) | 355.66 | 28.86 | 23025 | 5 | 3709613.5 | 4577451 | 5032152 | 6125284 | 4589026.5 | 3582682.5 | 3033850 | 4060605.5 | 5583753 | 4813462 | 8 | **0.877** |
| Q5U3K5 | Rabl6 | Rab-like protein 6 (Mus musculus) | 46.21 | 1.52 | 79831 | 1 | 396914.09 | 604230 | 660527.12 | 517060.38 | 593919.5 | 562741.12 | 620929.75 | 553521.44 | 586306.06 | 606012.25 | 11 | **1.057** |
| Q5U4D9 | Thoc6 | THO complex subunit 6 homolog (Mus musculus) | 117.54 | 10.56 | 37315 | 2 | 0 | 349316 | 543202.62 | 579133.44 | 724458.75 | 836785.62 | 781211.94 | 846243.75 | 912779.69 | 1182800.38 | 0 | **2.076** |
| Q5XFV8 | Cox14 | Cytochrome c oxidase assembly protein COX14 (Rattus norvegicus) | 39.73 | 14.04 | 6361 | 1 | 1963666.38 | 1017849 | 2146204.25 | 1893694.62 | 0 | 1923366.88 | 2671845 | 1754711.88 | 1765790.12 | 1868271.12 | 11 | **1.422** |
| Q5XFW8 | Sec13 | Protein SEC13 homolog (Rattus norvegicus) | 350.58 | 23.91 | 35548 | 5 | 11320647 | 13781997 | 11758546 | 10726740 | 9249950 | 12773796 | 11073979 | 12211736 | 11880495 | 13326334 | 7 | **1.078** |
| Q5XI13 | Grwd1 | Glutamate-rich WD repeat-containing protein 1 (Rattus norvegicus) | 141.76 | 5.62 | 49169 | 2 | 1434757.75 | 2586458.75 | 2568630 | 2164185 | 2919474.5 | 1752495 | 1827623.25 | 2775969 | 2557062 | 2286236.5 | 10 | **0.959** |
| Q5XI29 | Cpsf7 | Cleavage and polyadenylation specificity factor subunit 7 (Rattus norvegicus) | 84.49 | 4.11 | 51073 | 2 | 1511603.88 | 1963218 | 2224766.75 | 1713043.62 | 1854402 | 2228713 | 1433337.5 | 1450723.25 | 1926326.75 | 1848293 | 10 | **0.959** |
| Q5XI32 | Capzb | F-actin-capping protein subunit beta (Rattus norvegicus) | 935.45 | 58.46 | 30629 | 15 | 92935336 | 75285304 | 74209344 | 90150464 | 90580280 | 79034752 | 75479872 | 80193112 | 87583632 | 87739616 | 10 | **0.969** |
| Q5XI41 | Tram1 | Translocating chain-associated membrane protein 1 (Rattus norvegicus) | 129.71 | 5.61 | 43030 | 2 | 4011430.5 | 6169713 | 6844108 | 6244782 | 5099435.5 | 8707677 | 6412597 | 9114823 | 8448768 | 8784260 | 1 | **1.462** |
| Q5XI68 | Dr1 | Protein Dr1 (Rattus norvegicus) | 113.77 | 21.02 | 19430 | 2 | 1606009.12 | 0 | 919898.38 | 964893.31 | 1342570.88 | 1269903.88 | 1174181 | 1552409.88 | 1147275.62 | 1363664.88 | 8 | **1.346** |
| Q5XI72 | Eif4h | Eukaryotic translation initiation factor 4H (Rattus norvegicus) | 456.2 | 43.15 | 27324 | 7 | 21142800 | 27006452 | 29907240 | 24260752 | 30059556 | 30186788 | 28861636 | 28383218 | 26165992 | 32676704 | 7 | **1.105** |
| Q5XI73 | Arhgdia | Rho GDP-dissociation inhibitor 1 (Rattus norvegicus) | 792.84 | 77.45 | 23407 | 12 | 163098928 | 169422880 | 170488064 | 147493616 | 163370000 | 172248416 | 164883808 | 141981504 | 167749616 | 163328896 | 12 | **0.995** |
| Q5XIA1 | Ncln | Nicalin (Rattus norvegicus) | 193.84 | 5.86 | 62992 | 3 | 2151861.25 | 2287885 | 2645826.5 | 2304701 | 2396781.5 | 2263349.75 | 1904952.75 | 2412511.5 | 2489234.5 | 2156333.5 | 10 | **0.952** |
| Q5XIA4 | Cnot10 | CCR4-NOT transcription complex subunit 10 (Rattus norvegicus) | 91.82 | 1.61 | 81819 | 1 | 0 | 0 | 0 | 0 | 0 | 519746.88 | 682184.44 | 694753.06 | 442392.69 | 481712.62 | 10 | **#DIV/0 !** |
| Q5XIB1 | Tssc4 | Protein TSSC4 (Rattus norvegicus) | 41.46 | 3.15 | 33939 | 1 | 0 | 936112.94 | 668915.62 | 378701.78 | 1030592.38 | 908779.31 | 1032230.38 | 650236.75 | 861790.19 | 772565.5 | 9 | **1.402** |
| Q5XIC1 | Gmppa | Mannose-1-phosphate guanyltransferase alpha (Rattus norvegicus) | 136 | 10 | 46191 | 3 | 3349314 | 3003966.5 | 3072475.75 | 2673940.75 | 2395772 | 3208960.75 | 2601440.5 | 2537893 | 2710721.25 | 2460633.5 | 9 | **0.933** |
| Q5XIG4 | Ociad1 | OCIA domain-containing protein 1 (Rattus norvegicus) | 128.55 | 12.55 | 27659 | 2 | 679029.69 | 0 | 491221.84 | 0 | 297963.91 | 380109 | 566123.25 | 555079.25 | 430684.06 | 543132.56 | 8 | **1.686** |
| Q5XIG8 | Strap | Serine-threonine kinase receptor-associated protein (Rattus norvegicus) | 871.96 | 58 | 38456 | 14 | 36068916 | 39307032 | 39389216 | 41319880 | 40470708 | 37392080 | 35905520 | 37197752 | 40479596 | 41002372 | 10 | **0.977** |
| Q5XII9 | Mtfr1l | Mitochondrial fission regulator 1-like (Rattus norvegicus) | 45.34 | 5.88 | 31730 | 1 | 509495.31 | 554991.56 | 398020 | 0 | 0 | 446797.22 | 556061.38 | 683688.12 | 0 | 669943.81 | 8 | **1.611** |
| Q5XIK2 | Tmx2 | Thioredoxin-related transmembrane protein 2 (Rattus norvegicus) | 47.46 | 5.08 | 33866 | 1 | 199439.73 | 149146.69 | 130475.38 | 0 | 0 | 141908.67 | 175589.98 | 143247.98 | 140705.31 | 0 | 13 | **1.255** |
| Q5XIK8 | Ctdspl2 | CTD small phosphatase-like protein 2 (Rattus norvegicus) | 84.12 | 4.3 | 52783 | 2 | 1208769.25 | 1531539.38 | 1536944 | 1798418.25 | 988524.31 | 1579623.38 | 1627331.12 | 1020560 | 999835.62 | 1006276.5 | 11 | **0.882** |
| Q5XIN6 | Letm1 | Mitochondrial proton/calcium exchanger protein (Rattus norvegicus) | 50.42 | 1.89 | 83060 | 1 | 978937.06 | 1007472.31 | 935726.81 | 987526.25 | 0 | 1003973.38 | 890987.38 | 1062750.5 | 780675.25 | 818258.5 | 12 | **1.165** |
| Q5XIP1 | Pelo | Protein pelota homolog (Rattus norvegicus) | 71.89 | 4.68 | 43407 | 1 | 0 | 224693.91 | 0 | 290285.72 | 0 | 0 | 0 | 0 | 234332.2 | 239461.95 | 20 | **0.920** |
| Q5XIS9 | Prkd2 | Serine/threonine-protein kinase D2 (Rattus norvegicus) | 81.12 | 1.26 | 96497 | 1 | 0 | 287344.25 | 375569.72 | 234031.88 | 309881.44 | 0 | 280402.78 | 434820.34 | 312124.03 | 327273.56 | 10 | **1.122** |
| Q5XIU9 | Pgrmc2 | Membrane-associated progesterone receptor component 2 (Rattus norvegicus) | 58.33 | 6.45 | 23403 | 1 | 2447831.5 | 1316805.88 | 653150.12 | 0 | 0 | 2077234.38 | 2493819.75 | 1925278.75 | 2426638 | 1779440.62 | 5 | **2.423** |
| Q5XJY5 | Arcn1 | Coatomer subunit delta (Mus musculus) | 614.43 | 25.64 | 57229 | 12 | 25484876 | 27029068 | 23675650 | 27513160 | 24918984 | 22734278 | 21640090 | 26237080 | 27492982 | 27461936 | 11 | **0.976** |
| Q60520 | Sin3a | Paired amphipathic helix protein Sin3a (Mus musculus) | 112.38 | 1.96 | 145088 | 3 | 1471516.12 | 1557822 | 2138384 | 1508534.62 | 547404.12 | 1488570.25 | 0 | 2480150 | 2392834 | 2503167.5 | 8 | **1.227** |
| Q60591 | Nfatc2 | Nuclear factor of activated T-cells, cytoplasmic 2 (Mus musculus) | 122.21 | 5.39 | 100020 | 3 | 2248925.25 | 2150939.5 | 3245067.75 | 1991817.5 | 2718638.5 | 1129671.75 | 2304358.5 | 3204669.5 | 1705050.12 | 3468517.25 | 12 | **0.956** |
| Q60597 | Ogdh | 2-oxoglutarate dehydrogenase, mitochondrial (Mus musculus) | 833.66 | 23.46 | 116449 | 15 | 17757348 | 18024154 | 19733116 | 20889906 | 16472152 | 19513116 | 16981584 | 17879832 | 18341098 | 17843414 | 11 | **0.975** |
| Q60605 | Myl6 | Myosin light polypeptide 6 (Mus musculus) | 880.76 | 66.23 | 16930 | 13 | 136102688 | 129680048 | 136121552 | 143770608 | 149228064 | 121681680 | 119432856 | 153987264 | 162471136 | 171375264 | 10 | **1.049** |
| Q60631 | Grb2 | Growth factor receptor-bound protein 2 (Mus musculus) | 331.95 | 40.55 | 25238 | 7 | 15857944 | 13937070 | 16213742 | 13799542 | 12965350 | 18609592 | 15340028 | 19158688 | 18799744 | 19824746 | 2 | **1.261** |
| Q60634 | Flot2 | Flotillin-2 (Mus musculus) | 50.34 | 2.57 | 47038 | 1 | 0 | 671555.44 | 943507.62 | 642792.19 | 559660.38 | 601901.81 | 467587.47 | 786725.06 | 637410.5 | 1095420.62 | 11 | **1.274** |
| Q60649 | Clpb | Caseinolytic peptidase B protein homolog (Mus musculus) | 131.1 | 7.53 | 76004 | 3 | 0 | 154512.48 | 186423.11 | 1546862.88 | 0 | 0 | 1108477.75 | 101220.05 | 120450.31 | 95790.95 | 12 | **0.755** |
| Q60664 | Lrmp | Lymphoid-restricted membrane protein (Mus musculus) | 125.35 | 6.31 | 59588 | 3 | 4583877.5 | 2981663 | 4211195.5 | 2474549 | 3970251.25 | 4449480 | 4667361 | 4996272 | 4595842 | 4659045 | 1 | **1.282** |
| Q60668 | Hnrnpd | Heterogeneous nuclear ribonucleoprotein D0 (Mus musculus) | 273.16 | 16.9 | 38354 | 6 | 14585797 | 9919449 | 7003118 | 8595690 | 10174079 | 5524847.5 | 5265991 | 8404422 | 10207964 | 9142980 | 7 | **0.767** |
| Q60676 | Ppp5c | Serine/threonine-protein phosphatase 5 (Mus musculus) | 168.15 | 8.62 | 56877 | 3 | 2849329 | 3588032.5 | 4658247.5 | 2618619.5 | 1314270.75 | 3967146.5 | 3982677 | 1729192.25 | 3813954.5 | 2394023.25 | 11 | **1.057** |
| Q60680 | Chuk | Inhibitor of nuclear factor kappa-B kinase subunit alpha (Mus musculus) | 38.93 | 1.21 | 84729 | 1 | 643666.62 | 656465.62 | 736284.25 | 976410.38 | 496259.84 | 539055.81 | 0 | 508800.16 | 470718.75 | 539464.5 | 3 | **0.586** |
| Q60692 | Psmb6 | Proteasome subunit beta type-6 (Mus musculus) | 91.43 | 21.01 | 25379 | 2 | 493344.69 | 2869109.75 | 2257783.75 | 4129616.5 | 1981195.5 | 3241091 | 0 | 2660980 | 1008372 | 2183456.5 | 10 | **0.775** |
| Q60710 | Samhd1 | Deoxynucleoside triphosphate triphosphohydrolase SAMHD1 (Mus musculus) | 575.25 | 25.53 | 75893 | 12 | 14663633 | 15675412 | 17425166 | 17127408 | 16638098 | 17817376 | 15816074 | 21246180 | 16423023 | 16878340 | 8 | **1.082** |
| Q60716 | P4ha2 | Prolyl 4-hydroxylase subunit alpha-2 (Mus musculus) | 178.34 | 5.77 | 61002 | 2 | 48570.74 | 852836.62 | 1026416.75 | 1225657 | 1018634.5 | 959494.5 | 564928.44 | 1058338.25 | 1339925.5 | 1116042.38 | 9 | **1.208** |
| Q60749 | Khdrbs1 | KH domain-containing, RNA-binding, signal transduction-associated protein 1 (Mus muscu | 107.95 | 4.51 | 48371 | 2 | 9964621 | 5877386.5 | 6415137 | 6219882.5 | 6725410 | 8611029 | 8953002 | 9489886 | 8568263 | 7822416 | 5 | **1.234** |
| Q60766 | Irgm1 | Immunity-related GTPase family M protein 1 (Mus musculus) | 157.52 | 7.82 | 46552 | 3 | 632534.38 | 2410812.5 | 2614739.5 | 1948272.25 | 915818.44 | 1858175 | 2413985.5 | 1655436 | 960411.5 | 1616895.12 | 12 | **0.998** |
| Q60770 | Stxbp3 | Syntaxin-binding protein 3 (Mus musculus) | 80.35 | 2.87 | 67942 | 2 | 946200.44 | 1471518.12 | 1253451.75 | 746183.12 | 1151556.75 | 1130118.88 | 0 | 1423488.5 | 1690011.38 | 1521780.75 | 9 | **1.035** |
| Q60787 | Lcp2 | Lymphocyte cytosolic protein 2 (Mus musculus) | 61.23 | 3 | 60238 | 1 | 477509.38 | 757246.06 | 959794.44 | 634866.12 | 730329.25 | 981519.88 | 860115.31 | 864167.31 | 970856.25 | 1118908 | 2 | **1.347** |
| Q60809 | Cnot7 | CCR4-NOT transcription complex subunit 7 (Mus musculus) | 125.42 | 10.18 | 32718 | 2 | 0 | 1105625.25 | 1222019.12 | 912212.75 | 644952.94 | 736569.06 | 939646.19 | 853402.94 | 609706.81 | 826623.75 | 10 | **1.021** |
| Q60848 | Hells | Lymphocyte-specific helicase (Mus musculus) | 417.44 | 13.52 | 95126 | 8 | 6993636 | 4651362 | 5470245 | 5554529.5 | 3710439.5 | 6593816.5 | 4897779 | 7321116 | 6839119 | 7332374 | 5 | **1.250** |
| Q60854 | Serpinb6 | Serpin B6 (Mus musculus) 6 | 819.46 | 38.62 | 42599 | 14 | 51638212 | 51021092 | 51928304 | 49325096 | 45365480 | 50159488 | 48389708 | 46983004 | 52134768 | 53112756 | 11 | **1.006** |
| Q60862 | Orc2 | Origin recognition complex subunit 2 (Mus musculus) | 40.3 | 4.17 | 65893 | 1 | 167790.19 | 154276.64 | 134410.86 | 0 | 0 | 254899.08 | 0 | 181403.05 | 0 | 252412.12 | 11 | **1.509** |
| Q60864 | Stip1 | Stress-induced-phosphoprotein 1 (Mus musculus) | 1457.11 | 49.54 | 62582 | 26 | 111244288 | 92101776 | 95306784 | 87078864 | 124789656 | 97233184 | 82086912 | 94708664 | 98200032 | 97843256 | 11 | **0.921** |
| Q60866 | Pter | Phosphotriesterase-related protein (Mus musculus) | 98.26 | 10.03 | 39218 | 2 | 728451.75 | 727309.19 | 751607 | 1133702.75 | 2609426.25 | 1947433.62 | 1175061.75 | 1789463.75 | 1996451.5 | 2088501 | 5 | **1.512** |
| Q60875 | Arhgef2 | Rho guanine nucleotide exchange factor 2 (Mus musculus) | 748.72 | 18.68 | 111974 | 14 | 21338886 | 21217498 | 23995772 | 24383498 | 23428208 | 28929866 | 26329734 | 26858584 | 26179222 | 26262888 | 0 | **1.177** |
| Q60902 | Eps15l1 | Epidermal growth factor receptor substrate 15-like 1 (Mus musculus) | 252.7 | 7.17 | 99309 | 5 | 3266550 | 3902578.5 | 3807494 | 5354728 | 4824884 | 4571113 | 3641064 | 4866980.5 | 5300564.5 | 5226682.5 | 9 | **1.116** |
| Q60930 | Vdac2 | Voltage-dependent anion-selective channel protein 2 (Mus musculus) | 973.98 | 63.05 | 31733 | 15 | 155715024 | 179080688 | 222324704 | 247476976 | 217631616 | 196824960 | 217166800 | 235558688 | 234251344 | 235009856 | 9 | **1.094** |
| Q60931 | Vdac3 | Voltage-dependent anion-selective channel protein 3 (Mus musculus) | 606.01 | 35.69 | 30753 | 9 | 34453828 | 56728752 | 67925712 | 47648536 | 44167368 | 61501576 | 61291136 | 72060728 | 66012416 | 70310392 | 3 | **1.320** |
| Q60953 | Pml | Protein PML (Mus musculus) | 249.9 | 7.57 | 98242 | 5 | 320016.38 | 908674.62 | 1464902.25 | 949151.12 | 933650.25 | 933378 | 1227622 | 756342.88 | 875582.12 | 844598.19 | 9 | **1.013** |
| Q60967 | Papss1 | Bifunctional 3'-phosphoadenosine 5'-phosphosulfate synthase 1 (Mus musculus) | 129.26 | 8.49 | 70794 | 3 | 2681509.25 | 3083569.25 | 2198464.25 | 3391513.5 | 2468608.75 | 3199388.5 | 2486615.75 | 2157639 | 3729142 | 2768507.25 | 11 | **1.037** |
| Q60972 | Rbbp4 | Histone-binding protein RBBP4 (Mus musculus) | 468.01 | 25.18 | 47656 | 8 | 26488240 | 37374772 | 32792416 | 41779560 | 34185848 | 29010320 | 21711834 | 22645210 | 22523602 | 23709420 | 1 | **0.693** |
| Q60973 | Rbbp7 | Histone-binding protein RBBP7 (Mus musculus) | 343.25 | 14.12 | 47790 | 6 | 25712494 | 35155528 | 29611268 | 38458200 | 31410146 | 26492002 | 20377712 | 19717172 | 19272196 | 19492622 | 1 | **0.657** |
| Q60996 | Ppp2r5c | Serine/threonine-protein phosphatase 2A 56 kDa regulatory subunit gamma isoform (Mus n | 138.16 | 7.82 | 60824 | 3 | 4327191 | 3008082.5 | 3598678.75 | 2315386.5 | 4081819.5 | 3739965 | 3177883.75 | 3930318 | 2652505 | 2952182.75 | 10 | **0.949** |
| Q61024 | Asns | Asparagine synthetase [glutamine-hydrolyzing] (Mus musculus) | 911.97 | 30.3 | 64283 | 15 | 28678540 | 40583492 | 38508576 | 41846352 | 27654506 | 29099002 | 39311800 | 35599488 | 33592928 | 35596168 | 11 | **0.977** |
| Q61029 | Tmpo | Lamina-associated polypeptide 2, isoforms beta/delta/epsilon/gamma (Mus musculus) | 525.52 | 27.88 | 50373 | 9 | 20223160 | 16489923 | 14253640 | 19135690 | 21567270 | 14213561 | 18731324 | 14553014 | 19425332 | 15181426 | 7 | **0.896** |
| Q61033 | Tmpo | Lamina-associated polypeptide 2, isoforms alpha/zeta (Mus musculus) | 359.03 | 14.43 | 75168 | 7 | 11979311 | 6019266.5 | 5608220 | 3827734 | 12257405 | 9864797 | 11764940 | 7096628 | 11217026 | 7504728 | 10 | **1.195** |
| Q61035 | Hars | Histidine--tRNA ligase, cytoplasmic (Mus musculus) | 881.4 | 36.35 | 57432 | 16 | 30019590 | 30136350 | 33364684 | 31995350 | 29017518 | 32802780 | 34625296 | 36257340 | 36789048 | 38192296 | 1 | **1.156** |
| Q61074 | Ppm1g | Protein phosphatase 1G (Mus musculus) | 459.19 | 19.56 | 58728 | 8 | 12622728 | 18625848 | 24800592 | 17647108 | 20320544 | 17337768 | 23286128 | 20824862 | 19308376 | 26726186 | 8 | **1.143** |
| Q61081 | Cdc37 | Hsp90 co-chaperone Cdc37 (Mus musculus) | 590.54 | 28.5 | 44593 | 10 | 14420896 | 17617568 | 10979996 | 13776163 | 12267768 | 15115948 | 17121686 | 15654554 | 13613985 | 14631401 | 7 | **1.102** |
| Q61093 | Cybb | Cytochrome b-245 heavy chain (Mus musculus) | 184.24 | 8.77 | 65305 | 4 | 2681147.25 | 2157110 | 2962913 | 1128218 | 2490853 | 4124419.25 | 2980819 | 3490380.25 | 3515873.5 | 2854190 | 1 | **1.486** |
| Q61102 | Abcb7 | ATP-binding cassette sub-family B member 7, mitochondrial (Mus musculus) | 416.64 | 14.63 | 82581 | 7 | 2612623.75 | 2556418.75 | 2363881 | 1071030 | 2719353.25 | 2548570.5 | 2589569.75 | 5176230.5 | 4810160 | 4080775.5 | 5 | **1.696** |
| Q61103 | Dpf2 | Zinc finger protein ubi-d4 (Mus musculus) | 108.25 | 6.39 | 44230 | 2 | 2538021 | 1649279.25 | 1566193.75 | 1702666.12 | 0 | 2031056.38 | 0 | 1890369.38 | 1735043 | 1719932.75 | 9 | **0.989** |
| Q61107 | Gbp4 | Guanylate-binding protein 4 (Mus musculus) | 117.56 | 4.19 | 70801 | 2 | 396650.97 | 1083443.75 | 1130162.88 | 1441397.62 | 420899.78 | 692443.06 | 709033.56 | 732046.12 | 915409.88 | 670596.75 | 10 | **0.832** |
| Q61151 | Ppp2r5e | Serine/threonine-protein phosphatase 2A 56 kDa regulatory subunit epsilon isoform (Mus n | 199.21 | 11.56 | 54713 | 4 | 774782.88 | 3825615.25 | 2724508.5 | 1479514.75 | 1234819.75 | 1465449.88 | 527415.75 | 1695779.5 | 5913057 | 1810299.75 | 12 | **1.137** |
| Q61160 | Fadd | FAS-associated death domain protein (Mus musculus) | 45.78 | 8.29 | 22960 | 1 | 413449.19 | 333315.66 | 161261.59 | 122633.84 | 54750.42 | 349268.59 | 339934 | 271052.25 | 0 | 114418.37 | 12 | **0.990** |
| Q61164 | Ctcf | Transcriptional repressor CTCF (Mus musculus) | 75.99 | 2.45 | 83745 | 1 | 210274.72 | 162171.64 | 252968.56 | 231877.53 | 218771.02 | 225396.09 | 271423.41 | 231254.61 | 240141.89 | 0 | 10 | **0.900** |
| Q61166 | Mapre1 | Microtubule-associated protein RP/EB family member 1 (Mus musculus) | 638.65 | 45.15 | 30016 | 10 | 28485444 | 18879120 | 23396690 | 20569862 | 23026556 | 25589842 | 24943046 | 28074374 | 32680492 | 29458130 | 3 | **1.231** |
| Q61171 | Prdx2 | Peroxiredoxin-2 (Mus musculus) | 627.78 | 53.03 | 21779 | 10 | 116389776 | 115335776 | 110145408 | 100419088 | 101842728 | 119761528 | 88581960 | 109005320 | 106650080 | 109383560 | 11 | **0.980** |
| Q61183 | Papola | Poly(A) polymerase alpha (Mus musculus) | 90.17 | 3.38 | 82309 | 2 | 356947.56 | 0 | 720790.94 | 1139200.38 | 1074759.75 | 601371.62 | 0 | 0 | 485493.44 | 733839.25 | 10 | **0.553** |
| Q61187 | Tsg101 | Tumor susceptibility gene 101 protein (Mus musculus) | 127.63 | 9.72 | 44124 | 3 | 5242700 | 4060357 | 3158149 | 3095302 | 2356114.75 | 4758333.5 | 4944337 | 5115152 | 5078862 | 4766921 | 5 | **1.377** |
| Q61191 | Hcfc1 | Host cell factor 1 (Mus musculus) | 405.11 | 6.11 | 210437 | 7 | 7909553 | 5986460 | 6475377.5 | 3335270.5 | 4193311.5 | 7449761 | 7026291.5 | 4689894 | 6307393 | 6175874.5 | 9 | **1.134** |
| Q61194 | Pik3c2a | Phosphatidylinositol 4-phosphate 3-kinase C2 domain-containing subunit alpha (Mus muscu | 101.82 | 1.3 | 190758 | 2 | 859592.44 | 883436.38 | 919868.62 | 890543.62 | 890959.44 | 1081119.88 | 476153.53 | 966508.94 | 919903.94 | 896177.62 | 6 | **0.976** |
| Q61207 | Psap | Prosaposin (Mus musculus) | 351.59 | 10.23 | 61422 | 6 | 65557404 | 40204336 | 35689752 | 31093736 | 45800776 | 34817480 | 36369880 | 25759958 | 32140492 | 32619848 | 5 | **0.741** |
| Q61210 | Arhgef1 | Rho guanine nucleotide exchange factor 1 (Mus musculus) | 640.29 | 17.5 | 102805 | 13 | 15524944 | 12125941 | 10557637 | 11987524 | 10924321 | 11629742 | 12516276 | 11990378 | 12826930 | 11919506 | 10 | **0.996** |
| Q61233 | Lcp1 | Plastin-2 (Mus musculus) | 4443.479 | 87.72 | 70149 | 65 | 1236071808 | 1415134208 | 1613736576 | 1466943360 | 1650113792 | 1815926528 | 1503529728 | 1636105216 | 1628646912 | 1570863744 | 6 | **1.105** |
| Q61235 | Sntb2 | Beta-2-syntrophin (Mus musculus) | 69.82 | 1.73 | 56382 | 1 | 111258.27 | 502471.28 | 542167.81 | 0 | 82959.95 | 665255.25 | 792931.62 | 716638.25 | 758532.62 | 477693.94 | 2 | **2.753** |
| Q61239 | Fnta | Protein farnesyltransferase/geranylgeranyltransferase type-1 subunit alpha (Mus musculus) | 122.44 | 8.75 | 44013 | 3 | 0 | 800374 | 1304872.38 | 1071770.62 | 3200223.5 | 1955546.5 | 640219.12 | 1629811.5 | 1920254.25 | 2558767.5 | 8 | **1.365** |
| Q61263 | Soat1 | Sterol O-acyltransferase 1 (Mus musculus) | 240.35 | 10.56 | 63799 | 5 | 4406222 | 2973206 | 3564616.5 | 1800673.25 | 4428889 | 7085524 | 3469766.5 | 6436634 | 6384946.5 | 5645380.5 | 3 | **1.690** |
| Q61316 | Hspa4 | Heat shock 70 kDa protein 4 (Mus musculus) | 2621.66 | 59.22 | 94133 | 40 | 189859520 | 159784288 | 188259280 | 169531040 | 173828592 | 187591536 | 179928592 | 192491920 | 188148064 | 195329360 | 6 | **1.071** |
| Q61425 | Hadh | Hydroxyacyl-coenzyme A dehydrogenase, mitochondrial (Mus musculus) | 220.4 | 25.8 | 34464 | 5 | 6486097.5 | 8079990 | 3730738.5 | 10431408 | 6741539 | 6294982 | 7242530.5 | 6739521.5 | 6110906.5 | 6147306 | 8 | **0.917** |

| | | | | | | | | | | | | | | | | | |
|---|---|---|---|---|---|---|---|---|---|---|---|---|---|---|---|---|---|
| Q61462 | Cyba | Cytochrome b-245 light chain (Mus musculus) | 42.77 | 7.29 | 20748 | 1 | 574296.38 | 187052.45 | 324114.12 | 0 | 0 | 352973.75 | 378699.06 | 199453.48 | 0 | 0 | 15 | **0.858** |
| Q61490 | Alcam | CD166 antigen (Mus musculus) | 109.43 | 5.15 | 65092 | 2 | 3428168.5 | 2531297.5 | 1848106.38 | 3190617.25 | 2600547 | 2260435 | 2258019.25 | 1974147.88 | 2253602.25 | 1732283 | 4 | **0.771** |
| Q61510 | Trim25 | E3 ubiquitin/ISG15 ligase TRIM25 (Mus musculus) | 85.53 | 3.15 | 71726 | 2 | 1222550.75 | 1910856.5 | 2491087 | 1537731.25 | 1150712.25 | 1651976 | 1065712.75 | 1690212.25 | 2068651.5 | 2373305 | 11 | **1.065** |
| Q61543 | Glg1 | Golgi apparatus protein 1 (Mus musculus) | 657.57 | 10.04 | 133734 | 13 | 15104117 | 12010670 | 11713912 | 10917595 | 15872236 | 13177535 | 13358381 | 12657653 | 15952874 | 12305872 | 8 | **1.028** |
| Q61545 | Ewsr1 | RNA-binding protein EWS (Mus musculus) | 312.6 | 12.21 | 68462 | 6 | 12509667 | 10482950 | 12096581 | 7080004 | 12903158 | 13690740 | 8527290 | 13184370 | 8917994 | 14439372 | 8 | **1.067** |
| Q61584 | Fxr1 | Fragile X mental retardation syndrome-related protein 1 (Mus musculus) | 905.4 | 27.33 | 76222 | 14 | 29676034 | 30762316 | 30969514 | 42150328 | 40482008 | 37354116 | 31311810 | 36871668 | 42813336 | 42315296 | 6 | **1.096** |
| Q61595 | Ktn1 | Kinectin (Mus musculus) | 78.13 | 1.21 | 152592 | 1 | 118467.96 | 430717.44 | 408626.66 | 604136.38 | 982610.19 | 411644.78 | 570766 | 483316.78 | 742061.56 | 951072.62 | 9 | **1.241** |
| Q61599 | Arhgdib | Rho GDP-dissociation inhibitor 2 (Mus musculus) | 845.35 | 76.5 | 22851 | 15 | 88344248 | 194720528 | 205601184 | 167074848 | 170601696 | 161734688 | 147351744 | 159433456 | 150949040 | 150807680 | 5 | **0.932** |
| Q61655 | Ddx19a | ATP-dependent RNA helicase DDX19A (Mus musculus) | 519 | 26.15 | 53933 | 10 | 18245448 | 16758974 | 18663884 | 14861772 | 17758860 | 18671816 | 15520011 | 20431184 | 20418926 | 21634402 | 4 | **1.120** |
| Q61656 | Ddx5 | Probable ATP-dependent RNA helicase DDX5 (Mus musculus) | 1728.53 | 48.86 | 69290 | 28 | 258983136 | 242068832 | 243039984 | 231105248 | 230273472 | 220576752 | 202354608 | 239192128 | 245806112 | 240279600 | 8 | **0.952** |
| Q61686 | Cbx5 | Chromobox protein homolog 5 (Mus musculus) | 259.64 | 29.84 | 22186 | 5 | 11818309 | 8935919 | 7465371 | 11407333 | 5578252 | 8786149 | 7636152.5 | 5287452.5 | 8105733 | 7070972 | 7 | **0.816** |
| Q61687 | Atrx | Transcriptional regulator ATRX (Mus musculus) | 55.43 | 0.36 | 278587 | 1 | 0 | 0 | 0 | 81761.51 | 101272.65 | 127725.2 | 0 | 152710.36 | 141337.23 | 123317.09 | 8 | **2.978** |
| Q61699 | Hsph1 | Heat shock protein 105 kDa (Mus musculus) | 731.13 | 22.26 | 96407 | 14 | 24309856 | 32394042 | 33700680 | 33112902 | 30348410 | 24129114 | 19164516 | 26180822 | 24731164 | 24757972 | 3 | **0.773** |
| Q61702 | Itih1 | Inter-alpha-trypsin inhibitor heavy chain H1 (Mus musculus) | 69.28 | 1.43 | 101067 | 1 | 0 | 1688212.88 | 1327418.12 | 1784584.5 | 1670532 | 921234.19 | 0 | 682057.06 | 837222.38 | 583988.38 | 5 | **0.467** |
| Q61703 | Itih2 | Inter-alpha-trypsin inhibitor heavy chain H2 (Mus musculus) | 691.32 | 10.68 | 105928 | 10 | 80092088 | 154135456 | 119587656 | 190626800 | 145461728 | 68308360 | 81223432 | 69023712 | 80066984 | 73843368 | 1 | **0.540** |
| Q61704 | Itih3 | Inter-alpha-trypsin inhibitor heavy chain H3 (Mus musculus) | 192.29 | 4.05 | 99358 | 4 | 14952218 | 15293322 | 14856794 | 15055238 | 16118957 | 14852712 | 12881658 | 14339444 | 13430063 | 13637845 | 0 | **0.906** |
| Q61735 | Cd47 | Leukocyte surface antigen CD47 (Mus musculus) | 199.18 | 11.22 | 33098 | 3 | 1856878.62 | 1360091.12 | 1971137.5 | 1462390 | 1491463 | 2068888.88 | 2321879.25 | 2612556 | 2265504.5 | 2723765.5 | 0 | **1.473** |
| Q61749 | Eif2b4 | Translation initiation factor eIF-2B subunit delta (Mus musculus) | 131.72 | 6.3 | 57624 | 2 | 2879764 | 367553.69 | 352557.84 | 2084735.12 | 2397659.25 | 2035659.5 | 2417925.5 | 2390861.5 | 2603586.75 | 1946323.75 | 10 | **1.410** |
| Q61792 | Lasp1 | LIM and SH3 domain protein 1 (Mus musculus) | 624.73 | 41.83 | 29994 | 12 | 36957400 | 30174744 | 35071708 | 26855626 | 47154192 | 35708040 | 43042100 | 39756080 | 41861900 | 39647288 | 6 | **1.135** |
| Q61838 | Pzp | Pregnancy zone protein (Mus musculus) | 120.73 | 1.67 | 165853 | 3 | 43203140 | 59155500 | 54267400 | 78622840 | 54118824 | 53483560 | 55501232 | 57549268 | 46175924 | 54439616 | 11 | **0.923** |
| Q61881 | Mcm7 | DNA replication licensing factor MCM7 (Mus musculus) | 1162.48 | 38.94 | 81211 | 20 | 32743728 | 27580108 | 29226104 | 30527366 | 32424022 | 29843088 | 30428714 | 33429052 | 33595004 | 34738056 | 6 | **1.063** |
| Q61941 | Nnt | NAD(P) transhydrogenase, mitochondrial (Mus musculus) | 185 | 5.06 | 113838 | 4 | 1645045.88 | 1896381 | 2878010 | 2821818.5 | 2591511.75 | 2007668.75 | 2510074 | 1701202.25 | 2273369.5 | 3207132.75 | 12 | **0.989** |
| Q61990 | Pcbp2 | Poly(rC)-binding protein 2 (Mus musculus) | 734.15 | 46.96 | 38222 | 11 | 129606536 | 127542632 | 130549024 | 123716416 | 129456952 | 137509856 | 121852400 | 139270624 | 139708720 | 143482944 | 5 | **1.064** |
| Q62084 | Ppp1r14b | Protein phosphatase 1 regulatory subunit 14B (Mus musculus) b | 120.28 | 9.52 | 15957 | 2 | 354973.91 | 738512.69 | 817771.94 | 429206.19 | 191811.94 | 638197.25 | 266433.75 | 306192.5 | 556277.44 | 811629.75 | 12 | **1.018** |
| Q62087 | Pon3 | Serum paraoxonase/lactonase 3 (Mus musculus) | 136.58 | 7.63 | 39351 | 2 | 0 | 0 | 0 | 221355.36 | 167253.39 | 0 | 0 | 68052.98 | 47407.01 | 43842.28 | 16 | **0.410** |
| Q62093 | Srsf2 | Serine/arginine-rich splicing factor 2 (Mus musculus) | 356.73 | 28.51 | 25476 | 6 | 43016348 | 25575772 | 22620754 | 22425870 | 31209754 | 23165850 | 31640974 | 23930944 | 31199686 | 27496346 | 11 | **0.949** |
| Q62095 | Ddx3y | ATP-dependent RNA helicase DDX3Y (Mus musculus) | 1246.42 | 29.64 | 73428 | 20 | 73173472 | 74546760 | 89401408 | 77977720 | 99176792 | 84502584 | 70348200 | 95722464 | 91448992 | 93649200 | 10 | **1.052** |
| Q62159 | Rhoc | Rho-related GTP-binding protein RhoC (Mus musculus) | 639.96 | 61.66 | 22006 | 10 | 44768740 | 75929272 | 63297544 | 88260248 | 62178964 | 55651712 | 50646912 | 46957568 | 47885084 | 43296232 | 4 | **0.731** |
| Q62167 | Ddx3x | ATP-dependent RNA helicase DDX3X (Mus musculus) | 1544.12 | 39.27 | 73101 | 24 | 100465600 | 95854472 | 113133168 | 92231472 | 120180688 | 110523256 | 98840200 | 123719920 | 117432080 | 119261040 | 7 | **1.092** |
| Q62192 | Cd180 | CD180 antigen (Mus musculus) | 559.4 | 18.61 | 74302 | 9 | 4281750 | 8081371 | 6690938 | 6255928 | 7450838 | 9207954 | 3874752.75 | 8085499.5 | 8331419 | 8353012.5 | 5 | **1.155** |
| Q62193 | Rpa2 | Replication protein A 32 kDa subunit (Mus musculus) | 199.43 | 18.89 | 29718 | 3 | 2866413 | 4035694.5 | 4325762 | 3938593.75 | 4188095 | 4630575 | 1099795 | 3128231.75 | 4227264 | 4413633 | 10 | **0.904** |
| Q62261 | Sptbn1 | Spectrin beta chain, non-erythrocytic 1 (Mus musculus) | 281.04 | 3 | 274223 | 5 | 2647243.25 | 3743727 | 3402273.5 | 1643304.75 | 710290.81 | 2816572 | 2907833.5 | 3176382.5 | 5055512 | 4085491 | 6 | **1.485** |
| Q62318 | Trim28 | Transcription intermediary factor 1-beta (Mus musculus) | 992.04 | 30.46 | 88847 | 18 | 72128736 | 66953184 | 62696468 | 75650176 | 72742576 | 72141056 | 68235128 | 69814824 | 75189040 | 70549328 | 12 | **1.016** |
| Q62348 | Tsn | Translin (Mus musculus) | 588.41 | 45.61 | 26201 | 10 | 30770372 | 33443564 | 43807140 | 36563868 | 41321968 | 37756632 | 42205936 | 49571152 | 50574384 | 53533148 | 3 | **1.257** |
| Q62351 | Tfrc | Transferrin receptor protein 1 (Mus musculus) | 1060.72 | 31.06 | 85731 | 20 | 50407492 | 56730892 | 60393612 | 54076828 | 49038216 | 64450980 | 53906576 | 64544664 | 56252976 | 59999344 | 6 | **1.105** |
| Q62376 | Snrnp70 | U1 small nuclear ribonucleoprotein 70 kDa (Mus musculus) | 312.33 | 11.16 | 51992 | 5 | 12611767 | 10111568 | 12514822 | 8689346 | 12138076 | 19475058 | 14854798 | 15433239 | 14003354 | 17928290 | 0 | **1.457** |
| Q62384 | Zpr1 | Zinc finger protein ZPR1 (Mus musculus) | 180.99 | 10.02 | 50715 | 4 | 3206234.75 | 2032275.12 | 1611006.12 | 2627208 | 1639073.75 | 2009636.75 | 2322757 | 1416482.75 | 3988047.25 | 2539500.75 | 12 | **1.104** |
| Q62393 | Tpd52 | Tumor protein D52 (Mus musculus) | 253.74 | 29.46 | 24313 | 5 | 15876312 | 4788250 | 2096685.12 | 3713855 | 9085266 | 5277910.5 | 4897260.5 | 4498801 | 6847654.5 | 6111579.5 | 11 | **0.777** |
| Q62418 | Dbnl | Drebrin-like protein (Mus musculus) | 345.51 | 17.89 | 48700 | 5 | 7577654.5 | 3938723 | 5304213.5 | 5326833 | 6497200 | 4932544.5 | 5645384 | 5088765 | 4866032 | 5941079 | 9 | **0.924** |
| Q62422 | Ostf1 | Osteoclast-stimulating factor 1 (Mus musculus) | 208.1 | 26.05 | 23783 | 4 | 8977438 | 9831183 | 7761643.5 | 12145120 | 10340110 | 7154883 | 11156788 | 10547859 | 12298829 | 10873224 | 8 | **1.061** |
| Q62425 | Ndufa4 | Cytochrome c oxidase subunit NDUFA4 (Mus musculus) | 200.5 | 68.29 | 9327 | 4 | 14241844 | 11506135 | 9313389 | 9743226 | 9569176 | 7837423 | 10601600 | 8465712 | 4653588.5 | 7664189.5 | 3 | **0.721** |
| Q62426 | Cstb | Cystatin-B (Mus musculus) | 55.55 | 21.43 | 11046 | 1 | 3599607 | 938704.31 | 1424512.88 | 0 | 1227170.38 | 3556527.5 | 2495932.75 | 3202599.75 | 2031917.62 | 2530046.5 | 5 | **1.922** |
| Q62433 | Ndrg1 | Protein NDRG1 (Mus musculus) | 426.14 | 24.87 | 43009 | 7 | 9340474 | 9734781 | 9200961 | 9731754 | 8399910 | 10107543 | 10003538 | 10189747 | 9608176 | 9440149 | 4 | **1.063** |
| Q62446 | Fkbp3 | Peptidyl-prolyl cis-trans isomerase FKBP3 (Mus musculus) | 44.24 | 4.91 | 25148 | 1 | 584479.94 | 310390.84 | 602680 | 583023.94 | 2202360.25 | 0 | 0 | 0 | 0 | 0 | 10 | **0.000** |
| Q62448 | Eif4g2 | Eukaryotic translation initiation factor 4 gamma 2 (Mus musculus) | 611.01 | 16.56 | 102106 | 15 | 19486116 | 24652322 | 21183168 | 23504576 | 20354470 | 24649510 | 25173908 | 23378984 | 24676460 | 21980844 | 5 | **1.098** |
| Q62523 | Zyx | Zyxin (Mus musculus) | 237.91 | 10.11 | 60546 | 5 | 13858361 | 6430135 | 7846679.5 | 5840874.5 | 5602086 | 12974167 | 15565576 | 10104797 | 10915403 | 11064187 | 4 | **1.532** |
| Q62760 | Tomm20 | Mitochondrial import receptor subunit TOM20 homolog (Rattus norvegicus) | 70.56 | 13.79 | 16284 | 1 | 97037.02 | 321930.91 | 522966.84 | 402754.56 | 478780.31 | 631371.81 | 117754.52 | 474879.09 | 570323.75 | 531473.5 | 6 | **1.275** |
| Q62910 | Synj1 | Synaptojanin-1 (Rattus norvegicus) | 66.84 | 0.76 | 172881 | 1 | 329061.38 | 528498.06 | 546263.25 | 466603.56 | 332783.22 | 541412.81 | 509235.88 | 493288 | 493230.31 | 410316.97 | 10 | **1.111** |
| Q62975 | Serpina10 | Protein Z-dependent protease inhibitor (Rattus norvegicus) 10 | 55.2 | 2.29 | 50244 | 1 | 388463.12 | 2042192.88 | 1736123.5 | 2021535.12 | 1990973.38 | 1301179.75 | 1278014.12 | 1096021.5 | 1004830.06 | 829989.06 | 5 | **0.674** |
| Q62991 | Scfd1 | Sec1 family domain-containing protein 1 (Rattus norvegicus) | 245.75 | 11.62 | 72263 | 5 | 3393566.25 | 3043056 | 3636662.25 | 4292040 | 3444489.5 | 3386206.5 | 3408457 | 686743.06 | 3295106 | 3208103 | 5 | **0.785** |
| Q63009 | Prmt1 | Protein arginine N-methyltransferase 1 (Rattus norvegicus) | 971.43 | 50.14 | 40522 | 17 | 25179690 | 39443056 | 38821576 | 38141068 | 29609630 | 33801316 | 37558648 | 42107076 | 46454000 | 45160660 | 6 | **1.198** |
| Q63016 | Slc7a5 | Large neutral amino acids transporter small subunit 1 (Rattus norvegicus) | 91.67 | 8.79 | 55903 | 2 | 0 | 1602384.5 | 1672532.12 | 0 | 1789032.62 | 1878005 | 1792639.12 | 1716709.5 | 1746570.75 | 1790454.12 | 3 | **1.762** |
| Q63041 | A1m | Alpha-1-macroglobulin (Rattus norvegicus) | 133.37 | 1.93 | 167125 | 3 | 12317940 | 19108602 | 16604047 | 22163832 | 12327131 | 8186262 | 8170010 | 13184295 | 13344770 | 12896696 | 6 | **0.676** |
| Q63065 | Pdk1 | [Pyruvate dehydrogenase (acetyl-transferring)] kinase isozyme 1, mitochondrial (Rattus nor | 124.01 | 8.99 | 49081 | 2 | 607331.81 | 1232563.25 | 1681537.75 | 1241185.75 | 1201737.5 | 1026112 | 963021.56 | 220551.33 | 1374231 | 502280.78 | 6 | **0.685** |
| Q63068 | Phax | Phosphorylated adapter RNA export protein (Rattus norvegicus) | 66.52 | 2.86 | 43166 | 1 | 964697.81 | 869603 | 985611 | 985366.31 | 1483194.5 | 1177334.88 | 1185948.75 | 1232816.62 | 1438961.62 | 1382063.62 | 5 | **1.213** |
| Q63159 | Coq3 | Ubiquinone biosynthesis O-methyltransferase, mitochondrial (Rattus norvegicus) | 40.48 | 4.64 | 38708 | 1 | 840454.81 | 0 | 0 | 0 | 0 | 492986.25 | 516861.94 | 0 | 505912.78 | 0 | 19 | **1.804** |
| Q63187 | Eloa | Elongin-A (Rattus norvegicus) | 51.86 | 2.98 | 87203 | 1 | 370485 | 382267 | 261430.55 | 294026.78 | 260810.98 | 480897.47 | 536604.94 | 0 | 197514.61 | 201807.91 | 10 | **0.903** |
| Q63228 | Gmfb | Glia maturation factor beta (Rattus norvegicus) | 226.43 | 41.55 | 16736 | 4 | 16645724 | 13966267 | 11477186 | 14147139 | 16410504 | 15989424 | 13738037 | 15646429 | 14268489 | 14834795 | 12 | **1.025** |
| Q63369 | Nfkb1 | Nuclear factor NF-kappa-B p105 subunit (Rattus norvegicus) | 98.41 | 3.45 | 56554 | 2 | 1695964.88 | 2440074.5 | 2178244.5 | 2565049.75 | 2019869.75 | 2508693.5 | 1508831.12 | 1923650.12 | 1950606.5 | 1743081.12 | 7 | **0.884** |
| Q63413 | Ddx39b | Spliceosome RNA helicase Ddx39b (Rattus norvegicus) | 1220.32 | 54.21 | 49035 | 21 | 180738992 | 158531296 | 156215088 | 163603696 | 175411600 | 183324864 | 168837568 | 179734528 | 174710864 | 173574992 | 7 | **1.055** |
| Q63450 | Camk1 | Calcium/calmodulin-dependent protein kinase type 1 (Rattus norvegicus) | 69.7 | 2.67 | 41638 | 1 | 0 | 2857739.25 | 2851207.75 | 2204564.75 | 1906592.5 | 3197552.5 | 2294252.5 | 3314667 | 2294138.75 | 3059817 | 4 | **1.442** |
| Q63486 | Rraga | Ras-related GTP-binding protein A (Rattus norvegicus) | 170.25 | 15.34 | 36566 | 3 | 2893923.5 | 1918951.88 | 1885976.5 | 2961286 | 2618282.25 | 3069346 | 2610039.5 | 2988756.5 | 3179430.5 | 3087423 | 3 | **1.216** |
| Q63524 | Tmed2 | Transmembrane emp24 domain-containing protein 2 (Rattus norvegicus) | 131.68 | 12.44 | 22733 | 2 | 1727179.5 | 1741035.5 | 2036100.88 | 2473725.75 | 2066277.62 | 1676480.5 | 1053204.12 | 1768498 | 1104189.62 | 1518435.38 | 2 | **0.709** |
| Q63584 | Tmed10 | Transmembrane emp24 domain-containing protein 10 (Rattus norvegicus) | 480.03 | 25.57 | 24858 | 7 | 41505828 | 44608724 | 44039924 | 41271532 | 37238256 | 40906596 | 42755260 | 48168688 | 48013244 | 53140844 | 6 | **1.117** |
| Q63610 | Tpm3 | Tropomyosin alpha-3 chain (Rattus norvegicus) | 1163.18 | 56.05 | 29007 | 21 | 232066976 | 196457808 | 172754848 | 182122976 | 252671536 | 162659808 | 191423296 | 172547536 | 217235712 | 182049776 | 6 | **0.894** |
| Q63615 | Vps33a | Vacuolar protein sorting-associated protein 33A (Rattus norvegicus) | 143.52 | 6.53 | 67514 | 2 | 378827.59 | 349544.09 | 719419.75 | 346649.12 | 656075.5 | 629320.94 | 546869 | 857527 | 731168.19 | 712726.5 | 5 | **1.419** |
| Q63623 | Scaf8 | Protein SCAF8 (Rattus norvegicus) | 75.49 | 0.95 | 139504 | 1 | 1611112.75 | 1161440.5 | 1230587.88 | 1012992.06 | 885152.69 | 1148454.5 | 0 | 603323.5 | 1365239.5 | 1183388.75 | 9 | **0.729** |
| Q63635 | Stx6 | Syntaxin-6 (Rattus norvegicus) | 43.88 | 4.31 | 29057 | 1 | 2522984.75 | 1311813.38 | 1260501.12 | 2370445.5 | 1521626.25 | 1213811.62 | 2034076.5 | 1279720.75 | 1467005 | 1544546.88 | 9 | **0.839** |
| Q63746 | Nfkbia | NF-kappa-B inhibitor alpha (Rattus norvegicus) | 46.08 | 6.37 | 35017 | 1 | 199506.58 | 0 | 279251.44 | 202169.45 | 262263.91 | 165224.09 | 257975.98 | 307166.81 | 186974.83 | 211992.06 | 12 | **1.197** |
| Q63768 | Crk | Adapter molecule crk (Rattus norvegicus) | 53.32 | 3.95 | 33845 | 1 | 818212.56 | 1299213.12 | 699379.19 | 808280.81 | 858442.31 | 1202161.38 | 759944.56 | 1069173.5 | 914480.69 | 1112813.88 | 8 | **1.128** |
| Q63829 | Commd3 | COMM domain-containing protein 3 (Mus musculus) | 143.51 | 16.92 | 22037 | 2 | 422166.69 | 1321263.12 | 1175867.75 | 351336.03 | 1364168.25 | 513048.31 | 705923.75 | 882582.56 | 654139.44 | 1127990.38 | 10 | **0.838** |
| Q63945 | Set | Protein SET (Rattus norvegicus) | 442.47 | 33.56 | 33406 | 7 | 32288204 | 54585480 | 60472480 | 48233804 | 55795860 | 63630112 | 53627996 | 61834944 | 63680044 | 64639648 | 3 | **1.223** |
| Q64012 | Raly | RNA-binding protein Raly (Mus musculus) | 581.47 | 36.54 | 33188 | 11 | 46591560 | 29507376 | 28428116 | 24674232 | 28756140 | 39559476 | 40139252 | 43703856 | 45641196 | 41764128 | 5 | **1.335** |
| Q640N3 | Arhgap30 | Rho GTPase-activating protein 30 (Mus musculus) 0 | 291.71 | 5.09 | 120114 | 5 | 2316403.5 | 3099584.25 | 2810494 | 2041406.12 | 2065002.25 | 3006321.5 | 3957429.5 | 3712807.75 | 3460130.5 | 3091243 | 2 | **1.397** |
| Q64105 | Spr | Sepiapterin reductase (Mus musculus) | 618.78 | 49.81 | 27883 | 10 | 21494828 | 18939344 | 17465872 | 21963048 | 21474036 | 22761374 | 19671276 | 13978734 | 13591816 | 13702711 | 7 | **0.826** |
| Q64152 | Btf3 | Transcription factor BTF3 (Mus musculus) | 348.04 | 40.69 | 22031 | 6 | 29127656 | 14542995 | 13712994 | 14898115 | 13147333 | 11379325 | 14409282 | 10212162 | 14365370 | 10207032 | 4 | **0.709** |
| Q641Y8 | Ddx1 | ATP-dependent RNA helicase DDX1 (Rattus norvegicus) | 845.1 | 23.78 | 82497 | 14 | 33661792 | 29468252 | 26873478 | 25401736 | 23840694 | 34069860 | 30069248 | 26589112 | 27936800 | 27306206 | 8 | **1.048** |
| Q641Z6 | Ehd1 | EH domain-containing protein 1 (Rattus norvegicus) | 1101.47 | 44.19 | 60603 | 21 | 57252712 | 41130408 | 40835868 | 41430684 | 37483112 | 48882228 | 51768092 | 50515572 | 49777016 | 47676692 | 5 | **1.140** |
| Q64213 | Sf1 | Splicing factor 1 (Mus musculus) | 242.73 | 11.18 | 70403 | 5 | 15345217 | 8586038 | 9473810 | 13502342 | 7940786 | 11718684 | 12208648 | 9676971 | 10255852 | 10558766 | 10 | **0.992** |
| Q64232 | Tecr | Very-long-chain enoyl-CoA reductase (Rattus norvegicus) | 192.23 | 9.42 | 36123 | 3 | 3404840.75 | 2697328.5 | 4928238 | 3056768.25 | 4489798.5 | 6918285 | 5046014 | 8107817 | 7018635.5 | 7191101 | 0 | **1.845** |
| Q64240 | Ambp | Protein AMBP (Rattus norvegicus) | 39.26 | 2.01 | 38851 | 1 | 13092443 | 19914738 | 15190335 | 28142160 | 19279354 | 11746894 | 12901745 | 8794090 | 12182833 | 10521481 | 0 | **0.587** |
| Q64261 | Cdk6 | Cyclin-dependent kinase 6 (Mus musculus) | 144.78 | 11.35 | 37028 | 3 | 2605967 | 1635936.38 | 2384111 | 1503237 | 998169.75 | 1261815.38 | 2067282.12 | 1949946.75 | 1624013.75 | 1632293.75 | 11 | **0.935** |
| Q64281 | Lilrb4 | Leukocyte immunoglobulin-like receptor subfamily B member 4 (Mus musculus) | 158.06 | 8.66 | 37544 | 2 | 4344924 | 4550747.5 | 3170979.5 | 4137591 | 4774771 | 4033291.75 | 3932411.25 | 4386333.5 | 4606679.5 | 4902134.5 | 11 | **1.042** |
| Q642C0 | Dnajc8 | DnaJ homolog subfamily C member 8 (Rattus norvegicus) | 147.85 | 18.18 | 29813 | 3 | 7025889 | 6093753 | 3820715.5 | 5799480 | 5031670.5 | 3609048 | 4415517.5 | 3268578.5 | 4126296 | 3206729.25 | 2 | **0.671** |
| Q64310 | Surf4 | Surfeit locus protein 4 (Mus musculus) | 161.04 | 11.52 | 30381 | 2 | 0 | 1997440.25 | 2472123.5 | 0 | 0 | 3149822.5 | 3583743.75 | 4109240.5 | 3835024.75 | 3599830 | 3 | **4.089** |
| Q64324 | Stxbp2 | Syntaxin-binding protein 2 (Mus musculus) | 968.01 | 36.26 | 66357 | 16 | 18996106 | 11398588 | 17726692 | 11939922 | 12762599 | 19261004 | 21123428 | 22442302 | 16499246 | 20280518 | 2 | **1.368** |
| Q64337 | Sqstm1 | Sequestosome-1 (Mus musculus) | 556.84 | 34.62 | 48163 | 9 | 16571344 | 14889086 | 18614046 | 18021344 | 20522124 | 18487792 | 20886420 | 22632888 | 19498468 | 22222588 | 3 | **1.171** |
| Q64378 | Fkbp5 | Peptidyl-prolyl cis-trans isomerase FKBP5 (Mus musculus) | 475.07 | 24.56 | 50966 | 9 | 16970852 | 15378793 | 13295729 | 12349047 | 6548351 | 17659088 | 15596357 | 14952126 | 16485409 | 18985154 | 4 | **1.296** |

| | | | | | | | | | | | | | | | | | |
|---|---|---|---|---|---|---|---|---|---|---|---|---|---|---|---|---|---|
| Q64430 | Atp7a | Copper-transporting ATPase 1 (Mus musculus) | 122.55 | 4.09 | 161959 | 3 | 1543172.12 | 1483242.5 | 1151046.75 | 1895195.25 | 1616114.75 | 1894748.88 | 1646976 | 2223014.75 | 2142208 | 2153948.25 | 2 | **1.309** |
| Q64433 | Hspe1 | 10 kDa heat shock protein, mitochondrial (Mus musculus) | 364.62 | 60.78 | 10963 | 7 | 47728796 | 17487524 | 11830801 | 16608922 | 22048626 | 13334591 | 12815551 | 10915114 | 13925400 | 10049184 | 3 | **0.528** |
| Q64455 | Ptprj | Receptor-type tyrosine-protein phosphatase eta (Mus musculus) | 46.02 | 0.57 | 136769 | 1 | 830331.38 | 918884 | 856927.06 | 764201.31 | 862822.81 | 729345.81 | 838985.19 | 885682.69 | 949395.88 | 683639.12 | 11 | **0.965** |
| Q64511 | Top2b | DNA topoisomerase 2-beta (Mus musculus) | 198.37 | 2.54 | 181909 | 4 | 6332216.5 | 5432771 | 5854563.5 | 6288783 | 7153681 | 6275476 | 5591782.5 | 6121217 | 6007598 | 6638290.5 | 11 | **0.986** |
| Q64514 | Tpp2 | Tripeptidyl-peptidase 2 (Mus musculus) | 1841.9 | 38.27 | 139879 | 35 | 63416304 | 92806832 | 86503000 | 81342248 | 81838632 | 77318504 | 73246672 | 77341096 | 80515992 | 83250544 | 7 | **0.965** |
| Q64516 | Gk | Glycerol kinase (Mus musculus) | 257.38 | 11.99 | 61227 | 5 | 6342370 | 5567125 | 6574299 | 5250268 | 6894587 | 5277967.5 | 4943730.5 | 4982814 | 6493335.5 | 5619250 | 6 | **0.892** |
| Q64521 | Gpd2 | Glycerol-3-phosphate dehydrogenase, mitochondrial (Mus musculus) | 712.79 | 23.52 | 80954 | 12 | 12089437 | 10987854 | 12302110 | 11709918 | 12863244 | 9064719 | 12610966 | 12928290 | 14405851 | 12695782 | 7 | **1.029** |
| Q64522 | Hist2h2ab | Histone H2A type 2-B (Mus musculus) ab | 362.34 | 46.92 | 14013 | 4 | 601903168 | 191210944 | 309859008 | 104602568 | 165518688 | 543550080 | 371312896 | 417661600 | 437849472 | 448702400 | 5 | **1.616** |
| Q64559 | Acot7 | Cytosolic acyl coenzyme A thioester hydrolase (Rattus norvegicus) | 254.61 | 17.85 | 42735 | 5 | 10739042 | 9810430 | 11125309 | 5518940 | 3746235.5 | 12200860 | 13406530 | 10502910 | 6942721.5 | 10273700 | 7 | **1.303** |
| Q64568 | Atp2b3 | Plasma membrane calcium-transporting ATPase 3 (Rattus norvegicus) | 460.35 | 7.15 | 138559 | 8 | 4257239.5 | 4162519.75 | 4968704 | 5302598 | 4838082 | 6718033 | 5358854.5 | 6585834.5 | 6855326 | 7000164 | 0 | **1.382** |
| Q64620 | Ppp6c | Serine/threonine-protein phosphatase 6 catalytic subunit (Rattus norvegicus) | 223.7 | 25.57 | 35159 | 5 | 5213359.5 | 5808321 | 4595732.5 | 5162864 | 4114037.75 | 5069832 | 1232363.62 | 3843941.75 | 3481425.75 | 3420360 | 2 | **0.685** |
| Q64654 | Cyp51a1 | Lanosterol 14-alpha demethylase (Rattus norvegicus) | 60.79 | 4.17 | 56707 | 1 | 141473.11 | 79127.52 | 103795.01 | 0 | 0 | 214417.02 | 243388.11 | 0 | 0 | 330227.75 | 11 | **2.429** |
| Q64674 | Srm | Spermidine synthase (Mus musculus) | 376.28 | 36.09 | 33995 | 7 | 12010679 | 13183930 | 15002532 | 13124441 | 12789413 | 14396674 | 11162805 | 12455278 | 13173452 | 13234769 | 12 | **0.974** |
| Q64701 | Rbl1 | Retinoblastoma-like protein 1 (Mus musculus) | 94.87 | 1.51 | 119456 | 2 | 1121269.62 | 2157592.75 | 1380969.25 | 1938370 | 2062798.5 | 2367853.5 | 1728444.25 | 2079478.25 | 1889186.88 | 2239752 | 7 | **1.190** |
| Q64737 | Gart | Trifunctional purine biosynthetic protein adenosine-3 (Mus musculus) | 1974.32 | 43.56 | 107503 | 29 | 62144468 | 56660248 | 61666016 | 52310736 | 55338796 | 64023396 | 58916760 | 57941764 | 59844448 | 60654096 | 8 | **1.046** |
| Q64FW2 | Retsat | All-trans-retinol 13,14-reductase (Mus musculus) | 48.7 | 1.97 | 67334 | 1 | 1094583.5 | 794159.88 | 1097941 | 0 | 0 | 1275435.12 | 1153302.62 | 1094124.75 | 1010108.06 | 929048.19 | 7 | **1.829** |
| Q65Z40 | Wapl | Wings apart-like protein homolog (Mus musculus) | 239.93 | 6 | 134070 | 4 | 504604.31 | 1613579.38 | 1503509.5 | 1206931 | 1066350.5 | 634117 | 1015411.5 | 202897.41 | 1267959 | 1560849.5 | 9 | **0.794** |
| Q66H47 | Mrpl24 | 39S ribosomal protein L24, mitochondrial (Rattus norvegicus) | 79.81 | 7.41 | 25001 | 2 | 1525060.5 | 653717.88 | 406449.03 | 0 | 971641.38 | 1220101.88 | 1107184.75 | 966283.19 | 742039.56 | 1012064.44 | 7 | **1.419** |
| Q66H58 | Ints14 | Integrator complex subunit 14 (Rattus norvegicus) | 215.9 | 6.99 | 57144 | 4 | 448672.31 | 773806.62 | 905018.19 | 154074.73 | 821904.38 | 960578.75 | 727774.62 | 1021360.06 | 1037511.56 | 1010853.25 | 3 | **1.533** |
| Q66HA6 | Arl8b | ADP-ribosylation factor-like protein 8B (Rattus norvegicus) | 522.48 | 48.39 | 21539 | 10 | 19977360 | 20473012 | 24311932 | 18971378 | 20538468 | 20695596 | 15797918 | 20762400 | 20632866 | 22219986 | 9 | **0.960** |
| Q66HG8 | Ik | Protein Red (Rattus norvegicus) | 36.99 | 1.62 | 65588 | 1 | 1032700.75 | 1046689.38 | 811022.12 | 1290334.88 | 730851.81 | 569294.62 | 713990.06 | 549406.06 | 0 | 457031.47 | 0 | **0.466** |
| Q68FD5 | Cltc | Clathrin heavy chain 1 (Mus musculus) | 5546.32 | 62.09 | 191557 | 85 | 443175936 | 453603424 | 467018080 | 427371904 | 410131392 | 426287072 | 444849344 | 472847424 | 454573280 | 485937248 | 7 | **1.038** |
| Q68FH4 | Galk2 | N-acetylgalactosamine kinase (Mus musculus) | 166.17 | 10.04 | 50503 | 3 | 2387933.75 | 2609896.75 | 2087483.12 | 4547794.5 | 2254848.25 | 2680666 | 2632208 | 2586205 | 2687465.75 | 2508919 | 7 | **0.943** |
| Q68FL4 | Ahcyl2 | Putative adenosylhomocysteinase 3 (Mus musculus) | 358.81 | 12.56 | 66899 | 8 | 5577114 | 6885869.5 | 7360633 | 6300641 | 5786532 | 5618593 | 5298327 | 5412290 | 5224923 | 6423257 | 4 | **0.877** |
| Q68FL6 | Mars | Methionine--tRNA ligase, cytoplasmic (Mus musculus) | 922.62 | 27.27 | 101431 | 16 | 23501046 | 19264368 | 18046866 | 13421543 | 14684978 | 21939196 | 21988674 | 21272772 | 19783084 | 18657514 | 6 | **1.166** |
| Q68FQ7 | Rpap3 | RNA polymerase II-associated protein 3 (Rattus norvegicus) | 143.1 | 4.7 | 74645 | 2 | 984229.38 | 1587681.62 | 1518486.75 | 990975.69 | 1819051.25 | 1316168 | 1378349.75 | 1395982.25 | 1539271 | 1424175.12 | 11 | **1.022** |
| Q68FX0 | Idh3B | Isocitrate dehydrogenase [NAD] subunit beta, mitochondrial (Rattus norvegicus) | 340.72 | 16.1 | 42354 | 6 | 8554133 | 8770983 | 10103298 | 8445233 | 6668109 | 10781554 | 9751475 | 7467230 | 6885022 | 7641021.5 | 12 | **1.000** |
| Q68FX7 | Thoc5 | THO complex subunit 5 homolog (Rattus norvegicus) | 41.45 | 1.03 | 78660 | 1 | 259350.23 | 1465564.5 | 1502307.12 | 1241362 | 1320082.25 | 1701336.38 | 289635.19 | 1303216.75 | 1438091.62 | 1446108.5 | 11 | **1.067** |
| Q68G38 | Tor1a | Torsin-1A (Rattus norvegicus) | 94.46 | 8.41 | 37949 | 2 | 175571.39 | 1210556 | 1206684.62 | 648967.62 | 0 | 911932.38 | 1122116.25 | 1198220 | 1329055.5 | 1375987 | 6 | **1.831** |
| Q69ZK0 | Prex1 | Phosphatidylinositol 3,4,5-trisphosphate-dependent Rac exchanger 1 protein (Mus musculus | 677.16 | 11.39 | 184935 | 14 | 16031509 | 15019242 | 15900163 | 12105854 | 12594382 | 13541800 | 14181523 | 15406799 | 13476052 | 16768460 | 11 | **1.024** |
| Q69ZR2 | Hectd1 | E3 ubiquitin-protein ligase HECTD1 (Mus musculus) | 156.91 | 1.6 | 290086 | 3 | 786470.75 | 986330 | 860229.88 | 833191.12 | 1116955.88 | 507970.94 | 771188.88 | 321011.88 | 248066.69 | 227896.09 | 0 | **0.453** |
| Q6A028 | Swap70 | Switch-associated protein 70 (Mus musculus) | 702.07 | 27.01 | 68996 | 14 | 20340664 | 12146768 | 14668466 | 13444584 | 16743930 | 20681756 | 18176586 | 18725946 | 16625150 | 18489358 | 5 | **1.199** |
| Q6A0A2 | Larp4b | La-related protein 4B (Mus musculus) | 134.65 | 5.4 | 81627 | 3 | 498517.41 | 942268.69 | 1567366.25 | 1078518 | 1034613.62 | 1550041.5 | 1338928.62 | 1108055.25 | 1227365.25 | 910115.12 | 8 | **1.198** |
| Q6A0A9 | FAM120A | Constitutive coactivator of PPAR-gamma-like protein 1 (Mus musculus) | 374.32 | 9.17 | 121646 | 7 | 16756109 | 4590983 | 5828986.5 | 4952607.5 | 6959478.5 | 7025362.5 | 6123902 | 6490121 | 6186608 | 6858209 | 9 | **0.836** |
| Q6A0D4 | Rftn1 | Raftlin (Mus musculus) | 82.84 | 7.22 | 61537 | 2 | 0 | 697283.38 | 733868.5 | 396582.69 | 847693.62 | 485663.81 | 789533.38 | 362252.5 | 388606.75 | 648159.62 | 10 | **1.000** |
| Q6AXS3 | Dek | Protein DEK (Rattus norvegicus) | 362.57 | 20.37 | 42892 | 7 | 17035166 | 16069457 | 13276088 | 12409314 | 12207806 | 14993514 | 20927432 | 16519776 | 19536856 | 18669518 | 3 | **1.277** |
| Q6AXS5 | Serbp1 | Plasminogen activator inhibitor 1 RNA-binding protein (Rattus norvegicus) | 577.34 | 28.26 | 44754 | 9 | 47678948 | 28916110 | 45920468 | 26746068 | 35232164 | 40202308 | 43116020 | 38965016 | 36411540 | 39746640 | 10 | **1.076** |
| Q6AXT7 | Rbm42 | RNA-binding protein 42 (Rattus norvegicus) | 45.05 | 2.11 | 49905 | 1 | 1063492.12 | 613974.5 | 602457.31 | 553819.62 | 722885.12 | 631926.88 | 577247.69 | 880845.81 | 1366896.5 | 713622.12 | 9 | **1.173** |
| Q6AXY7 | Prpf38b | Pre-mRNA-splicing factor 38B (Rattus norvegicus) | 214.29 | 10.7 | 63932 | 4 | 543381 | 3386853 | 3500449 | 2427174.75 | 1883521.12 | 2326549 | 3011683.75 | 3898282.25 | 2092556.38 | 3216305.5 | 10 | **1.239** |
| Q6AY04 | Maip1 | m-AAA protease-interacting protein 1, mitochondrial (Rattus norvegicus) | 50.1 | 3.44 | 33090 | 1 | 0 | 795387.25 | 813827.94 | 619522.06 | 0 | 540035.88 | 499473.25 | 721633 | 717415.31 | 582568.19 | 12 | **1.373** |
| Q6AYA1 | Gar1 | H/ACA ribonucleoprotein complex subunit 1 (Rattus norvegicus) | 151.92 | 19.03 | 23011 | 4 | 4683194 | 8759990 | 6870344 | 8048133 | 6919556.5 | 7904734 | 8922720 | 7475355 | 8648386 | 8279606 | 6 | **1.169** |
| Q6AYC4 | Capg | Macrophage-capping protein (Rattus norvegicus) | 1091.15 | 54.15 | 38799 | 15 | 513798944 | 329410240 | 361621152 | 332138304 | 355899168 | 367705728 | 417563392 | 351203456 | 347692928 | 361232704 | 10 | **0.975** |
| Q6AYE2 | Sh3glb1 | Endophilin-B1 (Rattus norvegicus) | 135.93 | 4.11 | 40787 | 1 | 1967816.75 | 712111.62 | 1099744.75 | 629320.38 | 1292322.5 | 1635831.88 | 1699335 | 2027415 | 1850373.12 | 1798709.5 | 4 | **1.581** |
| Q6AYH5 | Dctn2 | Dynactin subunit 2 (Rattus norvegicus) | 492.33 | 27.11 | 44148 | 8 | 11875237 | 8554747 | 7865284.5 | 9902351 | 8184755.5 | 9265009 | 9950234 | 9337833 | 10645447 | 9829564 | 8 | **1.057** |
| Q6AYK3 | Isyna1 | Inositol-3-phosphate synthase 1 (Rattus norvegicus) | 547.97 | 22.26 | 60884 | 10 | 28869374 | 34230124 | 35442580 | 35589364 | 36639380 | 35481352 | 27944512 | 31871126 | 33110172 | 33760904 | 6 | **0.950** |
| Q6AYL5 | Sf3b4 | Splicing factor 3B subunit 4 (Rattus norvegicus) | 159.56 | 7.31 | 44356 | 2 | 0 | 786676.38 | 551665.69 | 642294.38 | 0 | 759419.69 | 49601.56 | 795406.75 | 631352.75 | 586178 | 9 | **1.425** |
| Q6AYS6 | Snx17 | Sorting nexin-17 (Rattus norvegicus) | 88.72 | 8.3 | 52882 | 2 | 0 | 106631.71 | 510677.66 | 1119651 | 0 | 506305.91 | 0 | 1087483 | 212198.52 | 467614.94 | 13 | **1.309** |
| Q6AYT3 | Rtcb | tRNA-splicing ligase RtcB homolog (Rattus norvegicus) | 374.63 | 18.22 | 55249 | 8 | 10922125 | 7022850.5 | 10388573 | 9154756 | 8249612 | 10675492 | 9791082 | 10581280 | 12574456 | 10937056 | 4 | **1.193** |
| Q6AYT7 | Abhd12 | Monoacylglycerol lipase ABHD12 (Rattus norvegicus) | 366.89 | 30.9 | 45296 | 8 | 8436548 | 7208941.5 | 6707258 | 9464938 | 5853029.5 | 7749662 | 7555788 | 7434140 | 6102858.5 | 6109870.5 | 11 | **0.928** |
| Q6AYY8 | Slc33a1 | Acetyl-coenzyme A transporter 1 (Rattus norvegicus) | 132.7 | 5.64 | 61101 | 3 | 0 | 536015.81 | 1965644.75 | 441271.72 | 1649886.5 | 1738522.25 | 1285752.62 | 538187.94 | 2599802 | 2331277.75 | 5 | **1.849** |
| Q6AZ50 | Atg3 | Ubiquitin-like-conjugating enzyme ATG3 (Rattus norvegicus) | 70.5 | 4.14 | 35822 | 1 | 1270716.62 | 0 | 0 | 0 | 0 | 1382488 | 1341777.75 | 1504295.12 | 1182122.75 | 1379877 | 7 | **5.344** |
| Q6BBI8 | Ufc1 | Ubiquitin-fold modifier-conjugating enzyme 1 (Rattus norvegicus) | 186.44 | 34.13 | 19492 | 4 | 5285583 | 3611718 | 3918638.5 | 1946599.25 | 2779851.75 | 4218345 | 4563336.5 | 3878060.5 | 3882348.5 | 3839399 | 8 | **1.162** |
| Q6DFW4 | Nop58 | Nucleolar protein 58 (Mus musculus) | 632.54 | 23.13 | 60343 | 11 | 25705086 | 29434476 | 23194788 | 19531106 | 21297088 | 22331594 | 29575174 | 22072542 | 20423196 | 21733566 | 12 | **0.975** |
| Q6DGG1 | Abhd14b | Protein ABHD14B (Rattus norvegicus) | 78.18 | 10 | 22618 | 2 | 764128 | 662701.69 | 576443.19 | 0 | 0 | 820134.69 | 767888 | 699924.44 | 0 | 646791.06 | 9 | **1.465** |
| Q6DID7 | Wls | Protein wntless homolog (Mus musculus) | 93.27 | 3.33 | 62188 | 2 | 1561253 | 872990.56 | 1306778.5 | 1365605.75 | 1621327.5 | 1462053.5 | 1618486 | 1766594.25 | 1556922.12 | 1127450.38 | 9 | **1.119** |
| Q6DKG0 | Naa35 | N-alpha-acetyltransferase 35, NatC auxiliary subunit (Rattus norvegicus) | 63.91 | 1.52 | 83210 | 1 | 0 | 142201.73 | 157634.97 | 0 | 0 | 259311.53 | 0 | 144482.44 | 0 | 178255.34 | 14 | **1.941** |
| Q6GQT1 | A2m | Alpha-2-macroglobulin-P (Mus musculus) | 275.57 | 2.85 | 164353 | 5 | 78741472 | 145454720 | 132000832 | 174227104 | 139502784 | 95236152 | 103241640 | 88835728 | 77017152 | 85582088 | 4 | **0.672** |
| Q6GQT9 | Nomo1 | Nodal modulator 1 (Mus musculus) | 721.65 | 18.45 | 133420 | 14 | 10724028 | 9922615 | 11912003 | 13191779 | 7571692.5 | 8468878 | 7660946.5 | 10526504 | 9687481 | 10406036 | 7 | **0.877** |
| Q6GX84 | Fignl1 | Fidgetin-like protein 1 (Rattus norvegicus) | 59.75 | 2.81 | 74197 | 2 | 3499187.5 | 2198051.25 | 1698311.75 | 2490282.5 | 1728726.5 | 2807376.25 | 1811512 | 2145687.25 | 2071640.25 | 2039134.38 | 12 | **0.936** |
| Q6I7R3 | Isoc1 | Isochorismatase domain-containing protein 1 (Rattus norvegicus) | 156.52 | 14.81 | 32002 | 3 | 195654.78 | 1062582.75 | 817751.94 | 1329983 | 1768246.38 | 869052.06 | 1778444.12 | 1711378.12 | 801148.25 | 2040318.5 | 8 | **1.392** |
| Q6IMY8 | Hnrnpu | Heterogeneous nuclear ribonucleoprotein U (Rattus norvegicus) | 1745.15 | 41.73 | 87732 | 30 | 326900576 | 297482944 | 274565344 | 340285568 | 283495968 | 271463296 | 214358560 | 240446576 | 224878944 | 238956000 | 0 | **0.782** |
| Q6IRK9 | Cpq | Carboxypeptidase Q (Rattus norvegicus) | 58.25 | 2.33 | 52042 | 1 | 3123518.5 | 1925399.88 | 1492330.5 | 1559764.25 | 1667023.25 | 1929934.12 | 1912329.5 | 1828895.5 | 1674004.38 | 2086695.88 | 8 | **0.966** |
| Q6IRU2 | Tpm4 | Tropomyosin alpha-4 chain (Mus musculus) | 1177.79 | 49.6 | 28468 | 18 | 95060024 | 93317096 | 90402880 | 80114072 | 102232376 | 79530936 | 88711368 | 70804200 | 83157368 | 78927712 | 2 | **0.870** |
| Q6MG49 | Bag6 | Large proline-rich protein BAG6 (Rattus norvegicus) | 173.08 | 3.4 | 120012 | 3 | 5545902 | 6512303 | 5397259 | 5033788 | 6280863.5 | 6513432 | 7374915 | 6546351 | 6512809 | 6477716.5 | 1 | **1.162** |
| Q6MG61 | Clic1 | Chloride intracellular channel protein 1 (Rattus norvegicus) | 1180.17 | 71.78 | 26981 | 16 | 114818200 | 108123376 | 99289480 | 117453768 | 110913496 | 101303520 | 111692640 | 112340216 | 116209656 | 116790536 | 10 | **1.014** |
| Q6MGA9 | Brd2 | Bromodomain-containing protein 2 (Rattus norvegicus) | 43.21 | 1.63 | 88051 | 1 | 597771.56 | 1431551.25 | 1332651 | 1724493.12 | 1425071.12 | 840435.56 | 1463202.88 | 958660.06 | 1208239.5 | 1048070.5 | 8 | **0.848** |
| Q6MGD0 | Cuta | Protein CutA (Rattus norvegicus) | 117.39 | 18.08 | 18659 | 2 | 3487921.5 | 8620422 | 8751589 | 2563012 | 3396748.5 | 9740528 | 8639913 | 3790826.5 | 3604576.75 | 3733227.25 | 7 | **1.100** |
| Q6NS46 | Pdcd11 | Protein RRP5 homolog (Mus musculus) | 652.59 | 8.43 | 207779 | 12 | 28488194 | 12330772 | 14281149 | 14792030 | 11950500 | 9047179 | 15241961 | 15010878 | 9523439 | 14211374 | 10 | **0.770** |
| Q6NSQ7 | Ltv1 | Protein LTV1 homolog (Mus musculus) | 85.02 | 6.17 | 54023 | 2 | 1623049 | 1504006.5 | 1589268.62 | 1646637.88 | 335405.66 | 1660681.38 | 1223553.62 | 1370047.38 | 1850802.25 | 2029372.25 | 8 | **1.214** |
| Q6NSR8 | Npepl1 | Probable aminopeptidase NPEPL1 (Mus musculus) | 151.43 | 7.06 | 55940 | 3 | 928812 | 3606108.75 | 3262994.5 | 3359475 | 4460097.5 | 3005590 | 4175296.5 | 4101861 | 2946895.5 | 4302196 | 11 | **1.187** |
| Q6NV83 | U2surp | U2 snRNP-associated SURP motif-containing protein (Mus musculus) | 370.58 | 8.84 | 118261 | 7 | 2281948.5 | 3144802.5 | 3629638.75 | 3707161.25 | 3789695 | 1819237.12 | 1546430.75 | 2390092 | 3644847.5 | 4192928.75 | 9 | **0.821** |
| Q6NVE8 | Wdr44 | WD repeat-containing protein 44 (Mus musculus) | 39.06 | 0.77 | 101555 | 1 | 3888677 | 0 | 697168.88 | 1333882.25 | 1718748.38 | 0 | 2338055.75 | 2665577.75 | 2389757.5 | 2225564 | 9 | **1.259** |
| Q6NVF9 | Cpsf6 | Cleavage and polyadenylation specificity factor subunit 6 (Mus musculus) | 301.4 | 12.52 | 59153 | 6 | 9952264 | 9126299 | 9942373 | 7981855.5 | 9888452 | 9690156 | 9317428 | 8356031 | 7592590 | 8595518 | 6 | **0.929** |
| Q6NWW9 | Fndc3b | Fibronectin type III domain-containing protein 3B (Mus musculus) | 98.09 | 1.16 | 132764 | 1 | 215929.27 | 333272.16 | 363312.44 | 296992.97 | 129171.73 | 332712.59 | 336827.06 | 209612.44 | 360924.12 | 262138.56 | 11 | **1.122** |
| Q6NXI6 | Rprd2 | Regulation of nuclear pre-mRNA domain-containing protein 2 (Mus musculus) | 69.41 | 1.7 | 156586 | 1 | 0 | 0 | 183492.48 | 149358.36 | 197623.09 | 0 | 0 | 175285.3 | 237217.45 | 139025.89 | 15 | **1.040** |
| Q6NZC7 | Sec23ip | SEC23-interacting protein (Mus musculus) | 42.27 | 2.1 | 110780 | 1 | 0 | 0 | 351641.97 | 501015.03 | 346185 | 0 | 0 | 188163.55 | 0 | 173702.42 | 13 | **0.302** |
| Q6NZJ6 | Eif4g1 | Eukaryotic translation initiation factor 4 gamma 1 (Mus musculus) | 1790.32 | 24.44 | 176077 | 30 | 47748780 | 52298560 | 50376380 | 51189552 | 50069100 | 57246608 | 44928872 | 60468220 | 57675920 | 56565060 | 5 | **1.100** |
| Q6P069 | Sri | Sorcin (Mus musculus) | 69.81 | 5.56 | 21627 | 1 | 5224541.5 | 3587447 | 3994575.25 | 3267365.5 | 4192207.5 | 3643675.25 | 4679711 | 4066767.25 | 4299711.5 | 4005643.75 | 9 | **1.021** |
| Q6P1B1 | Xpnpep1 | Xaa-Pro aminopeptidase 1 (Mus musculus) | 1115.91 | 43.98 | 69591 | 19 | 51844488 | 46475812 | 48326660 | 54250688 | 46171152 | 49221188 | 44433260 | 42359228 | 44159444 | 47447216 | 5 | **0.921** |
| Q6P2B1 | Tnpo3 | Transportin-3 (Mus musculus) | 299.02 | 8.56 | 104170 | 5 | 1012174 | 2389210 | 2535487.75 | 3072810 | 2205173 | 1486702 | 780089.5 | 2473916.5 | 2046796.75 | 2373168 | 7 | **0.817** |
| Q6P2K6 | Ppp4r3a | Serine/threonine-protein phosphatase 4 regulatory subunit 3A (Mus musculus) | 123.02 | 2.93 | 93842 | 2 | 317307.66 | 1529723.75 | 1753133.5 | 690514.62 | 938149.75 | 1454813.38 | 1160513 | 1899760.38 | 1495289.75 | 2232676 | 6 | **1.576** |
| Q6P542 | Abcf1 | ATP-binding cassette sub-family F member 1 (Mus musculus) | 370.81 | 11.47 | 94945 | 6 | 5995104 | 8192915 | 5519206 | 8725529 | 9551408 | 8831191 | 6068371 | 8117712 | 8061374 | 7161394 | 12 | **1.007** |
| Q6P549 | Inppl1 | Phosphatidylinositol 3,4,5-trisphosphate 5-phosphatase 2 (Mus musculus) | 283.25 | 6.6 | 138973 | 6 | 5922042.5 | 5671564.5 | 5428325.5 | 3566923.5 | 3887346.25 | 6563275 | 7398276.5 | 5840655 | 6492665.5 | 6197518.5 | 1 | **1.328** |
| Q6P5D8 | Smchd1 | Structural maintenance of chromosomes flexible hinge domain-containing protein 1 (Mus m | 674.13 | 8.17 | 225648 | 13 | 188497568 | 81201080 | 123299384 | 44220628 | 101926904 | 138209808 | 145345184 | 142618800 | 127717288 | 12280269 | 9 | **1.050** |
| Q6P5E4 | Uggt1 | UDP-glucose:glycoprotein glucosyltransferase 1 (Mus musculus) | 2092.78 | 35.14 | 176434 | 35 | 47125184 | 52568520 | 50924296 | 52908648 | 44401760 | 61946120 | 50968044 | 60287912 | 54879460 | 56645284 | 2 | **1.148** |
| Q6P5E6 | Gga2 | ADP-ribosylation factor-binding protein GGA2 (Mus musculus) | 109.42 | 5.64 | 66049 | 2 | 215705.12 | 461837 | 305688.91 | 1811921.88 | 2433340.75 | 297234.03 | 253684.44 | 128337.61 | 2308753.75 | 2261137 | 10 | **1.004** |
| Q6P5F9 | Xpo1 | Exportin-1 (Mus musculus) | 1260.33 | 28.1 | 123093 | 23 | 12623372 | 36621260 | 31492868 | 45776396 | 24289132 | 23288650 | 22884180 | 24745342 | 24160746 | 29734452 | 7 | **0.828** |
| Q6P5G6 | Ubxn7 | UBX domain-containing protein 7 (Mus musculus) | 44.49 | 2.78 | 52164 | 1 | 1270464.38 | 1006344.5 | 1217197.38 | 1472999 | 272574.12 | 666097.75 | 0 | 979045.88 | 1553372.88 | 752341.75 | 8 | **0.754** |

| | | | | | | | | | | | | | | | | | |
|---|---|---|---|---|---|---|---|---|---|---|---|---|---|---|---|---|---|
| Q6P5H2 | Nes | Nestin (Mus musculus) | 2575.19 | 30.58 | 207124 | 39 | 69833944 | 47796800 | 61008520 | 33898144 | 71421352 | 69106496 | 62513756 | 65963576 | 65252872 | 69809960 | 10 | **1.171** |
| Q6P6Q9 | Micu1 | Calcium uptake protein 1, mitochondrial (Rattus norvegicus) | 63.6 | 3.56 | 54195 | 1 | 165074.19 | 0 | 0 | 387084.72 | 0 | 197030.92 | 0 | 182691.31 | 241928.98 | 209593.64 | 12 | **1.505** |
| Q6P6S2 | Slc39a11 | Zinc transporter ZIP11 (Rattus norvegicus) 1 | 83.74 | 6.57 | 34868 | 1 | 257823.14 | 0 | 362723.34 | 276475.62 | 0 | 274959.47 | 342839.25 | 272918.03 | 284634.56 | 319993.38 | 8 | **1.667** |
| Q6P6S4 | Sil1 | Nucleotide exchange factor SIL1 (Rattus norvegicus) | 51.26 | 2.15 | 52350 | 1 | 620557.06 | 486242.56 | 413967.28 | 0 | 0 | 550255.75 | 570687.38 | 618009.25 | 0 | 0 | 14 | **1.143** |
| Q6P6T4 | Eml2 | Echinoderm microtubule-associated protein-like 2 (Rattus norvegicus) | 143.16 | 7.7 | 70711 | 3 | 2266448.5 | 2253104.75 | 2321121.5 | 1762449.88 | 4883611 | 6432911 | 2435132.25 | 3772062.5 | 3569897 | 3832611 | 4 | **1.486** |
| Q6P7P5 | Bzw1 | Basic leucine zipper and W2 domain-containing protein 1 (Rattus norvegicus) | 1278.49 | 64.44 | 48043 | 23 | 33261536 | 61683004 | 62472696 | 55738528 | 48634164 | 59071148 | 55240408 | 67991448 | 59780876 | 68182976 | 7 | **1.185** |
| Q6P7Q1 | Babam2 | BRISC and BRCA1-A complex member 2 (Rattus norvegicus) | 129.9 | 7.57 | 43559 | 2 | 740934.12 | 1893792.62 | 2321578.75 | 2456107.25 | 1028874.81 | 1337138.25 | 1772926 | 1824119.5 | 2344910 | 1505321.25 | 12 | **1.041** |
| Q6P8X1 | Snx6 | Sorting nexin-6 (Mus musculus) | 299.49 | 14.29 | 46649 | 5 | 3100572.75 | 7421194 | 10381109 | 6181412.5 | 3698197.75 | 2838160.25 | 4360444 | 3436549.5 | 4625579 | 3003151.5 | 5 | **0.593** |
| Q6P9L6 | Kif15 | Kinesin-like protein KIF15 (Mus musculus) | 56.53 | 0.79 | 160120 | 1 | 0 | 0 | 0 | 232938.61 | 201758.98 | 0 | 0 | 374034.03 | 432459.41 | 351981.09 | 13 | **2.665** |
| Q6P9P6 | Kif11 | Kinesin-like protein KIF11 (Mus musculus) | 245.7 | 5.7 | 118027 | 4 | 1744903.25 | 3671474 | 876250.5 | 2966220.5 | 3287476 | 2705673.5 | 2110549.75 | 3371410.5 | 3149200.5 | 3178179.5 | 11 | **1.157** |
| Q6P9Q4 | Fhod1 | FH1/FH2 domain-containing protein 1 (Mus musculus) | 51.6 | 1.09 | 129600 | 1 | 731776 | 224352.95 | 728697.88 | 991229.06 | 0 | 824399.88 | 768454.19 | 926894.75 | 228420.16 | 807308.56 | 7 | **1.329** |
| Q6P9R1 | Ddx51 | ATP-dependent RNA helicase DDX51 (Mus musculus) | 62.46 | 2.35 | 70368 | 1 | 0 | 0 | 165635.86 | 107120.23 | 179025.83 | 0 | 0 | 166536.72 | 149016.48 | 143374.97 | 15 | **1.016** |
| Q6P9R2 | Oxsr1 | Serine/threonine-protein kinase OSR1 (Mus musculus) | 276.15 | 13.47 | 58214 | 5 | 2935563.5 | 4431650.5 | 5133616 | 3034466 | 3711995.75 | 6859279 | 5021111.5 | 5441414.5 | 5086479 | 5106272.5 | 3 | **1.430** |
| Q6PA06 | Atl2 | Atlastin-2 (Mus musculus) | 128.11 | 4.12 | 66224 | 3 | 1136045.5 | 1038830.06 | 1190756.5 | 895330.56 | 1199783.62 | 1420327.38 | 1184000 | 1440825.25 | 1048177.19 | 1380750.38 | 5 | **1.186** |
| Q6PAR5 | Gapvd1 | GTPase-activating protein and VPS9 domain-containing protein 1 (Mus musculus) | 172.39 | 3.16 | 162402 | 4 | 441586.41 | 1364860.88 | 1679900.12 | 1509245.75 | 360295.09 | 1457027 | 1243755.62 | 1302760.25 | 1159464.75 | 337728.06 | 9 | **1.027** |
| Q6PB44 | Ptpn23 | Tyrosine-protein phosphatase non-receptor type 23 (Mus musculus) | 218.84 | 3.72 | 185216 | 4 | 1476694.88 | 2214965 | 2310879.25 | 1925667.25 | 1289455.25 | 2007808.88 | 1766701.25 | 2860592.25 | 2873981.5 | 2971534.25 | 5 | **1.354** |
| Q6PB66 | Lrpprc | Leucine-rich PPR motif-containing protein, mitochondrial (Mus musculus) | 1412.29 | 25.86 | 156615 | 26 | 22121898 | 35753040 | 38978200 | 39152100 | 33497498 | 33653560 | 33747564 | 35191204 | 36796896 | 38788372 | 12 | **1.051** |
| Q6PB93 | Galnt2 | Polypeptide N-acetylgalactosaminyltransferase 2 (Mus musculus) | 306.76 | 10.88 | 64514 | 6 | 4474865.5 | 5743809 | 6729541.5 | 5155620 | 6328759 | 4976230.5 | 4932300.5 | 6604530.5 | 6363851 | 7071074.5 | 10 | **1.053** |
| Q6PCT3 | Tpd52l2 | Tumor protein D54 (Rattus norvegicus) | 305.96 | 26.82 | 23992 | 5 | 10065032 | 7168793 | 6005493.5 | 3450960.5 | 9047944 | 10557544 | 9555978 | 8869043 | 10172726 | 9391890 | 4 | **1.358** |
| Q6PD03 | Ppp2r5a | Serine/threonine-protein phosphatase 2A 56 kDa regulatory subunit alpha isoform (Mus mu | 249.86 | 13.58 | 56347 | 5 | 941239.75 | 3632894.5 | 4763480.5 | 2332767.5 | 1512240.75 | 2215255.5 | 1727525.38 | 2372694.25 | 5252792 | 2571447.75 | 10 | **1.073** |
| Q6PD26 | Pigs | GPI transamidase component PIG-S (Mus musculus) | 308.03 | 13.15 | 61711 | 5 | 5423142 | 2653285 | 3068118.25 | 2960111.25 | 3167330 | 3990397.25 | 3357748.25 | 4118338.75 | 4337983 | 5119341.5 | 5 | **1.211** |
| Q6PDG5 | Smarcc2 | SWI/SNF complex subunit SMARCC2 (Mus musculus) | 304.8 | 7.83 | 132604 | 6 | 3626599.5 | 3640470.25 | 3140339.5 | 3880090.25 | 2539128.25 | 2773647.5 | 2948148.75 | 3287499.25 | 3539944 | 3539511.5 | 8 | **0.956** |
| Q6PDI5 | Ecpas | Proteasome adapter and scaffold protein ECM29 (Mus musculus) | 720.77 | 8.8 | 203703 | 12 | 6566925.5 | 10707576 | 9101243 | 13422881 | 9917540 | 7633656.5 | 8803280 | 7412982 | 7104517.5 | 7280950.5 | 5 | **0.769** |
| Q6PDM2 | Srsf1 | Serine/arginine-rich splicing factor 1 (Mus musculus) | 356.69 | 29.03 | 27745 | 7 | 20952860 | 21164014 | 11657652 | 39311116 | 22778838 | 16062991 | 9574256 | 11438428 | 10182023 | 11597571 | 1 | **0.508** |
| Q6PDQ2 | Chd4 | Chromodomain-helicase-DNA-binding protein 4 (Mus musculus) | 668.53 | 8.83 | 217751 | 15 | 14706768 | 16647910 | 15876824 | 18027274 | 11429570 | 13343331 | 14536501 | 14547979 | 14408956 | 13523204 | 5 | **0.917** |
| Q6PDV7 | Rpl10 | 60S ribosomal protein L10 (Rattus norvegicus) | 538.01 | 35.05 | 24604 | 8 | 45946304 | 27173170 | 39727192 | 18494318 | 41168152 | 55690872 | 56597908 | 56576288 | 54520872 | 55282632 | 0 | **1.615** |
| Q6PE01 | Snrnp40 | U5 small nuclear ribonucleoprotein 40 kDa protein (Mus musculus) | 43.88 | 3.35 | 39276 | 1 | 1349658.25 | 1181566.88 | 0 | 641035.75 | 0 | 0 | 1422879.75 | 1804169.62 | 0 | 2164842 | 11 | **1.700** |
| Q6PEC3 | Yif1b | Protein YIF1B (Rattus norvegicus) | 51.87 | 5.02 | 28418 | 1 | 747185.38 | 0 | 433915.44 | 443705.38 | 0 | 870316.75 | 641791.94 | 573456.31 | 677911.88 | 615644.94 | 5 | **2.080** |
| Q6PEC4 | Skp1 | S-phase kinase-associated protein 1 (Rattus norvegicus) | 551.05 | 65.64 | 18672 | 8 | 10144317 | 18456662 | 17192762 | 18519496 | 16491640 | 15307620 | 17252862 | 17969066 | 20612084 | 18898904 | 8 | **1.114** |
| Q6PFD9 | Nup98 | Nuclear pore complex protein Nup98-Nup96 (Mus musculus) | 298.59 | 5.01 | 197241 | 6 | 3313934 | 2551144.5 | 4969427 | 4305914 | 3451249.25 | 3577742 | 3177796.5 | 4114510.25 | 3626685 | 4387125 | 11 | **1.016** |
| Q6PFR5 | Tra2a | Transformer-2 protein homolog alpha (Mus musculus) | 190.64 | 16.01 | 32316 | 3 | 3353813.75 | 4847828.5 | 3905799 | 3779141.75 | 3107025.25 | 3995551 | 2598590.5 | 3807678.25 | 4160480.5 | 4540364 | 10 | **1.006** |
| Q6PGB6 | Naa50 | N-alpha-acetyltransferase 50 (Mus musculus) | 537.41 | 51.48 | 19414 | 10 | 12115995 | 17680974 | 21781444 | 18235504 | 19203004 | 20465260 | 17803200 | 20940126 | 18913902 | 18573084 | 9 | **1.086** |
| Q6PGC1 | Dhx29 | ATP-dependent RNA helicase DHX29 (Mus musculus) | 52.08 | 0.88 | 153975 | 1 | 0 | 752708.19 | 671089.81 | 713607.25 | 592747.31 | 646695 | 505715.31 | 570181.62 | 691644.12 | 563786 | 8 | **1.091** |
| Q6PGF3 | Med16 | Mediator of RNA polymerase II transcription subunit 16 (Mus musculus) | 43.65 | 1.21 | 91786 | 1 | 1718915.5 | 3844718.75 | 2619110.75 | 4143951 | 3799213.25 | 2021779.25 | 2059921.25 | 2046743.38 | 1521259.5 | 1997711 | 4 | **0.598** |
| Q6PGF7 | Exoc8 | Exocyst complex component 8 (Mus musculus) | 54.9 | 1.26 | 81035 | 1 | 170724.12 | 126557.28 | 44904.78 | 107813.12 | 248449.5 | 35496.38 | 40669.61 | 89879.04 | 151047.36 | 0 | 4 | **0.454** |
| Q6PGG2 | Gmip | GEM-interacting protein (Mus musculus) | 72.38 | 1.34 | 107545 | 1 | 889324.12 | 0 | 0 | 0 | 0 | 799558.56 | 1038865.69 | 747986 | 862418 | 772741.31 | 10 | **4.747** |
| Q6PGL7 | Washc2 | WASH complex subunit 2 (Mus musculus) | 183.18 | 2.92 | 145311 | 3 | 811541.25 | 1504555.12 | 1877692.5 | 936078.94 | 1907482.75 | 1377997.5 | 1843480.38 | 1476546.62 | 1884916.75 | 1582737.38 | 11 | **1.160** |
| Q6PIP5 | Nudcd1 | NudC domain-containing protein 1 (Mus musculus) | 315.79 | 17.53 | 66705 | 6 | 3924964 | 4514111.5 | 7061545.5 | 6566514.5 | 5557757.5 | 4722487 | 4334363 | 5478014.5 | 5397253 | 6328054.5 | 10 | **0.951** |
| Q6PIU9 | | Uncharacterized protein FLJ45252 homolog (Mus musculus) | 58.35 | 4.52 | 37705 | 1 | 0 | 970983.31 | 0 | 0 | 0 | 639244.62 | 659193.69 | 793115.44 | 795510.06 | 785123.88 | 11 | **3.782** |
| Q6PR54 | Rif1 | Telomere-associated protein RIF1 (Mus musculus) | 59.41 | 0.7 | 266228 | 1 | 711464.81 | 0 | 298600.44 | 0 | 0 | 296070.16 | 367146.53 | 347040.56 | 400770.06 | 346227.81 | 9 | **1.740** |
| Q6Q899 | Ddx58 | Probable ATP-dependent RNA helicase DDX58 (Mus musculus) | 141.06 | 5.94 | 105975 | 3 | 1144744.25 | 1611819.62 | 2211699.5 | 1415247.38 | 1488109.88 | 2171103.75 | 637729.5 | 1346275.75 | 2248857 | 2476778 | 10 | **1.128** |
| Q6QA69 | Abhd5 | 1-acylglycerol-3-phosphate O-acyltransferase ABHD5 (Rattus norvegicus) | 48.94 | 4.56 | 39104 | 1 | 537760.62 | 400881.03 | 561477.44 | 437403.25 | 0 | 451195.97 | 607705.38 | 503315.75 | 508313.12 | 476820.12 | 8 | **1.315** |
| Q6QI44 | Naa25 | N-alpha-acetyltransferase 25, NatB auxiliary subunit (Rattus norvegicus) | 233.65 | 6.19 | 111258 | 5 | 1464246.5 | 4020346 | 3234405.5 | 4152688 | 2656899.25 | 2627344.25 | 3556214 | 3686924.25 | 2167771 | 2869882.5 | 10 | **0.960** |
| Q6R5N8 | Tlr13 | Toll-like receptor 13 (Mus musculus) | 245.59 | 6.36 | 114443 | 4 | 1451025 | 535826.94 | 1955733.75 | 1779767.75 | 1249303.5 | 1918985.62 | 2326784.5 | 2528471.75 | 2073375.38 | 2431242 | 1 | **1.618** |
| Q6TEK3 | Vkorc1l1 | Vitamin K epoxide reductase complex subunit 1-like protein 1 (Rattus norvegicus) 1 | 61.67 | 6.25 | 19779 | 1 | 232709.2 | 246894.72 | 354725.97 | 379431.88 | 421643.25 | 431979.28 | 427549.25 | 470569 | 395827.28 | 350751.41 | 4 | **1.270** |
| Q6TUG0 | Dnajb11 | DnaJ homolog subfamily B member 11 (Rattus norvegicus) | 304.96 | 19.83 | 40495 | 5 | 11579732 | 8580338 | 9186313 | 5894812 | 6050657.5 | 9527958 | 9530410 | 9867872 | 9262364 | 7775125 | 7 | **1.113** |
| Q6UPE1 | Etfdh | Electron transfer flavoprotein-ubiquinone oxidoreductase, mitochondrial (Rattus norvegicus | 47.53 | 1.95 | 68198 | 1 | 1450633.62 | 823893.12 | 1242392.5 | 986576.06 | 1348692.5 | 1354783.38 | 1589130 | 1475990.75 | 2083374.88 | 1796484.75 | 1 | **1.418** |
| Q6URK4 | Hnrnpa3 | Heterogeneous nuclear ribonucleoprotein A3 (Rattus norvegicus) | 1379.86 | 44.06 | 39652 | 21 | 88436600 | 120112368 | 135562512 | 104023208 | 124051696 | 142285280 | 112936144 | 136916960 | 133085496 | 138586384 | 4 | **1.160** |
| Q6URW6 | Myh14 | Myosin-14 (Mus musculus) | 486.35 | 4.4 | 228586 | 8 | 74616480 | 63095568 | 60315128 | 68083096 | 62195564 | 64325076 | 58444412 | 59428664 | 61735480 | 59315260 | 4 | **0.924** |
| Q6VBQ8 | Pno1 | RNA-binding protein PNO1 (Rattus norvegicus) | 139.94 | 12.1 | 27472 | 2 | 1242929 | 2264735.5 | 2150548.5 | 2138783.75 | 1748934.12 | 1362469.25 | 1548761 | 2246555 | 1727772.88 | 1318309 | 8 | **0.859** |
| Q6VN19 | Ranbp10 | Ran-binding protein 10 (Mus musculus) | 75.39 | 1.94 | 67188 | 1 | 822369.19 | 631061 | 469659.84 | 747847.56 | 608468.19 | 709329.69 | 672020.44 | 467926.19 | 546595.88 | 534799.75 | 8 | **0.894** |
| Q6WKZ8 | Ubr2 | E3 ubiquitin-protein ligase UBR2 (Mus musculus) | 51.78 | 0.74 | 199155 | 1 | 0 | 627219.38 | 648001.56 | 392413.78 | 186615.38 | 0 | 0 | 512435.62 | 422642.69 | 521176.31 | 12 | **0.785** |
| Q6WVG3 | Kctd12 | BTB/POZ domain-containing protein KCTD12 (Mus musculus) | 545.1 | 43.12 | 35892 | 9 | 6394612.5 | 16186642 | 14535545 | 16914912 | 10795665 | 10008356 | 6497943 | 8717023 | 7275317 | 9206331 | 5 | **0.643** |
| Q6Y7W8 | Gigyf2 | GRB10-interacting GYF protein 2 (Mus musculus) | 239.27 | 5.65 | 149193 | 4 | 1038129.62 | 2225299 | 1670873.88 | 1633779 | 611437.38 | 1500366.62 | 1612830.75 | 1078464.75 | 1509223 | 1148632 | 10 | **0.954** |
| Q6ZPE2 | Sbf1 | Myotubularin-related protein 5 (Mus musculus) | 100.2 | 1.55 | 208693 | 2 | 725494.88 | 708593.88 | 658439.75 | 443211.38 | 498561.09 | 747486.44 | 928179.62 | 889962.12 | 566839.62 | 460571.12 | 7 | **1.184** |
| Q6ZPF4 | Fmnl3 | Formin-like protein 3 (Mus musculus) | 161.07 | 2.92 | 117169 | 3 | 1478677.75 | 1106506.5 | 676126.5 | 655391.19 | 455238.88 | 826700.5 | 1144825.75 | 642117.44 | 498212.69 | 0 | 9 | **0.712** |
| Q6ZPJ3 | Ube2o | (E3-independent) E2 ubiquitin-conjugating enzyme UBE2O (Mus musculus) | 49.27 | 1.16 | 140834 | 1 | 573832.81 | 577768.56 | 735730.38 | 528058.88 | 718984.62 | 849914.81 | 709376.56 | 716214.19 | 978623.94 | 736537.94 | 4 | **1.273** |
| Q6ZPY7 | Kdm3b | Lysine-specific demethylase 3B (Mus musculus) | 107.06 | 3.39 | 170875 | 3 | 894709.62 | 683118.19 | 832599.94 | 747631.75 | 1096078.25 | 811799.62 | 205954.52 | 870999.81 | 580106.12 | 1709636.88 | 10 | **0.982** |
| Q6ZPZ3 | Zc3h4 | Zinc finger CCCH domain-containing protein 4 (Mus musculus) | 44.92 | 1.23 | 140967 | 1 | 426593.12 | 228146.19 | 234027.36 | 169446.14 | 243452.88 | 203584.75 | 0 | 222943.69 | 236799.98 | 188856.17 | 6 | **0.655** |
| Q6ZQ08 | Cnot1 | CCR4-NOT transcription complex subunit 1 (Mus musculus) | 301.74 | 3.54 | 266808 | 7 | 941379.12 | 3459081.25 | 3260407.5 | 3896999.25 | 2794310.75 | 2796938.25 | 2573570 | 3663323.5 | 3355956.5 | 3625044.25 | 11 | **1.116** |
| Q6ZQ58 | Larp1 | La-related protein 1 (Mus musculus) | 542.02 | 11.1 | 121125 | 10 | 6742024 | 4397926 | 5096867 | 4530643 | 5932518.5 | 6205548 | 5479640 | 5463884 | 6286088 | 6655474.5 | 7 | **1.127** |
| Q6ZQ88 | Kdm1a | Lysine-specific histone demethylase 1A (Mus musculus) | 157.31 | 5.39 | 92851 | 3 | 0 | 554883.25 | 488819.12 | 965345.5 | 1332663 | 1352674 | 843680 | 597859.12 | 1512429.75 | 820495.62 | 6 | **1.534** |
| Q6ZQA0 | Nbeal2 | Neurobeachin-like protein 2 (Mus musculus) | 124.3 | 1.35 | 301924 | 3 | 466361.16 | 0 | 2021299 | 0 | 519807.12 | 540625.94 | 816847.56 | 1264471 | 1912514 | 1338394.38 | 6 | **1.953** |
| Q6ZQB6 | Ppip5k2 | Inositol hexakisphosphate and diphosphoinositol-pentakisphosphate kinase 2 (Mus musculu | 207.16 | 4.07 | 128427 | 4 | 577268.69 | 390517.75 | 395383.25 | 970593.25 | 939821.12 | 789206 | 1655937.25 | 1261161.25 | 1614895.25 | 1852465.12 | 2 | **2.191** |
| Q6ZQH8 | Nup188 | Nucleoporin NUP188 homolog (Mus musculus) | 103.7 | 1.48 | 196696 | 2 | 491154.19 | 97138.81 | 120123.32 | 90362.17 | 123442.93 | 324912.72 | 330845.62 | 394346.88 | 453991.41 | 428387.62 | 5 | **2.095** |
| Q6ZQL4 | Wdr43 | WD repeat-containing protein 43 (Mus musculus) | 349.15 | 13.44 | 75381 | 6 | 7379772.5 | 7304457 | 7972580 | 6194982.5 | 6619699.5 | 7355829.5 | 7674798.5 | 5669813.5 | 4572728.5 | 7434134 | 11 | **0.922** |
| Q6ZQM8 | Ugt1a7c | UDP-glucuronosyltransferase 1-7C (Mus musculus) | 1040.11 | 43.69 | 59758 | 20 | 57994828 | 65241960 | 62152608 | 78728800 | 50414808 | 55061044 | 62072384 | 64788568 | 67079096 | 64474116 | 12 | **0.997** |
| Q6ZWM4 | Lsm8 | U6 snRNA-associated Sm-like protein LSm8 (Mus musculus) | 238.34 | 57.29 | 10403 | 4 | 10502714 | 10666172 | 10603018 | 9565185 | 10696359 | 10757285 | 8663580 | 9336354 | 10887683 | 10619739 | 12 | **0.966** |
| Q6ZWQ7 | Spcs3 | Signal peptidase complex subunit 3 (Mus musculus) | 47.2 | 6.11 | 20313 | 1 | 1250247.5 | 1181256.12 | 1154966 | 1376555.12 | 1212961.25 | 1657545.88 | 1375058.62 | 1648304.75 | 1607736.38 | 2127076.75 | 1 | **1.363** |
| Q6ZWR6 | Syne1 | Nesprin-1 (Mus musculus) | 113.55 | 0.23 | 1009926 | 2 | 541038 | 865393.88 | 756714.19 | 937441.88 | 1067613.5 | 531457.69 | 830150.12 | 905137.25 | 970335.5 | 733440.06 | 10 | **0.953** |
| Q75Q40 | Tomm40 | Mitochondrial import receptor subunit TOM40 homolog (Rattus norvegicus) | 355.82 | 16.62 | 37919 | 5 | 3261141 | 21053388 | 15786607 | 6387929 | 5986126 | 7781842 | 6873703.5 | 16201997 | 17943792 | 18030776 | 7 | **1.274** |
| Q76MZ3 | Ppp2r1a | Serine/threonine-protein phosphatase 2A 65 kDa regulatory subunit A alpha isoform (Mus r | 2279.06 | 62.14 | 65323 | 32 | 86862904 | 104173520 | 98276088 | 102825560 | 88094064 | 94670208 | 90455704 | 100571880 | 85693176 | 97468384 | 9 | **0.976** |
| Q76N33 | Stambpl1 | AMSH-like protease (Mus musculus) 1 | 93.13 | 7.11 | 49640 | 2 | 2927159.25 | 2188490 | 2359345.5 | 785481.62 | 0 | 3031030 | 2565897 | 2594602.75 | 2251526.25 | 2238950 | 6 | **1.535** |
| Q78IK2 | Atp5md | Up-regulated during skeletal muscle growth protein 5 (Mus musculus) | 146.94 | 43.1 | 6382 | 2 | 3988656 | 3286536.75 | 2986828.75 | 4028452.5 | 2706885.5 | 1672012 | 2484547.25 | 2530898.25 | 2957842 | 3405169.5 | 4 | **0.768** |
| Q78IK4 | Apool | MICOS complex subunit Mic27 (Mus musculus) | 364.22 | 33.21 | 29261 | 6 | 11326233 | 7031175 | 7949862 | 8065786 | 7586598.5 | 6689788 | 9457782 | 7674828.5 | 7678455 | 8005355.5 | 11 | **0.942** |
| Q78JE5 | Fbxo22 | F-box only protein 22 (Mus musculus) | 209.19 | 15.42 | 44203 | 5 | 5092280 | 3400195.25 | 4193134.25 | 6811863 | 2038506 | 3751831.25 | 2946040.5 | 3297669 | 3812636.5 | 3584770.75 | 8 | **0.808** |
| Q78P75 | Dynll2 | Dynein light chain 2, cytoplasmic (Rattus norvegicus) | 84.68 | 20.22 | 10350 | 2 | 11312828 | 8991256 | 8392927 | 5441795 | 9405565 | 8887144 | 6500631 | 9872793 | 9446716 | 5337651.5 | 11 | **0.920** |
| Q78PB6 | Ndel1 | Nuclear distribution protein nudE-like 1 (Rattus norvegicus) | 61.6 | 3.48 | 38366 | 1 | 542876.44 | 241653.7 | 290373.88 | 393315.12 | 388964.78 | 414390.41 | 617422.81 | 557270.62 | 383888.25 | 300854.12 | 7 | **1.224** |
| Q78PY7 | Snd1 | Staphylococcal nuclease domain-containing protein 1 (Mus musculus) | 2281.55 | 51.98 | 102088 | 37 | 131920752 | 121943776 | 119835416 | 94553224 | 94836288 | 111913960 | 111868688 | 111807600 | 116954544 | 120937680 | 11 | **1.018** |
| Q78T54 | Vma21 | Vacuolar ATPase assembly integral membrane protein Vma21 (Mus musculus) | 49.26 | 9.9 | 11365 | 1 | 2305852.5 | 1087636.25 | 1061302.38 | 2529936.5 | 2597800 | 1895379.5 | 1044923.69 | 1875275.25 | 2199031.5 | 1977856.75 | 8 | **0.938** |
| Q78ZA7 | Nap1l4 | Nucleosome assembly protein 1-like 4 (Mus musculus) | 684.12 | 41.87 | 42679 | 11 | 25482166 | 26644142 | 28730476 | 25615648 | 26514372 | 23881436 | 25138950 | 24097952 | 25186942 | 24353602 | 0 | **0.922** |
| Q791T5 | Mtch1 | Mitochondrial carrier homolog 1 (Mus musculus) | 49.92 | 3.34 | 41565 | 1 | 0 | 129370.87 | 0 | 678511 | 427818.28 | 0 | 0 | 578529.06 | 642700.56 | 682522 | 13 | **1.541** |
| Q791V5 | Mtch2 | Mitochondrial carrier homolog 2 (Mus musculus) | 540.12 | 50.83 | 33499 | 10 | 15239935 | 12609204 | 13638062 | 14449011 | 14263456 | 12132626 | 18882672 | 16533925 | 18601402 | 16282498 | 5 | **1.174** |
| Q793I8 | Tifa | TRAF-interacting protein with FHA domain-containing protein A (Mus musculus) | 37.95 | 4.89 | 21560 | 1 | 625720.94 | 617368.06 | 861653.56 | 564969.62 | 0 | 848127.25 | 613938 | 819493.94 | 774309.81 | 675328.88 | 7 | **1.398** |
| Q794E4 | Hnrnpf | Heterogeneous nuclear ribonucleoprotein F (Rattus norvegicus) | 943.7 | 50.12 | 45730 | 13 | 70381568 | 62357552 | 61809296 | 63135592 | 57729152 | 61712688 | 58048392 | 62482744 | 69617600 | 76646576 | 11 | **1.042** |
| Q7TMB8 | Cyfip1 | Cytoplasmic FMR1-interacting protein 1 (Mus musculus) | 1602.59 | 22.59 | 145241 | 30 | 24494034 | 33896004 | 30843050 | 39114576 | 23380138 | 28118984 | 27806678 | 33344650 | 29205822 | 30528726 | 11 | **0.982** |
| Q7TMF3 | Ndufa12 | NADH dehydrogenase [ubiquinone] 1 alpha subcomplex subunit 12 (Mus musculus) | 112.05 | 17.24 | 17086 | 2 | 1934291.5 | 1354168 | 1162960.38 | 264477.56 | 731882.88 | 962467.75 | 1158761.25 | 1023432.88 | 1037316.5 | 512553.47 | 9 | **0.862** |
| Q7TMK9 | Syncrip | Heterogeneous nuclear ribonucleoprotein Q (Mus musculus) | 778 | 28.09 | 69633 | 14 | 47783240 | 41226784 | 37374900 | 38168988 | 37394964 | 43726800 | 42970336 | 42178652 | 41391712 | 39241636 | 6 | **1.037** |

| | | | | | | | | | | | | | | | | | |
|---|---|---|---|---|---|---|---|---|---|---|---|---|---|---|---|---|---|
| Q7TMR0 | Prcp | Lysosomal Pro-X carboxypeptidase (Mus musculus) | 145.15 | 9.57 | 55027 | 2 | 0 | 289063.59 | 805594.5 | 315681.59 | 317323.31 | 587373.69 | 127112.14 | 305307.16 | 392126.47 | 472782.19 | 10 | **1.091** |
| Q7TMY7 | Ipo8 | Importin-8 (Mus musculus) | 125.2 | 2.57 | 117078 | 2 | 2740633.5 | 187241.09 | 4535920 | 5144256 | 0 | 2928488 | 2471367.25 | 2615510 | 2406522.5 | 2152662.75 | 11 | **0.997** |
| Q7TMY8 | Huwe1 | E3 ubiquitin-protein ligase HUWE1 (Mus musculus) | 754.27 | 4.91 | 482635 | 14 | 40088968 | 24984356 | 8807937 | 24157722 | 35581704 | 37768384 | 9274814 | 9971884 | 25711366 | 33034620 | 12 | **0.866** |
| Q7TNC4 | Luc7l2 | Putative RNA-binding protein Luc7-like 2 (Mus musculus) | 369.66 | 22.19 | 46583 | 8 | 15953886 | 18434930 | 20102706 | 17534648 | 17917466 | 17068648 | 15964620 | 17697148 | 16668851 | 18797606 | 9 | **0.958** |
| Q7TNE3 | Spag7 | Sperm-associated antigen 7 (Mus musculus) | 41.94 | 10.13 | 25921 | 1 | 618501.62 | 420123.75 | 279169.31 | 484911.72 | 887150.06 | 437657.5 | 0 | 487632.84 | 286350.78 | 485046.81 | 9 | **0.631** |
| Q7TPJ0 | Ssr1 | Translocon-associated protein subunit alpha (Rattus norvegicus) | 123.95 | 7.21 | 35629 | 2 | 8488896 | 10362602 | 7811083.5 | 10721379 | 9193645 | 10042187 | 10257873 | 8918402 | 8669264 | 8792000 | 12 | **1.002** |
| Q7TPR4 | Actn1 | Alpha-actinin-1 (Mus musculus) | 1702.46 | 30.72 | 103068 | 26 | 67426464 | 77366416 | 76839544 | 74468936 | 75249656 | 81892144 | 77900288 | 77061240 | 80387024 | 74696656 | 4 | **1.055** |
| Q7TPV4 | Mybbp1a | Myb-binding protein 1A (Mus musculus) | 720.38 | 10.94 | 152037 | 10 | 9312836 | 8626739 | 8947684 | 10636028 | 10015083 | 7133762 | 9940156 | 7143808 | 7980702 | 6721491.5 | 3 | **0.819** |
| Q7TQ94 | Nit1 | Deaminated glutathione amidase (Rattus norvegicus) | 51.21 | 5.2 | 36094 | 1 | 0 | 320214.16 | 335147.12 | 243438.36 | 392877 | 0 | 0 | 377367.59 | 361477.72 | 331040.47 | 14 | **0.828** |
| Q7TQH0 | Atxn2l | Ataxin-2-like protein (Mus musculus) | 328.78 | 8.87 | 110649 | 7 | 8355028.5 | 8803118 | 8389726 | 8950738 | 11919496 | 9853273 | 11027946 | 9215324 | 7985675.5 | 8673146 | 11 | **1.007** |
| Q7TQH7 | Lrp10 | Low-density lipoprotein receptor-related protein 10 (Mus musculus) | 43.77 | 0.98 | 76461 | 1 | 603148.44 | 0 | 384286.81 | 0 | 0 | 1232649 | 880943.88 | 739692.44 | 833403.62 | 0 | 8 | **3.734** |
| Q7TQK1 | Ints7 | Integrator complex subunit 7 (Mus musculus) | 45.84 | 1.35 | 106861 | 1 | 235939.98 | 379172.75 | 287361.28 | 386590.72 | 267013.19 | 417654.62 | 470710.19 | 375320.72 | 355843.09 | 308023.94 | 6 | **1.239** |
| Q7TS74 | Ckap2l | Cytoskeleton-associated protein 2-like (Mus musculus) | 42.39 | 3.09 | 82999 | 1 | 0 | 119786.75 | 145557.83 | 105652.85 | 122279.08 | 0 | 0 | 133239.72 | 154542.56 | 132666.22 | 12 | **0.852** |
| Q7TSI3 | Ppp6r1 | Serine/threonine-protein phosphatase 6 regulatory subunit 1 (Mus musculus) | 350.51 | 10.98 | 94527 | 6 | 3480698.5 | 3908504.5 | 4906620.5 | 4838460.5 | 3676374.75 | 2264275.75 | 4388926 | 2792892.25 | 4407177 | 4621332 | 9 | **0.888** |
| Q7TSS2 | Ube2q1 | Ubiquitin-conjugating enzyme E2 Q1 (Mus musculus) | 90.55 | 6.16 | 46173 | 2 | 144417.78 | 1990203 | 2181290 | 1900658.25 | 1762853.5 | 1899449.62 | 1836734.75 | 1857184 | 2324844.5 | 1917840.88 | 11 | **1.233** |
| Q7TSV4 | Pgm2 | Phosphoglucomutase-2 (Mus musculus) | 38.97 | 1.61 | 68748 | 1 | 1680637.75 | 527525.12 | 1275280.38 | 1033047.5 | 1483162.75 | 1276886.88 | 918056.81 | 1972680.25 | 982320.75 | 1231841.62 | 12 | **1.064** |
| Q7TT45 | Rragd | Ras-related GTP-binding protein D (Mus musculus) | 257.72 | 16.7 | 51232 | 5 | 5831176.5 | 9464054 | 9870206 | 9020963 | 10908344 | 8638116 | 7583039.5 | 10411055 | 11234809 | 10797798 | 10 | **1.079** |
| Q80T85 | Dcaf5 | DDB1- and CUL4-associated factor 5 (Mus musculus) | 41.72 | 0.74 | 103676 | 1 | 518199.88 | 342910.78 | 363971.28 | 141998.27 | 441799.69 | 407748.06 | 394657.19 | 354943.53 | 374786.22 | 459440.12 | 10 | **1.101** |
| Q80TE0 | Rpap1 | RNA polymerase II-associated protein 1 (Mus musculus) | 52.52 | 0.92 | 155270 | 1 | 477521.12 | 348386.59 | 317781.09 | 0 | 93431.27 | 353303.09 | 374187.84 | 455864.03 | 0 | 0 | 13 | **0.957** |
| Q80TH2 | Erbin | Erbin (Mus musculus) | 93.98 | 2.43 | 157248 | 2 | 0 | 1346918.75 | 1465541.12 | 1483115.88 | 1081130.62 | 989836.44 | 0 | 1291288.25 | 1536123 | 1337761.25 | 11 | **0.959** |
| Q80TL7 | Mon2 | Protein MON2 homolog (Mus musculus) | 106.16 | 2.22 | 189078 | 2 | 1934862.88 | 1347423.75 | 1654557.62 | 1179537.38 | 950424.94 | 1513273.75 | 1877083.5 | 368836.03 | 1665047.88 | 1659063.88 | 10 | **1.002** |
| Q80TM9 | Nisch | Nischarin (Mus musculus) | 412.47 | 6.4 | 175012 | 8 | 4881959.5 | 4474445 | 3771633 | 4903333 | 5279293.5 | 5640946.5 | 4613032.5 | 5875659 | 5709454 | 5695903 | 3 | **1.181** |
| Q80TR8 | Dcaf1 | DDB1- and CUL4-associated factor 1 (Mus musculus) | 47.41 | 0.6 | 168931 | 1 | 606925.44 | 289005.59 | 423250.16 | 0 | 0 | 444783.22 | 413032.56 | 526017.5 | 0 | 0 | 15 | **1.049** |
| Q80TY0 | Fnbp1 | Formin-binding protein 1 (Mus musculus) | 670.03 | 20.13 | 71344 | 11 | 20915946 | 22780172 | 22787584 | 16423364 | 20589134 | 21629572 | 17759816 | 21009336 | 26147232 | 23353312 | 8 | **1.062** |
| Q80U72 | Scrib | Protein scribble homolog (Mus musculus) | 98.25 | 3.04 | 174059 | 3 | 1713180.38 | 589233.62 | 1581590.12 | 1692254 | 1685204.25 | 1395796 | 1471204 | 1494200.38 | 1643792 | 1415842.25 | 6 | **1.022** |
| Q80U87 | Usp8 | Ubiquitin carboxyl-terminal hydrolase 8 (Mus musculus) | 171.53 | 4.44 | 122611 | 4 | 2197891 | 1627862.5 | 1744086.75 | 1213044.62 | 0 | 2338775.25 | 2199139.25 | 2488814 | 1839140.12 | 1861559.5 | 2 | **1.582** |
| Q80U93 | Nup214 | Nuclear pore complex protein Nup214 (Mus musculus) | 260.26 | 3.36 | 212979 | 5 | 4608723.5 | 1139355.12 | 1998658.25 | 1640193.62 | 2734084.25 | 3522633.5 | 2864488 | 3188048 | 3902747.75 | 3571451.75 | 5 | **1.407** |
| Q80UG5 | Sept9 | Septin-9 (Mus musculus) | 374.74 | 15.61 | 65575 | 8 | 15269423 | 19812600 | 18373376 | 18339718 | 15551954 | 20697624 | 18185972 | 19262038 | 19960456 | 22248832 | 4 | **1.149** |
| Q80UJ7 | Rab3gap1 | Rab3 GTPase-activating protein catalytic subunit (Mus musculus) 1 | 317.32 | 9.48 | 110198 | 7 | 2375794.25 | 5972416.5 | 6683389.5 | 5953429 | 3767286.75 | 5681385 | 6012655.5 | 6135587.5 | 5797303 | 6585556.5 | 9 | **1.221** |
| Q80UM3 | Naa15 | N-alpha-acetyltransferase 15, NatA auxiliary subunit (Mus musculus) | 870.61 | 19.65 | 100961 | 17 | 15784429 | 32750048 | 30684406 | 39761072 | 26599764 | 20026796 | 15490138 | 21319550 | 22377006 | 25912092 | 4 | **0.722** |
| Q80UM7 | Mogs | Mannosyl-oligosaccharide glucosidase (Mus musculus) | 311.31 | 9.23 | 91831 | 5 | 7347752.5 | 5530353.5 | 4912012.5 | 5951837 | 5159473.5 | 5927950 | 6178643.5 | 5399464.5 | 5327486.5 | 5214195.5 | 12 | **0.970** |
| Q80UY2 | Kcmf1 | E3 ubiquitin-protein ligase KCMF1 (Mus musculus) | 134.47 | 8.14 | 41791 | 2 | 114636.93 | 450420.5 | 419232.88 | 366416.09 | 389317.88 | 343049.06 | 369753.66 | 349429.22 | 392708.69 | 326375.19 | 8 | **1.024** |
| Q80VA0 | Galnt7 | N-acetylgalactosaminyltransferase 7 (Mus musculus) | 41.05 | 1.67 | 75419 | 1 | 443468.22 | 1079684.5 | 703100.19 | 1398305.5 | 1103815.75 | 1015441.12 | 466538.97 | 1054163.75 | 1057692.75 | 1302404.25 | 11 | **1.036** |
| Q80VD1 | Fam98b | Protein FAM98B (Mus musculus) | 52.93 | 3.96 | 45349 | 1 | 142871.61 | 421646.75 | 52524.76 | 258614.17 | 352393.31 | 37592.82 | 255234.52 | 280513.72 | 284833.5 | 243655.62 | 10 | **0.897** |
| Q80W54 | Zmpste24 | CAAX prenyl protease 1 homolog Mus musculus | 48.35 | 2.32 | 54735 | 1 | 289725.62 | 670349.06 | 835199.25 | 791070.19 | 50150.98 | 287983.91 | 719632.06 | 588721.19 | 667458.81 | 52187.44 | 9 | **0.878** |
| Q80WJ7 | Mtdh | Protein LYRIC (Mus musculus) | 586.64 | 27.46 | 63846 | 10 | 17286944 | 10653678 | 8092196 | 6948055 | 12282488 | 7503656 | 12870029 | 10631050 | 11219143 | 8500972 | 12 | **0.918** |
| Q80WQ2 | Vac14 | Protein VAC14 homolog (Mus musculus) | 596.08 | 23.27 | 88048 | 12 | 4032363.75 | 8469553 | 7265155 | 8463176 | 7108478 | 5502694.5 | 5459020.5 | 8634659 | 8095667.5 | 6199867.5 | 11 | **0.959** |
| Q80WS3 | Fbll1 | rRNA/tRNA 2'-O-methyltransferase fibrillarin-like protein 1 (Mus musculus) | 138.8 | 11.46 | 33339 | 3 | 15148749 | 14759404 | 7323840.5 | 8346900 | 12553787 | 19241234 | 14336966 | 11755900 | 10246971 | 10465844 | 11 | **1.136** |
| Q80WW9 | Ddrgk1 | DDRGK domain-containing protein 1 (Mus musculus) | 87.3 | 5.4 | 35977 | 1 | 411739.62 | 821075.44 | 887087.06 | 1003863.44 | 854464.25 | 486039.66 | 46764.76 | 534509.81 | 1004041.44 | 633977.44 | 8 | **0.680** |
| Q80X41 | Vrk1 | Serine/threonine-protein kinase VRK1 (Mus musculus) | 89.87 | 5 | 49741 | 2 | 3177324.25 | 4766319.5 | 5332888.5 | 3965845 | 4844155 | 5718415 | 5070773 | 4637416.5 | 4771011 | 5023494.5 | 7 | **1.142** |
| Q80X50 | Ubap2l | Ubiquitin-associated protein 2-like (Mus musculus) | 432.14 | 7.59 | 116799 | 6 | 21480638 | 16359374 | 13572936 | 14582412 | 18643864 | 19450552 | 20050618 | 19126106 | 14718647 | 17662792 | 8 | **1.075** |
| Q80X82 | Sympk | Symplekin (Mus musculus) | 216.4 | 4.91 | 142284 | 4 | 760999.19 | 2835531.5 | 2960656.75 | 2339264.5 | 1955705.25 | 1696257.38 | 2244674 | 2258121.75 | 1453375.25 | 2378026.25 | 9 | **0.924** |
| Q80XI3 | Eif4g3 | Eukaryotic translation initiation factor 4 gamma 3 (Mus musculus) | 392.64 | 5.95 | 174890 | 8 | 17253344 | 17629468 | 17385840 | 18916544 | 17912190 | 18517566 | 10419470 | 18153672 | 19057738 | 19201944 | 7 | **0.958** |
| Q80XP9 | Wnk3 | Serine/threonine-protein kinase WNK3 (Mus musculus) | 115.29 | 1.65 | 193991 | 3 | 1022803.38 | 759475.69 | 1656417.5 | 892422.62 | 2213650 | 958016.25 | 2446953 | 626176.44 | 2615901.75 | 916904.94 | 11 | **1.156** |
| Q80XQ2 | Tbc1d5 | TBC1 domain family member 5 (Mus musculus) | 50.66 | 1.35 | 91837 | 1 | 1056527.25 | 1131692.25 | 1007833.5 | 912120.75 | 868380.75 | 752737.44 | 1023331.06 | 940367.88 | 1205792 | 1142857.62 | 10 | **1.018** |
| Q80Y14 | Glrx5 | Glutaredoxin-related protein 5, mitochondrial (Mus musculus) | 47.77 | 7.89 | 16292 | 1 | 1015601.69 | 0 | 744169.94 | 585456.12 | 415248.97 | 876935.56 | 1326746.62 | 523018.75 | 929344.25 | 812549.5 | 6 | **1.619** |
| Q80Y44 | Ddx10 | Probable ATP-dependent RNA helicase DDX10 (Mus musculus) | 75.78 | 2.97 | 100739 | 2 | 358676.66 | 373865.91 | 408017.25 | 346551.16 | 264444.88 | 297485.78 | 428672.75 | 364804.56 | 389117.34 | 347909.97 | 10 | **1.044** |
| Q80Y55 | Bsdc1 | BSD domain-containing protein 1 (Mus musculus) | 99.33 | 7.26 | 46953 | 2 | 287258.94 | 0 | 260652.27 | 695694.38 | 621259.5 | 171816.59 | 236637.12 | 346577.34 | 667896.75 | 178208.41 | 10 | **0.859** |
| Q80Y81 | Elac2 | Zinc phosphodiesterase ELAC protein 2 (Mus musculus) | 154.79 | 4.93 | 92719 | 3 | 648142.25 | 1265463.5 | 602818.12 | 2082322.88 | 1538325.88 | 670703.56 | 805156 | 1655296.38 | 1423698.75 | 1846151.25 | 10 | **1.043** |
| Q80YW0 | Cyth4 | Cytohesin-4 (Mus musculus) | 320.47 | 19.08 | 45285 | 6 | 4434418.5 | 3688656.5 | 4720602.5 | 3821218.5 | 4304535.5 | 4500536 | 4470701 | 4737262 | 5179209 | 5057384.5 | 2 | **1.142** |
| Q80Z70 | Sel1l | Protein sel-1 homolog 1 (Rattus norvegicus) | 56.16 | 1.39 | 88683 | 1 | 274354.62 | 694313.44 | 600642.12 | 662023.38 | 647626.19 | 536741.12 | 655498.75 | 349396.28 | 652578.94 | 323466.94 | 9 | **0.875** |
| Q80ZF7 | Rdh10 | Retinol dehydrogenase 10 (Rattus norvegicus) | 43.26 | 2.35 | 38061 | 1 | 622941.62 | 760759.62 | 645175.81 | 719206.31 | 548724.62 | 711211.31 | 630514.62 | 646911.62 | 855317.56 | 649914.44 | 9 | **1.060** |
| Q810A7 | Ddx42 | ATP-dependent RNA helicase DDX42 (Mus musculus) | 271.65 | 9.26 | 101965 | 5 | 5024733 | 3620294 | 6285649 | 5653363.5 | 5995461 | 4433727 | 3386142.25 | 5512055 | 6312112 | 6097408 | 12 | **0.968** |
| Q810B6 | Ankfy1 | Rabankyrin-5 (Mus musculus) | 498.66 | 10.27 | 128653 | 10 | 6148351 | 7346488 | 8980431 | 8360902.5 | 6762369 | 8172353 | 8559922 | 8252230 | 8010322 | 6360967.5 | 11 | **1.047** |
| Q810V0 | Mphosph10 | U3 small nucleolar ribonucleoprotein protein MPP10 (Mus musculus) 10 | 59.35 | 2.79 | 78735 | 1 | 0 | 1065882.25 | 0 | 1136417.25 | 1355985.38 | 0 | 0 | 0 | 910627.44 | 0 | 16 | **0.256** |
| Q810Y8 | Pramel7 | Preferentially expressed antigen in melanoma-like protein 7 (Mus musculus) | 48.14 | 1.46 | 55834 | 1 | 559665.88 | 1514801.12 | 426979.25 | 1760853 | 1002354.31 | 526118 | 396285.47 | 340689.91 | 360599.78 | 392073.09 | 1 | **0.383** |
| Q8BFQ4 | Wdr82 | WD repeat-containing protein 82 (Mus musculus) | 169.43 | 10.22 | 35079 | 2 | 1427644 | 2393363.5 | 2761715 | 2432327.5 | 2851447 | 4430388.5 | 3322999 | 3596114.5 | 2912888.5 | 2729337.5 | 2 | **1.432** |
| Q8BFQ8 | Gatd1 | Glutamine amidotransferase-like class 1 domain-containing protein 1 (Mus musculus) | 48.14 | 4.55 | 23277 | 1 | 1180692.25 | 1295204.88 | 1300452.75 | 1271740.25 | 1187332.12 | 1588939.12 | 1369094 | 1294271 | 1306583.88 | 1243630.88 | 5 | **1.091** |
| Q8BFR4 | Gns | N-acetylglucosamine-6-sulfatase (Mus musculus) | 323.3 | 11.58 | 61175 | 7 | 12993063 | 9778848 | 8773300 | 8682317 | 8672697 | 8461970 | 8216372.5 | 8494493 | 10212138 | 8704157 | 6 | **0.902** |
| Q8BFR5 | Tufm | Elongation factor Tu, mitochondrial (Mus musculus) | 301.19 | 18.58 | 49508 | 5 | 13183686 | 12703462 | 10854874 | 15238600 | 15100254 | 12273743 | 13037934 | 13274904 | 12953968 | 13415306 | 11 | **0.968** |
| Q8BFV2 | Pcid2 | PCI domain-containing protein 2 (Mus musculus) | 40.4 | 2.51 | 46132 | 1 | 0 | 811578.44 | 513601.19 | 501390.59 | 0 | 367279.81 | 0 | 399812.31 | 0 | 424384.41 | 11 | **0.652** |
| Q8BFY6 | Pef1 | Peflin (Mus musculus) | 40.81 | 4 | 29228 | 1 | 0 | 715097.25 | 794173.62 | 771001.12 | 550507.88 | 0 | 0 | 673606.31 | 797192.75 | 0 | 13 | **0.520** |
| Q8BFY9 | Tnpo1 | Transportin-1 (Mus musculus) | 1138.08 | 30.62 | 102357 | 21 | 39598964 | 41170592 | 39005996 | 38092828 | 38559760 | 39333812 | 40527452 | 43019848 | 39196100 | 42807648 | 5 | **1.043** |
| Q8BFZ9 | Erlin2 | Erlin-2 (Mus musculus) | 236.19 | 21.18 | 37873 | 5 | 4597689 | 4046860 | 2316891.5 | 4339051 | 4413910 | 2988999.25 | 3367805.5 | 3376346.5 | 2066231.5 | 1868932.62 | 3 | **0.693** |
| Q8BG07 | Pld4 | Phospholipase D4 (Mus musculus) | 762.2 | 35.59 | 56154 | 13 | 35268428 | 33078132 | 34448792 | 24068848 | 38366792 | 34450000 | 35744668 | 33074404 | 34941832 | 35327304 | 10 | **1.050** |
| Q8BG30 | Nelfa | Negative elongation factor A (Mus musculus) | 39.21 | 1.89 | 57585 | 1 | 0 | 117381.76 | 203212.86 | 455695.47 | 476055.44 | 221053.45 | 302461.53 | 248518.36 | 258770.94 | 272402.22 | 10 | **1.041** |
| Q8BG32 | Psmd11 | 26S proteasome non-ATPase regulatory subunit 11 (Mus musculus) | 1198.88 | 54.03 | 47437 | 20 | 84287536 | 68544256 | 82486072 | 56370148 | 67489192 | 92130992 | 79421688 | 89508904 | 79969672 | 83005208 | 5 | **1.181** |
| Q8BG48 | Stk17b | Serine/threonine-protein kinase 17B (Mus musculus) | 56.88 | 3.76 | 41983 | 1 | 0 | 518930.28 | 1008849.44 | 864747.12 | 990752.44 | 0 | 0 | 0 | 716305.94 | 653952 | 10 | **0.405** |
| Q8BG51 | Rhot1 | Mitochondrial Rho GTPase 1 (Mus musculus) | 113.23 | 4.44 | 72242 | 2 | 0 | 257161.61 | 66405.03 | 164873.89 | 147953.25 | 252518.27 | 0 | 312086.84 | 0 | 361816.06 | 11 | **1.456** |
| Q8BG73 | Sh3bgrl2 | SH3 domain-binding glutamic acid-rich-like protein 2 (Mus musculus) 2 | 127.73 | 24.3 | 12255 | 2 | 2853648 | 5503475 | 6004725 | 6200165.5 | 6029693 | 6392381 | 5278633.5 | 5775646 | 5754132.5 | 5139671.5 | 11 | **1.066** |
| Q8BGC4 | Zadh2 | Prostaglandin reductase-3 (Mus musculus) | 44.12 | 7.69 | 40529 | 1 | 355237.62 | 0 | 439520.06 | 129358.23 | 198947.09 | 227092.8 | 331856.41 | 276144.5 | 0 | 278634.44 | 13 | **0.992** |
| Q8BGD9 | Eif4b | Eukaryotic translation initiation factor 4B (Mus musculus) | 1102.82 | 33.06 | 68840 | 16 | 42670844 | 33034790 | 33688512 | 34157880 | 37956024 | 33212372 | 25436738 | 30534062 | 33974548 | 34820352 | 6 | **0.870** |
| Q8BGE6 | Atg4b | Cysteine protease ATG4B (Mus musculus) | 279.81 | 15.78 | 44375 | 4 | 847903.69 | 1683305.12 | 1797061.88 | 1867327.62 | 1107488.25 | 1915546.25 | 2026547.62 | 2135695.75 | 2327042.75 | 2124955.25 | 0 | **1.442** |
| Q8BGH2 | Samm50 | Sorting and assembly machinery component 50 homolog (Mus musculus) | 328.51 | 18.98 | 51864 | 8 | 14073690 | 16473099 | 16648828 | 18750782 | 18399028 | 15263902 | 12886869 | 21369128 | 21597020 | 16870750 | 11 | **1.043** |
| Q8BGQ7 | Aars | Alanine--tRNA ligase, cytoplasmic (Mus musculus) | 1749.64 | 42.87 | 106909 | 26 | 62273436 | 52460684 | 53172208 | 58509712 | 48221496 | 57230656 | 61026772 | 59500416 | 54860072 | 58553844 | 7 | **1.060** |
| Q8BGS7 | Cept1 | Choline/ethanolaminephosphotransferase 1 (Mus musculus) | 109.52 | 8.89 | 46434 | 2 | 576917.44 | 823944.12 | 744015.25 | 737564.19 | 161847.77 | 509019.53 | 794920.5 | 795760.44 | 738794.38 | 646443.62 | 11 | **1.145** |
| Q8BGV0 | Nars2 | Probable asparagine--tRNA ligase, mitochondrial (Mus musculus) | 43.7 | 6.08 | 53962 | 1 | 0 | 202721.64 | 214386.22 | 135571.5 | 238912.53 | 0 | 0 | 360288.16 | 344675.5 | 289274.12 | 10 | **1.256** |
| Q8BH04 | Pck2 | Phosphoenolpyruvate carboxykinase [GTP], mitochondrial (Mus musculus) | 859.86 | 30.16 | 70528 | 14 | 34131352 | 20925378 | 22733480 | 14336425 | 22654450 | 30956812 | 27027758 | 27198118 | 23978072 | 23063206 | 5 | **1.152** |
| Q8BH43 | Wasf2 | Wiskott-Aldrich syndrome protein family member 2 (Mus musculus) | 376.08 | 15.29 | 54074 | 6 | 11136790 | 8679173 | 9698083 | 8315517.5 | 8800363 | 11703187 | 10872095 | 11471886 | 10712848 | 10135500 | 3 | **1.177** |
| Q8BH59 | Slc25a12 | Calcium-binding mitochondrial carrier protein Aralar1 (Mus musculus) 2 | 1140.04 | 45.94 | 74570 | 21 | 38173864 | 31799552 | 29758806 | 38213600 | 31049336 | 31422974 | 33264672 | 30633894 | 30679418 | 32085138 | 10 | **0.935** |
| Q8BH61 | F13a1 | Coagulation factor XIII A chain (Mus musculus) | 481.3 | 17.62 | 83207 | 10 | 12713783 | 14438145 | 16586800 | 20758680 | 15574946 | 12513140 | 12625392 | 13079383 | 14386617 | 14088937 | 3 | **0.833** |
| Q8BH69 | Sephs1 | Selenide, water dikinase 1 (Mus musculus) | 211.5 | 10.2 | 42907 | 4 | 1482913 | 3431656.25 | 4163668 | 1956520.75 | 3820937.25 | 5305862 | 1824202 | 4339734 | 4365997.5 | 4030457.25 | 5 | **1.337** |
| Q8BH74 | Nup107 | Nuclear pore complex protein Nup107 (Mus musculus) | 288.05 | 6.05 | 106717 | 5 | 2275971.5 | 3807937.5 | 3972698.75 | 4219159.5 | 3538102.25 | 3640179.75 | 3666207.75 | 4118619.25 | 3865971.75 | 4058834.5 | 10 | **1.086** |
| Q8BH86 | Dglucy | D-glutamate cyclase, mitochondrial (Mus musculus) | 58.93 | 3.24 | 66366 | 1 | 0 | 602446.75 | 673361.88 | 532025.06 | 0 | 0 | 711351.69 | 413853.12 | 475087.72 | 569227.25 | 14 | **1.200** |
| Q8BHB4 | Wdr3 | WD repeat-containing protein 3 (Mus musculus) | 200.76 | 4.88 | 105775 | 4 | 2715784 | 3429874 | 3956322 | 2058313.62 | 1854183.12 | 3123944.75 | 3639702 | 3148194.75 | 1972915.75 | 3133443 | 11 | **1.072** |
| Q8BHC4 | Dcakd | Dephospho-CoA kinase domain-containing protein (Mus musculus) | 315.95 | 37.23 | 26476 | 7 | 8029793 | 6904150.5 | 8784550 | 5852678 | 5972169.5 | 6892639.5 | 6027466 | 7876737 | 7862331 | 6691566 | 12 | **0.995** |
| Q8BHD7 | Ptbp3 | Polypyrimidine tract-binding protein 3 (Mus musculus) | 438.29 | 17.4 | 56701 | 6 | 10382926 | 4683150 | 5334246.5 | 4634813.5 | 6762535.5 | 7804817.5 | 8199436 | 7722081 | 8345943 | 7018818 | 5 | **1.229** |
| Q8BHJ5 | Tbl1xr1 | F-box-like/WD repeat-containing protein TBL1XR1 (Mus musculus) | 171.99 | 10.31 | 55661 | 3 | 3567063.5 | 4859021 | 4834836 | 2366527.25 | 2075910.12 | 5765001.5 | 2489050.5 | 3313488 | 3001635 | 2691621.75 | 12 | **0.975** |

| | | | | | | | | | | | | | | | | | | |
|---|---|---|---|---|---|---|---|---|---|---|---|---|---|---|---|---|---|---|
| Q8BHL5 | Elmo2 | Engulfment and cell motility protein 2 (Mus musculus) | 139.18 | 7.38 | 83887 | 4 | 1059260.25 | 2679162.75 | 2318553 | 2574442.25 | 1511537.75 | 1111963.75 | 921047.19 | 1106150.5 | 896623.19 | 979580.75 | 2 | **0.494** |
| Q8BHL8 | Psmf1 | Proteasome inhibitor PI31 subunit (Mus musculus) | 87.73 | 4.43 | 29664 | 1 | 6212180.5 | 3845476.5 | 4679232 | 2624085 | 3243682 | 2992348.25 | 3714948 | 2602887.75 | 2927081.5 | 2986512.5 | 5 | **0.739** |
| Q8BHN3 | Ganab | Neutral alpha-glucosidase AB (Mus musculus) | 1191.52 | 28.92 | 106911 | 23 | 46779320 | 60615112 | 57171204 | 54700000 | 55713324 | 67828488 | 62007924 | 62550728 | 68005496 | 60189768 | 1 | **1.166** |
| Q8BHZ0 | Fam49a | Protein FAM49A (Mus musculus) | 202.71 | 14.86 | 37343 | 4 | 6829890 | 8070093 | 7593108 | 6424803 | 6130577 | 5626125 | 6897279.5 | 7155953.5 | 6944787.5 | 7451930 | 12 | **0.972** |
| Q8BI84 | Mia3 | Transport and Golgi organization protein 1 homolog (Mus musculus) | 86.36 | 1.45 | 213675 | 2 | 712883.69 | 2257234.25 | 1806186.75 | 1363883 | 1659505.5 | 1695602.25 | 1218998 | 1668619 | 759276.19 | 1605776 | 10 | **0.891** |
| Q8BIG7 | Comtd1 | Catechol O-methyltransferase domain-containing protein 1 (Mus musculus) | 117.26 | 22.14 | 28961 | 3 | 1725886.5 | 1313972.25 | 1300493.5 | 1569667 | 1655596 | 1383079.38 | 533135.38 | 1661302.25 | 1705349.25 | 1182646.75 | 10 | **0.855** |
| Q8BIJ6 | Iars2 | Isoleucine--tRNA ligase, mitochondrial (Mus musculus) | 626.66 | 16.21 | 112804 | 11 | 13700113 | 16684297 | 16526144 | 16060119 | 14898928 | 13941510 | 14251788 | 16342995 | 17466048 | 16578646 | 11 | **1.009** |
| Q8BIJ7 | Rufy1 | RUN and FYVE domain-containing protein 1 (Mus musculus) | 120.72 | 4.35 | 80376 | 3 | 3199382.25 | 2563904.5 | 2724615.75 | 2684632 | 3021813 | 1905232.75 | 2436750 | 1846024.5 | 1654439.75 | 1865970.75 | 0 | **0.684** |
| Q8BIW1 | Prune1 | Exopolyphosphatase PRUNE1 (Mus musculus) | 43.26 | 3.74 | 50239 | 1 | 380785.75 | 0 | 219637.53 | 0 | 425675.47 | 0 | 0 | 0 | 241064.58 | 253569.41 | 15 | **0.482** |
| Q8BJ48 | Nagpa | N-acetylglucosamine-1-phosphodiester alpha-N-acetylglucosaminidase (Mus musculus) | 262.13 | 8.32 | 56044 | 3 | 1634260.38 | 1722636.25 | 1117834.25 | 1020637.62 | 1557874.38 | 1031101.56 | 2218985.25 | 2100252.5 | 2063283 | 1942627.75 | 4 | **1.327** |
| Q8BJ71 | Nup93 | Nuclear pore complex protein Nup93 (Mus musculus) | 572.41 | 19.41 | 93281 | 10 | 15239164 | 16485206 | 14927002 | 15901748 | 16140615 | 13674236 | 14147587 | 15603232 | 14229367 | 17303086 | 7 | **0.953** |
| Q8BJS4 | Sun2 | SUN domain-containing protein 2 (Mus musculus) | 242.12 | 6.7 | 81605 | 4 | 664333.38 | 1315169.12 | 1938989.5 | 1235166.12 | 1263797.75 | 1410481.38 | 727308.25 | 2060254.5 | 953334.31 | 1735717.25 | 10 | **1.073** |
| Q8BJW5 | Nol11 | Nucleolar protein 11 (Mus musculus) | 249.1 | 7.61 | 80818 | 5 | 4230473 | 4246139.5 | 4207219.5 | 4359728 | 2584962.5 | 4118882 | 3977244.5 | 3960308.25 | 3892284.5 | 3686776.25 | 5 | **1.000** |
| Q8BJW6 | Eif2a | Eukaryotic translation initiation factor 2A (Mus musculus) | 284.93 | 13.25 | 64403 | 5 | 8288275.5 | 4986399.5 | 4650503 | 3321167 | 5721086 | 6072634.5 | 5655317 | 5323815 | 5347226 | 5221831 | 9 | **1.024** |
| Q8BJY1 | Psmd5 | 26S proteasome non-ATPase regulatory subunit 5 (Mus musculus) | 700.4 | 32.74 | 55972 | 13 | 18239960 | 13698720 | 15219554 | 12831861 | 14244302 | 12661436 | 14230265 | 17382864 | 16603165 | 13207090 | 11 | **0.998** |
| Q8BJZ4 | Mrps35 | 28S ribosomal protein S35, mitochondrial (Mus musculus) | 45.82 | 4.06 | 35975 | 1 | 580721.38 | 845921.19 | 885239.81 | 552994.75 | 655426.38 | 614826.62 | 842800.69 | 696185.31 | 632664.06 | 587484.75 | 12 | **0.958** |
| Q8BK64 | Ahsa1 | Activator of 90 kDa heat shock protein ATPase homolog 1 (Mus musculus) | 450.99 | 34.62 | 38117 | 9 | 17389434 | 31464522 | 29523276 | 29954456 | 22956226 | 19965136 | 23920764 | 24275374 | 24602118 | 22531658 | 8 | **0.878** |
| Q8BK67 | Rcc2 | Protein RCC2 (Mus musculus) | 1292.8 | 58.08 | 55983 | 22 | 69131120 | 65656116 | 62334928 | 58170664 | 104597568 | 72215448 | 58943844 | 67117376 | 61921140 | 66521744 | 12 | **0.908** |
| Q8BK72 | Mrps27 | 28S ribosomal protein S27, mitochondrial (Mus musculus) | 127.93 | 8.19 | 47779 | 3 | 1722513.88 | 2927081 | 2297690.5 | 3912309 | 2225373.75 | 2431347.5 | 1919750 | 2448159.5 | 1391002.75 | 2385625.5 | 10 | **0.808** |
| Q8BKC5 | Ipo5 | Importin-5 (Mus musculus) | 1660.71 | 38.83 | 123591 | 28 | 26398810 | 45463464 | 48447084 | 49557060 | 40404624 | 45267640 | 41930784 | 45533920 | 42893024 | 42533844 | 11 | **1.038** |
| Q8BKE6 | Cyp20a1 | Cytochrome P450 20A1 (Mus musculus) | 102.24 | 5.63 | 52149 | 2 | 1294498.12 | 1688746 | 985344.94 | 3167260 | 1408283.5 | 873211.38 | 711990.31 | 825666.81 | 1419437.5 | 1714908.75 | 7 | **0.649** |
| Q8BKS9 | Pum3 | Pumilio homolog 3 (Mus musculus) | 167.6 | 6.18 | 72800 | 3 | 0 | 949088.19 | 950274.5 | 1822290 | 1238372.5 | 420212.22 | 575664.38 | 811524.12 | 632226.88 | 648365.5 | 5 | **0.623** |
| Q8BKT3 | Gcfc2 | GC-rich sequence DNA-binding factor 2 (Mus musculus) | 89.4 | 2.47 | 87442 | 2 | 1501527 | 1151306.25 | 1202682.38 | 1440867.88 | 2850683.5 | 1545476.38 | 1620480.25 | 1330649.25 | 1494263 | 1453629.75 | 9 | **0.914** |
| Q8BKZ9 | Pdhx | Pyruvate dehydrogenase protein X component, mitochondrial (Mus musculus) | 72.71 | 3.39 | 53999 | 2 | 1429654.12 | 3015851 | 2380540 | 2606071 | 2670064.25 | 1601714 | 1570636.25 | 2589244 | 4092733 | 2583454.25 | 11 | **1.028** |
| Q8BL66 | Eea1 | Early endosome antigen 1 (Mus musculus) | 318.18 | 5.03 | 160915 | 6 | 36140980 | 9342870 | 9188473 | 6484792 | 7334677 | 11461674 | 12326484 | 11386452 | 10922858 | 11631029 | 5 | **0.843** |
| Q8BL97 | Srsf7 | Serine/arginine-rich splicing factor 7 (Mus musculus) | 191.01 | 16.48 | 30818 | 4 | 20577728 | 20369808 | 14241044 | 23487058 | 23429204 | 14534238 | 12316406 | 12556554 | 13309783 | 13546492 | 1 | **0.649** |
| Q8BLF1 | Nceh1 | Neutral cholesterol ester hydrolase 1 (Mus musculus) | 651.53 | 40.69 | 45740 | 11 | 5611267 | 27819202 | 32706882 | 24945496 | 15577125 | 29315426 | 31038390 | 25940948 | 29268866 | 32132486 | 6 | **1.385** |
| Q8BLN5 | Lss | Lanosterol synthase (Mus musculus) | 107.93 | 3.68 | 83141 | 2 | 897587.25 | 0 | 0 | 0 | 758017.81 | 1285520.75 | 771313.81 | 888930.81 | 877196.69 | 0 | 11 | **2.309** |
| Q8BMA6 | Srp68 | Signal recognition particle subunit SRP68 (Mus musculus) | 585.88 | 20.32 | 70574 | 10 | 9757512 | 12521727 | 12767195 | 11711125 | 8518076 | 10243191 | 12367617 | 12182344 | 13277793 | 12536366 | 8 | **1.096** |
| Q8BMD8 | Slc25a24 | Calcium-binding mitochondrial carrier protein SCaMC-1 (Mus musculus) 4 | 85.71 | 4.42 | 52902 | 2 | 3091811.75 | 2470227.75 | 2115855 | 2181599.5 | 2108379.25 | 2856257.25 | 3603355.75 | 2768635.25 | 3196243 | 2931327.5 | 3 | **1.283** |
| Q8BMF4 | Dlat | Dihydrolipoyllysine-residue acetyltransferase component of pyruvate dehydrogenase comple | 629.52 | 26.48 | 67942 | 11 | 48982736 | 49848232 | 44104200 | 65677960 | 51538776 | 38093992 | 40124152 | 40994576 | 46735536 | 42522808 | 1 | **0.801** |
| Q8BMG7 | Rab3gap2 | Rab3 GTPase-activating protein non-catalytic subunit (Mus musculus) 2 | 511.8 | 10.54 | 152535 | 11 | 12359269 | 10703619 | 11634493 | 11218653 | 6809260.5 | 11739132 | 11779181 | 10064339 | 11467429 | 10457572 | 12 | **1.053** |
| Q8BMJ2 | Lars | Leucine--tRNA ligase, cytoplasmic (Mus musculus) | 1587.36 | 30.22 | 134192 | 29 | 35931644 | 38235800 | 38966076 | 31734684 | 42145420 | 33897032 | 37946208 | 39800968 | 38413328 | 46584852 | 10 | **1.051** |
| Q8BMJ3 | Eif1ax | Eukaryotic translation initiation factor 1A, X-chromosomal (Mus musculus) | 350.66 | 33.33 | 16460 | 6 | 9601984 | 13471040 | 12650149 | 17031380 | 9368074 | 10267392 | 8373716 | 10045540 | 9659754 | 9590153 | 7 | **0.772** |
| Q8BMK4 | Ckap4 | Cytoskeleton-associated protein 4 (Mus musculus) | 1255.31 | 40.7 | 63692 | 19 | 27352868 | 31204870 | 30225846 | 29251008 | 27919638 | 27728038 | 29834356 | 30633982 | 30722740 | 29519294 | 10 | **1.017** |
| Q8BML9 | Qars | Glutamine--tRNA ligase (Mus musculus) | 553.6 | 16 | 87677 | 11 | 20976476 | 14701501 | 13635880 | 13724112 | 13798585 | 15116279 | 15690403 | 17760348 | 17023440 | 17097866 | 5 | **1.076** |
| Q8BMP6 | Acbd3 | Golgi resident protein GCP60 (Mus musculus) | 395.27 | 18.86 | 60181 | 6 | 14358169 | 7611865 | 6124310 | 6212337.5 | 5690857 | 4935023 | 6949018.5 | 4620421 | 5126822.5 | 4390598 | 3 | **0.651** |
| Q8BMS1 | Hadha | Trifunctional enzyme subunit alpha, mitochondrial (Mus musculus) | 841.74 | 26.74 | 82670 | 14 | 32748184 | 25368796 | 19637788 | 21233828 | 23900800 | 18005972 | 22835450 | 22382492 | 25755838 | 21486400 | 10 | **0.899** |
| Q8BN21 | Vrk2 | Serine/threonine-protein kinase VRK2 (Mus musculus) | 151.71 | 8.15 | 58119 | 3 | 1894329.75 | 1875576.62 | 2449994 | 1841781 | 1872859.75 | 2281688.25 | 2531973.5 | 1669658.5 | 1683316.5 | 1965872 | 12 | **1.020** |
| Q8BP27 | Sfr1 | Swi5-dependent recombination DNA repair protein 1 homolog (Mus musculus) | 53.67 | 3.76 | 35183 | 1 | 1155722.25 | 1026705.94 | 1442718.5 | 1038660.81 | 958218.44 | 1030540.06 | 813909.12 | 1108196.38 | 626974.31 | 1266624.25 | 9 | **0.862** |
| Q8BP40 | Acp6 | Lysophosphatidic acid phosphatase type 6 (Mus musculus) | 84.41 | 10.77 | 47625 | 2 | 334788.06 | 772199.31 | 0 | 0 | 67512.53 | 300301.84 | 667090.06 | 467767.5 | 507390.31 | 696574.88 | 7 | **2.247** |
| Q8BP47 | Nars | Asparagine--tRNA ligase, cytoplasmic (Mus musculus) | 1009.49 | 37.75 | 64279 | 19 | 45358020 | 57544516 | 53404236 | 52913768 | 50443980 | 67020892 | 55861628 | 67125168 | 60391128 | 66469680 | 1 | **1.220** |
| Q8BP48 | Metap1 | Methionine aminopeptidase 1 (Mus musculus) | 198.26 | 13.21 | 43221 | 4 | 8160583 | 5526571 | 2415235.25 | 3977371 | 6229797 | 7517363 | 2855359.75 | 7400464.5 | 4354810.5 | 7232086 | 10 | **1.116** |
| Q8BP92 | Rcn2 | Reticulocalbin-2 (Mus musculus) | 314.2 | 22.19 | 37271 | 4 | 1175125.12 | 4483165.5 | 2899440.75 | 4127694.5 | 2674610 | 2571595 | 1478788.75 | 1287987 | 2304042.25 | 2510829 | 5 | **0.661** |
| Q8BPA8 | Dpcd | Protein DPCD (Mus musculus) | 46.45 | 4.93 | 23034 | 1 | 262968.19 | 225307.36 | 302001.03 | 0 | 0 | 238006.3 | 177077.23 | 246608.12 | 250228.7 | 235761.97 | 12 | **1.452** |
| Q8BPU7 | Elmo1 | Engulfment and cell motility protein 1 (Mus musculus) | 377.08 | 14.58 | 83936 | 8 | 7983245 | 7277130 | 6455415 | 8203346 | 6792058 | 6902312.5 | 4481415 | 4178739.5 | 3468686.5 | 5653655 | 2 | **0.672** |
| Q8BPX9 | Slc15a3 | Solute carrier family 15 member 3 (Mus musculus) | 228.84 | 6.92 | 64051 | 4 | 11601103 | 4199966.5 | 3500977 | 3129776.5 | 4243716 | 7853122 | 5681777 | 7475991 | 6372523 | 4854175.5 | 5 | **1.209** |
| Q8BQ30 | Ppp1r18 | Phostensin (Mus musculus) | 141.29 | 7.91 | 65630 | 3 | 8803406 | 2992483.75 | 2692974.75 | 4977826 | 5917354 | 5408049.5 | 3753373.25 | 4544874 | 5162849 | 5062027.5 | 12 | **0.943** |
| Q8BQM4 | Heatr3 | HEAT repeat-containing protein 3 (Mus musculus) | 426.83 | 15.76 | 74305 | 9 | 8603242 | 7564890 | 7371899 | 7746691 | 7115301 | 6820442.5 | 7307206.5 | 7695399.5 | 7709946 | 7954072 | 11 | **0.976** |
| Q8BRT1 | Clasp2 | CLIP-associating protein 2 (Mus musculus) | 203.9 | 2.33 | 140739 | 3 | 1864144.75 | 1514934.38 | 1516501.88 | 835686.94 | 927732.38 | 1825078 | 1640010.75 | 1691751.25 | 859015.44 | 1882748 | 7 | **1.186** |
| Q8BSY0 | Asph | Aspartyl/asparaginyl beta-hydroxylase (Mus musculus) | 190.12 | 8.64 | 83042 | 4 | 2451310.08 | 3133471.75 | 914559.38 | 3133357.75 | 2773284.5 | 2936787.5 | 3466356.75 | 3558520 | 428286 | 3322259.5 | 6 | **1.344** |
| Q8BT07 | Cep55 | Centrosomal protein of 55 kDa (Mus musculus) | 53.92 | 2.81 | 53930 | 1 | 0 | 0 | 0 | 747228.69 | 607418.44 | 535868.69 | 484139.56 | 0 | 616170.75 | 589246.06 | 15 | **1.643** |
| Q8BT60 | Cpne3 | Copine-3 (Mus musculus) | 286.22 | 13.88 | 59585 | 6 | 5596008 | 7445646.5 | 9205044 | 7465642.5 | 6839168 | 8225616 | 6710400 | 6992378 | 7666504.5 | 7984012 | 10 | **1.028** |
| Q8BTI7 | Ankrd52 | Serine/threonine-protein phosphatase 6 regulatory ankyrin repeat subunit C (Mus musculus) | 85.16 | 1.21 | 115032 | 1 | 0 | 297656.69 | 226087.39 | 387601.16 | 253094.88 | 0 | 0 | 269656.03 | 244967.2 | 289077.16 | 11 | **0.690** |
| Q8BTI8 | Srrm2 | Serine/arginine repetitive matrix protein 2 (Mus musculus) | 218.06 | 2.22 | 294840 | 4 | 840778 | 1215966.38 | 1949703.5 | 1225781.12 | 1371627.75 | 768059.12 | 0 | 2259095.5 | 1728864.62 | 1824608.5 | 12 | **0.996** |
| Q8BTI9 | Pik3cb | Phosphatidylinositol 4,5-bisphosphate 3-kinase catalytic subunit beta isoform (Mus musculu | 39.04 | 0.66 | 121711 | 1 | 1596797.38 | 1155140.25 | 875780.25 | 1686547.25 | 1248761.38 | 823065.88 | 856347.75 | 838049.25 | 914807.44 | 985539 | 2 | **0.673** |
| Q8BTM8 | Flna | Filamin-A (Mus musculus) | 2594.71 | 24.56 | 281222 | 42 | 72252136 | 68382136 | 65112356 | 68257256 | 73262696 | 69192920 | 65007500 | 66198536 | 62131872 | 58017048 | 4 | **0.923** |
| Q8BTY2 | Slc4a7 | Sodium bicarbonate cotransporter 3 (Mus musculus) | 253.94 | 5.03 | 116514 | 5 | 2109903.25 | 2089166.62 | 2123523 | 2105377.75 | 2110024.25 | 1728562.12 | 1846547.5 | 2359097.25 | 1293359.38 | 1886051.62 | 5 | **0.865** |
| Q8BTZ7 | Gmppb | Mannose-1-phosphate guanyltransferase beta (Mus musculus) | 84.7 | 5 | 39917 | 1 | 990787.81 | 3670341.5 | 3967042.75 | 686227.25 | 1056573.75 | 1370853.75 | 2042263.88 | 1070275 | 1056390.25 | 1139064.12 | 11 | **0.644** |
| Q8BU03 | Pwp2 | Periodic tryptophan protein 2 homolog (Mus musculus) | 123.36 | 4.9 | 102910 | 3 | 948666 | 0 | 430966.97 | 296387.59 | 998680.69 | 1040045.12 | 351920.19 | 1111034.25 | 980182.25 | 991308.12 | 5 | **1.673** |
| Q8BU14 | Sec62 | Translocation protein SEC62 (Mus musculus) | 131.41 | 7.54 | 45581 | 3 | 8074492 | 984614.31 | 5075467 | 2531317 | 4643463 | 6889533 | 5533914.5 | 6777934.5 | 6707489 | 7332048.5 | 5 | **1.560** |
| Q8BU30 | Iars | Isoleucine--tRNA ligase, cytoplasmic (Mus musculus) | 592.96 | 12.36 | 144271 | 13 | 17948480 | 22253348 | 24075342 | 21935262 | 23505060 | 22107774 | 20994044 | 22626514 | 23453292 | 22760496 | 12 | **1.020** |
| Q8BU33 | Ilvbl | Acetolactate synthase-like protein (Mus musculus) | 64.29 | 3.48 | 68156 | 1 | 75476.17 | 0 | 303869.16 | 0 | 258851.03 | 0 | 0 | 381734.66 | 403223.22 | 411153.47 | 11 | **1.874** |
| Q8BV13 | Cops7b | COP9 signalosome complex subunit 7b (Mus musculus) | 72.96 | 6.82 | 29689 | 1 | 133150.75 | 309589.44 | 479462.59 | 373347.91 | 549953.38 | 323909.69 | 510677.16 | 449001.06 | 640110.94 | 505441.56 | 7 | **1.316** |
| Q8BVE3 | Atp6v1h | V-type proton ATPase subunit H (Mus musculus) | 709.04 | 38.51 | 55855 | 13 | 11215336 | 19883238 | 17293836 | 18808336 | 14509520 | 18108998 | 17372360 | 15431959 | 17571524 | 18878802 | 10 | **1.069** |
| Q8BVF2 | Pdcl3 | Phosducin-like protein 3 (Mus musculus) | 368.86 | 35.42 | 27581 | 7 | 6259189.5 | 8894749 | 7158099.5 | 8364687 | 7615527 | 6705374.5 | 4862082 | 6716042 | 6946609 | 6706936.5 | 4 | **0.834** |
| Q8BVG4 | Dpp9 | Dipeptidyl peptidase 9 (Mus musculus) | 84.07 | 1.97 | 98001 | 2 | 739294.69 | 1550724.38 | 506264.12 | 940545 | 920988.62 | 1558866.5 | 1382170.25 | 519807.34 | 213053.22 | 1618040 | 10 | **1.136** |
| Q8BVI4 | Qdpr | Dihydropteridine reductase (Mus musculus) | 395.54 | 47.72 | 25570 | 7 | 17848860 | 11184300 | 13028854 | 10676685 | 11635310 | 11292790 | 12060230 | 11498744 | 10440394 | 10586035 | 7 | **0.868** |
| Q8BVK9 | Sp110 | Sp110 nuclear body protein (Mus musculus) | 571.59 | 29.89 | 50140 | 10 | 9673949 | 14279232 | 14636949 | 7978838 | 10834764 | 8637691 | 9291809 | 7946584 | 9953913 | 10686746 | 6 | **0.810** |
| Q8BVW3 | Trim14 | Tripartite motif-containing protein 14 (Mus musculus) | 45.51 | 2.5 | 49641 | 1 | 0 | 1028337 | 729456.69 | 1305506.12 | 911814.81 | 0 | 666636.62 | 771882 | 756316.25 | 0 | 8 | **0.552** |
| Q8BVY0 | Rsl1d1 | Ribosomal L1 domain-containing protein 1 (Mus musculus) | 542.93 | 19.91 | 50421 | 9 | 21438816 | 13637190 | 15433164 | 11122829 | 11199044 | 19253422 | 18939508 | 16090341 | 15666250 | 16407846 | 5 | **1.186** |
| Q8BWD8 | Cdk19 | Cyclin-dependent kinase 19 (Mus musculus) | 42.44 | 4.39 | 56570 | 1 | 0 | 0 | 202820.8 | 0 | 169571.19 | 72669.45 | 0 | 142274.05 | 115730.93 | 129168.59 | 15 | **1.235** |
| Q8BWQ6 | | UPF0505 protein C16orf62 homolog (Mus musculus) | 211.92 | 4.78 | 109077 | 4 | 1875683.12 | 4232212.5 | 3059493.5 | 4635512.5 | 1402911.62 | 2262677 | 2305866.25 | 2754369.75 | 2803126.5 | 2805709.75 | 10 | **0.850** |
| Q8BWU5 | Osgep | Probable tRNA N6-adenosine threonylcarbamoyltransferase (Mus musculus) | 78.54 | 4.78 | 36301 | 1 | 0 | 449809.12 | 0 | 540730.25 | 468205.41 | 514683.66 | 396205.47 | 394164.16 | 429843.78 | 404039.59 | 12 | **1.466** |
| Q8BWW4 | Larp4 | La-related protein 4 (Mus musculus) | 188.71 | 5.29 | 79763 | 3 | 4196153.5 | 2881933.75 | 4532198 | 5041117 | 2333243.5 | 3744389 | 3647534 | 6398548 | 5320217 | 6182019 | 6 | **1.332** |
| Q8BX09 | Rbbp5 | Retinoblastoma-binding protein 5 (Mus musculus) | 52.88 | 1.67 | 59098 | 1 | 909857.31 | 1052147.12 | 886310.44 | 0 | 1130503.62 | 1154042 | 1283298.25 | 1383076.88 | 898153.06 | 909644.19 | 6 | **1.415** |
| Q8BX17 | Gemin5 | Gem-associated protein 5 (Mus musculus) | 69.93 | 0.73 | 166592 | 1 | 494607.94 | 643484.44 | 612260.31 | 484647.09 | 581778.75 | 381161.38 | 510333.38 | 479482.97 | 506804.31 | 412268.72 | 4 | **0.813** |
| Q8BX70 | Vps13c | Vacuolar protein sorting-associated protein 13C (Mus musculus) | 1932.58 | 13.34 | 420089 | 37 | 35397764 | 44709992 | 47245744 | 47790944 | 41244824 | 41399924 | 45975328 | 52995528 | 51936444 | 54134364 | 5 | **1.139** |
| Q8BX94 | Osbpl2 | Oxysterol-binding protein-related protein 2 (Mus musculus) | 130.89 | 8.68 | 55384 | 3 | 1533271.88 | 1471411 | 2340738.5 | 1933830.25 | 1689509.25 | 2036461.25 | 1279711.88 | 1827338.25 | 2062222.25 | 1831628.88 | 11 | **1.008** |
| Q8BXQ2 | Pigt | GPI transamidase component PIG-T (Mus musculus) | 44.81 | 1.37 | 65705 | 1 | 852596.75 | 2093534.88 | 1960939 | 1493167.38 | 1821104.5 | 1758385.75 | 1535154.75 | 1416666.38 | 1192009.12 | 1400319.12 | 7 | **0.888** |
| Q8BXZ1 | Tmx3 | Protein disulfide-isomerase TMX3 (Mus musculus) | 99.28 | 4.61 | 51848 | 2 | 1703210.88 | 2402894.5 | 2407076.5 | 2674182.25 | 2014284.88 | 2094199.75 | 4628058 | 4138965.5 | 3752312.5 | 3890608 | 3 | **1.652** |
| Q8BY71 | Hat1 | Histone acetyltransferase type B catalytic subunit (Mus musculus) | 364.22 | 20.43 | 49278 | 6 | 8115951.5 | 6157283.5 | 6723733 | 5560174 | 5664764.5 | 7001075 | 7896259 | 8952502 | 8119933 | 7577889.5 | 3 | **1.227** |
| Q8BY87 | Usp47 | Ubiquitin carboxyl-terminal hydrolase 47 (Mus musculus) | 445.39 | 7.05 | 157455 | 8 | 2778283.5 | 4807879.5 | 5115685 | 6127119.5 | 5420240.5 | 5460834.5 | 3625323.5 | 5303121 | 5483406.5 | 4988821.5 | 11 | **1.025** |
| Q8BYA0 | Tbcd | Tubulin-specific chaperone D (Mus musculus) | 204.96 | 3.85 | 133321 | 4 | 1305800.5 | 2535895 | 2441845.75 | 3486272 | 2588863.5 | 3557945.5 | 3534424 | 3408697 | 3149222 | 3944446.5 | 2 | **1.424** |
| Q8BYB9 | Poglut1 | Protein O-glucosyltransferase 1 (Mus musculus) | 80.87 | 3.57 | 46379 | 1 | 0 | 538300.5 | 713341.19 | 0 | 628111.56 | 622328.81 | 0 | 644361.56 | 444637.38 | 647891.25 | 13 | **1.255** |
| Q8BYH7 | Tbc1d17 | TBC1 domain family member 17 (Mus musculus) | 61.3 | 1.55 | 72860 | 1 | 859110.69 | 905617.88 | 980876.12 | 0 | 0 | 1185083.12 | 1003750.25 | 1303463.5 | 0 | 0 | 11 | **1.272** |
| Q8BYI6 | Lpcat2 | Lysophosphatidylcholine acyltransferase 2 (Mus musculus) | 484.85 | 21.14 | 60254 | 10 | 7876710.5 | 10795904 | 12472595 | 15669913 | 10264017 | 11302852 | 9621987 | 11590735 | 10213912 | 12441201 | 11 | **0.967** |
| Q8BYL4 | Yars2 | Tyrosine--tRNA ligase, mitochondrial (Mus musculus) | 47.28 | 2.54 | 52597 | 1 | 79030.72 | 308279.03 | 401261.78 | 295417.06 | 308192.28 | 326566.91 | 454464.22 | 385321.59 | 319820.66 | 307944.28 | 6 | **1.289** |

| | | | | | | | | | | | | | | | | | | |
|---|---|---|---|---|---|---|---|---|---|---|---|---|---|---|---|---|---|---|
| Q8BYR1 | Lcmt2 | tRNA wybutosine-synthesizing protein 4 (Mus musculus) | 57.71 | 2.04 | 75295 | 2 | 5900471.5 | 5127059 | 4408988 | 5580981 | 5776746 | 4336982 | 5583747 | 6302302 | 6527574 | 6775032.5 | 7 | **1.102** |
| Q8BYW1 | Arhgap25 | Rho GTPase-activating protein 25 (Mus musculus) 5 | 244.24 | 7.72 | 73383 | 4 | 5977453 | 3929924 | 3899751 | 4562439.5 | 7111347 | 6031178 | 7724519 | 6678002.5 | 5195623 | 5944911 | 6 | **1.239** |
| Q8BYZ1 | Abi3 | ABI gene family member 3 (Mus musculus) | 63.6 | 4.9 | 39107 | 1 | 0 | 561619.75 | 822672.25 | 0 | 0 | 288404.44 | 351835.19 | 294339.56 | 279050.84 | 0 | 15 | **0.877** |
| Q8BZA9 | Tigar | Fructose-2,6-bisphosphatase TIGAR (Mus musculus) | 184.33 | 19.33 | 29191 | 3 | 2076824.12 | 1106800.5 | 1265464.88 | 2287805.5 | 2641822 | 4166751.5 | 1668696.75 | 2135863 | 3860165 | 3918423.5 | 5 | **1.679** |
| Q8BZM1 | Glmn | Glomulin (Mus musculus) | 259.47 | 11.24 | 67756 | 5 | 1330946.5 | 2018042.5 | 1431140.88 | 1220603.12 | 833193.5 | 1241643.5 | 1414667 | 2027728 | 1497444.12 | 2047051.75 | 6 | **1.204** |
| Q8BZN6 | Dock10 | Dedicator of cytokinesis protein 10 (Mus musculus) | 73.61 | 0.74 | 245758 | 1 | 114023.24 | 310807.25 | 459560.88 | 410203 | 83055.59 | 0 | 104443.26 | 484061.22 | 81541.59 | 130089.34 | 8 | **0.581** |
| Q8BZS9 | Dhx32 | Putative pre-mRNA-splicing factor ATP-dependent RNA helicase DHX32 (Mus musculus) | 87.57 | 1.61 | 83941 | 1 | 0 | 161011.41 | 198555.41 | 199158.12 | 164751.14 | 216566.56 | 183185.08 | 250695.36 | 177398.83 | 192283.75 | 6 | **1.410** |
| Q8BZW8 | Nhlrc2 | NHL repeat-containing protein 2 (Mus musculus) | 123.26 | 3.03 | 78430 | 2 | 1506508.25 | 734235.88 | 369407.47 | 1357797 | 0 | 1315418.38 | 1057363.12 | 0 | 1060040.5 | 182151.08 | 11 | **0.911** |
| Q8C052 | Map1s | Microtubule-associated protein 1S (Mus musculus) | 58.53 | 1.03 | 102939 | 1 | 2036851.12 | 1257799.25 | 1503488.62 | 1696239.88 | 1542894.5 | 1857843 | 1462974.62 | 1639331.38 | 1805248 | 1662157.38 | 10 | **1.049** |
| Q8C079 | Strip1 | Striatin-interacting protein 1 (Mus musculus) | 54.25 | 0.84 | 95585 | 1 | 669809.62 | 614749.69 | 777179.81 | 334131.28 | 654636.94 | 770927.12 | 747285.19 | 782174.06 | 599782.94 | 745458.25 | 7 | **1.195** |
| Q8C092 | Taf5 | Transcription initiation factor TFIID subunit 5 (Mus musculus) | 68.75 | 1.37 | 87045 | 1 | 332213.84 | 238715.23 | 268912.41 | 232429.67 | 0 | 251822.41 | 275190.88 | 264307.62 | 240083.48 | 236917.23 | 10 | **1.183** |
| Q8C0C7 | Farsa | Phenylalanine--tRNA ligase alpha subunit (Mus musculus) | 311.39 | 15.35 | 57599 | 6 | 10398223 | 4449375 | 5659061 | 1978909.75 | 3265121.5 | 8080030 | 8691424 | 7313486 | 5577859 | 5372798 | 7 | **1.361** |
| Q8C0D5 | Efl1 | Elongation factor-like GTPase 1 (Mus musculus) | 261.38 | 5.86 | 125777 | 5 | 1839556.25 | 1374392.5 | 1728237 | 1542481.5 | 1458601 | 564668.12 | 1126146 | 1377889.75 | 1260958.25 | 1420385.62 | 2 | **0.724** |
| Q8C0E2 | Vps26b | Vacuolar protein sorting-associated protein 26B (Mus musculus) | 64.72 | 6.25 | 39125 | 2 | 859855.44 | 1902585.25 | 2547477.25 | 1934025.12 | 2079560.25 | 4542676 | 1847248.38 | 2575932.25 | 2176774.25 | 2539947.5 | 6 | **1.468** |
| Q8C0I1 | Agps | Alkyldihydroxyacetonephosphate synthase, peroxisomal (Mus musculus) | 655.49 | 26.2 | 71684 | 13 | 11559444 | 11515855 | 12591021 | 11662402 | 10881729 | 12886085 | 13146042 | 12505841 | 12802432 | 12170687 | 2 | **1.091** |
| Q8C0L0 | Tmx4 | Thioredoxin-related transmembrane protein 4 (Mus musculus) | 45.95 | 3.58 | 37131 | 1 | 1941304.88 | 1681015 | 1440515.12 | 680737.81 | 414663.94 | 1471169.25 | 1462291.88 | 1349558.62 | 2363496.75 | 1771883 | 8 | **1.367** |
| Q8C0L8 | Cog5 | Conserved oligomeric Golgi complex subunit 5 (Mus musculus) | 105.12 | 2.53 | 91391 | 2 | 442343.19 | 326478.53 | 1443331.38 | 1453436.25 | 1544725.12 | 1391974.62 | 1455555.12 | 1047818.25 | 1345557.75 | 1144903 | 12 | **1.226** |
| Q8C0M9 | Asrgl1 | Isoaspartyl peptidase/L-asparaginase (Mus musculus) | 158.84 | 10.74 | 33950 | 3 | 4816876 | 5681349 | 6586040.5 | 4252604 | 5410441 | 7222291 | 7147483 | 7047618 | 7526197.5 | 7861725 | 0 | **1.376** |
| Q8C129 | Lnpep | Leucyl-cystinyl aminopeptidase (Mus musculus) | 880.91 | 18.05 | 117304 | 16 | 11866470 | 17068328 | 20323560 | 13867710 | 16567412 | 18990984 | 18513824 | 18175510 | 20673402 | 18038272 | 4 | **1.184** |
| Q8C142 | Ldlrap1 | Low density lipoprotein receptor adapter protein 1 (Mus musculus) | 84.03 | 5.19 | 33975 | 1 | 876088.69 | 545245.81 | 1126052.62 | 323560.41 | 538929.62 | 820263.81 | 553641.94 | 805002.94 | 849620.12 | 751505.75 | 10 | **1.109** |
| Q8C147 | Dock8 | Dedicator of cytokinesis protein 8 (Mus musculus) | 482.36 | 7.05 | 238978 | 12 | 9540007 | 11248918 | 9435130 | 9027333 | 6930637 | 14286656 | 11773972 | 11349385 | 11746994 | 12228385 | 0 | **1.329** |
| Q8C156 | Ncaph | Condensin complex subunit 2 (Mus musculus) | 233.1 | 8.48 | 82303 | 4 | 4475026.5 | 3023802.5 | 4068749.5 | 4892171 | 4617017 | 3769339.25 | 3263068.5 | 3132590.75 | 4456051 | 5120966 | 10 | **0.937** |
| Q8C166 | Cpne1 | Copine-1 (Mus musculus) | 431.54 | 17.91 | 58887 | 8 | 13509331 | 13630927 | 13527415 | 12576890 | 12957423 | 15097363 | 14770170 | 11689569 | 13950021 | 14572795 | 5 | **1.059** |
| Q8C1A5 | Thop1 | Thimet oligopeptidase (Mus musculus) | 420.68 | 12.66 | 78026 | 8 | 5540076 | 8268267 | 8154129 | 9579206 | 10222119 | 7880748.5 | 3632862 | 7480454.5 | 7977808.5 | 8974328 | 6 | **0.861** |
| Q8C1Z7 | Bbs4 | Bardet-Biedl syndrome 4 protein homolog (Mus musculus) | 37.45 | 1.35 | 58255 | 1 | 2370527 | 5836531.5 | 4469477 | 7008241.5 | 4715483.5 | 2678007.25 | 2526038.75 | 2776929.25 | 2303871.75 | 2145396.25 | 3 | **0.509** |
| Q8C2E7 | Washc5 | WASH complex subunit 5 (Mus musculus) | 130.91 | 3.19 | 134110 | 3 | 1406764.12 | 2606726.75 | 2560832 | 2998637.5 | 2081108.12 | 2018687.25 | 2527007 | 2548282.25 | 2449047.75 | 2311517 | 9 | **1.017** |
| Q8C2Q3 | Rbm14 | RNA-binding protein 14 (Mus musculus) | 371.28 | 12.11 | 69449 | 7 | 13362374 | 8191330 | 8743728 | 8472373 | 7834088.5 | 9570711 | 11420520 | 8038262 | 8096801.5 | 8989766 | 11 | **0.990** |
| Q8C3J5 | Dock2 | Dedicator of cytokinesis protein 2 (Mus musculus) | 774.78 | 12.53 | 211704 | 18 | 15637594 | 19812556 | 23823998 | 22756366 | 17524294 | 17603670 | 17007606 | 20933226 | 19557656 | 18919174 | 10 | **0.944** |
| Q8C3X8 | Lmf2 | Lipase maturation factor 2 (Mus musculus) | 133.7 | 4.84 | 79997 | 2 | 0 | 99661.51 | 0 | 115225.23 | 0 | 414269.09 | 370413.72 | 428541.78 | 0 | 369849.56 | 8 | **7.367** |
| Q8C3Y4 | Kntc1 | Kinetochore-associated protein 1 (Mus musculus) | 50.99 | 0.59 | 250358 | 1 | 0 | 167512.94 | 155815.67 | 203014.36 | 208871.42 | 0 | 0 | 248657.08 | 190724.06 | 214818.28 | 12 | **0.890** |
| Q8C4J7 | Tbl3 | Transducin beta-like protein 3 (Mus musculus) | 210.11 | 4.99 | 88266 | 3 | 3163340 | 2578719.25 | 2927249.5 | 1642839.62 | 1459762.62 | 3198351.25 | 3695056.5 | 3134667.25 | 1903826 | 3250498.75 | 4 | **1.290** |
| Q8C4Y3 | Nelfb | Negative elongation factor B (Mus musculus) | 93.7 | 5.34 | 65636 | 2 | 185727.88 | 932492.25 | 620631.94 | 691527.19 | 811408.69 | 696677.88 | 514574.91 | 606481.06 | 488919.34 | 538517.94 | 7 | **0.878** |
| Q8C5N3 | Cwc22 | Pre-mRNA-splicing factor CWC22 homolog (Mus musculus) | 42.12 | 0.88 | 104773 | 1 | 0 | 1200016.62 | 1070498.12 | 0 | 0 | 712770.06 | 904381.69 | 891082.81 | 1024215.5 | 765093.75 | 13 | **1.893** |
| Q8C5Q4 | Grsf1 | G-rich sequence factor 1 (Mus musculus) | 102.93 | 7.1 | 53076 | 2 | 2478137.5 | 2214496.25 | 1797548.88 | 1623933.25 | 1448485.25 | 1685348.75 | 2168326.25 | 1632505.88 | 1637586.75 | 1588423.25 | 10 | **0.911** |
| Q8C6G8 | Wdr26 | WD repeat-containing protein 26 (Mus musculus) | 488.37 | 16.69 | 70543 | 9 | 15550126 | 11522170 | 11123432 | 11563308 | 12437276 | 16815974 | 12729216 | 14486203 | 15821884 | 15307593 | 3 | **1.208** |
| Q8C7K6 | Pcyox1l | Prenylcysteine oxidase-like (Mus musculus) | 58.74 | 2.22 | 54875 | 1 | 1964719.62 | 2477241.5 | 1330456.38 | 2289821.75 | 1777588.75 | 2671774.75 | 2607127.75 | 2534526.25 | 2337854 | 2407713 | 2 | **1.276** |
| Q8C7R4 | Uba6 | Ubiquitin-like modifier-activating enzyme 6 (Mus musculus) | 523.52 | 12.82 | 117966 | 11 | 5703967.5 | 11657004 | 9150952 | 10836511 | 7431760 | 4675597 | 10699197 | 8759108 | 8027178.5 | 10821287 | 10 | **0.960** |
| Q8C7V3 | Utp15 | U3 small nucleolar RNA-associated protein 15 homolog (Mus musculus) | 243.16 | 11.74 | 59375 | 5 | 5506715.5 | 3299477.75 | 4273339.5 | 1982100.25 | 3761248.75 | 5899276 | 5445147 | 4765831.5 | 4154400.5 | 3987026.25 | 6 | **1.288** |
| Q8C7X2 | Emc1 | ER membrane protein complex subunit 1 (Mus musculus) | 821.26 | 22.97 | 111605 | 15 | 16191814 | 16621164 | 20994746 | 13225174 | 14135030 | 21307180 | 19361768 | 17719086 | 15937653 | 18036912 | 6 | **1.138** |
| Q8C854 | Myef2 | Myelin expression factor 2 (Mus musculus) | 328.79 | 11 | 63295 | 6 | 10460159 | 11410724 | 8673994 | 12013570 | 11656122 | 9121322 | 10433854 | 10532419 | 8788554 | 11162525 | 7 | **0.923** |
| Q8C878 | Uba3 | NEDD8-activating enzyme E1 catalytic subunit (Mus musculus) | 449.76 | 31.17 | 51737 | 8 | 5759333 | 6805605 | 7276427.5 | 4457404.5 | 3220322.25 | 6857945 | 6597346 | 6253431.5 | 6956740 | 5368516.5 | 9 | **1.164** |
| Q8C9W3 | Adamts2 | A disintegrin and metalloproteinase with thrombospondin motifs 2 (Mus musculus) | 41.97 | 0.58 | 135299 | 1 | 711568.06 | 190601.11 | 13388835 | 7011694.5 | 342260.78 | 20533540 | 207777.05 | 14629239 | 14826417 | 14138520 | 4 | **2.972** |
| Q8CAQ8 | Immt | MICOS complex subunit Mic60 (Mus musculus) | 1560.71 | 46.63 | 83900 | 28 | 50250916 | 45673592 | 49253184 | 53535720 | 51389760 | 47891968 | 42293028 | 46487892 | 47756812 | 46908252 | 4 | **0.925** |
| Q8CAS9 | Parp9 | Poly [ADP-ribose] polymerase 9 (Mus musculus) | 39 | 1.96 | 96659 | 1 | 0 | 0 | 145086.83 | 0 | 50177.42 | 283123.97 | 270639.81 | 0 | 156421.75 | 135432.81 | 9 | **4.331** |
| Q8CAY6 | Acat2 | Acetyl-CoA acetyltransferase, cytosolic (Mus musculus) | 133.21 | 6.55 | 41298 | 2 | 3083416.5 | 1797598.25 | 1696922 | 746866.31 | 1174005.12 | 2506390.25 | 2406161 | 2366573 | 1275156.88 | 2286385.5 | 7 | **1.276** |
| Q8CB96 | Rassf4 | Ras association domain-containing protein 4 (Mus musculus) | 76.57 | 6.52 | 36934 | 2 | 3533183.5 | 2530076 | 2777699.25 | 2840606.5 | 2989788 | 2346229.25 | 2553095.25 | 2784816.75 | 2952795 | 2570675.75 | 7 | **0.900** |
| Q8CBY8 | Dctn4 | Dynactin subunit 4 (Mus musculus) | 126.91 | 5.78 | 53057 | 2 | 1200407.88 | 0 | 1152798.88 | 499692.38 | 740827.12 | 727937.75 | 788640.81 | 0 | 1026726.25 | 888475.81 | 12 | **0.955** |
| Q8CCB4 | Vps53 | Vacuolar protein sorting-associated protein 53 homolog (Mus musculus) | 41.25 | 2.88 | 94423 | 1 | 392048.12 | 0 | 216107.88 | 528276.38 | 535774.25 | 0 | 169505.77 | 406013.44 | 578138.88 | 432419.03 | 13 | **0.948** |
| Q8CCF0 | Prpf31 | U4/U6 small nuclear ribonucleoprotein Prp31 (Mus musculus) | 416.6 | 18.84 | 55430 | 8 | 6386337 | 4427191.5 | 5886495 | 7872635.5 | 4822138 | 7033423 | 6458315 | 6446926 | 6692585.5 | 6751540.5 | 5 | **1.136** |
| Q8CCH2 | Nhlrc3 | NHL repeat-containing protein 3 (Mus musculus) | 499.2 | 37.75 | 38195 | 8 | 9614741 | 6873191.5 | 7431759.5 | 4120116.75 | 8511324 | 7609364.5 | 8393210 | 9335178 | 7805494.5 | 8985157 | 8 | **1.153** |
| Q8CCI5 | Rybp | RING1 and YY1-binding protein (Mus musculus) | 122.34 | 29.39 | 24777 | 3 | 2995805.75 | 1831403.25 | 1918894.75 | 2322178 | 2080525.75 | 1737182.25 | 2159333 | 2223947.25 | 1751049 | 1441593.25 | 6 | **0.835** |
| Q8CCJ3 | Ufl1 | E3 UFM1-protein ligase 1 (Mus musculus) | 131.76 | 4.54 | 89520 | 2 | 111202.43 | 739979.62 | 869180.62 | 976743.88 | 1053623.38 | 366252.72 | 264593.72 | 842134.81 | 816634.94 | 1015721.62 | 10 | **0.881** |
| Q8CCP0 | Nemf | Nuclear export mediator factor Nemf (Mus musculus) | 127.2 | 2.82 | 121188 | 3 | 218737.72 | 1536970.25 | 930962 | 1493607.75 | 1204810.5 | 2036749.25 | 623170.25 | 1426989.12 | 507976.59 | 0 | 10 | **0.853** |
| Q8CCS6 | Pabpn1 | Polyadenylate-binding protein 2 (Mus musculus) | 193.29 | 25.17 | 32297 | 3 | 6452459.5 | 13231935 | 10652040 | 11188017 | 9533090 | 11300580 | 9844440 | 9952551 | 8891621 | 10186944 | 11 | **0.983** |
| Q8CD15 | Riox2 | Ribosomal oxygenase 2 (Mus musculus) | 46.18 | 4.3 | 53517 | 1 | 506637.88 | 0 | 587665.81 | 339319.69 | 392079.28 | 0 | 566046.94 | 343885.34 | 311415.25 | 321509.81 | 9 | **0.845** |
| Q8CD92 | Ttc27 | Tetratricopeptide repeat protein 27 (Mus musculus) | 84.61 | 2.48 | 96431 | 1 | 53979.25 | 0 | 242983.78 | 218818.89 | 116906.27 | 0 | 0 | 153571.17 | 194751 | 185853.34 | 12 | **0.844** |
| Q8CDG3 | Vcpip1 | Deubiquitinating protein VCIP135 (Mus musculus) | 146.6 | 3.44 | 134503 | 3 | 2284467.5 | 1639299.75 | 1669321 | 1610944 | 130852.46 | 1662083 | 1984020.12 | 1632339 | 1541269.38 | 1161441.5 | 11 | **1.088** |
| Q8CDN6 | Txnl1 | Thioredoxin-like protein 1 (Mus musculus) | 169.86 | 11.42 | 32237 | 3 | 7723082 | 6667929 | 4388500 | 9023080 | 3273178.75 | 5210500.5 | 4899255 | 2844909 | 4077495.25 | 3656967.25 | 6 | **0.666** |
| Q8CE96 | Trmt6 | tRNA (adenine(58)-N(1))-methyltransferase non-catalytic subunit TRM6 (Mus musculus) | 46.34 | 3.62 | 55518 | 1 | 327889.97 | 498734.91 | 0 | 596565.75 | 625285.31 | 0 | 365990.12 | 796968.81 | 673561.38 | 280512.5 | 12 | **1.033** |
| Q8CF89 | Tab1 | TGF-beta-activated kinase 1 and MAP3K7-binding protein 1 (Mus musculus) | 60.49 | 2.19 | 54616 | 1 | 0 | 623898.62 | 0 | 1292926.88 | 558037.19 | 698622.31 | 1046798.75 | 0 | 868760.44 | 0 | 14 | **1.056** |
| Q8CFD4 | Snx8 | Sorting nexin-8 (Mus musculus) | 163.1 | 5.45 | 52059 | 2 | 0 | 866943.56 | 1471517.88 | 0 | 1220882.25 | 1529562.25 | 1245905.25 | 1441420.5 | 1325567.5 | 1148621.38 | 6 | **1.880** |
| Q8CFE2 | Hpf1 | Histone PARylation factor 1 (Mus musculus) | 323.21 | 26.01 | 39291 | 6 | 5348505.5 | 5045796 | 7020946 | 3998406.5 | 6911479 | 5381382.5 | 7507542.5 | 6731727 | 4905764 | 6585139 | 10 | **1.098** |
| Q8CFE3 | Rcor1 | REST corepressor 1 (Mus musculus) | 89.5 | 7.5 | 52715 | 2 | 593810.75 | 705585.06 | 833613.56 | 1403501.5 | 949699.5 | 581498.94 | 632499.94 | 582823.56 | 722898.69 | 0 | 3 | **0.562** |
| Q8CFE4 | Scyl2 | SCY1-like protein 2 (Mus musculus) | 111.94 | 2.69 | 103317 | 2 | 614218.19 | 1393773.25 | 1369137.38 | 0 | 0 | 1384688.88 | 1768733.62 | 1530561.5 | 387920.59 | 1289858.5 | 7 | **1.884** |
| Q8CFI7 | Polr2b | DNA-directed RNA polymerase II subunit RPB2 (Mus musculus) | 286.24 | 5.71 | 133911 | 6 | 5393130 | 4410624.5 | 2696232.75 | 4683194.5 | 2864024 | 5443255.5 | 4557142.5 | 4403518.5 | 4220934.5 | 4194324 | 11 | **1.138** |
| Q8CFQ3 | Aqr | RNA helicase aquarius (Mus musculus) | 39.28 | 1.22 | 170294 | 1 | 3706394.25 | 0 | 0 | 0 | 0 | 0 | 0 | 651611.62 | 0 | 0 | 26 | **0.176** |
| Q8CFV9 | Rfk | Riboflavin kinase (Mus musculus) | 182.09 | 26.45 | 17437 | 4 | 3598573 | 2968309.75 | 4718794.5 | 2553053.75 | 4060742.25 | 3674241.25 | 3042313.5 | 3991149.25 | 4047998.75 | 4219876 | 10 | **1.060** |
| Q8CG47 | Smc4 | Structural maintenance of chromosomes protein 4 (Mus musculus) | 1098.75 | 21.54 | 146895 | 21 | 29018362 | 24534398 | 30410856 | 28072824 | 22033080 | 27489872 | 29598978 | 29900354 | 28746874 | 33034520 | 7 | **1.110** |
| Q8CG48 | Smc2 | Structural maintenance of chromosomes protein 2 (Mus musculus) | 647.17 | 11.92 | 134239 | 12 | 15951559 | 14968531 | 14848508 | 14855747 | 12854444 | 15016958 | 15590693 | 16422365 | 16193502 | 17001786 | 2 | **1.092** |
| Q8CG72 | Adprhl2 | Poly(ADP-ribose) glycohydrolase ARH3 (Mus musculus) | 82.89 | 9.46 | 39414 | 2 | 2898236 | 709345.19 | 1432012.75 | 11074674 | 1542150.12 | 1279420.88 | 1486162.88 | 1073471.5 | 1373590.5 | 463900.03 | 5 | **0.322** |
| Q8CG76 | Akr7a2 | Aflatoxin B1 aldehyde reductase member 2 (Mus musculus) | 122.74 | 7.08 | 40612 | 2 | 1805302.25 | 750787.56 | 1322537.5 | 609912.56 | 962388.38 | 1688559.12 | 1857709.25 | 1814916 | 2038652.75 | 1563539.88 | 2 | **1.644** |
| Q8CGC6 | Rbm28 | RNA-binding protein 28 (Mus musculus) | 130.77 | 4.13 | 84214 | 2 | 949924.81 | 917472.75 | 1378597.5 | 1251093.12 | 1496542 | 1091564.62 | 1436877.25 | 1318309.25 | 1468680.25 | 1445551.25 | 8 | **1.128** |
| Q8CGC7 | Eprs | Bifunctional glutamate/proline--tRNA ligase (Mus musculus) | 2576.22 | 40.74 | 170079 | 44 | 117354744 | 108361776 | 106903488 | 130798784 | 128075880 | 111301568 | 106235808 | 110187840 | 125086808 | 114720128 | 9 | **0.959** |
| Q8CGF7 | Tcerg1 | Transcription elongation regulator 1 (Mus musculus) | 68.32 | 1.09 | 123788 | 1 | 0 | 1276933 | 755610.38 | 1569108.25 | 694044 | 487007.44 | 147390.38 | 0 | 0 | 471954.44 | 6 | **0.258** |
| Q8CGK3 | Lonp1 | Lon protease homolog, mitochondrial (Mus musculus) | 1062.37 | 29.93 | 105843 | 20 | 21194564 | 16934524 | 18256272 | 15221628 | 17337396 | 22470952 | 18168296 | 21903332 | 20248820 | 15903539 | 7 | **1.110** |
| Q8CGS4 | Chmp3 | Charged multivesicular body protein 3 (Rattus norvegicus) | 60.1 | 5.38 | 25062 | 1 | 0 | 884905.38 | 1035253.06 | 900928.88 | 772835.38 | 1757502.5 | 1224130.88 | 1680384.62 | 1378109.75 | 1174866.25 | 0 | **2.008** |
| Q8CGZ0 | Cherp | Calcium homeostasis endoplasmic reticulum protein (Mus musculus) | 91.76 | 2.03 | 106169 | 2 | 577419.06 | 371554.38 | 592367.69 | 720006.94 | 698953.62 | 747243 | 779291.44 | 768420.75 | 789617.12 | 783849.12 | 0 | **1.307** |
| Q8CH18 | Ccar1 | Cell division cycle and apoptosis regulator protein 1 (Mus musculus) | 52.26 | 0.79 | 132060 | 1 | 2768102.25 | 2569168.5 | 1804796.12 | 2279307.75 | 2658877.5 | 2456142.5 | 2442338.75 | 2258293.5 | 2010095.12 | 1724637.75 | 6 | **0.902** |
| Q8CH72 | Trim32 | E3 ubiquitin-protein ligase TRIM32 (Mus musculus) | 41.58 | 2.29 | 72057 | 1 | 1117890.75 | 0 | 801313.81 | 0 | 968825.06 | 792573.88 | 1151373.5 | 1130472 | 833096 | 791735.94 | 9 | **1.627** |
| Q8CHH9 | Sept8 | Septin-8 (Mus musculus) | 297.77 | 23.31 | 49812 | 7 | 9479831 | 9426716 | 7302402 | 6405823 | 8078406 | 7297091 | 7845403 | 6123344.5 | 4964508.5 | 7261736 | 4 | **0.823** |
| Q8CHJ1 | Pigu | Phosphatidylinositol glycan anchor biosynthesis class U protein (Rattus norvegicus) | 73.01 | 3.91 | 49953 | 1 | 0 | 254036.48 | 396564.66 | 308403.69 | 310809.09 | 359477.06 | 215383.23 | 325217.25 | 291961.81 | 273397.88 | 12 | **1.154** |
| Q8CHK3 | Mboat7 | Lysophospholipid acyltransferase 7 (Mus musculus) | 93.13 | 6.77 | 53436 | 2 | 79178.33 | 601429.38 | 388369.06 | 0 | 472564.94 | 1076775.5 | 1212790.12 | 560690.5 | 0 | 312594.06 | 9 | **2.052** |
| Q8CHW4 | Eif2b5 | Translation initiation factor eIF-2B subunit epsilon (Mus musculus) | 184.03 | 9.21 | 80086 | 4 | 2064707.25 | 1997070.5 | 2566880.75 | 1975011.25 | 1718561.5 | 1664176.5 | 0 | 1979653.75 | 0 | 57075.43 | 3 | **0.359** |
| Q8CI11 | Gnl3 | Guanine nucleotide-binding protein-like 3 (Mus musculus) | 549.26 | 27.14 | 60786 | 10 | 9759250 | 7726324 | 8583164 | 5977804 | 8191639.5 | 10736771 | 11596406 | 10295735 | 9548130 | 11163072 | 1 | **1.326** |
| Q8CI51 | Pdlim5 | PDZ and LIM domain protein 5 (Mus musculus) | 323.2 | 12.86 | 63299 | 7 | 9835905 | 6890762 | 5746090.5 | 5098001.5 | 6671426.5 | 8703127 | 9977159 | 9190570 | 7802593 | 8914283 | 4 | **1.302** |
| Q8CI75 | Dis3l2 | DIS3-like exonuclease 2 (Mus musculus) | 169.13 | 3.68 | 97775 | 4 | 2834102 | 2005339.5 | 4323202 | 1257832.38 | 2175228.75 | 4476492.5 | 3839969 | 4729946.5 | 4020298.25 | 4901016 | 2 | **1.744** |
| Q8CI95 | Osbpl11 | Oxysterol-binding protein-related protein 11 (Mus musculus) | 50.64 | 1.33 | 83629 | 1 | 1715255.62 | 1165535.62 | 1021313.38 | 0 | 0 | 1216965 | 1281045.88 | 1170383.75 | 0 | 896447.81 | 12 | **1.170** |
| Q8CIE6 | Copa | Coatomer subunit alpha (Mus musculus) | 1721.65 | 31.05 | 138432 | 31 | 80911672 | 59160712 | 61926908 | 43568980 | 59335668 | 67415824 | 61046728 | 68601696 | 65076992 | 67358200 | 6 | **1.081** |

| | | | | | | | | | | | | | | | | | | |
|---|---|---|---|---|---|---|---|---|---|---|---|---|---|---|---|---|---|---|
| Q8CIG8 | Prmt5 | Protein arginine N-methyltransferase 5 (Mus musculus) | 193.47 | 5.34 | 72680 | 4 | 3390537.75 | 5055565.5 | 3981195 | 5570538 | 5651392 | 5368018 | 3866985 | 5010825 | 4928039 | 4069570.5 | 10 | **0.983** |
| Q8CIH5 | Plcg2 | 1-phosphatidylinositol 4,5-bisphosphate phosphodiesterase gamma-2 (Mus musculus) | 1019.14 | 22.45 | 147592 | 20 | 17842362 | 21268810 | 21628564 | 25872098 | 13828411 | 21601476 | 18667002 | 24177156 | 23125204 | 23112748 | 8 | **1.102** |
| Q8CII2 | Cdc123 | Cell division cycle protein 123 homolog (Mus musculus) | 188.3 | 11.31 | 38816 | 3 | 658463.75 | 4542398 | 5304724 | 4831966.5 | 4366777 | 3695516 | 3600418.75 | 3482138 | 3339378.25 | 3180963.5 | 5 | **0.878** |
| Q8CIM5 | Gpr84 | G-protein coupled receptor 84 (Mus musculus) | 275.9 | 16.92 | 43717 | 5 | 3256805.75 | 11043370 | 11109841 | 11293992 | 5845981 | 8031350.5 | 8910021 | 11313540 | 11615574 | 9934269 | 9 | **1.170** |
| Q8CIM8 | Ints4 | Integrator complex subunit 4 (Mus musculus) | 51.97 | 1.04 | 108193 | 1 | 809202.62 | 204130.25 | 395861.41 | 234412.12 | 491157.25 | 496481.16 | 719623.56 | 588261.5 | 387049.72 | 387671.59 | 9 | **1.208** |
| Q8CIN4 | Pak2 | Serine/threonine-protein kinase PAK 2 (Mus musculus) | 725.47 | 27.1 | 57930 | 11 | 17855324 | 15822756 | 14689041 | 10496700 | 17762336 | 14612513 | 14810549 | 15698805 | 17992330 | 16168585 | 12 | **1.035** |
| Q8CIN7 | Impa2 | Inositol monophosphatase 2 (Rattus norvegicus) | 111.05 | 7.59 | 31796 | 2 | 2448747.25 | 2410364.75 | 2354152.75 | 1722246.62 | 2251148.5 | 2159966 | 1473041.88 | 1758464.12 | 1696569.25 | 1658473.88 | 2 | **0.782** |
| Q8CJF7 | Ahctf1 | Protein ELYS (Mus musculus) | 49.12 | 0.98 | 247646 | 1 | 0 | 208340.47 | 0 | 0 | 0 | 195634.14 | 86539.35 | 183482.95 | 0 | 0 | 19 | **2.235** |
| Q8CJG0 | Ago2 | Protein argonaute-2 (Mus musculus) | 175.31 | 5 | 97304 | 3 | 137298.08 | 812460.19 | 1495810.25 | 1334930.25 | 856019 | 1100533.88 | 1617719.62 | 1298882.25 | 1210644 | 1001380.06 | 8 | **1.343** |
| Q8JZK9 | Hmgcs1 | Hydroxymethylglutaryl-CoA synthase, cytoplasmic (Mus musculus) | 392.81 | 20.77 | 57569 | 7 | 5114441.5 | 6735674.5 | 7228274 | 6823166 | 5716575 | 6120540.5 | 4295592.5 | 8503596 | 7297434 | 7081678 | 9 | **1.053** |
| Q8JZN5 | Acad9 | Acyl-CoA dehydrogenase family member 9, mitochondrial (Mus musculus) | 578.24 | 20 | 68722 | 10 | 2438406 | 7731912.5 | 6373878 | 7869430.5 | 6114970.5 | 6211625 | 7299031 | 6180861.5 | 6424811 | 8150672 | 10 | **1.122** |
| Q8JZQ9 | Eif3b | Eukaryotic translation initiation factor 3 subunit B (Mus musculus) | 1179.6 | 37.11 | 91370 | 23 | 30780442 | 34410176 | 40602432 | 34729500 | 34640068 | 45383744 | 38033080 | 41556560 | 40888108 | 46522716 | 1 | **1.212** |
| Q8JZR0 | Acsl5 | Long-chain-fatty-acid--CoA ligase 5 (Mus musculus) | 734.54 | 26.79 | 76206 | 14 | 22921910 | 24854402 | 27423914 | 28042922 | 24023772 | 28420406 | 28034180 | 29600874 | 26994950 | 29985224 | 3 | **1.124** |
| Q8JZX4 | Rbm17 | Splicing factor 45 (Mus musculus) | 105.87 | 6.91 | 45304 | 2 | 2697769.25 | 2226380 | 2277957.75 | 1677328.12 | 2807467.75 | 2491339.5 | 1935890.38 | 2464261.25 | 2138557.5 | 2332604 | 11 | **0.972** |
| Q8JZY2 | Commd10 | COMM domain-containing protein 10 (Mus musculus) | 55.75 | 5.94 | 22812 | 1 | 1204217.38 | 1226951.88 | 514371.41 | 650162.69 | 789802.31 | 895646.56 | 851580.12 | 1045558.19 | 935530.38 | 1004254.31 | 10 | **1.079** |
| Q8K019 | Bclaf1 | Bcl-2-associated transcription factor 1 (Mus musculus) | 81.19 | 2.39 | 106002 | 2 | 7014296 | 4485215 | 4696502.5 | 4235116 | 5212132 | 4801818 | 5425692 | 4783270.5 | 5208691 | 5774359.5 | 8 | **1.014** |
| Q8K0C9 | Gmds | GDP-mannose 4,6 dehydratase (Mus musculus) | 184.95 | 12.63 | 41985 | 3 | 2549958.75 | 3979518.25 | 3574213.25 | 3316721.5 | 3589704.5 | 986568 | 1230650.75 | 2667417.5 | 3312511 | 2855020.25 | 3 | **0.650** |
| Q8K0D5 | Gfm1 | Elongation factor G, mitochondrial (Mus musculus) | 250.5 | 9.99 | 83550 | 6 | 3501657.25 | 6256391 | 6304128.5 | 3925173.5 | 4572707 | 6688039 | 6531404.5 | 7124716 | 6315148.5 | 4666547 | 2 | **1.275** |
| Q8K0V4 | Cnot3 | CCR4-NOT transcription complex subunit 3 (Mus musculus) | 103.5 | 4.13 | 81946 | 2 | 303338.5 | 0 | 0 | 0 | 0 | 348721.47 | 249477.66 | 389239.72 | 416027.41 | 0 | 12 | **4.627** |
| Q8K124 | Plekho2 | Pleckstrin homology domain-containing family O member 2 (Mus musculus) | 314.79 | 22.22 | 53872 | 6 | 5877003 | 4079727 | 2754570.25 | 1083440 | 3905640.5 | 3980421 | 6401574.5 | 4827738.5 | 5481292 | 5260222.5 | 5 | **1.466** |
| Q8K183 | Pdxk | Pyridoxal kinase (Mus musculus) | 363.55 | 16.35 | 35015 | 6 | 5421159 | 7104171.5 | 9425047 | 6814149 | 8814731 | 5584792 | 8109211.5 | 8437886 | 9522407 | 8636825 | 10 | **1.072** |
| Q8K1B8 | Fermt3 | Fermitin family homolog 3 (Mus musculus) | 1802.91 | 57.29 | 75635 | 28 | 97356984 | 97559296 | 116614800 | 101618288 | 99615688 | 125796464 | 129982936 | 133540384 | 131254784 | 138248240 | 0 | **1.285** |
| Q8K1C9 | Lrrc41 | Leucine-rich repeat-containing protein 41 (Mus musculus) | 163.78 | 7.93 | 88180 | 3 | 944708 | 2009387 | 2098737.75 | 1779796.25 | 377403 | 1270873.75 | 1126295.25 | 1732870.62 | 460529.31 | 1373200 | 9 | **0.827** |
| Q8K1J6 | Trnt1 | CCA tRNA nucleotidyltransferase 1, mitochondrial (Mus musculus) | 217.8 | 9.91 | 49895 | 5 | 25273868 | 16202727 | 15205632 | 19270604 | 14676581 | 16826390 | 22282636 | 20023062 | 20843766 | 18184590 | 7 | **1.083** |
| Q8K1R3 | Pnpt1 | Polyribonucleotide nucleotidyltransferase 1, mitochondrial (Mus musculus) | 140.64 | 4.98 | 85683 | 3 | 1898192 | 3194317.75 | 3466719.5 | 3177227.5 | 2827186.5 | 3155128.5 | 2440950.5 | 3558024.5 | 3485001.5 | 4079617 | 7 | **1.148** |
| Q8K1R7 | Nek9 | Serine/threonine-protein kinase Nek9 (Mus musculus) | 137.72 | 3.56 | 107143 | 3 | 2702109.25 | 2247502.5 | 2930160 | 3401156 | 1897260 | 3462748 | 2947000.25 | 3472101.75 | 3203252 | 2543728.25 | 5 | **1.186** |
| Q8K1X1 | Wdr11 | WD repeat-containing protein 11 (Mus musculus) | 212.29 | 4.58 | 135937 | 3 | 235899.97 | 280145.94 | 193279.14 | 356440.34 | 0 | 206187.62 | 256996.17 | 305959.16 | 221970.56 | 255668.69 | 11 | **1.170** |
| Q8K1X4 | Nckap1l | Nck-associated protein 1-like (Mus musculus) | 1104.11 | 21.52 | 128905 | 21 | 23424484 | 25846258 | 28399808 | 23421548 | 20729148 | 27377292 | 27554070 | 30179064 | 26342216 | 29746782 | 3 | **1.159** |
| Q8K224 | Nat10 | RNA cytidine acetyltransferase (Mus musculus) | 321.24 | 7.71 | 115419 | 5 | 1185470.12 | 3002435.25 | 2498856.5 | 2031704.38 | 1873161.75 | 2805873 | 3099708.75 | 2884527.75 | 2686820.5 | 3290468.25 | 3 | **1.394** |
| Q8K296 | Mtmr3 | Myotubularin-related protein 3 (Mus musculus) | 100.58 | 2.42 | 133840 | 2 | 0 | 44540.85 | 49058.41 | 125322.03 | 121354.76 | 0 | 30513.84 | 198134.41 | 192025.84 | 207288.31 | 9 | **1.845** |
| Q8K297 | Colgalt1 | Procollagen galactosyltransferase 1 (Mus musculus) 1 | 374.85 | 13.29 | 71061 | 8 | 8208723.5 | 10977550 | 11324589 | 10741527 | 10060518 | 8684635 | 11880655 | 11928122 | 12025792 | 12694703 | 4 | **1.115** |
| Q8K298 | Anln | Anillin (Mus musculus) | 131.71 | 2.85 | 122794 | 2 | 1201599.75 | 1118871.62 | 1269263.25 | 674287 | 740557.06 | 1200473.5 | 1086062.25 | 774907.12 | 1066113.75 | 925765 | 11 | **1.010** |
| Q8K2B3 | Sdha | Succinate dehydrogenase [ubiquinone] flavoprotein subunit, mitochondrial (Mus musculus) | 677.61 | 20.48 | 72585 | 10 | 40665028 | 20774454 | 23848358 | 22295106 | 25114164 | 26425688 | 26612980 | 24957886 | 21403152 | 25616730 | 9 | **0.942** |
| Q8K2C9 | Hacd3 | Very-long-chain (3R)-3-hydroxyacyl-CoA dehydratase 3 (Mus musculus) | 51.75 | 2.76 | 43131 | 1 | 533772.38 | 662629.62 | 762020.94 | 444557.38 | 300526.41 | 673689.19 | 641554.88 | 986066.19 | 698390.38 | 796581.75 | 4 | **1.404** |
| Q8K2D3 | Edc3 | Enhancer of mRNA-decapping protein 3 (Mus musculus) | 52.87 | 4.53 | 55957 | 1 | 0 | 174005.53 | 162050.27 | 177284.48 | 141125.56 | 0 | 0 | 221427.27 | 274586.69 | 0 | 15 | **0.758** |
| Q8K2I4 | Manba | Beta-mannosidase (Mus musculus) | 399.31 | 10.81 | 100831 | 8 | 5179125.5 | 6194666 | 9878734 | 5240585 | 7015025.5 | 6866145.5 | 7433913 | 6653277 | 5143374.5 | 7512724.5 | 11 | **1.003** |
| Q8K2L8 | Trappc12 | Trafficking protein particle complex subunit 12 (Mus musculus) 2 | 40.91 | 1.13 | 87694 | 1 | 1067669.25 | 907994.69 | 934928.56 | 0 | 944426.12 | 1105292.62 | 1008191.62 | 879786.25 | 826014.56 | 798117.12 | 12 | **1.198** |
| Q8K2Q0 | Commd9 | COMM domain-containing protein 9 (Mus musculus) | 55.42 | 8.08 | 21850 | 1 | 249160.7 | 508837.88 | 332058.88 | 180359.3 | 423595.91 | 270142.31 | 420896.5 | 264441.97 | 303557.34 | 291413.81 | 11 | **0.915** |
| Q8K2Q7 | Brox | BRO1 domain-containing protein BROX (Mus musculus) | 302.09 | 17.76 | 46202 | 6 | 8339041.5 | 6588113.5 | 10142438 | 7900088 | 3339790 | 9850802 | 10463721 | 9461201 | 9105892 | 9894054 | 4 | **1.343** |
| Q8K2T1 | Nmral1 | NmrA-like family domain-containing protein 1 (Mus musculus) | 88.84 | 11.97 | 34376 | 2 | 3192421.25 | 555065 | 1166108.5 | 1008623.31 | 2549948.75 | 3218119.75 | 3441596.5 | 3003998 | 3301908 | 976285.88 | 5 | **1.646** |
| Q8K2V6 | Ipo11 | Importin-11 (Mus musculus) | 188.5 | 6.77 | 112416 | 4 | 439502.81 | 2150723.25 | 1896617 | 2675393.5 | 2210779 | 1344626.75 | 702376.38 | 2408685 | 1770169.5 | 1787656.5 | 8 | **0.855** |
| Q8K2X3 | Stn1 | CST complex subunit STN1 (Mus musculus) | 109.97 | 6.61 | 43486 | 2 | 2690917 | 1585224.5 | 1760101 | 2266021.5 | 1895347.38 | 1501374.62 | 1433193 | 1678454 | 1995695.5 | 1854650 | 6 | **0.830** |
| Q8K2Z4 | Ncapd2 | Condensin complex subunit 1 (Mus musculus) | 273.34 | 5.03 | 155665 | 6 | 3405925.5 | 6063135 | 5819587.5 | 6761356 | 3386562 | 5937553 | 5651811.5 | 6290935 | 4897930.5 | 5807015.5 | 12 | **1.124** |
| Q8K301 | Ddx52 | Probable ATP-dependent RNA helicase DDX52 (Mus musculus) | 96.63 | 5.85 | 67474 | 2 | 473462 | 866220.94 | 778143 | 580437.75 | 365363.25 | 729465.56 | 816410.69 | 892489.81 | 461065.53 | 351719.84 | 12 | **1.061** |
| Q8K310 | Matr3 | Matrin-3 (Mus musculus) | 354.65 | 11.58 | 94630 | 7 | 11933473 | 11576359 | 10471009 | 10498839 | 13311472 | 11923887 | 12059776 | 12668371 | 11360250 | 12574720 | 8 | **1.048** |
| Q8K363 | Ddx18 | ATP-dependent RNA helicase DDX18 (Mus musculus) | 736.12 | 26.21 | 74181 | 11 | 12525212 | 9477523 | 8783961 | 7319419.5 | 10376472 | 10182154 | 13465012 | 12424243 | 12045759 | 11681268 | 5 | **1.233** |
| Q8K368 | Fanci | Fanconi anemia group I protein homolog (Mus musculus) | 105.22 | 1.35 | 149326 | 2 | 0 | 222148.33 | 213229.16 | 149986.97 | 93398.73 | 0 | 116923.12 | 157597.16 | 181104.2 | 161474.66 | 12 | **0.909** |
| Q8K3G9 | Appl2 | DCC-interacting protein 13-beta (Mus musculus) | 108.12 | 4.68 | 73854 | 2 | 2103850.5 | 1606969.62 | 1531819.25 | 1009537.06 | 1297722.12 | 2037872 | 1774184.38 | 2019963.75 | 1675858.75 | 1649498.5 | 5 | **1.213** |
| Q8K3J1 | Ndufs8 | NADH dehydrogenase [ubiquinone] iron-sulfur protein 8, mitochondrial (Mus musculus) | 111.68 | 9.43 | 24038 | 2 | 628977.44 | 626398.12 | 809470 | 1238229.5 | 1683526.88 | 1674808.75 | 1874390.25 | 2291224 | 1282851.38 | 1917800 | 2 | **1.813** |
| Q8K400 | Stxbp5 | Syntaxin-binding protein 5 (Mus musculus) | 42.39 | 1.04 | 127651 | 1 | 715876.25 | 0 | 0 | 561907.44 | 696897.75 | 903560.5 | 735196.62 | 881419.38 | 829901.06 | 541556.12 | 4 | **1.971** |
| Q8K411 | Pitrm1 | Presequence protease, mitochondrial (Mus musculus) | 61.84 | 1.54 | 117372 | 1 | 613700.19 | 0 | 800444.75 | 0 | 0 | 778772.19 | 620212.31 | 660707.75 | 792519.81 | 838692.75 | 7 | **2.610** |
| Q8K4I3 | Arhgef6 | Rho guanine nucleotide exchange factor 6 (Mus musculus) | 149.7 | 4.67 | 87051 | 3 | 0 | 429542.5 | 681568.25 | 0 | 500639.31 | 209678.28 | 883534.38 | 674080.5 | 477882.56 | 760134.38 | 7 | **1.865** |
| Q8K4Q0 | Rptor | Regulatory-associated protein of mTOR (Mus musculus) | 69.85 | 1.42 | 149471 | 1 | 424078.09 | 519782.91 | 493835.31 | 473014.41 | 463861.25 | 508778.53 | 532091.69 | 642180.19 | 430008.44 | 474359.53 | 7 | **1.090** |
| Q8K4Q8 | Colec12 | Collectin-12 (Mus musculus) | 617.91 | 15.9 | 81304 | 11 | 34665272 | 32040668 | 32598034 | 33821788 | 29044056 | 34319224 | 31183560 | 33941920 | 34056712 | 36175760 | 7 | **1.046** |
| Q8K4Z5 | Sf3a1 | Splicing factor 3A subunit 1 (Mus musculus) | 813.23 | 26.68 | 88545 | 16 | 40536500 | 33661636 | 32063712 | 33565896 | 31877576 | 32264684 | 31685604 | 31706980 | 32040892 | 33916204 | 7 | **0.941** |
| Q8K581 | Txndc9 | Thioredoxin domain-containing protein 9 (Rattus norvegicus) | 41.45 | 3.98 | 26272 | 1 | 3506580 | 2743171.75 | 2669357.25 | 3197708.75 | 2102496.5 | 4154339 | 3411770.25 | 2910958.75 | 3169442.5 | 2289966.25 | 9 | **1.121** |
| Q8QZS1 | Hibch | 3-hydroxyisobutyryl-CoA hydrolase, mitochondrial (Mus musculus) | 160.06 | 11.43 | 43038 | 3 | 1222847.25 | 1520938.88 | 2631867.5 | 4690909.5 | 3314901 | 2060985.5 | 1853398.25 | 2424456.75 | 2894611.5 | 2468634.5 | 11 | **0.874** |
| Q8QZT1 | Acat1 | Acetyl-CoA acetyltransferase, mitochondrial (Mus musculus) | 695.59 | 34.43 | 44816 | 11 | 47797176 | 24916394 | 31058404 | 22640516 | 31066684 | 40306112 | 36461416 | 39569936 | 39225188 | 41821472 | 5 | **1.253** |
| Q8QZY1 | Eif3l | Eukaryotic translation initiation factor 3 subunit L (Mus musculus) | 1243.61 | 39.18 | 66613 | 19 | 16601864 | 36995104 | 32954694 | 39645552 | 27915340 | 32720476 | 25917614 | 31252520 | 31720626 | 35812208 | 10 | **1.021** |
| Q8R016 | Blmh | Bleomycin hydrolase (Mus musculus) | 530.75 | 32.97 | 52511 | 11 | 21455928 | 14804096 | 12225815 | 15986508 | 12413886 | 23218560 | 19288688 | 18699474 | 18189524 | 18073352 | 4 | **1.268** |
| Q8R050 | Gspt1 | Eukaryotic peptide chain release factor GTP-binding subunit ERF3A (Mus musculus) | 481.58 | 20.28 | 68625 | 8 | 20163406 | 19401406 | 18575852 | 19161814 | 17029720 | 17025802 | 20307518 | 15990679 | 16699068 | 18045154 | 6 | **0.934** |
| Q8R060 | Zwilch | Protein zwilch homolog (Mus musculus) | 75.45 | 1.7 | 66839 | 1 | 739580.25 | 714957.69 | 631045.5 | 657743.19 | 797058.5 | 818074.44 | 679087.69 | 839695.44 | 879627.56 | 771445.38 | 4 | **1.126** |
| Q8R0F3 | Sumf1 | Formylglycine-generating enzyme (Mus musculus) | 96.32 | 8.6 | 40659 | 2 | 1302399.88 | 1216378.38 | 1232199.88 | 835851.56 | 1135175.25 | 1166348.88 | 1109760.5 | 1017787.25 | 998492.44 | 1226941.5 | 8 | **0.965** |
| Q8R0F6 | Ilkap | Integrin-linked kinase-associated serine/threonine phosphatase 2C (Mus musculus) | 143.84 | 7.4 | 42774 | 3 | 3583390.25 | 1881357.25 | 1633394.88 | 2379962.5 | 2304557.25 | 2715986.25 | 3029296.5 | 2002046 | 2006045 | 1532594.75 | 12 | **0.958** |
| Q8R0G9 | Nup133 | Nuclear pore complex protein Nup133 (Mus musculus) | 418.98 | 10.48 | 128620 | 7 | 1156218 | 1862064 | 2456994 | 3775920.75 | 2929527.75 | 4509538 | 3383138 | 3978905.25 | 3798858.5 | 4191582.5 | 1 | **1.631** |
| Q8R0H9 | Gga1 | ADP-ribosylation factor-binding protein GGA1 (Mus musculus) | 133.7 | 5.83 | 69972 | 2 | 215705.12 | 321153.34 | 383992.19 | 2009137.75 | 2028978.88 | 321834.16 | 247341.28 | 279065.34 | 2251067.25 | 2503024.5 | 11 | **1.130** |
| Q8R0W0 | Eppk1 | Epiplakin (Mus musculus) | 202.52 | 1.88 | 724630 | 4 | 7404321 | 13169506 | 10896575 | 12284232 | 10593701 | 10347846 | 11912169 | 10952048 | 11376798 | 10933286 | 12 | **1.022** |
| Q8R0X7 | Sgpl1 | Sphingosine-1-phosphate lyase 1 (Mus musculus) | 432.94 | 19.37 | 63677 | 8 | 7432917 | 6359404 | 6193373.5 | 5437047 | 4349817 | 8488538 | 7337472 | 6966844.5 | 6714103 | 6516144 | 4 | **1.210** |
| Q8R123 | Flad1 | FAD synthase (Mus musculus) | 44.91 | 2.64 | 54766 | 1 | 413826.34 | 518412 | 364418.88 | 144874.66 | 338168.69 | 399716.16 | 392713.31 | 374882.25 | 384565.41 | 333194.69 | 12 | **1.059** |
| Q8R127 | Sccpdh | Saccharopine dehydrogenase-like oxidoreductase (Mus musculus) | 255.66 | 17.02 | 47129 | 5 | 3220017.75 | 2563060.5 | 2406588.75 | 2716091.5 | 3718444 | 2477550 | 3953402 | 3177154.5 | 2497744 | 1903807.5 | 10 | **0.958** |
| Q8R146 | Apeh | Acylamino-acid-releasing enzyme (Mus musculus) | 116.29 | 3.01 | 81581 | 2 | 2008281 | 3181040.5 | 3444972.5 | 885949.56 | 2747420.25 | 3316044.25 | 2575919.75 | 2691519.25 | 2809861 | 2625488.5 | 12 | **1.143** |
| Q8R164 | Bphl | Valacyclovir hydrolase (Mus musculus) | 68.77 | 6.87 | 32851 | 1 | 0 | 173812.17 | 140515.69 | 0 | 199760.92 | 182402.53 | 254703.39 | 217433.62 | 162066.61 | 125694.25 | 7 | **1.833** |
| Q8R180 | Ero1a | ERO1-like protein alpha (Mus musculus) | 988.1 | 42.24 | 54084 | 16 | 38574724 | 40042600 | 40335780 | 35194888 | 40161032 | 47505636 | 42997684 | 44930304 | 45186976 | 45939628 | 0 | **1.166** |
| Q8R1A4 | Dock7 | Dedicator of cytokinesis protein 7 (Mus musculus) | 84.8 | 1.46 | 241438 | 2 | 159083.16 | 713434.81 | 435968.44 | 235316.88 | 177967.86 | 130068.54 | 681667.19 | 793260.88 | 98122.79 | 824343.38 | 11 | **1.468** |
| Q8R1F1 | Fam129b | Niban-like protein 1 (Mus musculus) | 382.12 | 13.75 | 84819 | 7 | 5611662 | 6151221 | 8467062 | 6080701.5 | 7514323.5 | 8272091.5 | 9697897 | 8698157 | 8613213 | 8574286 | 1 | **1.297** |
| Q8R1F5 | Hyi | Putative hydroxypyruvate isomerase (Mus musculus) | 98.43 | 10.47 | 30449 | 2 | 0 | 353478 | 236329.69 | 279376.16 | 216507.41 | 0 | 0 | 41113.61 | 54358.23 | 66737.26 | 6 | **0.149** |
| Q8R1I1 | Uqcr10 | Cytochrome b-c1 complex subunit 9 (Mus musculus) | 50.56 | 10.94 | 7446 | 1 | 3101289.25 | 2438258 | 2835532.25 | 0 | 2196647 | 3532666 | 3910460.5 | 3306670.75 | 3820523.25 | 3146834 | 0 | **1.676** |
| Q8R1Q8 | Dync1li1 | Cytoplasmic dynein 1 light intermediate chain 1 (Mus musculus) 1 | 290.18 | 11.09 | 56614 | 5 | 8463263 | 11080932 | 10994810 | 16218793 | 10806518 | 9997054 | 11048271 | 11210875 | 12787583 | 15897336 | 9 | **1.059** |
| Q8R1S4 | Mtss1 | Metastasis suppressor protein 1 (Mus musculus) | 202.58 | 7.64 | 82408 | 4 | 5824279 | 1703488.5 | 2362279.75 | 791122.44 | 4475268.5 | 5724760.5 | 4050104.75 | 5053899 | 4642277 | 4265654 | 7 | **1.566** |
| Q8R1T1 | Chmp7 | Charged multivesicular body protein 7 (Mus musculus) | 61.81 | 3.77 | 50633 | 1 | 396327.06 | 248449.45 | 227235.81 | 0 | 279025.5 | 202479.23 | 287424.53 | 33805.91 | 241451.27 | 280312.34 | 12 | **0.908** |
| Q8R1V4 | Tmed4 | Transmembrane emp24 domain-containing protein 4 (Mus musculus) | 126.11 | 12.33 | 26022 | 2 | 3208289 | 3900714 | 3612902.25 | 3891101.25 | 3404396.5 | 2762168 | 3146432.5 | 3098104 | 3068116.5 | 3078101.75 | 0 | **0.841** |
| Q8R1X6 | Spart | Spartin (Mus musculus) | 130.8 | 9.54 | 72655 | 3 | 2877756.25 | 1533336 | 1403964.12 | 2979199.75 | 3597816.25 | 2559164.25 | 2238802.75 | 2433695 | 2857537.5 | 3436477.75 | 12 | **1.091** |
| Q8R2E9 | Ero1b | ERO1-like protein beta (Mus musculus) | 281.85 | 13.28 | 53518 | 5 | 11278067 | 9894715 | 10934055 | 9760840 | 11036819 | 10887593 | 12362283 | 10792161 | 11330684 | 11343491 | 6 | **1.072** |
| Q8R2K1 | Fuom | Fucose mutarotase (Mus musculus) | 134.84 | 17.65 | 16805 | 2 | 163293.06 | 1799300.12 | 1375393.75 | 1721390.38 | 1041749.56 | 176347.02 | 372607.88 | 657602.81 | 351030.69 | 367250.75 | 5 | **0.315** |
| Q8R2N2 | Utp4 | U3 small nucleolar RNA-associated protein 4 homolog (Mus musculus) | 67.05 | 3.06 | 76909 | 1 | 0 | 442100.22 | 361520.03 | 314380.47 | 235592.55 | 442561.97 | 99282.55 | 476823.12 | 335743.75 | 230409.67 | 10 | **1.171** |
| Q8R2U0 | Seh1l | Nucleoporin SEH1 (Mus musculus) | 63.84 | 5 | 39775 | 2 | 1612536.12 | 1179644.88 | 1418205.12 | 1536632.88 | 1817874 | 2241012 | 415911.31 | 1531525 | 1654818.75 | 1352897.38 | 12 | **0.951** |
| Q8R2W9 | Pank3 | Pantothenate kinase 3 (Mus musculus) | 89.02 | 5.41 | 41120 | 2 | 2354291.75 | 1362725.62 | 1266687.12 | 1271814 | 1455872.38 | 2376328.5 | 1867329.62 | 1192662.75 | 1477265 | 1244082.12 | 12 | **1.058** |
| Q8R2Y8 | Ptrh2 | Peptidyl-tRNA hydrolase 2, mitochondrial (Mus musculus) | 115.76 | 19.89 | 19527 | 2 | 1477312.5 | 1163753 | 992000.69 | 474497.75 | 0 | 1303827 | 1364898.75 | 889731 | 1027663.62 | 739245.62 | 10 | **1.296** |

| | | | | | | | | | | | | | | | | | |
|---|---|---|---|---|---|---|---|---|---|---|---|---|---|---|---|---|---|
| Q8R307 | Vps18 | Vacuolar protein sorting-associated protein 18 homolog (Mus musculus) | 60.3 | 1.85 | 110219 | 1 | 0 | 186804.19 | 0 | 351346.56 | 221221.41 | 42005.78 | 173954.84 | 0 | 317599.47 | 284166.62 | 14 | **1.077** |
| Q8R311 | Ctage5 | Endoplasmic reticulum export factor CTAGE5 (Mus musculus) | 98.3 | 3.47 | 87718 | 2 | 99166.52 | 1181604.38 | 1422194.5 | 1028261 | 838791.31 | 675608.62 | 854865.69 | 1168750.5 | 1212304.75 | 1256754.38 | 11 | **1.131** |
| Q8R313 | Exoc6 | Exocyst complex component 6 (Mus musculus) | 192.01 | 5.86 | 93077 | 4 | 2115859.5 | 2109036.5 | 2402905.25 | 1608645.75 | 1786543.25 | 1944030.75 | 1521034.12 | 1910751.75 | 2046461.75 | 2084097.5 | 8 | **0.948** |
| Q8R323 | Rfc3 | Replication factor C subunit 3 (Mus musculus) | 361.12 | 16.29 | 40526 | 5 | 3367347 | 3486049 | 4227290.5 | 2690401 | 2139675.25 | 3620058 | 4186438.75 | 2600886.5 | 3948206.5 | 3464339.25 | 9 | **1.120** |
| Q8R3C0 | Mcmbp | Mini-chromosome maintenance complex-binding protein (Mus musculus) | 547.09 | 21.96 | 72891 | 9 | 11145805 | 12333293 | 7351883 | 9457109 | 8572677 | 9887828 | 7455373 | 9881316 | 11501494 | 10377224 | 11 | **1.005** |
| Q8R3C6 | Rbm19 | Probable RNA-binding protein 19 (Mus musculus) | 173.26 | 5.46 | 106083 | 4 | 1483509.75 | 2458983 | 1768648.12 | 105818.85 | 1337486.75 | 1310325 | 1761674.88 | 2751989.5 | 1425827.5 | 2033781.5 | 10 | **1.298** |
| Q8R3H7 | Hs2st1 | Heparan sulfate 2-O-sulfotransferase 1 (Mus musculus) | 248.59 | 18.82 | 41813 | 4 | 1132574.88 | 2610502 | 1561326.5 | 2962999.25 | 1988897 | 2316445.25 | 1034331 | 2720175.5 | 1398156.5 | 3079525.5 | 12 | **1.029** |
| Q8R3N1 | Nop14 | Nucleolar protein 14 (Mus musculus) | 80.12 | 3.84 | 98769 | 2 | 199062.28 | 655055.75 | 709691.31 | 799810.62 | 350904.56 | 333644.28 | 352677.66 | 754317.19 | 735842.75 | 683719.75 | 11 | **1.054** |
| Q8R3N6 | Thoc1 | THO complex subunit 1 (Mus musculus) | 136.17 | 4.72 | 75436 | 3 | 3595440.25 | 3427751.25 | 1601796.75 | 3364017.5 | 2775374.5 | 2883855.5 | 3199871.25 | 2844682.5 | 2875114.5 | 2985400.75 | 10 | **1.002** |
| Q8R3S6 | Exoc1 | Exocyst complex component 1 (Mus musculus) | 40.31 | 2.46 | 101889 | 1 | 115154.36 | 481025.47 | 207800.98 | 0 | 363442.5 | 148049.91 | 307447.69 | 174557.67 | 211748.47 | 233270.19 | 12 | **0.921** |
| Q8R404 | Mic13 | MICOS complex subunit MIC13 (Mus musculus) | 114.52 | 22.69 | 13373 | 2 | 5753747 | 4217147.5 | 4070110.5 | 4499802 | 4215501 | 284352.88 | 4864205 | 3574306 | 3691041.75 | 3890656.75 | 4 | **0.716** |
| Q8R418 | Dicer1 | Endoribonuclease Dicer (Mus musculus) | 55.71 | 0.52 | 216821 | 1 | 0 | 519319.5 | 352566.84 | 347145.59 | 254404.28 | 325726 | 324185.09 | 317184.38 | 360605.66 | 317467.16 | 12 | **1.117** |
| Q8R424 | Stambp | STAM-binding protein (Rattus norvegicus) | 47.03 | 4.72 | 48512 | 1 | 1504606 | 1113132.5 | 902520.25 | 1178702 | 1173683.62 | 1336330.25 | 1157601 | 1325464.5 | 1178960.75 | 883290.06 | 11 | **1.002** |
| Q8R480 | Nup85 | Nuclear pore complex protein Nup85 (Mus musculus) | 152.46 | 5.34 | 74776 | 2 | 194485.61 | 513302.12 | 268128.66 | 234157 | 186503.25 | 172866.39 | 383324.31 | 230671.47 | 116333.43 | 339032.28 | 10 | **0.889** |
| Q8R491 | Ehd3 | EH domain-containing protein 3 (Rattus norvegicus) | 686.92 | 25.42 | 60791 | 15 | 26450998 | 23691832 | 23766442 | 22669970 | 18609376 | 26768132 | 26608716 | 27820790 | 26214320 | 24887792 | 2 | **1.149** |
| Q8R4R6 | Nup35 | Nucleoporin NUP35 (Mus musculus) | 115.32 | 8.62 | 34786 | 2 | 780937.62 | 458989.91 | 469833.62 | 457313.34 | 828674.38 | 457780.03 | 800751.44 | 566225.88 | 614135.31 | 693145.81 | 11 | **1.045** |
| Q8R4Y8 | Rttn | Rotatin (Mus musculus) | 41.7 | 0.31 | 248489 | 1 | 1868501.25 | 1058635.38 | 55552884 | 34922152 | 40657224 | 332731.41 | 423238.41 | 51281008 | 50881028 | 44605372 | 12 | **1.100** |
| Q8R570 | Snap47 | Synaptosomal-associated protein 47 (Mus musculus) | 90.32 | 7.02 | 46524 | 2 | 0 | 392078.44 | 333858.72 | 341121.97 | 258091.88 | 0 | 0 | 282389.38 | 316531.59 | 213494.12 | 8 | **0.613** |
| Q8R5A6 | Tbc1d22a | TBC1 domain family member 22A (Mus musculus) a | 45.61 | 1.55 | 59362 | 1 | 0 | 482038.97 | 504999.34 | 0 | 468769.12 | 0 | 148132.31 | 0 | 495986.84 | 0 | 15 | **0.442** |
| Q8R5F7 | Ifih1 | Interferon-induced helicase C domain-containing protein 1 (Mus musculus) | 60.08 | 1.17 | 115971 | 1 | 0 | 352920.5 | 0 | 0 | 162515.86 | 0 | 0 | 92681.15 | 291649.94 | 264868.5 | 17 | **1.260** |
| Q8R5H1 | Usp15 | Ubiquitin carboxyl-terminal hydrolase 15 (Mus musculus) | 309.14 | 8.56 | 112325 | 7 | 3529183.5 | 3100872.75 | 4125773.25 | 3899605.5 | 3779905.75 | 4001110 | 3939680 | 3271187 | 4061862.25 | 4512016 | 7 | **1.073** |
| Q8R5J9 | Arl6ip5 | PRA1 family protein 3 (Mus musculus) | 41.3 | 5.85 | 21558 | 1 | 895266.12 | 0 | 0 | 0 | 0 | 2084234.88 | 1745388.5 | 1686900.75 | 1771756 | 0 | 11 | **8.141** |
| Q8R5K4 | Nol6 | Nucleolar protein 6 (Mus musculus) | 45.35 | 0.61 | 129228 | 1 | 138399.69 | 670603.31 | 664420.31 | 1253887 | 590587.56 | 0 | 649601.5 | 1265090.25 | 1030366.5 | 938935.56 | 10 | **1.171** |
| Q8VBT0 | Tmx1 | Thioredoxin-related transmembrane protein 1 (Mus musculus) | 191.31 | 13.31 | 31396 | 3 | 5216532 | 8945320 | 8226320.5 | 9987873 | 8654013 | 8941000 | 7013927 | 8183820 | 7434533 | 8027627.5 | 7 | **0.965** |
| Q8VBT6 | Apobr | Apolipoprotein B receptor (Mus musculus) | 671.01 | 16.88 | 102705 | 12 | 18869924 | 18795624 | 15759059 | 18220176 | 19504328 | 14407709 | 14199821 | 16097994 | 14954116 | 15528605 | 1 | **0.825** |
| Q8VBT9 | Aspscr1 | Tether containing UBX domain for GLUT4 (Mus musculus) | 45.78 | 2.91 | 59796 | 1 | 494471.38 | 461134.56 | 353281.28 | 0 | 106791.16 | 316176.56 | 452470.91 | 278450.56 | 0 | 342642.41 | 10 | **0.982** |
| Q8VBV3 | Exosc2 | Exosome complex component RRP4 (Mus musculus) | 65.14 | 5.12 | 32632 | 1 | 1866366.25 | 0 | 0 | 845305.38 | 1298177 | 1931319.88 | 1614217.5 | 1790636.38 | 1685233.12 | 1659091.38 | 5 | **2.165** |
| Q8VBW6 | Nae1 | NEDD8-activating enzyme E1 regulatory subunit (Mus musculus) | 517.92 | 27.9 | 60274 | 9 | 10039916 | 6877148 | 7889351 | 7390069.5 | 4660933 | 6480966 | 6835003.5 | 6381914 | 7166724 | 8308205.5 | 9 | **0.954** |
| Q8VBZ3 | Clptm1 | Cleft lip and palate transmembrane protein 1 homolog (Mus musculus) | 296.82 | 14.91 | 75291 | 5 | 5130690 | 4978695 | 3892109.5 | 3448561.75 | 1787061.38 | 4403296 | 4541368.5 | 4362799 | 4740963 | 4177178 | 10 | **1.155** |
| Q8VC03 | Eml3 | Echinoderm microtubule-associated protein-like 3 (Mus musculus) | 48.32 | 1.78 | 95695 | 1 | 870948.81 | 491904.53 | 0 | 0 | 591726.56 | 570128.12 | 722344.88 | 592219.81 | 542884.44 | 550194 | 9 | **1.523** |
| Q8VC04 | Tmem106a | Transmembrane protein 106A (Mus musculus) a | 57.31 | 6.9 | 29109 | 1 | 0 | 2469937.5 | 2407956 | 2941212.75 | 3251823.5 | 2272143.25 | 2577363.75 | 3029092.5 | 2625790.25 | 2654969.25 | 11 | **1.189** |
| Q8VC85 | Lsm1 | U6 snRNA-associated Sm-like protein LSm1 (Mus musculus) | 39.93 | 5.26 | 15236 | 1 | 1089221.5 | 806560.5 | 709568.5 | 947139.5 | 840918.06 | 1214802.12 | 952527.06 | 949706.62 | 1014246.81 | 766369.38 | 7 | **1.115** |
| Q8VCF0 | Mavs | Mitochondrial antiviral-signaling protein (Mus musculus) | 201.68 | 8.55 | 53399 | 3 | 3582780.25 | 1356707.75 | 2275202 | 1032169.81 | 1260316.25 | 2508060.75 | 2764998.25 | 2197776 | 2786789 | 2804493 | 6 | **1.374** |
| Q8VCH8 | Ubxn4 | UBX domain-containing protein 4 (Mus musculus) | 48.88 | 2.77 | 56472 | 1 | 0 | 991715.19 | 528474.12 | 1653552.62 | 701430.06 | 0 | 0 | 280120.59 | 357119.75 | 0 | 8 | **0.164** |
| Q8VCN5 | Cth | Cystathionine gamma-lyase (Mus musculus) | 637.47 | 38.44 | 43567 | 11 | 17113654 | 21175776 | 18188864 | 24434584 | 16760536 | 18996936 | 18140400 | 17386318 | 15815738 | 16219205 | 7 | **0.886** |
| Q8VCN9 | Tbcc | Tubulin-specific chaperone C (Mus musculus) | 67.69 | 4.4 | 38125 | 1 | 112413.36 | 84703.91 | 228156.75 | 196841.36 | 68533.8 | 319362.41 | 283624.5 | 194546.83 | 58268.71 | 202906.44 | 8 | **1.533** |
| Q8VCT3 | Rnpep | Aminopeptidase B (Mus musculus) | 1099.95 | 42.77 | 72416 | 19 | 37543616 | 39896060 | 39859532 | 40367988 | 43980264 | 46521512 | 43551884 | 45382620 | 43506696 | 43252744 | 3 | **1.102** |
| Q8VCW4 | Unc93b1 | Protein unc-93 homolog B1 (Mus musculus) | 159.54 | 8.36 | 66981 | 3 | 1641689.25 | 1176171.5 | 2417022.5 | 413106.09 | 1138194.75 | 3504548.5 | 2032838.75 | 3562829.5 | 2060364.75 | 2185261 | 3 | **1.967** |
| Q8VCW8 | Acsf2 | Acyl-CoA synthetase family member 2, mitochondrial (Mus musculus) | 42.53 | 2.44 | 67951 | 1 | 2013130.62 | 1301908.88 | 1272454.12 | 1626654.25 | 988704.88 | 1442607.12 | 1669674.38 | 1211094.75 | 1487094.12 | 1757294.62 | 10 | **1.051** |
| Q8VD75 | Hip1 | Huntingtin-interacting protein 1 (Mus musculus) | 63.42 | 1.94 | 115202 | 1 | 95937.62 | 0 | 452547.69 | 0 | 0 | 228851.47 | 0 | 424107.81 | 0 | 324006.59 | 16 | **1.781** |
| Q8VDD5 | Myh9 | Myosin-9 (Mus musculus) | 8025.52 | 59.59 | 226372 | 117 | 732211072 | 690678976 | 684426816 | 683302720 | 743336896 | 676265792 | 616595968 | 648994048 | 673755776 | 645019776 | 0 | **0.923** |
| Q8VDF2 | Uhrf1 | E3 ubiquitin-protein ligase UHRF1 (Mus musculus) | 608.95 | 17.52 | 88304 | 11 | 2318648.25 | 6951086.5 | 6302897.5 | 8530499 | 7549758.5 | 3910169.25 | 3508799.75 | 5164247 | 6831831.5 | 6219281 | 6 | **0.810** |
| Q8VDJ3 | Hdlbp | Vigilin (Mus musculus) | 1445.2 | 26.74 | 141743 | 25 | 52119616 | 45024600 | 48312688 | 43109372 | 47509160 | 53281004 | 49005576 | 49950712 | 46107084 | 49952108 | 6 | **1.052** |
| Q8VDL4 | Adpgk | ADP-dependent glucokinase (Mus musculus) | 350.89 | 18.55 | 53902 | 6 | 15208655 | 14304656 | 12950385 | 11645482 | 9990232 | 12451157 | 12909954 | 12049621 | 11799233 | 11919136 | 10 | **0.954** |
| Q8VDM4 | Psmd2 | 26S proteasome non-ATPase regulatory subunit 2 (Mus musculus) | 1768.22 | 39.21 | 100203 | 28 | 59458132 | 68582032 | 64331120 | 70685160 | 50794668 | 47057408 | 52498596 | 53908604 | 51968680 | 53945444 | 4 | **0.826** |
| Q8VDM6 | Hnrnpul1 | Heterogeneous nuclear ribonucleoprotein U-like protein 1 (Mus musculus) 1 | 175.45 | 5.47 | 96002 | 4 | 5363235.5 | 2488971.5 | 3791635.25 | 3295594.5 | 3102315.5 | 3624096 | 4528011 | 4286889.5 | 4265966 | 4265177 | 6 | **1.162** |
| Q8VDP4 | Ccar2 | Cell cycle and apoptosis regulator protein 2 (Mus musculus) | 141.86 | 5.1 | 103002 | 3 | 3127964 | 2471909.5 | 2336096.25 | 1597874.62 | 2287027.5 | 2694107 | 2833584.5 | 3021392 | 2560098.25 | 2536681 | 5 | **1.154** |
| Q8VDP6 | Cdipt | CDP-diacylglycerol--inositol 3-phosphatidyltransferase (Mus musculus) | 226.13 | 24.88 | 23599 | 5 | 4776480 | 4624201.5 | 3179155.25 | 2116387.25 | 1899961.75 | 5643121.5 | 6362668 | 6144849 | 5467401.5 | 5867800.5 | 0 | **1.777** |
| Q8VDV3 | Rab3il1 | Guanine nucleotide exchange factor for Rab-3A (Mus musculus) | 97.31 | 8.88 | 42713 | 2 | 1570346.75 | 452226.69 | 546776.56 | 0 | 119986.94 | 326481.34 | 0 | 1121619.12 | 6155680 | 902868.19 | 10 | **3.163** |
| Q8VDW0 | Ddx39a | ATP-dependent RNA helicase DDX39A (Mus musculus) | 873.99 | 49.41 | 49067 | 17 | 155376352 | 134881392 | 131562760 | 133910144 | 142143632 | 159357104 | 139754672 | 151615680 | 146542592 | 146370352 | 5 | **1.066** |
| Q8VE10 | Naa40 | N-alpha-acetyltransferase 40 (Mus musculus) | 57.63 | 5.06 | 27229 | 1 | 1751114.5 | 1509969.5 | 1749963.62 | 1981886 | 1799503.75 | 1640435.88 | 1588761.88 | 1524381.62 | 1401055.25 | 1282335.12 | 3 | **0.846** |
| Q8VE22 | Mrps23 | 28S ribosomal protein S23, mitochondrial (Mus musculus) | 86.21 | 11.3 | 20348 | 2 | 955057.56 | 1082272.88 | 1432959.5 | 0 | 1270801.38 | 1150152 | 1137567.5 | 1369085.62 | 1483312.25 | 1310529.62 | 6 | **1.361** |
| Q8VE37 | Rcc1 | Regulator of chromosome condensation (Mus musculus) | 297.19 | 20.67 | 44931 | 6 | 11338368 | 16990230 | 13780166 | 10947922 | 9559262 | 12395880 | 12409474 | 10375770 | 12343265 | 11668013 | 12 | **0.945** |
| Q8VE88 | Fam114a2 | Protein FAM114A2 (Mus musculus) 2 | 113.25 | 7.65 | 54045 | 2 | 1502610.62 | 1227279.38 | 1082813.25 | 1084332.75 | 1316078.38 | 1570883.75 | 1697267.25 | 886850.81 | 1410542 | 1217239.5 | 9 | **1.092** |
| Q8VE97 | Srsf4 | Serine/arginine-rich splicing factor 4 (Mus musculus) | 250.39 | 9.82 | 55979 | 5 | 46939200 | 40967540 | 46074112 | 44268272 | 14533026 | 36885136 | 36354948 | 40695988 | 42840756 | 41154400 | 7 | **1.027** |
| Q8VE99 | Ccdc115 | Coiled-coil domain-containing protein 115 (Mus musculus) | 164.22 | 31.11 | 19743 | 3 | 1565236 | 807147.38 | 899901.12 | 860186.06 | 745030.38 | 1182617 | 1150429.88 | 1006843.94 | 729740.06 | 1153154.38 | 9 | **1.071** |
| Q8VEB4 | Pla2g15 | Group XV phospholipase A2 (Mus musculus) | 103.93 | 5.1 | 47307 | 2 | 2747560.75 | 3486074.25 | 2787485.25 | 3359958 | 3446273.5 | 2579940 | 2903902 | 2277146.75 | 2644375.75 | 2353637.25 | 2 | **0.806** |
| Q8VEE4 | Rpa1 | Replication protein A 70 kDa DNA-binding subunit (Mus musculus) | 623.84 | 22.79 | 69037 | 13 | 15913625 | 20138026 | 21625658 | 19743940 | 17440818 | 22318244 | 18198898 | 18515644 | 20688536 | 22178966 | 7 | **1.074** |
| Q8VEH3 | Arl8a | ADP-ribosylation factor-like protein 8A (Mus musculus) | 423.62 | 47.31 | 21390 | 8 | 12946490 | 11674934 | 16445471 | 9170743 | 13986508 | 14410406 | 15476588 | 14783878 | 13334219 | 15925761 | 6 | **1.151** |
| Q8VEH6 | Cbwd1 | COBW domain-containing protein 1 (Mus musculus) | 89.21 | 5.6 | 43772 | 1 | 281662.62 | 275935.16 | 131691.28 | 300515.5 | 242163.41 | 358930.91 | 425976.59 | 439254.25 | 365226.62 | 414490.66 | 0 | **1.627** |
| Q8VEJ4 | Nle1 | Notchless protein homolog 1 (Mus musculus) | 186.86 | 10.31 | 53134 | 3 | 599094.69 | 1959568.25 | 1507012.25 | 1175815.38 | 1316437.25 | 508858.53 | 1453740.62 | 1607203.5 | 1435239.5 | 1605691.5 | 11 | **1.008** |
| Q8VEL2 | Mtmr14 | Myotubularin-related protein 14 (Mus musculus) | 37.18 | 1.39 | 72443 | 1 | 263616 | 561798.44 | 981803.56 | 0 | 0 | 663314 | 697848.88 | 795435.25 | 0 | 646773.81 | 10 | **1.551** |
| Q8VH49 | Higd1a | HIG1 domain family member 1A, mitochondrial (Rattus norvegicus) | 164.59 | 43.01 | 10301 | 2 | 1235671.75 | 1705492.75 | 1339173.38 | 758322.44 | 1516519.5 | 842254.94 | 1742864.5 | 2091389.88 | 1894093.25 | 1947617.88 | 4 | **1.299** |
| Q8VH51 | Rbm39 | RNA-binding protein 39 (Mus musculus) | 724.18 | 31.89 | 59407 | 11 | 19205616 | 16834676 | 19533276 | 16479602 | 17739686 | 18541298 | 21561618 | 21082828 | 19031944 | 19967292 | 4 | **1.116** |
| Q8VHI3 | Pofut2 | GDP-fucose protein O-fucosyltransferase 2 (Mus musculus) | 143.48 | 8.39 | 49429 | 3 | 3062032 | 1156795.25 | 3703098.75 | 1141095.75 | 3348158.75 | 3153571.25 | 2659498.75 | 836978.12 | 3092565.25 | 3772370 | 12 | **1.089** |
| Q8VHR5 | Gatad2b | Transcriptional repressor p66-beta (Mus musculus) | 39.15 | 2.19 | 65411 | 1 | 1529200.25 | 0 | 861130.81 | 0 | 876489.88 | 1078579.25 | 0 | 827311.31 | 724947.81 | 722067.75 | 12 | **1.026** |
| Q8VI36 | Pxn | Paxillin (Mus musculus) | 214.36 | 13.71 | 64476 | 5 | 1325196.38 | 1815868.25 | 1808702 | 1609963.5 | 2167805.25 | 430107.5 | 1519493.12 | 2549192.25 | 2533835 | 2747805.5 | 9 | **1.121** |
| Q8VI75 | Ipo4 | Importin-4 (Mus musculus) | 160.6 | 2.59 | 119275 | 3 | 1062598.12 | 1184517.38 | 1084328.38 | 1228505.25 | 1016664.62 | 1039231.62 | 1352874.25 | 1205580.12 | 1147466 | 1005221.88 | 12 | **1.031** |
| Q8VI93 | Oas3 | 2'-5'-oligoadenylate synthase 3 (Mus musculus) | 373.21 | 8.61 | 126333 | 8 | 5226069 | 3488270.5 | 4718349 | 4817912.5 | 4028115.25 | 5125047 | 5754890 | 5774000 | 5653836.5 | 6051684.5 | 1 | **1.273** |
| Q8VID1 | Dhrs4 | Dehydrogenase/reductase SDR family member 4 (Rattus norvegicus) | 95.62 | 6.09 | 29822 | 2 | 2193177.75 | 1362641.75 | 1416142.5 | 997530.5 | 1090789.88 | 1900029.38 | 374223 | 1555451.5 | 1678074.38 | 1853487.5 | 9 | **1.043** |
| Q8VIJ5 | Oga | Protein O-GlcNAcase (Rattus norvegicus) | 50.23 | 1.42 | 102918 | 1 | 1121744.38 | 908673.94 | 1226213.5 | 854612.69 | 0 | 959102.12 | 1236122.88 | 837015.06 | 654050.75 | 555640 | 11 | **1.032** |
| Q8VIJ6 | Sfpq | Splicing factor, proline- and glutamine-rich (Mus musculus) | 974.08 | 25.89 | 75442 | 15 | 121556840 | 99574584 | 108281104 | 78828032 | 92610672 | 92952360 | 103712032 | 114061464 | 119275672 | 100336896 | 9 | **1.059** |
| Q8WTY4 | Ciapin1 | Anamorsin (Mus musculus) | 232.69 | 18.77 | 33429 | 4 | 5100436 | 2214696.25 | 5154759 | 1943155.25 | 2549446 | 4495976.5 | 4910793 | 3433371.75 | 4944742 | 5157158.5 | 8 | **1.353** |
| Q91V41 | Rab14 | Ras-related protein Rab-14 (Mus musculus) | 840.34 | 70.7 | 23897 | 15 | 25389106 | 33037090 | 36170056 | 42428020 | 37318284 | 37562812 | 38056812 | 38350776 | 38902276 | 39912660 | 5 | **1.106** |
| Q91V61 | Sfxn3 | Sideroflexin-3 (Mus musculus) | 206.51 | 18.38 | 35406 | 3 | 5391655.5 | 316514.5 | 1887570.38 | 429045 | 3688469.75 | 1920639.75 | 1905931.25 | 4285761 | 1969305.25 | 3530599.5 | 9 | **1.162** |
| Q91V83 | Tti1 | TELO2-interacting protein 1 homolog (Mus musculus) | 67.31 | 1.75 | 120814 | 1 | 173816.95 | 210100.27 | 218980.11 | 61703.09 | 0 | 170727 | 225361.78 | 0 | 79298.1 | 30015.87 | 11 | **0.760** |
| Q91V92 | Acly | ATP-citrate synthase (Mus musculus) | 1758.01 | 31.81 | 119728 | 26 | 66102884 | 61729232 | 61909492 | 60426196 | 55157080 | 64479100 | 68141792 | 67980272 | 72057368 | 69932824 | 1 | **1.122** |
| Q91VA6 | Poldip2 | Polymerase delta-interacting protein 2 (Mus musculus) | 50.84 | 3.26 | 41870 | 1 | 796925 | 1335283.75 | 1046863.12 | 768385.81 | 787704.88 | 839739.62 | 789195.81 | 897522.69 | 761707.19 | 902458.25 | 11 | **0.885** |
| Q91VC3 | Eif4a3 | Eukaryotic initiation factor 4A-III (Mus musculus) | 1218.98 | 53.28 | 46840 | 22 | 95079080 | 93340928 | 106687616 | 89142488 | 100367088 | 101720696 | 101530792 | 102183080 | 106712024 | 112666352 | 3 | **1.083** |
| Q91VC9 | Ghitm | Growth hormone-inducible transmembrane protein (Mus musculus) | 114.83 | 10.12 | 37275 | 2 | 3326453.75 | 1835393.75 | 2886252.75 | 1888453.62 | 1967052.5 | 2265989 | 2036501.5 | 2441200 | 2682873 | 2383951.5 | 10 | **0.992** |
| Q91VD9 | Ndufs1 | NADH-ubiquinone oxidoreductase 75 kDa subunit, mitochondrial (Mus musculus) | 640.63 | 23.11 | 79777 | 12 | 15498604 | 8834320 | 10945300 | 8898156 | 11779450 | 14614764 | 14704964 | 14966609 | 14451499 | 13652676 | 5 | **1.294** |
| Q91VE0 | Slc27a4 | Long-chain fatty acid transport protein 4 (Mus musculus) | 56.91 | 1.4 | 72319 | 1 | 1011314.62 | 697817.88 | 602900.44 | 632878.69 | 0 | 924769.44 | 737585.5 | 955541.62 | 790793.88 | 851603.31 | 5 | **1.447** |
| Q91VE6 | Nifk | MKI67 FHA domain-interacting nucleolar phosphoprotein (Mus musculus) | 345.08 | 24.61 | 36265 | 6 | 4578400.5 | 3645391.5 | 4830219 | 3293783.5 | 4775198 | 5990912 | 5456316 | 6518318.5 | 5843093 | 5979826.5 | 0 | **1.410** |
| Q91VH2 | Snx9 | Sorting nexin-9 (Mus musculus) | 150.98 | 7.39 | 66546 | 3 | 2955183 | 3112432.75 | 3134434.5 | 3898332 | 3415633.5 | 3232021.25 | 2516880.25 | 2968419.75 | 2928517.25 | 3029544 | 5 | **0.889** |
| Q91VI7 | Rnh1 | Ribonuclease inhibitor (Mus musculus) | 2196.7 | 85.96 | 49816 | 26 | 119746576 | 117752600 | 102600872 | 120257104 | 105134664 | 103621056 | 103520344 | 107605528 | 112603848 | 108473320 | 8 | **0.948** |
| Q91VJ1 | Aim2 | Interferon-inducible protein AIM2 (Mus musculus) | 178.32 | 10.45 | 40155 | 3 | 2460313.25 | 2477537.25 | 2700849.25 | 2837936 | 2593378.5 | 2330645.75 | 2380114.5 | 2511745 | 3082556.5 | 3033946.5 | 12 | **1.021** |
| Q91VM9 | Ppa2 | Inorganic pyrophosphatase 2, mitochondrial (Mus musculus) | 103.21 | 8.18 | 38115 | 2 | 1269759.25 | 2014260.88 | 1883951.5 | 2958451.25 | 3858061.5 | 1506152.75 | 1516472.88 | 2356870 | 1735694 | 2548033.5 | 9 | **0.806** |
| Q91VR2 | Atp5f1c | ATP synthase subunit gamma, mitochondrial (Mus musculus) | 540.7 | 38.26 | 32886 | 11 | 23091386 | 14560628 | 17546888 | 9691430 | 19066806 | 23271726 | 9080008 | 14128959 | 15023960 | 22415886 | 12 | **1.000** |

| | | | | | | | | | | | | | | | | | | |
|---|---|---|---|---|---|---|---|---|---|---|---|---|---|---|---|---|---|---|
| Q91VU0 | Fam3c | Protein FAM3C (Mus musculus) | 173.43 | 19.82 | 24753 | 3 | 1697554.5 | 1199581.75 | 237156.23 | 1109657.75 | 245629.16 | 554485.88 | 1442525 | 1079771 | 1290574.88 | 992619.19 | 11 | **1.194** |
| Q91VU7 | Pus7 | Pseudouridylate synthase 7 homolog (Mus musculus) | 109.16 | 4.39 | 74793 | 2 | 1299566.12 | 699016.06 | 1039924.38 | 1047376.19 | 0 | 1273369.5 | 1535923 | 1428311.38 | 1172052.62 | 1348892.88 | 2 | **1.654** |
| Q91VW3 | Sh3bgrl3 | SH3 domain-binding glutamic acid-rich-like protein 3 (Mus musculus) 3 | 154.55 | 66.67 | 10477 | 3 | 8468669 | 4232319.5 | 6368633.5 | 4317160.5 | 6273189 | 6585991.5 | 8088373.5 | 6542868 | 7218815 | 6903690.5 | 5 | **1.191** |
| Q91VY9 | Znf622 | Zinc finger protein 622 (Mus musculus) | 112.16 | 5.46 | 53451 | 2 | 0 | 0 | 901629.75 | 1606294.88 | 1401353.12 | 0 | 0 | 1472207 | 0 | 303161.19 | 15 | **0.454** |
| Q91VZ6 | Smap1 | Stromal membrane-associated protein 1 (Mus musculus) | 46.65 | 2.73 | 47660 | 1 | 1860404 | 0 | 0 | 0 | 959702.12 | 921402.44 | 0 | 893866.75 | 732242.94 | 787309.75 | 15 | **1.183** |
| Q91W50 | Csde1 | Cold shock domain-containing protein E1 (Mus musculus) | 930.01 | 24.81 | 88791 | 17 | 34736200 | 40607776 | 46295220 | 40909868 | 40708240 | 43204368 | 27672276 | 39020976 | 40934492 | 42625916 | 12 | **0.952** |
| Q91W89 | Man2c1 | Alpha-mannosidase 2C1 (Mus musculus) | 188.13 | 3.95 | 115688 | 3 | 1221183 | 900497.44 | 1436979.38 | 662884.19 | 702580.62 | 1484345 | 1441747.75 | 1385684.5 | 929320.75 | 1451959.5 | 3 | **1.359** |
| Q91W90 | Txndc5 | Thioredoxin domain-containing protein 5 (Mus musculus) | 410.23 | 21.1 | 46415 | 6 | 10548518 | 10296940 | 8359495.5 | 10873278 | 10139004 | 6018436 | 3967890.5 | 8738028 | 9050132 | 8859624 | 3 | **0.730** |
| Q91W97 | Hkdc1 | Putative hexokinase HKDC1 (Mus musculus) | 193.9 | 2.95 | 102259 | 4 | 19943890 | 16667797 | 18186238 | 15291586 | 15653522 | 21351074 | 19595232 | 19404072 | 18203994 | 19242358 | 4 | **1.141** |
| Q91WC0 | Setd3 | Histone-lysine N-methyltransferase setd3 (Mus musculus) | 93.37 | 6.23 | 67176 | 2 | 693645.25 | 1189710.75 | 903931.75 | 844749.44 | 851339.69 | 707939.25 | 686088.25 | 735739.62 | 859524.38 | 579581.88 | 5 | **0.796** |
| Q91WD5 | Ndufs2 | NADH dehydrogenase [ubiquinone] iron-sulfur protein 2, mitochondrial (Mus musculus) | 168.73 | 9.5 | 52626 | 3 | 5195535.5 | 4320866 | 2698895 | 3193987.25 | 3062251.75 | 3664020.5 | 3064297.25 | 3247268.5 | 3137073 | 2646469.5 | 10 | **0.853** |
| Q91WE2 | Fam192a | PSME3-interacting protein (Mus musculus) | 63.82 | 5.51 | 28702 | 2 | 1908011.88 | 6238930 | 8685856 | 5919485.5 | 6718169.5 | 7809006 | 6760993 | 6974059.5 | 7861759 | 7813272 | 5 | **1.263** |
| Q91WK2 | Eif3h | Eukaryotic translation initiation factor 3 subunit H (Mus musculus) | 628.8 | 44.32 | 39832 | 11 | 13769276 | 19469906 | 21746996 | 22984144 | 18997610 | 18327582 | 17503512 | 21177950 | 17305668 | 19312050 | 8 | **0.966** |
| Q91WM3 | Rrp9 | U3 small nucleolar RNA-interacting protein 2 (Mus musculus) | 67.34 | 3.79 | 52107 | 1 | 335028.25 | 404096.69 | 394961.72 | 84178.21 | 355119.97 | 315865.84 | 432581.38 | 597554.81 | 618796.69 | 836161.06 | 4 | **1.780** |
| Q91WQ3 | Yars | Tyrosine--tRNA ligase, cytoplasmic (Mus musculus) | 644.55 | 33.14 | 59105 | 14 | 13442360 | 17190332 | 21169514 | 23012626 | 16183973 | 22156760 | 21442102 | 20761050 | 25723376 | 21460294 | 5 | **1.226** |
| Q91WR3 | Ascc2 | Activating signal cointegrator 1 complex subunit 2 (Mus musculus) | 111.38 | 4.81 | 85653 | 2 | 349260.91 | 0 | 0 | 293371.38 | 420677.5 | 338660.34 | 179562.98 | 467457.56 | 588850.75 | 473736.19 | 6 | **1.926** |
| Q91XB0 | Trex1 | Three-prime repair exonuclease 1 (Mus musculus) | 120.05 | 9.55 | 33675 | 2 | 506477.34 | 700065.31 | 433952.25 | 0 | 0 | 271236.5 | 388889.44 | 351542.22 | 446924.75 | 289386.53 | 11 | **1.066** |
| Q91XC9 | Pex16 | Peroxisomal membrane protein PEX16 (Mus musculus) | 90.44 | 7.14 | 38677 | 2 | 0 | 91140.59 | 285370.56 | 0 | 0 | 424474.66 | 348586.31 | 217282.7 | 0 | 0 | 14 | **2.630** |
| Q91XU1 | Qki | Protein quaking (Rattus norvegicus) | 121.09 | 9.68 | 37643 | 2 | 2102168.5 | 2204797.75 | 1015729.31 | 1241310.62 | 0 | 938771.44 | 2063837.5 | 1252704.75 | 2091332 | 774846.19 | 11 | **1.085** |
| Q91XV3 | Basp1 | Brain acid soluble protein 1 (Mus musculus) | 618.72 | 61.95 | 22087 | 9 | 44443752 | 15059807 | 6688641 | 13973098 | 27488138 | 8433696 | 20314726 | 6717164 | 15440953 | 7591896.5 | 9 | **0.543** |
| Q91YE6 | Ipo9 | Importin-9 (Mus musculus) | 1234.54 | 26.03 | 116052 | 20 | 31753936 | 46133820 | 47003736 | 52860940 | 41405212 | 41837684 | 44428180 | 47588068 | 46056244 | 49699184 | 11 | **1.048** |
| Q91YH5 | Atl3 | Atlastin-3 (Mus musculus) | 596.67 | 26.06 | 60575 | 12 | 9094138 | 11317889 | 11119208 | 10439397 | 10343381 | 10943914 | 12645889 | 10520156 | 8068865 | 11307307 | 10 | **1.022** |
| Q91YI0 | Asl | Argininosuccinate lyase (Mus musculus) | 328.19 | 17.46 | 51739 | 6 | 6334618 | 5224060 | 5895187.5 | 7243024 | 7431294.5 | 7419717 | 6765132 | 7017562.5 | 7835508 | 6387089 | 7 | **1.103** |
| Q91YJ2 | Snx4 | Sorting nexin-4 (Mus musculus) | 51.28 | 2 | 51778 | 1 | 842574.81 | 1115483.38 | 1134318.12 | 1550117.38 | 1062628.75 | 854329.31 | 1010602.75 | 817900.56 | 1171626.62 | 885571.19 | 7 | **0.831** |
| Q91YK2 | Rrp1b | Ribosomal RNA processing protein 1 homolog B (Mus musculus) | 404.23 | 13.95 | 80582 | 7 | 2510345.25 | 5818303 | 6926537 | 4734004 | 4215074 | 4638100 | 4712596 | 5511526 | 5258183 | 4168400 | 11 | **1.003** |
| Q91YM4 | Tbrg4 | FAST kinase domain-containing protein 4 (Mus musculus) | 44.11 | 2.7 | 71513 | 1 | 0 | 378340.03 | 326721.56 | 0 | 0 | 0 | 342691.56 | 0 | 0 | 0 | 22 | **0.486** |
| Q91YN5 | Uap1 | UDP-N-acetylhexosamine pyrophosphorylase (Mus musculus) | 179.9 | 8.05 | 58609 | 3 | 3768332.75 | 3533894 | 2496724.25 | 3359900 | 3722725.25 | 3907250.25 | 3473397.5 | 4942580.5 | 4804025 | 4335315 | 3 | **1.271** |
| Q91YP3 | Dera | Deoxyribose-phosphate aldolase (Mus musculus) | 528.49 | 42.14 | 34975 | 9 | 12077179 | 8485389 | 10013590 | 6686048 | 6776831 | 11249890 | 10655416 | 11133658 | 10532910 | 11365185 | 5 | **1.247** |
| Q91YQ5 | Rpn1 | Dolichyl-diphosphooligosaccharide--protein glycosyltransferase subunit 1 (Mus musculus) | 1390.68 | 48.03 | 68528 | 22 | 112556736 | 109514984 | 109770704 | 96547128 | 103479608 | 124444312 | 112816272 | 112678120 | 116713496 | 115937768 | 0 | **1.095** |
| Q91YR1 | Twf1 | Twinfilin-1 (Mus musculus) | 223.68 | 11.71 | 40079 | 3 | 10042178 | 8993530 | 11084526 | 9775708 | 9535441 | 2634897 | 11071388 | 10624781 | 9847630 | 10582940 | 10 | **0.906** |
| Q91YR9 | Ptgr1 | Prostaglandin reductase 1 (Mus musculus) | 982.27 | 64.13 | 35560 | 17 | 25567216 | 31761794 | 41896740 | 33736740 | 32025610 | 37413388 | 29956606 | 34347416 | 31611162 | 34419872 | 11 | **1.017** |
| Q91YT0 | Ndufv1 | NADH dehydrogenase [ubiquinone] flavoprotein 1, mitochondrial (Mus musculus) | 270.75 | 17.67 | 50834 | 5 | 11072687 | 7579753.5 | 7237282.5 | 6245110.5 | 6444836 | 9738693 | 6201003.5 | 8046599 | 7113468 | 6920208 | 12 | **0.985** |
| Q91YT7 | Ythdf2 | YTH domain-containing family protein 2 (Mus musculus) | 116.5 | 4.32 | 62280 | 2 | 2618742 | 656221 | 1157064.75 | 0 | 395040.47 | 1702118.88 | 1730188.75 | 1341419.25 | 1412998 | 1653977 | 5 | **1.624** |
| Q91YU8 | Ppan | Suppressor of SWI4 1 homolog (Mus musculus) | 71.11 | 2.13 | 52756 | 1 | 1015332.62 | 606279.31 | 691218.62 | 538643.69 | 321091.66 | 525947.88 | 707077.5 | 608235.56 | 540793.12 | 599666.88 | 12 | **0.940** |
| Q91YW3 | Dnajc3 | DnaJ homolog subfamily C member 3 (Mus musculus) | 43.57 | 3.77 | 57464 | 1 | 890613.94 | 1446646.62 | 936579 | 1257989.25 | 1041574.5 | 1020748.12 | 1006332.81 | 1220545.75 | 1156484.12 | 933429.75 | 11 | **0.958** |
| Q91YX0 | Themis2 | Protein THEMIS2 (Mus musculus) | 156.61 | 5.88 | 74378 | 2 | 412218.16 | 813856.75 | 751413.62 | 936511.69 | 673608.12 | 671187.62 | 818649.12 | 731012.75 | 712013.38 | 398806.09 | 9 | **0.929** |
| Q91YY4 | Atpaf2 | ATP synthase mitochondrial F1 complex assembly factor 2 (Mus musculus) | 58.54 | 4.15 | 33289 | 1 | 0 | 0 | 0 | 0 | 0 | 242854.53 | 147891.41 | 193386.39 | 273354.47 | 209523.81 | 10 | **#DIV/0 !** |
| Q91Z53 | Grhpr | Glyoxylate reductase/hydroxypyruvate reductase (Mus musculus) | 175.75 | 12.5 | 35329 | 4 | 3864489.5 | 2306182 | 2548946.5 | 2807991.75 | 4689751.5 | 3210473.75 | 3298694 | 4393672 | 3626448.5 | 3558054 | 9 | **1.115** |
| Q91ZA3 | Pcca | Propionyl-CoA carboxylase alpha chain, mitochondrial (Mus musculus) | 169.78 | 6.08 | 79922 | 3 | 42718.17 | 743186.31 | 1424955 | 1473868 | 1726385.12 | 180784.47 | 1481351 | 749722.5 | 607439.25 | 623364.31 | 9 | **0.673** |
| Q91ZJ5 | Ugp2 | UTP--glucose-1-phosphate uridylyltransferase (Mus musculus) | 354.77 | 16.34 | 56979 | 7 | 7923870 | 8634859 | 9967498 | 5521948 | 7767739.5 | 10288406 | 9094718 | 8586866 | 9586116 | 9625652 | 5 | **1.185** |
| Q91ZQ0 | Vmp1 | Vacuole membrane protein 1 (Rattus norvegicus) | 85.59 | 3.94 | 45901 | 2 | 0 | 651305.88 | 790518 | 0 | 208994.66 | 223796.58 | 689533.38 | 1126329.12 | 801916.69 | 880544.12 | 4 | **2.255** |
| Q91ZR2 | Snx18 | Sorting nexin-18 (Mus musculus) | 392.13 | 16.12 | 67904 | 8 | 10536917 | 9662248 | 10715133 | 10150066 | 10889144 | 10820795 | 6552042 | 12471790 | 11351963 | 12074094 | 6 | **1.025** |
| Q91ZR3 | Crebzf | CREB/ATF bZIP transcription factor (Mus musculus) | 71.28 | 3.91 | 37825 | 2 | 102272.81 | 14067474 | 12983682 | 9648784 | 10741500 | 99641.36 | 172422.56 | 12110022 | 9706302 | 12483100 | 9 | **0.727** |
| Q91ZV7 | Plxdc1 | Plexin domain-containing protein 1 (Mus musculus) | 56.74 | 3 | 55635 | 1 | 100492.33 | 1431056.88 | 133970.7 | 1215483 | 1599707.38 | 169122.83 | 1142434.5 | 1341472.38 | 1334726.75 | 996699.5 | 12 | **1.112** |
| Q91ZW3 | Smarca5 | SWI/SNF-related matrix-associated actin-dependent regulator of chromatin subfamily A me | 564.65 | 12.27 | 121627 | 14 | 13989636 | 17108058 | 16110962 | 20456326 | 14797316 | 15332912 | 14650205 | 14491844 | 16719324 | 16449214 | 10 | **0.942** |
| Q91ZX7 | Lrp1 | Prolow-density lipoprotein receptor-related protein 1 (Mus musculus) | 3737.55 | 20.04 | 504742 | 66 | 130040296 | 125010152 | 132964608 | 130908488 | 116438136 | 133733800 | 137926416 | 132842624 | 137192848 | 134313712 | 1 | **1.064** |
| Q920A5 | Scpep1 | Retinoid-inducible serine carboxypeptidase (Mus musculus) | 246.04 | 11.06 | 50965 | 4 | 8579905 | 10627568 | 10458332 | 8193697.5 | 11742171 | 8977426 | 8838706 | 8612921 | 12099648 | 11701582 | 10 | **1.013** |
| Q920A7 | Afg3l1 | AFG3-like protein 1 (Mus musculus) | 49.34 | 1.77 | 87047 | 1 | 496944.03 | 374461.56 | 499093.81 | 95065.09 | 322009.91 | 387149.56 | 81928.12 | 225601.81 | 365786.94 | 0 | 6 | **0.593** |
| Q920B9 | Supt16h | FACT complex subunit SPT16 (Mus musculus) | 1009.86 | 22.83 | 119825 | 19 | 30459230 | 32072288 | 29354540 | 30381540 | 19437382 | 29130720 | 31708668 | 31521220 | 30784792 | 32108208 | 7 | **1.096** |
| Q920E5 | Fdps | Farnesyl pyrophosphate synthase (Mus musculus) | 1033.42 | 49.01 | 40582 | 15 | 50070596 | 60101456 | 62609900 | 59262244 | 53857372 | 63797156 | 61540656 | 60042280 | 60700628 | 56033460 | 7 | **1.057** |
| Q920L1 | Fads1 | Fatty acid desaturase 1 (Mus musculus) | 38.85 | 4.03 | 52323 | 1 | 61017.19 | 0 | 119454.45 | 0 | 48693.61 | 173194.81 | 144364.78 | 0 | 0 | 0 | 16 | **1.386** |
| Q920Q4 | Vps16 | Vacuolar protein sorting-associated protein 16 homolog (Mus musculus) | 242.48 | 9.06 | 94928 | 4 | 2462781.75 | 2523617 | 838509 | 1953288.62 | 1157707.75 | 2571672.25 | 2572334.75 | 2173334.25 | 2225768 | 2114776.5 | 6 | **1.305** |
| Q920Q8 | Ivns1abp | Influenza virus NS1A-binding protein homolog (Mus musculus) p | 43.44 | 1.56 | 71581 | 1 | 1595140.88 | 3579185 | 2920918.75 | 819508.94 | 3722077 | 860037.69 | 2319790 | 1653701.38 | 2685090.75 | 616426.06 | 7 | **0.644** |
| Q921E2 | Rab31 | Ras-related protein Rab-31 (Mus musculus) | 202.71 | 18.04 | 21331 | 3 | 4789092 | 1845793.5 | 1980994.25 | 1948055.25 | 2806269.25 | 4443718 | 4456726 | 3827718 | 3934626 | 4028955.5 | 5 | **1.548** |
| Q921F2 | Tardbp | TAR DNA-binding protein 43 (Mus musculus) | 587.93 | 35.27 | 44548 | 9 | 24916448 | 17376632 | 14528356 | 20796632 | 16791546 | 18514068 | 17617718 | 15710576 | 16694100 | 16989732 | 10 | **0.906** |
| Q921F4 | Hnrnpll | Heterogeneous nuclear ribonucleoprotein L-like (Mus musculus) | 392.24 | 15.74 | 64125 | 7 | 10017222 | 10859545 | 10136510 | 6618693 | 5427909 | 11036881 | 7123040 | 7723231 | 8197463.5 | 8166615.5 | 12 | **0.981** |
| Q921G8 | Tubgcp2 | Gamma-tubulin complex component 2 (Mus musculus) | 83.98 | 2.32 | 103223 | 2 | 0 | 1135775.5 | 815638.38 | 278346.09 | 338503.22 | 682332 | 459560.62 | 276032.5 | 986367.62 | 318334.91 | 12 | **1.060** |
| Q921H8 | Acaa1a | 3-ketoacyl-CoA thiolase A, peroxisomal (Mus musculus) | 820.72 | 55.19 | 43953 | 16 | 23762716 | 23946500 | 26136500 | 20264308 | 25463104 | 25485104 | 25764254 | 24994240 | 19731516 | 26761700 | 9 | **1.026** |
| Q921I2 | Klhdc4 | Kelch domain-containing protein 4 (Mus musculus) | 42.41 | 2.57 | 64860 | 1 | 2397834.25 | 1523641 | 1721986.75 | 1597748.12 | 1185597.25 | 1525453.75 | 1595488.38 | 1473083.75 | 1206041.25 | 1520190.38 | 7 | **0.869** |
| Q921I9 | Exosc4 | Exosome complex component RRP41 (Mus musculus) | 45.94 | 5.71 | 26250 | 1 | 916382.94 | 625010.19 | 695996.75 | 716128.06 | 937051.88 | 1034227.38 | 544543.44 | 765012.88 | 703745.88 | 659302.25 | 11 | **0.953** |
| Q921J2 | Rheb | GTP-binding protein Rheb (Mus musculus) | 301.72 | 32.61 | 20451 | 7 | 6463154 | 10167071 | 11117605 | 9687883 | 9383678 | 6050436.5 | 7504779 | 10869039 | 10254580 | 10438903 | 12 | **0.964** |
| Q921M3 | Sf3b3 | Splicing factor 3B subunit 3 (Mus musculus) | 1661.11 | 31.47 | 135550 | 28 | 43763152 | 47330820 | 43780304 | 49160280 | 42188100 | 42662312 | 41352568 | 40043988 | 39066748 | 44873520 | 4 | **0.919** |
| Q921M7 | Fam49b | Protein FAM49B (Mus musculus) | 947.08 | 59.88 | 36776 | 16 | 37333528 | 28721712 | 36465880 | 34161856 | 36094096 | 38169368 | 37350096 | 43558440 | 43241944 | 40122408 | 0 | **1.172** |
| Q921N6 | Ddx27 | Probable ATP-dependent RNA helicase DDX27 (Mus musculus) | 38.99 | 1.71 | 85939 | 1 | 832621.5 | 626519.81 | 632977.38 | 316124.47 | 893393.38 | 769743.44 | 874743.5 | 707077.88 | 750699.44 | 571667.44 | 11 | **1.113** |
| Q921T2 | Tor1aip1 | Torsin-1A-interacting protein 1 (Mus musculus) 1 | 501.75 | 21.51 | 66781 | 9 | 11035773 | 13540498 | 12712089 | 9630994 | 17225940 | 15483193 | 14692032 | 16979852 | 17616304 | 14809799 | 4 | **1.241** |
| Q921W4 | Cryzl1 | Quinone oxidoreductase-like protein 1 (Mus musculus) | 44.35 | 5.75 | 38725 | 1 | 0 | 2656077.5 | 1862301.88 | 2309556 | 1730424.5 | 1546915.25 | 0 | 1771330.88 | 1798731.75 | 1876973.38 | 9 | **0.817** |
| Q921Z5 | Tnfaip8 | Tumor necrosis factor alpha-induced protein 8 (Mus musculus) | 104.89 | 10.1 | 22960 | 1 | 0 | 561882.12 | 514049.44 | 432214.34 | 458510.72 | 0 | 489235.91 | 489032.88 | 36063.66 | 495720.19 | 11 | **0.768** |
| Q922B2 | Dars | Aspartate--tRNA ligase, cytoplasmic (Mus musculus) | 1374.44 | 54.49 | 57147 | 25 | 95989856 | 98736616 | 105459328 | 99411984 | 87583240 | 104365976 | 81922592 | 116258136 | 112696680 | 119869288 | 6 | **1.098** |
| Q922D4 | Ppp6r3 | Serine/threonine-protein phosphatase 6 regulatory subunit 3 (Mus musculus) | 427.05 | 10.9 | 94653 | 9 | 1818963.75 | 5034187.5 | 5369386 | 4936475 | 5281975.5 | 6031754.5 | 4403599.5 | 4794173.5 | 4324082.5 | 5047268 | 11 | **1.096** |
| Q922D8 | Mthfd1 | C-1-tetrahydrofolate synthase, cytoplasmic (Mus musculus) | 1014.6 | 22.89 | 101200 | 20 | 46172084 | 35085536 | 39193896 | 40507644 | 42259020 | 43544904 | 40869004 | 42974988 | 43105760 | 44542160 | 6 | **1.058** |
| Q922F4 | Tubb6 | Tubulin beta-6 chain (Mus musculus) | 1547.71 | 63.31 | 50090 | 25 | 387108192 | 377633248 | 301420896 | 350248992 | 305979168 | 358752672 | 339173952 | 384551968 | 395131424 | 383706304 | 7 | **1.081** |
| Q922H2 | Pdk3 | [Pyruvate dehydrogenase (acetyl-transferring)] kinase isozyme 3, mitochondrial (Mus musc | 109.96 | 6.27 | 47923 | 2 | 5198534 | 1449384.38 | 3380638 | 5289052 | 1433852.88 | 2478435 | 1581401.88 | 1871935.12 | 3082007.75 | 1458559.5 | 10 | **0.625** |
| Q922J3 | Clip1 | CAP-Gly domain-containing linker protein 1 (Mus musculus) | 126.11 | 2.08 | 155814 | 2 | 0 | 265792.19 | 317426.78 | 338596.62 | 290849.22 | 0 | 71826.45 | 202149.56 | 205709.31 | 37331.61 | 5 | **0.426** |
| Q922K7 | Nop2 | Probable 28S rRNA (cytosine-C(5))-methyltransferase (Mus musculus) | 528.07 | 18.92 | 86752 | 10 | 26321102 | 21347146 | 22711492 | 22774814 | 20759234 | 19056568 | 23149634 | 20988040 | 23151788 | 23102476 | 12 | **0.961** |
| Q922L6 | Nelfcd | Negative elongation factor D (Mus musculus) | 138.03 | 7.11 | 66276 | 3 | 1698184.38 | 1933276 | 1449279.25 | 1670418.5 | 1520831.25 | 1364611 | 1639913.75 | 1732784.88 | 717573.5 | 642666.81 | 6 | **0.737** |
| Q922Q1 | Marc2 | Mitochondrial amidoxime reducing component 2 (Mus musculus) | 286.18 | 18.93 | 38194 | 6 | 14643590 | 7076998 | 9732721 | 5550139.5 | 8035766.5 | 13654203 | 12511439 | 13277167 | 10360692 | 9127169 | 6 | **1.308** |
| Q922Q4 | Pycr2 | Pyrroline-5-carboxylate reductase 2 (Mus musculus) | 683.57 | 44.06 | 33659 | 9 | 35081680 | 20344404 | 21345752 | 19843330 | 18871096 | 24166708 | 22413550 | 20791458 | 19595736 | 21011956 | 10 | **0.935** |
| Q922Q8 | Lrrc59 | Leucine-rich repeat-containing protein 59 (Mus musculus) | 519.29 | 41.69 | 34877 | 9 | 64515536 | 29352848 | 32134804 | 32091602 | 38656516 | 45888816 | 55099792 | 50015524 | 51242020 | 52708852 | 5 | **1.296** |
| Q922R5 | Ppp4r3b | Serine/threonine-protein phosphatase 4 regulatory subunit 3B (Mus musculus) | 115.77 | 4.15 | 93934 | 2 | 0 | 272642.56 | 470050.03 | 253520.39 | 379787.12 | 286676.97 | 244585.25 | 493157.19 | 585159.19 | 511456.47 | 6 | **1.541** |
| Q922R8 | Pdia6 | Protein disulfide-isomerase A6 (Mus musculus) | 1137.22 | 45.23 | 48100 | 16 | 82887664 | 115675984 | 104594272 | 121259968 | 92088016 | 105136832 | 109239960 | 112835568 | 106323808 | 106958800 | 10 | **1.046** |
| Q922S8 | Kif2c | Kinesin-like protein KIF2C (Mus musculus) | 85.03 | 3.19 | 81085 | 2 | 732349.44 | 601551.06 | 572556.12 | 50871.07 | 0 | 869862.19 | 828225.06 | 578946.88 | 491542.19 | 0 | 10 | **1.414** |
| Q922V4 | Plrg1 | Pleiotropic regulator 1 (Mus musculus) | 304.49 | 17.74 | 56938 | 6 | 5957888.5 | 6635818.5 | 6244546 | 7220558.5 | 6312278.5 | 7668122.5 | 4824683 | 6324757 | 4640075 | 5942253.5 | 8 | **0.908** |
| Q923D2 | Blvrb | Flavin reductase (NADPH) (Mus musculus) | 401.47 | 51.94 | 22197 | 5 | 6135705 | 7885618 | 4841874.5 | 15719861 | 6455308.5 | 6094378.5 | 1637195 | 5982957.5 | 5104322 | 4279561.5 | 3 | **0.563** |
| Q923G2 | Polr2h | DNA-directed RNA polymerases I, II, and III subunit RPABC3 (Mus musculus) | 239.08 | 21.33 | 17143 | 3 | 2270610 | 2034414.5 | 2984128.5 | 1846876.25 | 1013318.88 | 5593045.5 | 944906.75 | 1955546.75 | 2106741.5 | 4728976.5 | 10 | **1.510** |
| Q923V8 | Selenof | Selenoprotein F (Rattus norvegicus) | 97.83 | 25.31 | 17822 | 2 | 527550.25 | 276335.22 | 1403206.25 | 0 | 372335.22 | 500485.03 | 162325.81 | 536561.5 | 734179.25 | 1192665.75 | 9 | **1.212** |
| Q924C1 | Xpo5 | Exportin-5 (Mus musculus) | 493.99 | 10.55 | 136973 | 9 | 6385318.5 | 9392989 | 7967240 | 8494590 | 6247779.5 | 6181309.5 | 7938616 | 7945965.5 | 6718663 | 6814346 | 8 | **0.925** |
| Q924T2 | Mrps2 | 28S ribosomal protein S2, mitochondrial (Mus musculus) | 172.71 | 15.81 | 32313 | 3 | 1495170.25 | 2867341.5 | 2647197 | 1726645.62 | 1678089.5 | 2337937 | 2365184 | 2085669.25 | 1863333.5 | 1673218.62 | 12 | **0.991** |
| Q924Z4 | Cers2 | Ceramide synthase 2 (Mus musculus) | 44.12 | 5.26 | 45024 | 1 | 0 | 0 | 0 | 0 | 0 | 0 | 0 | 475841.62 | 561412.5 | 461003.47 | 20 | **#DIV/0 !** |

| | | | | | | | | | | | | | | | | | | |
|---|---|---|---|---|---|---|---|---|---|---|---|---|---|---|---|---|---|---|
| Q93092 | Taldo1 | Transaldolase (Mus musculus) | 1713.72 | 64.99 | 37387 | 29 | 500573120 | 592965504 | 667387904 | 553527680 | 608817920 | 644327616 | 647976000 | 634404928 | 618319424 | 654216128 | 5 | **1.094** |
| Q99020 | Hnrnpab | Heterogeneous nuclear ribonucleoprotein A/B (Mus musculus) | 374.87 | 23.16 | 30831 | 7 | 26113080 | 19523394 | 19933084 | 22044102 | 18792290 | 15431141 | 18588040 | 18195396 | 18404018 | 18860910 | 1 | **0.841** |
| Q99104 | Myo5a | Unconventional myosin-Va (Mus musculus) | 114.31 | 1.73 | 215538 | 2 | 0 | 366239 | 748745 | 306011.03 | 333742.84 | 229937.2 | 225389.94 | 835274.12 | 700067.75 | 703062.75 | 10 | **1.535** |
| Q99J09 | Wdr77 | Methylosome protein 50 (Mus musculus) | 100.68 | 4.97 | 36943 | 2 | 1972618.12 | 2881536.25 | 1901881 | 2640365.75 | 2106087.25 | 666330.44 | 1833114.88 | 1600296.88 | 1617394.88 | 1499994.38 | 0 | **0.627** |
| Q99J23 | Ghdc | GH3 domain-containing protein (Mus musculus) | 77.02 | 2.44 | 58508 | 1 | 245407.64 | 137524.8 | 134712.8 | 83156.61 | 277464.94 | 317808.38 | 183474.61 | 302513.78 | 260457.66 | 223810.52 | 5 | **1.467** |
| Q99J36 | Thumpd1 | THUMP domain-containing protein 1 (Mus musculus) | 50.3 | 3.14 | 38885 | 1 | 663170.38 | 513415.53 | 505182.22 | 566071.12 | 0 | 581702.25 | 1096361.38 | 589097.69 | 1013554.38 | 747135.69 | 2 | **1.792** |
| Q99J62 | Rfc4 | Replication factor C subunit 4 (Mus musculus) | 150.55 | 7.69 | 39867 | 3 | 3151412.25 | 2919733.5 | 2216625.5 | 1583544.88 | 3498956.5 | 2983163.5 | 2718234 | 3808770.5 | 3949389 | 4728781 | 5 | **1.360** |
| Q99J77 | Nans | Sialic acid synthase (Mus musculus) | 379.33 | 28.13 | 40024 | 7 | 9023272 | 11133002 | 10535539 | 10560016 | 10094408 | 9793452 | 9484223 | 10630360 | 10325880 | 9128745 | 9 | **0.961** |
| Q99J93 | Ifitm2 | Interferon-induced transmembrane protein 2 (Mus musculus) | 151.94 | 16.67 | 15743 | 2 | 31892844 | 13980997 | 13986901 | 14671817 | 12317865 | 24710450 | 31596452 | 26514892 | 26959264 | 23615622 | 5 | **1.536** |
| Q99JB2 | Stoml2 | Stomatin-like protein 2, mitochondrial (Mus musculus) | 382.23 | 25.21 | 38385 | 7 | 6292050.5 | 7400238 | 4854161 | 7409037 | 3781724.5 | 6604077 | 6084104 | 6776460 | 7444721.5 | 6528199 | 9 | **1.124** |
| Q99JF8 | Psip1 | PC4 and SFRS1-interacting protein (Mus musculus) | 255.63 | 10.98 | 59697 | 5 | 7646717 | 5772780 | 7304590 | 5818042.5 | 6899212.5 | 5582827.5 | 1813819.12 | 5725922 | 5381208 | 4823107.5 | 0 | **0.698** |
| Q99JI4 | Psmd6 | 26S proteasome non-ATPase regulatory subunit 6 (Mus musculus) | 1254.74 | 63.24 | 45536 | 21 | 38342888 | 43963096 | 48554544 | 39110708 | 46170316 | 48837268 | 46975232 | 51426416 | 50830024 | 51472044 | 1 | **1.155** |
| Q99JI6 | Rap1b | Ras-related protein Rap-1b (Mus musculus) | 875.93 | 72.28 | 20825 | 13 | 80741872 | 84125064 | 87651616 | 86329376 | 80352624 | 84162272 | 51481056 | 71140480 | 71226232 | 72309560 | 3 | **0.836** |
| Q99JR1 | Sfxn1 | Sideroflexin-1 (Mus musculus) | 322.58 | 26.4 | 35649 | 5 | 13584478 | 7246131 | 7392594 | 7485075 | 10438745 | 7148368 | 8729632 | 9207212 | 7331724.5 | 9295561 | 10 | **0.904** |
| Q99JW2 | Acy1 | Aminoacylase-1 (Mus musculus) | 158.84 | 7.6 | 45781 | 3 | 15082656 | 7949979 | 8064280.5 | 11041590 | 6882507 | 4952781 | 2430309.5 | 6053959 | 7640625 | 6164442.5 | 1 | **0.556** |
| Q99JX3 | Gorasp2 | Golgi reassembly-stacking protein 2 (Mus musculus) | 253.09 | 15.52 | 47038 | 4 | 2849209.5 | 1861137.25 | 2590774.5 | 1218820 | 3315269.5 | 2751430 | 2857840.5 | 3384441.5 | 3639539.5 | 2986121.5 | 4 | **1.320** |
| Q99JX4 | Eif3m | Eukaryotic translation initiation factor 3 subunit M (Mus musculus) | 657.64 | 39.04 | 42517 | 10 | 7911695.5 | 13468985 | 16424337 | 16333527 | 14972003 | 14297950 | 9088261 | 15724661 | 14581342 | 16369399 | 12 | **1.014** |
| Q99JX7 | Nxf1 | Nuclear RNA export factor 1 (Mus musculus) | 200.3 | 7.77 | 70300 | 3 | 1438941 | 1662686.5 | 2162989.5 | 1932070.88 | 1735853.5 | 2440552.75 | 2084434 | 2315568.5 | 2781399.75 | 2968249 | 1 | **1.409** |
| Q99JY0 | Hadhb | Trifunctional enzyme subunit beta, mitochondrial (Mus musculus) | 410.3 | 19.16 | 51386 | 8 | 12468172 | 10840924 | 10735863 | 10890634 | 9311724 | 9886077 | 11265536 | 11009183 | 12402923 | 11215583 | 8 | **1.028** |
| Q99K01 | Pdxdc1 | Pyridoxal-dependent decarboxylase domain-containing protein 1 (Mus musculus) | 215.07 | 8.13 | 87335 | 4 | 1519587.12 | 1555311.38 | 1632770.38 | 1168945.75 | 1098230 | 1362157.12 | 1683460 | 1572674.5 | 225809.02 | 1694271.62 | 9 | **0.937** |
| Q99K23 | Ufsp2 | Ufm1-specific protease 2 (Mus musculus) | 84.19 | 6.07 | 52515 | 2 | 2097885.5 | 2016418.75 | 2690214.5 | 0 | 2192782 | 2453210 | 2119486 | 2472533 | 2412042 | 2343131.5 | 6 | **1.312** |
| Q99K28 | Arfgap2 | ADP-ribosylation factor GTPase-activating protein 2 (Mus musculus) | 75.77 | 3.85 | 56598 | 2 | 1015494.5 | 2274403 | 2384860.5 | 1409649.62 | 1728236.12 | 1581884 | 1385655.38 | 2273102 | 2304055 | 1927844.25 | 12 | **1.075** |
| Q99K51 | Pls3 | Plastin-3 (Mus musculus) | 1429.48 | 47.3 | 70742 | 23 | 154466176 | 311149344 | 352296672 | 334656512 | 323675776 | 383940320 | 312790112 | 348469120 | 331643776 | 347451008 | 7 | **1.168** |
| Q99K70 | Rragc | Ras-related GTP-binding protein C (Mus musculus) | 363.5 | 26.38 | 44121 | 7 | 7468410.5 | 12231275 | 13248762 | 11455862 | 13530086 | 11931488 | 10713786 | 13047508 | 13905386 | 13271022 | 10 | **1.085** |
| Q99K85 | Psat1 | Phosphoserine aminotransferase (Mus musculus) | 918.24 | 41.89 | 40473 | 15 | 71773904 | 87686720 | 83914104 | 80673952 | 80804456 | 98449792 | 79771128 | 88022664 | 83169856 | 85515656 | 7 | **1.074** |
| Q99KC8 | Vwa5a | von Willebrand factor A domain-containing protein 5A (Mus musculus) | 1167.9 | 33.17 | 87143 | 19 | 68334872 | 55723684 | 55021340 | 58981864 | 56160852 | 61478460 | 64758292 | 56972472 | 57180592 | 59550780 | 7 | **1.019** |
| Q99KE1 | Me2 | NAD-dependent malic enzyme, mitochondrial (Mus musculus) | 554.77 | 24.96 | 65799 | 9 | 19970452 | 12480658 | 8809399 | 8894317 | 8588038 | 14788060 | 13180080 | 10958473 | 10923330 | 8089651 | 11 | **0.986** |
| Q99KH8 | Stk24 | Serine/threonine-protein kinase 24 (Mus musculus) | 321.36 | 19.03 | 47954 | 5 | 4355188.5 | 3329060 | 4403477.5 | 2781065.75 | 4644023.5 | 4968207.5 | 4992944 | 4002356.25 | 4821604 | 4470403 | 4 | **1.192** |
| Q99KI0 | Aco2 | Aconitate hydratase, mitochondrial (Mus musculus) | 1608.92 | 41.54 | 85464 | 27 | 110291520 | 93250448 | 99393880 | 86583552 | 89815616 | 120311656 | 107436976 | 111739728 | 110929696 | 102643288 | 2 | **1.154** |
| Q99KK2 | Cmas | N-acylneuraminate cytidylyltransferase (Mus musculus) | 401.15 | 19.68 | 48058 | 8 | 10323455 | 11775142 | 10251500 | 11759364 | 8481665 | 13510601 | 13728653 | 13276094 | 12482178 | 14187706 | 0 | **1.278** |
| Q99KK7 | Dpp3 | Dipeptidyl peptidase 3 (Mus musculus) | 563.51 | 24.53 | 82898 | 11 | 14155748 | 50657188 | 25610196 | 12753589 | 9065198 | 16842952 | 14869704 | 17010230 | 15674134 | 14897252 | 10 | **0.706** |
| Q99KK9 | Hars2 | Probable histidine--tRNA ligase, mitochondrial (Mus musculus) | 195.19 | 9.9 | 56985 | 5 | 12360387 | 7509973.5 | 8097056.5 | 8203669 | 6879306 | 11590608 | 12823273 | 11391848 | 13173172 | 13853412 | 2 | **1.460** |
| Q99KN9 | Clint1 | Clathrin interactor 1 (Mus musculus) | 318.61 | 9.03 | 68513 | 6 | 21226396 | 16853488 | 16178878 | 13677315 | 15072354 | 16194932 | 17410420 | 14717955 | 17079648 | 12572914 | 12 | **0.939** |
| Q99KP3 | Cryl1 | Lambda-crystallin homolog (Mus musculus) | 230.93 | 18.81 | 35209 | 4 | 4369064.5 | 4187513.5 | 2951753 | 4985573.5 | 2535859 | 4782041 | 5128175.5 | 2777332 | 5363062 | 5795616 | 5 | **1.253** |
| Q99KP6 | Prpf19 | Pre-mRNA-processing factor 19 (Mus musculus) | 504.47 | 30.16 | 55239 | 9 | 27867270 | 28233722 | 26815484 | 25021124 | 18479492 | 29411540 | 17830398 | 26402400 | 25520312 | 25687762 | 11 | **0.988** |
| Q99KQ4 | Nampt | Nicotinamide phosphoribosyltransferase (Mus musculus) | 647.48 | 24.64 | 55447 | 10 | 19380394 | 16572482 | 16972854 | 12765550 | 8638193 | 17309354 | 17924580 | 16986630 | 15657058 | 15534760 | 9 | **1.122** |
| Q99KR3 | Lactb2 | Endoribonuclease LACTB2 (Mus musculus) | 145.32 | 10.42 | 32754 | 3 | 1869130.5 | 3085324 | 2488319.75 | 1460977.88 | 2522661.25 | 2074095 | 2605666.5 | 2685712.5 | 2421179 | 2697988 | 9 | **1.093** |
| Q99L04 | Dhrs1 | Dehydrogenase/reductase SDR family member 1 (Mus musculus) | 262.12 | 25.88 | 34005 | 5 | 1476104.38 | 4548469 | 37850868 | 4725937.5 | 38727380 | 3928930.25 | 2653953.5 | 2623690 | 4297967.5 | 4123089 | 5 | **0.202** |
| Q99L13 | Hibadh | 3-hydroxyisobutyrate dehydrogenase, mitochondrial (Mus musculus) | 116.51 | 9.55 | 35440 | 2 | 1230565.12 | 669356.44 | 637227.56 | 904383 | 983003.06 | 831445.06 | 954095.06 | 912963.94 | 588197.5 | 757736 | 10 | **0.914** |
| Q99L27 | Gmpr2 | GMP reductase 2 (Mus musculus) | 40.15 | 3.45 | 38019 | 1 | 2666275 | 1763533.62 | 1704204.75 | 1921515.38 | 2239456 | 1237139.62 | 1218128 | 2375414.75 | 1935360.62 | 2468857.75 | 11 | **0.897** |
| Q99L45 | Eif2s2 | Eukaryotic translation initiation factor 2 subunit 2 (Mus musculus) | 466.75 | 33.23 | 38092 | 9 | 8139071.5 | 8403338 | 8854106 | 7312299 | 8852806 | 13300126 | 10767633 | 10309363 | 10831080 | 7891886.5 | 4 | **1.278** |
| Q99L47 | St13 | Hsc70-interacting protein (Mus musculus) | 567.16 | 22.1 | 41656 | 10 | 56510288 | 36266820 | 33663104 | 35957544 | 38869796 | 46171108 | 41750184 | 41689736 | 43567376 | 41874580 | 5 | **1.068** |
| Q99L48 | Nmd3 | 60S ribosomal export protein NMD3 (Mus musculus) | 112.17 | 4.17 | 57611 | 2 | 2659139.75 | 1093734.88 | 2826293.25 | 1436413.25 | 532098.75 | 1216058.12 | 2822844.5 | 2786781 | 2259127.5 | 2747833.75 | 8 | **1.384** |
| Q99LC3 | Ndufa10 | NADH dehydrogenase [ubiquinone] 1 alpha subcomplex subunit 10, mitochondrial (Mus m | 209.62 | 12.68 | 40603 | 3 | 4567147 | 4061940.75 | 3316403.25 | 4897834 | 4107749 | 3547814.25 | 2775376.75 | 3516266.25 | 3785550.5 | 2209306 | 3 | **0.756** |
| Q99LC5 | Etfa | Electron transfer flavoprotein subunit alpha, mitochondrial (Mus musculus) | 1059.13 | 65.17 | 35009 | 16 | 57536996 | 48581608 | 58083480 | 47884008 | 57859060 | 51506644 | 50680144 | 54592008 | 56538500 | 57530704 | 10 | **1.003** |
| Q99LC8 | Eif2b1 | Translation initiation factor eIF-2B subunit alpha (Mus musculus) | 252.36 | 21.31 | 33816 | 4 | 1733949.5 | 2641224.75 | 2479005.5 | 1448936.12 | 980278.94 | 1703719.25 | 2115421.25 | 2072779.5 | 1679226.5 | 1769995.62 | 12 | **1.006** |
| Q99LD9 | Eif2b2 | Translation initiation factor eIF-2B subunit beta (Mus musculus) | 120.68 | 7.41 | 38898 | 3 | 4150014.5 | 1610957.75 | 1417733.12 | 1096538 | 1253185 | 1461883.5 | 3548543 | 1408725.5 | 2828896.5 | 1414309.38 | 10 | **1.119** |
| Q99LE6 | Abcf2 | ATP-binding cassette sub-family F member 2 (Mus musculus) | 118.64 | 6.53 | 71782 | 3 | 649713.19 | 2210849 | 1226692.62 | 1933380.38 | 2005147.12 | 1841826.5 | 1932190 | 1699219.38 | 1818558.5 | 1689536 | 10 | **1.119** |
| Q99LG2 | Tnpo2 | Transportin-2 (Mus musculus) | 515.72 | 11.72 | 100456 | 10 | 24194732 | 19592404 | 19635862 | 18182618 | 18435114 | 19571260 | 22681880 | 22571122 | 20975380 | 23509530 | 7 | **1.093** |
| Q99LH1 | Gnl2 | Nucleolar GTP-binding protein 2 (Mus musculus) | 49.95 | 1.79 | 83345 | 1 | 0 | 617523.88 | 456711.81 | 0 | 639221.06 | 455188.53 | 0 | 0 | 628492.19 | 0 | 15 | **0.632** |
| Q99LI2 | Clcc1 | Chloride channel CLIC-like protein 1 (Mus musculus) | 260.6 | 12.06 | 60621 | 4 | 562881.25 | 1760468.25 | 2205356.5 | 1614650.5 | 1352641.75 | 2327058.75 | 1874118.25 | 2136777.5 | 1889865.75 | 2438051.5 | 3 | **1.423** |
| Q99LI7 | Cstf3 | Cleavage stimulation factor subunit 3 (Mus musculus) | 313.2 | 13.39 | 82877 | 8 | 3458558 | 4241310.5 | 4516377 | 4306962 | 4291899.5 | 4958071 | 1527708.25 | 3104069 | 4825973 | 4452137 | 11 | **0.906** |
| Q99LJ0 | Cttnbp2nl | CTTNBP2 N-terminal-like protein (Mus musculus) nl | 144.79 | 5.49 | 69841 | 2 | 1041003.38 | 1027483.56 | 1018094.38 | 519047.03 | 623845.06 | 421181.72 | 1083433.5 | 565417.75 | 503988.16 | 653218.56 | 8 | **0.763** |
| Q99LM2 | Cdk5rap3 | CDK5 regulatory subunit-associated protein 3 (Mus musculus) 3 | 47.7 | 1.39 | 56991 | 1 | 133306184 | 1442050.62 | 1523536.88 | 1583155.38 | 1744535.25 | 132105368 | 1896311.75 | 1753782.25 | 1935830.5 | 1934968.5 | 5 | **1.000** |
| Q99LT0 | Dpy30 | Protein dpy-30 homolog (Mus musculus) | 48.82 | 16.16 | 11213 | 1 | 0 | 0 | 0 | 650923.44 | 1545847.88 | 0 | 563912.56 | 259623.22 | 1071803.25 | 444606.06 | 15 | **1.065** |
| Q99LX0 | Park7 | Protein/nucleic acid deglycase DJ-1 (Mus musculus) | 825.55 | 88.36 | 20021 | 14 | 66188672 | 50280728 | 65111592 | 43621908 | 54789348 | 73087440 | 73269168 | 57472572 | 62078836 | 62377040 | 6 | **1.172** |
| Q99M31 | Hspa14 | Heat shock 70 kDa protein 14 (Mus musculus) | 427.3 | 23.77 | 54650 | 8 | 7703953.5 | 5668461 | 6209080 | 6269161 | 4896831 | 6019456 | 5845610 | 7807020.5 | 6747370 | 6513098 | 8 | **1.071** |
| Q99M87 | Dnaja3 | DnaJ homolog subfamily A member 3, mitochondrial (Mus musculus) | 60.97 | 4.79 | 52443 | 1 | 377208.5 | 353633.91 | 343481.25 | 0 | 0 | 300145.69 | 420069.06 | 0 | 109494.73 | 0 | 14 | **0.772** |
| Q99MD9 | Nasp | Nuclear autoantigenic sperm protein (Mus musculus) | 964.37 | 27.43 | 83954 | 17 | 34222972 | 40995612 | 45379248 | 47274944 | 37626776 | 48652776 | 38170472 | 42756144 | 45707128 | 49072148 | 6 | **1.092** |
| Q99ME9 | Gtpbp4 | Nucleolar GTP-binding protein 1 (Mus musculus) | 178.98 | 8.04 | 74113 | 4 | 2389711 | 3296778.5 | 3079070.75 | 3817420 | 2890768.5 | 3169823.25 | 2348150.75 | 3358539.5 | 3675256.75 | 3443807.5 | 10 | **1.034** |
| Q99MN1 | Kars | Lysine--tRNA ligase (Mus musculus) | 957.92 | 35.29 | 67840 | 18 | 35569216 | 43304596 | 41927788 | 42372924 | 35315876 | 36758864 | 36533856 | 35219344 | 35697760 | 39572148 | 8 | **0.926** |
| Q99MR6 | Srrt | Serrate RNA effector molecule homolog (Mus musculus) | 348.83 | 10.51 | 100452 | 8 | 11748840 | 9569540 | 9667387 | 8976306 | 10029820 | 10620209 | 11031846 | 11659776 | 11816919 | 9739383 | 5 | **1.098** |
| Q99MS7 | Ehbp1l1 | EH domain-binding protein 1-like protein 1 (Mus musculus) | 126.33 | 2.04 | 184834 | 2 | 431970.53 | 316130.28 | 1060637.75 | 569065 | 494222.03 | 921312.25 | 885532.38 | 427532.34 | 870692.12 | 429417.31 | 11 | **1.231** |
| Q99N27 | Snx1 | Sorting nexin-1 (Rattus norvegicus) | 395.69 | 15.9 | 59044 | 7 | 13247040 | 7685032 | 5986345 | 7539753 | 6280757.5 | 5975976 | 17566564 | 14605307 | 17962894 | 13508622 | 5 | **1.709** |
| Q99N69 | Lpxn | Leupaxin (Mus musculus) | 473.67 | 26.68 | 43478 | 8 | 12960576 | 10254100 | 8059292.5 | 9828612 | 9408454 | 12019072 | 11896353 | 13008258 | 10396535 | 11670607 | 4 | **1.168** |
| Q99N96 | Mrpl1 | 39S ribosomal protein L1, mitochondrial (Mus musculus) | 39.88 | 6.25 | 37597 | 1 | 0 | 0 | 0 | 505910.19 | 0 | 158362.58 | 0 | 603523.5 | 436613.62 | 0 | 18 | **2.369** |
| Q99NB9 | Sf3b1 | Splicing factor 3B subunit 1 (Mus musculus) | 420.36 | 7.82 | 145816 | 8 | 1539452.75 | 4176109.5 | 4933006 | 5742156 | 3361791 | 4010634.25 | 3572001.25 | 4866424.5 | 4245677 | 3904421.25 | 12 | **1.043** |
| Q99NF8 | Ranbp17 | Ran-binding protein 17 (Mus musculus) | 50.12 | 1.01 | 124054 | 1 | 0 | 232795.25 | 313646.47 | 457351.75 | 325061.94 | 325529.59 | 0 | 253893.19 | 235999.81 | 340698.03 | 13 | **0.870** |
| Q99P31 | Hspbp1 | Hsp70-binding protein 1 (Mus musculus) | 396.81 | 24.65 | 39167 | 6 | 15572829 | 16648936 | 15135134 | 16674818 | 14696916 | 17444946 | 14111812 | 16331096 | 16265420 | 15114473 | 12 | **1.007** |
| Q99P72 | Rtn4 | Reticulon-4 (Mus musculus) | 891.51 | 21.69 | 126613 | 14 | 45816880 | 58522240 | 56102604 | 62642624 | 50772680 | 53750636 | 53537292 | 59837696 | 57757176 | 65475396 | 9 | **1.060** |
| Q99P75 | Rab9a | Ras-related protein Rab-9A (Rattus norvegicus) | 54.27 | 5.47 | 22896 | 1 | 3000558.5 | 1572662.12 | 1506127.5 | 1586334.62 | 2292861.5 | 2083796.88 | 2084933 | 2183530.5 | 2227373.75 | 2087182.75 | 10 | **1.071** |
| Q99P88 | Nup155 | Nuclear pore complex protein Nup155 (Mus musculus) | 783.91 | 12.15 | 155118 | 13 | 11182187 | 12447760 | 11095671 | 10858871 | 11117034 | 12973682 | 10414991 | 12427567 | 13311084 | 12429459 | 7 | **1.086** |
| Q99PD4 | Arpc1a | Actin-related protein 2/3 complex subunit 1A (Rattus norvegicus) | 130.88 | 9.46 | 41599 | 2 | 6035380.5 | 6787036 | 7923336 | 6472612 | 7558356 | 9062375 | 6632706 | 7470189 | 6419342 | 7105123 | 11 | **1.055** |
| Q99PG2 | Ogfr | Opioid growth factor receptor (Mus musculus) | 102.53 | 8.85 | 70679 | 2 | 521176.38 | 0 | 502617.75 | 0 | 953401.94 | 508800.19 | 460648.72 | 473626.31 | 1102910.62 | 992146.12 | 9 | **1.789** |
| Q99PK0 | Xab2 | Pre-mRNA-splicing factor SYF1 (Rattus norvegicus) | 62.48 | 1.99 | 99998 | 1 | 132857.44 | 0 | 0 | 0 | 0 | 277672 | 229168.89 | 319614.47 | 190883.62 | 167058.12 | 6 | **8.915** |
| Q99PL5 | Rrbp1 | Ribosome-binding protein 1 (Mus musculus) | 1563.16 | 38.5 | 172879 | 29 | 64885532 | 55760372 | 44423072 | 54862152 | 59104580 | 48938280 | 52552184 | 54266524 | 63195668 | 58371832 | 10 | **0.994** |
| Q99PV0 | Prpf8 | Pre-mRNA-processing-splicing factor 8 (Mus musculus) | 758.85 | 7.15 | 273616 | 16 | 486507968 | 13969936 | 15964204 | 15242156 | 801006464 | 16723350 | 13393897 | 18160926 | 17272190 | 19922660 | 12 | **0.064** |
| Q9CPN8 | Igf2bp3 | Insulin-like growth factor 2 mRNA-binding protein 3 (Mus musculus) | 406.79 | 17.44 | 63575 | 7 | 4909105.5 | 9030501 | 7496375 | 8158534.5 | 7229757 | 7450758 | 7182306 | 5941963 | 5580435 | 5982077 | 6 | **0.873** |
| Q9CPP0 | Npm3 | Nucleoplasmin-3 (Mus musculus) | 94.76 | 9.14 | 19023 | 1 | 3319986 | 2820344.75 | 2644307.75 | 2678283.5 | 2332057.75 | 2381816.5 | 2776435 | 2249629.25 | 2343969.25 | 2039326.25 | 5 | **0.855** |
| Q9CPQ1 | Cox6c | Cytochrome c oxidase subunit 6C (Mus musculus) | 106.48 | 19.74 | 8469 | 2 | 21094392 | 18046158 | 15667896 | 30155534 | 7423071 | 15562008 | 14642746 | 14845371 | 16029314 | 7082901.5 | 5 | **0.738** |
| Q9CPQ3 | Tomm22 | Mitochondrial import receptor subunit TOM22 homolog (Mus musculus) | 124.35 | 22.54 | 15537 | 2 | 4343423.5 | 5181914 | 5186045 | 4550509 | 4267228.5 | 4410889 | 5283629 | 4458340 | 4421576.5 | 4580708 | 11 | **0.984** |
| Q9CPR5 | Mrpl15 | 39S ribosomal protein L15, mitochondrial (Mus musculus) | 111.17 | 7.46 | 33542 | 2 | 2371457.5 | 1508126.25 | 1028735.94 | 1515901.5 | 1257038.25 | 1196945.62 | 1935527.38 | 2295281 | 1013390.81 | 2108854.5 | 12 | **1.113** |
| Q9CPT4 | Mydgf | Myeloid-derived growth factor (Mus musculus) | 208.96 | 28.31 | 17982 | 4 | 7283248.5 | 10134368 | 10376450 | 10233379 | 10994415 | 4304942 | 9240033 | 10858968 | 11151473 | 10870402 | 11 | **0.947** |
| Q9CPT5 | Nop16 | Nucleolar protein 16 (Mus musculus) | 139.7 | 16.85 | 21139 | 2 | 1147313.38 | 1331243.25 | 1670305.75 | 1341552.38 | 1860874.75 | 1156373 | 1870424.12 | 1269050.75 | 1650001.38 | 1606115.5 | 12 | **1.027** |
| Q9CPU0 | Glo1 | Lactoylglutathione lyase (Mus musculus) | 335.4 | 33.15 | 20810 | 7 | 31343350 | 22311566 | 18502672 | 49628440 | 32033424 | 19816388 | 10980174 | 19227870 | 22145792 | 20715206 | 4 | **0.604** |
| Q9CPU4 | Mgst3 | Microsomal glutathione S-transferase 3 (Mus musculus) | 258.97 | 40.52 | 16958 | 4 | 7873490 | 7880625 | 8046954 | 9220037 | 8375414 | 9288208 | 8157046 | 10196396 | 10051416 | 9972051 | 2 | **1.151** |
| Q9CPV4 | Glod4 | Glyoxalase domain-containing protein 4 (Mus musculus) | 618.83 | 51.34 | 33317 | 12 | 30235516 | 24462858 | 21367424 | 23760952 | 23552250 | 31033778 | 27422508 | 23903274 | 24425824 | 24773730 | 6 | **1.066** |
| Q9CPV5 | Pmf1 | Polyamine-modulated factor 1 (Mus musculus) | 64.65 | 5.94 | 23121 | 1 | 509188.28 | 0 | 0 | 325431.84 | 354425.91 | 370470.09 | 378448.59 | 463704.06 | 384254.44 | 385550.34 | 6 | **1.667** |

| | | | | | | | | | | | | | | | | | |
|---|---|---|---|---|---|---|---|---|---|---|---|---|---|---|---|---|---|
| Q9CPW4 | Arpc5 | Actin-related protein 2/3 complex subunit 5 (Mus musculus) | 434.44 | 39.07 | 16288 | 6 | 71049464 | 52425888 | 46085056 | 57201964 | 50704144 | 60112972 | 53818044 | 54430560 | 64910232 | 62115496 | 7 | **1.065** |
| Q9CPY7 | Lap3 | Cytosol aminopeptidase (Mus musculus) | 626.14 | 31.98 | 56141 | 12 | 13409480 | 20250166 | 19906172 | 17165248 | 16343308 | 12194322 | 16127598 | 14561162 | 13949388 | 14774241 | 4 | **0.822** |
| Q9CQ54 | Ndufc2 | NADH dehydrogenase [ubiquinone] 1 subunit C2 (Mus musculus) | 41.77 | 10.83 | 14164 | 1 | 1045378.62 | 1132764.5 | 953640 | 1775177.5 | 1348816.5 | 1061914.12 | 1208689.5 | 1054949.62 | 1128712.12 | 1270865 | 12 | **0.915** |
| Q9CQ60 | Pgls | 6-phosphogluconolactonase (Mus musculus) | 392.27 | 47.86 | 27254 | 7 | 20699124 | 13954372 | 10299925 | 14906457 | 12015145 | 16443679 | 15241248 | 19065316 | 13538552 | 13141846 | 9 | **1.077** |
| Q9CQ62 | Decr1 | 2,4-dienoyl-CoA reductase, mitochondrial (Mus musculus) | 162.14 | 11.04 | 36214 | 4 | 540787.44 | 2834693.75 | 859571.62 | 64839.73 | 2338887.75 | 1312149.25 | 1129316.62 | 1844415.5 | 2222646.25 | 3158420 | 8 | **1.456** |
| Q9CQ65 | Mtap | S-methyl-5'-thioadenosine phosphorylase (Mus musculus) | 484.46 | 44.17 | 31062 | 8 | 15089797 | 21921236 | 24425776 | 24689394 | 26273898 | 25025646 | 15949322 | 25784286 | 26170438 | 26202882 | 8 | **1.060** |
| Q9CQ71 | Rpa3 | Replication protein A 14 kDa subunit (Mus musculus) | 238.23 | 51.24 | 13584 | 4 | 3075260.25 | 2460618.5 | 1729436.38 | 2619752.25 | 1871982 | 3661612 | 3433842.25 | 3251912 | 3553032 | 3497001.25 | 0 | **1.480** |
| Q9CQ75 | Ndufa2 | NADH dehydrogenase [ubiquinone] 1 alpha subcomplex subunit 2 (Mus musculus) | 71.34 | 14.14 | 10916 | 1 | 3110744.25 | 2076563.75 | 2393681.5 | 1537964.62 | 1029100.81 | 1146113.25 | 1901423 | 1552219.25 | 1342123.12 | 1659472.88 | 8 | **0.749** |
| Q9CQ86 | Mien1 | Migration and invasion enhancer 1 (Mus musculus) | 40.69 | 9.57 | 12295 | 1 | 608642.5 | 1535380.75 | 1172826.5 | 1203453.38 | 1591012.62 | 973502.5 | 860928.25 | 1153229.5 | 0 | 0 | 4 | **0.489** |
| Q9CQ88 | Tspan31 | Tetraspanin-31 (Mus musculus) | 69.44 | 5.71 | 22694 | 1 | 0 | 761594.56 | 1024109.69 | 810266.38 | 701734.12 | 799321.25 | 1060527.38 | 1028808.81 | 785979.06 | 779775.25 | 6 | **1.351** |
| Q9CQA1 | Trappc5 | Trafficking protein particle complex subunit 5 (Mus musculus) | 99.27 | 10.11 | 20795 | 2 | 3209936.75 | 1065323.12 | 1064606.62 | 1222885.12 | 1002624.44 | 2836750.5 | 2297566 | 2359332 | 2461132.75 | 0 | 9 | **1.316** |
| Q9CQA3 | Sdhb | Succinate dehydrogenase [ubiquinone] iron-sulfur subunit, mitochondrial (Mus musculus) | 417.47 | 30.14 | 31814 | 8 | 11368674 | 9281094 | 7117027 | 8758237 | 7503903 | 9048792 | 13666248 | 12100678 | 14750155 | 14371498 | 2 | **1.452** |
| Q9CQB5 | Cisd2 | CDGSH iron-sulfur domain-containing protein 2 (Mus musculus) | 139.96 | 25.19 | 15242 | 3 | 4097184.5 | 3533409.5 | 2986784 | 4836630 | 4493424.5 | 4189925.75 | 4768560 | 3762898.5 | 3846023.25 | 4147939.75 | 11 | **1.038** |
| Q9CQD1 | Rab5a | Ras-related protein Rab-5A (Mus musculus) | 316.34 | 30.23 | 23599 | 5 | 12268211 | 28844208 | 19191204 | 23963984 | 17034404 | 21966848 | 18665918 | 22563012 | 20521560 | 23613548 | 11 | **1.060** |
| Q9CQE1 | Nipsnap3b | Protein NipSnap homolog 3B (Mus musculus) 3b | 51.97 | 4.45 | 28308 | 1 | 264524.09 | 595678.38 | 0 | 595349.19 | 742821.69 | 881717.31 | 620318.94 | 529422.25 | 539596.12 | 591232.25 | 10 | **1.438** |
| Q9CQE3 | Mrps17 | 28S ribosomal protein S17, mitochondrial (Mus musculus) | 40.26 | 7.5 | 13382 | 1 | 1369420.88 | 656782.31 | 959469.19 | 0 | 962920.5 | 1738710.38 | 1159993.25 | 1344940.38 | 1465009.62 | 1398791.75 | 2 | **1.800** |
| Q9CQE5 | Rgs10 | Regulator of G-protein signaling 10 (Mus musculus) | 65.62 | 9.39 | 21151 | 1 | 367570.75 | 337842.38 | 332317.97 | 487593.31 | 499158.75 | 259497.23 | 368157.28 | 271342.94 | 367826.97 | 308425.47 | 6 | **0.778** |
| Q9CQE8 | RTRAF | RNA transcription, translation and transport factor protein (Mus musculus) | 254.6 | 25.82 | 28152 | 4 | 1699404.38 | 1898349.12 | 967999.75 | 1224428.88 | 1885529.5 | 2053089.88 | 2218021.25 | 1542123.75 | 1714525.75 | 1168535.38 | 9 | **1.133** |
| Q9CQF9 | Pcyox1 | Prenylcysteine oxidase (Mus musculus) | 381.22 | 19.41 | 56495 | 7 | 6892295 | 7626065.5 | 6864704.5 | 8032805 | 7116890.5 | 9268172 | 9239152 | 11645286 | 10430830 | 11689953 | 0 | **1.431** |
| Q9CQG2 | Mettl16 | U6 small nuclear RNA (adenine-(43)-N(6))-methyltransferase (Mus musculus) | 126.59 | 5.42 | 62341 | 2 | 684657.19 | 65593.02 | 781252.88 | 844666.62 | 833397.38 | 857821 | 787645.31 | 799760.25 | 842192.12 | 1040571.88 | 5 | **1.348** |
| Q9CQH3 | Ndufb5 | NADH dehydrogenase [ubiquinone] 1 beta subcomplex subunit 5, mitochondrial (Mus musc | 174.28 | 16.4 | 21710 | 3 | 2802826.5 | 8662226 | 11043240 | 6822584 | 699354.19 | 4469555.5 | 4397600.5 | 11819106 | 10786934 | 14749999 | 7 | **1.539** |
| Q9CQI6 | Cotl1 | Coactosin-like protein (Mus musculus) | 606.9 | 76.76 | 15944 | 12 | 150418640 | 151537408 | 131495136 | 177060720 | 130922136 | 157522608 | 122173592 | 130152712 | 141231376 | 134520480 | 8 | **0.925** |
| Q9CQI7 | Snrpb2 | U2 small nuclear ribonucleoprotein B'' (Mus musculus) | 92.77 | 12 | 25323 | 2 | 1767373.25 | 3758408.25 | 2300881.75 | 3964264.25 | 4018227 | 798945.25 | 385133.72 | 2311499.5 | 3513420 | 2827085.75 | 6 | **0.622** |
| Q9CQJ6 | Denr | Density-regulated protein (Mus musculus) | 192.5 | 21.72 | 22166 | 3 | 4138450.75 | 2496802.5 | 2833473 | 2412917.25 | 3322617.75 | 4089865.75 | 3608414 | 3426734.5 | 3530424.25 | 3484462 | 5 | **1.193** |
| Q9CQK7 | Rwdd1 | RWD domain-containing protein 1 (Mus musculus) | 109.47 | 17.28 | 27785 | 2 | 589740.81 | 201323.64 | 286797.31 | 560000.5 | 703177.19 | 1031149.25 | 878814.12 | 986013.31 | 752585.12 | 656689.88 | 1 | **1.839** |
| Q9CQM5 | Txndc17 | Thioredoxin domain-containing protein 17 (Mus musculus) | 214.32 | 39.84 | 14015 | 4 | 16364702 | 13356868 | 12355688 | 10710824 | 13275750 | 13665620 | 13069894 | 14191216 | 14617703 | 14317228 | 7 | **1.057** |
| Q9CQM9 | Glrx3 | Glutaredoxin-3 (Mus musculus) | 519.03 | 42.73 | 37778 | 8 | 14396173 | 11350011 | 7729766.5 | 9542693 | 11866800 | 13651358 | 14057772 | 13548795 | 10264004 | 12196846 | 7 | **1.161** |
| Q9CQN1 | Trap1 | Heat shock protein 75 kDa, mitochondrial (Mus musculus) | 862.59 | 29.75 | 80209 | 16 | 32524462 | 25465564 | 24408638 | 22926796 | 16997520 | 25081724 | 20315512 | 22917318 | 28394912 | 24456546 | 12 | **0.991** |
| Q9CQN6 | Tmem14c | Transmembrane protein 14C (Mus musculus) | 385.4 | 92.11 | 11642 | 6 | 1114018 | 7346104.5 | 7366443.5 | 7029066 | 4501439.5 | 5924313.5 | 4046965.5 | 7147412.5 | 5869912 | 6659107 | 10 | **1.084** |
| Q9CQQ7 | Atp5pb | ATP synthase F(0) complex subunit B1, mitochondrial (Mus musculus) | 289.89 | 21.48 | 28949 | 5 | 21317676 | 22798760 | 28688532 | 34661544 | 22568712 | 13849307 | 17841796 | 13842794 | 26095972 | 14443022 | 3 | **0.662** |
| Q9CQQ8 | Lsm7 | U6 snRNA-associated Sm-like protein LSm7 (Mus musculus) | 96.21 | 15.53 | 11636 | 2 | 4432946 | 2172027 | 2598312.25 | 2973901.5 | 4517110.5 | 3129700.5 | 5190150 | 2113139.5 | 2449444 | 4257672.5 | 12 | **1.027** |
| Q9CQR4 | Acot13 | Acyl-coenzyme A thioesterase 13 (Mus musculus) | 43.35 | 7.14 | 15183 | 1 | 2350318 | 2300637.25 | 2317188.5 | 2516848 | 689587.19 | 1891232.38 | 1935178.5 | 0 | 1380975.88 | 2143949 | 4 | **0.723** |
| Q9CQS8 | Sec61b | Protein transport protein Sec61 subunit beta (Mus musculus) | 130.92 | 26.04 | 9958 | 2 | 10415725 | 6512419.5 | 7987371 | 6252709 | 7950428 | 9107331 | 9960442 | 7555273 | 9062972 | 8810269 | 7 | **1.137** |
| Q9CQT1 | Mri1 | Methylthioribose-1-phosphate isomerase (Mus musculus) | 310.61 | 20.6 | 39411 | 5 | 7683135 | 6945606.5 | 5958579 | 7625509 | 6173886 | 7331405 | 8098666 | 6899673.5 | 6336974 | 6389172 | 11 | **1.019** |
| Q9CQT2 | Rbm7 | RNA-binding protein 7 (Mus musculus) | 51.68 | 6.79 | 30148 | 1 | 113288.78 | 218322.41 | 0 | 205124.56 | 135139.28 | 321334.56 | 249859.88 | 269806.84 | 282269.69 | 286990.66 | 0 | **2.099** |
| Q9CQV5 | Mrps24 | 28S ribosomal protein S24, mitochondrial (Mus musculus) | 128.26 | 17.37 | 18901 | 2 | 1004762.56 | 794578.38 | 803018.56 | 833196.62 | 837810.5 | 648034.19 | 903432.06 | 1030394.94 | 946055.12 | 643132.75 | 12 | **0.976** |
| Q9CQV8 | Ywhab | 14-3-3 protein beta/alpha (Mus musculus) | 1470.17 | 73.17 | 28086 | 22 | 211338032 | 421808864 | 553469440 | 432647744 | 314257024 | 290761088 | 257243152 | 275770336 | 345204128 | 351240832 | 7 | **0.786** |
| Q9CQW1 | Ykt6 | Synaptobrevin homolog YKT6 (Mus musculus) | 305.42 | 29.8 | 22314 | 6 | 7204078.5 | 13856776 | 13202056 | 12365305 | 12364993 | 10531946 | 12231920 | 13062408 | 11153522 | 13297451 | 10 | **1.022** |
| Q9CQW9 | Ifitm3 | Interferon-induced transmembrane protein 3 (Mus musculus) | 437.4 | 48.91 | 14954 | 5 | 43844404 | 19577750 | 19529572 | 21081752 | 18840788 | 34581264 | 41405292 | 38811552 | 37279812 | 34364804 | 5 | **1.517** |
| Q9CQX2 | Cyb5b | Cytochrome b5 type B (Mus musculus) | 66.2 | 8.22 | 16318 | 1 | 2640448.25 | 2288399.5 | 2204771.25 | 2475343.75 | 2478909.5 | 1955339 | 2083769.88 | 1672124.12 | 2440531 | 2210081.75 | 3 | **0.857** |
| Q9CR16 | Ppid | Peptidyl-prolyl cis-trans isomerase D (Mus musculus) | 639.14 | 36.49 | 40743 | 13 | 41659696 | 32369040 | 50393832 | 41306436 | 49812404 | 50582292 | 51662556 | 52744068 | 51158892 | 53046500 | 0 | **1.203** |
| Q9CR27 | Washc3 | WASH complex subunit 3 (Mus musculus) | 61.83 | 9.28 | 21093 | 1 | 347727.12 | 587945.56 | 736059.5 | 0 | 667506 | 631538 | 607678.75 | 569883.38 | 0 | 0 | 11 | **0.773** |
| Q9CR29 | Ccdc43 | Coiled-coil domain-containing protein 43 (Mus musculus) | 82.82 | 11.71 | 25049 | 1 | 179652.45 | 331975.22 | 355201.5 | 322569 | 323792.75 | 314094.31 | 408817.97 | 372309.22 | 505899.75 | 466222.09 | 4 | **1.366** |
| Q9CR51 | Atp6v1g1 | V-type proton ATPase subunit G 1 (Mus musculus) 1 | 157.99 | 22.03 | 13724 | 2 | 1482906 | 2844779 | 1065511.62 | 2168409.25 | 2895523.5 | 4195065 | 3591799.5 | 1268889.38 | 3155619.75 | 3388566.75 | 4 | **1.492** |
| Q9CR57 | Rpl14 | 60S ribosomal protein L14 (Mus musculus) | 345.72 | 25.81 | 23564 | 5 | 112028944 | 80873568 | 73920256 | 96565704 | 90532888 | 99460256 | 96275480 | 95675104 | 96853048 | 88985440 | 9 | **1.051** |
| Q9CR60 | Golt1b | Vesicle transport protein GOT1B (Mus musculus) | 157.01 | 10.87 | 15422 | 2 | 1616039.25 | 1400524 | 1945564.88 | 1366455.5 | 1708437 | 2280811.75 | 1871167 | 1435560.88 | 1808201.38 | 1429692.75 | 8 | **1.098** |
| Q9CR62 | Slc25a11 | Mitochondrial 2-oxoglutarate/malate carrier protein (Mus musculus) 1 | 710.25 | 48.73 | 34155 | 14 | 23603856 | 27491832 | 26932738 | 31055646 | 21351958 | 25016368 | 22673472 | 30451170 | 29687182 | 31024056 | 10 | **1.065** |
| Q9CR64 | Tmem167a | Protein kish-A (Mus musculus) a | 57.5 | 12.5 | 8074 | 1 | 877972.5 | 490482.53 | 735381.94 | 390407.94 | 661882.88 | 987581.19 | 974726.75 | 823104.44 | 687526.94 | 719444.75 | 5 | **1.328** |
| Q9CR67 | Tmem33 | Transmembrane protein 33 (Mus musculus) | 123.55 | 8.91 | 28031 | 2 | 10065258 | 5819804 | 5621442.5 | 6438515 | 5366103 | 8102109 | 8813931 | 8259984 | 7390081 | 8015256 | 5 | **1.218** |
| Q9CR68 | Uqcrfs1 | Cytochrome b-c1 complex subunit Rieske, mitochondrial (Mus musculus) | 243.43 | 25.91 | 29368 | 5 | 8888792 | 9792534 | 10887452 | 9955205 | 10268088 | 8529122 | 10527899 | 19293138 | 15192976 | 21988226 | 6 | **1.517** |
| Q9CR95 | Necap1 | Adaptin ear-binding coat-associated protein 1 (Mus musculus) | 159.14 | 19.27 | 29639 | 3 | 5198836.5 | 5108206 | 5146017.5 | 4475055 | 5836264 | 6528992 | 5861236 | 5582994 | 6944971.5 | 8353821 | 1 | **1.291** |
| Q9CRA5 | Golph3 | Golgi phosphoprotein 3 (Mus musculus) | 177.19 | 9.73 | 33752 | 2 | 539239.88 | 455333.5 | 363709.81 | 260713.3 | 686089.19 | 301430.03 | 374902.25 | 454688.91 | 453717.12 | 448034.12 | 9 | **0.882** |
| Q9CRA8 | Exosc5 | Exosome complex component RRP46 (Mus musculus) | 90.24 | 9.79 | 25194 | 2 | 1728609.75 | 1799424.88 | 1806845.75 | 1409796.88 | 1649902.75 | 1609271.25 | 1171255.38 | 1799856.88 | 1221595.38 | 1288393 | 5 | **0.845** |
| Q9CRB2 | Nhp2 | H/ACA ribonucleoprotein complex subunit 2 (Mus musculus) | 125.05 | 16.99 | 17247 | 2 | 3211164.75 | 493133.47 | 740985 | 416939.59 | 799840.88 | 3106828.5 | 3333431 | 1493188.12 | 2794678 | 980864.31 | 4 | **2.068** |
| Q9CRB9 | Chchd3 | MICOS complex subunit Mic19 (Mus musculus) | 196.98 | 15.42 | 26335 | 3 | 7983322 | 4768500.5 | 4916917 | 5166007 | 6526365 | 3568731.75 | 6183645.5 | 5185212 | 6033158 | 5530000 | 12 | **0.903** |
| Q9CRC8 | Lrrc40 | Leucine-rich repeat-containing protein 40 (Mus musculus) | 225.03 | 9.3 | 68076 | 4 | 2583905 | 3814866.5 | 4153568 | 2637148.75 | 3577901.5 | 3122844.25 | 3057639.5 | 3882059 | 2158478.75 | 1797292.75 | 8 | **0.836** |
| Q9CRD0 | Ociad1 | OCIA domain-containing protein 1 (Mus musculus) | 128.55 | 12.55 | 27610 | 2 | 679029.69 | 0 | 491221.84 | 0 | 297963.91 | 380109 | 566123.25 | 555079.25 | 430684.06 | 543132.56 | 8 | **1.686** |
| Q9CRT8 | Xpot | Exportin-T (Mus musculus) | 283.19 | 7.58 | 109734 | 6 | 1343875.5 | 3111725.25 | 4028265 | 2387883 | 2025572.25 | 3430075.25 | 3375975 | 3255413.5 | 2514240.5 | 3499845.5 | 6 | **1.246** |
| Q9CSH3 | Dis3 | Exosome complex exonuclease RRP44 (Mus musculus) | 151.07 | 3.76 | 108838 | 3 | 2414432.25 | 1820696.25 | 1459494.75 | 1825859.62 | 1918338.75 | 3642321.25 | 2449488.75 | 2863788.75 | 3002330 | 3000953.5 | 0 | **1.585** |
| Q9CSN1 | Snw1 | SNW domain-containing protein 1 (Mus musculus) | 112.03 | 5.41 | 61475 | 2 | 2802648.5 | 1425823.75 | 1417772.25 | 1332470.25 | 1829828.88 | 1656317.25 | 1867767.5 | 1360345.62 | 918263 | 1676841 | 11 | **0.849** |
| Q9CT10 | Ranbp3 | Ran-binding protein 3 (Mus musculus) | 423.3 | 18.53 | 52573 | 8 | 6660741 | 10985549 | 12498015 | 12893815 | 13497025 | 11606323 | 7308438 | 12377914 | 12766257 | 11170220 | 10 | **0.977** |
| Q9CU62 | Smc1a | Structural maintenance of chromosomes protein 1A (Mus musculus) | 321.02 | 8.03 | 143235 | 8 | 5446286 | 6592218.5 | 6468164 | 8481374 | 5739094 | 6853970.5 | 1787076.5 | 3010859.5 | 5623278.5 | 3664608 | 5 | **0.640** |
| Q9CV28 | MINDY3 | Ubiquitin carboxyl-terminal hydrolase MINDY-3 (Mus musculus) | 43.13 | 2.25 | 49612 | 1 | 0 | 676294.12 | 488769.78 | 635481.5 | 746643.06 | 0 | 0 | 903947.38 | 544941.31 | 0 | 13 | **0.569** |
| Q9CVB6 | Arpc2 | Actin-related protein 2/3 complex subunit 2 (Mus musculus) | 959.86 | 62 | 34357 | 16 | 62546368 | 87909112 | 87676712 | 86902696 | 77169184 | 84461352 | 67535096 | 81288248 | 82614184 | 91172696 | 12 | **1.012** |
| Q9CWG1 | Glipr1 | Glioma pathogenesis-related protein 1 (Mus musculus) | 53.19 | 5.49 | 29129 | 1 | 456548.62 | 913856.75 | 313281.16 | 0 | 0 | 169889.95 | 592312.56 | 545913.75 | 781356 | 313185.66 | 10 | **1.427** |
| Q9CWI3 | Bccip | BRCA2 and CDKN1A-interacting protein (Mus musculus) | 99.58 | 6.33 | 35942 | 2 | 1837276.62 | 3304052.75 | 3541547.5 | 3061363.75 | 2621491.5 | 2776857 | 3603490.5 | 2943430.5 | 3180598.75 | 3341156.25 | 9 | **1.103** |
| Q9CWJ9 | Atic | Bifunctional purine biosynthesis protein PURH (Mus musculus) | 2108.76 | 76.01 | 64217 | 32 | 140019184 | 111050624 | 127240280 | 103969248 | 119658144 | 160310688 | 132893920 | 143069312 | 151185024 | 150056608 | 1 | **1.225** |
| Q9CWK8 | Snx2 | Sorting nexin-2 (Mus musculus) | 955.69 | 44.32 | 58471 | 16 | 67583920 | 53456304 | 48991496 | 55543988 | 48486356 | 54772892 | 67436400 | 60668168 | 61810564 | 59963756 | 6 | **1.112** |
| Q9CWM4 | Pfdn1 | Prefoldin subunit 1 (Mus musculus) | 99.83 | 14.75 | 14255 | 2 | 3842645.25 | 2769300 | 2412802.5 | 2181594.5 | 2137758.75 | 2784954 | 3267057 | 2581610 | 2779780.75 | 2583674.5 | 7 | **1.049** |
| Q9CWR2 | Smyd3 | Histone-lysine N-methyltransferase SMYD3 (Mus musculus) | 42.08 | 4.91 | 49126 | 1 | 0 | 242969.84 | 231974.83 | 127005.41 | 134075.59 | 0 | 0 | 223790.28 | 154988.86 | 142603.77 | 12 | **0.708** |
| Q9CWU6 | Uqcc1 | Ubiquinol-cytochrome-c reductase complex assembly factor 1 (Mus musculus) | 57.61 | 4.07 | 34300 | 1 | 953780.88 | 610892.38 | 0 | 750761.06 | 0 | 668337.38 | 671048.69 | 774758.69 | 827160.94 | 859217.75 | 8 | **1.641** |
| Q9CWU9 | Nup37 | Nucleoporin Nup37 (Mus musculus) | 74.48 | 4.91 | 36732 | 1 | 798960.88 | 499201.47 | 616610.62 | 0 | 738644.62 | 627236.75 | 777977.19 | 460871.53 | 514881.84 | 403442.81 | 11 | **1.049** |
| Q9CWV0 | Malsu1 | Mitochondrial assembly of ribosomal large subunit protein 1 (Mus musculus) | 76.21 | 4.82 | 26020 | 1 | 0 | 339099.09 | 212951.58 | 286653.41 | 205491.89 | 155349.64 | 230695.39 | 151746.08 | 203500.98 | 174674.56 | 7 | **0.877** |
| Q9CWX9 | Ddx47 | Probable ATP-dependent RNA helicase DDX47 (Mus musculus) | 197.62 | 9.01 | 50639 | 3 | 7267906 | 5867251.5 | 6447483 | 20041004 | 13110208 | 6024638 | 5685044 | 9079100 | 8601678 | 8850202 | 10 | **0.725** |
| Q9CWZ7 | Napg | Gamma-soluble NSF attachment protein (Mus musculus) | 246.34 | 19.23 | 34732 | 5 | 10962052 | 5621424 | 7098498 | 5431298 | 10863155 | 8923849 | 5960009.5 | 12835385 | 11784595 | 11064537 | 5 | **1.265** |
| Q9CX34 | Sugt1 | Protein SGT1 homolog (Mus musculus) | 370.66 | 22.32 | 38159 | 5 | 1108764.75 | 2632485 | 3351879.5 | 2250812 | 3019070 | 2242049.5 | 3255818.75 | 2840295.75 | 3117599.25 | 3220806.75 | 9 | **1.187** |
| Q9CX56 | Psmd8 | 26S proteasome non-ATPase regulatory subunit 8 (Mus musculus) | 172.18 | 8.5 | 39930 | 3 | 1805178.62 | 1387584.62 | 1735092.5 | 1763565.88 | 1528873.5 | 1733114.12 | 2064466.5 | 1786391.5 | 2204234.5 | 2256274.5 | 4 | **1.222** |
| Q9CX86 | Hnrnpa0 | Heterogeneous nuclear ribonucleoprotein A0 (Mus musculus) | 572.24 | 38.36 | 30530 | 10 | 23357144 | 33862500 | 32961934 | 31042768 | 31943410 | 34163744 | 29647784 | 33153778 | 32103968 | 32840386 | 9 | **1.057** |
| Q9CXE7 | Tmed5 | Transmembrane emp24 domain-containing protein 5 (Mus musculus) | 227.02 | 24.02 | 26172 | 4 | 2205268.5 | 4116746.25 | 3726442.5 | 2622828.5 | 640654.06 | 3040643 | 4089347.5 | 3978910.75 | 1416248.75 | 3181781.75 | 10 | **1.180** |
| Q9CXF4 | Tbc1d15 | TBC1 domain family member 15 (Mus musculus) | 388.27 | 16.84 | 76527 | 8 | 4927086.5 | 7003759 | 6645591.5 | 7698582 | 5419636.5 | 7151216 | 7213941 | 7619623.5 | 7350306 | 8145407 | 4 | **1.183** |
| Q9CXF7 | Chd1l | Chromodomain-helicase-DNA-binding protein 1-like (Mus musculus) | 42.58 | 0.89 | 101438 | 1 | 581463.25 | 235970.42 | 365981.91 | 0 | 0 | 214051.44 | 521741.47 | 460702.28 | 401483.44 | 478082.62 | 8 | **1.754** |
| Q9CXK4 | Slc50a1 | Sugar transporter SWEET1 (Mus musculus) | 47.74 | 9.05 | 24649 | 1 | 957010.69 | 1288114.88 | 1350496 | 0 | 0 | 1167290.88 | 1872490.5 | 0 | 0 | 0 | 17 | **0.845** |
| Q9CXK8 | Nip7 | 60S ribosome subunit biogenesis protein NIP7 homolog (Mus musculus) | 51 | 7.78 | 20452 | 1 | 0 | 1032699.62 | 1011588.06 | 955821.31 | 1176357.12 | 1063046 | 1369235.38 | 1528731.88 | 1218545.75 | 1527116.88 | 1 | **1.606** |
| Q9CXL3 | | Uncharacterized protein C7orf50 homolog (Mus musculus) | 43.48 | 12.31 | 22168 | 1 | 347687.78 | 231196.38 | 235973.94 | 0 | 0 | 383839.38 | 569830.38 | 436099.09 | 0 | 264015 | 7 | **2.030** |
| Q9CXR1 | Dhrs7 | Dehydrogenase/reductase SDR family member 7 (Mus musculus) | 196.72 | 17.46 | 38167 | 4 | 0 | 926259136 | 593093312 | 1750078.88 | 1963938.62 | 0 | 526060256 | 2364125.75 | 2443241 | 1671129.25 | 11 | **0.350** |
| Q9CXV1 | Sdhd | Succinate dehydrogenase [ubiquinone] cytochrome b small subunit, mitochondrial (Mus mu | 74.43 | 16.35 | 17014 | 1 | 633490 | 500021.06 | 466687.66 | 245505.58 | 492538.28 | 196542.86 | 737750.88 | 312846.16 | 398958.44 | 319156.09 | 8 | **0.840** |
| Q9CXW3 | Cacybp | Calcyclin-binding protein (Mus musculus) | 687.09 | 55.9 | 26510 | 12 | 47929868 | 47314848 | 51411084 | 40950408 | 61759380 | 52396956 | 41942344 | 49218428 | 55300516 | 49983824 | 10 | **0.998** |
| Q9CXY6 | Ilf2 | Interleukin enhancer-binding factor 2 (Mus musculus) | 503.65 | 34.36 | 43062 | 9 | 11812674 | 19565306 | 15598460 | 21229928 | 14029301 | 12271469 | 14012394 | 12995568 | 12739020 | 13591972 | 5 | **0.798** |

| | | | | | | | | | | | | | | | | |
|---|---|---|---|---|---|---|---|---|---|---|---|---|---|---|---|---|
| Q9CXY9 | Pigk | GPI-anchor transamidase (Mus musculus) | 62.14 | 3.04 | 44895 | 1 | 2138747.25 | 1220754.5 | 1730883.5 | 0 | 2198978.75 | 1684602 | 1792214 | 1499706.38 | 1591990.12 | 1386871.62 | 11 | **1.091** |
| Q9CY16 | Mrps28 | 28S ribosomal protein S28, mitochondrial (Mus musculus) | 117.31 | 17.74 | 20520 | 2 | 2373291.5 | 2306918.75 | 1451857.75 | 1642238.62 | 1999273.75 | 409357.09 | 1756940.88 | 2030332.62 | 1908983.5 | 1837747 | 9 | **0.813** |
| Q9CY21 | Bud23 | Probable 18S rRNA (guanine-N(7))-methyltransferase (Mus musculus) | 40.48 | 4.98 | 31587 | 1 | 0 | 98218.77 | 195778.52 | 106916.13 | 108682.51 | 0 | 0 | 28970.82 | 151868.8 | 147375.7 | 12 | **0.644** |
| Q9CY64 | Blvra | Biliverdin reductase A (Mus musculus) | 712.06 | 43.39 | 33525 | 11 | 37978756 | 31394310 | 37498984 | 29714908 | 40354500 | 45465744 | 39386104 | 42020936 | 45371616 | 41404132 | 1 | **1.207** |
| Q9CY73 | Mrpl44 | 39S ribosomal protein L44, mitochondrial (Mus musculus) | 118.1 | 11.71 | 37527 | 2 | 254988.25 | 288553.34 | 256667.09 | 0 | 0 | 251743.73 | 472353.12 | 396906.78 | 0 | 287047.84 | 10 | **1.760** |
| Q9CYA6 | Zcchc8 | Zinc finger CCHC domain-containing protein 8 (Mus musculus) | 90.74 | 4.65 | 78026 | 2 | 1798671.75 | 1296171.75 | 1688788.75 | 608881.12 | 981253.12 | 1003193.56 | 1775096.5 | 683769.44 | 1691871.5 | 1868056.25 | 9 | **1.102** |
| Q9CYG7 | Tomm34 | Mitochondrial import receptor subunit TOM34 (Mus musculus) | 379.07 | 27.51 | 34278 | 7 | 5287659.5 | 5990328 | 5758038 | 5090597 | 6729947 | 5405581 | 5779508.5 | 5489584 | 5694582 | 5039532 | 9 | **0.950** |
| Q9CYI4 | Luc7l | Putative RNA-binding protein Luc7-like 1 (Mus musculus) | 161.51 | 10.51 | 43934 | 4 | 7813986.5 | 8376556 | 9208730 | 7518894.5 | 7470441.5 | 8373859.5 | 7860780 | 8878388 | 8152167 | 9221264 | 7 | **1.052** |
| Q9CYN2 | Spcs2 | Signal peptidase complex subunit 2 (Mus musculus) | 232.52 | 18.58 | 24978 | 3 | 1528556.88 | 4077244.25 | 5966314 | 4619397.5 | 1853927 | 4236097 | 4411643.5 | 4717997 | 5516179 | 3875768.5 | 9 | **1.261** |
| Q9CYR0 | Ssbp1 | Single-stranded DNA-binding protein, mitochondrial (Mus musculus) | 227.82 | 30.26 | 17319 | 4 | 10558350 | 5674872 | 6556359 | 9835778 | 10156011 | 11468485 | 12550331 | 11595112 | 11997526 | 12367304 | 0 | **1.402** |
| Q9CZ04 | Cops7a | COP9 signalosome complex subunit 7a (Mus musculus) | 176.11 | 10.91 | 30224 | 2 | 1489736.62 | 1203717.25 | 1216569.88 | 908845.88 | 1182364.5 | 1174319.88 | 1486544.75 | 1228669 | 1375076.5 | 1153486.25 | 11 | **1.069** |
| Q9CZ13 | Uqcrc1 | Cytochrome b-c1 complex subunit 1, mitochondrial (Mus musculus) | 708.63 | 33.75 | 52852 | 13 | 39447500 | 35801444 | 37517368 | 35211288 | 38139896 | 42716932 | 41470508 | 39756084 | 39110492 | 40235836 | 1 | **1.092** |
| Q9CZ30 | Ola1 | Obg-like ATPase 1 (Mus musculus) | 578.52 | 36.36 | 44730 | 10 | 46892232 | 33715716 | 33995528 | 35471488 | 31192036 | 32075734 | 25406304 | 30398128 | 31117650 | 31116110 | 1 | **0.828** |
| Q9CZ44 | Nsfl1c | NSFL1 cofactor p47 (Mus musculus) | 334.86 | 19.73 | 40710 | 5 | 11258613 | 6085267 | 3890711.5 | 3824070 | 9168937 | 8732631 | 4945907 | 6745618.5 | 8747398 | 7470486 | 11 | **1.071** |
| Q9CZB0 | Sdhc | Succinate dehydrogenase cytochrome b560 subunit, mitochondrial (Mus musculus) | 191.36 | 28.99 | 18382 | 4 | 4360575 | 6329942 | 4552827.5 | 6264042 | 3698726.5 | 5801643 | 6907908 | 6623424 | 6539664 | 7168187 | 2 | **1.311** |
| Q9CZD3 | Gars | Glycine--tRNA ligase (Mus musculus) | 1668.94 | 45.95 | 81878 | 27 | 123322168 | 141775696 | 145889008 | 139078976 | 128201096 | 141099024 | 126942824 | 144285536 | 157748672 | 152628224 | 7 | **1.066** |
| Q9CZE3 | Rab32 | Ras-related protein Rab-32 (Mus musculus) | 142.99 | 13.45 | 25068 | 2 | 1133520.75 | 1184054 | 1655727.88 | 260520.16 | 1443850.25 | 1726312.75 | 1274475.62 | 1625874.38 | 1384813 | 1865663.25 | 5 | **1.387** |
| Q9CZH3 | Psmg3 | Proteasome assembly chaperone 3 (Mus musculus) | 154.43 | 42.62 | 13331 | 3 | 4598740 | 4345437 | 4264644 | 858858.56 | 1985079.5 | 4120903.25 | 4825888 | 4747701.5 | 4563190 | 4739398 | 4 | **1.433** |
| Q9CZN7 | Shmt2 | Serine hydroxymethyltransferase, mitochondrial (Mus musculus) | 1257.81 | 52.58 | 55759 | 21 | 125134192 | 90537816 | 84856464 | 80625984 | 89011984 | 112366080 | 97797808 | 96169424 | 101347488 | 101000816 | 5 | **1.082** |
| Q9CZR8 | Tsfm | Elongation factor Ts, mitochondrial (Mus musculus) | 289.3 | 22.84 | 35334 | 4 | 1688526.25 | 1354263.25 | 3101802.75 | 2505798.25 | 1507014.88 | 2602099.75 | 2564861 | 2685007 | 2474380.25 | 3428523.75 | 5 | **1.354** |
| Q9CZT5 | Vasn | Vasorin (Mus musculus) | 45.61 | 1.04 | 72261 | 1 | 2858921 | 1120527.12 | 2464469.25 | 987255.88 | 616053.38 | 4330316.5 | 3641829.25 | 3293795.75 | 3180477.5 | 344836.88 | 5 | **1.838** |
| Q9CZU3 | Mtrex | Exosome RNA helicase MTR4 (Mus musculus) | 568.6 | 15.38 | 117636 | 11 | 6828429 | 8022319 | 9188359 | 8636967 | 6923829.5 | 7606824.5 | 8758524 | 9527950 | 9618344 | 8697657 | 5 | **1.116** |
| Q9CZU6 | Cs | Citrate synthase, mitochondrial (Mus musculus) | 621.85 | 24.57 | 51737 | 11 | 89370168 | 54291752 | 59740464 | 52423896 | 58466644 | 79175080 | 84516472 | 85340912 | 83059608 | 90327488 | 4 | **1.344** |
| Q9CZW4 | Acsl3 | Long-chain-fatty-acid--CoA ligase 3 (Mus musculus) | 356.95 | 15.56 | 80492 | 8 | 83003192 | 29116630 | 28337008 | 32654952 | 37937344 | 36716600 | 38713868 | 41454628 | 32731964 | 39923880 | 7 | **0.898** |
| Q9CZW5 | Tomm70 | Mitochondrial import receptor subunit TOM70 (Mus musculus) | 409.63 | 13.58 | 67590 | 8 | 39135660 | 11471456 | 14471948 | 10896938 | 11313010 | 14143182 | 16306971 | 18645468 | 20341480 | 20437232 | 6 | **1.030** |
| Q9CZX0 | Elp3 | Elongator complex protein 3 (Mus musculus) | 167.51 | 7.86 | 62385 | 3 | 1651721 | 1839202.5 | 1914725.5 | 1887541.25 | 2103118.5 | 2380261.5 | 1696412.75 | 2077409.88 | 1942415 | 2065128.62 | 7 | **1.081** |
| Q9CZY3 | Ube2v1 | Ubiquitin-conjugating enzyme E2 variant 1 (Mus musculus) | 314.78 | 29.93 | 16355 | 5 | 16063625 | 14739749 | 15228930 | 15767990 | 10339162 | 14037532 | 16345233 | 12695826 | 8638682 | 9233414 | 7 | **0.845** |
| Q9D051 | Pdhb | Pyruvate dehydrogenase E1 component subunit beta, mitochondrial (Mus musculus) | 445.3 | 25.91 | 38937 | 7 | 14621108 | 11256998 | 10815982 | 5362771 | 8875350 | 11075552 | 13257043 | 13755598 | 15934298 | 11969607 | 5 | **1.296** |
| Q9D071 | Mms19 | MMS19 nucleotide excision repair protein homolog (Mus musculus) | 272.94 | 7.95 | 113089 | 5 | 1650653.5 | 4095054 | 3507063.5 | 3154009 | 2429273 | 2278036.75 | 3031979.75 | 3867409.5 | 2827703.5 | 3486585.75 | 12 | **1.044** |
| Q9D0B6 | Pbdc1 | Protein PBDC1 (Mus musculus) | 321.41 | 34.34 | 22223 | 5 | 6003040 | 8773359 | 7584549 | 10417232 | 8938170 | 8616212 | 9920527 | 8041557 | 10512320 | 10512782 | 7 | **1.141** |
| Q9D0E1 | Hnrnpm | Heterogeneous nuclear ribonucleoprotein M (Mus musculus) | 2051.8 | 53.5 | 77649 | 34 | 156367936 | 173828880 | 163489168 | 193025904 | 187974432 | 140570944 | 148661680 | 152429200 | 155468336 | 158675584 | 1 | **0.864** |
| Q9D0F3 | Lman1 | Protein ERGIC-53 (Mus musculus) | 322.81 | 17.02 | 57789 | 5 | 11734565 | 11346201 | 9644640 | 8895177 | 7760906 | 13058697 | 11397721 | 12183071 | 11567285 | 9662642 | 4 | **1.172** |
| Q9D0F6 | Rfc5 | Replication factor C subunit 5 (Mus musculus) | 187.29 | 15.63 | 38096 | 4 | 4361422.5 | 3706438.75 | 3656631 | 3451797.5 | 2940182.75 | 6000710 | 4754865.5 | 3573185.25 | 4972895 | 3934130.5 | 4 | **1.283** |
| Q9D0F9 | Pgm1 | Phosphoglucomutase-1 (Mus musculus) | 1295.99 | 45.91 | 61418 | 22 | 28949478 | 38391368 | 56085016 | 36086804 | 44855464 | 42156308 | 49251948 | 42583168 | 47192556 | 50666532 | 7 | **1.134** |
| Q9D0G0 | Mrps30 | 28S ribosomal protein S30, mitochondrial (Mus musculus) | 172.3 | 13.57 | 49939 | 3 | 471290.5 | 652426.25 | 542749.75 | 933014.25 | 537391.88 | 329323.78 | 288304.38 | 901084.5 | 953447.25 | 709460.06 | 12 | **1.014** |
| Q9D0I8 | Mrto4 | mRNA turnover protein 4 homolog (Mus musculus) | 349.62 | 28.03 | 27546 | 7 | 15341257 | 13605968 | 15906383 | 13562809 | 12096718 | 14508187 | 16050313 | 15323617 | 15634754 | 15401710 | 6 | **1.091** |
| Q9D0I9 | Rars | Arginine--tRNA ligase, cytoplasmic (Mus musculus) | 1682.38 | 47.73 | 75674 | 28 | 98783128 | 107241912 | 106862320 | 99917120 | 94590288 | 99763312 | 100268352 | 108089096 | 95841736 | 109589336 | 9 | **1.012** |
| Q9D0K2 | Oxct1 | Succinyl-CoA:3-ketoacid coenzyme A transferase 1, mitochondrial (Mus musculus) | 1323.33 | 51.54 | 55989 | 20 | 47288076 | 48169672 | 59467952 | 38807556 | 42972856 | 52996100 | 56716632 | 53169700 | 54239216 | 57630616 | 5 | **1.161** |
| Q9D0M1 | Prpsap1 | Phosphoribosyl pyrophosphate synthase-associated protein 1 (Mus musculus) | 489.72 | 34.83 | 39432 | 10 | 11337061 | 12313177 | 11485010 | 12160554 | 11677477 | 10560452 | 9846667 | 12159180 | 13665312 | 13732036 | 12 | **1.017** |
| Q9D0M3 | Cyc1 | Cytochrome c1, heme protein, mitochondrial (Mus musculus) | 341.28 | 41.85 | 35328 | 7 | 11914544 | 18260688 | 12857846 | 15524271 | 8189644.5 | 14291036 | 24425578 | 15872434 | 16873184 | 15802474 | 5 | **1.307** |
| Q9D0Q7 | Mrpl45 | 39S ribosomal protein L45, mitochondrial (Mus musculus) | 65.55 | 5.23 | 35411 | 1 | 731558.56 | 214585.81 | 739424.38 | 656115.06 | 755645.88 | 404197.78 | 425049.38 | 848156.56 | 959022.25 | 645175.56 | 12 | **1.059** |
| Q9D0R2 | Tars | Threonine--tRNA ligase, cytoplasmic (Mus musculus) | 1489.17 | 42.38 | 83356 | 27 | 59244424 | 72189960 | 63363428 | 61666044 | 63977788 | 77910656 | 61682312 | 68207928 | 68983200 | 67704640 | 6 | **1.075** |
| Q9D0R4 | Ddx56 | Probable ATP-dependent RNA helicase DDX56 (Mus musculus) | 72.86 | 3.66 | 61212 | 1 | 236257.73 | 500954.5 | 263478.44 | 0 | 0 | 425460.78 | 408318.34 | 439002.62 | 337709.56 | 349187.56 | 6 | **1.958** |
| Q9D0R8 | Lsm12 | Protein LSM12 homolog (Mus musculus) | 55.15 | 6.67 | 21701 | 1 | 1366956.75 | 1714730.12 | 1666196.5 | 1659093.88 | 1534665.5 | 1286731.25 | 1373223.75 | 1688798.12 | 1573520.25 | 1511039.5 | 8 | **0.936** |
| Q9D172 | Gatd3a | Glutamine amidotransferase-like class 1 domain-containing protein 3A, mitochondrial (Mus | 129.38 | 18.8 | 28090 | 3 | 1257440.75 | 2278378.75 | 2147417 | 927408.69 | 2561618 | 3042086 | 1675360.25 | 1232327.62 | 2417809.5 | 2320947 | 9 | **1.165** |
| Q9D1A2 | Cndp2 | Cytosolic non-specific dipeptidase (Mus musculus) | 1553.46 | 64.84 | 52767 | 25 | 40843788 | 32846842 | 29729168 | 32993232 | 40777376 | 47492036 | 43132420 | 41806476 | 42391748 | 43153800 | 0 | **1.230** |
| Q9D1C1 | Ube2c | Ubiquitin-conjugating enzyme E2 C (Mus musculus) | 66.73 | 9.5 | 19606 | 1 | 1051253.88 | 1300491.38 | 432174.62 | 989820.81 | 1439072.38 | 918737.94 | 1115728.25 | 944538.06 | 979375.44 | 737894.5 | 7 | **0.901** |
| Q9D1E6 | Tbcb | Tubulin-folding cofactor B (Mus musculus) | 264.08 | 27.05 | 27386 | 5 | 3793388.75 | 1377305 | 2189447.25 | 0 | 325699.19 | 2781780.5 | 2741609.25 | 2596368 | 3456293.5 | 2172016 | 6 | **1.789** |
| Q9D1G2 | Pmvk | Phosphomevalonate kinase (Mus musculus) | 67.06 | 9.9 | 21916 | 1 | 436507.91 | 71282.36 | 0 | 237526.12 | 467199.53 | 488700.41 | 517456.88 | 490638.41 | 575328.88 | 486799.53 | 0 | **2.110** |
| Q9D1H7 | Get4 | Golgi to ER traffic protein 4 homolog (Mus musculus) | 114.91 | 8.56 | 36525 | 2 | 312966.78 | 712722.69 | 541378.69 | 1114606.5 | 877480.94 | 851575.12 | 0 | 649748.88 | 906260.81 | 867255.81 | 12 | **0.920** |
| Q9D1J1 | Necap2 | Adaptin ear-binding coat-associated protein 2 (Mus musculus) | 282.59 | 31.95 | 28598 | 5 | 1695925.62 | 2610268.25 | 3547412 | 3171858.75 | 2097561.5 | 2743244.75 | 2465619.5 | 4023249.5 | 3522896 | 4057936.5 | 6 | **1.281** |
| Q9D1L0 | Chchd2 | Coiled-coil-helix-coiled-coil-helix domain-containing protein 2 (Mus musculus) | 97.4 | 22.22 | 15661 | 2 | 3872163.5 | 1594808.25 | 1056036 | 1359696.62 | 2681977.25 | 1418046.88 | 1341644.5 | 1092113 | 1146327.38 | 1122849.12 | 6 | **0.579** |
| Q9D1M4 | Eef1e1 | Eukaryotic translation elongation factor 1 epsilon-1 (Mus musculus) | 51.14 | 5.75 | 19859 | 1 | 1161065.62 | 1019326.56 | 0 | 980937.31 | 1711013.25 | 0 | 780155.75 | 939638.12 | 1144107.88 | 837218.44 | 7 | **0.760** |
| Q9D1N9 | Mrpl21 | 39S ribosomal protein L21, mitochondrial (Mus musculus) | 72.08 | 8.61 | 23366 | 1 | 0 | 0 | 0 | 510788.09 | 394105.06 | 0 | 0 | 594943.12 | 591399.31 | 369601.25 | 15 | **1.719** |
| Q9D1Q6 | Erp44 | Endoplasmic reticulum resident protein 44 (Mus musculus) | 43.71 | 5.67 | 46853 | 1 | 637296.31 | 0 | 0 | 0 | 0 | 751841.62 | 449379.31 | 546566.69 | 735348.69 | 400846.03 | 9 | **4.525** |
| Q9D1R9 | Rpl34 | 60S ribosomal protein L34 (Mus musculus) | 226.42 | 29.06 | 13293 | 5 | 35191500 | 16088982 | 25409756 | 18719594 | 19159812 | 56500032 | 36953464 | 46456224 | 38945148 | 40233272 | 0 | **1.912** |
| Q9D270 | Zdhhc21 | Probable palmitoyltransferase ZDHHC21 (Mus musculus) | 41.04 | 3.02 | 31331 | 1 | 370004.12 | 512610.25 | 336071.09 | 438490.31 | 0 | 485099.31 | 478242.97 | 435913.09 | 595235.88 | 412749.22 | 6 | **1.453** |
| Q9D289 | Trappc6b | Trafficking protein particle complex subunit 6B (Mus musculus) b | 41.78 | 12.03 | 17936 | 1 | 285709.44 | 540391.62 | 225710.94 | 154459.3 | 446839.38 | 364179.75 | 359027.22 | 440237.34 | 518765.53 | 384215.81 | 9 | **1.250** |
| Q9D2C2 | Saal1 | Protein SAAL1 (Mus musculus) | 66.08 | 3.59 | 52769 | 1 | 532328.81 | 384401.59 | 236828.11 | 0 | 0 | 421480.09 | 515300.56 | 0 | 180263.34 | 121474.27 | 14 | **1.074** |
| Q9D2E2 | Toe1 | Target of EGR1 protein 1 (Mus musculus) | 54.63 | 2.54 | 56871 | 1 | 0 | 0 | 37556.82 | 0 | 600686.5 | 0 | 651412.12 | 120918.13 | 111361.49 | 175394.05 | 11 | **1.659** |
| Q9D2L9 | Fam111a | Protein FAM111A (Mus musculus) | 38.55 | 2.61 | 69949 | 1 | 280033.34 | 174074.33 | 156179.53 | 244432.25 | 234217.56 | 316555.47 | 314728.28 | 254261.78 | 417277.81 | 224571.89 | 4 | **1.403** |
| Q9D2M8 | Ube2v2 | Ubiquitin-conjugating enzyme E2 variant 2 (Mus musculus) | 300.84 | 30.34 | 16367 | 5 | 13601286 | 12108126 | 11845814 | 13915817 | 7557853 | 11742678 | 13305932 | 9787882 | 5968530 | 5425380 | 5 | **0.783** |
| Q9D2R0 | Aacs | Acetoacetyl-CoA synthetase (Mus musculus) | 38.55 | 1.49 | 75200 | 1 | 1814872 | 1758693.12 | 1213616.88 | 2165056 | 0 | 1320520.25 | 1314083.38 | 1116717.88 | 918974.44 | 916874.31 | 7 | **0.804** |
| Q9D2V7 | Coro7 | Coronin-7 (Mus musculus) | 440.81 | 21.91 | 100812 | 10 | 14642482 | 12090700 | 11306987 | 11932313 | 10469210 | 13501460 | 11726728 | 12892554 | 11489810 | 11082028 | 12 | **1.004** |
| Q9D338 | Mrpl19 | 39S ribosomal protein L19, mitochondrial (Mus musculus) | 93.81 | 7.53 | 33578 | 2 | 2929650.25 | 3157912.25 | 3209930.25 | 2853142.25 | 3356790 | 2912061 | 0 | 3317402 | 3067125.5 | 3101953.5 | 9 | **0.800** |
| Q9D3D9 | Atp5f1d | ATP synthase subunit delta, mitochondrial (Mus musculus) | 216.87 | 37.5 | 17600 | 4 | 25009322 | 24277390 | 25162350 | 27635934 | 22348664 | 24779072 | 22454126 | 24860680 | 26612074 | 28096320 | 11 | **1.019** |
| Q9D3E6 | Stag1 | Cohesin subunit SA-1 (Mus musculus) | 68.12 | 1.83 | 144440 | 2 | 731476.75 | 386399.12 | 381003.03 | 483243.47 | 322871.62 | 75133.74 | 83592.81 | 347771.53 | 441631.03 | 470428.72 | 7 | **0.615** |
| Q9D554 | Sf3a3 | Splicing factor 3A subunit 3 (Mus musculus) | 494.17 | 24.35 | 58842 | 9 | 7921522 | 11782683 | 10806888 | 13947638 | 6981294.5 | 8020571 | 13189979 | 8967293 | 12141296 | 11661746 | 10 | **1.049** |
| Q9D5V6 | Syap1 | Synapse-associated protein 1 (Mus musculus) | 96.82 | 6.58 | 41350 | 2 | 1238118.88 | 1314318.75 | 2067040.25 | 1542715.5 | 652299.44 | 2490666.5 | 2507211.25 | 841101.81 | 1527281.25 | 1348335.25 | 8 | **1.279** |
| Q9D662 | Sec23b | Protein transport protein Sec23B (Mus musculus) | 460.52 | 12.78 | 86437 | 7 | 7462519 | 9784504 | 11007727 | 7153493 | 7540811 | 9833328 | 10382845 | 11649153 | 9938915 | 8952665 | 5 | **1.182** |
| Q9D6F9 | Tubb4a | Tubulin beta-4A chain (Mus musculus) | 1759.53 | 64.64 | 49586 | 27 | 648486912 | 619394432 | 526855136 | 634318592 | 570193216 | 632390272 | 605804800 | 639961728 | 601063744 | 573519104 | 12 | **1.018** |
| Q9D6J1 | Cers4 | Ceramide synthase 4 (Mus musculus) | 42.45 | 4.58 | 46017 | 1 | 40538.86 | 40802.6 | 189926.02 | 78147.7 | 104320.41 | 0 | 0 | 0 | 139877.31 | 181840.73 | 11 | **0.709** |
| Q9D6J6 | Ndufv2 | NADH dehydrogenase [ubiquinone] flavoprotein 2, mitochondrial (Mus musculus) | 42.17 | 4.03 | 27285 | 1 | 31217.55 | 558904.19 | 537795 | 426905.88 | 347711.72 | 721191.5 | 617131.69 | 1074035.25 | 90468.4 | 715493.56 | 4 | **1.692** |
| Q9D6K7 | Ttc33 | Tetratricopeptide repeat protein 33 (Mus musculus) | 40.52 | 4.96 | 29371 | 1 | 763382 | 1148161.38 | 1290168.75 | 0 | 1323743.62 | 1288804.62 | 0 | 1228398.12 | 1433931.38 | 1407304.5 | 9 | **1.184** |
| Q9D6K8 | Fundc2 | FUN14 domain-containing protein 2 (Mus musculus) | 101.49 | 14.57 | 16564 | 2 | 895954.5 | 1014404.25 | 812565.19 | 1421926.75 | 515776.97 | 602607.31 | 1007207.62 | 1197223 | 891957.69 | 1387298.5 | 11 | **1.091** |
| Q9D6K9 | Cers5 | Ceramide synthase 5 (Mus musculus) | 54.07 | 4.35 | 48167 | 1 | 0 | 0 | 0 | 84805.97 | 175444.44 | 0 | 0 | 244343.75 | 212986.14 | 191152.19 | 13 | **2.492** |
| Q9D6N1 | Ca13 | Carbonic anhydrase 13 (Mus musculus) | 49.1 | 3.44 | 29522 | 1 | 2160063.5 | 0 | 0 | 3477349.25 | 3028591.5 | 2616628 | 2739253 | 2639423.75 | 2905776.75 | 2996871.5 | 11 | **1.604** |
| Q9D6R2 | Idh3a | Isocitrate dehydrogenase [NAD] subunit alpha, mitochondrial (Mus musculus) | 787.3 | 41.53 | 39639 | 15 | 48030756 | 43530060 | 47437704 | 40414020 | 46436408 | 53273396 | 47348536 | 49000648 | 49057168 | 49279848 | 2 | **1.098** |
| Q9D6T0 | Nosip | Nitric oxide synthase-interacting protein (Mus musculus) | 118.89 | 11.96 | 33209 | 2 | 964943.94 | 562685 | 1119474.88 | 449635 | 763310.94 | 409155.28 | 558385.62 | 780158.06 | 770902.75 | 832216.56 | 10 | **0.868** |
| Q9D6Y7 | Msra | Mitochondrial peptide methionine sulfoxide reductase (Mus musculus) | 56.57 | 6.87 | 25988 | 1 | 6816524.5 | 741263.12 | 677653.19 | 570734.06 | 536954.19 | 396724.41 | 661232.94 | 616420.25 | 447592.38 | 621770 | 6 | **0.294** |
| Q9D6Y9 | Gbe1 | 1,4-alpha-glucan-branching enzyme (Mus musculus) | 155.68 | 6.55 | 80364 | 3 | 1215485.88 | 1547712 | 1817326.5 | 837949.38 | 1928767.5 | 1118619.62 | 1155042.88 | 1816791.62 | 1379266.12 | 1682656.88 | 10 | **0.973** |
| Q9D6Z1 | Nop56 | Nucleolar protein 56 (Mus musculus) | 1505.61 | 58.45 | 64464 | 24 | 59561168 | 43700244 | 54060680 | 45817804 | 41057000 | 59343344 | 60509532 | 60481156 | 56625644 | 55788720 | 3 | **1.199** |
| Q9D753 | Exosc8 | Exosome complex component RRP43 (Mus musculus) | 178.77 | 21.38 | 29949 | 4 | 5575221.5 | 3513799.5 | 5074184 | 5977970 | 3217464.5 | 3291953 | 4170644.5 | 4786561.5 | 5096619.5 | 4878510.5 | 10 | **0.951** |
| Q9D771 | Tmem206 | Transmembrane protein 206 (Mus musculus) | 80.85 | 7.43 | 40178 | 2 | 1958589.88 | 2466115.25 | 2904188 | 1039178.81 | 1580072.75 | 2440233 | 3298134 | 2112876.75 | 1606403.5 | 1770295.88 | 10 | **1.129** |
| Q9D7A6 | Srp19 | Signal recognition particle 19 kDa protein (Mus musculus) | 67.6 | 17.36 | 16191 | 1 | 713358.25 | 588844.81 | 545800.56 | 468795.19 | 599639.12 | 395505.31 | 313037.28 | 529058.56 | 380234.25 | 440326.66 | 1 | **0.706** |
| Q9D7H3 | RtcA | RNA 3'-terminal phosphate cyclase (Mus musculus) | 146.71 | 10.11 | 39254 | 3 | 4448603 | 4830908 | 4914782 | 4207992 | 4617650 | 5133808 | 4493307 | 5375398.5 | 5857653.5 | 6019043.5 | 3 | **1.168** |
| Q9D7J4 | Cox20 | Cytochrome c oxidase assembly protein COX20, mitochondrial (Mus musculus) | 71.45 | 19.66 | 13163 | 1 | 607706.69 | 349559.5 | 354359.62 | 237257.2 | 266005.03 | 456446.16 | 577108.19 | 73346.64 | 55231.49 | 296305.28 | 10 | **0.804** |
| Q9D7N3 | Mrps9 | 28S ribosomal protein S9, mitochondrial (Mus musculus) | 51.89 | 3.33 | 44929 | 1 | 0 | 0 | 1022099.31 | 0 | 0 | 1060693.38 | 794561.38 | 835198.81 | 674837.69 | 409941.62 | 10 | **3.694** |

| | | | | | | | | | | | | | | | | | |
|---|---|---|---|---|---|---|---|---|---|---|---|---|---|---|---|---|---|
| Q9D7N9 | Apmap | Adipocyte plasma membrane-associated protein (Mus musculus) | 193.43 | 18.07 | 46434 | 3 | 1134530.5 | 1663021.25 | 1276361.75 | 1619905 | 779337.75 | 1323693.75 | 1187480.25 | 1230677.88 | 1548128.5 | 1167521.25 | 12 | **0.998** |
| Q9D7S7 | Rpl22l1 | 60S ribosomal protein L22-like 1 (Mus musculus) | 313.79 | 45.08 | 14467 | 4 | 31242984 | 30091402 | 38768304 | 31085326 | 31636140 | 43451032 | 41273304 | 43880660 | 42499220 | 44173108 | 0 | **1.322** |
| Q9D7V9 | Naaa | N-acylethanolamine-hydrolyzing acid amidase (Mus musculus) | 44.27 | 4.7 | 40075 | 1 | 469937.66 | 59616.38 | 305818.12 | 0 | 126036.9 | 240029.39 | 331341.44 | 76503.88 | 0 | 0 | 12 | **0.674** |
| Q9D7X3 | Dusp3 | Dual specificity protein phosphatase 3 (Mus musculus) | 397.58 | 49.73 | 20472 | 5 | 18493048 | 14128720 | 12806150 | 12087806 | 12623356 | 15909867 | 18858684 | 16667017 | 16146473 | 16781450 | 4 | **1.203** |
| Q9D819 | Ppa1 | Inorganic pyrophosphatase (Mus musculus) | 659.83 | 32.18 | 32667 | 9 | 19088324 | 20587636 | 20453538 | 19195740 | 21358206 | 15049428 | 15508187 | 20618826 | 23894350 | 22609932 | 11 | **0.970** |
| Q9D853 | Eef1akmt2 | EEF1A lysine methyltransferase 2 (Mus musculus) t2 | 38.51 | 3.69 | 26864 | 1 | 0 | 831658.75 | 1186659.25 | 980168.19 | 1123959.25 | 721996.44 | 710231.5 | 704566 | 1116358.75 | 1044190.38 | 9 | **1.042** |
| Q9D868 | Ppih | Peptidyl-prolyl cis-trans isomerase H (Mus musculus) | 37.26 | 4.26 | 20464 | 1 | 686331.94 | 1427586.25 | 1254591.5 | 1183384.38 | 1200602.12 | 0 | 1011622.19 | 1534362.88 | 1270835.88 | 1448512.12 | 10 | **0.915** |
| Q9D880 | Timm50 | Mitochondrial import inner membrane translocase subunit TIM50 (Mus musculus) | 454.72 | 26.35 | 39776 | 7 | 16861088 | 20350960 | 14845125 | 17987738 | 19877348 | 19217256 | 18425240 | 17938920 | 20776600 | 24681550 | 7 | **1.124** |
| Q9D883 | U2af1 | Splicing factor U2AF 35 kDa subunit (Mus musculus) | 315.43 | 25.1 | 27815 | 5 | 15764134 | 15602804 | 15586025 | 15969072 | 14838639 | 15285043 | 20705796 | 15690777 | 17667300 | 16757701 | 6 | **1.107** |
| Q9D892 | Itpa | Inosine triphosphate pyrophosphatase (Mus musculus) | 78.12 | 9.09 | 21897 | 2 | 2494576.5 | 1057719.75 | 990727.31 | 1256858.12 | 1546713.62 | 1394043.75 | 1858093.5 | 1441677.75 | 1598598.12 | 1586772.38 | 7 | **1.072** |
| Q9D8B3 | Chmp4b | Charged multivesicular body protein 4b (Mus musculus) | 125.25 | 10.71 | 24936 | 2 | 4662462 | 610795.31 | 0 | 830408.62 | 1727357.25 | 774001.19 | 2567879.25 | 870310.06 | 1367313.5 | 1174911.38 | 10 | **0.863** |
| Q9D8B4 | Ndufa11 | NADH dehydrogenase [ubiquinone] 1 alpha subcomplex subunit 11 (Mus musculus) | 52.22 | 7.8 | 14982 | 1 | 0 | 1699231.75 | 1460835.5 | 1350821 | 1100844.25 | 1639156.5 | 1314720.38 | 1893906.38 | 1582968.38 | 1819322 | 5 | **1.470** |
| Q9D8E6 | Rpl4 | 60S ribosomal protein L4 (Mus musculus) | 1383.84 | 47.97 | 47154 | 22 | 332671488 | 191079552 | 296308064 | 144261248 | 289881856 | 364349600 | 369732064 | 377829760 | 357688288 | 361410592 | 0 | **1.460** |
| Q9D8M4 | Rpl7l1 | 60S ribosomal protein L7-like 1 (Mus musculus) | 207.9 | 24.39 | 28544 | 4 | 3161741.75 | 2893555 | 3827651.25 | 3668494 | 4051142.75 | 4428128 | 2515422.25 | 4934208 | 4873312 | 4982404.5 | 5 | **1.235** |
| Q9D8N0 | Eef1g | Elongation factor 1-gamma (Mus musculus) | 1737.8 | 82.15 | 50061 | 31 | 335517728 | 346887680 | 388007424 | 342977056 | 328014464 | 312971488 | 333311648 | 369226208 | 358208736 | 376914144 | 12 | **1.005** |
| Q9D8N2 | Fam45a | Protein FAM45A (Mus musculus) | 72.67 | 5.04 | 40421 | 2 | 422185.06 | 576814.94 | 512883.75 | 875917 | 673990.44 | 470309.19 | 462897.47 | 574120.62 | 593536.31 | 653663 | 10 | **0.900** |
| Q9D8T2 | Gsdmdc1 | Gasdermin-D (Mus musculus) | 249.19 | 15.4 | 53238 | 5 | 12066900 | 14135467 | 14673462 | 16084483 | 14122643 | 11428936 | 12897154 | 12613025 | 12490078 | 13319656 | 4 | **0.883** |
| Q9D8U8 | Snx5 | Sorting nexin-5 (Mus musculus) | 741.95 | 38.86 | 46797 | 15 | 43740996 | 59356700 | 60910836 | 61365920 | 56343432 | 58403192 | 54793940 | 52695756 | 55791996 | 56632708 | 7 | **0.988** |
| Q9D8V0 | Hm13 | Minor histocompatibility antigen H13 (Mus musculus) | 237.72 | 10.05 | 41748 | 5 | 8133464.5 | 10022128 | 9351380 | 11985259 | 7344616.5 | 9009909 | 10130079 | 8628366 | 10536787 | 10860822 | 9 | **1.050** |
| Q9D8W5 | Psmd12 | 26S proteasome non-ATPase regulatory subunit 12 (Mus musculus) | 1213.36 | 55.92 | 52895 | 22 | 45609652 | 73114512 | 68184752 | 70332296 | 59295060 | 49643372 | 52906068 | 53957300 | 52527744 | 55767848 | 5 | **0.837** |
| Q9D8Y0 | Efhd2 | EF-hand domain-containing protein D2 (Mus musculus) | 984.37 | 70.42 | 26791 | 17 | 109714944 | 126828952 | 122015312 | 134026872 | 118941256 | 125063648 | 94328832 | 120332264 | 129340080 | 121793456 | 11 | **0.966** |
| Q9D8Y7 | Tnfaip8l2 | Tumor necrosis factor alpha-induced protein 8-like protein 2 (Mus musculus) l2 | 235.26 | 40.22 | 20615 | 4 | 2701229.75 | 2323800.5 | 3614726.25 | 1120284.38 | 1433721.5 | 3871081.5 | 4704362.5 | 5122376 | 4379149.5 | 4954088.5 | 0 | **2.057** |
| Q9D903 | Ebna1bp2 | Probable rRNA-processing protein EBP2 (Mus musculus) 2 | 68.34 | 3.92 | 34703 | 1 | 2489997.5 | 1000416.88 | 967692 | 830810.5 | 943327.38 | 1126479.5 | 1318667 | 1205494.38 | 1127161.38 | 1170864.62 | 5 | **0.954** |
| Q9D906 | Atg7 | Ubiquitin-like modifier-activating enzyme ATG7 (Mus musculus) | 811.36 | 22.64 | 77520 | 14 | 12381179 | 15649192 | 15572828 | 14830766 | 13162145 | 15380705 | 13400865 | 16566591 | 16670470 | 16451187 | 5 | **1.096** |
| Q9D938 | Tmem160 | Transmembrane protein 160 (Mus musculus) | 59.55 | 3.72 | 19587 | 1 | 1458697.75 | 1352966.88 | 1255509.88 | 1032782.38 | 1189620.25 | 1302788.12 | 1169517.12 | 1141031.75 | 1577316.75 | 1755456.12 | 10 | **1.104** |
| Q9D964 | Gatm | Glycine amidinotransferase, mitochondrial (Mus musculus) | 869.03 | 44.44 | 48297 | 16 | 37702348 | 52210332 | 54513468 | 63367180 | 48935376 | 45931368 | 40161936 | 46389184 | 45711512 | 51505496 | 6 | **0.895** |
| Q9D967 | Mdp1 | Magnesium-dependent phosphatase 1 (Mus musculus) | 64.63 | 10.98 | 18582 | 1 | 0 | 234925.81 | 341659.97 | 236514.89 | 396269.34 | 143982.23 | 160127.28 | 292686.25 | 217488.66 | 237728.27 | 9 | **0.870** |
| Q9D9V3 | Echdc1 | Ethylmalonyl-CoA decarboxylase (Mus musculus) | 92.76 | 14.91 | 35467 | 2 | 0 | 0 | 0 | 662790 | 0 | 325878.59 | 141796.77 | 191824.94 | 115070.01 | 172093.73 | 11 | **1.428** |
| Q9DAP7 | Asf1b | Histone chaperone ASF1B (Mus musculus) | 115.41 | 23.76 | 22464 | 2 | 405995.59 | 812325.44 | 858860.56 | 722633.62 | 890415.75 | 873188 | 556786.38 | 1032506.19 | 928495.38 | 842275.94 | 7 | **1.148** |
| Q9DAR7 | Dcps | m7GpppX diphosphatase (Mus musculus) | 590.92 | 39.05 | 38988 | 10 | 12339499 | 19395118 | 18663238 | 12000020 | 12895672 | 14880678 | 12550011 | 14838347 | 16035951 | 15283393 | 11 | **0.977** |
| Q9DAW6 | Prpf4 | U4/U6 small nuclear ribonucleoprotein Prp4 (Mus musculus) | 49.69 | 2.69 | 58370 | 1 | 0 | 889506.19 | 575706.25 | 0 | 705445.19 | 878151.69 | 366413.22 | 632648.88 | 642717.5 | 842800.06 | 10 | **1.549** |
| Q9DB20 | Atp5po | ATP synthase subunit O, mitochondrial (Mus musculus) | 350.21 | 33.8 | 23364 | 5 | 6065941.5 | 2447946.5 | 2177607.5 | 3788872.75 | 3080987.25 | 2977259.75 | 3988475 | 3573684.5 | 1974912 | 3313603.5 | 12 | **0.901** |
| Q9DB25 | Alg5 | Dolichyl-phosphate beta-glucosyltransferase (Mus musculus) | 126.73 | 10.49 | 36791 | 3 | 2293847.75 | 2941315 | 3068866.75 | 2946837.75 | 3170412 | 3163700.25 | 3445748.5 | 3780036 | 4290355.5 | 4225805.5 | 1 | **1.311** |
| Q9DB34 | Chmp2a | Charged multivesicular body protein 2a (Mus musculus) | 38.41 | 3.15 | 25134 | 1 | 702341 | 323327.5 | 359384.81 | 401221.62 | 211176.47 | 540832.38 | 677704.75 | 514252.31 | 529708.69 | 459861.31 | 5 | **1.363** |
| Q9DB43 | Zfpl1 | Zinc finger protein-like 1 (Mus musculus) | 50.87 | 3.87 | 34155 | 1 | 302786.84 | 0 | 321109.41 | 214290.67 | 0 | 401178.91 | 389919.62 | 308415.19 | 0 | 471661.38 | 7 | **1.874** |
| Q9DB73 | Cyb5r1 | NADH-cytochrome b5 reductase 1 (Mus musculus) | 43.23 | 8.85 | 34135 | 1 | 392932.38 | 0 | 358253.16 | 0 | 385629.53 | 380644.22 | 418748.06 | 601457.19 | 349038.44 | 427994.41 | 6 | **1.916** |
| Q9DB77 | Uqcrc2 | Cytochrome b-c1 complex subunit 2, mitochondrial (Mus musculus) | 1097.12 | 50.77 | 48235 | 18 | 58989688 | 47461280 | 55094100 | 48944728 | 46774816 | 64172832 | 61842268 | 63001944 | 60695300 | 64401856 | 0 | **1.221** |
| Q9DBB8 | Dhdh | Trans-1,2-dihydrobenzene-1,2-diol dehydrogenase (Mus musculus) | 49.75 | 3.6 | 36301 | 1 | 872698.81 | 675441.12 | 804595.12 | 292173.75 | 515564.5 | 732024.5 | 631903.31 | 640321.12 | 519842.34 | 721149.19 | 12 | **1.027** |
| Q9DBC7 | Prkar1a | cAMP-dependent protein kinase type I-alpha regulatory subunit (Mus musculus) | 538.65 | 29.13 | 43185 | 10 | 21150504 | 19031326 | 20981610 | 21430218 | 16478848 | 19718866 | 18775838 | 14288780 | 19433984 | 13164461 | 5 | **0.862** |
| Q9DBE9 | Ftsj3 | pre-rRNA processing protein FTSJ3 (Mus musculus) | 238.78 | 11.81 | 95532 | 5 | 2690970.25 | 605144.5 | 1356219.5 | 3279850.75 | 1217543.12 | 3760712.5 | 2239601 | 4041368.75 | 3458972.5 | 3698316 | 2 | **1.880** |
| Q9DBG1 | Cyp27a1 | Sterol 26-hydroxylase, mitochondrial (Mus musculus) | 64.42 | 4.5 | 60720 | 1 | 486007.94 | 587228.12 | 607891.44 | 639842.75 | 387037.16 | 514644.09 | 162121.08 | 571176.19 | 711711.25 | 650807.62 | 11 | **0.964** |
| Q9DBG5 | Plin3 | Perilipin-3 (Mus musculus) | 492.51 | 26.32 | 47262 | 8 | 19322244 | 18876352 | 22624796 | 19802590 | 22629274 | 16069559 | 20074150 | 17351656 | 17890412 | 17528700 | 3 | **0.861** |
| Q9DBG6 | Rpn2 | Dolichyl-diphosphooligosaccharide--protein glycosyltransferase subunit 2 (Mus musculus) | 868.21 | 29.48 | 69063 | 13 | 14386919 | 24510510 | 23339812 | 24862338 | 21441198 | 21704980 | 20007464 | 22248900 | 23752482 | 21498182 | 10 | **1.006** |
| Q9DBG7 | Srpra | Signal recognition particle receptor subunit alpha (Mus musculus) | 434.27 | 14.15 | 69623 | 7 | 9424932 | 10775670 | 11747683 | 6799212 | 6584732 | 10976633 | 10817764 | 11162297 | 7948228 | 10562410 | 8 | **1.135** |
| Q9DBH5 | Lman2 | Vesicular integral-membrane protein VIP36 (Mus musculus) | 137.7 | 12.01 | 40430 | 3 | 9897397 | 4150870 | 3551699.25 | 8723679 | 6722121.5 | 7118127 | 9356951 | 6508909.5 | 7398993.5 | 7495478.5 | 10 | **1.146** |
| Q9DBL1 | Acadsb | Short/branched chain specific acyl-CoA dehydrogenase, mitochondrial (Mus musculus) | 176.35 | 10.42 | 47874 | 3 | 1373654.88 | 1895461.5 | 4674146.5 | 3909757.5 | 4288589 | 2683624.25 | 1687095.5 | 3331434.25 | 4101676 | 4616018 | 12 | **1.017** |
| Q9DBL7 | Coasy | Bifunctional coenzyme A synthase (Mus musculus) | 200.59 | 11.55 | 62023 | 4 | 1670392.62 | 1889542.25 | 2207663 | 1754391.5 | 1556788.75 | 2064896.5 | 2161144.5 | 2355057.5 | 3129444.25 | 2170449.25 | 3 | **1.309** |
| Q9DBQ7 | Scyl3 | Protein-associating with the carboxyl-terminal domain of ezrin (Mus musculus) | 91.5 | 3.95 | 81333 | 2 | 1023612.38 | 1237203.88 | 883329.88 | 944190.75 | 1135637.25 | 1172632.75 | 0 | 1165567.12 | 1135899.5 | 973417.25 | 11 | **0.851** |
| Q9DBR0 | Akap8 | A-kinase anchor protein 8 (Mus musculus) | 50.39 | 1.89 | 76294 | 1 | 1314324.5 | 722969.81 | 844393.69 | 994426.88 | 1129766 | 1188251.62 | 1153852.5 | 1043073.56 | 1108232.62 | 1329784.25 | 6 | **1.163** |
| Q9DBR1 | Xrn2 | 5'-3' exoribonuclease 2 (Mus musculus) | 420.31 | 13.77 | 108687 | 8 | 9167107 | 11345977 | 8284278 | 13073392 | 9639055 | 10342709 | 11459150 | 12067224 | 10940559 | 8390328 | 10 | **1.033** |
| Q9DBS1 | Tmem43 | Transmembrane protein 43 (Mus musculus) | 573.85 | 36.5 | 44783 | 11 | 14442531 | 15168784 | 15548083 | 11202367 | 11085079 | 18130786 | 19455686 | 20175854 | 17130098 | 18406530 | 0 | **1.383** |
| Q9DBZ5 | Eif3k | Eukaryotic translation initiation factor 3 subunit K (Mus musculus) | 398.71 | 27.98 | 25087 | 5 | 8018207 | 11260811 | 9282338 | 6398295 | 4719562.5 | 8811993 | 8073010.5 | 6489230 | 6490981 | 6660977 | 12 | **0.921** |
| Q9DC16 | Ergic1 | Endoplasmic reticulum-Golgi intermediate compartment protein 1 (Mus musculus) | 174.21 | 13.1 | 32562 | 4 | 4933159 | 5063799 | 3857376 | 7082475 | 4934960 | 4945470.5 | 6170263.5 | 4878633 | 4256971.5 | 8051981.5 | 11 | **1.094** |
| Q9DC40 | Telo2 | Telomere length regulation protein TEL2 homolog (Mus musculus) | 181.6 | 6.19 | 93313 | 3 | 0 | 271496.47 | 364638.78 | 383889.62 | 232406.89 | 56652.96 | 67401.25 | 422246.41 | 136676.44 | 323068.56 | 11 | **0.803** |
| Q9DC69 | Ndufa9 | NADH dehydrogenase [ubiquinone] 1 alpha subcomplex subunit 9, mitochondrial (Mus mu | 500.72 | 33.16 | 42525 | 9 | 7923756 | 9273070 | 8397072 | 7347219.5 | 7697210.5 | 7459134.5 | 5048419.5 | 8840302 | 7331076 | 9246889 | 9 | **0.933** |
| Q9DC70 | Ndufs7 | NADH dehydrogenase [ubiquinone] iron-sulfur protein 7, mitochondrial (Mus musculus) | 186.11 | 14.29 | 24683 | 3 | 78768288 | 3138693.5 | 5093172 | 18101880 | 5862284 | 8516976 | 10290866 | 6762733 | 6472670 | 5816445 | 11 | **0.341** |
| Q9DCD0 | Pgd | 6-phosphogluconate dehydrogenase, decarboxylating (Mus musculus) | 2140.21 | 69.77 | 53247 | 32 | 371830880 | 329049184 | 375248256 | 285429504 | 354306656 | 413262272 | 352167744 | 420341504 | 412762816 | 408323360 | 3 | **1.170** |
| Q9DCE5 | Pak1ip1 | p21-activated protein kinase-interacting protein 1 (Mus musculus) | 119.78 | 5.76 | 42116 | 2 | 2279136.25 | 3177222.5 | 4961358 | 3744265 | 3390008 | 5231421.5 | 3575353.5 | 4600114.5 | 4885271 | 4538164.5 | 5 | **1.301** |
| Q9DCH4 | Eif3f | Eukaryotic translation initiation factor 3 subunit F (Mus musculus) | 841 | 38.78 | 37984 | 13 | 35091616 | 50888404 | 46468080 | 54302168 | 42681176 | 52168596 | 54072808 | 59514956 | 54370760 | 54638520 | 2 | **1.198** |
| Q9DCL9 | Paics | Multifunctional protein ADE2 (Mus musculus) | 885.55 | 44 | 47006 | 16 | 36054624 | 39495188 | 36479020 | 37901476 | 34699892 | 42703596 | 36781804 | 37574624 | 36957796 | 39965296 | 6 | **1.051** |
| Q9DCM0 | Ethe1 | Persulfide dioxygenase ETHE1, mitochondrial (Mus musculus) | 185.58 | 20.08 | 27739 | 4 | 9624646 | 5151898 | 4296389.5 | 7860841 | 4246804 | 6290347 | 4980713.5 | 5139157.5 | 5288472.5 | 5750932 | 12 | **0.880** |
| Q9DCM2 | Gstk1 | Glutathione S-transferase kappa 1 (Mus musculus) | 134.67 | 15.49 | 25704 | 3 | 3587680 | 2467348.75 | 3534005.25 | 2373629.5 | 1278452.5 | 2792947.5 | 1764256 | 3548294.75 | 2912006.75 | 2940335 | 11 | **1.054** |
| Q9DCN2 | Cyb5r3 | NADH-cytochrome b5 reductase 3 (Mus musculus) | 564.08 | 36.21 | 34128 | 9 | 23794934 | 10531544 | 9401964 | 14247532 | 15777402 | 21701068 | 20965370 | 19856730 | 18365352 | 16469244 | 5 | **1.320** |
| Q9DCS2 | Mettl26 | Methyltransferase-like 26 (Mus musculus) | 72.72 | 8.82 | 22687 | 1 | 302250.69 | 298982.59 | 239024.67 | 297736.09 | 318849.81 | 357829.06 | 370933.53 | 266176.44 | 286000.03 | 331822.72 | 8 | **1.107** |
| Q9DCS3 | Mecr | Enoyl-[acyl-carrier-protein] reductase, mitochondrial (Mus musculus) | 43.67 | 3.49 | 40343 | 1 | 587784.25 | 493615.19 | 646125.44 | 386972.47 | 0 | 702875.62 | 528645.81 | 234614.14 | 507284.75 | 436189.34 | 11 | **1.140** |
| Q9DCS9 | Ndufb10 | NADH dehydrogenase [ubiquinone] 1 beta subcomplex subunit 10 (Mus musculus) | 51.41 | 6.25 | 21024 | 1 | 601127.38 | 321315.59 | 786876.44 | 835747.56 | 180339.59 | 502205.31 | 617502.44 | 415219.72 | 481397.84 | 282574.97 | 10 | **0.844** |
| Q9DCT2 | Ndufs3 | NADH dehydrogenase [ubiquinone] iron-sulfur protein 3, mitochondrial (Mus musculus) | 318.25 | 31.94 | 30149 | 6 | 4015010 | 8326685 | 7707812.5 | 9598967 | 6913494 | 5276958.5 | 7426687.5 | 6942631.5 | 6014344 | 6758088 | 7 | **0.887** |
| Q9DCW4 | Etfb | Electron transfer flavoprotein subunit beta (Mus musculus) | 758.26 | 52.94 | 27623 | 13 | 62241552 | 68326624 | 59606936 | 73582432 | 60387740 | 60506344 | 54782708 | 61183272 | 59287108 | 61812596 | 6 | **0.918** |
| Q9DCX2 | Atp5pd | ATP synthase subunit d, mitochondrial (Mus musculus) | 203.13 | 36.65 | 18749 | 4 | 10225387 | 7535505.5 | 6398604.5 | 8008104 | 8875092 | 7046340.5 | 7131207.5 | 7456293.5 | 8360935.5 | 7590274.5 | 8 | **0.916** |
| Q9DCZ1 | Gmpr | GMP reductase 1 (Mus musculus) | 224.13 | 12.46 | 37482 | 4 | 2106583.75 | 1639273.12 | 1842785.12 | 1390299.12 | 1808494.25 | 1418572.12 | 1741124.12 | 1472970.5 | 6256448.5 | 1530639.75 | 10 | **1.413** |
| Q9DCZ4 | Apoo | MICOS complex subunit Mic26 (Mus musculus) | 44.09 | 5.05 | 22604 | 1 | 0 | 0 | 561788.31 | 580504.44 | 500750.81 | 637002.94 | 634637.88 | 721263.31 | 744884.81 | 812919.06 | 1 | **2.161** |
| Q9EP69 | Sacm1l | Phosphatidylinositide phosphatase SAC1 (Mus musculus) | 512.38 | 19.25 | 66944 | 10 | 10169969 | 8282677 | 9665329 | 8882212 | 7165364 | 11752325 | 8943954 | 11022274 | 11124661 | 9265069 | 4 | **1.180** |
| Q9EPB4 | Pycard | Apoptosis-associated speck-like protein containing a CARD (Mus musculus) | 610.75 | 68.91 | 21459 | 11 | 20393110 | 23985482 | 26789270 | 22357620 | 23756564 | 25033694 | 24137220 | 21108352 | 24921512 | 24519210 | 8 | **1.021** |
| Q9EPE9 | Atp13a1 | Endoplasmic reticulum transmembrane helix translocase (Mus musculus) | 185.02 | 3.5 | 132388 | 3 | 6913363 | 5236947 | 4076177 | 6906654 | 4501023 | 3281468.5 | 3510868 | 3414033 | 4683930 | 3850200.5 | 2 | **0.678** |
| Q9EPK2 | Rp2 | Protein XRP2 (Mus musculus) | 80.2 | 2.88 | 39377 | 1 | 652222.5 | 1009515.56 | 1226761.12 | 1028272.06 | 925020.44 | 747182.06 | 964944.19 | 1138749.75 | 1130216.62 | 1236820.12 | 9 | **1.078** |
| Q9EPL8 | Ipo7 | Importin-7 (Mus musculus) | 834.47 | 18.59 | 119486 | 15 | 7774594 | 15350954 | 18391328 | 23520212 | 10381935 | 13623747 | 12028838 | 11208154 | 11972342 | 12059622 | 10 | **0.807** |
| Q9EPU0 | Upf1 | Regulator of nonsense transcripts 1 (Mus musculus) | 387.73 | 8.63 | 123967 | 7 | 10247295 | 10831346 | 10855605 | 12115177 | 11377842 | 8056908 | 10044702 | 13577734 | 10960248 | 11850387 | 12 | **0.983** |
| Q9EQ06 | Hsd17b11 | Estradiol 17-beta-dehydrogenase 11 (Mus musculus) 1 | 44.29 | 5.7 | 32881 | 1 | 0 | 37541.6 | 318639.44 | 454423.88 | 513368 | 0 | 0 | 550126.69 | 508328.78 | 379955.16 | 13 | **1.086** |
| Q9EQ32 | Pik3ap1 | Phosphoinositide 3-kinase adapter protein 1 (Mus musculus) | 548.06 | 14.3 | 90928 | 8 | 7241367.5 | 12525120 | 13607058 | 13048404 | 12072537 | 11990475 | 8737115 | 10874585 | 11877935 | 12850023 | 7 | **0.963** |
| Q9EQ61 | Pes1 | Pescadillo homolog (Mus musculus) | 342.32 | 11.47 | 67796 | 7 | 6509522 | 3093606 | 3879265.5 | 2041339.12 | 2428558.5 | 4269622.5 | 5242289.5 | 5582012.5 | 6144280 | 5483363.5 | 5 | **1.488** |
| Q9EQ80 | Nif3l1 | NIF3-like protein 1 (Mus musculus) | 294.62 | 15.43 | 41746 | 6 | 5188247.5 | 6131996.5 | 6154424 | 5904863 | 5214755 | 6469102 | 6695526.5 | 6180281.5 | 5738354 | 5779727 | 6 | **1.079** |
| Q9EQH2 | Erap1 | Endoplasmic reticulum aminopeptidase 1 (Mus musculus) | 896.8 | 25.7 | 106599 | 18 | 22789078 | 23353684 | 23779472 | 21443120 | 21028840 | 25887184 | 23078916 | 24705628 | 25470128 | 27646392 | 2 | **1.128** |
| Q9EQH3 | Vps35 | Vacuolar protein sorting-associated protein 35 (Mus musculus) | 1415.93 | 35.05 | 91713 | 22 | 31842898 | 56039860 | 50947072 | 51482320 | 37177100 | 48615136 | 48816804 | 45422992 | 47360760 | 48901760 | 10 | **1.051** |
| Q9EQK5 | Mvp | Major vault protein (Mus musculus) | 1135.36 | 35.66 | 95924 | 19 | 46948372 | 52032216 | 47570180 | 44041864 | 43007288 | 48673032 | 49253808 | 45171792 | 45304988 | 45829108 | 11 | **1.003** |
| Q9EQP2 | Ehd4 | EH domain-containing protein 4 (Mus musculus) | 782.59 | 27.73 | 61481 | 15 | 28835550 | 27479528 | 28670274 | 31207252 | 20852498 | 24117324 | 27573882 | 28636236 | 30039374 | 27710460 | 11 | **1.008** |
| Q9EQW7 | Kif13a | Kinesin-like protein KIF13A (Mus musculus) | 42.1 | 0.74 | 195813 | 1 | 0 | 0 | 0 | 0 | 0 | 72148.31 | 0 | 97664.68 | 0 | 0 | 25 | **#DIV/0 !** |
| Q9ER38 | Tor3a | Torsin-3A (Mus musculus) | 181.68 | 12.21 | 43814 | 3 | 1259272.75 | 1017855.25 | 1284029.25 | 1420058 | 1540502.75 | 951526.25 | 1198189.12 | 966010.25 | 721861.88 | 1383800.62 | 4 | **0.801** |
| Q9ER41 | Tor1b | Torsin-1B (Mus musculus) | 163.19 | 8.33 | 37818 | 2 | 1986934.25 | 1162589.75 | 0 | 1027100 | 833861.38 | 2029131.25 | 1120383.25 | 1871769.25 | 1791272 | 1742512.75 | 5 | **1.707** |

| | | | | | | | | | | | | | | | | | |
|---|---|---|---|---|---|---|---|---|---|---|---|---|---|---|---|---|---|
| Q9ER72 | Cars | Cysteine--tRNA ligase, cytoplasmic (Mus musculus) | 1030.06 | 27.56 | 94860 | 19 | 20823504 | 26802330 | 30586790 | 30285074 | 30967974 | 34636464 | 29754456 | 34412352 | 35751748 | 36047388 | 3 | **1.223** |
| Q9ER88 | Dap3 | 28S ribosomal protein S29, mitochondrial (Mus musculus) | 347.04 | 24.3 | 44699 | 6 | 4423307.5 | 3992351 | 5483586 | 2791870 | 4234525 | 3023622.5 | 4439055 | 4209696 | 3816954.75 | 3764433.5 | 9 | **0.920** |
| Q9ERD8 | Parvg | Gamma-parvin (Mus musculus) | 95.47 | 6.34 | 37603 | 2 | 1084697.12 | 1172885.12 | 1880566.5 | 1342733.25 | 1400099.75 | 1057962.38 | 1534646.75 | 1243951.5 | 1352827.88 | 1258892.38 | 11 | **0.937** |
| Q9ERE7 | Mesd | LRP chaperone MESD (Mus musculus) | 80.56 | 10.71 | 25207 | 2 | 1620808.38 | 7503759 | 6774286 | 10635807 | 8872669 | 5323105 | 4775720 | 5395889 | 8019641 | 6883402 | 8 | **0.859** |
| Q9ERF3 | Wdr61 | WD repeat-containing protein 61 (Mus musculus) | 391.22 | 32.13 | 33773 | 5 | 2786819 | 5070363.5 | 7466751 | 3382885.5 | 6610166.5 | 6037577.5 | 4086499.75 | 7684086.5 | 4411290.5 | 7083357 | 9 | **1.157** |
| Q9ERG0 | Lima1 | LIM domain and actin-binding protein 1 (Mus musculus) | 123.72 | 5.84 | 84060 | 3 | 0 | 1556541.25 | 1632841.12 | 1300382 | 4574982.5 | 2269033 | 2136453.75 | 1650453.38 | 1393296.5 | 1873381.38 | 7 | **1.028** |
| Q9ERK4 | Cse1l | Exportin-2 (Mus musculus) | 1085.57 | 26.06 | 110455 | 18 | 14304702 | 22312918 | 22914538 | 30060616 | 18912684 | 19016568 | 14426441 | 19269086 | 21334176 | 19745362 | 9 | **0.864** |
| Q9ERL7 | Gmfg | Glia maturation factor gamma (Mus musculus) | 157.08 | 23.24 | 16748 | 3 | 17218546 | 10964300 | 9381190 | 11622567 | 12309890 | 12634224 | 11039356 | 13734539 | 12452133 | 12285443 | 8 | **1.011** |
| Q9ERN0 | Scamp2 | Secretory carrier-associated membrane protein 2 (Mus musculus) | 47.52 | 3.65 | 36465 | 1 | 0 | 0 | 828669.19 | 0 | 511518.28 | 0 | 540198.81 | 597210.81 | 568910.06 | 0 | 16 | **1.273** |
| Q9ERS2 | Ndufa13 | NADH dehydrogenase [ubiquinone] 1 alpha subcomplex subunit 13 (Mus musculus) | 200.89 | 21.53 | 16860 | 4 | 8439171 | 8389824 | 8602358 | 7429383.5 | 6822204.5 | 7977019.5 | 8004923 | 8064127.5 | 7464905 | 7981005 | 10 | **0.995** |
| Q9ERS5 | Plekha2 | Pleckstrin homology domain-containing family A member 2 (Mus musculus) | 53.67 | 3.76 | 47380 | 2 | 761026.69 | 0 | 0 | 588904.25 | 447199.72 | 535373.06 | 0 | 651484.88 | 487688.59 | 543167.5 | 13 | **1.234** |
| Q9ERU9 | Ranbp2 | E3 SUMO-protein ligase RanBP2 (Mus musculus) | 931.12 | 8.58 | 341121 | 20 | 17014032 | 14392849 | 16145400 | 12626104 | 14372915 | 15517058 | 15379884 | 16015011 | 18228068 | 19342486 | 6 | **1.133** |
| Q9ES00 | Ube4b | Ubiquitin conjugation factor E4 B (Mus musculus) | 186.55 | 4.94 | 133318 | 4 | 2302798.5 | 1391740.12 | 1855607.88 | 480393.5 | 986204.25 | 1930097.62 | 1562874 | 2485508.5 | 1614108.12 | 1292134 | 8 | **1.266** |
| Q9ES46 | Parvb | Beta-parvin (Mus musculus) | 152.71 | 13.15 | 41669 | 3 | 749182.19 | 295121.91 | 1323592.38 | 1281058.5 | 442912.12 | 736007.38 | 0 | 427804.25 | 581574.81 | 509669.72 | 7 | **0.551** |
| Q9ES52 | Inpp5d | Phosphatidylinositol 3,4,5-trisphosphate 5-phosphatase 1 (Mus musculus) | 1437.9 | 29.14 | 133542 | 27 | 58668100 | 55688120 | 56605980 | 53412548 | 52070144 | 59897512 | 55876992 | 55241676 | 56576692 | 57071564 | 8 | **1.030** |
| Q9ES97 | Rtn3 | Reticulon-3 (Mus musculus) | 235.09 | 7.37 | 103879 | 4 | 2511065.75 | 3925842.5 | 5371157 | 1302872.25 | 3123168.75 | 5337591.5 | 4939654 | 5960081.5 | 4556706 | 5791282 | 3 | **1.638** |
| Q9ESE1 | Lrba | Lipopolysaccharide-responsive and beige-like anchor protein (Mus musculus) | 98.84 | 0.63 | 317064 | 2 | 0 | 0 | 209011.27 | 363463.59 | 272739.28 | 345756.88 | 347246.75 | 409885.34 | 306261.5 | 269049.28 | 6 | **1.986** |
| Q9ESL4 | Map3k20 | Mitogen-activated protein kinase kinase kinase 20 (Mus musculus) | 192.92 | 5.86 | 91720 | 4 | 3003872.5 | 2728614.5 | 1507209.62 | 2559041 | 3806134.5 | 3471052.75 | 2535414.25 | 5115288.5 | 4700910 | 3303233.75 | 6 | **1.406** |
| Q9ESP1 | Sdf2l1 | Stromal cell-derived factor 2-like protein 1 (Mus musculus) | 131.44 | 13.12 | 23648 | 3 | 6324777.5 | 4002337 | 5742476 | 2979709 | 2993019.5 | 7431375 | 4754852.5 | 6237995.5 | 4669479.5 | 5445995 | 7 | **1.295** |
| Q9EST4 | Psmg2 | Proteasome assembly chaperone 2 (Mus musculus) | 82.22 | 7.2 | 29524 | 2 | 678928.75 | 0 | 448096.38 | 387501.72 | 454025.34 | 531180.69 | 676237.44 | 594410.69 | 507930.62 | 468686.47 | 5 | **1.411** |
| Q9EST5 | Anp32b | Acidic leucine-rich nuclear phosphoprotein 32 family member B (Mus musculus) | 298.85 | 21.69 | 31079 | 5 | 45578064 | 21919178 | 33570956 | 22376028 | 28791898 | 47434688 | 41525976 | 52367216 | 44056048 | 49637084 | 2 | **1.544** |
| Q9ESV0 | Ddx24 | ATP-dependent RNA helicase DDX24 (Mus musculus) | 107.75 | 4.08 | 96429 | 2 | 1310922.5 | 352245.59 | 333852.81 | 0 | 0 | 1551843 | 1427275.5 | 1256433.5 | 680680.88 | 1299967.5 | 4 | **3.113** |
| Q9ESW4 | Agk | Acylglycerol kinase, mitochondrial (Mus musculus) | 123.21 | 9.03 | 46976 | 2 | 1530097.5 | 2598619.5 | 1930064.88 | 2676398.25 | 3313504 | 2808777 | 1602420.25 | 2089660.75 | 2310053.5 | 1939909.5 | 11 | **0.892** |
| Q9ESX5 | Dkc1 | H/ACA ribonucleoprotein complex subunit DKC1 (Mus musculus) | 473.92 | 25.54 | 57402 | 9 | 9356584 | 9224773 | 8857899 | 6322551.5 | 6523830.5 | 8749040 | 10132898 | 9564367 | 9004378 | 7812159.5 | 8 | **1.124** |
| Q9ESY9 | Ifi30 | Gamma-interferon-inducible lysosomal thiol reductase (Mus musculus) | 108.44 | 10.89 | 27784 | 2 | 3720531 | 4729702.5 | 3744236.5 | 4117700.25 | 3847611 | 2717665 | 3203230.5 | 3196909 | 3672765.5 | 2868286 | 0 | **0.777** |
| Q9ET30 | Tm9sf3 | Transmembrane 9 superfamily member 3 (Mus musculus) | 284.73 | 12.95 | 67545 | 6 | 6335846.5 | 3625885 | 7569219 | 5553265.5 | 1544283.25 | 9203437 | 7731864 | 8603540 | 7734260 | 10971680 | 0 | **1.796** |
| Q9JHI5 | Ivd | Isovaleryl-CoA dehydrogenase, mitochondrial (Mus musculus) | 113.18 | 5.66 | 46325 | 2 | 4521884.5 | 3538388 | 2690661 | 4428663 | 3125442 | 2669389.5 | 2421895.5 | 2333521 | 2766178.5 | 2821765.5 | 2 | **0.711** |
| Q9JHI7 | Exosc9 | Exosome complex component RRP45 (Mus musculus) | 153.39 | 8.45 | 48937 | 3 | 5185326.5 | 4457357 | 4426722.5 | 5894238.5 | 5210094 | 4449778 | 6131491 | 4682616 | 4615051 | 4982963 | 12 | **0.988** |
| Q9JHJ0 | Tmod3 | Tropomodulin-3 (Mus musculus) | 197.91 | 15.06 | 39503 | 3 | 6707798 | 2962895 | 2663320.5 | 2458859.5 | 3679346.25 | 3802108 | 4356752 | 4966743.5 | 4774641 | 3795843.25 | 5 | **1.175** |
| Q9JHJ3 | Glmp | Glycosylated lysosomal membrane protein (Mus musculus) | 56.05 | 4.95 | 43804 | 1 | 1683804.12 | 563587.19 | 847973.56 | 462082.28 | 0 | 1304189.38 | 1346927.12 | 1125278.12 | 909011.19 | 1144360.75 | 5 | **1.639** |
| Q9JHK5 | Plek | Pleckstrin (Mus musculus) | 607.92 | 52.86 | 39901 | 11 | 10864186 | 16983086 | 10213953 | 14899201 | 14095854 | 14676571 | 12709646 | 14398981 | 15512771 | 11176850 | 11 | **1.021** |
| Q9JHP7 | Kdelc1 | Protein O-glucosyltransferase 2 (Mus musculus) | 137.57 | 6.37 | 57985 | 3 | 724535.12 | 1167634 | 376807.69 | 1133516.62 | 1368895.5 | 1525504.62 | 1435818.62 | 1917034.62 | 1277537.75 | 1631870.75 | 1 | **1.632** |
| Q9JHR7 | Ide | Insulin-degrading enzyme (Mus musculus) | 404.51 | 9.91 | 117772 | 9 | 8148025 | 8371598 | 10059256 | 11069980 | 8414852 | 8429227 | 8734144 | 9610364 | 9917549 | 10416547 | 9 | **1.023** |
| Q9JHS3 | Lamtor2 | Ragulator complex protein LAMTOR2 (Mus musculus) | 172.84 | 32.8 | 13480 | 3 | 1190038.38 | 1930229.38 | 1600376.38 | 1794462.38 | 1758349.88 | 1685136.88 | 596525.38 | 1736465.5 | 1670181.5 | 1911277.5 | 10 | **0.919** |
| Q9JHU4 | Dync1h1 | Cytoplasmic dynein 1 heavy chain 1 (Mus musculus) | 5693.959 | 30.28 | 532045 | 106 | 168765584 | 258110816 | 235908000 | 189210192 | 198767872 | 168802832 | 151631712 | 173420944 | 156460816 | 215755376 | 5 | **0.824** |
| Q9JHU5 | Arfip1 | Arfaptin-1 (Rattus norvegicus) | 79.58 | 3.28 | 40779 | 1 | 1332245.38 | 1636989 | 1297277.62 | 1463829.88 | 1182445.62 | 1401708.25 | 1294830.12 | 1150398.62 | 1437445.62 | 1434662.25 | 10 | **0.972** |
| Q9JHW2 | Nit2 | Omega-amidase NIT2 (Mus musculus) | 290.96 | 22.83 | 30502 | 4 | 5680939 | 2766343 | 4122472 | 4978461 | 2625701 | 5939881 | 5428010 | 4970575 | 6823314 | 4849747.5 | 5 | **1.389** |
| Q9JHW4 | Eefsec | Selenocysteine-specific elongation factor (Mus musculus) | 170.64 | 14.07 | 63539 | 4 | 1609432.12 | 1446285.38 | 1485029.25 | 1264092 | 1083492.5 | 2174604.25 | 2003280.25 | 1930322 | 1818255.75 | 1546983 | 1 | **1.375** |
| Q9JI10 | Stk3 | Serine/threonine-protein kinase 3 (Mus musculus) | 517.03 | 26.76 | 56855 | 11 | 13644397 | 17024538 | 17978442 | 15650761 | 14942632 | 17979914 | 16377508 | 18504722 | 17295954 | 16303961 | 5 | **1.091** |
| Q9JI11 | Stk4 | Serine/threonine-protein kinase 4 (Mus musculus) | 465.61 | 20.12 | 55541 | 8 | 7453869 | 11344853 | 13302543 | 10508456 | 11598648 | 14632691 | 12224529 | 14296318 | 13796708 | 10923443 | 4 | **1.215** |
| Q9JI13 | Utp3 | Something about silencing protein 10 (Mus musculus) | 108.15 | 7.25 | 53399 | 2 | 841757.25 | 229948.83 | 180530.72 | 192633.55 | 118124.09 | 653599.69 | 868899 | 286291.66 | 813598.56 | 1110790 | 3 | **2.388** |
| Q9JI18 | Lrp1b | Low-density lipoprotein receptor-related protein 1B (Mus musculus) | 75.47 | 0.37 | 513634 | 2 | 6497460.5 | 5705095.5 | 5642515.5 | 5692482.5 | 4771052.5 | 5714746 | 6703499.5 | 5043521.5 | 5666632.5 | 5860563.5 | 9 | **1.024** |
| Q9JI75 | Nqo2 | Ribosyldihydronicotinamide dehydrogenase [quinone] (Mus musculus) | 59.05 | 6.93 | 26248 | 1 | 626101.44 | 966488.69 | 1207905 | 1119210.38 | 1424730.62 | 92660.94 | 903811.5 | 1043864.88 | 1145940.12 | 1228815 | 10 | **0.826** |
| Q9JIA7 | Sphk2 | Sphingosine kinase 2 (Mus musculus) | 212.87 | 7.13 | 65618 | 3 | 3425155.75 | 2462618.25 | 2752920.75 | 2446392.5 | 2548285.5 | 3058018.5 | 3234496 | 2641442.75 | 2416231.25 | 2487860 | 12 | **1.015** |
| Q9JIF7 | Copb1 | Coatomer subunit beta (Mus musculus) | 1547.09 | 39.45 | 107066 | 30 | 38915776 | 60330800 | 54027428 | 66845372 | 45696260 | 46194196 | 45982692 | 51191480 | 50086252 | 54938828 | 11 | **0.934** |
| Q9JIH2 | Nup50 | Nuclear pore complex protein Nup50 (Mus musculus) | 291.65 | 13.3 | 49485 | 4 | 2789073 | 2987750.5 | 5761799.5 | 3749965.75 | 3028608.75 | 5884480 | 4098838 | 4827378.5 | 5235002 | 5396418 | 4 | **1.389** |
| Q9JII5 | Dazap1 | DAZ-associated protein 1 (Mus musculus) | 178.18 | 11.33 | 43214 | 3 | 3006523 | 3545635.5 | 3686293.5 | 3414365.5 | 3082283 | 1875162 | 2452774.5 | 2431534.25 | 2146500 | 1748794.88 | 0 | **0.637** |
| Q9JII6 | Akr1a1 | Alcohol dehydrogenase [NADP(+)] (Mus musculus) | 466.85 | 32.92 | 36587 | 10 | 21714854 | 14193023 | 13009048 | 11267840 | 15342526 | 17001432 | 15230420 | 14550016 | 14852763 | 9938818 | 12 | **0.948** |
| Q9JIK5 | Ddx21 | Nucleolar RNA helicase 2 (Mus musculus) | 1875.77 | 46.77 | 93551 | 32 | 103932488 | 106046992 | 86638816 | 81534576 | 69265256 | 85427120 | 87840072 | 83474480 | 76541152 | 83369136 | 10 | **0.931** |
| Q9JIK9 | Mrps34 | 28S ribosomal protein S34, mitochondrial (Mus musculus) | 87.48 | 6.88 | 25827 | 2 | 2335009.75 | 1310741.12 | 1361780.38 | 1640054.25 | 1877087 | 2132504.25 | 2069345.38 | 2439220 | 2168242.25 | 2379274.25 | 3 | **1.312** |
| Q9JIX8 | Acin1 | Apoptotic chromatin condensation inducer in the nucleus (Mus musculus) | 273.14 | 5.53 | 150719 | 6 | 6436735 | 9201810 | 9730738 | 8616594 | 5810445 | 4648736 | 3778697.5 | 7320343 | 8059906.5 | 6241184.5 | 5 | **0.755** |
| Q9JIY5 | Htra2 | Serine protease HTRA2, mitochondrial (Mus musculus) | 185.22 | 14.19 | 49348 | 3 | 2212753.75 | 3519716 | 2631708.25 | 4442565 | 3458404.75 | 2774195 | 2584784.75 | 2646665.75 | 4021712 | 3204068 | 11 | **0.936** |
| Q9JJ28 | Flii | Protein flightless-1 homolog (Mus musculus) | 757.09 | 12.82 | 144803 | 12 | 21788588 | 16940744 | 17415056 | 18717012 | 19362812 | 21816702 | 20739050 | 19292656 | 16984022 | 28253244 | 7 | **1.136** |
| Q9JJ80 | Rpf2 | Ribosome production factor 2 homolog (Mus musculus) | 229.34 | 16.99 | 35364 | 4 | 2862407.75 | 3553879.5 | 2994985.75 | 3414476.5 | 5385272.5 | 6602882.5 | 2427975.25 | 6063933.5 | 4376355.5 | 5671723.5 | 6 | **1.381** |
| Q9JJA4 | Wdr12 | Ribosome biogenesis protein WDR12 (Mus musculus) | 346.85 | 23.17 | 47347 | 6 | 3065441 | 9143611 | 9620045 | 6465578 | 7639866 | 7724665.5 | 7670386 | 7876105.5 | 7960073 | 9015661 | 10 | **1.120** |
| Q9JJT0 | Rcl1 | RNA 3'-terminal phosphate cyclase-like protein (Mus musculus) | 76.73 | 4.02 | 40841 | 1 | 0 | 665122.62 | 596632.5 | 572585.19 | 0 | 0 | 0 | 640543 | 0 | 457357.91 | 15 | **0.599** |
| Q9JJU8 | Sh3bgrl | SH3 domain-binding glutamic acid-rich-like protein (Mus musculus) | 337.62 | 55.26 | 12811 | 5 | 23033020 | 14360847 | 15444038 | 13291282 | 18125896 | 19195598 | 19093856 | 19513972 | 19649836 | 20584700 | 5 | **1.164** |
| Q9JJY4 | Ddx20 | Probable ATP-dependent RNA helicase DDX20 (Mus musculus) | 109.39 | 2.91 | 91710 | 2 | 88743.87 | 1355936.75 | 658172.25 | 0 | 454709.38 | 1304106 | 1483474.75 | 1757362.12 | 934980.12 | 1620780.75 | 2 | **2.776** |
| Q9JJZ4 | Ube2j1 | Ubiquitin-conjugating enzyme E2 J1 (Mus musculus) | 131.39 | 11.01 | 34990 | 3 | 2954302 | 3160160.5 | 3113269.5 | 2545406.75 | 3999890.25 | 1868412.88 | 2746682 | 3720748.25 | 3640816 | 3709218 | 12 | **0.994** |
| Q9JK23 | Psmg1 | Proteasome assembly chaperone 1 (Mus musculus) | 288.67 | 20.07 | 33104 | 5 | 8831279 | 6946882.5 | 7451736 | 6993773 | 7357287.5 | 11400424 | 9181466 | 10070964 | 10222900 | 9556267 | 0 | **1.342** |
| Q9JK38 | Gnpnat1 | Glucosamine 6-phosphate N-acetyltransferase (Mus musculus) | 75.32 | 5.98 | 20791 | 1 | 2661412 | 2474580.75 | 2693614 | 2763716.25 | 3889806.75 | 3693301.5 | 2837265.75 | 2258614 | 3314839 | 2123333.25 | 12 | **0.982** |
| Q9JK81 | Myg1 | UPF0160 protein MYG1, mitochondrial (Mus musculus) | 161.1 | 11.32 | 42723 | 4 | 3294616.25 | 3758939.75 | 3268202.5 | 1991718.38 | 1824434.12 | 2272021.5 | 3513218 | 2826797.25 | 5045825 | 2437091 | 10 | **1.138** |
| Q9JK91 | Mlh1 | DNA mismatch repair protein Mlh1 (Mus musculus) | 49.7 | 1.05 | 84670 | 1 | 1077383.38 | 575763.5 | 552188.12 | 651208.94 | 0 | 556045.56 | 535639.88 | 690092 | 616402.12 | 572813.25 | 12 | **1.040** |
| Q9JKB3 | Ybx3 | Y-box-binding protein 3 (Mus musculus) | 285.58 | 19.94 | 38814 | 4 | 3336067.5 | 1544372 | 10449169 | 3135558.5 | 8974133 | 3744814 | 2509302.5 | 7741403 | 7713885 | 8237101 | 12 | **1.091** |
| Q9JKF1 | Iqgap1 | Ras GTPase-activating-like protein IQGAP1 (Mus musculus) | 5232.991 | 59.08 | 188742 | 82 | 262882112 | 288994336 | 318926208 | 285131808 | 296354656 | 342866144 | 313387744 | 325381184 | 322622304 | 328476576 | 1 | **1.124** |
| Q9JKF7 | Mrpl39 | 39S ribosomal protein L39, mitochondrial (Mus musculus) | 176.16 | 11.31 | 38549 | 3 | 1620793.25 | 3000814.5 | 2525744 | 1808270.75 | 1811542.12 | 2447173 | 2671516.5 | 2365854 | 902014.12 | 1961680 | 12 | **0.961** |
| Q9JKL8 | Adnp | Activity-dependent neuroprotector homeobox protein (Rattus norvegicus) | 42.87 | 0.82 | 123626 | 1 | 475694.84 | 0 | 0 | 0 | 0 | 530835.38 | 635827.69 | 459384.97 | 0 | 0 | 17 | **3.418** |
| Q9JKN1 | Slc30a7 | Zinc transporter 7 (Mus musculus) | 39.4 | 3.97 | 41790 | 1 | 821925.06 | 831741.31 | 1108567.25 | 0 | 1048832 | 607354.56 | 993504.75 | 717952.94 | 806735.56 | 867753.19 | 9 | **1.048** |
| Q9JKP5 | Mbnl1 | Muscleblind-like protein 1 (Mus musculus) | 250.37 | 21.99 | 36976 | 4 | 15648503 | 14622969 | 12562127 | 12871019 | 13943086 | 11761713 | 9526042 | 10140380 | 10345586 | 9675524 | 0 | **0.739** |
| Q9JKR6 | Hyou1 | Hypoxia up-regulated protein 1 (Mus musculus) | 1757.98 | 37.84 | 111181 | 28 | 89806992 | 76598368 | 71505576 | 79272552 | 67870928 | 80122536 | 89475296 | 77334840 | 83872024 | 74602248 | 8 | **1.053** |
| Q9JKV1 | Adrm1 | Proteasomal ubiquitin receptor ADRM1 (Mus musculus) | 96.02 | 7.62 | 42060 | 2 | 5516537.5 | 2849727.25 | 3435112 | 1990207.25 | 3318559.5 | 4883134 | 4822599 | 2984563.5 | 3483185.75 | 5046095.5 | 7 | **1.240** |
| Q9JKX6 | Nudt5 | ADP-sugar pyrophosphatase (Mus musculus) | 301.01 | 43.12 | 23984 | 6 | 8595751 | 11686395 | 10142004 | 9009768 | 10217419 | 10671610 | 12226422 | 10846684 | 12125471 | 11668045 | 3 | **1.159** |
| Q9JL26 | Fmnl1 | Formin-like protein 1 (Mus musculus) | 1465.79 | 33.36 | 122060 | 27 | 49654980 | 52735944 | 56492260 | 56462224 | 56794928 | 51993464 | 55640672 | 58230456 | 55813820 | 58191276 | 10 | **1.028** |
| Q9JL35 | Hmgn5 | High mobility group nucleosome-binding domain-containing protein 5 (Mus musculus) | 199.63 | 11.33 | 45344 | 4 | 12940037 | 6959587 | 6683532.5 | 6372452 | 8746998 | 10982060 | 8757449 | 10236912 | 12981259 | 12230278 | 4 | **1.323** |
| Q9JLB0 | Mpp6 | MAGUK p55 subfamily member 6 (Mus musculus) | 55.42 | 2.17 | 62631 | 1 | 0 | 0 | 0 | 1686042.88 | 0 | 0 | 0 | 663026.94 | 1053110.38 | 885701.88 | 19 | **1.543** |
| Q9JLC8 | Sacs | Sacsin (Mus musculus) | 61.36 | 0.35 | 520685 | 2 | 11963284 | 11789488 | 10390196 | 12975868 | 12502982 | 8051224.5 | 7975492 | 4157240.25 | 7762141 | 6517474 | 0 | **0.578** |
| Q9JLI6 | Scly | Selenocysteine lyase (Mus musculus) | 213.9 | 6.71 | 47174 | 3 | 3054296.75 | 2022220.25 | 1932180.88 | 1818658.25 | 2540946.25 | 2624228.5 | 2054423.75 | 2196904.75 | 1850463.62 | 2326300.5 | 11 | **0.972** |
| Q9JLI8 | Sart3 | Squamous cell carcinoma antigen recognized by T-cells 3 (Mus musculus) | 79.85 | 2.18 | 109619 | 2 | 3266683 | 2870014.75 | 3258437.75 | 2656413.5 | 3634927 | 3095853.5 | 2661179.5 | 3081492 | 2874290.75 | 2901079 | 9 | **0.932** |
| Q9JLJ2 | Aldh9a1 | 4-trimethylaminobutyraldehyde dehydrogenase (Mus musculus) | 1196.92 | 58.91 | 53515 | 20 | 72352784 | 73709064 | 74641232 | 56423076 | 65866992 | 84215976 | 73313744 | 68181048 | 73269792 | 67059368 | 10 | **1.067** |
| Q9JLJ5 | Elovl1 | Elongation of very long chain fatty acids protein 1 (Mus musculus) | 92.7 | 7.53 | 32678 | 2 | 0 | 249801 | 453619.66 | 164137.02 | 158999.8 | 473893.44 | 382109 | 863168.38 | 614524.81 | 616150.5 | 1 | **2.874** |
| Q9JLQ0 | Cd2ap | CD2-associated protein (Mus musculus) | 272.31 | 11.46 | 70450 | 6 | 4788245.5 | 2330274.5 | 4289956 | 2965666.75 | 3222561 | 5304131.5 | 4792175.5 | 4744992 | 4756933 | 4779801.5 | 3 | **1.385** |
| Q9JLQ2 | Git2 | ARF GTPase-activating protein GIT2 (Mus musculus) | 51.29 | 2.54 | 78766 | 1 | 499475.06 | 539753.06 | 494329.72 | 0 | 0 | 403102.84 | 526547.88 | 392579.66 | 129430.52 | 0 | 12 | **0.947** |
| Q9JLV6 | Pnkp | Bifunctional polynucleotide phosphatase/kinase (Mus musculus) | 148.15 | 8.81 | 57223 | 3 | 6598094.5 | 2626046 | 2584892 | 2870635 | 781549.38 | 2381746 | 4417226.5 | 2459684.75 | 2414671 | 2572055.5 | 8 | **0.921** |
| Q9JM14 | Nt5c | 5'(3')-deoxyribonucleotidase, cytosolic type (Mus musculus) | 251.12 | 38.5 | 23076 | 4 | 3038828.75 | 4488557 | 4209418 | 4481618 | 2256615.25 | 3285299.5 | 3352519.5 | 3032405.5 | 3312161 | 3021025 | 8 | **0.866** |
| Q9JM76 | Arpc3 | Actin-related protein 2/3 complex subunit 3 (Mus musculus) | 445.11 | 49.44 | 20525 | 8 | 66247268 | 31179762 | 41574216 | 14627242 | 26343044 | 51939036 | 52107860 | 45201556 | 49782000 | 48722236 | 5 | **1.377** |
| Q9JM90 | Stap1 | Signal-transducing adaptor protein 1 (Mus musculus) | 331.83 | 36.7 | 34628 | 7 | 7034405 | 10788832 | 8133742.5 | 10860467 | 6478564.5 | 9247212 | 7925570 | 9920222 | 8119982 | 7764662 | 12 | **0.993** |
| Q9JMA1 | Usp14 | Ubiquitin carboxyl-terminal hydrolase 14 (Mus musculus) | 877.76 | 35.09 | 56002 | 16 | 28433840 | 33815912 | 32931168 | 32630184 | 31153548 | 32373990 | 32725676 | 37066620 | 26460450 | 35957348 | 10 | **1.035** |
| Q9JMD0 | Znf207 | BUB3-interacting and GLEBS motif-containing protein ZNF207 (Mus musculus) | 152.64 | 6.67 | 52793 | 3 | 2966680.25 | 1078623.12 | 2015200.25 | 536692.12 | 1482453.38 | 3081776.25 | 1775661 | 1721023.5 | 2612464.5 | 1826531.88 | 7 | **1.364** |

| | | | | | | | | | | | | | | | | | | |
|---|---|---|---|---|---|---|---|---|---|---|---|---|---|---|---|---|---|---|
| Q9JMH6 | Txnrd1 | Thioredoxin reductase 1, cytoplasmic (Mus musculus) | 1005.5 | 41.11 | 67084 | 15 | 38007252 | 43256372 | 44651728 | 45805612 | 41426752 | 47096208 | 47184376 | 47558532 | 45008784 | 44926072 | 2 | **1.087** |
| Q9JMH9 | Myo18a | Unconventional myosin-XVIIIa (Mus musculus) | 267.25 | 2.83 | 232755 | 5 | 2500831.5 | 3201123.5 | 3636395.5 | 2431576 | 2570533.5 | 1630314.5 | 2912820.5 | 2293012.75 | 4296879.5 | 2968760.5 | 11 | **0.983** |
| Q9QUH0 | Glrx | Glutaredoxin-1 (Mus musculus) | 343.66 | 57.01 | 11871 | 5 | 17896298 | 15843557 | 21054932 | 13543004 | 21908702 | 23023972 | 19054942 | 18173434 | 18253054 | 19149112 | 8 | **1.082** |
| Q9QUJ7 | Acsl4 | Long-chain-fatty-acid--CoA ligase 4 (Mus musculus) | 922.86 | 31.93 | 79077 | 18 | 102557992 | 54358928 | 54434024 | 55753804 | 62249816 | 60735304 | 64072236 | 72398968 | 64856728 | 70750312 | 6 | **1.011** |
| Q9QUM9 | Psma6 | Proteasome subunit alpha type-6 (Mus musculus) | 682.35 | 56.1 | 27372 | 12 | 78785000 | 78899720 | 88358208 | 62492956 | 77732312 | 73036168 | 69486864 | 71482280 | 71181784 | 81240944 | 8 | **0.949** |
| Q9QUN7 | Tlr2 | Toll-like receptor 2 (Mus musculus) | 243.1 | 9.31 | 89449 | 5 | 840126.25 | 3041644.25 | 3941297 | 2751767.5 | 2680343 | 1594734.88 | 2970489 | 4706178.5 | 4758188 | 5204565.5 | 6 | **1.451** |
| Q9QUR6 | Prep | Prolyl endopeptidase (Mus musculus) | 1076.04 | 36.62 | 80752 | 19 | 12635505 | 18766020 | 17443842 | 18897068 | 19318228 | 21582384 | 18710922 | 20587838 | 20603728 | 19493962 | 3 | **1.160** |
| Q9QUR7 | Pin1 | Peptidyl-prolyl cis-trans isomerase NIMA-interacting 1 (Mus musculus) | 46.34 | 9.09 | 18370 | 1 | 499883.72 | 0 | 0 | 0 | 847897.75 | 554552.5 | 637339.12 | 663910.88 | 778831.19 | 647042.19 | 8 | **2.435** |
| Q9QVP9 | Ptk2b | Protein-tyrosine kinase 2-beta (Mus musculus) | 719.37 | 20.42 | 115794 | 14 | 11993760 | 11846782 | 13928520 | 11522341 | 16021622 | 15323338 | 14305219 | 15489043 | 15768023 | 15368232 | 5 | **1.168** |
| Q9QWF0 | Chaf1a | Chromatin assembly factor 1 subunit A (Mus musculus) | 51.26 | 0.99 | 101936 | 1 | 1386686.5 | 1415201.38 | 3365308.5 | 1474608.38 | 1406559.12 | 1071439.75 | 4184630.5 | 1741663.12 | 1934676.25 | 1741239 | 8 | **1.180** |
| Q9QWR8 | Naga | Alpha-N-acetylgalactosaminidase (Mus musculus) | 416.14 | 20 | 47235 | 7 | 7608687 | 12253323 | 11789407 | 9678773 | 9352969 | 8608025 | 8560474 | 8590995 | 8575622 | 11790759 | 8 | **0.910** |
| Q9QX47 | Son | Protein SON (Mus musculus) | 246.71 | 4.26 | 265651 | 5 | 4652330 | 5140802 | 5239939 | 5108272 | 3310664.5 | 3668042 | 4438562.5 | 4191450.5 | 3963826.5 | 4254826 | 5 | **0.875** |
| Q9QXB9 | Drg2 | Developmentally-regulated GTP-binding protein 2 (Mus musculus) | 463.54 | 25.82 | 40718 | 8 | 8672917 | 11766404 | 12967976 | 10926342 | 7175209.5 | 12958193 | 11704447 | 13255659 | 13395102 | 10355583 | 6 | **1.197** |
| Q9QXG2 | Chm | Rab proteins geranylgeranyltransferase component A 1 (Mus musculus) | 136.05 | 4.36 | 73977 | 3 | 1901511 | 1740534.12 | 2152419.25 | 2063346.12 | 1133914.12 | 2151019.75 | 1632441 | 1352299.12 | 1262218.5 | 1916078.25 | 10 | **0.925** |
| Q9QXG4 | Acss2 | Acetyl-coenzyme A synthetase, cytoplasmic (Mus musculus) | 47.97 | 2.28 | 78862 | 1 | 0 | 0 | 0 | 0 | 0 | 283430.22 | 0 | 0 | 534892.56 | 595881.44 | 20 | **#DIV/0 !** |
| Q9QXK2 | Rad18 | E3 ubiquitin-protein ligase RAD18 (Mus musculus) | 42.37 | 4.72 | 57412 | 1 | 788972.75 | 738341.31 | 795459 | 965156.19 | 1002663.12 | 741428.38 | 863521.81 | 741885.06 | 653998.69 | 767813.75 | 6 | **0.878** |
| Q9QXK3 | Copg2 | Coatomer subunit gamma-2 (Mus musculus) | 715.35 | 21.47 | 97681 | 13 | 15255092 | 19062912 | 18131068 | 22025118 | 15806239 | 17434620 | 16132532 | 16280716 | 15529880 | 20909412 | 11 | **0.956** |
| Q9QXK7 | Cpsf3 | Cleavage and polyadenylation specificity factor subunit 3 (Mus musculus) | 147.66 | 7.89 | 77505 | 4 | 5096421 | 4822719 | 4702245.5 | 3713710.75 | 4420068.5 | 5425597 | 5671296.5 | 6035158.5 | 5204604.5 | 5232873.5 | 0 | **1.212** |
| Q9QXS1 | Plec | Plectin (Mus musculus) | 7918.789 | 37.39 | 534188 | 139 | 364743520 | 347232064 | 334147904 | 343994272 | 328968032 | 356974208 | 332121120 | 349480480 | 360825536 | 352311520 | 8 | **1.019** |
| Q9QXT0 | Cnpy2 | Protein canopy homolog 2 (Mus musculus) | 66.18 | 8.79 | 20767 | 1 | 1543191.62 | 1042412.25 | 837924.19 | 0 | 1331852.88 | 0 | 895325.25 | 755811.06 | 0 | 909918.12 | 8 | **0.539** |
| Q9QXW9 | Slc7a8 | Large neutral amino acids transporter small subunit 2 (Mus musculus) | 49.01 | 2.07 | 57873 | 1 | 0 | 0 | 207224.12 | 0 | 211013.62 | 306919.06 | 329197.47 | 364093.59 | 417380.91 | 328070.31 | 3 | **4.174** |
| Q9QXX4 | Slc25a13 | Calcium-binding mitochondrial carrier protein Aralar2 (Mus musculus) 3 | 1482.75 | 49.26 | 74467 | 24 | 46786200 | 47790664 | 42580404 | 49993524 | 37923556 | 41392212 | 43049272 | 41234752 | 43271056 | 44740736 | 8 | **0.949** |
| Q9QXZ0 | Macf1 | Microtubule-actin cross-linking factor 1 (Mus musculus) | 557.13 | 2.11 | 831878 | 11 | 3903796.5 | 5296611.5 | 4648007 | 3994152.5 | 4210420 | 3484477.5 | 4375237 | 3828528 | 3056218.5 | 3704707.25 | 3 | **0.837** |
| Q9QY76 | Vapb | Vesicle-associated membrane protein-associated protein B (Mus musculus) | 336.04 | 24.28 | 26946 | 6 | 6644623 | 4824924 | 6327744 | 4464134 | 6717954 | 6892839 | 8609200 | 10190532 | 10238523 | 9504947 | 0 | **1.568** |
| Q9QY81 | Nup210 | Nuclear pore membrane glycoprotein 210 (Mus musculus) | 491.96 | 7.95 | 204101 | 10 | 6690803.5 | 7706714.5 | 9236003 | 6366034 | 5160776 | 4707846 | 5540403.5 | 6610017 | 6425683.5 | 9173525 | 9 | **0.923** |
| Q9QYB1 | Clic4 | Chloride intracellular channel protein 4 (Mus musculus) | 1008.15 | 67.19 | 28729 | 14 | 34211532 | 38653336 | 35810908 | 40713680 | 37635036 | 40618160 | 38738608 | 41295436 | 35322936 | 38858380 | 7 | **1.042** |
| Q9QYF1 | Rdh11 | Retinol dehydrogenase 11 (Mus musculus) | 163.53 | 11.08 | 35148 | 3 | 9601927 | 5247675.5 | 6260237.5 | 5327430.5 | 5592704 | 8707266 | 7737484 | 7134321 | 5898295 | 7140122 | 6 | **1.143** |
| Q9QYI3 | Dnajc7 | DnaJ homolog subfamily C member 7 (Mus musculus) | 454.66 | 20.65 | 56476 | 8 | 10920561 | 9648754 | 10872103 | 9316711 | 6883634.5 | 11898399 | 12623760 | 12346370 | 8843088 | 13631655 | 4 | **1.246** |
| Q9QYJ0 | Dnaja2 | DnaJ homolog subfamily A member 2 (Mus musculus) | 570.18 | 44.9 | 45746 | 10 | 11681613 | 26518130 | 24543936 | 25904616 | 24740310 | 28155156 | 21111000 | 24252120 | 25099386 | 25434672 | 12 | **1.094** |
| Q9QYJ3 | Dnajb1 | DnaJ homolog subfamily B member 1 (Mus musculus) | 135.68 | 10.88 | 38167 | 4 | 4829990 | 6707438 | 4362186 | 4408623 | 5252682.5 | 4305168 | 3997078 | 4016600 | 5114635.5 | 4283194.5 | 3 | **0.850** |
| Q9QZ08 | Nagk | N-acetyl-D-glucosamine kinase (Mus musculus) | 101.03 | 5.25 | 37268 | 2 | 794177.75 | 1976510 | 2162333.5 | 1521759.75 | 1569490 | 2116899.75 | 1859989.5 | 2228902.5 | 2892267.75 | 1733013.75 | 5 | **1.350** |
| Q9QZ23 | Nfu1 | NFU1 iron-sulfur cluster scaffold homolog, mitochondrial (Mus musculus) | 192.08 | 16.47 | 28568 | 3 | 4545679 | 2947013 | 3315283 | 2645614 | 2900477.5 | 2932642.25 | 3531820.5 | 2887008 | 3328171.5 | 3303536 | 11 | **0.977** |
| Q9QZ73 | Dcun1d1 | DCN1-like protein 1 (Mus musculus) | 110.16 | 6.56 | 30097 | 1 | 736720.19 | 807107.31 | 467208.5 | 0 | 356617.12 | 412845.59 | 536448.62 | 693599.88 | 579126.5 | 382440.53 | 12 | **1.100** |
| Q9QZ88 | Vps29 | Vacuolar protein sorting-associated protein 29 (Mus musculus) | 223.16 | 29.67 | 20496 | 4 | 16989602 | 10247851 | 15111680 | 9259341 | 11382006 | 16803276 | 15297844 | 14316493 | 15837938 | 16146442 | 6 | **1.245** |
| Q9QZB7 | Actr10 | Actin-related protein 10 (Mus musculus) | 148.51 | 10.55 | 46208 | 3 | 1092676.5 | 767906.06 | 672576.44 | 596555.19 | 731921.94 | 1120681.88 | 1033292.88 | 671601.06 | 1075818 | 675336.44 | 9 | **1.185** |
| Q9QZD8 | Slc25a10 | Mitochondrial dicarboxylate carrier (Mus musculus) | 327.72 | 28.92 | 31715 | 5 | 4036830.75 | 5585938 | 5307387.5 | 4369161 | 5219476.5 | 6396479.5 | 6882465 | 6710438 | 5859767 | 6074167 | 0 | **1.302** |
| Q9QZD9 | Eif3i | Eukaryotic translation initiation factor 3 subunit I (Mus musculus) | 489.74 | 40.31 | 36461 | 10 | 10611700 | 11382407 | 11189722 | 13976795 | 11704008 | 9892327 | 10198566 | 8286151 | 10627191 | 11544301 | 4 | **0.859** |
| Q9QZE5 | Copg1 | Coatomer subunit gamma-1 (Mus musculus) | 1188.09 | 38.1 | 97513 | 22 | 28450378 | 27077588 | 26188578 | 23764988 | 20416238 | 29275850 | 29483708 | 25873082 | 20173954 | 23530874 | 12 | **1.019** |
| Q9QZE7 | Tsnax | Translin-associated protein X (Mus musculus) | 372.16 | 28.97 | 32926 | 6 | 3842143.5 | 3549835 | 5299757 | 6377124.5 | 5865807.5 | 4746434.5 | 4752095.5 | 5424709 | 5114239 | 5260175.5 | 11 | **1.015** |
| Q9QZL0 | Ripk3 | Receptor-interacting serine/threonine-protein kinase 3 (Mus musculus) | 519.04 | 36.01 | 53322 | 10 | 5288255 | 6819345 | 8782628 | 5988696.5 | 6505368 | 8422389 | 7499201.5 | 9794815 | 8538059 | 8996390 | 3 | **1.296** |
| Q9QZQ8 | H2afy | Core histone macro-H2A.1 (Mus musculus) | 406.43 | 32.8 | 39735 | 8 | 12460421 | 14853766 | 14213304 | 8793063 | 13738841 | 15951191 | 13062817 | 16414326 | 16279392 | 15325208 | 3 | **1.203** |
| Q9R059 | Fhl3 | Four and a half LIM domains protein 3 (Mus musculus) | 157.19 | 13.84 | 31795 | 3 | 9038762 | 5977058 | 4222119.5 | 4884137 | 7035311 | 6414717.5 | 7357525.5 | 6831012 | 7854536 | 7899222 | 7 | **1.167** |
| Q9R061 | Nubp2 | Cytosolic Fe-S cluster assembly factor NUBP2 (Mus musculus) | 65.55 | 6.55 | 29518 | 1 | 1565332 | 1380689.75 | 1791734 | 1042798.56 | 1281507.12 | 1109751.25 | 0 | 1337073 | 1566602.5 | 1406096.88 | 10 | **0.767** |
| Q9R099 | Tbl2 | Transducin beta-like protein 2 (Mus musculus) | 58.98 | 2.94 | 49584 | 1 | 0 | 0 | 1409986 | 0 | 1116980.62 | 0 | 272852.41 | 1161057.12 | 1158845.38 | 1279160.12 | 13 | **1.532** |
| Q9R0C8 | Vav3 | Guanine nucleotide exchange factor VAV3 (Mus musculus) | 498.24 | 16.53 | 97968 | 10 | 7103368.5 | 9849146 | 9743843 | 8686404 | 6478430 | 8277696.5 | 7201261 | 10222108 | 10090053 | 10162272 | 6 | **1.098** |
| Q9R0E1 | Plod3 | Multifunctional procollagen lysine hydroxylase and glycosyltransferase LH3 (Mus musculu | 143 | 5.8 | 84922 | 3 | 4499867 | 2723531.25 | 3369832.5 | 3284556.5 | 4691309.5 | 3674400 | 4269370 | 3305661 | 3702448.5 | 3992991.5 | 11 | **1.020** |
| Q9R0E2 | Plod1 | Procollagen-lysine,2-oxoglutarate 5-dioxygenase 1 (Mus musculus) | 196.76 | 7.55 | 83595 | 5 | 2966008.25 | 4995521.5 | 3464665.75 | 4940184.5 | 3459487.75 | 4244092.5 | 3585477 | 3236602 | 3789119.5 | 4227415 | 12 | **0.963** |
| Q9R0H0 | Acox1 | Peroxisomal acyl-coenzyme A oxidase 1 (Mus musculus) | 46.79 | 2.42 | 74649 | 1 | 119678.76 | 183337.75 | 175921 | 37064.62 | 26635.17 | 141901.95 | 216973.64 | 31191.83 | 54552.93 | 53837.02 | 12 | **0.919** |
| Q9R0L6 | Pcm1 | Pericentriolar material 1 protein (Mus musculus) | 67.76 | 1.43 | 228847 | 2 | 1431934.5 | 1618692.12 | 936585.75 | 1214718.5 | 1191061.38 | 919387.69 | 832221.25 | 1169244.62 | 599293.06 | 969213.38 | 2 | **0.702** |
| Q9R0N0 | Galk1 | Galactokinase (Mus musculus) | 475.76 | 32.4 | 42295 | 9 | 21391628 | 15705395 | 16063801 | 20044938 | 17511892 | 19785314 | 19509254 | 17653840 | 14846528 | 15749437 | 10 | **0.965** |
| Q9R0P3 | Esd | S-formylglutathione hydrolase (Mus musculus) | 984.8 | 68.44 | 31320 | 14 | 116118864 | 98588280 | 81708928 | 93193256 | 102583168 | 147383392 | 115428744 | 127883888 | 117502032 | 124613480 | 1 | **1.286** |
| Q9R0P4 | Smap | Small acidic protein (Mus musculus) | 101.85 | 8.84 | 20046 | 1 | 411829.62 | 0 | 480614.44 | 0 | 606805.75 | 354189.19 | 348152.62 | 429377.69 | 458460.03 | 499336.75 | 12 | **1.394** |
| Q9R0P5 | Dstn | Destrin (Mus musculus) | 613.77 | 52.12 | 18522 | 11 | 35306564 | 37236536 | 29346344 | 37536804 | 44299540 | 32459674 | 31304302 | 30084150 | 27633764 | 32929300 | 4 | **0.840** |
| Q9R0Q8 | Clec4e | C-type lectin domain family 4 member E (Mus musculus) | 122.39 | 12.15 | 24431 | 2 | 1777447.75 | 1325933.5 | 1669135 | 1328812 | 1802028.38 | 1049469.62 | 1822932.75 | 2166761.75 | 1956459.38 | 2115906.25 | 5 | **1.153** |
| Q9R0Q9 | Mpdu1 | Mannose-P-dolichol utilization defect 1 protein (Mus musculus) | 85.97 | 10.12 | 26498 | 2 | 2074227.25 | 760033.5 | 980003.56 | 1573424.12 | 966401.5 | 561392.5 | 940974.56 | 2192059 | 1200045.12 | 2027282.5 | 12 | **1.089** |
| Q9R0T8 | Ikbke | Inhibitor of nuclear factor kappa-B kinase subunit epsilon (Mus musculus) | 105.77 | 3.35 | 80953 | 2 | 1607899.62 | 1026246.75 | 1077920.25 | 1026225 | 840521.75 | 1202663.75 | 1415609.75 | 1291846.38 | 1429926 | 1356119.75 | 5 | **1.200** |
| Q9R0U0 | Srsf10 | Serine/arginine-rich splicing factor 10 (Mus musculus) | 237.81 | 18.7 | 31301 | 4 | 10027316 | 8387574.5 | 7510103.5 | 8525130 | 11471611 | 7860096.5 | 8215503.5 | 7521458 | 9308765 | 7671660.5 | 7 | **0.884** |
| Q9R112 | Sqor | Sulfide:quinone oxidoreductase, mitochondrial (Mus musculus) | 926.17 | 42 | 50282 | 14 | 40808648 | 31684136 | 39885096 | 38344684 | 29270636 | 34385496 | 36886632 | 42040716 | 40827212 | 38263944 | 9 | **1.069** |
| Q9R190 | Mta2 | Metastasis-associated protein MTA2 (Mus musculus) | 73.93 | 2.69 | 75030 | 2 | 5103447 | 3419922.5 | 3975993.75 | 2240922.5 | 2941331.75 | 5804106 | 4591037 | 4659588 | 4357788 | 3590493.5 | 5 | **1.301** |
| Q9R1C7 | Prpf40a | Pre-mRNA-processing factor 40 homolog A (Mus musculus) | 327.34 | 7.35 | 108481 | 5 | 6334788.5 | 6424074.5 | 7233328 | 8873647 | 7065689.5 | 5739360 | 8215178.5 | 6895936.5 | 6881132 | 7281048 | 12 | **0.974** |
| Q9R1J0 | Nsdhl | Sterol-4-alpha-carboxylate 3-dehydrogenase, decarboxylating (Mus musculus) | 203.4 | 16.57 | 40686 | 4 | 2100879.5 | 2803046 | 3262353.5 | 2059708.62 | 2406606.5 | 2787237.5 | 2655707.5 | 2311711.5 | 2649353 | 2885659.75 | 10 | **1.052** |
| Q9R1P1 | Psmb3 | Proteasome subunit beta type-3 (Mus musculus) | 720.54 | 49.76 | 22965 | 11 | 31777002 | 38111624 | 34497924 | 32182358 | 36847844 | 35766048 | 31344782 | 32393916 | 33368222 | 32024312 | 8 | **0.951** |
| Q9R1P3 | Psmb2 | Proteasome subunit beta type-2 (Mus musculus) | 478.36 | 66.17 | 22906 | 9 | 33693668 | 21455228 | 34334620 | 22590626 | 32548966 | 36513744 | 34618028 | 32782904 | 34288536 | 30979886 | 6 | **1.170** |
| Q9R1P4 | Psma1 | Proteasome subunit alpha type-1 (Mus musculus) | 613.29 | 65.02 | 29547 | 13 | 42823060 | 35305892 | 34309424 | 38341128 | 47692076 | 44316528 | 24048410 | 36764680 | 39201412 | 37288904 | 11 | **0.915** |
| Q9R1Q6 | Tmem176b | Transmembrane protein 176B (Mus musculus) b | 50.5 | 5.7 | 28367 | 1 | 427012.16 | 398030.12 | 382346.59 | 0 | 0 | 568241.69 | 1121387.62 | 677720.5 | 52287.91 | 75655.11 | 7 | **2.067** |
| Q9R1S3 | Pign | GPI ethanolamine phosphate transferase 1 (Mus musculus) | 59.96 | 1.29 | 105045 | 1 | 141103.14 | 145846.34 | 167419.3 | 429081.62 | 464395.41 | 157587.95 | 242117 | 857189.56 | 731136.19 | 623320.06 | 5 | **1.937** |
| Q9R1T2 | Sae1 | SUMO-activating enzyme subunit 1 (Mus musculus) | 573.01 | 26.86 | 38620 | 9 | 17956360 | 22488628 | 20965186 | 27157372 | 18316458 | 14280738 | 16826058 | 16255768 | 16682310 | 15636784 | 0 | **0.745** |
| Q9R1T4 | Sept6 | Septin-6 (Mus musculus) | 236.25 | 16.59 | 49620 | 5 | 8458874 | 9450570 | 9811591 | 7335221 | 8775534 | 8082690 | 5653885 | 9085331 | 8573510 | 8388490 | 7 | **0.908** |
| Q9R233 | Tapbp | Tapasin (Mus musculus) | 74.15 | 2.15 | 49736 | 1 | 5323044 | 2878588.5 | 2145491.25 | 3473144.75 | 3159772.75 | 1833276.25 | 2768517 | 1880638.12 | 5343270 | 0 | 6 | **0.696** |
| Q9WTI7 | Myo1c | Unconventional myosin-Ic (Mus musculus) | 1087.13 | 22.58 | 121944 | 19 | 9883394 | 6057990 | 11036076 | 5059040 | 12361700 | 18284760 | 12902204 | 14849386 | 13437869 | 13066853 | 0 | **1.634** |
| Q9WTJ8 | Fam50b | Protein FAM50B (Mus musculus) | 45.75 | 3.89 | 39594 | 1 | 1189674.62 | 1406542.25 | 1560053.75 | 1739657.25 | 1266977.38 | 1361698.5 | 1130950.88 | 1218990.25 | 1371262.62 | 1018051.69 | 5 | **0.852** |
| Q9WTK3 | Gpaa1 | Glycosylphosphatidylinositol anchor attachment 1 protein (Mus musculus) | 41.19 | 2.42 | 67949 | 1 | 0 | 0 | 382138.34 | 228646.27 | 254048.77 | 0 | 0 | 426730.31 | 553122.81 | 520040.09 | 11 | **1.734** |
| Q9WTK5 | Nfkb2 | Nuclear factor NF-kappa-B p100 subunit (Mus musculus) | 148.44 | 4.23 | 96832 | 3 | 3762418.75 | 2861340.5 | 3243930.5 | 2434206 | 2121875.75 | 3269981 | 3917656 | 3495248.75 | 3331639.25 | 3468407 | 4 | **1.212** |
| Q9WTM5 | Ruvbl2 | RuvB-like 2 (Mus musculus) | 874.97 | 40.6 | 51113 | 15 | 49207408 | 51495044 | 46660328 | 52274380 | 43486460 | 44556720 | 46222248 | 46570360 | 49434584 | 50791540 | 9 | **0.977** |
| Q9WTP6 | Ak2 | Adenylate kinase 2, mitochondrial (Mus musculus) | 172.77 | 20.08 | 26469 | 3 | 2496893.5 | 1441898.88 | 278502.09 | 484724.56 | 295846.03 | 2079205.88 | 1451056.62 | 1569754.25 | 1551135 | 1633691.25 | 5 | **1.658** |
| Q9WTR1 | Trpv2 | Transient receptor potential cation channel subfamily V member 2 (Mus musculus) | 65.52 | 1.85 | 85965 | 1 | 147477.14 | 0 | 0 | 161971.67 | 247087.05 | 0 | 0 | 494969.44 | 475390.44 | 388332.62 | 11 | **2.441** |
| Q9WTX6 | Cul1 | Cullin-1 (Mus musculus) | 93.79 | 3.74 | 89692 | 2 | 181402.34 | 367619.91 | 470933 | 1072252.25 | 319405.19 | 97395.62 | 943957.75 | 133134.77 | 80989.57 | 190021.11 | 5 | **0.599** |
| Q9WTZ0 | Uxt | Protein UXT (Mus musculus) | 40.13 | 7.01 | 18187 | 1 | 707634.44 | 1009423.31 | 747646.88 | 680102.56 | 737471.44 | 0 | 625399 | 696516.94 | 772481.69 | 700629.75 | 6 | **0.720** |
| Q9WU28 | Pfdn5 | Prefoldin subunit 5 (Mus musculus) | 234.15 | 38.31 | 17356 | 4 | 5780808.5 | 5976750.5 | 5938291.5 | 5053369 | 7823889 | 6048173 | 6353364 | 6385857 | 7718845 | 6225746 | 5 | **1.071** |
| Q9WU40 | Lemd3 | Inner nuclear membrane protein Man1 (Mus musculus) | 43.82 | 1.09 | 100307 | 1 | 706284.69 | 0 | 0 | 430811.5 | 0 | 0 | 567551.19 | 737426.69 | 601773.06 | 612453.19 | 11 | **2.215** |
| Q9WU62 | Incenp | Inner centromere protein (Mus musculus) | 132.98 | 3.18 | 101209 | 2 | 849047.31 | 434374.97 | 530157.56 | 401946.78 | 378250.81 | 635897.81 | 485362.91 | 621384.06 | 606166 | 571257.69 | 6 | **1.126** |
| Q9WU78 | Pdcd6ip | Programmed cell death 6-interacting protein (Mus musculus) | 1378.76 | 36.82 | 96024 | 21 | 41407172 | 34016476 | 35994172 | 35888332 | 40063312 | 43941256 | 45209812 | 43492868 | 46448048 | 41724260 | 0 | **1.179** |
| Q9WU81 | Slc37a2 | Glucose-6-phosphate exchanger SLC37A2 (Mus musculus) | 177.56 | 7.78 | 55073 | 3 | 2638900.75 | 2220164.5 | 2914908.25 | 1110559.88 | 1502380.5 | 4086873 | 3152658.5 | 3165144.25 | 2235582.5 | 2979446.75 | 2 | **1.504** |
| Q9WU84 | Ccs | Copper chaperone for superoxide dismutase (Mus musculus) | 56.08 | 4.74 | 28912 | 1 | 0 | 0 | 1781565.75 | 0 | 1984972.88 | 1421105.38 | 1527911.5 | 1449454.12 | 1536184.38 | 1796871.62 | 12 | **2.053** |
| Q9WUA2 | Farsb | Phenylalanine--tRNA ligase beta subunit (Mus musculus) | 740.29 | 29.2 | 65697 | 15 | 31682156 | 29414728 | 24800750 | 34367896 | 31492170 | 29916256 | 28489446 | 32701374 | 35032308 | 34183616 | 9 | **1.056** |
| Q9WUA3 | Pfkp | ATP-dependent 6-phosphofructokinase, platelet type (Mus musculus) | 1988.73 | 51.79 | 85455 | 29 | 77172800 | 74244264 | 86133736 | 72855352 | 78196120 | 81613168 | 82788176 | 92388288 | 87926152 | 96642040 | 2 | **1.136** |
| Q9WUD1 | Stub1 | STIP1 homology and U box-containing protein 1 (Mus musculus) | 57.84 | 3.95 | 34909 | 1 | 1781134 | 0 | 0 | 2208738.25 | 1712180.38 | 1339463.25 | 1526290.25 | 1476697.5 | 2125022.25 | 1912921.88 | 12 | **1.470** |
| Q9WUD8 | Faim | Fas apoptotic inhibitory molecule 1 (Mus musculus) | 63.72 | 9.5 | 20201 | 1 | 0 | 0 | 411653.62 | 244695.64 | 153465.88 | 217410.39 | 227658.89 | 393590.19 | 214307.61 | 319945.12 | 9 | **1.695** |

| | | | | | | | | | | | | | | | | | | |
|---|---|---|---|---|---|---|---|---|---|---|---|---|---|---|---|---|---|---|
| Q9WUK4 | Rfc2 | Replication factor C subunit 2 (Mus musculus) | 80.96 | 4.87 | 38725 | 2 | 7796811 | 6072110.5 | 5900851.5 | 4796432 | 4840054 | 4428107 | 5701688.5 | 4409298 | 4515534.5 | 2760148 | 2 | **0.742** |
| Q9WUL7 | Arl3 | ADP-ribosylation factor-like protein 3 (Mus musculus) | 329.41 | 51.1 | 20487 | 6 | 9346350 | 6420690.5 | 7647848 | 5604276.5 | 4973434 | 7798498.5 | 7978065 | 8340991 | 5444963 | 6765640 | 9 | **1.069** |
| Q9WUM4 | Coro1c | Coronin-1C (Mus musculus) | 775.59 | 37.97 | 53121 | 12 | 46038756 | 30460538 | 29837682 | 26612000 | 31497052 | 28338740 | 28284206 | 36741536 | 32290304 | 36802912 | 11 | **0.988** |
| Q9WUM5 | Suclg1 | Succinate--CoA ligase [ADP/GDP-forming] subunit alpha, mitochondrial (Mus musculus) | 119.17 | 7.8 | 36155 | 2 | 3597614.75 | 3229985.25 | 2949982.75 | 2296672 | 2448063.75 | 2712947 | 3006589 | 2351977.75 | 1900972 | 2332497 | 7 | **0.847** |
| Q9WUN2 | Tbk1 | Serine/threonine-protein kinase TBK1 (Mus musculus) | 48.98 | 1.65 | 83425 | 1 | 0 | 1035709.75 | 976211.19 | 1072336 | 668137.62 | 0 | 654372.19 | 677453.25 | 520362.91 | 760331.25 | 7 | **0.696** |
| Q9WUP7 | Uchl5 | Ubiquitin carboxyl-terminal hydrolase isozyme L5 (Mus musculus) | 310.62 | 23.71 | 37617 | 7 | 22637658 | 31603516 | 23387700 | 25072202 | 21982862 | 22643896 | 14510080 | 25307856 | 24624936 | 25603026 | 12 | **0.904** |
| Q9WUQ2 | Preb | Prolactin regulatory element-binding protein (Mus musculus) | 102.65 | 6 | 45437 | 2 | 2292041.5 | 2832547.25 | 1869848.88 | 2217876 | 2429296.5 | 2057161.75 | 2186933 | 2226289.5 | 2251017.75 | 2367953.75 | 9 | **0.953** |
| Q9WUR2 | Eci2 | Enoyl-CoA delta isomerase 2, mitochondrial (Mus musculus) | 121.4 | 7.16 | 43268 | 3 | 1402203.25 | 4053156.25 | 3690132.75 | 4089951 | 2351925 | 2537036.25 | 2564425.25 | 2155126.75 | 3403484.5 | 2488285.75 | 9 | **0.844** |
| Q9WUT3 | Rps6ka2 | Ribosomal protein S6 kinase alpha-2 (Mus musculus) | 166.93 | 4.5 | 83157 | 3 | 1757934.25 | 3131786.5 | 3227295.75 | 3287222 | 3808104 | 3582409.5 | 1894869.75 | 3768915 | 3736732.25 | 3256384.25 | 9 | **1.068** |
| Q9WUU7 | Ctsz | Cathepsin Z (Mus musculus) | 586.35 | 42.16 | 33996 | 10 | 73740048 | 74503336 | 68969760 | 49477696 | 83006416 | 48859664 | 59426240 | 46864256 | 60971288 | 59359336 | 3 | **0.788** |
| Q9WUZ9 | Entpd5 | Ectonucleoside triphosphate diphosphohydrolase 5 (Mus musculus) | 45.01 | 3.04 | 47102 | 1 | 0 | 162123.47 | 246982.64 | 220669.03 | 37024.22 | 340088.25 | 0 | 254413.27 | 293273.34 | 263564.62 | 5 | **1.727** |
| Q9WV32 | Arpc1b | Actin-related protein 2/3 complex subunit 1B (Mus musculus) | 812.53 | 50.54 | 41064 | 14 | 71265616 | 105961488 | 90702336 | 105319424 | 84658800 | 78562768 | 89937560 | 106455072 | 95819952 | 108193312 | 9 | **1.046** |
| Q9WV54 | Asah1 | Acid ceramidase (Mus musculus) | 847.05 | 43.4 | 44670 | 14 | 22002280 | 29657796 | 29237900 | 25148948 | 27813624 | 28954854 | 29788472 | 25463942 | 24569792 | 26520052 | 12 | **1.011** |
| Q9WV55 | Vapa | Vesicle-associated membrane protein-associated protein A (Mus musculus) | 303.56 | 30.92 | 27855 | 6 | 11738999 | 6591822.5 | 7143075 | 5427515.5 | 9483958 | 11860646 | 12774784 | 11703102 | 11516706 | 12580332 | 2 | **1.496** |
| Q9WV68 | Decr2 | Peroxisomal 2,4-dienoyl-CoA reductase (Mus musculus) | 44.48 | 6.16 | 31300 | 1 | 0 | 176232.41 | 187526.45 | 162485.41 | 130814.55 | 0 | 82032.71 | 0 | 0 | 0 | 11 | **0.125** |
| Q9WV85 | Nme3 | Nucleoside diphosphate kinase 3 (Mus musculus) | 50.43 | 10.06 | 19099 | 1 | 1639813.62 | 1126347.38 | 1119909.75 | 453183.41 | 627215.88 | 894008.62 | 1232365.12 | 1154149.88 | 1342716.38 | 1223879.25 | 7 | **1.177** |
| Q9WVA3 | Bub3 | Mitotic checkpoint protein BUB3 (Mus musculus) | 416.91 | 25.77 | 36955 | 8 | 39556276 | 26891598 | 32328436 | 24772508 | 28144384 | 34833860 | 31765832 | 34585056 | 33106406 | 35383752 | 6 | **1.119** |
| Q9WVA4 | Tagln2 | Transgelin-2 (Mus musculus) | 1015.91 | 77.39 | 22395 | 15 | 219596736 | 121886976 | 124870688 | 121706768 | 195559024 | 147615760 | 175019824 | 138340784 | 171364896 | 153781552 | 10 | **1.003** |
| Q9WVE8 | Pacsin2 | Protein kinase C and casein kinase substrate in neurons protein 2 (Mus musculus) | 396.15 | 19.96 | 55833 | 9 | 19663038 | 13931334 | 17467966 | 13708189 | 15898650 | 17557884 | 15428834 | 16815036 | 15238135 | 15917621 | 11 | **1.004** |
| Q9WVJ2 | Psmd13 | 26S proteasome non-ATPase regulatory subunit 13 (Mus musculus) | 762.99 | 34.04 | 42809 | 14 | 23025250 | 29061910 | 27354062 | 35789812 | 30369686 | 32889386 | 36782656 | 30016878 | 35486388 | 31148976 | 5 | **1.142** |
| Q9WVL2 | Stat2 | Signal transducer and activator of transcription 2 (Mus musculus) | 98.03 | 2.6 | 105417 | 2 | 779932.25 | 870098.12 | 797678.81 | 685587.62 | 638842.38 | 822889.88 | 0 | 1042108.5 | 854542.81 | 996668.06 | 7 | **0.985** |
| Q9WVQ5 | Apip | Methylthioribulose-1-phosphate dehydratase (Mus musculus) | 88.25 | 10.37 | 26949 | 2 | 475476.66 | 1667569.38 | 2090216.62 | 2237800.75 | 2442184 | 891456.75 | 632705.19 | 2209511.5 | 2379947.5 | 1822742.12 | 11 | **0.890** |
| Q9WVR4 | Fxr2 | Fragile X mental retardation syndrome-related protein 2 (Mus musculus) | 157.21 | 5.94 | 73743 | 3 | 12231247 | 10918731 | 9237197 | 11777968 | 10948514 | 10561091 | 11311493 | 10839517 | 10301745 | 11565011 | 9 | **0.990** |
| Q9Z0F8 | Adam17 | Disintegrin and metalloproteinase domain-containing protein 17 (Mus musculus) | 193.34 | 3.75 | 93056 | 2 | 210046.67 | 321654.22 | 314867.81 | 990069.75 | 606645.12 | 333180.91 | 998058.81 | 380486.19 | 1076791.38 | 1159691 | 4 | **1.616** |
| Q9Z0H3 | Smarcb1 | SWI/SNF-related matrix-associated actin-dependent regulator of chromatin subfamily B mei | 134.32 | 7.79 | 44141 | 2 | 3788868.75 | 4404790 | 1240414.38 | 1746817.75 | 944807 | 3611123.5 | 4907852 | 4476786.5 | 1390909.38 | 936575.94 | 10 | **1.264** |
| Q9Z0J0 | Npc2 | NPC intracellular cholesterol transporter 2 (Mus musculus) | 179.96 | 32.21 | 16442 | 4 | 17660444 | 7979192.5 | 10145604 | 6587539 | 16196390 | 15609922 | 17012678 | 12950448 | 15495765 | 13239047 | 9 | **1.269** |
| Q9Z0K8 | Vnn1 | Pantetheinase (Mus musculus) | 75.81 | 2.54 | 57091 | 1 | 3495649.5 | 5063249.5 | 5366602.5 | 6489542.5 | 5733571 | 2895996 | 2851758.5 | 3005498.5 | 3336449.75 | 3867966.25 | 1 | **0.610** |
| Q9Z0M5 | Lipa | Lysosomal acid lipase/cholesteryl ester hydrolase (Mus musculus) | 386.73 | 21.66 | 45325 | 7 | 13355938 | 18441054 | 18773882 | 18274772 | 14964363 | 27717948 | 22709558 | 28635640 | 22002850 | 25241798 | 0 | **1.507** |
| Q9Z0N1 | Eif2s3x | Eukaryotic translation initiation factor 2 subunit 3, X-linked (Mus musculus) | 765.82 | 40.04 | 51065 | 13 | 50434692 | 37313924 | 45814052 | 35639676 | 44954236 | 49761056 | 40586584 | 41119552 | 43165720 | 42696140 | 12 | **1.015** |
| Q9Z0P5 | Twf2 | Twinfilin-2 (Mus musculus) | 366.84 | 28.65 | 39471 | 7 | 8593346 | 10991893 | 8334126 | 10188249 | 8097551.5 | 11681725 | 8461992 | 9132221 | 10169220 | 9333715 | 9 | **1.056** |
| Q9Z0R0 | Haspin | Serine/threonine-protein kinase haspin (Mus musculus) | 42.08 | 2.92 | 84181 | 1 | 0 | 120372.14 | 149867.03 | 125962.27 | 100606.53 | 0 | 0 | 180062.44 | 162054.2 | 181230.61 | 10 | **1.053** |
| Q9Z0S1 | Bpnt1 | 3'(2'),5'-bisphosphate nucleotidase 1 (Mus musculus) | 648.47 | 52.6 | 33196 | 12 | 25054600 | 18933462 | 22150696 | 17078128 | 17698046 | 23080272 | 22407744 | 23574058 | 21439856 | 21048648 | 7 | **1.105** |
| Q9Z0U1 | Tjp2 | Tight junction protein ZO-2 (Mus musculus) | 100.42 | 2.49 | 131280 | 2 | 1335806.88 | 864205.75 | 972634.25 | 0 | 789660.12 | 1199705.25 | 1153831.62 | 1131629 | 992067.5 | 1646032.62 | 4 | **1.545** |
| Q9Z0W3 | Nup160 | Nuclear pore complex protein Nup160 (Mus musculus) | 149.83 | 3.28 | 158232 | 3 | 657622.5 | 1641126.62 | 1700536.88 | 1692152 | 1462990.25 | 1753829.75 | 1579297.25 | 1683707.5 | 1585964 | 1153925.88 | 12 | **1.084** |
| Q9Z0X1 | Aifm1 | Apoptosis-inducing factor 1, mitochondrial (Mus musculus) | 533.22 | 27.45 | 66765 | 11 | 12972144 | 12460118 | 14397180 | 10578283 | 11288768 | 13986777 | 12359488 | 12110226 | 12650192 | 13090325 | 10 | **1.041** |
| Q9Z110 | Aldh18a1 | Delta-1-pyrroline-5-carboxylate synthase (Mus musculus) 1 | 797.96 | 17.61 | 87266 | 13 | 26337372 | 24590678 | 25065040 | 22254740 | 24552480 | 29585924 | 25806250 | 31074016 | 25646900 | 26752596 | 2 | **1.131** |
| Q9Z120 | Mettl1 | tRNA (guanine-N(7)-)-methyltransferase (Mus musculus) | 72.72 | 5.6 | 30603 | 1 | 491449.72 | 225561.08 | 314293.44 | 388731.91 | 295971.38 | 437277.5 | 399582.03 | 309335.44 | 348067.69 | 573750.06 | 7 | **1.205** |
| Q9Z127 | Slc7a5 | Large neutral amino acids transporter small subunit 1 (Mus musculus) | 91.67 | 8.79 | 55872 | 2 | 0 | 1602384.5 | 1672532.12 | 0 | 1789032.62 | 1878005 | 1792639.12 | 1716709.5 | 1746570.75 | 1790454.12 | 3 | **1.762** |
| Q9Z160 | Cog1 | Conserved oligomeric Golgi complex subunit 1 (Mus musculus) | 46.96 | 0.92 | 109050 | 1 | 0 | 516803.44 | 0 | 578075.31 | 408390.5 | 0 | 0 | 630212.62 | 514662.88 | 536092.25 | 14 | **1.118** |
| Q9Z1B5 | Mad2l1 | Mitotic spindle assembly checkpoint protein MAD2A (Mus musculus) | 154.21 | 18.05 | 23598 | 3 | 0 | 321352.47 | 510880.75 | 222879.86 | 284181.12 | 58859.25 | 61740.68 | 452996.25 | 381092.22 | 458965.47 | 11 | **1.056** |
| Q9Z1D1 | Eif3g | Eukaryotic translation initiation factor 3 subunit G (Mus musculus) | 378.16 | 21.88 | 35638 | 7 | 23513204 | 18246612 | 21946648 | 16837184 | 18631732 | 19364452 | 20656704 | 18976788 | 18854994 | 19927332 | 10 | **0.986** |
| Q9Z1F9 | Uba2 | SUMO-activating enzyme subunit 2 (Mus musculus) | 1155.66 | 46.87 | 70569 | 18 | 16652275 | 32117038 | 26856532 | 27842934 | 23964792 | 25869388 | 22345746 | 31668070 | 28631566 | 29680908 | 10 | **1.084** |
| Q9Z1I6 | Arhgef1 | Rho guanine nucleotide exchange factor 1 (Rattus norvegicus) | 491.74 | 11.64 | 102597 | 10 | 9408666 | 7694599 | 6471443 | 7506225.5 | 7306608 | 8040926 | 7565349 | 7816112 | 7450435 | 7447503 | 10 | **0.998** |
| Q9Z1K5 | Arih1 | E3 ubiquitin-protein ligase ARIH1 (Mus musculus) | 60.16 | 2.52 | 64017 | 1 | 694813.19 | 1923876.25 | 2483447.25 | 2492661.25 | 2162455.25 | 1556262.38 | 2116802 | 1678485.5 | 1775833.25 | 2288459.75 | 8 | **0.965** |
| Q9Z1Q9 | Vars | Valine--tRNA ligase (Mus musculus) | 2193.35 | 37.93 | 140215 | 36 | 98429824 | 97584064 | 94710288 | 94464112 | 93805248 | 103976656 | 94000536 | 98386752 | 97818504 | 101386136 | 6 | **1.035** |
| Q9Z1T1 | Ap3b1 | AP-3 complex subunit beta-1 (Mus musculus) | 199.31 | 3.08 | 122740 | 3 | 5958597.5 | 3389056 | 3711203.5 | 3076624.5 | 2561259 | 5662181 | 3290883.25 | 4434419.5 | 4506315.5 | 4543937 | 7 | **1.200** |
| Q9Z1X4 | Ilf3 | Interleukin enhancer-binding factor 3 (Mus musculus) | 295.89 | 6.46 | 96021 | 5 | 7515828 | 5206334 | 6346688 | 5527725 | 5694356 | 6279394.5 | 7629831 | 6708191 | 6779538 | 6988130 | 5 | **1.135** |
| Q9Z1Z0 | Uso1 | General vesicular transport factor p115 (Mus musculus) | 421.16 | 11.37 | 106983 | 8 | 8076696.5 | 8716710 | 6908912.5 | 11338782 | 8634423 | 8536984 | 6770254 | 9889918 | 11206920 | 11465211 | 10 | **1.096** |
| Q9Z204 | Hnrnpc | Heterogeneous nuclear ribonucleoproteins C1/C2 (Mus musculus) | 1064.68 | 41.21 | 34385 | 17 | 84492512 | 100814048 | 94382184 | 115689976 | 98046664 | 74422008 | 76320984 | 91827688 | 95856672 | 93505848 | 4 | **0.875** |
| Q9Z277 | Baz1b | Tyrosine-protein kinase BAZ1B (Mus musculus) | 278.83 | 5.07 | 170651 | 6 | 3245889 | 4332817.5 | 5687674.5 | 4080735.5 | 2766526.25 | 2559054.5 | 3590606.75 | 3027553 | 3856106.5 | 5117005 | 9 | **0.902** |
| Q9Z2D0 | Mtmr9 | Myotubularin-related protein 9 (Mus musculus) | 86.45 | 3.49 | 62908 | 2 | 651688.62 | 1816891 | 1511830.25 | 887704.19 | 1344762 | 1578181.75 | 1019102.44 | 1495410.25 | 760534.88 | 1664802 | 11 | **1.049** |
| Q9Z2D1 | Mtmr2 | Myotubularin-related protein 2 (Mus musculus) | 42.64 | 1.09 | 73232 | 1 | 0 | 949016.88 | 717365.56 | 0 | 824028.19 | 0 | 0 | 722892.94 | 779624.56 | 579112 | 13 | **0.836** |
| Q9Z2D8 | Mbd3 | Methyl-CpG-binding domain protein 3 (Mus musculus) | 132.68 | 8.77 | 32168 | 2 | 1187487.25 | 1583757.88 | 1459949.5 | 1734008.25 | 902228.5 | 596785.38 | 1639489.88 | 1451947.12 | 1982749.5 | 1768411.25 | 9 | **1.083** |
| Q9Z2G9 | Htatip2 | Oxidoreductase HTATIP2 (Mus musculus) | 273.41 | 30.58 | 26870 | 7 | 10074734 | 11572392 | 12642872 | 7915706.5 | 7561492 | 12644772 | 15085542 | 14027538 | 13727557 | 14747075 | 0 | **1.411** |
| Q9Z2I8 | Suclg2 | Succinate--CoA ligase [GDP-forming] subunit beta, mitochondrial (Mus musculus) | 595.09 | 27.25 | 46840 | 8 | 16302236 | 20646682 | 17773196 | 26895016 | 12970909 | 15280318 | 18182646 | 17674902 | 13332263 | 13385020 | 8 | **0.823** |
| Q9Z2I9 | Sucla2 | Succinate--CoA ligase [ADP-forming] subunit beta, mitochondrial (Mus musculus) | 537.23 | 22.89 | 50114 | 9 | 9677398 | 12043428 | 12390471 | 10690346 | 10718453 | 12642059 | 10882806 | 10852793 | 11084569 | 11319672 | 8 | **1.023** |
| Q9Z2L0 | Vdac1 | Voltage-dependent anion-selective channel protein 1 (Rattus norvegicus) | 1436.91 | 77.74 | 30756 | 18 | 50192508 | 89414704 | 95325312 | 93019360 | 75711008 | 80267736 | 68464728 | 89079936 | 86142424 | 88292056 | 9 | **1.021** |
| Q9Z2L6 | Minpp1 | Multiple inositol polyphosphate phosphatase 1 (Mus musculus) | 56.73 | 2.29 | 54537 | 1 | 659025.69 | 0 | 360120.34 | 387678.59 | 0 | 553166.25 | 592045.38 | 420151.44 | 686769.5 | 354800.03 | 7 | **1.853** |
| Q9Z2L7 | Crlf3 | Cytokine receptor-like factor 3 (Mus musculus) | 215.99 | 16.06 | 49559 | 4 | 2884265.5 | 3588812.5 | 3414358.5 | 2062919.25 | 4562605.5 | 3682122.5 | 3759091 | 5390304.5 | 5643194.5 | 4934821 | 2 | **1.418** |
| Q9Z2M7 | Pmm2 | Phosphomannomutase 2 (Mus musculus) | 41.44 | 3.31 | 27657 | 1 | 774851.81 | 1096833.25 | 992341.88 | 1236871.75 | 1155678.25 | 921230.25 | 885841.88 | 942557.31 | 1091279.5 | 1024551.25 | 7 | **0.926** |
| Q9Z2N8 | Actl6a | Actin-like protein 6A (Mus musculus) | 572.78 | 33.8 | 47448 | 10 | 17041994 | 19245460 | 21941654 | 14642392 | 18593930 | 18378306 | 17216262 | 15058089 | 19784556 | 19199262 | 12 | **0.980** |
| Q9Z2U0 | Psma7 | Proteasome subunit alpha type-7 (Mus musculus) | 483.45 | 40.73 | 27855 | 8 | 44097392 | 55815600 | 54892144 | 55523936 | 63810272 | 66986600 | 57659824 | 53848136 | 56544680 | 56251868 | 7 | **1.063** |
| Q9Z2U1 | Psma5 | Proteasome subunit alpha type-5 (Mus musculus) | 531.54 | 47.3 | 26411 | 8 | 51317544 | 64463504 | 51708608 | 60438224 | 54156184 | 56614004 | 50971124 | 48048052 | 51094324 | 42165952 | 3 | **0.882** |
| Q9Z2W0 | Dnpep | Aspartyl aminopeptidase (Mus musculus) | 185.68 | 12.47 | 52207 | 4 | 11569765 | 3998574 | 4925197 | 3258869 | 4335763.5 | 12330895 | 13040008 | 11082149 | 9913981 | 9701616 | 3 | **1.996** |
| Q9Z2X2 | Psmd10 | 26S proteasome non-ATPase regulatory subunit 10 (Mus musculus) | 66.83 | 3.46 | 25084 | 1 | 1695794.5 | 1834463.38 | 1405376.75 | 2033275.5 | 2021015.5 | 1270029 | 1214131.38 | 1254115.62 | 1814389.88 | 1493670.12 | 3 | **0.784** |
| Q9Z2Y8 | Plpbp | Pyridoxal phosphate homeostasis protein (Mus musculus) | 52.02 | 4.74 | 30049 | 1 | 1517642.12 | 1228206.12 | 1298476.12 | 403055.5 | 1372549 | 1764501.5 | 1258098.88 | 1380296.62 | 1152033 | 1280507.38 | 11 | **1.174** |
| Q9Z315 | Sart1 | U4/U6.U5 tri-snRNP-associated protein 1 (Mus musculus) | 288.52 | 9.43 | 90885 | 5 | 5902484 | 3599132.25 | 4329052 | 3237930 | 3015247.5 | 4391996.5 | 4750546.5 | 4476887 | 4287346.5 | 4133126 | 7 | **1.097** |
| Q9Z330 | Dnmt1 | DNA (cytosine-5)-methyltransferase 1 (Rattus norvegicus) | 815.81 | 11.47 | 182774 | 15 | 24731732 | 20373776 | 21012054 | 21142584 | 20350292 | 22354316 | 19247906 | 22457526 | 21542494 | 21931658 | 9 | **0.999** |
| | | | | | | | 134965689929 | 126808415358 | 130110388306 | 123735532133 | 126815581159 | 135324373152 | 125446246555 | 129677428277 | 130774370580 | 131838874368 | | |
| | | | | | | | | | 128487121377 | | | | | 130612258586 | | | | **#DIV/0 !** |
