## supplemental Table 2 for "Biobased, Biodegradable but not bio-neutral: about the effects of polylactic acid nanoparticles on macrophages"

Supplementary table 2: list of modulated proteins, selected by the Mann Whitney U test (p<0.05)

| acc number | gene_name | description | protein_set_score | coverage | MW | #peptides | abund._Ctrl1 | abund._ctrl2 | abund._ctrl3 | abund._ctrl4 | abund._ctrl5 | abund._PLA1 | abund._PLA2 | abund._PLA3 | abund._PLA4 | abund._PLA5 | U PLA | ratio PLA/ctrl |
|---|---|---|---|---|---|---|---|---|---|---|---|---|---|---|---|---|---|---|
| P35821 | Ptpn1 | Tyrosine-protein phosphatase non-receptor type 1 (Mus musculus) | 319.23 | 16.67 | 49593 | 6 | 6173386.5 | 5347889 | 4585074 | 4436308 | 4585390 | 9322156 | 9237753 | 8835321 | 8582800 | 7997213 | 0 | 1.7500461586 |
| P10852 | Slc3a2 | 4F2 cell-surface antigen heavy chain (Mus musculus) | 684.55 | 29.85 | 58337 | 12 | 35131460 | 29372646 | 33723908 | 32223972 | 31917884 | 43692020 | 38452636 | 42802300 | 42271232 | 42973972 | 0 | 1.2945268725 |
| A1L314 | Mpeg1 | Macrophage-expressed gene 1 protein (Mus musculus) | 922.75 | 40.81 | 78390 | 17 | 41420120 | 38384404 | 40366676 | 35873584 | 34609164 | 47871508 | 52164452 | 53467204 | 46966720 | 50276012 | 0 | 1.3151885845 |
| P97858 | Slc35b1 | Solute carrier family 35 member B1 (Mus musculus) | 67.6 | 3.11 | 35883 | 1 | 960815.94 | 1012539.81 | 1057267.25 | 1053471.38 | 909420.5 | 1597779.88 | 1334737.62 | 1345261.5 | 1467780.12 | 1394650 | 0 | 1.429896434 |
| P70248 | Myo1f | Unconventional myosin-If (Mus musculus) | 491.15 | 8.55 | 125947 | 8 | 8799587 | 9474237 | 9687511 | 9019736 | 9407401 | 11333380 | 11035171 | 12668711 | 11679069 | 11706360 | 0 | 1.2594226212 |
| Q9JII5 | Dazap1 | DAZ-associated protein 1 (Mus musculus) | 178.18 | 11.33 | 43214 | 3 | 3006523 | 3545635.5 | 3686293.5 | 3414365.5 | 3082283 | 1875162 | 2452774.5 | 2431534.25 | 2146500 | 1748794.88 | 0 | 0.6366717445 |
| O08992 | Sdcbp | Syntenin-1 (Mus musculus) | 224.11 | 17.73 | 32379 | 3 | 1817779 | 1783844 | 2342503.25 | 831902.31 | 1628373.88 | 4090364.75 | 4056952 | 3978907 | 5898522 | 4764695.5 | 0 | 2.7116075667 |
| P47968 | Rpia | Ribose-5-phosphate isomerase (Mus musculus) | 164.85 | 10.89 | 32451 | 4 | 2552192.25 | 3596075.5 | 3591613.5 | 3679499.5 | 2910767.5 | 1036262.75 | 1257973.62 | 1216294 | 1012087.88 | 1490179.5 | 0 | 0.3682022758 |
| Q8K1B8 | Fermt3 | Fermitin family homolog 3 (Mus musculus) | 1802.91 | 57.29 | 75635 | 28 | 97356984 | 97559296 | 116614800 | 101618288 | 99615688 | 125796464 | 129982936 | 133540384 | 131254784 | 138248240 | 0 | 1.2848434196 |
| P41105 | Rpl28 | 60S ribosomal protein L28 (Mus musculus) | 363.84 | 45.99 | 15733 | 8 | 75073712 | 56685020 | 80793944 | 51264476 | 73146128 | 103587728 | 103633664 | 110740744 | 121299488 | 105383216 | 0 | 1.6163329132 |
| E9Q634 | Myo1e | Unconventional myosin-Ie (Mus musculus) | 854.97 | 18.43 | 126818 | 16 | 17856690 | 15635516 | 19211716 | 13847555 | 16946492 | 23137486 | 21423142 | 24289554 | 23303390 | 24706076 | 0 | 1.3995507843 |
| P24668 | M6pr | Cation-dependent mannose-6-phosphate receptor (Mus musculus) | 323.25 | 24.82 | 31172 | 4 | 9403911 | 11008829 | 10391964 | 12275437 | 11470308 | 4984128.5 | 7456842 | 7402903.5 | 7188308 | 7946741 | 0 | 0.6412215415 |
| P20108 | Prdx3 | Thioredoxin-dependent peroxide reductase, mitochondrial (Mus musculus) | 386.69 | 36.19 | 28127 | 7 | 32942818 | 30321688 | 28302956 | 33121882 | 30182782 | 36242400 | 38828284 | 36547288 | 36405884 | 39131900 | 0 | 1.2084534566 |
| P97822 | Anp32e | Acidic leucine-rich nuclear phosphoprotein 32 family member E (Mus musculus) | 589.8 | 30.38 | 29622 | 7 | 21962734 | 24678106 | 28632266 | 18881974 | 26902646 | 38945192 | 32865268 | 37635564 | 36352756 | 36551836 | 0 | 1.5063112618 |
| Q9JLC8 | Sacs | Sacsin (Mus musculus) | 61.36 | 0.35 | 520685 | 2 | 11963284 | 11789488 | 10390196 | 12975868 | 12502982 | 8051224.5 | 7975492 | 4157240.25 | 7762141 | 6517474 | 0 | 0.5780362442 |
| P60843 | Eif4a1 | Eukaryotic initiation factor 4A-I (Mus musculus) | 2306.16 | 71.92 | 46154 | 33 | 441859424 | 420403840 | 459000864 | 436052512 | 440689184 | 498719680 | 485018848 | 472670848 | 475270624 | 475272576 | 0 | 1.0950619647 |
| P11440 | Cdk1 | Cyclin-dependent kinase 1 (Mus musculus) | 764.52 | 49.16 | 34107 | 14 | 22547130 | 17055754 | 21492424 | 15336100 | 21691624 | 25143346 | 29980882 | 30479112 | 30259872 | 29939620 | 0 | 1.4859185354 |
| Q8R180 | Ero1a | ERO1-like protein alpha (Mus musculus) | 988.1 | 42.24 | 54084 | 16 | 38574724 | 40042600 | 40335780 | 35194888 | 40161032 | 47505636 | 42997684 | 44930304 | 45186976 | 45939628 | 0 | 1.1659789306 |
| Q8BIJ7 | Rufy1 | RUN and FYVE domain-containing protein 1 (Mus musculus) | 120.72 | 4.35 | 80376 | 3 | 3199382.25 | 2563904.5 | 2724615.75 | 2684632 | 3021813 | 1905232.75 | 2436750 | 1846024.5 | 1654439.75 | 1865970.75 | 0 | 0.6839636517 |
| Q9JK23 | Psmg1 | Proteasome assembly chaperone 1 (Mus musculus) | 288.67 | 20.07 | 33104 | 5 | 8831279 | 6946882.5 | 7451736 | 6993773 | 7357287.5 | 11400424 | 9181466 | 10070964 | 10222900 | 9556267 | 0 | 1.3419567697 |
| Q9JKP5 | Mbnl1 | Muscleblind-like protein 1 (Mus musculus) | 250.37 | 21.99 | 36976 | 4 | 15648503 | 14622969 | 12562127 | 12871019 | 13943086 | 11761713 | 9526042 | 10140380 | 10345586 | 9675524 | 0 | 0.7387069788 |
| Q3UEB3 | Puf60 | Poly(U)-binding-splicing factor PUF60 (Mus musculus) | 790.96 | 33.33 | 60249 | 13 | 24548658 | 26645376 | 26608820 | 20076234 | 26007418 | 34078100 | 30628748 | 33765948 | 32338582 | 33656128 | 0 | 1.3275659417 |
| P63004 | Pafah1b1 | Platelet-activating factor acetylhydrolase IB subunit alpha (Rattus norvegicus) 1 | 607.97 | 36.1 | 46670 | 12 | 24098346 | 22178674 | 26668980 | 20817088 | 23883970 | 35999868 | 28456912 | 32016546 | 29843050 | 32858630 | 0 | 1.3529875605 |
| Q9Z0M5 | Lipa | Lysosomal acid lipase/cholesteryl ester hydrolase (Mus musculus) | 386.73 | 21.66 | 45325 | 7 | 13355938 | 18441054 | 18773882 | 18274772 | 14964363 | 27717948 | 22709558 | 28635640 | 22002850 | 25241798 | 0 | 1.5070729082 |
| G5E829 | Atp2b1 | Plasma membrane calcium-transporting ATPase 1 (Mus musculus) | 671.03 | 13.93 | 134747 | 12 | 7816870 | 7632992 | 8968369 | 8379600.5 | 8474393 | 10707785 | 8999851 | 10356291 | 10555959 | 10510940 | 0 | 1.2388677039 |
| O35343 | Kpna4 | Importin subunit alpha-3 (Mus musculus) | 501.02 | 25.91 | 57923 | 9 | 11019406 | 12167423 | 11949402 | 11075872 | 9863869 | 17805050 | 16195701 | 15403492 | 15043157 | 13494040 | 0 | 1.3899257957 |
| P62274 | Rps29 | 40S ribosomal protein S29 (Mus musculus) | 90.72 | 32.14 | 6677 | 2 | 2440522.75 | 894117 | 1911949.5 | 1211972.5 | 1812306.12 | 3590463.25 | 3193292.5 | 3789460.5 | 3318915.5 | 3244327.25 | 0 | 2.0719057866 |
| Q64232 | Tecr | Very-long-chain enoyl-CoA reductase (Rattus norvegicus) | 192.23 | 9.42 | 36123 | 3 | 3404840.75 | 2697328.5 | 4928238 | 3056768.25 | 4489798.5 | 6918285 | 5046014 | 8107817 | 7018635.5 | 7191101 | 0 | 1.8453948689 |
| Q64568 | Atp2b3 | Plasma membrane calcium-transporting ATPase 3 (Rattus norvegicus) | 460.35 | 7.15 | 138559 | 8 | 4257239.5 | 4162519.75 | 4968704 | 5302598 | 4838082 | 6718033 | 5358854.5 | 6585834.5 | 6855326 | 7000164 | 0 | 1.3820397817 |
| Q9D7S7 | Rpl22l1 | 60S ribosomal protein L22-like 1 (Mus musculus) | 313.79 | 45.08 | 14467 | 4 | 31242984 | 30091402 | 38768304 | 31085326 | 31636140 | 43451032 | 41273304 | 43880660 | 42499220 | 44173108 | 0 | 1.3221461071 |
| Q60875 | Arhgef2 | Rho guanine nucleotide exchange factor 2 (Mus musculus) | 748.72 | 18.68 | 111974 | 14 | 21338886 | 21217498 | 23995772 | 24383498 | 23428208 | 28929866 | 26329734 | 26858584 | 26179222 | 26262888 | 0 | 1.1765980236 |
| P50516 | Atp6v1a | V-type proton ATPase catalytic subunit A (Mus musculus) | 2251.45 | 66.45 | 68326 | 34 | 123593824 | 120118504 | 131405120 | 106950920 | 107660016 | 142176208 | 144122560 | 145873040 | 138922928 | 158581376 | 0 | 1.2373087879 |
| Q61735 | Cd47 | Leukocyte surface antigen CD47 (Mus musculus) | 199.18 | 11.22 | 33098 | 3 | 1856878.62 | 1360091.12 | 1971137.5 | 1462390 | 1491463 | 2068888.88 | 2321879.25 | 2612556 | 2265504.5 | 2723765.5 | 0 | 1.4729369558 |
| P16036 | Slc25a3 | Phosphate carrier protein, mitochondrial (Rattus norvegicus) | 748.41 | 39.33 | 39445 | 13 | 73843920 | 83044224 | 86774984 | 83256488 | 67842736 | 91299040 | 94364648 | 106143904 | 103554408 | 106365792 | 0 | 1.2709616038 |
| P17439 | Gba | Glucosylceramidase (Mus musculus) | 57.18 | 2.52 | 57622 | 1 | 3171942.75 | 4108049.25 | 3780039.75 | 3426538.75 | 3655244.25 | 2102779.75 | 2411952.5 | 2273608 | 3052751.75 | 2861801.75 | 0 | 0.7001997278 |
| Q9CSH3 | Dis3 | Exosome complex exonuclease RRP44 (Mus musculus) | 151.07 | 3.76 | 108838 | 3 | 2414432.25 | 1820696.25 | 1459494.75 | 1825859.62 | 1918338.75 | 3642321.25 | 2449488.75 | 2863788.75 | 3002330 | 3000953.5 | 0 | 1.5848251882 |
| Q9CQF9 | Pcyox1 | Prenylcysteine oxidase (Mus musculus) | 381.22 | 19.41 | 56495 | 7 | 6892295 | 7626065.5 | 6864704.5 | 8032805 | 7116890.5 | 9268172 | 9239152 | 11645286 | 10430830 | 11689953 | 0 | 1.4308634848 |
| O54692 | Zw10 | Centromere/kinetochore protein zw10 homolog (Mus musculus) | 251.03 | 9.88 | 88063 | 5 | 1520201 | 2221459.5 | 2020323.5 | 1419322.5 | 1411225.12 | 2528023.75 | 2935945.5 | 2750049 | 2643862 | 2636045 | 0 | 1.5704248581 |
| P09581 | Csf1r | Macrophage colony-stimulating factor 1 receptor (Mus musculus) | 601.33 | 17.4 | 109179 | 12 | 21018306 | 18947928 | 22020824 | 16419172 | 16340135 | 24114492 | 25730374 | 25241064 | 32219154 | 28738920 | 0 | 1.4358757088 |
| P98086 | C1qa | Complement C1q subcomponent subunit A (Mus musculus) | 145.06 | 11.84 | 25974 | 2 | 3400937 | 3782547.5 | 3328614.5 | 3221944.75 | 3323420.75 | 3206552.5 | 2664780.5 | 2762225.25 | 2604307.5 | 2690862.5 | 0 | 0.8165767104 |
| Q91VE6 | Nifk | MKI67 FHA domain-interacting nucleolar phosphoprotein (Mus musculus) | 345.08 | 24.61 | 36265 | 6 | 4578400.5 | 3645391.5 | 4830219 | 3293783.5 | 4775198 | 5990912 | 5456316 | 6518318.5 | 5843093 | 5979826.5 | 0 | 1.4102389138 |
| P58252 | Eef2 | Elongation factor 2 (Mus musculus) | 3513.26 | 68.18 | 95314 | 55 | 600991424 | 658130496 | 691897088 | 621285312 | 630809280 | 799924032 | 726322496 | 726290816 | 726312320 | 705636800 | 0 | 1.1502827948 |
| Q9D1R9 | Rpl34 | 60S ribosomal protein L34 (Mus musculus) | 226.42 | 29.06 | 13293 | 5 | 35191500 | 16088982 | 25409756 | 18719594 | 19159812 | 56500032 | 36953464 | 46456224 | 38945148 | 40233272 | 0 | 1.9122704091 |
| P05063 | Aldoc | Fructose-bisphosphate aldolase C (Mus musculus) | 990.22 | 46.28 | 39395 | 17 | 289117024 | 224758496 | 256275712 | 229579776 | 269885632 | 333192608 | 325243232 | 299800960 | 326499136 | 318251200 | 0 | 1.2625757142 |
| P10810 | Cd14 | Monocyte differentiation antigen CD14 (Mus musculus) | 515.2 | 20.49 | 39204 | 7 | 16801956 | 20428604 | 18556168 | 21670858 | 19473342 | 15003135 | 13969148 | 13923828 | 16672033 | 15345096 | 0 | 0.7728517775 |

| | | | | | | | | | | | | | | | | | |
|---|---|---|---|---|---|---|---|---|---|---|---|---|---|---|---|---|---|
| Q9DBS1 | Tmem43 | Transmembrane protein 43 (Mus musculus) | 573.85 | 36.5 | 44783 | 11 | 14442531 | 15168784 | 15548083 | 11202367 | 11085079 | 18130786 | 19455686 | 20175854 | 17130098 | 18406530 | 0 | 1.3832960665 |
| Q9D8Y7 | Tnfaip8l2 | Tumor necrosis factor alpha-induced protein 8-like protein 2 (Mus musculus) l2 | 235.26 | 40.22 | 20615 | 4 | 2701229.75 | 2323800.5 | 3614726.25 | 1120284.38 | 1433721.5 | 3871081.5 | 4704362.5 | 5122376 | 4379149.5 | 4954088.5 | 0 | 2.0574903431 |
| Q6IMY8 | Hnrnpu | Heterogeneous nuclear ribonucleoprotein U (Rattus norvegicus) | 1745.15 | 41.73 | 87732 | 30 | 326900576 | 297482944 | 274565344 | 340285568 | 283495968 | 271463296 | 214358560 | 240446576 | 224878944 | 238956000 | 0 | 0.7815588209 |
| Q9QY76 | Vapb | Vesicle-associated membrane protein-associated protein B (Mus musculus) | 336.04 | 24.28 | 26946 | 6 | 6644623 | 4824924 | 6327744 | 4464134 | 6717954 | 6892839 | 8609200 | 10190532 | 10238523 | 9504947 | 0 | 1.5678749017 |
| B9EJ86 | Osbpl8 | Oxysterol-binding protein-related protein 8 (Mus musculus) | 952.48 | 25.08 | 101269 | 16 | 17439232 | 15670853 | 17732248 | 14771491 | 16407440 | 22990444 | 18855972 | 22578428 | 20475144 | 19126650 | 0 | 1.2682886477 |
| O55101 | Syngr2 | Synaptogyrin-2 (Mus musculus) | 54.27 | 4.46 | 24778 | 1 | 3110087.5 | 4368730.5 | 4182091.75 | 4224840 | 2969239 | 6056100.5 | 5033463.5 | 5334115 | 5359933 | 4792124.5 | 0 | 1.409480369 |
| Q8VEH6 | Cbwd1 | COBW domain-containing protein 1 (Mus musculus) | 89.21 | 5.6 | 43772 | 1 | 281662.62 | 275935.16 | 131691.28 | 300515.5 | 242163.41 | 358930.91 | 425976.59 | 439254.25 | 365226.62 | 414490.66 | 0 | 1.6265674748 |
| P84817 | Fis1 | Mitochondrial fission 1 protein (Rattus norvegicus) | 251.57 | 25.66 | 16995 | 3 | 1330152.25 | 1862384.5 | 1978001.25 | 1624005.62 | 2071632.75 | 2204155.5 | 2245717.25 | 2766192.5 | 2566265.5 | 2547056 | 0 | 1.3906092362 |
| P14685 | Psmd3 | 26S proteasome non-ATPase regulatory subunit 3 (Mus musculus) | 382.69 | 15.47 | 60718 | 7 | 8208220 | 5889570.5 | 9070015 | 9180806 | 10315656 | 13768109 | 11647393 | 12172161 | 11776871 | 11873422 | 0 | 1.4353453039 |
| Q9QZD8 | Slc25a10 | Mitochondrial dicarboxylate carrier (Mus musculus) | 327.72 | 28.92 | 31715 | 5 | 4036830.75 | 5585938 | 5307387.5 | 4369161 | 5219476.5 | 6396479.5 | 6882465 | 6710438 | 5859767 | 6074167 | 0 | 1.3019937614 |
| Q69ZR2 | Hectd1 | E3 ubiquitin-protein ligase HECTD1 (Mus musculus) | 156.91 | 1.6 | 290086 | 3 | 786470.75 | 986330 | 860229.88 | 833191.12 | 1116955.88 | 507970.94 | 771188.88 | 321011.88 | 248066.69 | 227896.09 | 0 | 0.4529901845 |
| Q62376 | Snrnp70 | U1 small nuclear ribonucleoprotein 70 kDa (Mus musculus) | 312.33 | 11.16 | 51992 | 5 | 12611767 | 10111568 | 12514822 | 8689346 | 12138076 | 19475058 | 14854798 | 15433239 | 14003354 | 17928290 | 0 | 1.4571282498 |
| Q8C0M9 | Asrgl1 | Isoaspartyl peptidase/L-asparaginase (Mus musculus) | 158.84 | 10.74 | 33950 | 3 | 4816876 | 5681349 | 6586040.5 | 4252604 | 5410441 | 7222291 | 7147483 | 7047618 | 7526197.5 | 7861725 | 0 | 1.3760379572 |
| P15791 | Camk2d | Calcium/calmodulin-dependent protein kinase type II subunit delta (Rattus norvegicus) | 304.22 | 14.45 | 60081 | 6 | 7907762.5 | 5995462.5 | 8686771 | 5127845.5 | 6703307 | 10411372 | 9885130 | 9753898 | 10177372 | 10862253 | 0 | 1.4842626474 |
| P35564 | Canx | Calnexin (Mus musculus) | 1288.68 | 45.52 | 67278 | 22 | 77678680 | 86358784 | 79166520 | 88791504 | 96543488 | 72865120 | 74561656 | 62470120 | 70893888 | 63852496 | 0 | 0.8042285517 |
| Q9WU78 | Pdcd6ip | Programmed cell death 6-interacting protein (Mus musculus) | 1378.76 | 36.82 | 96024 | 21 | 41407172 | 34016476 | 35994172 | 35888332 | 40063312 | 43941256 | 45209812 | 43492868 | 46448048 | 41724260 | 0 | 1.1785071019 |
| O88455 | Dhcr7 | 7-dehydrocholesterol reductase (Mus musculus) | 117.23 | 7.64 | 53919 | 3 | 653118.44 | 2097323.5 | 1698143.5 | 978678.88 | 437895.47 | 3139201.5 | 2721117.5 | 2518221.5 | 2825251.5 | 2458481.5 | 0 | 2.3293949337 |
| P45376 | Akr1b1 | Aldose reductase (Mus musculus) | 914.68 | 57.28 | 35732 | 17 | 200260416 | 211552176 | 208355568 | 230077408 | 188115200 | 185150976 | 162397440 | 175486128 | 181519920 | 169752496 | 0 | 0.8420069276 |
| Q99KK2 | Cmas | N-acylneuraminate cytidylyltransferase (Mus musculus) | 401.15 | 19.68 | 48058 | 8 | 10323455 | 11775142 | 10251500 | 11759364 | 8481665 | 13510601 | 13728653 | 13276094 | 12482178 | 14187706 | 0 | 1.2775013032 |
| P61087 | Ube2k | Ubiquitin-conjugating enzyme E2 K (Mus musculus) | 421.2 | 53.5 | 22407 | 7 | 12666450 | 14290062 | 13655464 | 13882630 | 12918791 | 9376058 | 11453376 | 9987658 | 12171281 | 11731070 | 0 | 0.8116998317 |
| O70503 | Hsd17b12 | Very-long-chain 3-oxoacyl-CoA reductase (Mus musculus) | 754.77 | 41.35 | 34742 | 13 | 35440676 | 35069836 | 34023676 | 32048882 | 31321556 | 38877088 | 35591736 | 38682968 | 38036008 | 36553204 | 0 | 1.1181407474 |
| P06745 | Gpi | Glucose-6-phosphate isomerase (Mus musculus) | 2093.14 | 61.83 | 62767 | 34 | 240326224 | 205848752 | 242658032 | 221203296 | 244266256 | 265550224 | 252102112 | 270212128 | 279561632 | 261053536 | 0 | 1.1508937761 |
| P47964 | Rpl36 | 60S ribosomal protein L36 (Mus musculus) | 87.03 | 18.1 | 12216 | 2 | 22056376 | 20064696 | 23002440 | 17040904 | 21473388 | 25654976 | 25622186 | 24047528 | 27286908 | 25125134 | 0 | 1.232530284 |
| P13020 | Gsn | Gelsolin (Mus musculus) | 1472.83 | 42.56 | 85942 | 24 | 137309024 | 121233416 | 117093624 | 129330256 | 144686176 | 111846048 | 111196136 | 102997000 | 104405832 | 100769120 | 0 | 0.8176896714 |
| P09405 | Ncl | Nucleolin (Mus musculus) | 2877.89 | 46.39 | 76723 | 48 | 646970880 | 737028160 | 748263296 | 632975744 | 739612416 | 797281856 | 748400128 | 855526400 | 842248256 | 862190976 | 0 | 1.1714187583 |
| P13439 | Umps | Uridine 5'-monophosphate synthase (Mus musculus) | 764.21 | 33.89 | 52292 | 13 | 16793204 | 17240022 | 17915862 | 14988556 | 17422246 | 19153388 | 19969980 | 18963840 | 18851082 | 19160568 | 0 | 1.1391534294 |
| Q9DB77 | Uqcrc2 | Cytochrome b-c1 complex subunit 2, mitochondrial (Mus musculus) | 1097.12 | 50.77 | 48235 | 18 | 58989688 | 47461280 | 55094100 | 48944728 | 46774816 | 64172832 | 61842268 | 63001944 | 60695300 | 64401856 | 0 | 1.2209771004 |
| P24547 | Impdh2 | Inosine-5'-monophosphate dehydrogenase 2 (Mus musculus) | 1099.27 | 30.93 | 55815 | 15 | 74971072 | 89268904 | 74491744 | 95979424 | 80457944 | 64838928 | 60690248 | 60187756 | 63744396 | 63975000 | 0 | 0.7549606583 |
| O88456 | Capns1 | Calpain small subunit 1 (Mus musculus) | 561.39 | 63.94 | 28463 | 10 | 13662510 | 13555646 | 13892038 | 13751943 | 13166243 | 14412270 | 16968294 | 17418950 | 16691464 | 18259780 | 0 | 1.2311149847 |
| Q9ESY9 | Ifi30 | Gamma-interferon-inducible lysosomal thiol reductase (Mus musculus) | 108.44 | 10.89 | 27784 | 2 | 3720531 | 4729702.5 | 3744236.5 | 4117700.25 | 3847611 | 2717665 | 3203230.5 | 3196909 | 3672765.5 | 2868286 | 0 | 0.7767373964 |
| Q4V898 | Rbmx | RNA-binding motif protein, X chromosome (Rattus norvegicus) | 240.57 | 14.1 | 42258 | 5 | 10834805 | 11528639 | 9023996 | 12974274 | 9924338 | 8433307 | 7292068 | 7069242 | 7543161.5 | 8794702 | 0 | 0.7208569984 |
| Q9CQ71 | Rpa3 | Replication protein A 14 kDa subunit (Mus musculus) | 238.23 | 51.24 | 13584 | 4 | 3075260.25 | 2460618.5 | 1729436.38 | 2619752.25 | 1871982 | 3661612 | 3433842.25 | 3251912 | 3553032 | 3497001.25 | 0 | 1.4797419776 |
| Q8C147 | Dock8 | Dedicator of cytokinesis protein 8 (Mus musculus) | 482.36 | 7.05 | 238978 | 12 | 9540007 | 11248918 | 9435130 | 9027333 | 6930637 | 14286656 | 11773972 | 11349385 | 11746994 | 12228385 | 0 | 1.3292052915 |
| O70318 | Epb41l2 | Band 4.1-like protein 2 (Mus musculus) | 362.89 | 10.63 | 109940 | 8 | 7466906 | 7358596.5 | 7394816 | 6191099 | 6093989 | 8208987 | 9529214 | 10044652 | 7998844 | 8041848.5 | 0 | 1.2700486661 |
| P24049 | Rpl17 | 60S ribosomal protein L17 (Rattus norvegicus) | 519.85 | 40.22 | 21397 | 9 | 68248096 | 51718272 | 67720912 | 39396828 | 58072240 | 68284160 | 82296536 | 85728816 | 77283832 | 86099560 | 0 | 1.4016623049 |
| Q8R1V4 | Tmed4 | Transmembrane emp24 domain-containing protein 4 (Mus musculus) | 126.11 | 12.33 | 26022 | 2 | 3208289 | 3900714 | 3612902.25 | 3891101.25 | 3404396.5 | 2762168 | 3146432.5 | 3098104 | 3068116.5 | 3078101.75 | 0 | 0.8410159194 |
| P54001 | P4ha1 | Prolyl 4-hydroxylase subunit alpha-1 (Rattus norvegicus) | 463.4 | 18.73 | 60898 | 8 | 8633017 | 9917150 | 10224831 | 8430589 | 10225026 | 10528331 | 10617057 | 11567671 | 11891528 | 12961898 | 0 | 1.2136989459 |
| P56480 | Atp5f1b | ATP synthase subunit beta, mitochondrial (Mus musculus) | 2673.85 | 79.58 | 56300 | 38 | 404212128 | 406746432 | 499733536 | 423109856 | 465373824 | 504886208 | 502116032 | 528180928 | 517253664 | 542219200 | 0 | 1.179831126 |
| O08795 | Prkcsh | Glucosidase 2 subunit beta (Mus musculus) | 409.18 | 18.23 | 58793 | 7 | 14825971 | 13577900 | 12878836 | 12407479 | 13973062 | 11904046 | 11560019 | 11202170 | 11984042 | 11919059 | 0 | 0.8656004217 |
| P60229 | Eif3e | Eukaryotic translation initiation factor 3 subunit E (Mus musculus) | 1158.44 | 48.09 | 52221 | 20 | 58898800 | 62009772 | 67856560 | 55119684 | 54408684 | 70420104 | 71584424 | 69852352 | 68584512 | 72131216 | 0 | 1.1819654401 |
| P06795 | Abcb1b | Multidrug resistance protein 1B (Mus musculus) | 76.99 | 1.72 | 140994 | 1 | 129495.48 | 119119.99 | 187363.22 | 0 | 109211.33 | 220433.33 | 217494.84 | 293832.41 | 250583.89 | 190829.22 | 0 | 2.1518620058 |
| Q66HG8 | Ik | Protein Red (Rattus norvegicus) | 36.99 | 1.62 | 65588 | 1 | 1032700.75 | 1046689.38 | 811022.12 | 1290334.88 | 730851.81 | 569294.62 | 713990.06 | 549406.06 | 0 | 457031.47 | 0 | 0.4661867221 |
| Q9Z2G9 | Htatip2 | Oxidoreductase HTATIP2 (Mus musculus) | 273.41 | 30.58 | 26870 | 7 | 10074734 | 11572392 | 12642872 | 7915706.5 | 7561492 | 12644772 | 15085542 | 14027538 | 13727557 | 14747075 | 0 | 1.4112204211 |
| P62996 | Tra2b | Transformer-2 protein homolog beta (Mus musculus) | 336.01 | 23.61 | 33666 | 5 | 10031407 | 10442159 | 9484616 | 12879595 | 9426679 | 7578763 | 7777982 | 7569084.5 | 7418938 | 7933221 | 0 | 0.732390451 |
| Q61704 | Itih3 | Inter-alpha-trypsin inhibitor heavy chain H3 (Mus musculus) | 192.29 | 4.05 | 99358 | 4 | 14952218 | 15293322 | 14856794 | 15055238 | 16118957 | 14852712 | 12881658 | 14339444 | 13430063 | 13637845 | 0 | 0.9064613048 |

| | | | | | | | | | | | | | | | | | |
|---|---|---|---|---|---|---|---|---|---|---|---|---|---|---|---|---|---|
| Q8VDP6 | Cdipt | CDP-diacylglycerol--inositol 3-phosphatidyltransferase (Mus musculus) | 226.13 | 24.88 | 23599 | 5 | 4776480 | 4624201.5 | 3179155.25 | 2116387.25 | 1899961.75 | 5643121.5 | 6362668 | 6144849 | 5467401.5 | 5867800.5 | 0 | 1.7766636831 |
| P61028 | Rab8b | Ras-related protein Rab-8B (Mus musculus) | 646.44 | 49.28 | 23603 | 11 | 13176939 | 11049601 | 13994494 | 12992504 | 12936688 | 14158461 | 14862581 | 15294314 | 14070254 | 16447691 | 0 | 1.1665321491 |
| Q9QXK7 | Cpsf3 | Cleavage and polyadenylation specificity factor subunit 3 (Mus musculus) | 147.66 | 7.89 | 77505 | 4 | 5096421 | 4822719 | 4702245.5 | 3713710.75 | 4420068.5 | 5425597 | 5671296.5 | 6035158.5 | 5204604.5 | 5232873.5 | 0 | 1.211572419 |
| D4AE41 | Rbmxl1 | RNA binding motif protein, X-linked-like-1 (Rattus norvegicus) | 354.06 | 19.33 | 42250 | 6 | 9483069 | 13205487 | 8633114 | 14336086 | 9781665 | 7158564.5 | 5755399.5 | 5660710.5 | 6195424.5 | 6983642.5 | 0 | 0.5727646669 |
| Q8VDD5 | Myh9 | Myosin-9 (Mus musculus) | 8025.52 | 59.59 | 226372 | 117 | 732211072 | 690678976 | 684426816 | 683302720 | 743336896 | 676265792 | 616595968 | 648994048 | 673755776 | 645019776 | 0 | 0.9226574743 |
| Q5FWK3 | Arhgap1 | Rho GTPase-activating protein 1 (Mus musculus) | 443.01 | 27.56 | 50411 | 8 | 6215351 | 7722780.5 | 8901978 | 4484900 | 3780669.5 | 10750000 | 9104083 | 10603899 | 9572534 | 10604433 | 0 | 1.6278361581 |
| O08547 | Sec22b | Vesicle-trafficking protein SEC22b (Mus musculus) | 286.61 | 28.84 | 24741 | 5 | 10928426 | 14928759 | 13852077 | 19014588 | 11611427 | 9694480 | 8581584 | 6525659 | 8643225 | 8019364.5 | 0 | 0.5895236966 |
| P56183 | Rrp1 | Ribosomal RNA processing protein 1 homolog A (Mus musculus) | 220.17 | 13.36 | 54777 | 4 | 1719924.88 | 1716124.88 | 2034712.5 | 295834.59 | 421134.38 | 2516586 | 2695452.25 | 2720812.75 | 3089063 | 2650115 | 0 | 2.2095382769 |
| Q3KRC5 | Dus3l | tRNA-dihydrouridine(47) synthase [NAD(P)(+)]-like (Rattus norvegicus) | 124.27 | 3.75 | 71540 | 2 | 993125.88 | 1081720.75 | 887292.62 | 790809.94 | 1037361.62 | 785925.25 | 329268.41 | 646542.44 | 716692.25 | 276667.44 | 0 | 0.5751392549 |
| Q07813 | Bax | Apoptosis regulator BAX (Mus musculus) | 579.63 | 51.56 | 21395 | 8 | 19731658 | 20532910 | 20252940 | 21050764 | 19609354 | 21733714 | 21285334 | 24968738 | 24826328 | 24664158 | 0 | 1.1611091962 |
| Q8CGS4 | Chmp3 | Charged multivesicular body protein 3 (Rattus norvegicus) | 60.1 | 5.38 | 25062 | 1 | 0 | 884905.38 | 1035253.06 | 900928.88 | 772835.38 | 1757502.5 | 1224130.88 | 1680384.62 | 1378109.75 | 1174866.25 | 0 | 2.00755403 |
| Q6PDV7 | Rpl10 | 60S ribosomal protein L10 (Rattus norvegicus) | 538.01 | 35.05 | 24604 | 8 | 45946304 | 27173170 | 39727192 | 18494318 | 41168152 | 55690872 | 56597908 | 56576288 | 54520872 | 55282632 | 0 | 1.6153844281 |
| P11680 | Cfp | Properdin (Mus musculus) | 141.6 | 5.6 | 50327 | 2 | 2010312 | 1652909 | 1698509.88 | 2088846.5 | 2311353.5 | 1347231.25 | 1518248.75 | 1339010.5 | 1528172.25 | 1627252.75 | 0 | 0.7539405462 |
| Q9WTI7 | Myo1c | Unconventional myosin-Ic (Mus musculus) | 1087.13 | 22.58 | 121944 | 19 | 9883394 | 6057990 | 11036076 | 5059040 | 12361700 | 18284760 | 12902204 | 14849386 | 13437869 | 13066853 | 0 | 1.6338741661 |
| P37913 | Lig1 | DNA ligase 1 (Mus musculus) | 567.63 | 16.81 | 102290 | 12 | 14394593 | 11726776 | 13635073 | 9504897 | 10018383 | 15964004 | 15390490 | 16686196 | 14452246 | 15630995 | 0 | 1.317886258 |
| P47199 | Cryz | Quinone oxidoreductase (Mus musculus) | 99.8 | 10.27 | 35269 | 2 | 1223611.12 | 1148962.12 | 1197404.12 | 1110846.38 | 1129542.38 | 1383507.88 | 1938565.12 | 2411163.25 | 2821552.5 | 2705563 | 0 | 1.9379762854 |
| P11152 | Lpl | Lipoprotein lipase (Mus musculus) | 42.24 | 3.59 | 53109 | 1 | 367128.19 | 295677.62 | 310697.94 | 518800.19 | 289952.03 | 278275.28 | 223802.84 | 176484.03 | 120022.11 | 156361.39 | 0 | 0.5358072387 |
| P35601 | Rfc1 | Replication factor C subunit 1 (Mus musculus) | 56.85 | 1.06 | 125985 | 1 | 319207.03 | 436636.66 | 409769.75 | 383541.66 | 319218.44 | 313999.5 | 111147.67 | 229361.92 | 211382.39 | 300486.72 | 0 | 0.6242746298 |
| Q5U4D9 | Thoc6 | THO complex subunit 6 homolog (Mus musculus) | 117.54 | 10.56 | 37315 | 2 | 0 | 349316 | 543202.62 | 579133.44 | 724458.75 | 836785.62 | 781211.94 | 846243.75 | 912779.69 | 1182800.38 | 0 | 2.0763166227 |
| P14115 | Rpl27a | 60S ribosomal protein L27a (Mus musculus) | 280.9 | 22.97 | 16605 | 5 | 104491960 | 61935052 | 92013640 | 47929644 | 107449208 | 145907136 | 123162736 | 125160640 | 119520352 | 125282816 | 0 | 1.5442328692 |
| Q99J09 | Wdr77 | Methylosome protein 50 (Mus musculus) | 100.68 | 4.97 | 36943 | 2 | 1972618.12 | 2881536.25 | 1901881 | 2640365.75 | 2106087.25 | 666330.44 | 1833114.88 | 1600296.88 | 1617394.88 | 1499994.38 | 0 | 0.627440883 |
| Q9ET30 | Tm9sf3 | Transmembrane 9 superfamily member 3 (Mus musculus) | 284.73 | 12.95 | 67545 | 6 | 6335846.5 | 3625885 | 7569219 | 5553265.5 | 1544283.25 | 9203437 | 7731864 | 8603540 | 7734260 | 10971680 | 0 | 1.7964870921 |
| Q9CQT2 | Rbm7 | RNA-binding protein 7 (Mus musculus) | 51.68 | 6.79 | 30148 | 1 | 113288.78 | 218322.41 | 0 | 205124.56 | 135139.28 | 321334.56 | 249859.88 | 269806.84 | 282269.69 | 286990.66 | 0 | 2.0989939602 |
| Q3V0A6 | Gm5592 | Uncharacterized protein C2orf78 homolog (Mus musculus) | 38.24 | 0.82 | 93729 | 1 | 3204431.75 | 2292052 | 3643453.5 | 1887739.38 | 3988163.75 | 4440378.5 | 4255606 | 4982961 | 4012531.5 | 4620919.5 | 0 | 1.4859239267 |
| O09110 | Map2k3 | Dual specificity mitogen-activated protein kinase kinase 3 (Mus musculus) | 205.79 | 14.41 | 39296 | 4 | 3581602.5 | 893733.31 | 3420099.25 | 1810836.25 | 2531529.75 | 4373626 | 3832487.5 | 4156146.25 | 4221544 | 4934146 | 0 | 1.7583183159 |
| P29351 | Ptpn6 | Tyrosine-protein phosphatase non-receptor type 6 (Mus musculus) | 2114.3 | 67.06 | 67559 | 34 | 122960872 | 112083472 | 119935088 | 84941992 | 106584472 | 134939328 | 138383008 | 131659712 | 129615992 | 137961712 | 0 | 1.2306541557 |
| Q921M7 | Fam49b | Protein FAM49B (Mus musculus) | 947.08 | 59.88 | 36776 | 16 | 37333528 | 28721712 | 36465880 | 34161856 | 36094096 | 38169368 | 37350096 | 43558440 | 43241944 | 40122408 | 0 | 1.1716963001 |
| Q9D1A2 | Cndp2 | Cytosolic non-specific dipeptidase (Mus musculus) | 1553.46 | 64.84 | 52767 | 25 | 40843788 | 32846842 | 29729168 | 32993232 | 40777376 | 47492036 | 43132420 | 41806476 | 42391748 | 43153800 | 0 | 1.2301821804 |
| Q78ZA7 | Nap1l4 | Nucleosome assembly protein 1-like 4 (Mus musculus) | 684.12 | 41.87 | 42679 | 11 | 25482166 | 26644142 | 28730476 | 25615648 | 26514372 | 23881436 | 25138950 | 24097952 | 25186942 | 24353602 | 0 | 0.9223387457 |
| P62897 | Cycs | Cytochrome c, somatic (Mus musculus) | 159.35 | 40.95 | 11605 | 4 | 35206976 | 35992556 | 35044396 | 37254512 | 47193508 | 28160890 | 27197172 | 29130132 | 33288050 | 33042912 | 0 | 0.7909046899 |
| Q91YQ5 | Rpn1 | Dolichyl-diphosphooligosaccharide--protein glycosyltransferase subunit 1 (Mus musculus) | 1390.68 | 48.03 | 68528 | 22 | 112556736 | 109514984 | 109770704 | 96547128 | 103479608 | 124444312 | 112816272 | 112678120 | 116713496 | 115937768 | 0 | 1.0953633183 |
| P62751 | Rpl23a | 60S ribosomal protein L23a (Mus musculus) | 395.31 | 41.03 | 17695 | 9 | 49045152 | 46638444 | 35049904 | 83678200 | 53910048 | 24310240 | 32218388 | 21121768 | 28703338 | 23479546 | 0 | 0.4838716242 |
| P61027 | Rab10 | Ras-related protein Rab-10 (Mus musculus) | 452.67 | 43.5 | 22541 | 9 | 14393470 | 9581477 | 15355400 | 7725858 | 15018065 | 18835202 | 19114772 | 17025148 | 15457588 | 17774304 | 0 | 1.4209915638 |
| Q9CYR0 | Ssbp1 | Single-stranded DNA-binding protein, mitochondrial (Mus musculus) | 227.82 | 30.26 | 17319 | 4 | 10558350 | 5674872 | 6556359 | 9835778 | 10156011 | 11468485 | 12550331 | 11595112 | 11997526 | 12367304 | 0 | 1.4019831062 |
| Q04692 | Smarcad1 | SWI/SNF-related matrix-associated actin-dependent regulator of chromatin subfamily A containing DEAD/H box 1 (Mus musculus) | 132.98 | 2.94 | 116451 | 3 | 1303545 | 1224065 | 783457.31 | 350832.59 | 458192.78 | 1401886.75 | 1330087.5 | 1627761 | 1383895.25 | 1602792.12 | 0 | 1.7830721759 |
| P17095 | Hmga1 | High mobility group protein HMG-I/HMG-Y (Mus musculus) | 51.23 | 14.95 | 11614 | 1 | 1336059 | 753171.94 | 950016.62 | 875054.62 | 1232856.12 | 696213.94 | 675834.38 | 611730.88 | 625623.19 | 680603.69 | 0 | 0.6391888277 |
| Q3U1J4 | Ddb1 | DNA damage-binding protein 1 (Mus musculus) | 1751.3 | 34.39 | 126853 | 33 | 70929504 | 85254432 | 86696392 | 72983608 | 86193680 | 96732704 | 86977600 | 90916792 | 90156560 | 93162400 | 0 | 1.1390060473 |
| Q9R1T2 | Sae1 | SUMO-activating enzyme subunit 1 (Mus musculus) | 573.01 | 26.86 | 38620 | 9 | 17956360 | 22488628 | 20965186 | 27157372 | 18316458 | 14280738 | 16826058 | 16255768 | 16682310 | 15636784 | 0 | 0.7454965665 |
| P70318 | Tial1 | Nucleolysin TIAR (Mus musculus) | 108.93 | 4.85 | 43389 | 2 | 3680462.5 | 4843167 | 3579978 | 5741653 | 4532267 | 2825753 | 3167625 | 3111571 | 3383272 | 3431324.5 | 0 | 0.7114077058 |
| Q8BGE6 | Atg4b | Cysteine protease ATG4B (Mus musculus) | 279.81 | 15.78 | 44375 | 4 | 847903.69 | 1683305.12 | 1797061.88 | 1867327.62 | 1107488.25 | 1915546.25 | 2026547.62 | 2135695.75 | 2327042.75 | 2124955.25 | 0 | 1.4418270321 |
| Q9D8E6 | Rpl4 | 60S ribosomal protein L4 (Mus musculus) | 1383.84 | 47.97 | 47154 | 22 | 332671488 | 191079552 | 296308064 | 144261248 | 289881856 | 364349600 | 369732064 | 377829760 | 357688288 | 361410592 | 0 | 1.4599003991 |
| Q64240 | Ambp | Protein AMBP (Rattus norvegicus) | 39.26 | 2.01 | 38851 | 1 | 13092443 | 19914738 | 15190335 | 28142160 | 19279354 | 11746894 | 12901745 | 8794090 | 12182833 | 10521481 | 0 | 0.587195279 |
| Q08879 | Fbln1 | Fibulin-1 (Mus musculus) | 81.78 | 2.7 | 78033 | 2 | 2710085.5 | 6079276 | 3334981 | 10005648 | 6245238.5 | 1898972.25 | 1038047.06 | 1824472.5 | 1832106.5 | 1732128.12 | 0 | 0.2934153035 |
| Q8CGZ0 | Cherp | Calcium homeostasis endoplasmic reticulum protein (Mus musculus) | 91.76 | 2.03 | 106169 | 2 | 577419.06 | 371554.38 | 592367.69 | 720006.94 | 698953.62 | 747243 | 779291.44 | 768420.75 | 789617.12 | 783849.12 | 0 | 1.3067659432 |
| Q9D1G2 | Pmvk | Phosphomevalonate kinase (Mus musculus) | 67.06 | 9.9 | 21916 | 1 | 436507.91 | 71282.36 | 0 | 237526.12 | 467199.53 | 488700.41 | 517456.88 | 490638.41 | 575328.88 | 486799.53 | 0 | 2.1104251646 |
| Q99JF8 | Psip1 | PC4 and SFRS1-interacting protein (Mus musculus) | 255.63 | 10.98 | 59697 | 5 | 7646717 | 5772780 | 7304590 | 5818042.5 | 6899212.5 | 5582827.5 | 1813819.12 | 5725922 | 5381208 | 4823107.5 | 0 | 0.6975462923 |
| P48722 | Hspa4l | Heat shock 70 kDa protein 4L (Mus musculus) | 229.89 | 6.44 | 94382 | 4 | 11495837 | 12815690 | 13208098 | 13010715 | 11893420 | 10418190 | 4304682.5 | 11232996 | 10210708 | 9732598 | 0 | 0.7352837205 |
| P23780 | Glb1 | Beta-galactosidase (Mus musculus) | 463.79 | 15.77 | 73121 | 8 | 22764638 | 23385564 | 23645104 | 22185520 | 23333822 | 25374072 | 27860266 | 25617050 | 24268578 | 33071734 | 0 | 1.181044233 |

| | | | | | | | | | | | | | | | | |
|---|---|---|---|---|---|---|---|---|---|---|---|---|---|---|---|---|
| Q9CR16 | Ppid | Peptidyl-prolyl cis-trans isomerase D (Mus musculus) | 639.14 | 36.49 | 40743 | 13 | 41659696 | 32369040 | 50393832 | 41306436 | 49812404 | 50582292 | 51662556 | 52744068 | 51158892 | 53046500 | 0 | 1.202526746 |
| Q8R1I1 | Uqcr10 | Cytochrome b-c1 complex subunit 9 (Mus musculus) | 50.56 | 10.94 | 7446 | 1 | 3101289.25 | 2438258 | 2835532.25 | 0 | 2196647 | 3532666 | 3910460.5 | 3306670.75 | 3820523.25 | 3146834 | 0 | 1.6758998164 |
| P08228 | Sod1 | Superoxide dismutase [Cu-Zn] (Mus musculus) | 207.36 | 25.32 | 15943 | 3 | 35989644 | 21394508 | 23559204 | 25583612 | 37062376 | 19427500 | 20034056 | 20427674 | 20957460 | 21156188 | 0 | 0.710379163 |
| P36860 | Ralb | Ras-related protein Ral-B (Rattus norvegicus) | 145.14 | 15.53 | 23317 | 3 | 1622788.88 | 2668082.5 | 1653143.12 | 3632086.5 | 1954599.88 | 1497320.38 | 1512595.5 | 1445566.5 | 1585637.12 | 1327120 | 0 | 0.639010549 |
| B5DEN9 | Vps28 | Vacuolar protein sorting-associated protein 28 homolog (Rattus norvegicus) | 61.14 | 6.58 | 26110 | 1 | 2709986.5 | 890031 | 1067848.62 | 865771.38 | 946301.19 | 690484.19 | 615722 | 649359.75 | 622875.38 | 736974.12 | 0 | 0.5116430261 |
| Q4KLH4 | Pspc1 | Paraspeckle component 1 (Rattus norvegicus) | 203.28 | 9 | 58760 | 3 | 14239966 | 9143534 | 11932610 | 9736154 | 12998823 | 16818524 | 14027440 | 17913566 | 17902590 | 18752540 | 1 | 1.4713705533 |
| P83868 | Ptges3 | Prostaglandin E synthase 3 (Rattus norvegicus) | 192.13 | 26.25 | 18721 | 3 | 13611028 | 14572796 | 17552614 | 12527433 | 13294126 | 23001664 | 17376210 | 22000266 | 18184640 | 22433478 | 1 | 1.4393395891 |
| P84104 | Srsf3 | Serine/arginine-rich splicing factor 3 (Mus musculus) | 285.07 | 31.71 | 19330 | 5 | 35770416 | 39939236 | 28566510 | 43552932 | 36416392 | 28696608 | 24137610 | 23705708 | 14943968 | 23141692 | 1 | 0.6221351116 |
| P35441 | Thbs1 | Thrombospondin-1 (Mus musculus) | 41.89 | 0.6 | 129647 | 1 | 9623173 | 12527290 | 7505677.5 | 14171931 | 13968960 | 5076641.5 | 6605606 | 0 | 5831837.5 | 0 | 1 | 0.3030274141 |
| Q60973 | Rbbp7 | Histone-binding protein RBBP7 (Mus musculus) | 343.25 | 14.12 | 47790 | 6 | 25712494 | 35155528 | 29611268 | 38458200 | 31410146 | 26492002 | 20377712 | 19717172 | 19272196 | 19492622 | 1 | 0.6570206249 |
| Q61035 | Hars | Histidine--tRNA ligase, cytoplasmic (Mus musculus) | 881.4 | 36.35 | 57432 | 16 | 30019590 | 30136350 | 33364684 | 31995350 | 29017518 | 32802780 | 34625296 | 36257340 | 36789048 | 38192296 | 1 | 1.1561685282 |
| Q8VBT6 | Apobr | Apolipoprotein B receptor (Mus musculus) | 671.01 | 16.88 | 102705 | 12 | 18869924 | 18795624 | 15759059 | 18220176 | 19504328 | 14407709 | 14199821 | 16097994 | 14954116 | 15528605 | 1 | 0.8248927957 |
| Q8BP47 | Nars | Asparagine--tRNA ligase, cytoplasmic (Mus musculus) | 1009.49 | 37.75 | 64279 | 19 | 45358020 | 57544516 | 53404236 | 52913768 | 50443980 | 67020892 | 55861628 | 67125168 | 60391128 | 66469680 | 1 | 1.2202995465 |
| Q5XI41 | Tram1 | Translocating chain-associated membrane protein 1 (Rattus norvegicus) | 129.71 | 5.61 | 43030 | 2 | 4011430.5 | 6169713 | 6844108 | 6244782 | 5099435.5 | 8707677 | 6412597 | 9114823 | 8448768 | 8784260 | 1 | 1.4617166433 |
| P23385 | Grm1 | Metabotropic glutamate receptor 1 (Rattus norvegicus) | 38.59 | 0.58 | 133236 | 1 | 147772.77 | 455547.31 | 464401.31 | 0 | 0 | 833028.38 | 624443.75 | 785761.69 | 747558.69 | 549459.38 | 1 | 3.3157075649 |
| P45377 | Akr1b8 | Aldose reductase-related protein 2 (Mus musculus) | 1109.12 | 64.87 | 36121 | 18 | 108310480 | 104183856 | 120352712 | 106592312 | 108640512 | 124519248 | 116796752 | 131509056 | 130770264 | 141474400 | 1 | 1.176962981 |
| Q9JHW4 | Eefsec | Selenocysteine-specific elongation factor (Mus musculus) | 170.64 | 14.07 | 63539 | 4 | 1609432.12 | 1446285.38 | 1485029.25 | 1264092 | 1083492.5 | 2174604.25 | 2003280.25 | 1930322 | 1818255.75 | 1546983 | 1 | 1.3752888626 |
| Q3UW53 | Fam129a | Protein Niban (Mus musculus) | 1701.28 | 44.38 | 102649 | 31 | 81109184 | 89051328 | 101453448 | 92577976 | 88006408 | 99086768 | 101830768 | 112417904 | 107088328 | 115568224 | 1 | 1.1853028635 |
| P30993 | C5ar1 | C5a anaphylatoxin chemotactic receptor 1 (Mus musculus) | 108.98 | 6.27 | 39023 | 2 | 3566930.5 | 4103693.75 | 3866001.5 | 4359313 | 3918211 | 5081924.5 | 4751952 | 4281760.5 | 4566730.5 | 4557687 | 1 | 1.1729019309 |
| Q99JX7 | Nxf1 | Nuclear RNA export factor 1 (Mus musculus) | 200.3 | 7.77 | 70300 | 3 | 1438941 | 1662686.5 | 2162989.5 | 1932070.88 | 1735853.5 | 2440552.75 | 2084434 | 2315568.5 | 2781399.75 | 2968249 | 1 | 1.4094761462 |
| P42125 | Eci1 | Enoyl-CoA delta isomerase 1, mitochondrial (Mus musculus) | 239.95 | 18.69 | 32250 | 3 | 4917455 | 4546290 | 3644504.5 | 5110930 | 4108446.75 | 3278440 | 3939527 | 2366267 | 2937554.5 | 2029824.75 | 1 | 0.6517313165 |
| Q9R0P3 | Esd | S-formylglutathione hydrolase (Mus musculus) | 984.8 | 68.44 | 31320 | 14 | 116118864 | 98588280 | 81708928 | 93193256 | 102583168 | 147383392 | 115428744 | 127883888 | 117502032 | 124613480 | 1 | 1.2856992765 |
| P62270 | Rps18 | 40S ribosomal protein S18 (Mus musculus) | 577.44 | 63.16 | 17719 | 11 | 187163264 | 96340576 | 149741472 | 89378752 | 150802352 | 258723488 | 177854336 | 213596160 | 210569680 | 219841328 | 1 | 1.6046073726 |
| Q9JLJ5 | Elovl1 | Elongation of very long chain fatty acids protein 1 (Mus musculus) | 92.7 | 7.53 | 32678 | 2 | 0 | 249801 | 453619.66 | 164137.02 | 158999.8 | 473893.44 | 382109 | 863168.38 | 614524.81 | 616150.5 | 1 | 2.873532352 |
| P49615 | Cdk5 | Cyclin-dependent-like kinase 5 (Mus musculus) | 132.7 | 8.22 | 33288 | 3 | 680543.5 | 680670.56 | 835709.75 | 1203185.12 | 1481614 | 1856611.12 | 1480266.5 | 1535710.5 | 1760438 | 1704054.5 | 1 | 1.7078151996 |
| O35379 | Abcc1 | Multidrug resistance-associated protein 1 (Mus musculus) | 272.02 | 5.37 | 171185 | 6 | 1556851.88 | 1396159.25 | 2259833.5 | 1450025.38 | 1222834 | 2994807.25 | 1933289 | 3601483.5 | 2435304 | 3243533.75 | 1 | 1.8017944222 |
| Q8CI11 | Gnl3 | Guanine nucleotide-binding protein-like 3 (Mus musculus) | 549.26 | 27.14 | 60786 | 10 | 9759250 | 7726324 | 8583164 | 5977804 | 8191639.5 | 10736771 | 11596406 | 10295735 | 9548130 | 11163072 | 1 | 1.3256094588 |
| P28650 | Adssl1 | Adenylosuccinate synthetase isozyme 1 (Mus musculus) | 332.35 | 15.1 | 50254 | 5 | 3789175.25 | 2498592.5 | 4206216.5 | 5149086 | 4224750 | 5893679.5 | 5148959 | 6038140.5 | 5684797 | 6299952.5 | 1 | 1.4629450103 |
| Q9CWJ9 | Atic | Bifunctional purine biosynthesis protein PURH (Mus musculus) | 2108.76 | 76.01 | 64217 | 32 | 140019184 | 111050624 | 127240280 | 103969248 | 119658144 | 160310688 | 132893920 | 143069312 | 151185024 | 150056608 | 1 | 1.2252361358 |
| P53395 | Dbt | Lipoamide acyltransferase component of branched-chain alpha-keto acid dehydrogenase complex, mitochondrial (Mus musculus) | 135.42 | 6.02 | 53247 | 2 | 1651408.75 | 1420769.62 | 2035327.38 | 1022555.81 | 1376893.75 | 2248312 | 2405552 | 1999527.75 | 2318225 | 2093684.5 | 1 | 1.4740065437 |
| Q9CZ13 | Uqcrc1 | Cytochrome b-c1 complex subunit 1, mitochondrial (Mus musculus) | 708.63 | 33.75 | 52852 | 13 | 39447500 | 35801444 | 37517368 | 35211288 | 38139896 | 42716932 | 41470508 | 39756084 | 39110492 | 40235836 | 1 | 1.0922662102 |
| Q2NL51 | Gsk3a | Glycogen synthase kinase-3 alpha (Mus musculus) | 154.03 | 8.16 | 51661 | 3 | 1609454 | 1978615.25 | 2147777 | 2117494 | 2381284.25 | 2297706 | 2426602 | 2808592.5 | 2750711.75 | 2726932 | 1 | 1.2712282947 |
| Q8JZQ9 | Eif3b | Eukaryotic translation initiation factor 3 subunit B (Mus musculus) | 1179.6 | 37.11 | 91370 | 23 | 30780442 | 34410176 | 40602432 | 34729500 | 34640068 | 45383744 | 38033080 | 41556560 | 40888108 | 46522716 | 1 | 1.2124973378 |
| Q60972 | Rbbp4 | Histone-binding protein RBBP4 (Mus musculus) | 468.01 | 25.18 | 47656 | 8 | 26488240 | 37374772 | 32792416 | 41779560 | 34185848 | 29010320 | 21711834 | 22645210 | 22523602 | 23709420 | 1 | 0.6928502304 |
| P63276 | Rps17 | 40S ribosomal protein S17 (Mus musculus) | 617.33 | 60 | 15524 | 10 | 53709456 | 39998872 | 51002076 | 33950156 | 46372416 | 68977208 | 62889144 | 64148724 | 55409628 | 52745572 | 1 | 1.3516697926 |
| Q91V92 | Acly | ATP-citrate synthase (Mus musculus) | 1758.01 | 31.81 | 119728 | 26 | 66102884 | 61729232 | 61909492 | 60426196 | 55157080 | 64479100 | 68141792 | 67980272 | 72057368 | 69932824 | 1 | 1.1220551418 |
| P61924 | Copz1 | Coatomer subunit zeta-1 (Mus musculus) | 314.78 | 35.03 | 20198 | 6 | 8374864 | 7623195 | 10032201 | 9490974 | 8926332 | 9797798 | 11161667 | 13634442 | 11006759 | 12690767 | 1 | 1.3114651317 |
| P70388 | Rad50 | DNA repair protein RAD50 (Mus musculus) | 145.74 | 4.04 | 153488 | 3 | 333198.03 | 541809.75 | 367633.53 | 234292.31 | 470133.56 | 825844 | 521437.72 | 628440.19 | 555867 | 647313.19 | 1 | 1.6326617451 |
| Q9Z0K8 | Vnn1 | Pantetheinase (Mus musculus) | 75.81 | 2.54 | 57091 | 1 | 3495649.5 | 5063249.5 | 5366602.5 | 6489542.5 | 5733571 | 2895996 | 2851758.5 | 3005498.5 | 3336449.75 | 3867966.25 | 1 | 0.6102682303 |
| Q8BL97 | Srsf7 | Serine/arginine-rich splicing factor 7 (Mus musculus) | 191.01 | 16.48 | 30818 | 4 | 20577728 | 20369808 | 14241044 | 23487058 | 23429204 | 14534238 | 12316406 | 12556554 | 13309783 | 13546492 | 1 | 0.6489748351 |
| Q9DB25 | Alg5 | Dolichyl-phosphate beta-glucosyltransferase (Mus musculus) | 126.73 | 10.49 | 36791 | 3 | 2293847.75 | 2941315 | 3068866.75 | 2946837.75 | 3170412 | 3163700.25 | 3445748.5 | 3780036 | 4290355.5 | 4225805.5 | 1 | 1.3109548343 |
| P29418 | Atp5f1e | ATP synthase subunit epsilon, mitochondrial (Rattus norvegicus) | 43.37 | 15.69 | 5767 | 1 | 3374864.75 | 2991149 | 2933104.75 | 2777129.5 | 2643576.25 | 3653230.75 | 3398464.25 | 3583707 | 3156163 | 3447913 | 1 | 1.1711741735 |
| Q9CQK7 | Rwdd1 | RWD domain-containing protein 1 (Mus musculus) | 109.47 | 17.28 | 27785 | 2 | 589740.81 | 201323.64 | 286797.31 | 560000.5 | 703177.19 | 1031149.25 | 878814.12 | 986013.31 | 752585.12 | 656689.88 | 1 | 1.839034229 |

| | | | | | | | | | | | | | | | | | |
|---|---|---|---|---|---|---|---|---|---|---|---|---|---|---|---|---|---|
| Q9JKF1 | Iqgap1 | Ras GTPase-activating-like protein IQGAP1 (Mus musculus) | 5232.991 | 59.08 | 188742 | 82 | 262882112 | 288994336 | 318926208 | 285131808 | 296354656 | 342866144 | 313387744 | 325381184 | 322622304 | 328476576 | 1 | 1.1242485601 |
| O70133 | Dhx9 | ATP-dependent RNA helicase A (Mus musculus) | 1821.86 | 26.16 | 149475 | 31 | 64600044 | 90062208 | 85556192 | 85777144 | 81483184 | 93586888 | 89941752 | 102238248 | 105379328 | 109645208 | 1 | 1.2290000324 |
| Q9D7A6 | Srp19 | Signal recognition particle 19 kDa protein (Mus musculus) | 67.6 | 17.36 | 16191 | 1 | 713358.25 | 588844.81 | 545800.56 | 468795.19 | 599639.12 | 395505.31 | 313037.28 | 529058.56 | 380234.25 | 440326.66 | 1 | 0.7057109081 |
| Q8VI93 | Oas3 | 2'-5'-oligoadenylate synthase 3 (Mus musculus) | 373.21 | 8.61 | 126333 | 8 | 5226069 | 3488270.5 | 4718349 | 4817912.5 | 4028115.25 | 5125047 | 5754890 | 5774000 | 5653836.5 | 6051684.5 | 1 | 1.2729395034 |
| Q8BHN3 | Ganab | Neutral alpha-glucosidase AB (Mus musculus) | 1191.52 | 28.92 | 106911 | 23 | 46779320 | 60615112 | 57171204 | 54700000 | 55713324 | 67828488 | 62007924 | 62550728 | 68005496 | 60189768 | 1 | 1.165843394 |
| P59999 | Arpc4 | Actin-related protein 2/3 complex subunit 4 (Mus musculus) | 620.23 | 75.6 | 19667 | 10 | 122111408 | 156472144 | 140645488 | 190976544 | 138710496 | 123081648 | 99155448 | 101584352 | 104115128 | 93652256 | 1 | 0.6964583161 |
| D4A208 | Srgap2 | SLIT-ROBO Rho GTPase-activating protein 2 (Rattus norvegicus) | 375.76 | 6.91 | 120878 | 7 | 4558247 | 4084222.25 | 3907515.5 | 6070298 | 4629985.5 | 4945878 | 6315725 | 7336522 | 6473749 | 6845962 | 1 | 1.3727943117 |
| Q8R1F1 | Fam129b | Niban-like protein 1 (Mus musculus) | 382.12 | 13.75 | 84819 | 7 | 5611662 | 6151221 | 8467062 | 6080701.5 | 7514323.5 | 8272091.5 | 9697897 | 8698157 | 8613213 | 8574286 | 1 | 1.2965464419 |
| P36993 | Ppm1b | Protein phosphatase 1B (Mus musculus) | 134.62 | 10.26 | 42795 | 3 | 2703121.75 | 2926719.5 | 3095647 | 2445665 | 2228373.25 | 3386224 | 3428101.5 | 2949653 | 3109145.25 | 3571932.25 | 1 | 1.2272863522 |
| P62918 | Rpl8 | 60S ribosomal protein L8 (Mus musculus) | 463.6 | 33.07 | 28025 | 10 | 159883504 | 61436336 | 106301568 | 49602768 | 117352520 | 188220448 | 133291328 | 171669632 | 187292112 | 184569968 | 1 | 1.749058326 |
| Q6ZWQ7 | Spcs3 | Signal peptidase complex subunit 3 (Mus musculus) | 47.2 | 6.11 | 20313 | 1 | 1250247.5 | 1181256.12 | 1154966 | 1376555.12 | 1212961.25 | 1657545.88 | 1375058.62 | 1648304.75 | 1607736.38 | 2127076.75 | 1 | 1.3626524402 |
| Q9CY64 | Blvra | Biliverdin reductase A (Mus musculus) | 712.06 | 43.39 | 33525 | 11 | 37978756 | 31394310 | 37498984 | 29714908 | 40354500 | 45465744 | 39386104 | 42020936 | 45371616 | 41404132 | 1 | 1.2074532131 |
| P70698 | Ctps1 | CTP synthase 1 (Mus musculus) | 395.86 | 14.21 | 66682 | 6 | 7857447.5 | 8452078 | 6328662 | 7763257 | 7634030 | 9410839 | 7863897 | 9971275 | 9083334 | 8889502 | 1 | 1.1888598103 |
| P61314 | Rpl15 | 60S ribosomal protein L15 (Rattus norvegicus) | 828.49 | 53.92 | 24146 | 16 | 126769208 | 92650016 | 162411872 | 61326404 | 104152648 | 181754352 | 166057856 | 178336944 | 149728624 | 176729904 | 1 | 1.5578144917 |
| P62281 | Rps11 | 40S ribosomal protein S11 (Mus musculus) | 438.46 | 50.63 | 18431 | 9 | 126808624 | 62411640 | 76966960 | 39773296 | 96685944 | 146796192 | 107006320 | 142986144 | 130696304 | 136590720 | 1 | 1.6492773174 |
| Q6R5N8 | Tlr13 | Toll-like receptor 13 (Mus musculus) | 245.59 | 6.36 | 114443 | 4 | 1451025 | 535826.94 | 1955733.75 | 1779767.75 | 1249303.5 | 1918985.62 | 2326784.5 | 2528471.75 | 2073375.38 | 2431242 | 1 | 1.6178161586 |
| Q5FVM3 | Retreg1 | Reticulophagy regulator 1 (Rattus norvegicus) | 100.57 | 4.79 | 52782 | 2 | 1096852.62 | 1293803.88 | 3189037.5 | 821258.06 | 788327.69 | 3228096 | 1628529.88 | 3847404.5 | 3555591 | 3330852.25 | 1 | 2.1685723984 |
| Q6UPE1 | Etfdh | Electron transfer flavoprotein-ubiquinone oxidoreductase, mitochondrial (Rattus norvegicus) | 47.53 | 1.95 | 68198 | 1 | 1450633.62 | 823893.12 | 1242392.5 | 986576.06 | 1348692.5 | 1354783.38 | 1589130 | 1475990.75 | 2083374.88 | 1796484.75 | 1 | 1.4182326411 |
| Q8R0G9 | Nup133 | Nuclear pore complex protein Nup133 (Mus musculus) | 418.98 | 10.48 | 128620 | 7 | 1156218 | 1862064 | 2456994 | 3775920.75 | 2929527.75 | 4509538 | 3383138 | 3978905.25 | 3798858.5 | 4191582.5 | 1 | 1.6306109091 |
| Q9D0E1 | Hnrnpm | Heterogeneous nuclear ribonucleoprotein M (Mus musculus) | 2051.8 | 53.5 | 77649 | 34 | 156367936 | 173828880 | 163489168 | 193025904 | 187974432 | 140570944 | 148661680 | 152429200 | 155468336 | 158675584 | 1 | 0.8640877612 |
| P05201 | Got1 | Aspartate aminotransferase, cytoplasmic (Mus musculus) | 1336.71 | 60.29 | 46248 | 22 | 117103864 | 134346400 | 118649360 | 137585264 | 109520224 | 110442216 | 103308520 | 99980704 | 105250800 | 105921064 | 1 | 0.850451971 |
| Q61093 | Cybb | Cytochrome b-245 heavy chain (Mus musculus) | 184.24 | 8.77 | 65305 | 4 | 2681147.25 | 2157110 | 2962913 | 1128218 | 2490853 | 4124419.25 | 2980819 | 3490380.25 | 3515873.5 | 2854190 | 1 | 1.4855800003 |
| Q99JI4 | Psmd6 | 26S proteasome non-ATPase regulatory subunit 6 (Mus musculus) | 1254.74 | 63.24 | 45536 | 21 | 38342888 | 43963096 | 48554544 | 39110708 | 46170316 | 48837268 | 46975232 | 51426416 | 50830024 | 51472044 | 1 | 1.1545257341 |
| Q9JHP7 | Kdelc1 | Protein O-glucosyltransferase 2 (Mus musculus) | 137.57 | 6.37 | 57985 | 3 | 724535.12 | 1167634 | 376807.69 | 1133516.62 | 1368895.5 | 1525504.62 | 1435818.62 | 1917034.62 | 1277537.75 | 1631870.75 | 1 | 1.6321801627 |
| P19253 | Rpl13a | 60S ribosomal protein L13a (Mus musculus) | 382.13 | 33.5 | 23464 | 8 | 129596624 | 41287772 | 103043784 | 21551392 | 82081200 | 176754832 | 145114224 | 130725104 | 132589856 | 122500672 | 1 | 1.8743596806 |
| Q61703 | Itih2 | Inter-alpha-trypsin inhibitor heavy chain H2 (Mus musculus) | 691.32 | 10.68 | 105928 | 10 | 80092088 | 154135456 | 119587656 | 190626800 | 145461728 | 68308360 | 81223432 | 69023712 | 80066984 | 73843368 | 1 | 0.5398809151 |
| P61255 | Rpl26 | 60S ribosomal protein L26 (Mus musculus) | 539.21 | 46.21 | 17258 | 10 | 121931128 | 55334548 | 95767352 | 35705252 | 78441152 | 148068928 | 129112016 | 122781304 | 114250808 | 124847072 | 1 | 1.6505528837 |
| Q6MG49 | Bag6 | Large proline-rich protein BAG6 (Rattus norvegicus) | 173.08 | 3.4 | 120012 | 3 | 5545902 | 6512303 | 5397259 | 5033788 | 6280863.5 | 6513432 | 7374915 | 6546351 | 6512809 | 6477716.5 | 1 | 1.161803591 |
| Q6P549 | Inppl1 | Phosphatidylinositol 3,4,5-trisphosphate 5-phosphatase 2 (Mus musculus) | 283.25 | 6.6 | 138973 | 6 | 5922042.5 | 5671564.5 | 5428325.5 | 3566923.5 | 3887346.25 | 6563275 | 7398276.5 | 5840655 | 6492665.5 | 6197518.5 | 1 | 1.3275094791 |
| Q9CR95 | Necap1 | Adaptin ear-binding coat-associated protein 1 (Mus musculus) | 159.14 | 19.27 | 29639 | 3 | 5198836.5 | 5108206 | 5146017.5 | 4475055 | 5836264 | 6528992 | 5861236 | 5582994 | 6944971.5 | 8353821 | 1 | 1.2913959424 |
| O70194 | Eif3d | Eukaryotic translation initiation factor 3 subunit D (Mus musculus) | 902.62 | 43.98 | 63989 | 16 | 30257812 | 28733616 | 31908856 | 24466882 | 35363644 | 35905060 | 35441632 | 36722988 | 35843788 | 34077192 | 1 | 1.1808512142 |
| Q99JW2 | Acy1 | Aminoacylase-1 (Mus musculus) | 158.84 | 7.6 | 45781 | 3 | 15082656 | 7949979 | 8064280.5 | 11041590 | 6882507 | 4952781 | 2430309.5 | 6053959 | 7640625 | 6164442.5 | 1 | 0.5557232625 |
| Q8BMF4 | Dlat | Dihydrolipoyllysine-residue acetyltransferase component of pyruvate dehydrogenase complex, mitochondrial (Mus musculus) | 629.52 | 26.48 | 67942 | 11 | 48982736 | 49848232 | 44104200 | 65677960 | 51538776 | 38093992 | 40124152 | 40994576 | 46735536 | 42522808 | 1 | 0.8013436027 |
| O35226 | Psmd4 | 26S proteasome non-ATPase regulatory subunit 4 (Mus musculus) | 244.27 | 14.1 | 40704 | 4 | 8758211 | 7135502 | 9490966 | 6921804.5 | 2990188.5 | 9950875 | 12271609 | 9334207 | 10221968 | 9533990 | 1 | 1.4537531754 |
| Q9DCZ4 | Apoo | MICOS complex subunit Mic26 (Mus musculus) | 44.09 | 5.05 | 22604 | 1 | 0 | 0 | 561788.31 | 580504.44 | 500750.81 | 637002.94 | 634637.88 | 721263.31 | 744884.81 | 812919.06 | 1 | 2.161055304 |
| Q91ZX7 | Lrp1 | Prolow-density lipoprotein receptor-related protein 1 (Mus musculus) | 3737.55 | 20.04 | 504742 | 66 | 130040296 | 125010152 | 132964608 | 130908488 | 116438136 | 133733800 | 137926416 | 132842624 | 137192848 | 134313712 | 1 | 1.0639757185 |
| O08715 | Akap1 | A-kinase anchor protein 1, mitochondrial (Mus musculus) | 99.93 | 3.27 | 92195 | 2 | 427368.44 | 433210.28 | 405868.62 | 0 | 0 | 464461.16 | 476194.62 | 436089.59 | 577892.31 | 710110.75 | 1 | 2.1041130932 |
| P47245 | Nrdc | Nardilysin (Rattus norvegicus) | 484.72 | 12.06 | 132976 | 11 | 16209932 | 7184836 | 9914651 | 9984664 | 9546030 | 6287344 | 6508002.5 | 5740381 | 8236811.5 | 6232689 | 1 | 0.624624478 |
| Q60664 | Lrmp | Lymphoid-restricted membrane protein (Mus musculus) | 125.35 | 6.31 | 59588 | 3 | 4583877.5 | 2981663 | 4211195.5 | 2474549 | 3970251.25 | 4449480 | 4667361 | 4996272 | 4595842 | 4659045 | 1 | 1.2824385211 |
| O35303 | Dnm1l | Dynamin-1-like protein (Rattus norvegicus) | 459.75 | 15.23 | 83908 | 7 | 2699549 | 6644695 | 7589436.5 | 6980706 | 6932898 | 8140386.5 | 8592178 | 9146497 | 8832190 | 7542495 | 1 | 1.3697719973 |
| Q6PDM2 | Srsf1 | Serine/arginine-rich splicing factor 1 (Mus musculus) | 356.69 | 29.03 | 27745 | 7 | 20952860 | 21164014 | 11657652 | 39311116 | 22778838 | 16062991 | 9574256 | 11438428 | 10182023 | 11597571 | 1 | 0.5079664536 |
| Q99020 | Hnrnpab | Heterogeneous nuclear ribonucleoprotein A/B (Mus musculus) | 374.87 | 23.16 | 30831 | 7 | 26113080 | 19523394 | 19933084 | 22044102 | 18792290 | 15431141 | 18588040 | 18195396 | 18404018 | 18860910 | 1 | 0.8409257659 |
| Q810Y8 | Pramel7 | Preferentially expressed antigen in melanoma-like protein 7 (Mus musculus) | 48.14 | 1.46 | 55834 | 1 | 559665.88 | 1514801.12 | 426979.25 | 1760853 | 1002354.31 | 526118 | 396285.47 | 340689.91 | 360599.78 | 392073.09 | 1 | 0.3828867801 |

| | | | | | | | | | | | | | | | | | |
|---|---|---|---|---|---|---|---|---|---|---|---|---|---|---|---|---|---|
| Q9CXK8 | Nip7 | 60S ribosome subunit biogenesis protein NIP7 homolog (Mus musculus) | 51 | 7.78 | 20452 | 1 | 0 | 1032699.62 | 1011588.06 | 955821.31 | 1176357.12 | 1063046 | 1369235.38 | 1528731.88 | 1218545.75 | 1527116.88 | 1 | 1.6058255265 |
| Q9CZ30 | Ola1 | Obg-like ATPase 1 (Mus musculus) | 578.52 | 36.36 | 44730 | 10 | 46892232 | 33715716 | 33995528 | 35471488 | 31192036 | 32075734 | 25406304 | 30398128 | 31117650 | 31116110 | 1 | 0.8281370906 |
| P67984 | Rpl22 | 60S ribosomal protein L22 (Mus musculus) | 361.87 | 40.62 | 14759 | 5 | 36282100 | 37211976 | 45418824 | 35698700 | 37412744 | 44792368 | 51265056 | 45413856 | 46996028 | 48804476 | 2 | 1.2356338736 |
| Q60631 | Grb2 | Growth factor receptor-bound protein 2 (Mus musculus) | 331.95 | 40.55 | 25238 | 7 | 15857944 | 13937070 | 16213742 | 13799542 | 12965350 | 18609592 | 15340028 | 19158688 | 18799744 | 19824746 | 2 | 1.2605221879 |
| Q640N3 | Arhgap30 | Rho GTPase-activating protein 30 (Mus musculus) 0 | 291.71 | 5.09 | 120114 | 5 | 2316403.5 | 3099584.25 | 2810494 | 2041406.12 | 2065002.25 | 3006321.5 | 3957429.5 | 3712807.75 | 3460130.5 | 3091243 | 2 | 1.3969095713 |
| P36371 | Tap2 | Antigen peptide transporter 2 (Mus musculus) | 284.82 | 9.4 | 77445 | 5 | 2132111 | 2500809.75 | 2584775.5 | 1387628.88 | 1609249.62 | 2436308.5 | 2998408 | 3508412 | 3163531 | 3181686.5 | 2 | 1.4967187939 |
| P51912 | Slc1a5 | Neutral amino acid transporter B(0) (Mus musculus) | 322.9 | 12.12 | 58483 | 6 | 7707645 | 7925704 | 8275023.5 | 6870187.5 | 9555320 | 11217212 | 9603862 | 10337615 | 9190203 | 9548128 | 2 | 1.237099431 |
| Q6ZQB6 | Ppip5k2 | Inositol hexakisphosphate and diphosphoinositol-pentakisphosphate kinase 2 (Mus musculus) | 207.16 | 4.07 | 128427 | 4 | 577268.69 | 390517.75 | 395383.25 | 970593.25 | 939821.12 | 789206 | 1655937.25 | 1261161.25 | 1614895.25 | 1852465.12 | 2 | 2.1913794601 |
| P53563 | Bcl2l1 | Bcl-2-like protein 1 (Rattus norvegicus) | 102.75 | 8.15 | 26158 | 2 | 3761144.5 | 3458364.25 | 3755021.5 | 3309615 | 3519322.5 | 3618658 | 4708578 | 4480489 | 4783961 | 4752746.5 | 2 | 1.2550606889 |
| Q99KK9 | Hars2 | Probable histidine--tRNA ligase, mitochondrial (Mus musculus) | 195.19 | 9.9 | 56985 | 5 | 12360387 | 7509973.5 | 8097056.5 | 8203669 | 6879306 | 11590608 | 12823273 | 11391848 | 13173172 | 13853412 | 2 | 1.4595061759 |
| Q8BPU7 | Elmo1 | Engulfment and cell motility protein 1 (Mus musculus) | 377.08 | 14.58 | 83936 | 8 | 7983245 | 7277130 | 6455415 | 8203346 | 6792058 | 6902312.5 | 4481415 | 4178739.5 | 3468686.5 | 5653655 | 2 | 0.672405493 |
| P08249 | Mdh2 | Malate dehydrogenase, mitochondrial (Mus musculus) | 1294.35 | 65.68 | 35611 | 20 | 184394672 | 251144736 | 234576336 | 245180528 | 223067872 | 195104528 | 154962544 | 179700592 | 189820672 | 183278000 | 2 | 0.7931261194 |
| O55142 | Rpl35a | 60S ribosomal protein L35a (Mus musculus) | 149.86 | 31.82 | 12554 | 4 | 19679548 | 27364646 | 32233748 | 35912268 | 22422812 | 23301492 | 11864810 | 14075509 | 14870770 | 13835862 | 2 | 0.5664321724 |
| Q8C0I1 | Agps | Alkyldihydroxyacetonephosphate synthase, peroxisomal (Mus musculus) | 655.49 | 26.2 | 71684 | 13 | 11559444 | 11515855 | 12591021 | 11662402 | 10881729 | 12886085 | 13146042 | 12505841 | 12802432 | 12170687 | 2 | 1.0910598683 |
| P57716 | Ncstn | Nicastrin (Mus musculus) | 351.91 | 10.17 | 78492 | 7 | 28305288 | 13573066 | 14016807 | 13562004 | 16860382 | 28499644 | 30216056 | 27631942 | 29551486 | 27819056 | 2 | 1.6649938395 |
| Q61235 | Sntb2 | Beta-2-syntrophin (Mus musculus) | 69.82 | 1.73 | 56382 | 1 | 111258.27 | 502471.28 | 542167.81 | 0 | 82959.95 | 665255.25 | 792931.62 | 716638.25 | 758532.62 | 477693.94 | 2 | 2.7533854403 |
| Q9EST5 | Anp32b | Acidic leucine-rich nuclear phosphoprotein 32 family member B (Mus musculus) | 298.85 | 21.69 | 31079 | 5 | 45578064 | 21919178 | 33570956 | 22376028 | 28791898 | 47434688 | 41525976 | 52367216 | 44056048 | 49637084 | 2 | 1.5437926678 |
| Q9EQH2 | Erap1 | Endoplasmic reticulum aminopeptidase 1 (Mus musculus) | 896.8 | 25.7 | 106599 | 18 | 22789078 | 23353684 | 23779472 | 21443120 | 21028840 | 25887184 | 23078916 | 24705628 | 25470128 | 27646392 | 2 | 1.128067594 |
| P27008 | Parp1 | Poly [ADP-ribose] polymerase 1 (Rattus norvegicus) | 1103.1 | 26.82 | 112660 | 19 | 33632480 | 29519652 | 32013806 | 31745962 | 33501988 | 37560104 | 35456028 | 32426152 | 37514352 | 38853228 | 2 | 1.1333798231 |
| Q8K3J1 | Ndufs8 | NADH dehydrogenase [ubiquinone] iron-sulfur protein 8, mitochondrial (Mus musculus) | 111.68 | 9.43 | 24038 | 2 | 628977.44 | 626398.12 | 809470 | 1238229.5 | 1683526.88 | 1674808.75 | 1874390.25 | 2291224 | 1282851.38 | 1917800 | 2 | 1.8130732087 |
| Q9JJY4 | Ddx20 | Probable ATP-dependent RNA helicase DDX20 (Mus musculus) | 109.39 | 2.91 | 91710 | 2 | 88743.87 | 1355936.75 | 658172.25 | 0 | 454709.38 | 1304106 | 1483474.75 | 1757362.12 | 934980.12 | 1620780.75 | 2 | 2.7763561728 |
| Q9CQA3 | Sdhb | Succinate dehydrogenase [ubiquinone] iron-sulfur subunit, mitochondrial (Mus musculus) | 417.47 | 30.14 | 31814 | 8 | 11368674 | 9281094 | 7117027 | 8758237 | 7503903 | 9048792 | 13666248 | 12100678 | 14750155 | 14371498 | 2 | 1.4521671033 |
| P17078 | Rpl35 | 60S ribosomal protein L35 (Rattus norvegicus) | 141.22 | 21.95 | 14553 | 3 | 37742420 | 8061734 | 23956934 | 5304917 | 12978617 | 54858616 | 39872980 | 39386296 | 32307782 | 33601092 | 2 | 2.271879434 |
| Q8VEB4 | Pla2g15 | Group XV phospholipase A2 (Mus musculus) | 103.93 | 5.1 | 47307 | 2 | 2747560.75 | 3486074.25 | 2787485.25 | 3359958 | 3446273.5 | 2579940 | 2903902 | 2277146.75 | 2644375.75 | 2353637.25 | 2 | 0.806136235 |
| O55012 | Picalm | Phosphatidylinositol-binding clathrin assembly protein (Rattus norvegicus) | 972.19 | 37.19 | 69286 | 16 | 30591922 | 38556988 | 41113356 | 32466186 | 35503640 | 40225940 | 40329552 | 48798456 | 42583884 | 43808300 | 2 | 1.2104785933 |
| P54311 | Gnb1 | Guanine nucleotide-binding protein G(I)/G(S)/G(T) subunit beta-1 (Rattus norvegicus) | 783.17 | 47.35 | 37377 | 12 | 41000780 | 40645532 | 56381436 | 32301790 | 34523988 | 50590956 | 60532348 | 56560448 | 54373636 | 59714928 | 2 | 1.3754818943 |
| P54728 | Rad23b | UV excision repair protein RAD23 homolog B (Mus musculus) | 144.87 | 9.86 | 43513 | 3 | 2841036.75 | 3262418 | 2752214 | 3969022.5 | 2341452.5 | 1837409.75 | 2438585.5 | 1033679.69 | 2519630.25 | 1089020.75 | 2 | 0.5880417651 |
| O08734 | Bak1 | Bcl-2 homologous antagonist/killer (Mus musculus) | 156.37 | 19.62 | 23295 | 3 | 1231320.88 | 2089871 | 3528660 | 986332.69 | 729239.69 | 3218961.5 | 2421250 | 4028904.5 | 3898303 | 4415748 | 2 | 2.0995068609 |
| Q8CI75 | Dis3l2 | DIS3-like exonuclease 2 (Mus musculus) | 169.13 | 3.68 | 97775 | 4 | 2834102 | 2005339.5 | 4323202 | 1257832.38 | 2175228.75 | 4476492.5 | 3839969 | 4729946.5 | 4020298.25 | 4901016 | 2 | 1.7440645756 |
| Q64430 | Atp7a | Copper-transporting ATPase 1 (Mus musculus) | 122.55 | 4.09 | 161959 | 3 | 1543172.12 | 1483242.5 | 1151046.75 | 1895195.25 | 1616114.75 | 1894748.88 | 1646976 | 2223014.75 | 2142208 | 2153948.25 | 2 | 1.3085180188 |
| Q63524 | Tmed2 | Transmembrane emp24 domain-containing protein 2 (Rattus norvegicus) | 131.68 | 12.44 | 22733 | 2 | 1727179.5 | 1741035.5 | 2036100.88 | 2473725.75 | 2066277.62 | 1676480.5 | 1053204.12 | 1768498 | 1104189.62 | 1518435.38 | 2 | 0.7089387984 |
| P19139 | Csnk2a1 | Casein kinase II subunit alpha (Rattus norvegicus) | 527.81 | 27.37 | 45073 | 8 | 15374872 | 15528830 | 18372356 | 16062830 | 17269048 | 19344792 | 16762326 | 19236556 | 19296768 | 21095606 | 2 | 1.1589207119 |
| Q5SSL4 | Abr | Active breakpoint cluster region-related protein (Mus musculus) | 263.97 | 7.92 | 97667 | 5 | 3349216 | 3320120.5 | 3619804 | 2797682.5 | 2774822 | 4030968.25 | 3475745.25 | 3502494 | 3830673.25 | 4105693.5 | 2 | 1.1944268233 |
| Q6P5E4 | Uggt1 | UDP-glucose:glycoprotein glucosyltransferase 1 (Mus musculus) | 2092.78 | 35.14 | 176434 | 35 | 47125184 | 52568520 | 50924296 | 52908648 | 44401760 | 61946120 | 50968044 | 60287912 | 54879460 | 56645284 | 2 | 1.1484235401 |
| Q9WUA3 | Pfkp | ATP-dependent 6-phosphofructokinase, platelet type (Mus musculus) | 1988.73 | 51.79 | 85455 | 29 | 77172800 | 74244264 | 86133736 | 72855352 | 78196120 | 81613168 | 82788176 | 92388288 | 87926152 | 96642040 | 2 | 1.1357571888 |
| P09411 | Pgk1 | Phosphoglycerate kinase 1 (Mus musculus) | 3209.94 | 78.42 | 44550 | 49 | 1085991424 | 964527232 | 916579456 | 893784064 | 973289856 | 1147771008 | 1004080256 | 1053363648 | 1111173888 | 1132560896 | 2 | 1.1271733112 |
| Q5U2Q3 | | Ester hydrolase C11orf54 homolog (Rattus norvegicus) | 109.61 | 8.89 | 34993 | 2 | 950291.75 | 2581437 | 1574954.25 | 2347289.5 | 2487450.75 | 901992.81 | 0 | 932614.31 | 1231680.88 | 1226933.5 | 2 | 0.4318517975 |
| P97298 | Serpinf1 | Pigment epithelium-derived factor (Mus musculus) 1 | 301.99 | 15.35 | 46234 | 5 | 6232210 | 12452724 | 10537165 | 14646486 | 12543689 | 6632242.5 | 7889910 | 5501916.5 | 6152079.5 | 5326374 | 2 | 0.5584338348 |
| Q9WV55 | Vapa | Vesicle-associated membrane protein-associated protein A (Mus musculus) | 303.56 | 30.92 | 27855 | 6 | 11738999 | 6591822.5 | 7143075 | 5427515.5 | 9483958 | 11860646 | 12774784 | 11703102 | 11516706 | 12580332 | 2 | 1.4964718659 |
| P55770 | Snu13 | NHP2-like protein 1 (Rattus norvegicus) | 297.49 | 47.66 | 14174 | 6 | 7882679 | 1799852.88 | 5333933.5 | 5459888.5 | 6285971 | 10954834 | 10088873 | 9575441 | 7288532 | 6998806.5 | 2 | 1.6779740438 |
| P17136 | Snrpb | Small nuclear ribonucleoprotein-associated protein B (Rattus norvegicus) | 214.41 | 13.42 | 23656 | 4 | 23788106 | 26357236 | 18829104 | 31934980 | 18690836 | 20294394 | 13428986 | 14447216 | 14935335 | 13789072 | 2 | 0.6429333993 |
| Q99KI0 | Aco2 | Aconitate hydratase, mitochondrial (Mus musculus) | 1608.92 | 41.54 | 85464 | 27 | 110291520 | 93250448 | 99393880 | 86583552 | 89815616 | 120311656 | 107436976 | 111739728 | 110929696 | 102643288 | 2 | 1.1538096019 |

| | | | | | | | | | | | | | | | | | |
|---|---|---|---|---|---|---|---|---|---|---|---|---|---|---|---|---|---|
| P62900 | Rpl31 | 60S ribosomal protein L31 (Mus musculus) | 94.84 | 16.8 | 14463 | 2 | 25741710 | 7417131 | 13461023 | 8084467 | 17349888 | 29347004 | 19683800 | 30651364 | 26755888 | 23623280 | 2 | 1.805048168 |
| P21107 | Tpm3 | Tropomyosin alpha-3 chain (Mus musculus) | 515.89 | 23.16 | 32994 | 10 | 147905312 | 133635584 | 121570352 | 122139024 | 155233120 | 107267856 | 120087392 | 99904976 | 127599144 | 103653104 | 2 | 0.8207584176 |
| Q642C0 | Dnajc8 | DnaJ homolog subfamily C member 8 (Rattus norvegicus) | 147.85 | 18.18 | 29813 | 3 | 7025889 | 6093753 | 3820715.5 | 5799480 | 5031670.5 | 3609048 | 4415517.5 | 3268578.5 | 4126296 | 3206729.25 | 2 | 0.6706934766 |
| P20069 | Pmpca | Mitochondrial-processing peptidase subunit alpha (Rattus norvegicus) | 45.74 | 2.1 | 58608 | 1 | 431796.97 | 567034.69 | 577726.62 | 367372.84 | 305874.03 | 802898.81 | 639472.31 | 1030475.25 | 653124.69 | 557675.81 | 2 | 1.6373181784 |
| Q8CIN7 | Impa2 | Inositol monophosphatase 2 (Rattus norvegicus) | 111.05 | 7.59 | 31796 | 2 | 2448747.25 | 2410364.75 | 2354152.75 | 1722246.62 | 2251148.5 | 2159966 | 1473041.88 | 1758464.12 | 1696569.25 | 1658473.88 | 2 | 0.7818701231 |
| P42227 | Stat3 | Signal transducer and activator of transcription 3 (Mus musculus) | 523.78 | 21.04 | 88054 | 10 | 4857057 | 5909592.5 | 6851488.5 | 5195042 | 5455524 | 8408922 | 7185163 | 5860043 | 8697097 | 6947801.5 | 2 | 1.3123709704 |
| P62334 | Psmc6 | 26S proteasome regulatory subunit 10B (Mus musculus) | 541.38 | 26.48 | 44173 | 9 | 15664647 | 12366426 | 12818725 | 10502664 | 15533898 | 16603395 | 17135284 | 16917364 | 17501328 | 15265783 | 2 | 1.2472371646 |
| Q64324 | Stxbp2 | Syntaxin-binding protein 2 (Mus musculus) | 968.01 | 36.26 | 66357 | 16 | 18996106 | 11398588 | 17726692 | 11939922 | 12762599 | 19261004 | 21123428 | 22442302 | 16499246 | 20280518 | 2 | 1.3677719598 |
| O35864 | Cops5 | COP9 signalosome complex subunit 5 (Mus musculus) | 349.26 | 21.26 | 37549 | 7 | 10847557 | 10901956 | 13385044 | 9139152 | 7315845 | 15038064 | 13684781 | 13169667 | 13612852 | 12810917 | 2 | 1.3242270131 |
| P14733 | Lmnb1 | Lamin-B1 (Mus musculus) | 1312.09 | 42.01 | 66786 | 23 | 55711460 | 64499460 | 66322732 | 63252284 | 55981752 | 51290252 | 56346776 | 54807004 | 54479140 | 52467972 | 2 | 0.881032086 |
| O09111 | Ndufb11 | NADH dehydrogenase [ubiquinone] 1 beta subcomplex subunit 11, mitochondrial (Mus musculus) | 48.13 | 14.57 | 17444 | 1 | 544344.19 | 595573.25 | 424509.75 | 533108.19 | 561995.69 | 371308.31 | 542629.69 | 377717.31 | 393806.53 | 391157.78 | 2 | 0.7808217183 |
| P80314 | Cct2 | T-complex protein 1 subunit beta (Mus musculus) | 2533.16 | 71.96 | 57477 | 35 | 127843808 | 136972192 | 143661136 | 128805512 | 153406448 | 164674176 | 144381872 | 158076128 | 155357584 | 149504816 | 2 | 1.1177164667 |
| Q8CG76 | Akr7a2 | Aflatoxin B1 aldehyde reductase member 2 (Mus musculus) | 122.74 | 7.08 | 40612 | 2 | 1805302.25 | 750787.56 | 1322537.5 | 609912.56 | 962388.38 | 1688559.12 | 1857709.25 | 1814916 | 2038652.75 | 1563539.88 | 2 | 1.6443762583 |
| Q9CPU4 | Mgst3 | Microsomal glutathione S-transferase 3 (Mus musculus) | 258.97 | 40.52 | 16958 | 4 | 7873490 | 7880625 | 8046954 | 9220037 | 8375414 | 9288208 | 8157046 | 10196396 | 10051416 | 9972051 | 2 | 1.1514281152 |
| P00397 | Mtco1 | Cytochrome c oxidase subunit 1 (Mus musculus) | 57.91 | 2.72 | 56910 | 1 | 211920.12 | 1038802.31 | 1178125 | 465283.16 | 636615.38 | 1453482.5 | 833832.12 | 1535903.5 | 1230163.25 | 1440565 | 2 | 1.839256187 |
| Q9DBE9 | Ftsj3 | pre-rRNA processing protein FTSJ3 (Mus musculus) | 238.78 | 11.81 | 95532 | 5 | 2690970.25 | 605144.5 | 1356219.5 | 3279850.75 | 1217543.12 | 3760712.5 | 2239601 | 4041368.75 | 3458972.5 | 3698316 | 2 | 1.8797247879 |
| P40142 | Tkt | Transketolase (Mus musculus) | 2152.29 | 62.12 | 67630 | 33 | 429328896 | 397428512 | 438889440 | 417566464 | 438042144 | 493665056 | 435067040 | 450342752 | 456969824 | 450808160 | 2 | 1.0780657396 |
| Q3UFY7 | Nt5c3b | 7-methylguanosine phosphate-specific 5'-nucleotidase (Mus musculus) | 144.21 | 16.67 | 34425 | 2 | 780870.62 | 1055232.75 | 1169098.62 | 1430541.62 | 1088567.62 | 1931237 | 1264819 | 1222412.62 | 1678097 | 1805563.12 | 2 | 1.4304278689 |
| P04184 | Tk1 | Thymidine kinase, cytosolic (Mus musculus) | 250.66 | 24.89 | 25776 | 5 | 7086316.5 | 8309174 | 7564118.5 | 7012133 | 7669179 | 6511499.5 | 7510161 | 5741195 | 6269874.5 | 6676438 | 2 | 0.8689789498 |
| Q8BHL5 | Elmo2 | Engulfment and cell motility protein 2 (Mus musculus) | 139.18 | 7.38 | 83887 | 4 | 1059260.25 | 2679162.75 | 2318553 | 2574442.25 | 1511537.75 | 1111963.75 | 921047.19 | 1106150.5 | 896623.19 | 979580.75 | 2 | 0.4944678238 |
| Q4LDD4 | Arap1 | Arf-GAP with Rho-GAP domain, ANK repeat and PH domain-containing protein 1 (Mus musculus) | 166.2 | 3.72 | 162276 | 4 | 2632927.75 | 2355005 | 3327428.5 | 2054825.12 | 2393750.75 | 3005829 | 2867770.75 | 3446299 | 4504738 | 3763128 | 2 | 1.3779263079 |
| Q8BFQ4 | Wdr82 | WD repeat-containing protein 82 (Mus musculus) | 169.43 | 10.22 | 35079 | 2 | 1427644 | 2393363.5 | 2761715 | 2432327.5 | 2851447 | 4430388.5 | 3322999 | 3596114.5 | 2912888.5 | 2729337.5 | 2 | 1.4319076641 |
| Q5U1X1 | Rexo2 | Oligoribonuclease, mitochondrial (Rattus norvegicus) | 194.54 | 15.19 | 26752 | 3 | 4393494 | 6141397 | 6697391 | 3730915.5 | 3869754.25 | 6784821.5 | 5545643 | 7485478.5 | 6965166 | 7763162 | 2 | 1.3910658446 |
| O35295 | Purb | Transcriptional activator protein Pur-beta (Mus musculus) | 117.75 | 15.12 | 33901 | 2 | 2042624.5 | 642832.69 | 598863.31 | 0 | 1724727.88 | 2389338.75 | 2015820.5 | 2504713.5 | 2190395.5 | 1846649.5 | 2 | 2.1854286323 |
| P62717 | Rpl18a | 60S ribosomal protein L18a (Mus musculus) | 427.4 | 44.89 | 20732 | 9 | 95099880 | 62016392 | 81118952 | 66292820 | 87760032 | 134338560 | 95750448 | 104621480 | 93815296 | 93176264 | 2 | 1.3298952477 |
| Q60787 | Lcp2 | Lymphocyte cytosolic protein 2 (Mus musculus) | 61.23 | 3 | 60238 | 1 | 477509.38 | 757246.06 | 959794.44 | 634866.12 | 730329.25 | 981519.88 | 860115.31 | 864167.31 | 970856.25 | 1118908 | 2 | 1.3471657136 |
| Q6IRU2 | Tpm4 | Tropomyosin alpha-4 chain (Mus musculus) | 1177.79 | 49.6 | 28468 | 18 | 95060024 | 93317096 | 90402880 | 80114072 | 102232376 | 79530936 | 88711368 | 70804200 | 83157368 | 78927712 | 2 | 0.8698949838 |
| P17426 | Ap2a1 | AP-2 complex subunit alpha-1 (Mus musculus) | 723.09 | 20.47 | 107664 | 15 | 22033118 | 24017876 | 25352158 | 17020830 | 17845682 | 26340406 | 23894972 | 27950960 | 26210074 | 27218944 | 2 | 1.2385035489 |
| P11928 | Oas1a | 2'-5'-oligoadenylate synthase 1A (Mus musculus) | 85.12 | 10.08 | 42429 | 2 | 1742244.25 | 624245.75 | 1723920.75 | 0 | 856371.38 | 2041777.5 | 2201820 | 1293864.75 | 2506547.25 | 2098335.75 | 2 | 2.0502914791 |
| P62944 | Ap2b1 | AP-2 complex subunit beta (Rattus norvegicus) | 1675.99 | 38.21 | 104553 | 28 | 36320704 | 45696172 | 51452096 | 46959536 | 44654064 | 49902248 | 47225652 | 58886612 | 54680268 | 55738036 | 2 | 1.1837114426 |
| Q9R0L6 | Pcm1 | Pericentriolar material 1 protein (Mus musculus) | 67.76 | 1.43 | 228847 | 2 | 1431934.5 | 1618692.12 | 936585.75 | 1214718.5 | 1191061.38 | 919387.69 | 832221.25 | 1169244.62 | 599293.06 | 969213.38 | 2 | 0.702231416 |
| P46471 | Psmc2 | 26S proteasome regulatory subunit 7 (Mus musculus) | 956.74 | 42.03 | 48648 | 16 | 25161098 | 27779066 | 26059404 | 24639772 | 25546876 | 27489546 | 26269146 | 31963046 | 31624096 | 33968688 | 2 | 1.1712899927 |
| Q8BYA0 | Tbcd | Tubulin-specific chaperone D (Mus musculus) | 204.96 | 3.85 | 133321 | 4 | 1305800.5 | 2535895 | 2441845.75 | 3486272 | 2588863.5 | 3557945.5 | 3534424 | 3408697 | 3149222 | 3944446.5 | 2 | 1.4236746665 |
| P86182 | Ccdc22 | Coiled-coil domain-containing protein 22 (Rattus norvegicus) | 44.87 | 2.07 | 70855 | 1 | 345777.66 | 186280.12 | 262239.94 | 258424.48 | 241253.44 | 379464.25 | 414181.81 | 357731.38 | 309000.75 | 277456.28 | 2 | 1.3430194636 |
| Q9D6R2 | Idh3a | Isocitrate dehydrogenase [NAD] subunit alpha, mitochondrial (Mus musculus) | 787.3 | 41.53 | 39639 | 15 | 48030756 | 43530060 | 47437704 | 40414020 | 46436408 | 53273396 | 47348536 | 49000648 | 49057168 | 49279848 | 2 | 1.0979001594 |
| Q9EPE9 | Atp13a1 | Endoplasmic reticulum transmembrane helix translocase (Mus musculus) | 185.02 | 3.5 | 132388 | 3 | 6913363 | 5236947 | 4076177 | 6906654 | 4501023 | 3281468.5 | 3510868 | 3414033 | 4683930 | 3850200.5 | 2 | 0.6781641739 |
| P62774 | Mtpn | Myotrophin (Mus musculus) | 304.58 | 51.69 | 12861 | 6 | 15650788 | 10371032 | 11989412 | 14364844 | 13511479 | 17411326 | 15666806 | 15289413 | 16119808 | 15160860 | 2 | 1.2088506395 |
| P11438 | Lamp1 | Lysosome-associated membrane glycoprotein 1 (Mus musculus) | 623.69 | 28.33 | 43865 | 9 | 51990440 | 72632288 | 85978304 | 69628408 | 88067248 | 93866744 | 83007336 | 89963920 | 98172808 | 100342136 | 2 | 1.2635273657 |
| Q4KLL4 | Tm9sf4 | Transmembrane 9 superfamily member 4 (Rattus norvegicus) | 71.26 | 2.95 | 74675 | 1 | 454580.28 | 650301.38 | 871451.25 | 659275.81 | 0 | 744500.56 | 913805.25 | 1293832 | 853878.06 | 1084226.5 | 2 | 1.8554508235 |
| P14069 | S100a6 | Protein S100-A6 (Mus musculus) | 127.6 | 26.97 | 10051 | 2 | 20374508 | 12076886 | 5154510.5 | 14591929 | 17146406 | 9874882 | 3194262.5 | 0 | 11217901 | 3154057.5 | 2 | 0.395722892 |
| P06801 | Me1 | NADP-dependent malic enzyme (Mus musculus) | 199.32 | 11.54 | 63954 | 3 | 2092376 | 1468467.5 | 1811592.12 | 1660795.5 | 1773850.75 | 2884075 | 2215113.25 | 2397950 | 2035250.25 | 1883195.12 | 2 | 1.2961822983 |
| Q9CZB0 | Sdhc | Succinate dehydrogenase cytochrome b560 subunit, mitochondrial (Mus musculus) | 191.36 | 28.99 | 18382 | 4 | 4360575 | 6329942 | 4552827.5 | 6264042 | 3698726.5 | 5801643 | 6907908 | 6623424 | 6539664 | 7168187 | 2 | 1.3108259096 |
| P84228 | Hist1h3b | Histone H3.2 (Mus musculus) b | 448.87 | 50.74 | 15388 | 8 | 1453005568 | 924865088 | 1236678400 | 1008597696 | 1197097344 | 1004275392 | 959959680 | 859621888 | 895114368 | 741153664 | 2 | 0.7663123605 |

| | | | | | | | | | | | | | | | | | |
|---|---|---|---|---|---|---|---|---|---|---|---|---|---|---|---|---|---|
| Q9JHI5 | Ivd | Isovaleryl-CoA dehydrogenase, mitochondrial (Mus musculus) | 113.18 | 5.66 | 46325 | 2 | 4521884.5 | 3538388 | 2690661 | 4428663 | 3125442 | 2669389.5 | 2421895.5 | 2333521 | 2766178.5 | 2821765.5 | 2 | 0.7108835089 |
| Q4V7C7 | Actr3 | Actin-related protein 3 (Rattus norvegicus) | 1321.91 | 73.92 | 47357 | 23 | 198307456 | 205025344 | 215385904 | 228014192 | 206483632 | 194967616 | 191728432 | 192524128 | 192542800 | 206241056 | 2 | 0.9285878127 |
| Q80YW0 | Cyth4 | Cytohesin-4 (Mus musculus) | 320.47 | 19.08 | 45285 | 6 | 4434418.5 | 3688656.5 | 4720602.5 | 3821218.5 | 4304535.5 | 4500536 | 4470701 | 4737262 | 5179209 | 5057384.5 | 2 | 1.1419047054 |
| O70551 | Srpk1 | SRSF protein kinase 1 (Mus musculus) | 544.63 | 20.52 | 73088 | 9 | 7640265 | 7949920.5 | 8513782 | 8951432 | 8041706.5 | 9075918 | 8371455 | 11364323 | 9599410 | 11106507 | 2 | 1.2048929431 |
| Q9Z110 | Aldh18a1 | Delta-1-pyrroline-5-carboxylate synthase (Mus musculus) 1 | 797.96 | 17.61 | 87266 | 13 | 26337372 | 24590678 | 25065040 | 22254740 | 24552480 | 29585924 | 25806250 | 31074016 | 25646900 | 26752596 | 2 | 1.1308252072 |
| O08528 | Hk2 | Hexokinase-2 (Mus musculus) | 1428.24 | 31.41 | 102535 | 22 | 58215216 | 55515092 | 56726132 | 52468956 | 48778392 | 58966864 | 56162244 | 60414516 | 58757920 | 61771996 | 2 | 1.0896923528 |
| Q8BTI9 | Pik3cb | Phosphatidylinositol 4,5-bisphosphate 3-kinase catalytic subunit beta isoform (Mus musculus) | 39.04 | 0.66 | 121711 | 1 | 1596797.38 | 1155140.25 | 875780.25 | 1686547.25 | 1248761.38 | 823065.88 | 856347.75 | 838049.25 | 914807.44 | 985539 | 2 | 0.6731359859 |
| P45952 | Acadm | Medium-chain specific acyl-CoA dehydrogenase, mitochondrial (Mus musculus) | 182.98 | 10.93 | 46481 | 4 | 11569391 | 10128812 | 9682616 | 10659141 | 9518494 | 10073664 | 9216589 | 8794430 | 8898458 | 9192944 | 2 | 0.8956064703 |
| Q9Z2L7 | Crlf3 | Cytokine receptor-like factor 3 (Mus musculus) | 215.99 | 16.06 | 49559 | 4 | 2884265.5 | 3588812.5 | 3414358.5 | 2062919.25 | 4562605.5 | 3682122.5 | 3759091 | 5390304.5 | 5643194.5 | 4934821 | 2 | 1.4176460022 |
| Q3TCN2 | Plbd2 | Putative phospholipase B-like 2 (Mus musculus) | 198.44 | 5.72 | 66289 | 4 | 817454.81 | 1542982.12 | 0 | 840471.12 | 1213164.75 | 1040069.69 | 1654590.5 | 1762398.5 | 1851449.75 | 1792600.38 | 2 | 1.8352911669 |
| P56383 | Atp5mc2 | ATP synthase F(0) complex subunit C2, mitochondrial (Mus musculus) | 57.58 | 21.23 | 15476 | 1 | 308962.47 | 244382.19 | 271012.88 | 129464.12 | 134610.95 | 0 | 236036.5 | 51000.69 | 83735.15 | 85013.41 | 2 | 0.4187542213 |
| Q8C0D5 | Efl1 | Elongation factor-like GTPase 1 (Mus musculus) | 261.38 | 5.86 | 125777 | 5 | 1839556.25 | 1374392.5 | 1728237 | 1542481.5 | 1458601 | 564668.12 | 1126146 | 1377889.75 | 1260958.25 | 1420385.62 | 2 | 0.7238894066 |
| Q04899 | Cdk18 | Cyclin-dependent kinase 18 (Mus musculus) | 185.5 | 7.54 | 51848 | 3 | 516443.78 | 353274.47 | 390966.78 | 400542 | 485665.94 | 545340.75 | 777635.44 | 543893.38 | 551789.75 | 468779.47 | 2 | 1.3449383972 |
| Q9CQE3 | Mrps17 | 28S ribosomal protein S17, mitochondrial (Mus musculus) | 40.26 | 7.5 | 13382 | 1 | 1369420.88 | 656782.31 | 959469.19 | 0 | 962920.5 | 1738710.38 | 1159993.25 | 1344940.38 | 1465009.62 | 1398791.75 | 2 | 1.799994478 |
| P63174 | Rpl38 | 60S ribosomal protein L38 (Rattus norvegicus) | 63.08 | 17.14 | 8218 | 1 | 592478.31 | 0 | 679462.75 | 144588.17 | 893722.44 | 1378028.12 | 780213.06 | 955536.19 | 932789.62 | 706693.62 | 2 | 2.057464419 |
| Q9WU81 | Slc37a2 | Glucose-6-phosphate exchanger SLC37A2 (Mus musculus) | 177.56 | 7.78 | 55073 | 3 | 2638900.75 | 2220164.5 | 2914908.25 | 1110559.88 | 1502380.5 | 4086873 | 3152658.5 | 3165144.25 | 2235582.5 | 2979446.75 | 2 | 1.5037868977 |
| Q9JMH6 | Txnrd1 | Thioredoxin reductase 1, cytoplasmic (Mus musculus) | 1005.5 | 41.11 | 67084 | 15 | 38007252 | 43256372 | 44651728 | 45805612 | 41426752 | 47096208 | 47184376 | 47558532 | 45008784 | 44926072 | 2 | 1.0873866084 |
| P50518 | Atp6v1e1 | V-type proton ATPase subunit E 1 (Mus musculus) 1 | 222.25 | 19.47 | 26157 | 4 | 8998090 | 3051607.5 | 6215975 | 2546323.25 | 5736166.5 | 6903900 | 7648374.5 | 9083834 | 9555134 | 9302507 | 2 | 1.600628665 |
| Q9DCH4 | Eif3f | Eukaryotic translation initiation factor 3 subunit F (Mus musculus) | 841 | 38.78 | 37984 | 13 | 35091616 | 50888404 | 46468080 | 54302168 | 42681176 | 52168596 | 54072808 | 59514956 | 54370760 | 54638520 | 2 | 1.1975936481 |
| Q8C7K6 | Pcyox1l | Prenylcysteine oxidase-like (Mus musculus) | 58.74 | 2.22 | 54875 | 1 | 1964719.62 | 2477241.5 | 1330456.38 | 2289821.75 | 1777588.75 | 2671774.75 | 2607127.75 | 2534526.25 | 2337854 | 2407713 | 2 | 1.2763430164 |
| Q8R491 | Ehd3 | EH domain-containing protein 3 (Rattus norvegicus) | 686.92 | 25.42 | 60791 | 15 | 26450998 | 23691832 | 23766442 | 22669970 | 18609376 | 26768132 | 26608716 | 27820790 | 26214320 | 24887792 | 2 | 1.1485488089 |
| P97576 | Grpel1 | GrpE protein homolog 1, mitochondrial (Rattus norvegicus) | 209.36 | 14.29 | 24297 | 4 | 3708725.75 | 4078263.5 | 4067755.5 | 3597665.75 | 5262176 | 4089841 | 5419328.5 | 5621821 | 6099237 | 4737838 | 2 | 1.253612545 |
| P63038 | Hspd1 | 60 kDa heat shock protein, mitochondrial (Mus musculus) | 3435.89 | 76.79 | 60955 | 49 | 506479328 | 615583744 | 554901888 | 681385920 | 551899008 | 525978176 | 486942272 | 501036768 | 482019520 | 509978464 | 2 | 0.861079047 |
| Q99J36 | Thumpd1 | THUMP domain-containing protein 1 (Mus musculus) | 50.3 | 3.14 | 38885 | 1 | 663170.38 | 513415.53 | 505182.22 | 566071.12 | 0 | 581702.25 | 1096361.38 | 589097.69 | 1013554.38 | 747135.69 | 2 | 1.7918769725 |
| O35075 | Dscr3 | Down syndrome critical region protein 3 homolog (Mus musculus) | 51.46 | 6.4 | 32970 | 1 | 650354.25 | 587723.38 | 476224.53 | 624933.44 | 709926.19 | 465299.62 | 407144.88 | 0 | 0 | 572671.56 | 2 | 0.4739387935 |
| P18596 | Atp2a3 | Sarcoplasmic/endoplasmic reticulum calcium ATPase 3 (Rattus norvegicus) | 673.77 | 10.18 | 116284 | 9 | 30412192 | 27498088 | 28449190 | 21252536 | 20660818 | 27919684 | 31170804 | 30928662 | 32519616 | 31818752 | 2 | 1.2033532372 |
| Q8CG48 | Smc2 | Structural maintenance of chromosomes protein 2 (Mus musculus) | 647.17 | 11.92 | 134239 | 12 | 15951559 | 14968531 | 14848508 | 14855747 | 12854444 | 15016958 | 15590693 | 16422365 | 16193502 | 17001786 | 2 | 1.0918158164 |
| P31266 | Rbpj | Recombining binding protein suppressor of hairless (Mus musculus) | 416.87 | 19.01 | 58537 | 8 | 33263554 | 36306948 | 30352102 | 32918280 | 29600056 | 31827160 | 28944510 | 28900452 | 28174276 | 29329920 | 2 | 0.9060297115 |
| Q64620 | Ppp6c | Serine/threonine-protein phosphatase 6 catalytic subunit (Rattus norvegicus) | 223.7 | 25.57 | 35159 | 5 | 5213359.5 | 5808321 | 4595732.5 | 5162864 | 4114037.75 | 5069832 | 1232363.62 | 3843941.75 | 3481425.75 | 3420360 | 2 | 0.6848119055 |
| P80318 | Cct3 | T-complex protein 1 subunit gamma (Mus musculus) | 1805.03 | 55.6 | 60630 | 33 | 217010480 | 188533056 | 215274224 | 162401952 | 189723056 | 220839072 | 205859264 | 223914608 | 218148064 | 230878800 | 2 | 1.1302204448 |
| P70704 | Atp8a1 | Phospholipid-transporting ATPase IA (Mus musculus) | 40.92 | 0.69 | 131413 | 1 | 8997087 | 3798856.5 | 2904947 | 5176719.5 | 4579527 | 2432238 | 3021963.25 | 1812270 | 2240359 | 2921013.25 | 2 | 0.4881870063 |
| P61222 | Abce1 | ATP-binding cassette sub-family E member 1 (Mus musculus) | 1140.1 | 42.9 | 67314 | 20 | 31509978 | 35673000 | 35584104 | 38097528 | 34945692 | 44934236 | 40108204 | 36035056 | 36346680 | 39193476 | 2 | 1.1183511419 |
| Q9WUK4 | Rfc2 | Replication factor C subunit 2 (Mus musculus) | 80.96 | 4.87 | 38725 | 2 | 7796811 | 6072110.5 | 5900851.5 | 4796432 | 4840054 | 4428107 | 5701688.5 | 4409298 | 4515534.5 | 2760148 | 2 | 0.7418412522 |
| Q91VU7 | Pus7 | Pseudouridylate synthase 7 homolog (Mus musculus) | 109.16 | 4.39 | 74793 | 2 | 1299566.12 | 699016.06 | 1039924.38 | 1047376.19 | 0 | 1273369.5 | 1535923 | 1428311.38 | 1172052.62 | 1348892.88 | 2 | 1.6541222041 |
| P60603 | Romo1 | Reactive oxygen species modulator 1 (Mus musculus) | 68.71 | 21.52 | 8183 | 1 | 1085766.12 | 1096754.25 | 1224121.12 | 1058383.62 | 1356971 | 1100306.38 | 1453951.25 | 1401712.88 | 1542871.62 | 2015828.25 | 2 | 1.2907377879 |
| Q8K0D5 | Gfm1 | Elongation factor G, mitochondrial (Mus musculus) | 250.5 | 9.99 | 83550 | 6 | 3501657.25 | 6256391 | 6304128.5 | 3925173.5 | 4572707 | 6688039 | 6531404.5 | 7124716 | 6315148.5 | 4666547 | 2 | 1.2754797223 |
| Q80U87 | Usp8 | Ubiquitin carboxyl-terminal hydrolase 8 (Mus musculus) | 171.53 | 4.44 | 122611 | 4 | 2197891 | 1627862.5 | 1744086.75 | 1213044.62 | 0 | 2338775.25 | 2199139.25 | 2488814 | 1839140.12 | 1861559.5 | 2 | 1.5815435947 |
