## supplemental Table 3 for "Biobased, Biodegradable but not bio-neutral: about the effects of polylactic acid nanoparticles on macrophages"

Supplementary Table 3: results of the pathway analysis by the David tool on the proteins modulated in response to PLA beads

| Annotation Cluster 1 | Enrichment Score: 9.368300008141553 | | | | | |
|---|---|---|---|---|---|---|
| Category | Term | Count | % | PValue | Genes | FDR |
| GOTERM_BP_DIRECT | GO:0006412~translation | 31 | 8.7078652 | 1.75093E-16 | P62918, Q9D7S7, P62717, O70194, Q9DCH4, P19253, P62281, P14115, Q8K0D5, Q61035, P67984, P62751, P62274, Q9D1R9, Q8C0D5, P62270, Q8JZQ9, P63276, O55142, Q8BP47, P61255, P58252, Q9D8E6, P62900, P60229, Q9JHW4, Q99KK9, P47964, P41105, P60843, Q9CQE3 | 3.4090576E-13 |
| GOTERM_BP_DIRECT | GO:0002181~cytoplasmic translation | 19 | 5.3370787 | 1.30116E-15 | P62918, Q9D7S7, P62717, Q9D1R9, P19253, P62281, P62270, P14115, P63276, O55142, P67984, P62751, P61255, Q9D8E6, P62274, P62900, Q9CQK7, P41105, P47964 | 1.266679E-12 |
| GOTERM_CC_DIRECT | GO:0022626~cytosolic ribosome | 16 | 4.494382 | 5.6014E-13 | P62918, P62717, Q9D1R9, P19253, P62281, P62270, P14115, P63276, O55142, P67984, P62751, P61255, Q9D8E6, P62900, P41105, P47964 | 8.1220271E-11 |
| GOTERM_CC_DIRECT | GO:0005840~ribosome | 21 | 5.8988764 | 5.78761E-12 | P62918, P11928, Q9D7S7, P62717, Q9D1R9, P19253, P35564, P62281, P62270, P14115, P63276, O55142, P67984, P62751, P61255, Q9D8E6, P62274, P62900, P41105, P47964, Q9CQE3 | 6.2940266E-10 |
| GOTERM_CC_DIRECT | GO:0022625~cytosolic large ribosomal subunit | 13 | 3.6516854 | 2.69666E-10 | P62918, P62717, Q9D1R9, P19253, P14115, O55142, P67984, P62751, P61255, Q9D8E6, P62900, P41105, P47964 | 1.6757841E-08 |
| GOTERM_MF_DIRECT | GO:0003735~structural constituent of ribosome | 19 | 5.3370787 | 8.67998E-10 | P62918, Q9D7S7, P62717, Q9D1R9, P19253, P62281, P62270, P14115, P63276, O55142, P67984, P62751, P61255, Q9D8E6, P62274, P62900, P41105, P47964, Q9CQE3 | 1.9124894E-07 |
| KEGG_PATHWAY | mmu05171:Coronavirus disease - COVID-19 | 25 | 7.0224719 | 2.29093E-09 | P62918, P11928, Q9D7S7, P62717, Q8BTI9, P19253, P62281, P14115, P67984, P62751, P62274, P98086, P42227, Q8VI93, Q9D1R9, P62270, P30993, P63276, O55142, P61255, Q61093, Q9D8E6, P62900, P47964, P41105 | 5.7044048E-07 |
| UP_KW_MOLECULAR_FUNCTION | KW-0687~Ribonucleoprotein | 24 | 6.741573 | 5.13169E-09 | P62918, Q9D7S7, P62717, Q9D1R9, P19253, P62281, Q9D7A6, P62270, P14115, Q62376, Q3UEB3, P63276, O55142, P67984, P62751, P61255, Q9D8E6, P62274, P62900, Q9QXK7, Q9D0E1, P41105, P47964, Q9CQE3 | 2.6541449E-07 |
| UP_KW_MOLECULAR_FUNCTION | KW-0689~Ribosomal protein | 19 | 5.3370787 | 8.04286E-09 | P62918, Q9D7S7, P62717, Q9D1R9, P19253, P62281, P62270, P14115, P63276, O55142, P67984, P62751, P61255, Q9D8E6, P62274, P62900, P41105, P47964, Q9CQE3 | 2.6541449E-07 |

| | | | | | | |
|---|---|---|---|---|---|---|
| KEGG_PATHWAY | mmu03010:Ribosome | 19 | 5.3370787 | 1.11694E-07 | P62918, Q9D7S7, P62717, Q9D1R9, P19253, P62281, P62270, P14115, P63276, O55142, P67984, P62751, P61255, Q9D8E6, P62274, P62900, P41105, P47964, Q9CQE3 | 9.2706324E-06 |
| GOTERM_CC_DIRECT | GO:0098793~presynapse | 18 | 5.0561798 | 6.35201E-07 | P62918, Q9D1R9, P19253, Q9CR95, P35564, P62281, P14115, Q64324, O55142, P67984, P62751, P61255, O08992, Q9D8E6, P61028, P41105, P47964, P49615 | 2.7631264E-05 |
| GOTERM_CC_DIRECT | GO:0045202~synapse | 35 | 9.8314607 | 3.56897E-06 | P62918, Q61235, P62717, Q5SSL4, O70194, Q9DCH4, P19253, P10852, P62281, P14115, P67984, O35864, P70704, P62751, P50518, O08992, G5E829, P49615, P98086, P57716, Q9D1R9, P35564, Q64430, P62270, Q8JZQ9, P63276, O55101, O55142, P61255, P58252, Q9D8E6, P62900, Q9D0E1, P47964, P41105 | 0.00011942339 |
| GOTERM_CC_DIRECT | GO:0098794~postsynapse | 17 | 4.7752809 | 3.93723E-06 | P62918, Q9D1R9, P19253, P62281, P62270, P14115, P63276, O55142, P67984, P62751, P61255, Q9D8E6, P62900, P98086, P41105, P47964, P49615 | 0.00012233546 |

| Annotation Cluster 2 | Enrichment Score: 6.992234447958213 | | | | | |
|---|---|---|---|---|---|---|
| Category | Term | Count | % | PValue | Genes | FDR |
| GOTERM_MF_DIRECT | GO:0000166~nucleotide binding | 64 | 17.977528 | 5.94397E-11 | Q8CG48, Q8BTI9, O70133, P46471, P70388, P37913, P11440, Q8K0D5, Q61035, Q8VEH6, P70704, P61222, P61028, P09411, P48722, P61027, P56480, P49615, P28650, Q9WUK4, Q64430, P70698, P35601, Q8BP47, P09581, Q9WTI7, P06795, E9Q634, Q9JJY4, O70551, Q2NL51, Q9CY64, O08528, P50516, P70248, P36371, Q8VDD5, Q91V92, Q04899, P61087, P04184, G5E829, P62334, Q8CI11, P80314, Q8VI93, Q9CZ30, Q8C0D5, Q6ZQB6, Q04692, O35379, P58252, O09110, P24547, Q9EPE9, Q9Z110, Q9D1G2, P63038, Q9WUA3, P80318, Q9JHW4, Q99KK9, P60843, Q3UFY7 | 1.9644822E-08 |
| GOTERM_MF_DIRECT | GO:0005524~ATP binding | 53 | 14.88764 | 4.83696E-08 | Q8CG48, P11928, Q8BTI9, O70133, P46471, P70388, P37913, P11440, Q61035, P70704, P61222, P09411, P48722, P56480, P49615, Q9WUK4, Q64430, P70698, P35601, Q8BP47, P09581, Q9WTI7, P06795, E9Q634, Q9JJY4, O70551, Q2NL51, O08528, P50516, P70248, P36371, Q8VDD5, Q91V92, Q04899, P61087, P04184, G5E829, P62334, P80314, Q8VI93, Q9CZ30, Q6ZQB6, Q04692, O35379, O09110, Q9EPE9, Q9Z110, Q9D1G2, P63038, Q9WUA3, P80318, Q99KK9, P60843 | 6.3944586E-06 |

| | | | | | | |
|---|---|---|---|---|---|---|
| UP_KW_LIGAND | KW-0547~Nucleotide-binding | 63 | 17.696629 | 2.08501E-07 | Q8CG48, P11928, Q8BTI9, O70133, P46471, P70388, P37913, P11440, Q8K0D5, Q61035, Q8VEH6, P70704, P61222, P61028, P09411, P48722, P61027, P56480, P49615, P28650, Q9WUK4, Q64430, P70698, P35601, Q8BP47, P09581, Q9WTI7, P06795, E9Q634, Q9JJY4, O70551, Q2NL51, O08528, P50516, P70248, P36371, Q8VDD5, Q91V92, Q04899, P61087, P04184, G5E829, P62334, Q8CI11, P80314, Q8VI93, Q9CZ30, Q8C0D5, Q6ZQB6, Q04692, O35379, P58252, O09110, Q9EPE9, Q9Z110, Q9D1G2, P63038, Q9WUA3, P80318, Q9JHW4, Q99KK9, P60843, Q3UFY7 | 4.5599432E-06 |
| UP_KW_LIGAND | KW-0067~ATP-binding | 53 | 14.88764 | 3.64795E-07 | Q8CG48, P11928, Q8BTI9, O70133, P46471, P70388, P37913, P11440, Q61035, P70704, P61222, P09411, P48722, P56480, P49615, Q9WUK4, Q64430, P70698, P35601, Q8BP47, P09581, Q9WTI7, P06795, E9Q634, Q9JJY4, O70551, Q2NL51, O08528, P50516, P70248, P36371, Q8VDD5, Q91V92, Q04899, P61087, P04184, G5E829, P62334, P80314, Q8VI93, Q9CZ30, Q6ZQB6, Q04692, O35379, O09110, Q9EPE9, Q9Z110, Q9D1G2, P63038, Q9WUA3, P80318, Q99KK9, P60843 | 4.5599432E-06 |
| GOTERM_MF_DIRECT | GO:0016887~ATPase activity | 21 | 5.8988764 | 1.0894E-06 | P80314, Q8CG48, P46471, O70133, Q9CZ30, P70388, Q9WUK4, Q64430, P35601, Q04692, P70704, P48722, Q9EPE9, G5E829, P56480, P62334, P63038, P80318, E9Q634, Q9JJY4, P60843 | 0.00012001558 |
| INTERPRO | IPR027417:P-loop containing nucleoside triphosphate hydrolase | 33 | 9.2696629 | 4.67304E-06 | Q8CG48, O70133, P46471, Q9JKF1, P50516, P70388, P70248, Q8K0D5, P36371, Q8VDD5, P61222, P61028, P04184, P61027, P62334, P56480, Q8CI11, P28650, Q9CZ30, Q8C0D5, Q9WUK4, P70698, P35601, Q04692, O35379, P58252, Q9WTI7, Q9D1G2, P06795, Q9JJY4, Q9JHW4, E9Q634, P60843 | 0.00165425601 |

| Annotation Cluster 3 | Enrichment Score: 5.913199641887708 | | | | | |
|---|---|---|---|---|---|---|
| Category | Term | Count | % | PValue | Genes | FDR |
| UP_KW_DOMAIN | KW-0809~Transit peptide | 25 | 7.0224719 | 5.08189E-08 | Q8K3J1, Q8K0D5, P42125, Q9CZB0, Q9CZ13, P56383, P45952, P56480, P08249, Q9DB77, Q9D6R2, Q9CYR0, P53395, Q8CG76, O08715, Q8BMF4, Q9JHI5, Q8C0I1, P20108, Q9CQA3, O09111, Q99KI0, P63038, Q99KK9, Q9CQE3 | 1.0671968E-06 |
| UP_SEQ_FEATURE | TRANSIT:Mitochondrion | 24 | 6.741573 | 5.09651E-07 | Q9D6R2, Q8K3J1, Q9CYR0, P53395, Q8CG76, O08715, Q9JHI5, Q8BMF4, Q8K0D5, P20108, P42125, Q9CQA3, O09111, Q9CZB0, Q99KI0, Q9CZ13, P56383, P45952, P56480, P63038, P08249, Q9DB77, Q99KK9, Q9CQE3 | 0.00016376794 |

| | | | | | | |
|---|---|---|---|---|---|---|
| UP_KW_CELLULAR_COMPONENT | KW-0496~Mitochondrion | 39 | 10.955056 | 7.03239E-05 | P11928, Q8K3J1, O08528, P11440, Q8K0D5, P42125, P61222, Q9CZB0, Q9CZ13, Q9DCZ4, P56383, P45952, P56480, P08249, Q8R1I1, Q9DB77, P62897, Q9D6R2, Q921M7, Q9QZD8, Q9CYR0, P53395, Q8CG76, O08715, Q8BMF4, Q9JHI5, O08734, P20108, Q8BGE6, Q9CQA3, O09111, Q07813, Q99KI0, Q9Z110, P63038, P60603, Q99KK9, Q9CQE3, P00397 | 0.00093765164 |
| Annotation Cluster 4 | Enrichment Score: 4.8231446763019905 | | | | | |
| Category | Term | Count | % | PValue | Genes | FDR |
| KEGG_PATHWAY | mmu01200:Carbon metabolism | 16 | 4.494382 | 1.062E-07 | Q9D6R2, P06801, P06745, Q9R0P3, O08528, Q8BMF4, P47968, Q9CQA3, P40142, Q9CZB0, Q99KI0, P05063, P09411, Q9WUA3, P08249, P05201 | 9.2706324E-06 |
| UP_KW_BIOLOGICAL_PROCESS | KW-0816~Tricarboxylic acid cycle | 6 | 1.6853933 | 4.609E-05 | Q9D6R2, Q9CQA3, Q9CZB0, Q99KI0, Q8BMF4, P08249 | 0.0013366095 |
| KEGG_PATHWAY | mmu00020:Citrate cycle (TCA cycle) | 7 | 1.9662921 | 7.72258E-05 | Q9D6R2, Q9CQA3, Q91V92, Q9CZB0, Q99KI0, Q8BMF4, P08249 | 0.00197905107 |
| GOTERM_BP_DIRECT | GO:0006099~tricarboxylic acid cycle | 6 | 1.6853933 | 0.000134874 | Q9D6R2, Q9CQA3, Q9CZB0, Q99KI0, Q8BMF4, P08249 | 0.0437664787 |
| Annotation Cluster 5 | Enrichment Score: 4.704741333913531 | | | | | |
| Category | Term | Count | % | PValue | Genes | FDR |
| UP_SEQ_FEATURE | CROSSLNK:Glycyl lysine isopeptide (Lys-Gly) (interchain with G-Cter in SUMO2) | 36 | 10.11236 | 2.6101E-07 | P62918, P09405, P62717, O70133, Q3U1J4, P14733, P10852, P97822, P11440, P62751, O35226, P40142, Q99JF8, P62996, Q8CI11, Q9DBE9, P80314, Q60973, Q9D1R9, Q6PDM2, Q99020, P62270, Q62376, Q8CGZ0, Q91VE6, Q3UEB3, P35601, Q04692, P61255, Q9D8E6, P24547, Q9D0E1, P63038, P80318, P41105, P60843 | 0.00012580665 |
| UP_KW_PTM | KW-0832~Ubl conjugation | 73 | 20.505618 | 0.000159532 | P62918, P09405, Q8BL97, P62717, O70133, P46471, Q3U1J4, Q9JMH6, P97822, P11440, Q9QY76, Q80U87, Q9R0L6, P62751, P61222, P61027, A1L314, P84228, P62996, P06745, Q91YQ5, Q9D1R9, Q6PDM2, P35564, Q9CQ71, Q62376, P35601, P14685, Q61093, P09581, Q9QXK7, P14733, P10852, Q8VDD5, O35226, P40142, Q91V92, P61087, P17095, P04184, Q99JF8, Q8CI11, Q9DBE9, P80314, Q60972, Q60973, Q921M7, Q99020, P36993, P62270, Q8CGZ0, Q91VE6, Q3UEB3, Q04692, P63276, Q8BGE6, P58252, P61255, Q9D8E6, P24547, Q9D0E1, P63038, P80318, P41105, P60843 | 0.00066462281 |

| | | | | | | |
|---|---|---|---|---|---|---|
| UP_KW_PTM | KW-1017~Isopeptide bond | 54 | 15.168539 | 0.000184617 | P62918, P09405, Q8BL97, P62717, O70133, Q3U1J4, P97822, P11440, Q9QY76, P62751, P61027, P62996, Q91YQ5, Q9D1R9, Q6PDM2, Q9CQ71, Q62376, P35601, P14685, Q61093, Q9QXK7, P14733, P10852, O35226, P40142, Q91V92, P61087, P17095, Q99JF8, Q8CI11, Q9DBE9, P80314, Q60972, Q60973, Q921M7, Q99020, P36993, P62270, Q8CGZ0, Q91VE6, Q3UEB3, Q04692, P63276, Q8BGE6, P58252, P61255, Q9D8E6, P24547, Q9D0E1, P63038, P80318, P41105, P60843 | 0.00066462281 |
| Annotation Cluster 6 | Enrichment Score: 4.24596503935281 | | | | | |
| Category | Term | Count | % | PValue | Genes | FDR |
| GOTERM_CC_DIRECT | GO:0005783~endoplasmic reticulum | 49 | 13.764045 | 1.88325E-06 | Q60664, P11928, Q9JLJ5, P14115, O08547, Q9QY76, O08795, P70704, O08992, P62274, P61027, Q9DCZ4, O88455, Q8BHN3, Q91YQ5, Q9D1R9, O08715, P35564, P17439, Q64430, Q8R180, Q9DB25, O54692, Q9JK23, Q07813, Q61093, Q9DBS1, Q9WV55, O70551, Q9CPU4, P36371, Q6ZWQ7, Q9JHP7, P35441, Q9WU81, Q9EQH2, P57716, Q8CGZ0, O08734, P97858, P35821, Q8VDP6, Q6P5E4, Q8BGE6, Q8R1V4, B9EJ86, Q9EPE9, O70503, P47964 | 7.4473923E-05 |
| UP_KW_CELLULAR_COMPONENT | KW-0256~Endoplasmic reticulum | 41 | 11.516854 | 1.26401E-05 | Q60664, P11928, Q9WV55, O70551, Q9JLJ5, O08547, Q9QY76, Q9CPU4, O08795, P36371, P70704, Q6ZWQ7, Q9JHP7, P35441, O08992, P62274, P61027, Q9DCZ4, Q9WU81, O88455, Q8BHN3, Q9EQH2, Q9D1R9, Q91YQ5, O08715, P35564, Q8CGZ0, P97858, Q8R180, P35821, Q8VDP6, Q6P5E4, Q8BGE6, Q9DB25, O54692, Q9JK23, Q8R1V4, B9EJ86, Q9EPE9, O70503, Q9DBS1 | 0.00025280251 |
| GOTERM_CC_DIRECT | GO:0005789~endoplasmic reticulum membrane | 25 | 7.0224719 | 0.007681496 | Q60664, Q9WV55, Q9JLJ5, O08547, Q9QY76, Q9CPU4, P36371, P40142, O08992, P61027, Q9DCZ4, Q9WU81, O88455, Q9EQH2, Q91YQ5, P35564, P97858, Q8R180, Q8VDP6, Q9DB25, O54692, Q07813, Q8R1V4, B9EJ86, Q9EPE9 | 0.09030948567 |
| Annotation Cluster 7 | Enrichment Score: 4.186778511904929 | | | | | |
| Category | Term | Count | % | PValue | Genes | FDR |
| GOTERM_MF_DIRECT | GO:0016491~oxidoreductase activity | 27 | 7.5842697 | 4.68133E-06 | Q8K3J1, Q9CY64, Q9JMH6, Q8C7K6, Q9ESY9, Q9CPU4, Q9CQF9, P45376, P45377, O88455, P45952, P08249, Q9D6R2, P06801, P08228, P47199, Q8CG76, Q9JHI5, Q8R180, Q8C0I1, P20108, Q9CQA3, Q61093, Q9Z2G9, P24547, Q9Z110, O70503 | 0.00044205154 |

| | | | | | | |
|---|---|---|---|---|---|---|
| UP_KW_MOLECULAR_FUNCTION | KW-0560~Oxidoreductase | 27 | 7.5842697 | 8.63966E-05 | Q8K3J1, Q9CY64, Q9JMH6, Q8C7K6, Q9ESY9, Q9CPU4, Q9CQF9, P45376, P45377, O88455, P45952, P08249, Q9D6R2, P06801, P08228, P47199, Q8CG76, Q9JHI5, Q8R180, P20108, Q9CQA3, Q61093, Q9Z2G9, P24547, Q9Z110, O70503, P00397 | 0.00095036298 |
| UP_KW_LIGAND | KW-0521~NADP | 12 | 3.3707865 | 0.000680453 | P06801, P47199, Q61093, Q9CY64, Q8CG76, Q9Z2G9, Q9JMH6, P45376, O88455, Q9Z110, P45377, O70503 | 0.0056704419 |

| Annotation Cluster 8 | Enrichment Score: 3.7635340237459953 | | | | | |
|---|---|---|---|---|---|---|
| Category | Term | Count | % | PValue | Genes | FDR |
| GOTERM_MF_DIRECT | GO:0003723~RNA binding | 49 | 13.764045 | 1.10561E-13 | P62918, P09405, Q8BL97, Q9D7S7, Q6R5N8, O70133, O70194, P62281, O35295, Q9JKP5, O08795, P67984, P62751, P84104, Q9CSH3, P17095, P62996, Q8VI93, Q60973, P47199, Q8CI75, Q6PDM2, O08715, Q99J36, Q99020, Q9D7A6, P62270, Q62376, Q8CGZ0, Q9JII5, Q91VE6, Q3UEB3, Q99JX7, Q9CQT2, Q8JZQ9, P70318, Q9CXK8, Q5U4D9, P58252, P61255, Q9D8E6, Q9QXK7, P24547, Q9D0E1, Q91VU7, Q9JJY4, P60843, Q9CQE3 | 7.3080929E-11 |
| UP_KW_MOLECULAR_FUNCTION | KW-0694~RNA-binding | 37 | 10.393258 | 1.67834E-08 | P62918, P11928, P09405, Q8BL97, Q6R5N8, O70133, O70194, P62281, Q9JKP5, P67984, P62751, Q9CSH3, P84104, P62996, Q8VI93, P47199, Q8CI75, Q6PDM2, O08715, Q9D7A6, Q99020, P62270, Q62376, Q9JII5, Q91VE6, Q3UEB3, Q99JX7, Q9CQT2, Q8JZQ9, P70318, Q9CXK8, Q5U4D9, P24547, Q9QXK7, Q9D0E1, P60843, Q9CQE3 | 3.6923401E-07 |
| UP_SEQ_FEATURE | DOMAIN:RRM | 15 | 4.2134831 | 2.78508E-06 | P09405, Q8BL97, Q6PDM2, Q99020, Q62376, Q9JII5, Q91VE6, Q3UEB3, Q99JX7, Q9CQT2, Q8JZQ9, P70318, P84104, Q9D0E1, P62996 | 0.00053696315 |
| SMART | SM00360:RRM | 14 | 3.9325843 | 9.46225E-06 | P09405, Q8BL97, Q6PDM2, Q99020, Q62376, Q9JII5, Q91VE6, Q3UEB3, Q9CQT2, Q8JZQ9, P70318, P84104, Q9D0E1, P62996 | 0.00058665931 |
| INTERPRO | IPR012677:Nucleotide-binding, alpha-beta plait | 16 | 4.494382 | 1.18415E-05 | P09405, Q8BL97, Q6PDM2, Q99020, Q62376, Q9JII5, Q91VE6, Q3UEB3, Q99JX7, Q9CQT2, Q8JZQ9, P70318, P62751, P84104, Q9D0E1, P62996 | 0.00279458871 |
| INTERPRO | IPR000504:RNA recognition motif domain | 14 | 3.9325843 | 4.43036E-05 | P09405, Q8BL97, Q6PDM2, Q99020, Q62376, Q9JII5, Q91VE6, Q3UEB3, Q9CQT2, Q8JZQ9, P70318, P84104, Q9D0E1, P62996 | 0.0072113923 |
| GOTERM_MF_DIRECT | GO:0003676~nucleic acid binding | 23 | 6.4606742 | 7.40358E-05 | P09405, Q8BL97, O70133, Q3U1J4, Q6PDM2, O08715, P62270, Q99020, Q62376, Q8CGZ0, Q9JII5, Q91VE6, Q3UEB3, Q99JX7, Q9CQT2, Q8JZQ9, P70318, Q8BP47, P84104, Q9D0E1, Q9JJY4, P62996, P60843 | 0.00611720948 |

| Annotation Cluster 9 | Enrichment Score: 3.477831332166054 | | | | | |
|---|---|---|---|---|---|---|
| Category | Term | Count | % | PValue | Genes | FDR |
| UP_SEQ_FEATURE | LIPID:N6-decanoyllysine | 8 | 2.247191 | 1.78707E-10 | P84228 | 1.7227368E-07 |
| SMART | SM00428:H3 | 8 | 2.247191 | 6.17874E-10 | P84228 | 7.6616402E-08 |
| INTERPRO | IPR000164:Histone H3 | 8 | 2.247191 | 1.22785E-09 | P84228 | 8.693177E-07 |
| UP_KW_PTM | KW-0164~Citrullination | 15 | 4.2134831 | 1.17666E-08 | Q9DBE9, P62751, Q9D8E6, P19253, P62281, P84228, Q99JF8, Q91VE6 | 7.0599853E-08 |
| GOTERM_BP_DIRECT | GO:0040029~regulation of gene expression, epigenetic | 8 | 2.247191 | 2.12865E-06 | P84228 | 0.00103612214 |
| UP_SEQ_FEATURE | DOMAIN:Histone H2A/H2B/H3 | 8 | 2.247191 | 2.2727E-06 | P84228 | 0.00053696315 |
| GOTERM_BP_DIRECT | GO:0006334~nucleosome assembly | 11 | 3.0898876 | 6.53341E-06 | Q60972, Q78ZA7, Q9EST5, P84228 | 0.00254411153 |
| INTERPRO | IPR007125:Histone core | 8 | 2.247191 | 9.4574E-05 | P84228 | 0.01115973651 |
| GOTERM_CC_DIRECT | GO:0000785~chromatin | 18 | 5.0561798 | 0.000206828 | Q60972, Q8CG48, Q8BFQ4, O35864, Q78ZA7, O70551, P17095, P70388, P84228, P42227 | 0.00449850639 |
| UP_KW_CELLULAR_COMPONENT | KW-0544~Nucleosome core | 8 | 2.247191 | 0.000616416 | P84228 | 0.00616416025 |
| UP_KW_PTM | KW-0013~ADP-ribosylation | 10 | 2.8089888 | 0.000654427 | P17095, P84228 | 0.00196328195 |
| UP_SEQ_FEATURE | LIPID:S-palmitoyl cysteine | 12 | 3.3707865 | 0.000909053 | P08228, P35564, P62281, P84228, Q9CPU4 | 0.08763272931 |
| GOTERM_MF_DIRECT | GO:0046982~protein heterodimerization activity | 16 | 4.494382 | 0.001136358 | Q9WV55, Q9R1T2, Q07813, O08992, Q61093, Q6IRU2, O08734, P84228, Q9QY76 | 0.06259440522 |
| UP_KW_PTM | KW-0379~Hydroxylation | 12 | 3.3707865 | 0.001399025 | P62918, P09411, P14115, P84228, P98086 | 0.00314780535 |

| Annotation Cluster 10 | Enrichment Score: 3.42643241519865 | | | | | |
|---|---|---|---|---|---|---|
| Category | Term | Count | % | PValue | Genes | FDR |
| KEGG_PATHWAY | mmu05020:Prion disease | 21 | 5.8988764 | 3.51395E-06 | P08228, P46471, Q8BTI9, Q8K3J1, P14685, Q99JI4, Q9CQA3, O35226, O09111, Q07813, Q61093, Q9CZB0, Q9CZ13, P56383, P56480, P62334, Q8R1I1, Q9DB77, P98086, P62897, P00397 | 0.00021874329 |
| UP_KW_BIOLOGICAL_PROCESS | KW-0249~Electron transport | 11 | 3.0898876 | 1.16712E-05 | Q9CQA3, O09111, Q61093, Q9CZB0, Q8K3J1, Q9CZ13, Q8R1I1, Q9DB77, Q8R180, P62897, P00397 | 0.00061912631 |
| KEGG_PATHWAY | mmu05014:Amyotrophic lateral sclerosis | 24 | 6.741573 | 1.31115E-05 | P08228, Q8BL97, Q8R0G9, P46471, Q8K3J1, Q99JX7, P14685, Q9QY76, Q99JI4, Q9CQA3, O35226, O09111, Q07813, O09110, Q9CZB0, P84104, Q9CZ13, P56383, P56480, P62334, Q8R1I1, Q9DB77, P62897, P00397 | 0.00065295464 |
| KEGG_PATHWAY | mmu00190:Oxidative phosphorylation | 13 | 3.6516854 | 6.3789E-05 | Q8K3J1, P50516, Q9CQA3, O09111, P50518, Q9CZB0, Q9CZ13, P56383, P56480, Q8R1I1, Q9DB77, P62897, P00397 | 0.00197905107 |

| | | | | | | |
|---|---|---|---|---|---|---|
| KEGG_PATHWAY | mmu05010:Alzheimer disease | 23 | 6.4606742 | 7.06135E-05 | P57716, Q91ZX7, P46471, Q8BTI9, Q8K3J1, Q9CR16, P14685, Q99JI4, Q9CQA3, O35226, P11152, O09111, Q61093, Q9CZB0, Q9CZ13, P56383, P56480, P62334, Q8R1I1, Q9DB77, P62897, P49615, P00397 | 0.00197905107 |
| KEGG_PATHWAY | mmu05012:Parkinson disease | 18 | 5.0561798 | 0.000125524 | P08228, P46471, Q8K3J1, P14685, Q99JI4, Q9CQA3, O35226, O09111, Q07813, Q9CZB0, Q9CZ13, P56383, P56480, P62334, Q8R1I1, Q9DB77, P62897, P00397 | 0.00284141034 |
| KEGG_PATHWAY | mmu05016:Huntington disease | 19 | 5.3370787 | 0.000208078 | P08228, P46471, Q8K3J1, P17426, P14685, Q99JI4, Q9CQA3, O35226, O09111, Q07813, Q9CZB0, Q9CZ13, P56383, P56480, P62334, Q8R1I1, Q9DB77, P62897, P00397 | 0.00431762676 |
| GOTERM_CC_DIRECT | GO:0070469~respiratory chain | 7 | 1.9662921 | 0.000243472 | O09111, Q8K3J1, Q9CZ13, Q8R1I1, Q9DB77, P62897, P00397 | 0.00504335103 |
| KEGG_PATHWAY | mmu05022:Pathways of neurodegeneration - multiple diseases | 24 | 6.741573 | 0.000525299 | P08228, P46471, Q8K3J1, Q9CR16, O08734, P14685, Q9QY76, Q99JI4, Q9CQA3, O35226, O09111, Q07813, O09110, Q61093, Q9CZB0, Q9CZ13, P56383, P56480, P62334, Q8R1I1, Q9DB77, P62897, P49615, P00397 | 0.01006149919 |
| KEGG_PATHWAY | mmu05208:Chemical carcinogenesis - reactive oxygen species | 15 | 4.2134831 | 0.000613365 | Q60631, P08228, Q8BTI9, Q8K3J1, P35821, Q9CPU4, Q9CQA3, O09111, Q9CZB0, Q9CZ13, P56383, P56480, Q8R1I1, Q9DB77, P00397 | 0.01090913552 |
| UP_KW_BIOLOGICAL_PROCESS | KW-0679~Respiratory chain | 7 | 1.9662921 | 0.000616075 | O09111, Q8K3J1, Q9CZ13, Q8R1I1, Q9DB77, P62897, P00397 | 0.01071970068 |
| KEGG_PATHWAY | mmu04932:Non-alcoholic fatty liver disease | 12 | 3.3707865 | 0.000951092 | Q9CQA3, O09111, Q07813, Q8BTI9, Q9CZB0, Q2NL51, Q8K3J1, Q9CZ13, Q8R1I1, Q9DB77, P62897, P00397 | 0.01480136181 |
| GOTERM_CC_DIRECT | GO:0005743~mitochondrial inner membrane | 18 | 5.0561798 | 0.001485937 | Q9QZD8, Q8K3J1, P42125, Q9CQA3, O09111, Q9CZB0, Q9CZ13, Q9DCZ4, Q9Z110, P56480, P63038, P60603, P08249, Q8R1I1, Q9DB77, P42227, P00397, Q9CQE3 | 0.02394009673 |
| UP_KW_CELLULAR_COMPONENT | KW-0999~Mitochondrion inner membrane | 13 | 3.6516854 | 0.005255569 | Q9QZD8, Q8K3J1, Q9CQA3, O09111, Q9CZB0, Q9CZ13, Q9DCZ4, Q9Z110, P56480, P60603, Q8R1I1, Q9DB77, P00397 | 0.03003182093 |

| Annotation Cluster 11 | Enrichment Score: 3.082004013922556 | | | | | |
|---|---|---|---|---|---|---|
| Category | Term | Count | % | PValue | Genes | FDR |
| UP_KW_BIOLOGICAL_PROCESS | KW-0249~Electron transport | 11 | 3.0898876 | 1.16712E-05 | Q9CQA3, O09111, Q61093, Q9CZB0, Q8K3J1, Q9CZ13, Q8R1I1, Q9DB77, Q8R180, P62897, P00397 | 0.00061912631 |
| GOTERM_CC_DIRECT | GO:0070469~respiratory chain | 7 | 1.9662921 | 0.000243472 | O09111, Q8K3J1, Q9CZ13, Q8R1I1, Q9DB77, P62897, P00397 | 0.00504335103 |
| GOTERM_CC_DIRECT | GO:0005750~mitochondrial respiratory chain complex III | 4 | 1.1235955 | 0.000564085 | Q9CZ13, Q8R1I1, Q9DB77, P00397 | 0.01066855759 |
| UP_KW_BIOLOGICAL_PROCESS | KW-0679~Respiratory chain | 7 | 1.9662921 | 0.000616075 | O09111, Q8K3J1, Q9CZ13, Q8R1I1, Q9DB77, P62897, P00397 | 0.01071970068 |

| | | | | | | |
|---|---|---|---|---|---|---|
| Annotation Cluster 12 | Enrichment Score: 2.6449171107640685 | | | | | |
| Category | Term | Count | % | PValue | Genes | FDR |
| UP_KW_BIOLOGICAL_PROCESS | KW-0648~Protein biosynthesis | 12 | 3.3707865 | 1.42328E-05 | P58252, Q8BP47, Q9DCH4, O70194, Q8C0D5, P60229, Q8K0D5, Q9JHW4, P60843, Q99KK9, Q61035, Q8JZQ9 | 0.00061912631 |
| GOTERM_CC_DIRECT | GO:0033290~eukaryotic 48S preinitiation complex | 4 | 1.1235955 | 0.001130424 | Q9DCH4, O70194, P60229, Q8JZQ9 | 0.02048892849 |
| GOTERM_CC_DIRECT | GO:0005852~eukaryotic translation initiation factor 3 complex | 4 | 1.1235955 | 0.001376767 | Q9DCH4, O70194, P60229, Q8JZQ9 | 0.02394009673 |
| GOTERM_BP_DIRECT | GO:0001732~formation of cytoplasmic translation initiation complex | 4 | 1.1235955 | 0.001561436 | Q9DCH4, O70194, P60229, Q8JZQ9 | 0.19318868947 |
| GOTERM_CC_DIRECT | GO:0016282~eukaryotic 43S preinitiation complex | 4 | 1.1235955 | 0.001654348 | Q9DCH4, O70194, P60229, Q8JZQ9 | 0.02570147671 |
| Annotation Cluster 13 | Enrichment Score: 2.5479359384965816 | | | | | |
| Category | Term | Count | % | PValue | Genes | FDR |
| GOTERM_MF_DIRECT | GO:0017124~SH3 domain binding | 9 | 2.5280899 | 0.001030467 | Q60631, Q9WU78, P29351, Q8BIJ7, Q5FWK3, Q6P549, Q8BHL5, Q8BPU7, Q80U87 | 0.06192171506 |
| Annotation Cluster 14 | Enrichment Score: 2.420377273906477 | | | | | |
| Category | Term | Count | % | PValue | Genes | FDR |
| GOTERM_CC_DIRECT | GO:0022624~proteasome accessory complex | 5 | 1.4044944 | 8.13869E-05 | Q99JI4, O35226, P46471, P62334, P14685 | 0.00208254785 |
| UP_KW_CELLULAR_COMPONENT | KW-0647~Proteasome | 6 | 1.6853933 | 0.001433291 | Q99JI4, O35226, P46471, P54728, P62334, P14685 | 0.01146632984 |
| GOTERM_CC_DIRECT | GO:0000502~proteasome complex | 6 | 1.6853933 | 0.001850621 | Q99JI4, O35226, P46471, P54728, P62334, P14685 | 0.02775930834 |
| Annotation Cluster 15 | Enrichment Score: 2.1450240188303296 | | | | | |
| Category | Term | Count | % | PValue | Genes | FDR |
| UP_KW_BIOLOGICAL_PROCESS | KW-0324~Glycolysis | 5 | 1.4044944 | 0.002224592 | P06745, P05063, O08528, P09411, Q9WUA3 | 0.03225658557 |
| Annotation Cluster 16 | Enrichment Score: 1.7525834387892805 | | | | | |
| Category | Term | Count | % | PValue | Genes | FDR |

| | | | | | | |
|---|---|---|---|---|---|---|
| UP_KW_LIGAND | KW-0274~FAD | 8 | 2.247191 | 0.009548578 | Q8C0I1, Q9CQF9, Q61093, Q9JMH6, Q8C7K6, P45952, Q9JHI5, Q8R180 | 0.05412698994 |
| UP_KW_LIGAND | KW-0285~Flavoprotein | 8 | 2.247191 | 0.010825398 | Q8C0I1, Q9CQF9, Q61093, Q9JMH6, Q8C7K6, P45952, Q9JHI5, Q8R180 | 0.05412698994 |
| Annotation Cluster 17 | Enrichment Score: 1.6746771580234787 | | | | | |
| Category | Term | Count | % | PValue | Genes | FDR |
| INTERPRO | IPR017900:4Fe-4S ferredoxin, iron-sulphur binding, conserved site | 3 | 0.8426966 | 0.001217217 | Q9CQA3, P61222, Q8K3J1 | 0.09631275432 |
| Annotation Cluster 18 | Enrichment Score: 1.6476167193215445 | | | | | |
| Category | Term | Count | % | PValue | Genes | FDR |
| UP_KW_BIOLOGICAL_PROCESS | KW-0509~mRNA transport | 7 | 1.9662921 | 0.003806516 | Q5U4D9, Q8BL97, Q8R0G9, O70133, Q6PDM2, P84104, Q99JX7 | 0.04730955162 |
| Annotation Cluster 19 | Enrichment Score: 1.5583078211177934 | | | | | |
| Category | Term | Count | % | PValue | Genes | FDR |
| GOTERM_CC_DIRECT | GO:0005764~lysosome | 16 | 4.494382 | 0.003315368 | P08228, P57716, P23780, Q9Z0M5, P11438, P10852, Q9ESY9, P50516, P17439, P70318, Q9CQF9, P24668, P45377, Q3TCN2, Q8VEB4, P05201 | 0.04652210261 |
| Annotation Cluster 20 | Enrichment Score: 1.5036738713049727 | | | | | |
| Category | Term | Count | % | PValue | Genes | FDR |
| GOTERM_CC_DIRECT | GO:0015629~actin cytoskeleton | 11 | 3.0898876 | 0.003808661 | P13020, P21107, Q8VDD5, O70133, Q9WTI7, Q6IRU2, Q9JKF1, P59999, P70248, E9Q634, O70318 | 0.05177398992 |
| Annotation Cluster 21 | Enrichment Score: 1.4497999795318788 | | | | | |
| Category | Term | Count | % | PValue | Genes | FDR |
| UP_KW_BIOLOGICAL_PROCESS | KW-0249~Electron transport | 11 | 3.0898876 | 1.16712E-05 | Q9CQA3, O09111, Q61093, Q9CZB0, Q8K3J1, Q9CZ13, Q8R1I1, Q9DB77, Q8R180, P62897, P00397 | 0.00061912631 |
| Annotation Cluster 22 | Enrichment Score: 1.4168460123066215 | | | | | |
| Category | Term | Count | % | PValue | Genes | FDR |

| | | | | | | |
|---|---|---|---|---|---|---|
| SMART | SM00382:AAA | 9 | 2.5280899 | 0.00065347 | P36371, O35379, P61222, P46471, Q9WUK4, P62334, P56480, P06795, P35601 | 0.02701010409 |
| INTERPRO | IPR003593:AAA+ ATPase domain | 9 | 2.5280899 | 0.001224315 | P36371, O35379, P61222, P46471, Q9WUK4, P62334, P56480, P06795, P35601 | 0.09631275432 |

| Annotation Cluster 23 | Enrichment Score: 1.2694767495088586 | | | | | |
|---|---|---|---|---|---|---|
| Category | Term | Count | % | PValue | Genes | FDR |
| KEGG_PATHWAY | mmu03430:Mismatch repair | 5 | 1.4044944 | 0.001455378 | Q9CYR0, Q9CQ71, P37913, Q9WUK4, P35601 | 0.02131701364 |
| KEGG_PATHWAY | mmu03420:Nucleotide excision repair | 6 | 1.6853933 | 0.0029459 | Q3U1J4, P54728, Q9CQ71, P37913, Q9WUK4, P35601 | 0.04075161853 |
| UP_KW_BIOLOGICAL_PROCESS | KW-0235~DNA replication | 7 | 1.9662921 | 0.004466235 | Q60972, Q60973, Q9CYR0, Q9CQ71, P37913, Q9WUK4, P35601 | 0.0485703026 |
| KEGG_PATHWAY | mmu03030:DNA replication | 5 | 1.4044944 | 0.008247428 | Q9CYR0, Q9CQ71, P37913, Q9WUK4, P35601 | 0.0977909279 |
