## supplemental Table 4 for "Biobased, Biodegradable but not bio-neutral: about the effects of polylactic acid nanoparticles on macrophages"

Supplementary Table 4: list of mitochondrial proteins modulated in response to PLA beads

| Uniprot accession | Protein name | ratio PLA treated/control | Mann Whitney U value |
|---|---|---|---|
| O08528 | Hexokinase-2 | 1.09 | 2 |
| O08715 | A-kinase anchor protein 1, mitochondrial | 2.1 | 1 |
| O08734 | Bcl-2 homologous antagonist/killer | 2.1 | 2 |
| O09111 | NADH dehydrogenase [ubiquinone] 1 beta subcomplex subunit 11, mitochondrial | 0.78 | 2 |
| P00397 | Cytochrome c oxidase subunit 1 | 1.84 | 2 |
| P08249 | Malate dehydrogenase, mitochondrial | 0.79 | 2 |
| P11440 | Cyclin-dependent kinase 1 | 1.48 | 0 |
| P11928 | 2'-5'-oligoadenylate synthase 1A | 2.05 | 2 |
| P20108 | Peroxiredoxin 3 | 1.21 | 0 |
| P42125 | Enoyl-CoA delta isomerase 1, mitochondrial | 0.65 | 1 |
| P45952 | Medium-chain specific acyl-CoA dehydrogenase, mitochondrial | 0.89 | 2 |
| P53395 | Lipoamide acyltransferase component of branched-chain alpha-keto acid dehydrogenase complex, mitochondrial | 1.47 | 1 |
| P56383 | ATP synthase F(0) complex subunit C2, mitochondrial | 0.42 | 2 |
| P56480 | ATP synthase subunit beta, mitochondrial | 1.18 | 0 |
| P60603 | Reactive oxygen species modulator 1 | 1.29 | 2 |
| P61222 | ATP-binding cassette sub-family E member 1 | 1.12 | 2 |
| P62897 | Cytochrome c, somatic | 0.79 | 0 |
| P63038 | 60 kDa heat shock protein, mitochondrial | 0.86 | 2 |
| Q07813 | Apoptosis regulator BAX | 1.16 | 0 |
| Q6UPE1 | Electron transfer flavoprotein-ubiquinone oxidoreductase, mitochondrial | 1.42 | 1 |
| Q8BGE6 | Cysteine protease ATG4B | 1.44 | 0 |
| Q8BMF4 | Dihydrolipoyllysine-residue acetyltransferase component of pyruvate dehydrogenase complex, mitochondrial | 0.8 | 1 |
| Q8CG76 | Aflatoxin B1 aldehyde reductase member 2 | 1.64 | 2 |
| Q8K0D5 | Elongation factor G, mitochondrial | 1.27 | 2 |
| Q8K3J1 | NADH dehydrogenase [ubiquinone] iron-sulfur protein 8, mitochondrial | 1.81 | 2 |
| Q8R1I1 | Cytochrome b-c1 complex subunit 9 | 1.68 | 0 |
| Q91V92 | ATP-citrate synthase | 1.12 | 1 |
| Q921M7 | Protein FAM49B | 1.17 | 0 |

| | | | |
|---|---|---|---|
| Q99KI0 | Aconitate hydratase, mitochondrial | 1.15 | 2 |
| Q99KK9 | Probable histidine--tRNA ligase, mitochondrial | 1.46 | 2 |
| Q9CQA3 | Succinate dehydrogenase [ubiquinone] iron-sulfur subunit, mitochondrial | 1.45 | 2 |
| Q9CQE3 | 28S ribosomal protein S17, mitochondrial | 1.8 | 2 |
| Q9CYR0 | Single-stranded DNA-binding protein, mitochondrial | 1.4 | 0 |
| Q9CZ13 | Cytochrome b-c1 complex subunit 1, mitochondrial | 1.09 | 1 |
| Q9CZB0 | Succinate dehydrogenase cytochrome b560 subunit, mitochondrial | 1.31 | 2 |
| Q9D6R2 | Isocitrate dehydrogenase [NAD] subunit alpha, mitochondrial | 1.1 | 2 |
| Q9DB77 | Cytochrome b-c1 complex subunit 2, mitochondrial | 1.22 | 0 |
| Q9DCZ4 | MICOS complex subunit Mic26 | 2.16 | 1 |
| Q9JHI5 | Isovaleryl-CoA dehydrogenase, mitochondrial | 0.71 | 2 |
| Q9QZD8 | Mitochondrial dicarboxylate carrier | 1.3 | 0 |
| Q9Z110 | Delta-1-pyrroline-5-carboxylate synthase 1 | 1.13 | 2 |
