## supplemental Table 5 for "Biobased, Biodegradable but not bio-neutral: about the effects of polylactic acid nanoparticles on macrophages"

Supplementary Table 5: list of ER proteins modulated in response to PLA beac

| Uniprot accession | Protein name | ratio PLA treated/control | Mann Whitney U value |
|---|---|---|---|
| B9EJ86 | Oxysterol-binding protein-related protein 8 | 1.27 | 0 |
| O08547 | Vesicle-trafficking protein SEC22b | 0.59 | 0 |
| O08795 | Glucosidase 2 subunit beta | 0.87 | 0 |
| O08992 | Syntenin-1 | 2.71 | |
| O54692 | Centromere/kinetochore protein zw10 homolog | 1.57 | 0 |
| O70503 | Very-long-chain 3-oxoacyl-CoA reductase | 1.12 | 0 |
| O88455 | 7-dehydrocholesterol reductase | 2.33 | 0 |
| P35564 | Calnexin | 0.8 | 0 |
| P35821 | Tyrosine-protein phosphatase non-receptor type 1 | 1.75 | 0 |
| P61027 | Ras-related protein Rab-10 | 1.42 | 0 |
| P70704 | Phospholipid-transporting ATPase IA | 0.49 | 2 |
| P97858 | Solute carrier family 35 member B1 | 1.43 | 0 |
| Q64430 | Copper-transporting ATPase 1 | 1.31 | 2 |
| Q6P5E4 | UDP-glucose:glycoprotein glucosyltransferase 1 | 1.15 | 2 |
| Q6ZWQ7 | Signal peptidase complex subunit 3 | 1.36 | 1 |
| Q8BHN3 | Neutral alpha-glucosidase AB | 1.17 | 1 |
| Q8CGZ0 | Calcium homeostasis ER protein | 1.31 | 0 |
| Q8R1V4 | Transmembrane emp24 domain-containing protein 4 | 0.84 | 0 |
| Q8VDP6 | CDP-diacylglycerol--inositol 3-phosphatidyltransferase | 1.78 | 0 |
| Q91YQ5 | Dolichyl-diphosphooligosaccharide--protein glycosyltransferase subunit 1 | 1.1 | 0 |
| Q9CPU4 | Microsomal glutathione S-transferase 3 | 1.15 | 2 |
| Q9DB25 | Dolichyl-phosphate beta-glucosyltransferase | 1.31 | 1 |
| Q9DBS1 | Transmembrane protein 43 | 1.38 | 0 |
| Q9EPE9 | Endoplasmic reticulum transmembrane helix translocase | 0.68 | 2 |
| Q9EQH2 | Endoplasmic reticulum aminopeptidase 1 | 1.13 | 2 |
| Q9JHP7 | Protein O-glucosyltransferase 2 | 1.63 | 1 |
| Q9JLJ5 | Elongation of very long chain fatty acids protein 1 | 2.87 | 1 |
| Q9QY76 | Vesicle-associated membrane protein-associated protein B | 1.57 | 0 |
| Q9WU81 | Glucose-6-phosphate exchanger SLC37A2 | 1.5 | 2 |
| Q9WV55 | Vesicle-associated membrane protein-associated protein A | 1.5 | 2 |
