## supplemental Table 6 for "Biobased, Biodegradable but not bio-neutral: about the effects of polylactic acid nanoparticles on macrophages"

Supplementary Table 6: list of lysosomal/endosomal proteins modulated in response to PLA beads

| Uniprot accession | Protein name | ratio PLA treated/control | Mann Whitney U value |
|---|---|---|---|
| Q8CGS4 | Charged multivesicular body protein 3 | 2.01 | |
| Q3TCN2 | Putative phospholipase B-like 2 | 1.84 | 2 |
| P57716 | Nicastrin | 1.66 | 2 |
| P50518 | V-type proton ATPase subunit E 1 | 1.6 | 2 |
| Q9Z0M5 | Lysosomal acid lipase/cholesteryl ester hydrolase | 1.51 | |
| Q61093 | Cytochrome b-245 heavy chain | 1.49 | 1 |
| Q9CQF9 | Prenylcysteine oxidase 1 | 1.43 | |
| P61027 | Ras-related protein Rab-10 | 1.42 | |
| Q64430 | Copper-transporting ATPase 1 | 1.31 | 2 |
| P11438 | Lysosome-associated membrane glycoprotein 1 | 1.26 | 2 |
| P50516 | V-type proton ATPase catalytic subunit A | 1.24 | |
| P17426 | AP-2 complex subunit alpha-1 | 1.24 | 2 |
| P23780 | Beta-galactosidase | 1.18 | |
| P45377 | Aldose reductase-related protein 2 | 1.18 | 1 |
| P62944 | AP-2 complex subunit beta | 1.18 | 2 |
| P61028 | Ras-related protein Rab-8B | 1.17 | |
| Q8VEB4 | Phospholipase A2 group XV | 0.81 | 2 |
| Q63524 | Transmembrane emp24 domain-containing protein 2 | 0.71 | 2 |
| P17439 | Lysosomal acid glucosylceramidase | 0.7 | |
| P24668 | Cation-dependent mannose-6-phosphate receptor | 0.64 | |
| B5DEN9 | Vacuolar protein sorting-associated protein 28 homolog | 0.51 | |
| Q9WTI7 | Unconventional myosin-Ic | 1.63 | |
| Q5FWK3 | Rho GTPase-activating protein 1 | 1.63 | |
| E9Q634 | Unconventional myosin-Ie | 1.4 | |
| Q4LDD4 | Arf-GAP with Rho-GAP domain, ANK repeat and PH domain-containing protein 1 | 1.38 | 2 |
| O70318 | Band 4.1-like protein 2 | 1.27 | |
| P70248 | Unconventional myosin-If | 1.26 | |
| Q9JKF1 | Ras GTPase-activating-like protein IQGAP1 | 1.12 | 1 |
| Q4V7C7 | Actin-related protein 3 | 0.93 | 2 |
| Q8VDD5 | Myosin-9 | 0.92 | |
| Q6IRU2 | Tropomyosin alpha-4 chain | 0.87 | 2 |
| P13020 | Gelsolin | 0.82 | |
| P21107 | Tropomyosin alpha-3 chain | 0.82 | 2 |
| P59999 | Actin-related protein 2/3 complex subunit 4 | 0.7 | 1 |
| Q8BPU7 | Engulfment and cell motility protein 1 | 0.67 | 2 |
| Q8BHL5 | Engulfment and cell motility protein 2 | 0.42 | 2 |
