## supplemental Table 7 for "Biobased, Biodegradable but not bio-neutral: about the effects of polylactic acid nanoparticles on macrophages"

Supplementary Table 7: List of proteins linked to immune functions modulated in response to PLA beads

| Uniprot accession | Protein name | ratio PLA treated/control | Mann Whitney U value |
|---|---|---|---|
| A1L314 | MPEG-1 | 1.32 | 0 |
| O70133 | ATP-dependent RNA helicase A | 1.23 | 1 |
| P09581 | Macrophage colony-stimulating factor 1 receptor | 1.44 | 0 |
| P10810 | Monocyte differentiation antigen CD14 | 0.77 | 0 |
| P11680 | Properdin | 0.75 | 0 |
| P11928 | 2'-5'-oligoadenylate synthase 1A | 2.05 | 2 |
| P30993 | C5a anaphylatoxin chemotactic receptor 1 | 1.17 | 1 |
| P36371 | Antigen peptide transporter 2 | 1.49 | 2 |
| P83868 | Prostaglandin E synthase 3 | 1.44 | 1 |
| P98086 | Complement C1q subcomponent subunit A | 0.82 | 0 |
| Q60664 | Inositol 1,4,5-triphosphate receptor associated 2 | 1.28 | 1 |
| Q60875 | Rho guanine nucleotide exchange factor 2 | 1.18 | 0 |
| Q61093 | Cytochrome b-245 heavy chain | 1.49 | 1 |
| Q6P549 | Phosphatidylinositol 3,4,5-trisphosphate 5-phosphatase 2 | 1.33 | 1 |
| Q6R5N8 | Toll-like receptor 13 | 1.62 | 1 |
| Q8VI93 | 2'-5'-oligoadenylate synthase 3 | 1.27 | 1 |
| Q9D8Y7 | Tumor necrosis factor alpha-induced protein 8-like protein 2 | 2.06 | 0 |
| Q9DBS1 | Transmembrane protein 43 | 1.38 | 0 |
| Q9EQH2 | Endoplasmic reticulum aminopeptidase 1 | 1.13 | 2 |
| Q9ESY9 | Gamma-interferon-inducible lysosomal thiol reductase | 0.78 | 0 |
| P61027 | Ras-related protein Rab-10 | 1.42 | 0 |
| Q64324 | Syntaxin-binding protein 2 | 1.37 | 2 |
