## supplemental Table 8 for "Biobased, Biodegradable but not bio-neutral: about the effects of polylactic acid nanoparticles on macrophages"

Supplementary Table 8: List of proteins linked to redox homeostasis modulated in response to PLA beads

| Uniprot accession | Protein name | ratio PLA treated/control | Mann Whitney U value |
|---|---|---|---|
| P08228 | Superoxide dismutase [Cu-Zn] | 0.71 | 0 |
| P20108 | Peroxiredoxin 3 | 1.21 | 0 |
| P45376 | Aldo-keto reductase family 1 member B1 | 0.84 | 0 |
| P45377 | Aldose reductase-related protein 2 | 1.18 | 1 |
| P47199 | Quinone oxidoreductase | 1.94 | 0 |
| Q61093 | Cytochrome b-245 heavy chain | 1.49 | 1 |
| Q8C7K6 | Prenylcysteine oxidase-like | 1.28 | 2 |
| Q8CG76 | Aflatoxin B1 aldehyde reductase member 2 | 1.64 | 2 |
| Q9CQF9 | Prenylcysteine oxidase 1 | 1.43 | 0 |
| Q9CY64 | Biliverdin reductase A | 1.21 | 1 |
| Q9JMH6 | Thioredoxin reductase 1, cytoplasmic | 1.09 | 2 |
| Q9Z2G9 | Oxidoreductase HTATIP2 | 1.41 | 0 |
| P06795 | Multidrug resistance protein 1B | 2.15 | 0 |
