## supplemental methods (shotgun proteomic analysis) for "Biobased, Biodegradable but not bio-neutral: about the effects of polylactic acid nanoparticles on macrophages"

### Supplementary methods: shotgun proteomics

The gel plugs were washed three times with 50 µL of 25 mM ammonium hydrogen carbonate ($NH_4HCO_3$) and 50 µL of acetonitrile. The cysteine residues were reduced by 50 µL of 10 mM dithiothreitol at 57°C and alkylated by 50 µL of 55 mM iodoacetamide. After two washes with $NH_4HCO_3$ and acetonitrile, the gel plugs were dehydrated by acetonitrile. The digestion of proteins was done in gel with 20 µL of 12 ng/µL of modified porcine trypsin (Promega, Madison, WI, USA) in 25 mM NH4HCO3. The digestion was performed overnight at room temperature. The generated peptides were extracted with 40 µL of 60% acetonitrile in 0.1% formic acid. Acetonitrile was evaporated under vacuum and samples were resuspended with 40 µL of a solution of 1% acetonitrile and 0.1% formic acid in order to obtain a final concentration of 500 ng/µL.

NanoLC-MS/MS analysis was performed using a nanoACQUITY Ultra-Performance-LC (Waters Corporation, Milford, USA) coupled to a Q-Exactive Plus mass spectrometer (Thermo Fisher Scientific, Bremen, Germany). The nanoLC system was composed of a nanoEase™ M/Z Peptide BEH C18 column (130Å, 250 mm x 75 µm with a 1.7 µm particle size, Waters Corporation, Milford, USA) and a nanoEase TM M/Z Symmetry C18 precolumn (100Å, 20 mm × 180 µm with a 5 µm particle size, Waters Corporation, Milford, USA). The solvent system consisted of 0.1% formic acid in water (solvent A) and 0.1% formic acid in acetonitrile (solvent B). 1.5 µL of sample was loaded into the enrichment column during 3 min at 5 µL/min with 99% of solvent A and 1% of solvent B. Elution of the peptides was performed at a flow rate of 350 nL/min with a 1-35% linear gradient of solvent B in 79 minutes. The Q-Exactive Plus was operated in data-dependent acquisition mode by automatically switching between full MS and consecutive MS/MS acquisitions. Full-scan MS spectra were collected from 300 - 1,800 m/z at a resolution of 70,000 at 200 m/z with an automatic gain control target fixed at $3 \times 10^6$ ions and a maximum injection time of 50 ms. The top 10 precursor ions with an intensity exceeding $2 \times 10^5$ ions and charge states ≥ 2 were selected from each MS spectrum for fragmentation by higher-energy collisional dissociation. Spectra were collected at a resolution of 17,500 at 200 m/z with a fixed first mass of 100 m/z, an automatic gain control target fixed at $1 \times 10^5$ ions and a maximum injection time of 100 ms. Dynamic exclusion time was set to 60 s. Raw data were converted into mgf files using the MSConvert tool from ProteomeWizard (v3.0.6090; http://proteowizard.sourceforge.net/).

For protein identification, the MS/MS data were interpreted using a local Mascot server with MASCOT 2.6.2 algorithm (Matrix Science, London, UK) against a database containing all *Mus musculus* and *Rattus norvegicus* entries from UniProtKB/SwissProt (version 2019_10, 25,156 sequences) and the corresponding 25,156 reverse entries. The database was generated using an in-house developed Fasta toolbox (https://iphc-galaxy.u-strasbg.fr/) allowing to automate fasta files manipulations such as extracting sequences from UniProtKB, removing duplicate entries, adding contaminants or decoy entries. Spectra were searched with a mass tolerance of 10 ppm for MS and 0.05 Da for MS/MS data, allowing a maximum of one trypsin missed cleavage. Trypsin was specified as enzyme. Acetylation of protein n-termini, carbamidomethylation of cysteine residues and oxidation of methionine residues were specified as variable modifications. Identification results were imported into Proline software version 2.2 (http://profiproteomics.fr/proline/) for validation. Peptide Spectrum Matches (PSM) with pretty rank equal to one and length greater than 7 amino acids were retained. False Discovery Rate was then optimized to be below 1% at PSM level using Mascot Adjusted E-value and below 1% at Protein Level using Mascot Standard score.

Peptides

Peptides abundances were extracted thanks to Proline software version 2.2 (http://profiproteomics.fr/proline/) using a m/z tolerance of 10 ppm. Alignment of the LC-MS runs was performed using Loess smoothing. Cross Assignment was performed within groups only. Protein Abundances were computed by sum of peptides abundances (normalized using the median).

Proteins were considered as significantly different if their U value in the Mann-Whitney U test was ≤2 in the control vs. sample comparison.
